# CGC1, a new reference genome for *Caenorhabditis elegans*

**DOI:** 10.1101/2024.12.04.626850

**Authors:** Kazuki Ichikawa, Massa J. Shoura, Karen L. Artiles, Dae-Eun Jeong, Chie Owa, Haruka Kobayashi, Yoshihiko Suzuki, Manami Kanamori, Yu Toyoshima, Yuichi Iino, Ann E. Rougvie, Lamia Wahba, Andrew Z. Fire, Erich M. Schwarz, Shinichi Morishita

## Abstract

The original 100.3 Mb reference genome for *Caenorhabditis elegans*, generated from the wild-type laboratory strain N2, has been crucial for analysis of *C. elegans* since 1998 and has been considered complete since 2005. Unexpectedly, this long-standing reference was shown to be incomplete in 2019 by a genome assembly from the N2-derived strain VC2010. Moreover, genetically divergent versions of N2 have arisen over decades of research and hindered reproducibility of *C. elegans* genetics and genomics. Here we provide a 106.4 Mb gap-free, telomere-to-telomere genome assembly of *C. elegans*, generated from CGC1, an isogenic derivative of the N2 strain. We used improved long-read sequencing and manual assembly of 43 recalcitrant genomic regions to overcome deficiencies of prior N2 and VC2010 assemblies, and to assemble tandem repeat loci including a 772-kb sequence for the 45S rRNA genes. While many differences from earlier assemblies came from repeat regions, unique additions to the genome were also found. Of 19,972 protein-coding genes in the N2 assembly, 19,790 (99.1%) encode products that are unchanged in the CGC1 assembly. The CGC1 assembly also may encode 183 new protein-coding and 163 new ncRNA genes. CGC1 thus provides both a completely defined reference genome and corresponding isogenic wild-type strain for *C. elegans*, allowing unique opportunities for model and systems biology.

## Introduction

The nematode *Caenorhabditis elegans* is a model for biology ranging from mechanistic functions of individual proteins and RNAs to multicellular interactions of development and neurobiology (BRENNER 1974; CORSI *et al*. 2015). For any organism, a key element of modern biological understanding is to sequence and characterize its genome. *C. elegans* was the first animal to have its genome sequenced in 1998 (CONSORTIUM 1998); by 2005, its genome was considered complete and gap-free (HILLIER *et al*. 2005). It was thus surprising when, 14 years later, our attempt to reproduce a perfect isogenic copy of the reference *C. elegans* genome instead showed that it was neither complete nor gap-free (YOSHIMURA *et al*. 2019), as discrepancies observed by others were by then also suggesting (HILLIER *et al*. 2005; LI *et al*. 2015; TYSON *et al*. 2018).

*C. elegans*’ original reference genome (Table 1) was generated from the standard wild-type strain, N2 (BRENNER 1974). Unfortunately, N2 was probably genetically polymorphic even when it was first frozen in 1969 (STERKEN *et al*. 2015); it continued to accumulate observable genetic polymorphisms between laboratories into the 2000s (GEMS AND RIDDLE 2000; VERGARA *et al*. 2009); and, at this point, no frozen stock representing its original genotype exists. This lack of a frozen isogenic *C. elegans* wild-type strain with an exactly matched genome motivated our first effort to reproduce the *C. elegans* reference genome from an isogenic derivative of N2 (Table 1), which we published in 2019 as VC2010 (YOSHIMURA *et al*. 2019). VC2010 reproduced 100.3 Mb of N2 sequences with 99.98% identity, but also contained an extra 1.8 Mb of genomic sequence, along with ten genomic regions we could not assemble fully: five regions with long complex tandem repeats; tandem arrays of 5S rRNA genes (980 nt), 45S rRNA genes (7,197 nt), and positioning sequence on X (pSX1) sequences (172 nt) (NELSON AND HONDA 1985; ELLIS *et al*. 1986; JOHNSON *et al*. 2006); and two telomeric regions on the right ends of chromosomes I and III. For *C. elegans* to have a completely accurate reference genome, the N2 reference assembly needed to be replaced, but the VC2010 assembly could not fully replace it.

**Table 1.** Chromosome lengths and tandem repeat lengths of different *C. elegans* genome assemblies.

| Chromosome lengths |  |  |  |  |  |
| --- | --- | --- | --- | --- | --- |
| Chromosome | CGC1 (nt) | VC2010 (nt) | Difference of<br>CGC1 from<br>VC2010 (nt) | N2 (nt) | Difference of<br>CGC1 from<br>N2 (nt) |
| chrI | 16,508,284 | 15,331,301 | 1,176,983 | 15,072,434 | 1,435,850 |
| chrII | 15,800,154 | 15,525,148 | 275,006 | 15,279,421 | 520,733 |
| chrIII | 14,624,068 | 14,108,536 | 515,532 | 13,783,801 | 840,267 |
| chrIV | 18,555,121 | 17,759,200 | 795,921 | 17,493,829 | 1,061,292 |
| chrV | 22,067,386 | 21,243,235 | 824,151 | 20,924,180 | 1,143,206 |
| chrX | 18,785,586 | 18,110,855 | 674,731 | 17,718,942 | 1,066,644 |
| chrM | 13,994 | 13,988 | 6 | 13,794 | 200 |
| Total | 106,354,593 | 102,092,263 | 4,262,330 | 100,286,401 | 6,068,192 |
| Length of 174 >5-kb tandem repeats in each chromosome |  |  |  |  |  |
| Chromosome | CGC1 (nt) | VC2010 (nt) | Difference of<br>CGC1 from<br>VC2010 (nt) | N2 (nt) | Difference of<br>CGC1 from<br>N2 (nt) |
| chrI | 1,448,169 | 300,262 | 1,147,907 | 98,083 | 1,350,086 |
| chrII | 560,095 | 285,098 | 274,997 | 100,630 | 459,465 |
| chrIII | 904,536 | 410,543 | 496,012 | 164,534 | 740,002 |
| chrIV | 1,151,459 | 353,570 | 797,889 | 113,950 | 1,037,509 |
| chrV | 1,168,580 | 432,266 | 736,314 | 152,978 | 1,015,602 |
| chrX | 1,111,883 | 448,391 | 663,527 | 137,334 | 974,549 |
| chrM | 0 | 0 | 0 | 0 | 0 |
| Total | 6,344,722 | 2,230,130 | 4,116,646 | 767,509 | 5,577,213 |

Since 2019, long-read sequencing has improved enough to allow *de novo* assembly of telomere-to-telomere gap-free sequences for complex eukaryotic genomes, including the first human haploid genome (NURK *et al*. 2022). Our VC2010 genome assembly relied on short Illumina reads, long PacBio RSII reads, and long Oxford Nanopore MinION reads, with respective mean read lengths of 73 nt, 8.8 kb, and 14.2 kb, and respective error rates of 0.1%, 15-20%, and 15-20% (YOSHIMURA *et al*. 2019). Since then, PacBio and Nanopore have improved both the length and error rate of their long reads: PacBio Sequel II generates HiFi reads with a sequencing error rate of 0.1% and a mean read length of 15-20 kb (WENGER *et al*. 2019), while Nanopore PromethION generates ultralong reads with an error rate of 1% and 10% of their total sequence output (N10) having lengths of ≥150 kb (SHAFIN *et al*. 2020). These advances have enabled complete genome assemblies for humans (NURK *et al*. 2022) and other multicellular eukaryotes (XIE *et al*. 2024), and motivated us to assemble a comparable *C. elegans* reference genome. We found that conventional HiFi-read assembly tools produced accurate and long contigs for nonrepetitive regions of the *C. elegans* genome, but incorrectly assembled contigs for extremely long tandem repeats (TRs) such as the 5S rDNA, 45S rDNA, and pSX1 arrays. To solve this, we first assembled nonrepetitive contigs automatically, and then assembled 43 TR-filled genomic regions by intensive manual inspection. Here we describe the resulting *C. elegans* assembly, which with its matching wild-type *C. elegans* strain we call CGC1.

## Results

### CGC1, an isogenic C. elegans strain derived from N2

As a *C. elegans* reference strain, we selected CGC1, an isogenic derivative of the original reference strain N2 (BRENNER 1974) by way of VC2010 (FLIBOTTE *et al*. 2010). We originally called this isogenic derivative “PD1074” while calling our genome assembly of it “VC2010” (YOSHIMURA *et al*. 2019); however, these two names were confusingly different from each other, and were both difficult to remember. We thus renamed PD1074 as CGC1. With a few exceptions noted below, CGC1 should be genetically equivalent to N2; but unlike N2, it is as close to perfectly isogenic as possible, and has been mass-frozen in a multitude of vials, which should make its genetics and genomics fully reproducible by *C. elegans* laboratories over many years. CGC1 is available at the Caenorhabditis Genetics Center stock center (https://cgc.umn.edu/strain/CGC1).

### Genome sequencing and assembly

We harvested genomic DNA from CGC1 and sequenced it in two sets apiece: HiFi reads (WENGER *et al*. 2019) with 85x and 218x genome coverage, and Nanopore reads (SHAFIN *et al*. 2020) with 123x and 164x coverage (Supplementary Table 1a). HiFi and Nanopore had complementary advantages: HiFi reads had a 10-fold lower error rate (0.1%) than Nanopore (1.0%), whereas Nanopore reads were 7.6-fold longer (N10 of 151,427 nt in set 1) than HiFi (N10 of 19,847 nt in set 1; Supplementary Table 1b). We used the first HiFi and Nanopore read sets to assemble CGC1, and the second read sets to confirm the assembly’s accuracy.

We first assembled HiFi reads into 80 contigs (Supplementary Table 2) with hiCanu (NURK *et al*. 2020) and purge_dups (GUAN *et al*. 2020), and further reduced them to 61 non-redundant contigs by manually inspecting their order after aligning them to the VC2010 assembly (Supplementary Figure 1). This left us with 54 gaps between neighboring contigs, all of which could be spanned by Nanopore ultralong reads. Eleven gaps could be filled with the consensus of Nanopore ultralong reads. The remaining 43 gaps were filled through manual assembly, by first using Nanopore ultralong reads to close gaps throughout the genome assembly and then using either Nanopore or HiFi reads to correct errors and generate the complete genome (Figure 1). Figure 2 shows two examples where a single Nanopore read spans a gap. All but one of the 43 gaps could be spanned by a single Nanopore read; one remaining gap could be covered by combining two neighboring Nanopore ultralong reads (Supplementary Figure 2). We thus could find a series of alternating contigs and Nanopore ultralong reads that fully spanned each chromosome.

**Figure 1.**
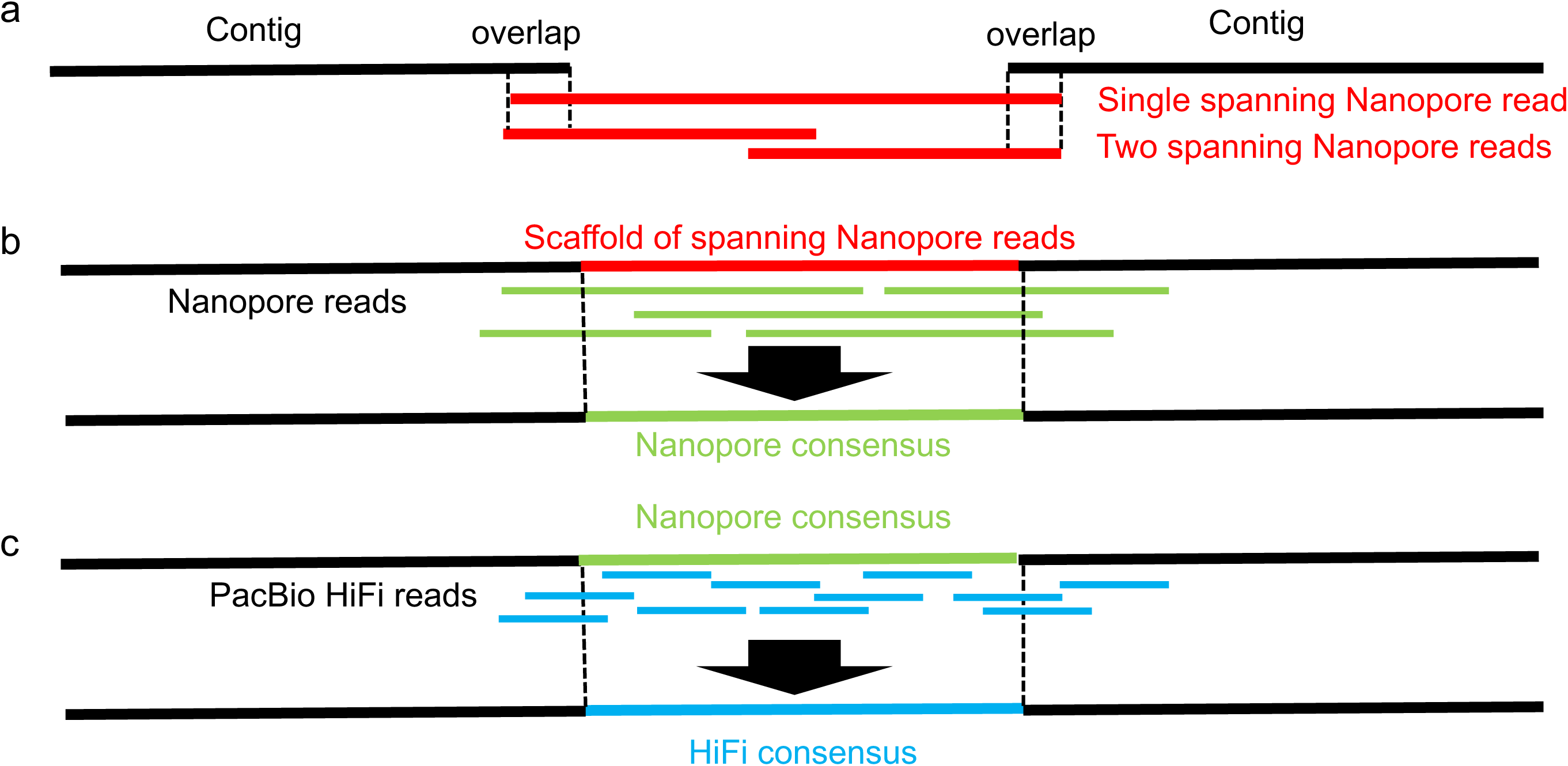
Overlap-layout-consensus approach of assembling tandem repeat regions. **a**, The overlap and layout step finds one or more Nanopore ultralong reads that span a focal gap, and lays out multiple reads properly. **b**, The consensus step corrects errors in the scaffold of Nanopore reads (red), aligns other Nanopore reads (green) to the scaffold, and calculates the consensus of the aligned reads. **c**, To further eliminate errors in the consensus sequence (green), HiFi reads (blue) are aligned to the consensus, and used to generate the consensus of mapped HiFi reads.

**Figure 2.**
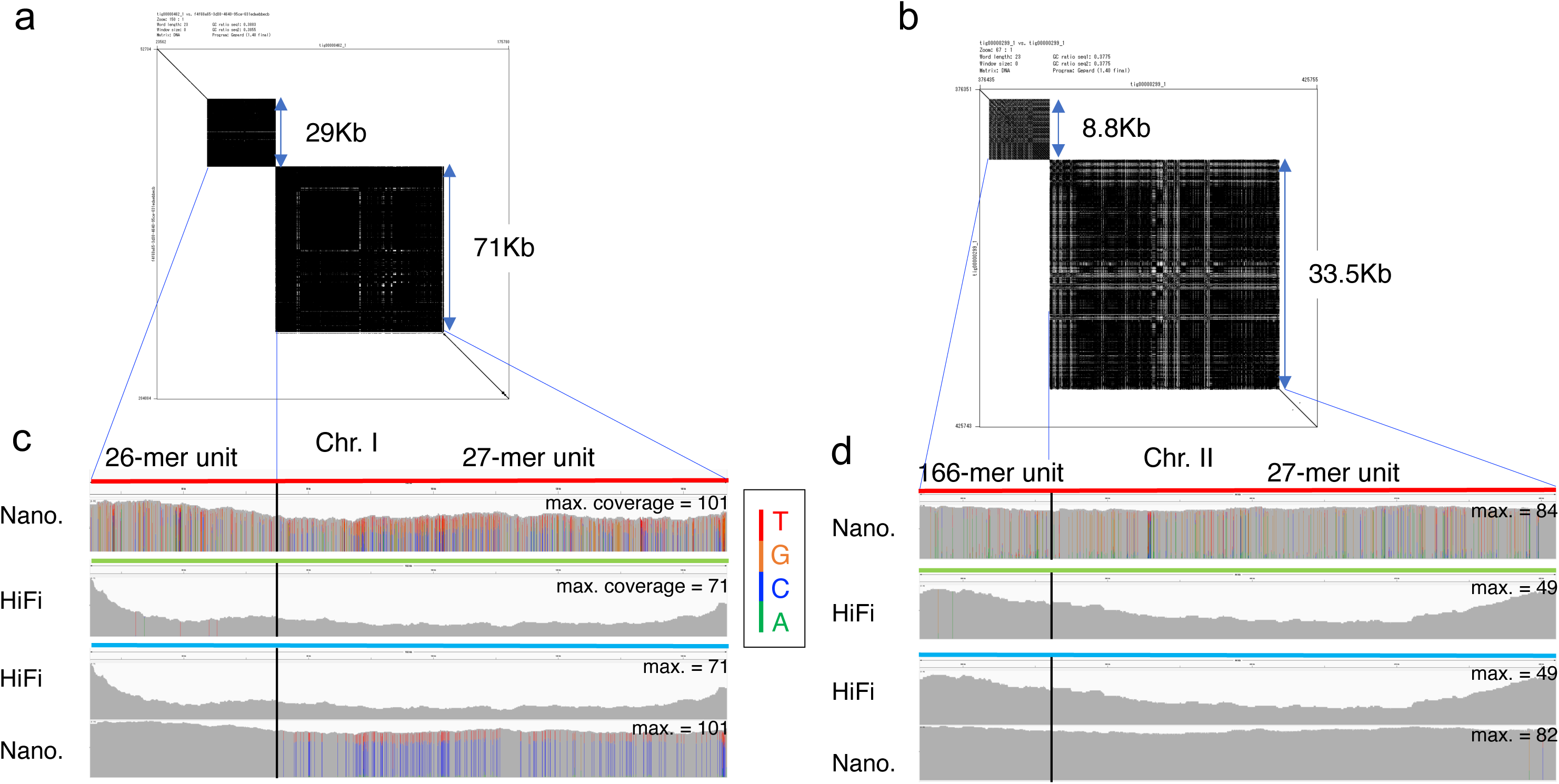
Polishing two complex tandem repeat regions in chromosomes I and II. **a-b**, Dot plots of two Nanopore reads that span neighboring tandem repeats in chromosome I (4,715,580-4,817,023) (**a**) and chromosome II (14,800,218-14,842,635) (**b**). **c**, The top row shows a spanning Nanopore read (colored red) and the coverage of other Nanopore reads mapped to the spanning read in chromosome I. The bar for each base represents the distribution of aligned bases in the consensus. If one nucleotide accounts for more than 80%, the bar is colored gray; otherwise, the four nucleotides A, C, G, and T are green, blue, orange, and red, respectively. The 2nd row shows coverage of HiFi reads mapped to the Nanopore consensus (light green). The third and fourth rows present coverages of HiFi reads and Nanopore reads to the HiFi consensus (light blue). Bars in the 3rd row are gray, showing the consistency of HiFi reads with the HiFi consensus. The 4th row shows that there are many bars with Cs (blue) as the majority and Ts (red) as the minority, but the base in the consensus of all bars is C. **d**, Read coverage of a spanning Nanopore read, Nanopore consensus, and HiFi consensus for complex tandem repeat regions in chromosome II.

### Error-correction of closed genomic gaps

Having closed 43 gaps with 1-2 Nanopore ultralong reads (Figure 1a), we had to correct errors of these reads by aligning other Nanopore reads to each gap and computing the consensus of Nanopore reads (Figure 1b). We then aligned HiFi reads to each Nanopore consensus and examined if sequencing errors could be corrected. For 25 out of 43 gaps, HiFi reads were useful in correcting sequencing errors, while for 13 of 43 gaps, Nanopore consensus sequences were sufficient. The remaining five of 43 gaps had complex regions with two different tandem repeats, making the process more complicated. We observed that 1.77%-3.59% of nucleotides in these five regions were corrected by the Nanopore consensus step (Table 2). However, we could further improve gaps by aligning HiFi reads to their Nanopore consensus and generating the consensus of aligned HiFi reads (Figure 1c, Supplementary Figure 3, and Materials and Methods for details). Overall, sequencing errors in 43 gaps were corrected by using consensus of Nanopore reads, consensus of HiFi reads, or a combination of both: the column of gaps in Supplementary Table 3 shows how each gap was filled, and Supplementary Figure 4 details the read coverage distribution of each gap by Nanopore and HiFi reads.

**Table 2.** Error correction by using the Nanopore and PacBio consensuses.

| Chromosome | Starting nt | Ending nt | Total length of tandem repeats | Nanopore corrections | HiFi corrections |
| --- | --- | --- | --- | --- | --- |
| I | 4,715,580 | 4,817,023 | 101,443 | 2,627 (2.59%) | 24 (0.024%) |
| II | 14,800,218 | 14,842,635 | 42,417 | 1,093 (2.58%) | 6 (0.014%) |
| III | 7,856,863 | 8,031,900 | 175,037 | 5,439 (3.11%) | 25 (0.014%) |
| X | 4,508,408 | 4,592,148 | 83,740 | 3,007 (3.59%) | NA |
| X | 5,581,121 | 5,673,280 | 92,159 | 1,633 (1.77%) | NA |

Nucleotide coordinates of these five gap regions are given for the final CGC1 genome assembly. Corrections for each read type (Nanopore or HiFi) include the total number of corrected bases, insertions, and deletions, and their ratio to the total length (in parentheses). "NA" in the last column implies that no HiFi corrections are made in chromosome X.

### Tandem repeats newly found in the CGC1 assembly

The CGC1 genome assembly is longer than the earlier VC2010 assembly (YOSHIMURA *et al*. 2019) by 3.8 Mb; 96% of this difference consists of 174 tandem repeats ≥5 kb in size (Table 1; Supplementary Table 3; Supplementary Data Files 1-3). Tandem repeats in CGC1 are noticeably larger than those also found in the VC2010 assembly; tandem repeats of ≥50 kb in size are observed only in the CGC1 assembly (Supplementary Figure 5a-c), presumably because previous methods underestimated them.

Of these 174 ≥5 kb tandem repeats, 50 were present in 43 gaps (seven gaps had two neighboring repeats), while the others are present in contigs assembled from HiFi reads by hiCanu. We tested how accurately we had assembled these repeat-containing contigs by mapping Nanopore ultralong reads, determining consensuses of the mapped Nanopore reads, and comparing the consensuses to the contigs. Nanopore read coverage was uniform in each tandem repeat and no significant structural discrepancies were found (Supplementary Figure 4).

Of these 174 ≥5 kb tandem repeats, some shared identical or near-identical units in common but were located on different chromosomes. A prominent instance of this was a set of 12 regions sharing the 27-mer tandem repeat 5’-ACT CTC TGT GGC TTC CCA CTA TAT TTT-3’, which motivated us to analyze their evolutionary proximity (Figure 3). Despite the evolutionary proximity of the tandem repeat regions in two groups of units, they are located far apart on different chromosomes. Supplementary Figure 6 and Supplementary Table 4 show the distribution of various units inside each region, implying that detailed analysis on the distributions of unit variants is informative to understand evolution of tandem repeats.

**Figure 3.**
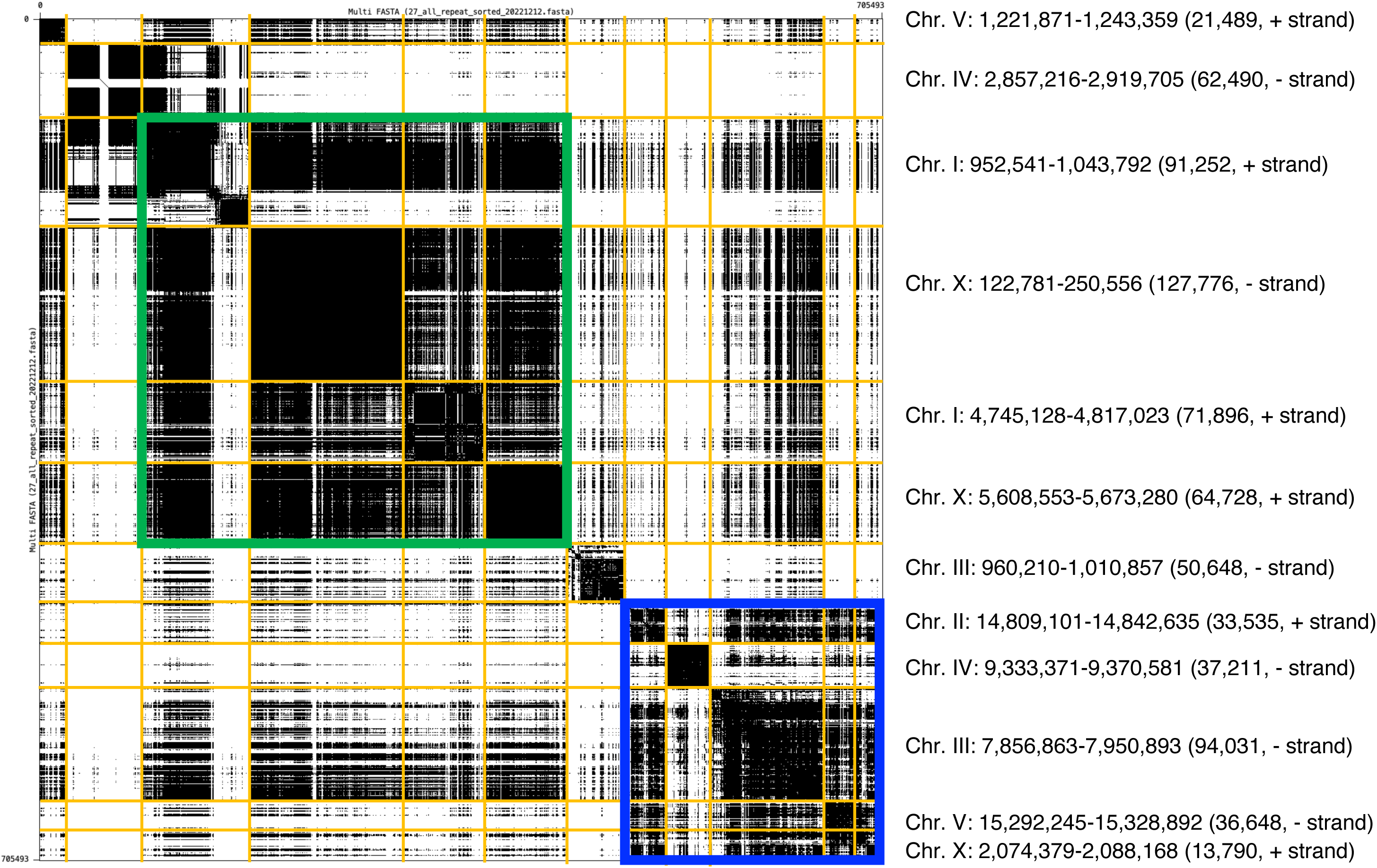
Similarity of prominent 27-mer tandem repeat regions. Dot plot of 12 prominent tandem repeat genomic regions sharing a common 27-mer repeat unit. For example, the top row shows a 21,489-nt genomic region of chromosome V (1,221,871-1,243,359) on the + (Watson) strand. Dots represent perfect matches of length 54 nt. For example, the dot plot for the fourth genomic region of the X chromosome (122,781-250,556) is very dense and appears black due to the high similarity between the 54-mers. In contrast, white areas indicate discordance. Regions are grouped by sequence similarity rather than their genomic position. The green box contains four genomic regions that share the most frequent version of the 27-mer unit (5’-ACT CTC TGT GGC TTC CCA CTA TAT TTT-3’), which we call type1. While, the blue box contains five genomic regions that share many copies of both type1 and another version of the 27-mer, type2 (5’-ACT CTC TGT GGC TTC CCA CCA TAT TTT-3’) that has one (underlined) T-to-C substitution with respect to type1.

### Comparison of CGC1 to automated GALA and hifiasm assemblies

Because alternative genome assembly methods are continually being developed, we tested our CGC1 assembly against two alternative approaches. Awad and Gan have recently described GALA, a program for gap-free chromosomal long-read *de novo* genome assembly, and have used it to produce an alternative genome assembly for *C. elegans* (AWAD AND GAN 2023). We compared our CGC1 assembly with their GALA assembly on four tandem repeat regions (Supplementary Figure 7). Three regions in the GALA assembly were much shorter than the corresponding regions in the CGC1 assembly, and another region in the GALA assembly was highly inconsistent with a Nanopore ultralong read and the CGC1 assembly. This problem with the GALA assembly is presumably due to their use of our limited PacBio and Nanopore sequence data reported in 2019 (YOSHIMURA *et al*. 2019), which was of lower quality than the new data in this study, demonstrating that the significant improvement in sequence quality of PacBio and Nanopore sequencing contributes to the high quality of CGC1 assemblies.

Cheng et al. (CHENG *et al*. 2021) have devised hifiasm, an assembler that can combine both HiFi and Nanopore reads into a single genome sequence. We used hifiasm 0.14 to assemble our two HiFi datasets with ≥20-kb reads and two Nanopore datasets with ≥50-kb reads (Supplementary Table 1) into a genome sequence that had 16 contigs for seven chromosomes, with an rDNA array divided into six smaller fragments (Supplementary Figure 8). Chromosome I, III, and X were split into two at large tandem repeats (chrI:952,541-1,043,792, chrIII:5,728,283-5,804,341, and chrX:12,540,490-12,624,876), and the hifiasm assembly’s pSX1 region was much shorter than that of CGC1 (Supplementary Figure 9). This demonstrates how highly tandemly repeated regions are still difficult to assemble automatically and require manual inspection.

### Assembly and analysis of the pSX1 and 5S rDNA tandem repeat regions

It has been challenging to assemble accurate sequences from genomic arrays of pSX1 (positioning sequence on X; 172 nt in size), 5S rDNA (976 nt), and 45S rDNA (7,197 nt) that form extremely large repetitive loci in the *C. elegans* genome (NELSON AND HONDA 1985; ELLIS *et al*. 1986; JOHNSON *et al*. 2006). In CGC1, we found that three ultralong reads and one ultralong read respectively spanned the pSX1 and 5S rDNA arrays of sizes 153 kb and 212 kb (Supplementary Figure 10a-b). We corrected sequencing errors in their spanning reads by aligning Nanopore reads and PacBio HiFi reads to the two spanning reads and determining the consensus sequences of aligned reads (Supplementary Figure 10c-d). In the resulting error-corrected pSX1 and 5S rDNA arrays, we respectively identified 887 and 198 copies of pSX1 and 5S rDNA elements (Supplementary Table 5); in the VC2010 assembly (YOSHIMURA *et al*. 2019), these had been respectively underestimated to have 167 and 144-145 copies. This discrepancy might be due to our lack of ultralong reads when assembling VC2010. Alternatively, it might reflect real genetic change: tandem repeat lengths at the rDNA locus of *C. elegans* can vary dynamically over time (WAHBA *et al*. 2021), and such variation might at least partially account for differences between our earlier VC2010 and later CGC1 genome assemblies. An independent assembly of the 5S rDNA locus with Nanopore reads from the N2 strain by Ding et al. estimated 167 consecutive 5S rDNA units (DING *et al*. 2022). These results demonstrate the importance of Nanopore ultralong reads for properly assembling large tandemly repeated genomic sequences.

The complete sequences of the pSX1 and 5S rDNA arrays provide an opportunity to study how they evolved (Figure 4 and Supplementary Figure 11). In both arrays, some variants of the repeated element appear to preferentially occur near one another significantly (according to the Wald–Wolfowitz runs test), suggesting that they might have expanded through replication slippage of a single progenitor variant. The 153-kb pSX1 array in chromosome X was homologous to another 115-kb pSX1 array in chromosome V (Supplementary Figure 12). In the X-chromosomal array, the most frequent 11 pSX1 variants matched the reference pSX1 element by >97%, and each variant had a unique set of substitutions and insertions (Figure 4). By contrast, in the chromosome V array, the most frequent 10 variants match the reference by ≤92%, and they lacked unique substitutions or deletions which would allow individual variants to be unambiguously distinguished (Supplementary Figure 13). One possible explanation for this pattern is that the X-chromosomal array arose by more recent transposition and amplification of a simpler pSX1 segment from the chromosome V array, which would give elements in the X-chromosomal array less time to diverge from one another.

**Figure 4.**
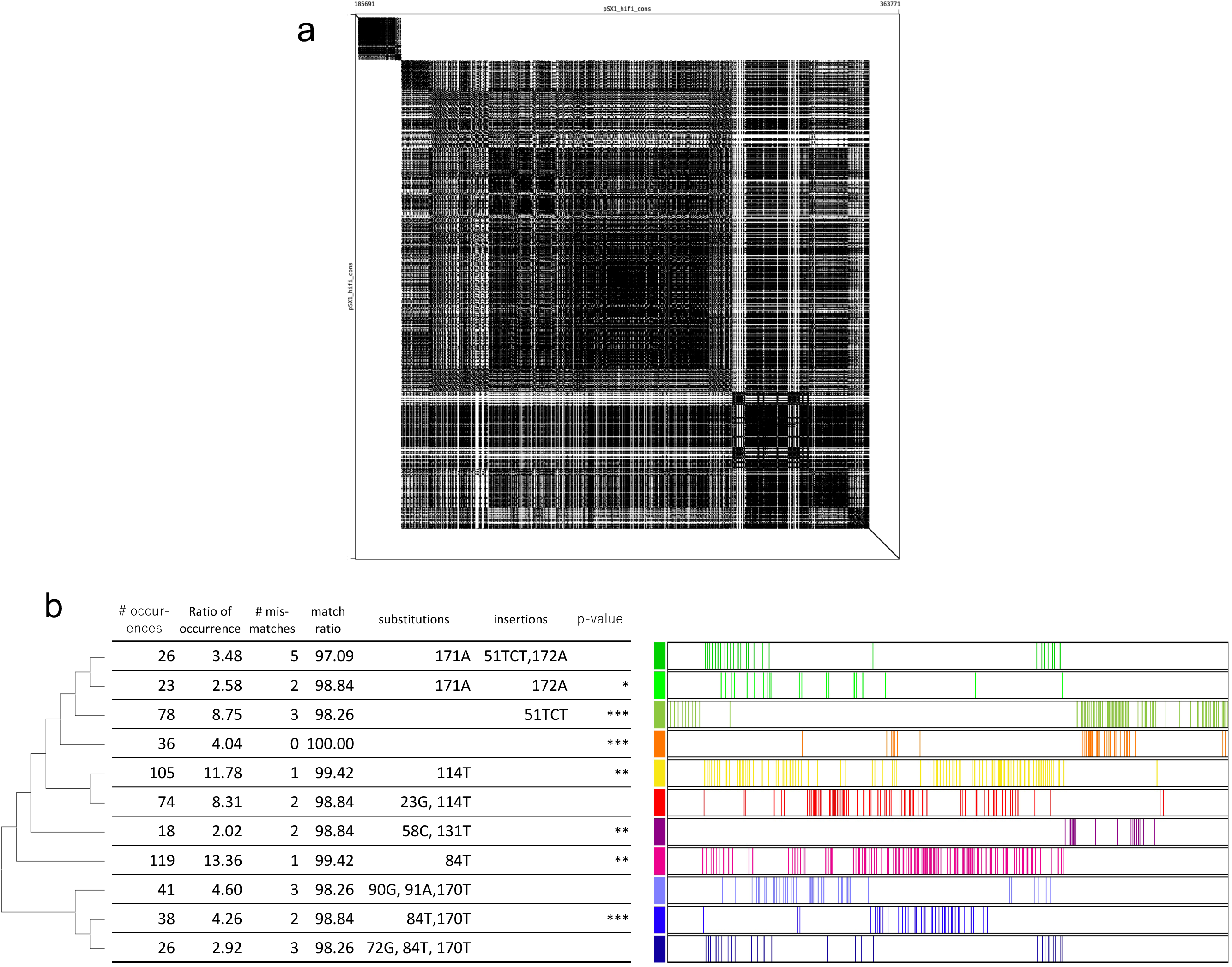
Structure of the pSX1 array. **a**, Self-dot plot of a region with the 153-kb pSX1 array in chromosome X (441,085-594,508). Dots represent perfect matches of substrings of length 100 nt. **b**, On the left side of the table, the phylogenetic tree shows the proximity of variants in terms of sequence similarity. On the right side of the table, the colored bars indicate the locations of the occurrences of each variant. Characteristics of eleven frequent variants of the reference pSX1 (172 nt, orange): the number of occurrences of each variant, the ratio of number of occurrences to total, number of mismatches with the reference pSX1, percentage match with the reference, positions of base substitutions (e.g., 171A means the base at position 171 is substituted with A), and positions with insertion bases (e.g., 51TCT shows TCT is inserted at position 51), and the statistical significance codes for p-values (***<=0.1%, **<=1%, and *<=5%) such that each variant occurs adjacent to each other preferentially according to the Wald– Wolfowitz runs test.

### Assembly and analysis of the 45S rDNA tandem repeat region

To assemble the 45S rDNA array encoding 18S, 5.8S, and 28S rRNAs, we first searched the 45S rDNA representative repeat unit of 7,197 nt (Supplementary Table 5) for single nucleotide variants (SNVs) that could distinguish the positions of each 45S rDNA unit variant within the array. We aligned 3,322 PacBio HiFi reads to the 45S rDNA representative repeat unit. Because the HiFi reads were long enough for the 45S rDNA unit to occur approximately twice, HiFi read coverage in the 45S rDNA unit averaged 6,691 at 7,197 positions. In six of the 7,197 positions, minor single-nucleotide variants were detected 60 or more times (Figure 5a); we thus used these six SNVs as markers to assemble Nanopore reads to construct a 45S rDNA array. To do this reliably with Nanopore reads that had an error rate of ∼5% for the six SNVs, we searched for positions that perfectly matched eleven bases centered on each of the six SNV sites. We also used a HiFi-validated insertion of GTCC at position 2811 as a marker. We thus extracted Nanopore reads with the six SNVs from both Nanopore ultralong read datasets (Supplementary Table 1), aligned them so that the distances of adjacent SNVs and insertions within reads were consistent between aligned reads (Figure 5b), and calculated an initial consensus from these aligned Nanopore reads (Figure 5c). To correct persistent errors, we aligned ≥100-kb Nanopore reads in the two Nanopore read datasets to the initial consensus and recalculated the final consensus of the aligned reads.

**Figure 5.**
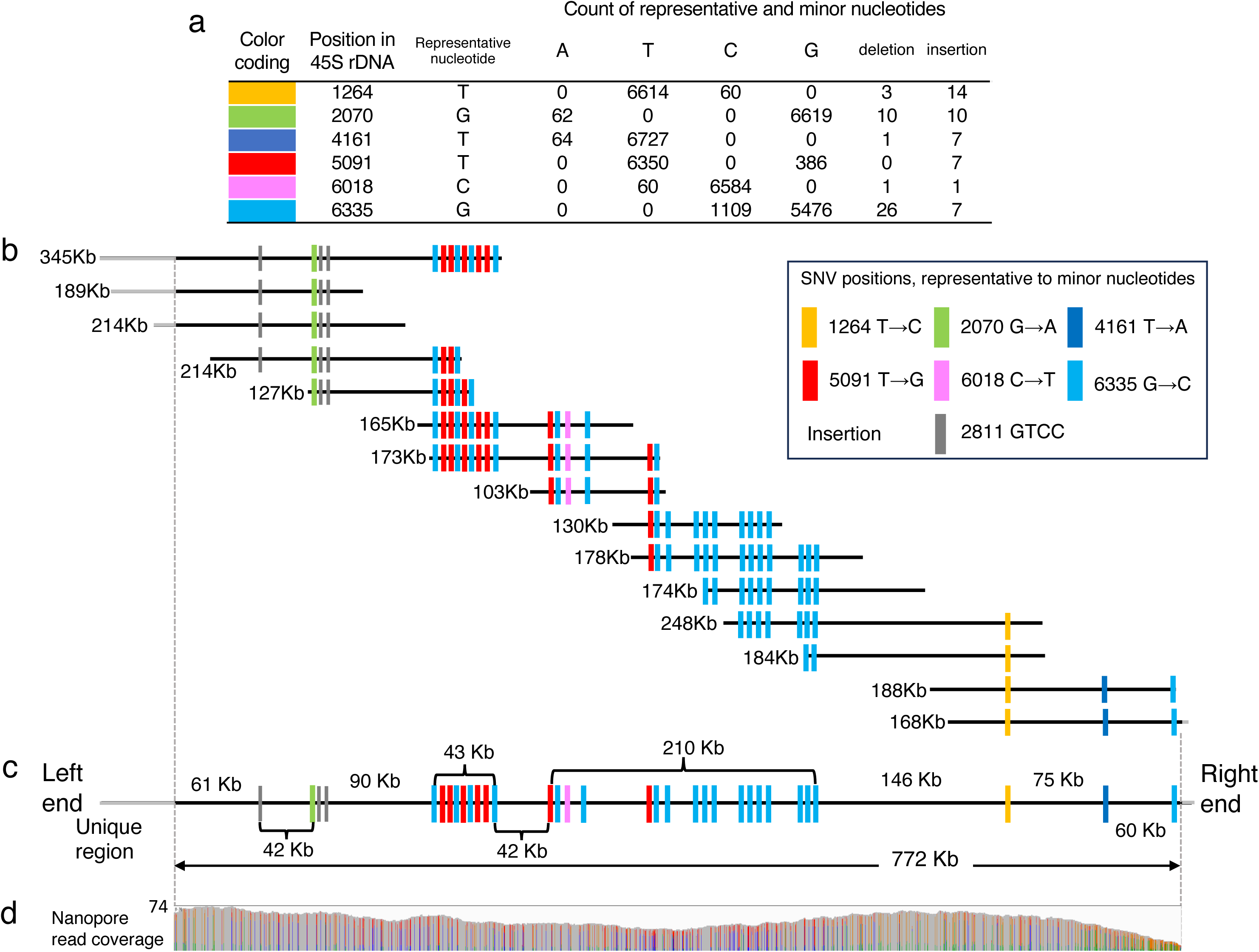
45S rDNA array at the right end of Chromosome I. **a**. Single nucleotide variants (SNVs) detected in the 45S rDNA representative repeat unit of size 7197 nt. 3322 PacBio HiFi reads were aligned to the 45S rDNA representative repeat unit. Because each HiFi read was long enough for the 45S rDNA unit to occur approximately twice, HiFi read coverage in the 45S rDNA unit averaged 6691 at 7197 positions. The table shows SNVs whose minor nucleotides are detected 60 or more times in PacBio HiFi reads. **b**, Nanopore reads are aligned so that distances of adjacent SNVs and insertions within Nanopore reads are consistent between aligned reads. **c**. Consensus of aligned Nanopore reads, an assembly of the 45S rDNA array with 107 units. Each position in the consensus is covered by two or more Nanopore reads shown in Figure **b**. **d**. Histogram displays read coverage for each position within the consensus by Nanopore reads >100 kb in length.

This yielded a final assembly of the 45S rDNA array with 107 units totaling 772 kb. This resembles both an independent size estimate of 799 kb from our own PacBio HiFi reads (Materials and Methods) and another study that used Nanopore sequencing to estimate a size of 705-820 kb for the 45S rDNA array in genomic DNA from a different laboratory population derived from N2 (DING *et al*. 2022). We further confirmed the accuracy of the variants in our assembled 45S rDNA assembly for CGC1 by mapping RNA-seq reads to the assembly (WAHBA *et al*. 2021) and verifying that they matched the six SNVs and GTCC insertion (Supplementary Table 6). We also tested the hypothesis that all 45S rDNA repeats occurred in a single array at the right end of chromosome I, by checking 497 Nanopore reads in the Nanopore dataset 1 that had BlastN hits of 45S rDNA and were ≥30 kb in size. After removing three chimeric reads, we found that the remaining 494 reads were consistent with the hypothesis: 442 of the 494 reads were filled with 45S rDNA occurrences, 36 had telomeric repeats (GGCTTA)_n_ at one end, and 16 had the unique region before the 45S rDNA array at one end (Supplementary Table 7).

Finally, because changes in the copy number of ribosomal RNA genes are observed in other species such as budding yeast and humans (HORI *et al*. 2023), we investigated how the 45S rDNA array of CGC1 differs from arrays in other *C. elegans* strains. For this, we analyzed publicly available PacBio HiFi read datasets from two strains “DLW N2 Bristol”, “DLW CB4856 [Hawaiian strain]” (BUSH *et al*. 2023), and from two strains named ALT1 and ALT2 derived from N2 and CB4856 (LEE *et al*. 2023), according to the same procedure of analyzing CGC1. Because Nanopore ultralong reads were not available for these four strains, we could not completely assemble their 45S rDNA arrays. However, we reliably detected SNVs within the 7,197-nt 45S rDNA representative repeat unit; three of the six SNVs found in CGC1 were also found in the DLW N2 population, while no SNVs were present in the other three strains (Supplementary Table 8). On the other hand, DLW CB4856, ALT1, and ALT2 each had SNVs absent from in the CGC1 and DLW N2 strains. Thus, SNVs in the 45S rDNA array evolutionarily diverged between wild-type strains of *C. elegans*.

### Telomeric repeats on the left end and right end of each chromosome

Because CGC1 included complete assemblies of the rDNA array and a second subtelomeric repeat region, it was also possible (unlike in the earlier VC2010 assembly) to fully assemble telomeric repeats at both ends of all chromosomes of CGC1. To illustrate this, Supplementary Figure 14 displays dot plots between pairs of 10-kb genomic sequences taken from the left end and right end of each chromosome in CGC1, and Supplementary Table 9 shows the starting and ending positions of each telomeric repeat.

### The CGC1 genome remains dynamic

While frozen stocks of CGC1 can be assumed to have a constant genome composition, it remains important to be aware of continued genetic variation. In examining sequencing data from different populations of strain CGC1 used in this work, we found a consistent site of genetic variation at position 9613 of the mitochondrial genome. This site is heteroplasmic, with a major allele (9613G) representing the majority of reads in the initial datasets and a minor allele (9613T) shown in a small fraction of sequencing reads (0-4.56%) (Supplementary Table 10). Strikingly, this fraction increased in samples derived more recently from animals that were propagated in the laboratory through multiple generations. While we do not know whether this allele varied because of adaptive selection or genetic drift, these results exemplify the biological dynamism unavoidable by any genome reference, and by CGC1 in particular.

### Genetic contents of CGC1

Protein-coding and ncRNA genes of *C. elegans*, along with repetitive DNA elements and pseudogenes, have been identified by manual curation of its N2 reference genome over the past 25 years (STERNBERG *et al*. 2024). For the new CGC1 reference genome to be useful to biologists, it must be annotated in a way that preserves as much of the older N2 annotation content as possible, while also indicating new genes missing in the earlier N2 assembly. To do this, we used LiftOff (SHUMATE AND SALZBERG 2021) to map N2 genes onto the CGC1 assembly (i.e., to lift the genes over from the N2 to the CGC1 assemblies; Supplementary Table 11; Supplementary Data Files 4-5), and determined which genes kept their contents unchanged in CGC1. We also used AUGUSTUS (STANKE *et al*. 2008) and StringTie2 (KOVAKA *et al*. 2019) to predict genes and transcripts in CGC1 (Supplementary Tables 12-13; Supplementary Data Files 6-9), and identified which were most likely to be genuinely new. To define a background of N2-shared versus CGC1-unique sequences in the CGC1 assembly to which genes or transcripts could be assigned, we identified uniquely aligned blocks between assemblies with MUMmer4 (MARCAIS *et al*. 2018).

To find common genes, we took nuclear (nonmitochondrial) N2 genes from the WS292 release of WormBase (STERNBERG *et al*. 2024) and mapped them to CGC1 as three sets: protein-coding, ncRNA, and pseudogenes. Of 19,972 protein-coding genes in N2 (WS292), all but 75 could be mapped onto the CGC1 genome. Of the mapped genes, 24 became untranslatable; another 83 could be translated after mapping, but no longer encoded a protein identical to N2. Nevertheless, 19,790 (99.1%) of N2 protein-coding genes encoded at least one unchanged product after mapping to CGC1. Of the 75 unmapped protein-coding genes, 49 encoded micropeptides of 9-15 amino acids (OLEXIOUK *et al*. 2018) that made them difficult to map; the other 26 unmapped N2 genes, which encoded proteins of ≥100 amino acids, all had orthologs among the AUGUSTUS gene predictions. This suggests that these 26 N2 genes might exist in the CGC1 assembly, despite not being identified by LiftOff, and might be identified by examining their AUGUSTUS orthologs in CGC1. Of the 24 untranslatable mapped genes, 22 (91.7%) overlapped AUGUSTUS gene predictions; of the 83 mapped genes with altered translation, 79 (95.2%) overlapped AUGUSTUS predictions. This suggests the 101 N2 genes with lost or altered translation in the CGC1 assembly can be corrected by examining their overlapping AUGUSTUS predictions. Overall, for protein-coding genes in N2, less than one out of 100 will require structural revision before analysis in CGC1, and there exist cognate AUGUSTUS gene predictions that should enable over 90% of revisions. Mapping of ncRNA genes and pseudogenes to CGC1 was equally successful: of 24,788 ncRNA genes in N2, only three could not be mapped to CGC1, and 24,591 (99.2%) encoded at least one transcript identical to N2; for 2,131 N2 pseudogenes, only four could not be mapped to CGC1, and 2,092 (98.2%) had at least one conceptual transcript identical to N2 (Supplementary Data File 10). Note that some genetic differences between the N2 and CGC1 genomes may be real, having arisen during the derivation of the CGC1 strain from the N2 strain; for instance, CGC1 carries a spontaneous partial deletion of *alh-2* (MAYDAN *et al*. 2010), and *alh-2* fell into our set of 83 genes with altered translation products. In addition, some N2 pseudogenes might correspond to functional genes in CGC1, because of mutations in CGC1 restoring the pseudogene to function (STEWART *et al*. 2005).

Long-read sequencing allows the assembly of tandemly repeated genomic regions that would otherwise be omitted by older technologies (ALKAN *et al*. 2011; DENTON *et al*. 2014; NURK *et al*. 2022), and this can reveal previously unobserved tandemly repeated genes. We found that 34 protein-coding genes in N2 had tandemly repeated copies in CGC1, totaling 60 extra genes (Supplementary Table 11). Notably, the telomerase-associated gene *pot-3* (YU *et al*. 2023) had seven tandem copies in CGC1 (Supplementary Table 11). Similarly, we found that 117 N2 ncRNA genes were tandemly repeated in CGC1 with 630 extra genes; most of these were copies of genes encoding 5S and 5.8S ribosomal RNAs (such as *rrn-2*, *rrn-4*, and *sls-1*).

Because CGC1 had 6.1 Mb of DNA missing from N2 that could encode new genes or exons, we used AUGUSTUS and StringTie2 to predict protein-coding and ncRNA genes *de novo*. To define CGC1 genomic coordinates for sequences either shared with N2 (Supplementary Data File 11) or unique to CGC1 (Supplementary Data File 12), we aligned the two assemblies with MUMmer4 (MARCAIS *et al*. 2018). We then predicted 21,238 protein-coding genes with AUGUSTUS. Of these, 19,232 (90.6%) overlapped 19,427 N2 protein-coding genes, showing that AUGUSTUS could repredict 97.3% of all N2 protein-coding genes that been lifted over into the CGC1 assembly. Of the non-N2-overlapping 2,006 AUGUSTUS predictions, 1,779 overlapped N2 genomic DNA and are described further below; but 227 at least partially overlapped CGC1-specific genomic DNA and might be new protein-coding genes of CGC1. In addition, 46 protein-coding N2 genes overlapped AUGUSTUS predictions that also encoded exons in local blocks of CGC1-specific genomic DNA; these may represent additional exons of the genes that were invisible without the CGC1 genome (Supplementary Table 11). Independently, we predicted 23,249 genomic loci encoding StringTie2 transcripts assembled from ribodepleted RNA-seq data (WANG *et al*. 2022). This allowed detection of 18,755 (93.8%) lifted-over N2 protein-coding genes and 19,746 (93.0%) AUGUSTUS protein-coding genes; as a further check on our protein-coding gene predictions, we mapped mass spectrometric data from three surveys of the *C. elegans* proteome (XIA *et al*. 2018; MÜLLER *et al*. 2020; CERON-NORIEGA *et al*. 2023), which yielded protein spectra for 9,508 (47.6%) of N2 genes and 9,640 (45.4%) of AUGUSTUS genes. StringTie2 also detected ncRNAs, with transcripts overlapping 5,173 (20.9%) of lifted-over N2 ncRNA genes, including long ncRNA genes such as *lep-5* (KIONTKE *et al*. 2019), *linc-4* (WEI *et al*. 2019; ISHTAYEH *et al*. 2021), and *tts-1* (ESSERS *et al*. 2015). Of the remaining 3,868 StringTie2 loci not overlapping these gene sets, 163 at least partly overlapped CGC1-specific genomic DNA and did not encode rRNA; these may represent novel ncRNA genes of CGC1 (Supplementary Table 14).

We used several criteria to identify which of the 2,006 new protein-coding genes predicted by AUGUSTUS represented real *C. elegans* genes. Out of the 227 that overlapped CGC1-specific genomic DNA, we discounted 14 whose protein products showed orthology to *C. elegans* transposon-encoded proteins. We also discounted 30 genes likely to be mispredictions from repetitive genomic DNA: 28 whose proteins consisted of ≥50% low-complexity sequence, and two whose proteins were predicted to encode 40 or 60 transmembrane domains. This left 183 genes with varying degrees of credibility: 150 had evidence for transcription (by overlapping StringTie2 transcripts), and 13 also had evidence for translation (by having protein spectra). Of the 33 non-StringTie2-associated genes, five showed orthology to genes in N2, *C. nigoni*, or *C. remanei.* Of the 137 genes with StringTie2 but not proteomic evidence, 31 encoded a variety of protein types, while 106 were tandem copies encoding a single putative ART2/RRT15 domain-containing protein with a coding sequence embedded in, but antisense to, tandemly repeated 26S ribosomal RNA genes. There is evidence that the sense strand of these 106 putative ART2/RRT15 genes may also be transcribed: they overlap with StringTie2 transcripts that were assembled from ribodepleted RNA-seq data, and they partially overlap an ncRNA antisense to 26S rRNA (F31C3.14) identified by the modENCODE consortium (LU *et al*. 2011). Homologs of these putative *C. elegans* ART2/RRT15 genes have been reported in diverse eukaryotes, including one *Saccharomyces cerevisiae* homolog (RRT15) that is also encoded antisense to rRNA and whose mutation lowers RNA polymerase I transcription (HONTZ *et al*. 2009). Failure to detect proteomic spectra for these putative ART2/RRT15 genes is inconclusive, because these spectra had a 53.4% false negative rate for known N2 genes. We conclude that some of these 183 genes are likely to be real, but defining their true status will require further analysis (Supplementary Table 15).

As a byproduct of predicting novel protein-coding genes and ncRNA transcripts in CGC1-specific genomic DNA, we also predicted genes and transcripts in CGC1 sequences shared with N2. WormBase curators have spent decades evaluating gene predictions in N2, and found many to be repetitive elements or pseudogenes; thus, new predictions in N2-shared genomic sequence must be treated with caution. In addition to repredicting 97.3% of known N2 protein-coding genes, AUGUSTUS also predicted 1,779 genes that fell into N2-shared genomic sequences but did not overlap known N2 genes. However, 1,465 of them corresponded with N2 pseudogenes or repetitive DNA elements. The remaining 314 genes (Supplementary Table 16) primarily encoded small proteins (median, 94 residues; range, 66-1,218 residues); 201 of these genes overlapped StringTie2 transcripts, and 12 had proteomic spectra. In addition to repredicting 20.9% of known N2 ncRNA genes, StringTie also predicted 3,300 transcripts (Supplementary Table 17) that fell into N2-shared genomic sequences but did not overlap known genes or pseudogenes, and had no ncRNA motifs from Rfam 14.10 (KALVARI *et al*. 2021). Like new gene predictions in CGC1-specific genomic DNA, new predictions in N2-shared genomic DNA are possibly but not certainly real.

### Transcription of long tandem repeat regions

In *C. elegans*, TRs could be transcriptionally active either because they fall inside regions of transcribed genes which harbor them, or because the TRs are themselves actively transcribed as non-coding RNA-encoding (ncRNA) genes. To address the former question, we examined the distribution of TRs in introns of protein-coding genes mapped from the N2 genome (Supplementary Table 18; Supplementary Figure 5d). Some introns were largely filled with TRs, but a number of introns were almost free from TRs. For example, among introns with a length of ≥1 kb, 113 consisted of ≥80% TR sequences, while 432 consisted of ≤20% (Supplementary Figure 5e). Thus, transcription of intronic TRs is commonplace, and some large introns tolerate being heavily occupied by TRs, but most large introns are predominantly non-TR sequence. To test further whether the 174 TRs of length ≥5 kb were transcribed, we mapped onto the CGC1 genome publicly available Nanopore RNA-seq data from different tissue types (mixed-stage embryo, L1, L2, L3, L4 larva, young adult, and gravid adult) in two independent studies (LI *et al*. 2020; ROACH *et al*. 2020). In four of the 174 long tandem repeats of length ≥5 kb (Supplementary Table 3), we observed RNA-seq reads from different tissue types that mapped onto the TRs with alternative splicing patterns outside the tandem repeats (Figure 6 and Supplementary Figure 15). Three of these instances fall into exons or introns of protein-coding genes; a fourth instance may be termination of mRNAs from adjacent genes. The possibility remains that TRs are specifically transcribed into ncRNAs (SUBIRANA AND MESSEGUER 2021), as is known to occur for some loci in the *Drosophila melanogaster* genome (BISWAS *et al*. 2024), but our long RNA-seq mappings could also be consistent with TR transcription in *C. elegans* being a byproduct of mRNA transcription.

**Figure 6.**
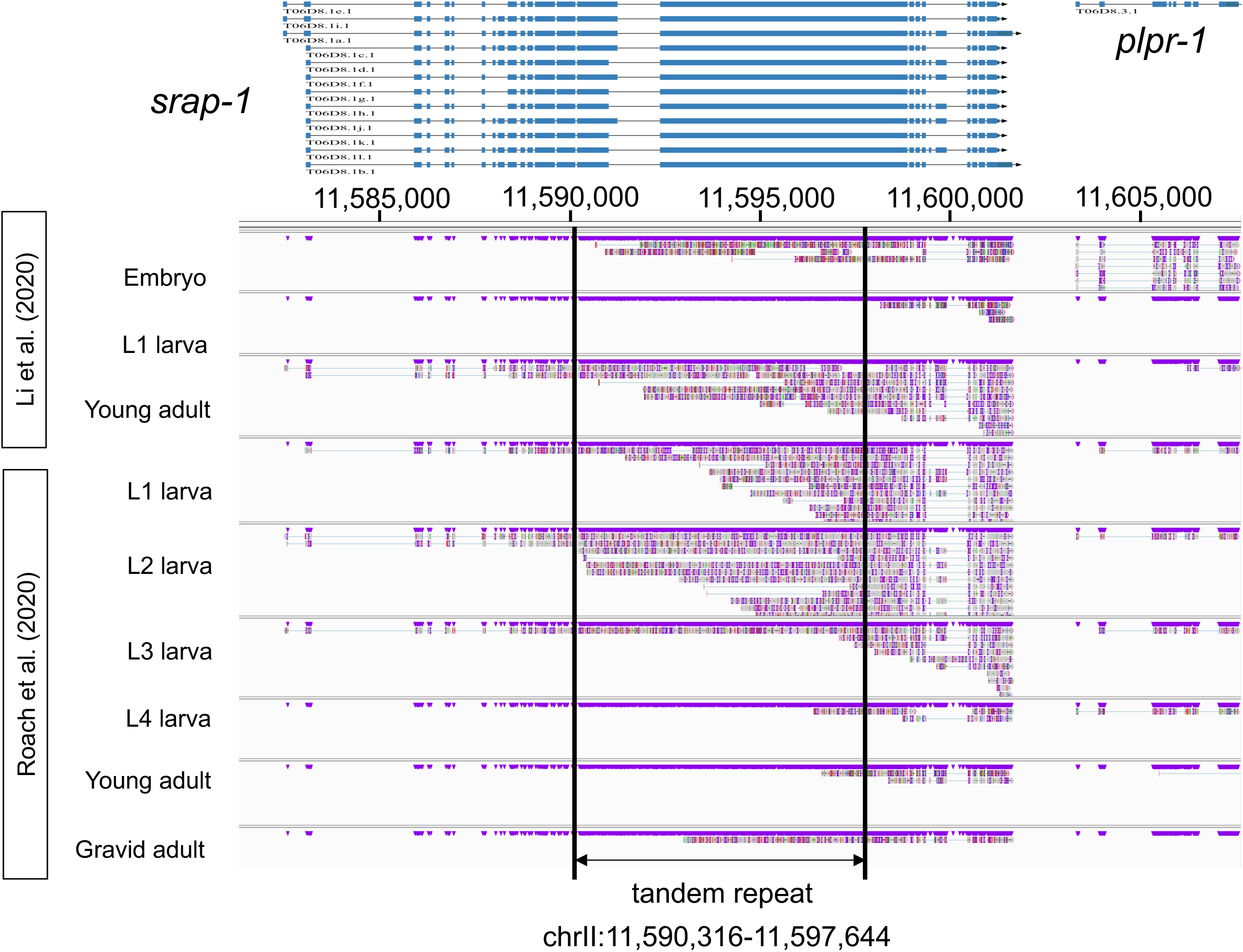
Transcription from a tandem repeat in various tissue types. This shows an alignment of long RNA-seq reads to a tandem repeat in chromosome II (nt 11,590,316-11,597,644). The topmost rows show mRNA isoform splicing for genes in the region, with *srap-1* containing the tandem repeat. Below, three tracks show alignments of tissue type data from Li et al. (LI *et al*. 2020) (GSE130044, NCBI Gene Expression Omnibus); six further tracks show tissue type data from Roach et al. (ROACH *et al*. 2020) (PRJEB31791, ENA). Alignment on tandem repeats and introns in *srap-1* were consistently observed in different tissue types, even though the RNA-seq data were collected in two independent studies, supporting the presence of transcription.

## Discussion

Using high-accuracy PacBio HiFi and ultralong Oxford Nanopore long-read sequencing, and augmenting our initial hiCanu assembly with manual assembly of 54 tandemly repeated regions, we have generated the first telomere-to-telomere gap-free genome assembly for the model organism *C. elegans*. This assembly encompasses the existing N2 reference genome and reproduces 99% of gene structures in N2, but adds 6 Mb of sequences. In addition to TRs, these added sequences encode additional interstitial exons for 46 protein-coding genes, 13-183 entirely new protein-coding genes, up to 163 new ncRNAs, and a 772-kb 45S rRNA gene array.

We found that automated general-purpose assemblers could generate reliable large contigs for much of the CGC1 genome, but that neither hiCanu nor hifiasm could correctly assemble CGC1’s longest TRs. Since *C. elegans* lies toward the lower end of genomic complexity for multicellular eukaryotes (GREGORY 2024), the putative TR sequences in automatic assemblies for other eukaryotic genomes should be viewed with caution; it is likely that large TRs like those of *C. elegans* exist in many other genomes but have been underassembled or missed entirely. Here, we have proposed a strategy of covering long TRs with Nanopore ultralong reads and then aligning PacBio HiFi reads to the ultralong reads to eliminate sequencing errors (Figure 1). When completing very long tandem repeats (e.g., the 45S rDNA array), we needed first to find positional markers manually, and then to overlap ultralong reads according to those positional markers to cover an entire tandem repeat (Figure 5). Future versions of genome assembly programs may find it possible and useful to automate this process.

The CGC1 genome assembly, with its matched clonal strain, should aid future *C. elegans* research in several ways. The most pragmatic of these is accuracy. When individual laboratories use classical and molecular genetics to dissect molecular and biological functions, they rely on their reference genome being completely correct, and since 2005 the general assumption has been that the N2 reference genome is completely correct (HILLIER *et al*. 2005). We show that this is not entirely true: 1% of N2 genes have some uncertainty that makes them impossible to map automatically into CGC1, or have cryptic exons only present in CGC1, or have tandem repeat copies that the N2 assembly fails to capture. The number of additional protein-coding genes encoded exclusively by CGC1 is small and uncertain, but not zero. Modern forward genetics of *C. elegans* often involves chemical mutagenesis and phenotypic screening followed by whole-genome sequencing to identify candidate alleles (DOITSIDOU *et al*. 2016): isogenicity of the CGC1 strain with completeness of the CGC1 genome should make mutant hunts more reliable and will both simplify and improve genetic and genomic studies.

CGC1 should also aid *C. elegans* research in population genomics, chromosomal organization, and the functions of highly repetitive genomic loci. Genetic polymorphism in wild isolates of *C. elegans* should be more reliably assigned to single-nucleotide variants and genomic islands of elevated diversity in the CGC1 assembly than in N2 (CROMBIE *et al*. 2019; LEE *et al*. 2021). Global traits of *C. elegans* chromosomes such as meiotic recombination rates and the densities of various genomic elements should be more correctly ascertained (CARLTON *et al*. 2022). The full sequencing of 45S rDNA array reported in this study should help determine the mechanism of maintaining rDNA clusters in *C. elegans*, as previously done in *S. cerevisiae* (KOBAYASHI AND GANLEY 2005); it might also clarify whether cellular senescence is correlated with rDNA stability, as suggested in mammalian cells (HORI *et al*. 2023). For many other TRs in *C. elegans*, studying their evolution and possible function only became feasible when CGC1 made them visible.

Finally, the CGC1 genome should enable synthetic biology in what is arguably the best understood and most experimentally tractable model metazoan, *C. elegans* (CORSI *et al*. 2015). The combination of powerful computational tools for predicting gene functions, efficient methods for large-scale DNA synthesis, and effective protocols for targeted genome mutagenesis (e.g., CRISPR) has made it practical to analyze a genome, devise a model for how that genome could be functionally modified (e.g., by streamlining or recoding it), synthesize an artificial revision of that genome, propagate the revised genome in a living cell, and experimentally test whether this confirms the functional model (VENTER *et al*. 2022; HOOSE *et al*. 2023). For such work, it is necessary (though not sufficient) that the organism’s genome assembly be entirely complete and accurate. Partly for that reason, attempts at such synthetic biology have focused heavily until now on modifying the genomes of single-celled organisms, such as the bacterium *Escherichia coli* or the baker’s yeast *Saccharomyces cerevisae* (PELLETIER *et al*. 2021; ZHAO *et al*. 2023; TIAN *et al*. 2024). An ultimate goal of synthetic biology is to scale this work upward to the most complex multicellular eukaryotes of interest, such as humans or other mammals, or such as the crop plants that sustain human life (BOEKE *et al*. 2016; TUNCEL *et al*. 2023). However, genome engineering faces a wide gap in difficulty between *E. coli* and *Homo sapiens*. *C. elegans* was selected as a model organism in part because its complexity is greater than that of a microbe, but less than that of a human. This intermediate complexity may make *C. elegans* an ideal test system for synthetic biology of animal genomes, and may make CGC1 the first *C. elegans* genome fully suited for such synthetic biology.

## Data Availability

The raw sequencing reads and genome assembly for *C. elegans* CGC1 are available under the accession number PRJDB19205 (Biosamples SAMD00841453-SAMD00841456). These data, along with predicted genes and transcripts, are also archived in the Open Science Framework (https://osf.io/n3fdy). We anticipate that gene predictions will also be added to GenBank after they have been curated by WormBase.

## Acknowledgments

This work was supported by the Japan Agency for Medical Research and Development (AMED) 24tm0424219h0004 and JSPS KAKENHI Grant Number JP22H04925 (PAGS) to SM, an Arnold O. Beckman Postdoctoral Independence Award and American Heart Association Postdoctoral Fellowship to MJS, NIH grant R35GM130366 to AZF, an NIH resource grant P40OD010440 for the Caenorhabditis Genetics Center to AER, and Cornell institutional funds to EMS. The authors would like to thank Jun Yoshimura for generating a previous VC2010 assembly, Yuta Suzuki for basecalling reads, and Information Technology Center, The University of Tokyo for permission to use their supercomputing facilities (Oakbridge-CX and mdx).

## Materials and Methods

### Orienting and visualizing genome assemblies with associated data

In this paper, we use the terms “left end” and “right end” of the *C. elegans* CGC1 reference chromosomes as designated in the original genetic map of Brenner (BRENNER 1974); the strands of CGC1 follow the original strand choice of the *C. elegans* N2 assembly (CONSORTIUM 1998). We drew dot plots between sequences of two chromosomes or chromosome sets with GenomeMatcher 3.0.6 (OHTSUBO *et al*. 2008) using the nucmer program in MUMmer 4.0.0rc1 (KURTZ *et al*. 2004); we drew dot plots between sequences of two tandem repeat regions with Gepard 2.1 (KRUMSIEK et al. 2007). We visualized multiple alignment of reads and read coverage of the CGC1 genome with the Integrative Genomics Viewer 2.12.3 (IGV) (ROBINSON et al. 2011). We visualized gene structures, StringTie2 transcripts, and other genome annotations with JBrowse2 (DIESH *et al*. 2023).

### C. elegans strain nomenclature

PD1074 (YOSHIMURA *et al*. 2019) is the original strain name of CGC1, which we renamed for two reasons: to have a strain name exactly identical to our telomere-to-telomere gap-free assembly name; and to have a strain/assembly name that would be easy for biologists to remember and use. CGC1 (i.e., PD1074) was derived from VC2010 by selfing an individual hermaphrodite and allowing the population to expand for approximately 10 generations before expansion and mass freezing at the CGC stock center. VC2010 was, in turn, a Moerman Gene Knockout Lab subculture of N2 obtained from the Caenorhabditis Genetics Center (https://cgc.umn.edu) in 2002 and first described by Flibotte et al. (FLIBOTTE *et al*. 2010).

### C. elegans genome and gene annotation versions

In analyzing genomic DNA from the N2 assembly, we have used the most recent stable version from WormBase, which has remained unchanged from WormBase release WS235 (made available by FTP on November 28, 2012) to at least WormBase release WS294 (made available by FTP on August 16, 2024; for full details of WormBase version releases, see https://wormbase.org/about/release_schedule). A specific copy of the N2 genome sequence that we used was downloaded from WormBase WS289 (Supplementary Table 20), but any version from WS235 to WS294 would have been identical. In analyzing annotations from the N2 assembly (either for genes or for repetitive genomic DNA regions; Supplementary Table 20), we have used WormBase release WS292 (made available by FTP on February 29, 2024). On one hand, the gene annotations in WS292 will not be exactly the same as those in any other version of Wormbase, so it might superficially seem that our analyses only apply to N2 genes from WS292 alone. However, gene structures from Wormbase have been manually curated for up to 25 years; by 2024, the gene annotations for N2 had become very heavily reviewed and quite stable, with relatively minor changes in overall gene content over time. For this reason, in the main text, we generally refer to “N2 genes” rather than “WS292 genes”. It is their origin from N2 that is most relevant to biologists; their being a specific snapshot from WS292 is technically important, but not of central biological relevance to our analyses here.

### Sample isolation

For PacBio and Nanopore DNA sequencing, *C. elegans* of the strain CGC1 was grown on enriched Nematode Growth Medium (NGM) plates seeded with an *E. coli* OP50 bacterial lawn (BRENNER 1974). Roughly synchronized adult animals were obtained by standard hypochlorite synchronization. The animals were washed off agar plates with cold buffer (50 mM NaCl, 5 mM EDTA, pH 7.5) into 15 ml conical tubes. Worms were centrifuged at 500x g for 3 minutes using a swinging bucket rotor in a tabletop centrifuge (Eppendorf Centrifuge 5702R). To decrease bacterial populations, pelleted animals were resuspended in 1 ml of cold buffer (50 mM NaCl, 5 mM EDTA, pH 7.5) and swirled persistently. Using a pipette, we gently transferred the worms to the top of a new tube with a 10 ml cold sucrose cushion (160 mM sucrose, 50 mM NaCl, 5 mM EDTA, pH 7.5), centrifuged at 100x g for two minutes in a tabletop centrifuge (Eppendorf Centrifuge 5702R) and the supernatant including bacteria was removed. Worm pellets at the bottom were used for further genomic DNA (gDNA) isolation.

### DNA isolation and PacBio DNA sequencing of C. elegans CGC1

For PacBio DNA sequencing, gDNA was extracted using Blood & Cell Culture DNA Mini Kit (Qiagen) including Genomic-tip 20/G according to the QIAGEN Genomic DNA Handbook. Briefly, 20 mg worm-pellets were homogenized mechanically in 2 ml Buffer G2 with Rnase A using four BioMasher II (Nippi, Japan) on dry ice. 100 μl Qiagen Protease stock solution was added to the sample and incubated at 50℃ for 2 hours. The sample containing particulate matter was centrifuged at 5000x g for 10 min at 4℃. Only the supernatant was transferred onto the QIAGEN Genomic-tip 20/G according to the Genomic-tip protocol. Genomic DNA collected in a new DNA Lo-Bind 2 ml tube from Genomic-tip was added 1.0X volume of AMPure PB beads (Pacific Biosciences) for purification. After mixing by pipetting 10X with wide bore tips, DNA was allowed to bind to beads at room temperature for 20 minutes with gentle shaking. The tube was placed in a magnetic rack to collect the beads to the side, and the cleared supernatant was slowly pipetted off. To the beads washed twice with freshly prepared 80% ethanol was added Qiagen buffer EB (10 mM Tris-Cl, pH 8.5) to elute the DNA at 37℃ for 15 minutes. The eluted DNA quality was checked using the dsDNA HS Assay Kit with a Qubit fluorometer (ThermoFisher Scientific), the NanoDrop spectrophotometer (ThermoFisher Scientific), and the Pippin Pulse system (Sage Science) using a 0.75% agarose gel and the 5-kb–430-kb protocol. HiFi PacBio SMRTbell library preparation required high-quality, high-molecular-weight input gDNA with a majority of the DNA fragments ≥50 kb as determined by Pippin Pulse.

PacBio SMRTbell libraries were prepared according to the manufacturer’s protocol. Briefly, gDNA was sheared twice using Diagenode’s Megaruptor 2 with a software setting of 25 kb and purified with 1.0X volume of AMPure PB beads (Pacific Biosciences). DNA sizing was checked on the FEMTO Pulse (Agilent) using the Genomic DNA 165 kb kit on extended mode. SMRTbell prep kit 3.0 (Pacific Biosciences) or SMRTbell express template prep kit 2.0 (Pacific Biosciences) was used to prepare SMRTbell libraries according to the manufacturer’s protocol. On the Sequel II system, binding kit 3.2 and cleanup beads or binding kit 2.2 (Pacific Biosciences) were needed for binding sequencing polymerase to SMRTbell libraries in preparation for sequencing. Sequel II sequencing kit 2.0 and SMRT Cell 8M (Pacific Biosciences) were used with 30 hours movie length.

### DNA isolation and Nanopore DNA sequencing of C. elegans CGC1

For Nanopore DNA sequencing, we aimed to extract gDNA from *C. elegans* with exceptionally low fragmentation and high-molecular-weight, according to the Nanopore protocol (Rapid sequencing gDNA-Ultra-Long Sequencing Kit, SQK-ULK001), and the ULK_9124_v110_revH_24Mar2021 version of this protocol was used in this paper. In our experiments, we were able to extract longer DNA using the new version protocol and this is described below. Briefly, approximately 1 g worm-pellets were gently homogenized in 10 ml of cold Cell Suspension Buffer (20mM Tris-HCl pH 7.5, 20mM EDTA, 350 mM sucrose) using the Qiagen TissueRuptor. The homogenate was passed through the 200μm, 100μm, 50μm and 30μm pluriStrainer (pluriSelect) in order. The approximate concentration of the nuclei in the purified homogenate was determined using a microscope and a 0.4 w/v% Trypan Blue solution stain. A volume corresponding to 6 million nuclei was used for further ultra-long DNA extraction using NEB Monarch HMW DNA Extraction Kit for Tissue according to the Nanopore protocol. The eluted ultra-long DNA quality was measured using the dsDNA BR Assay Kit with a Qubit fluorometer (ThermoFisher Scientific) and the Pippin Pulse system (Sage Science). Ultra-long DNA library preparation required high-quality, high-molecular-weight input DNA with a majority of the DNA fragments ≥150 kb as determined by the Pippin Pulse using a 0.75% agarose gel and the 5-kb-430-kb protocol.

Nanopore libraries were prepared using the Nanopore Ultra-Long Sequencing Kit (SQK-ULK001, ONT) and sequenced on a MinION sequencer (version Mk1B, ONT). Spot-ON Flow Cell R9 version (FLO-MIN 106D, ONT) was used with run length setting at 72 hours. To increase data output, a second library (at 24 hours after the run was started) and a third library (at 48 hours after the run was started) of ultra-long DNA were reloaded straight after flushing a flow cell using Nanopore Flow Cell Wash Kit (EXP-WSH004).

### Basecalling of PacBio and Nanopore reads

PacBio raw reads were processed into circular consensus sequence (CCS) reads (≥Q20; also termed HiFi reads) using the SMRT Link software v11.0 or v10.0. Nanopore fast5 files were basecalled via Guppy (version 5.0.16) with the arguments “*–flowcell FLO-MIN106 –kit SQK-ULK001*”.

### Genome assembly and refinement of the CGC1 genome

We first assembled HiFi reads into 602 contigs with hiCanu 2.1.1 (Nurk *et al*. 2020), consolidated them to 80 non-redundant contigs (Supplementary Table 2) with purge_dups 1.2.5 (Guan *et al*. 2020), and further reduced them to 61 non-redundant contigs by manually inspecting their order after aligning them to the VC2010 assembly with minimap2 v2.13 (Supplementary Figure 1). This left us with 54 gaps between neighboring contigs, all of which could be spanned by Nanopore ultralong reads. Eleven gaps were shorter than 1 kb, and could be determined by the consensus of Nanopore ultralong reads. However, the remaining 43 gaps were filled with long tandem repeats of ≥10 kb in size, making their correct sequence more difficult to determine. We therefore assembled these 43 gaps manually, by first using Nanopore ultralong reads to close gaps throughout the genome assembly and then using Nanopore and HiFi reads to correct errors and generate the complete genome (Figure 1).

### Identifying tandem repeats

We used mTR to identify tandem repeats within CGC1 and other *C. elegans* genome assemblies (MORISHITA *et al*. 2021). Tandem repeats are usually classified as short tandem repeats with a repeat unit length of 6 nt or less and minisatellites with a repeat unit length of 6 nt or longer.

### Filling gaps in five of 43 complex regions

This HiFi error correction step was successful in correcting errors in 0.014%-0.024% of sequence in three of the five complex regions (Table 2). However, in two other regions (Supplementary Figure 3), polishing the Nanopore consensus by HiFi reads increased a number of errors because the consensus at an erroneous base did not account for most bases and more than one base could be prevalent in the column. This suggested that, although HiFi reads are free from sequence bias in non-repetitive regions (WENGER *et al*. 2019), they may have sequence bias in tandem repeats. For example, because an adenine in a 27-mer unit on chromosome X (nt 5,581,121-5,673,280) was often called as a cytosine erroneously in HiFi reads (Supplementary Figure 3e), mapping HiFi reads to the Nanopore consensus caused an error rate of ∼4% (1/27). To correct for this error, we reasoned that HiFi reads are obtained as a consensus of subreads, that sequencing errors of subreads occur at random, and that mapping PacBio subreads instead of HiFi reads might help us determine which consensus was correct. Mapping PacBio subreads, we validated and chose the Nanopore consensus as the reference sequence (Supplementary Figure 3d-e).

### Identifying variants in tandemly repeated sequence elements

It is essential to ensure that tools used for assembly of tandemly repeated regions do not suffer from the artefact of regression to the mean, which would assign variants present in a minority of repeats to a majority sequence. Conversely, an approach that starts with identifying SNVs and indels that are characteristic of a minority of repeats and requiring a perfect alignment of these provides an opportunity for a definitive long-range assembly of such regions. Once a draft assembly has been produced through such an approach, a next challenge entails an assessment of reliability. One approach here would be to search a set of long reads (Supplementary Figure 16 shows ultralong reads that span the pSX1 array) for perfect matches (or, closest partial matches) of the variant repeat; however, this process is confounded in cases where the error rate is much higher than the variant rate. Nanopore’s ultralong reads have a sequence error rate of ∼5%); so, for example, ∼8 nt of the 172 nt pSX1 will on average be incorrect. We instead attempted to locate SNVs and indels unique to the variant, and listed perfect matches of a short motif string around a focal SNV (or indel) because a shorter string is less likely to include sequencing errors. Indeed, the occurrences of the short motif in two different Nanopore reads showed a clear pattern of reproducibility in the occurrences of the two variants in the assembled pSX1 array (Supplementary Figures 17-21).

### Analysis and error correction of the 45S rDNA array

Figure 5 shows how the 45S rDNA array was assembled from Nanopore reads. The size of the assembled 45S rDNA array reached to 772 kb, with confirmation from an independent dataset consisting of PacBio HiFi reads. Notable here, of the six SNV positions, SNVs at four positions (nt 1,264, 2,070, 4,161, and 6,018) occurred only once in the assembly. The minor nucleotides of these positions were covered by 60, 62, 64, and 60 HiFi reads, respectively, while the reference (majority) nucleotides were covered by 6,614, 6,619, 6,727, and 6,584 reads (Figure 5a). This is consistent with an array where one unit has the minor nucleotide and ∼110 units (e.g., 6,614/60) have a representative nucleotide within the 45S rDNA array assembly. Thus, the total length of the array approximates 799 kb (111 units times 7,197 nt per unit), which is close to 772 kb, the size of the assembled array. Another study estimated copy numbers of 98 to 114 units from the Nanopore reads of strain N2 (DING *et al*. 2022).

Figure 5d shows the read coverage distribution in the final consensus, and deletions were observed at 541 positions, all of which were homopolymers (strings of identical nucleotides) and likely generated from Nanopore sequencing errors. At 587 positions of the Nanopore consensus, over 20% of Nanopore reads had mismatches that did not match the consensus bases. This discrepancy is likely due to bias in Nanopore sequencing because thirteen bases centered on each mismatch did not match any PacBio HiFi reads while thirteen bases centered on the consensus base matched averaged 6,633 locations in PacBio HiFi reads. Overall, the final consensus had 107 copies of the 45S rDNA representative repeat unit with 30 SNVs and three insertions illustrated in Figure 5b.

### General manipulations of genomic data

Where possible, we used mamba to install and run version-controlled software environments from bioconda (GRUNING *et al*. 2018). For reformatting or parsing of computational results, we used Perl scripts either developed for general use or custom-coded for a given analysis. All such Perl scripts (named below with italics and the suffix “*.pl*”) were archived on GitHub (https://github.com/schwarzem/ems_perl). Internet sources (URLs) for other software are listed in Supplementary Table 19.

We extracted gene-encoded DNA or protein sequences from genomic DNA sequences and gene annotations in GTF/GFF format with gffread 0.12.7 (PERTEA AND PERTEA 2020) using the arguments ‘*-g [genomic sequence FASTA] -o /dev/null -C --keep-genes -P -V -H -y [translated protein sequence] -w [transcribed DNA sequence]*’. We identified overlaps between different features of the CGC1 assembly with the *intersect* tool of BEDTools 2.30.0 (QUINLAN AND HALL 2010) using the argument ‘*-loj*’. Before testing them for overlaps, genomic features were converted from GTF/GFF to BED format with *gff2bed* from BEDOPS 2.4.41 (NEPH *et al*. 2012) using the argument ‘*--do-not-sort*’.

### Obtaining published nematode genomic data

Published genome sequences, coding sequences, proteomes, and gene annotations were downloaded from WormBase (STERNBERG *et al*. 2024), ParaSite (HOWE *et al*. 2017), or the Open Science Framework (FOSTER AND DEARDORFF 2017) (Supplementary Table 20). Published ribodepleted RNA-seq data of *C. elegans* (WANG *et al*. 2022) were downloaded from the European Nucleotide Archive (Supplementary Table 20).

### Identifying shared and unique nucleotide sequences in the N2 and CGC1 assemblies

To determine nucleotide coordinates for regions of the CGC1 assembly that were either uniquely shared by the N2 assembly or not so shared, we first generated an alignment of uniquely matching sequences of the two assemblies with nucmer using the arguments ‘*--mum --breaklen 1 --maxgap 0 --batch 1*’, and then filtering their delta alignment with dnadiff, both nucmer and dnadiff being from MUMmer4 4.0.0rc1 (MARCAIS *et al*. 2018). We converted the resulting filtered delta alignment into MAF format with delta2maf from Mugsy 1.2.3 (ANGIUOLI AND SALZBERG 2011), and then from MAF to GFF format with maf-convert from LAST 1548 (KIELBASA *et al*. 2011) using the argument ‘*gff*’. Readable genomic nucleotide coordinate annotations were added to the GFF alignment with *id2nucmer_gffs_14jul2024.pl*. The GFF alignment was converted to BED format with gff2bed from BEDOPS 2.4.41 (NEPH *et al*. 2012) using the argument ‘*--do-not-sort*’, sorted with *sort* from Linux using the arguments ‘-k1,1 -k2,2n’, merged into nonredundant genomic regions with bedtools merge from BEDtools 2.30.0 using the argument ‘*-delim “|”*’, and converted back to GFF format with *convert_bed2gff.pl* from AGAT 1.3.3 (DAINAT *et al*. 2024). Source and sequence type annotations in the GFF alignment were converted from ‘data\tgene’ to ‘maf-convert\tregion’ with inline Perl. Alignment regions were given specific sequence-block names with *id2nucmer_gffs_26aug2024.pl*. The annotated GFF alignment was converted back to BAM format with gff2bed from BEDOPS 2.4.41 using the argument ‘*--do-not-sort*’, and genomically sorted with bedtools sort from BEDtools 2.30.0 using the argument ‘*-g [CGC1 genome assembly]*’. Coordinates of the CGC1 genome that were not part of the resulting N2 alignment were extracted into a BED file with bedtools complement from BEDtools 2.30.0 using the arguments ‘*-i [N2-CGC1 genome alignment] -g [CGC1 genome assembly]*’.

### Mapping N2 genetic content to the CGC1 assembly

We obtained canonical N2 gene annotations from WormBase WS292. Before mapping to them to CGC1, we split them into separate GFF3 annotation files with protein-coding genes, pseudogenes, and non-coding RNA genes, by selecting for the biotypes ‘protein_coding’, ‘pseudogene’, or any other biotypes (respectively); this first allowed us to use different mapping parameters where appropriate for different gene types, and later allowed us to compare *de novo* protein-coding genes separately to N2 protein-coding genes versus N2 pseudogenes. We mapped all three sets of canonical N2 WS292 gene annotations to the CGC1 assembly with Liftoff 1.6.3 (SHUMATE AND SALZBERG 2021) using the arguments ‘*-s 0.9 -cds - polish -copies -chroms [table of identical chromosomes]*’. For mapping of ncRNA genes, which were otherwise too small to map reliably, we also used the argument ‘*-flank 1.0*’.

### Prediction of protein-coding genes

We predicted protein-coding genes in the CGC1 assembly *de novo* with AUGUSTUS 3.5.0 (STANKE *et al*. 2008) using the arguments ‘*--strand=both --genemodel=partial --noInFrameStop=true -- singlestrand=false --maxtracks=3 --alternatives-from-sampling=true --alternatives-from-evidence=true --minexonintronprob=0.1 --minmeanexonintronprob=0.4 --uniqueGeneId=true --protein=on -- introns=on --start=on --stop=on --cds=on --codingseq=on --UTR=off --species=caenorhabditis -- extrinsicCfgFile=[augustus_dir]/config/extrinsic/extrinsic.ME.cfg --progress=true --gff3=on -- outfile=[gene_predictions].aug --hintsfile=[hints].gff*’. To generate hints for AUGUSTUS, we merged CDS DNA sequences both for N2 protein-coding genes from WormBase WS289 and for VC2010 v.1 protein-coding genes from WormBase WS268; we then mapped the CDS DNA sequences to CGC1 genomic DNA with BLAT v362 (KENT 2002) using the argument ‘*-minIdentity=92*’, filtered the BLAT alignments with pslCDnaFilter with the argument *’-maxAligns=1*’, and converted the alignments from PSL to GFF format with *blat2hints.pl* from AUGUSTUS.

### StringTie2 prediction of transcripts and transcribed genomic loci in CGC1

We predicted transcription units in the CGC1 assembly using paired-end rRNA-depleted *C. elegans* RNA-seq data from six N2 and *lpd-3* replicates (WANG *et al*. 2022) with StringTie 2.2.1 (KOVAKA *et al*. 2019) using the argument ‘-G *[AUGUSTUS gene predictions]*’ to identify transcripts corresponding with *de novo* protein-coding gene predictions (and, by exclusion, identify possible ncRNA transcripts). Before running StringTie2, we aggregated the RNA-seq reads into a single pair of read files; we used HISAT2 2.2.1 (KIM *et al*. 2019) to index the CGC1 genome with ‘*hisat2-build*’ and to map pooled RNA-seq reads to it with ‘*hisat2 --summary-file [mapping summary] -x [CGC1 HISAT2 index] -1 [read-pair end 1] -2 [read-pair end 2] -S [alignment in SAM format]*’; we then reformatted the resulting SAM alignment to a sorted, indexed BAM alignment with ‘*view -b*’, ‘*sort*’, and ‘*index*’ in SAMtools 1.17 (DANECEK *et al*. 2021).

### Long RNA-seq analysis of TRs

We mapped long RNA-seq reads to the CGC1 assembly with winnowmap2 (JAIN *et al*. 2022). Of 174 tandem repeat regions of ≥5-kb in CGC1, 10 regions had alignments; however, six alignments showed no splicing of the long RNA-seq reads, suggesting that they might arise from DNA contamination rather than mRNA. Four other alignments with splicing were correlated with gene structures via JBrowse2 to distinguish likely intronic transcripts from possible ncRNAs.

### Annotation of gene products

For protein-coding genes, we predicted both N-terminal signal sequences and transmembrane alpha-helical anchors with Phobius 1.01 (KÄLL *et al*. 2004), reformatting results with *tabulate_phobius_hits.pl*. We predicted coiled-coil domains with Ncoils 2002.08.22 (LUPAS 1996), reformatting results with *tabulate_ncoils_x.fa.pl*. We predicted low-complexity regions with PSEG 1999.06.10 (WOOTTON 1994) using the argument ‘*-l*’ and reformatting results with *summarize_psegs.pl*. We identified protein domains from the Pfam 37.0 database (MISTRY *et al*. 2021) with hmmscan in HMMER 3.3.2, using the arguments ‘*--cut_ga*’ to impose family-specific significance thresholds, ‘*-o /dev/null*’ to discard text outputs, and ‘*--tblout*’ to export tabular outputs; Pfam results were reformatted with *pfam_hmmscan2annot.pl*. We also identified protein domains from the InterProScan 5.57-90.0 database with *interproscan.sh* (PAYSAN-LAFOSSE *et al*. 2023) using the arguments *’-dp iprlookup goterms*’, and reformatting results with *tabulate_iprscan_tsv.pl*. We identified orthologies between protein-coding gene sets from *C. elegans* (N2 and CGC1), *C. nigoni*, *C. remanei*, and *C. elegans* N2 transposon-encoded proteins, with OrthoFinder 2.5.4 (EMMS AND KELLY 2019) using the arguments ‘*-S diamond_ultra_sens - og*’; results were reformatted with *prot2gene_ofind.pl* and *genes2omcls.pl*. For all annotation analyses except OrthoFinder, full proteomes were used; for OrthoFinder, we used maximum-isoform proteome subsets generated with *get_largest_isoforms.pl*. An overall annotation table was constructed from these and other gene annotations (such as overlaps between lifted-over N2 and CGC1 AUGUSTUS genes) with *add_tab_annots.pl*.

For StringTie2 transcripts, we identified ncRNA motifs from the Rfam 14.10 database (KALVARI *et al*. 2021) with cmsearch in INFERNAL 1.1.5 (NAWROCKI AND EDDY 2013) using the arguments ‘*--cut_ga*’ to impose family-specific significance thresholds, ‘*-o /dev/null*’ to discard text outputs, and ‘*--tblout*’ to export tabular outputs.

### Detection of proteomic spectra matching predicted C. elegans proteins

We downloaded mass spectrometric data in RAW file format (Supplementary Table 20) from the PRIDE database (PEREZ-RIVEROL *et al*. 2022) for three surveys of the *C. elegans* proteome (XIA *et al*. 2018; MÜLLER *et al*. 2020; CERON-NORIEGA *et al*. 2023). We reformatted these data files from RAW to mzML format with thermorawfileparser 1.4.4 (HULSTAERT *et al*. 2020), using the arguments *’--format 2 - -metadata 1*’. We mapped protein spectra in these files to *C. elegans* protein sequences (from either our *de novo* AUGUSTUS gene predictions in CGC1, or from the pre-liftover N2 protein-coding genes from WormBase WS292) with Comet (ENG *et al*. 2013) from Crux 4.1 (MCILWAIN *et al*. 2014), using the arguments ‘*--nucleotide_reading_frame 0 --decoy_search 1*’. We statistically analyzed these three mappings as a single collective data set with Percolator (KALL *et al*. 2007) from Crux 4.1, using the argument ‘*--decoy-prefix decoy_*’. We classified protein-coding genes (from either AUGUSTUS predictions from the N2 or CGC1 assembly) as being detected in protein spectra if they showed mapped spectra with a statistical significance (corrected for multiple hypothesis testing) of q ≤ 0.01. This threshold was chosen to limit the rate of false positives to 1%; it is necessarily arbitrary, and will by nature entail some false negatives as well as false positives.

### Observation of ART2/RRT15 homologs in eukaryotes

We detected homologs of putative *C. elegans* ART2/RRT15-domain encoding genes by examining proteins associated with their shared InterPro motif (https://www.ebi.ac.uk/interpro/entry/InterPro/ *IPR052997*) and by BlastP searches of the NCBI nonredundant protein database (nr; *https:// blast.ncbi.nlm.nih.gov/Blast.cgi?PROGRAM=blastp*). Partial overlap of the ART2/RRT15-domain encoding genes with the previously observed *C. elegans* ncRNA gene F31C3.14 (LU *et al*. 2011) was detected by BlastN of the *C. elegans* reference genome (https://wormbase.org/tools/blast_blat) and confirmed by StringTie2 transcript overlaps (Supplementary Table 13).

### Disqualifying novel AUGUSTUS gene predictions or StringTie2 transcript loci

Most novel AUGUSTUS predictions (Supplementary Table 12) were identified as those that did not overlap with lifted-over N2 protein-coding genes, and that at least partially overlapped with sequences specific to the CGC1 assembly (i.e., that were not shared with the N2 assembly). For AUGUSTUS predictions that at least partially overlapped sequences specific to the CGC1 assembly, we disqualified them if they met any of the following criteria: orthology to a transposon-encoded protein; consisting of ≥50% low-complexity residues; or encoding ≥40 transmembrane domains. For other AUGUSTUS predictions that overlapped only N2-shared genomic sequences but that still failed to overlap N2 protein-coding genes, we disqualified them both by the above criteria and also by the following criteria: overlap with an annotated N2 pseudogene; overlap with a repetitive DNA region annotated with ‘Transposon’; or encoding a protein with the transposon-associated InterPro domain “Transposase, Tc5, C-terminal [IPR007350]”.

For StringTie2 transcript loci, we identified those that at least partially overlapped sequences specific to the CGC1 assembly, and then disqualified them as follows: we excluded any overlap with any lifted-over N2 protein-coding gene; we excluded any that overlapped with a AUGUSTUS-predicted protein-coding gene; we excluded any that overlapped with a lifted-over N2 ncRNA gene; and we excluded any transcripts encoding rRNA motifs from Rfam. For StringTie2 transcript loci that overlapped only N2-shared genomic DNA, we disqualified them by the same criteria, but then also excluded any overlap with a lifted-over N2 pseudogene.

## Supplementary Figure Legends

**Supplementary Figure 1. Alignment of hiCanu non-redundant contigs from CGC1 in the vertical axis to the VC2010 v1 genome in the horizontal axis.** We aligned hiCanu contigs using nucmer in MUMmer4.0.0rc1 and generated the dot plots using mummerplot in MUMmer4.0.0rc1.

**Supplementary Figure 2. Assembly of a complex tandem repeat region in chromosome III (7,856,863-8,031,900). a**, The top two neighboring Nanopore reads, numbered 1 and 2, are laid out to determine their relative positions. The presence of the two reads is corroborated by the bottom three additional Nanopore reads in Nanopore dataset 2. The boundary of the two neighboring tandem repeats was found in each read, which allowed us to superimpose the two boundaries to layout the two reads as illustrated by the red directed edge. **b**, The result of merging the two reads to a single region with two neighboring tandem repeats. **c**, Read coverage of the merged Nanopore reads, Nanopore consensus, and HiFi consensus as in Supplementary Figure 2c for complex tandem repeat regions in chromosome III.

**Supplementary Figure 3. Assembly of two complex tandem repeat regions in chromosome X. a-b**, Dot plots of two Nanopore reads that span two tandem repeats in chromosome X, 4,508,408-4,592,148 (**a**) and 5,581,121-5,673,280 (**b**). **c**, The red box highlights that HiFi reads in the second row have a number of bases that disagree with the first-half of Nanopore consensus (light green); however, Nanopore reads in the 4th row nearly concord with the consensus, suggesting HiFi reads have sequence bias in the first half. Thus, we used the Nanopore consensus (light green) in the first-half and the HiFi consensus (light blue) in the last-half as the final consensus. **d**, The red box emphasizes that HiFi reads in the 2nd row have many bases that are inconsistent with the last half of Nanopore consensus (green) where Nanopore reads in the 3rd row and subreads in the 4th row are nearly consistent, suggesting sequence bias in HiFi reads (see Figure e). **e**, We thus mapped PacBio subreads to the Nanopore consensus, and confirmed the lack of sequence bias in the PacBio subreads and a high accuracy of the Nanopore consensus because the consensus in the subread multiple alignment was occupied by a single base at each column. Specifically, alignments of HiFi reads and subreads to the variant 27-mer unit such that C in the representative is substituted by A. Subreads free from sequence bias support A, but HiFi reads support C, which can be a false-positive call. Thus, we used the Nanopore consensus as the reference sequence in place of the HiFi consensus.

**Supplementary Figure 4. Confirmation of 173 tandem repeat sequences in 152 >5-kb repeat regions (excluding 45S rDNA) in the CGC1 assembly**. The upper shows the self dot plot of the tandem repeat in the region. The lower two tracks show the read coverages of HiFi reads and Nanopore ultralong reads that are aligned with the tandem repeat. Some of 152 regions can have more than one tandem repeat, and therefore, a total of 173 tandem repeats are shown in the figure. Nanopore’s ultralong reads have nearly uniform read coverage but discrepancies are found in some genomic regions due to recurrent sequencing errors. In contrast, the read coverage of HiFi reads can be uneven if the occurrences of tandem repeats are highly similar to each other and HiFi reads can map to different locations, thereby generating mismatches. The genomic region of chrI:952,541-1,043,792 on page 2 shows such an example. Many other example regions are found on pages 10, 13, 14, 16, 30, 38, 50, 52, 54, 55, 56, 57, 60, 62, 66, 70, 72, 77, 80, 81, 85, 86, 89, 92, 104, 113, 116, 118, 119, 121, 126, 127, 130, 132, 134, 135, 136, 138, 140, 144, 147, and 148. The consensus sequence of individual tandem repeat is selected if there is little discrepancy in one of the coverage information. Otherwise, a consensus was generated by manual inspection.

**Supplementary Figure 5. Statistics of tandem repeats (TRs). a-b**. TRs are clustered into two groups: one group has TRs shorter than 5 kb (green), and the other has the remaining longer TRs (yellow). Panel **a** shows the number in each group, while Panel **b** displays the total length of TRs in each group in the respective assemblies of CGC1 and VC2010 (v1). Of note, the longer TR group (length ≥5 kb) dominates the total length of TRs in the new CGC1 assembly. **c**. To highlight that CGC1 has longer TRs than the previous VC2010 assembly (Yoshimura et al, 2019), the histogram shows the length distribution of TRs of length 5 kb or more in the two assemblies. **d-e**. To examine whether TRs in introns dominate the sizes of introns, Panel **d** illustrates an intron with a tandem repeat. Each dot in Panel **e** shows the pair of the length of an intron (x-axis) and the ratio of the total length of TRs in the intron to the intron length (y-axis). Some introns are largely filled with TRs, but a number of introns are almost free from TRs. For example, among introns of length 1 kb or more, 113 have ratio 0.8 or more, while 432 have ratio 0.2 or less. Supplementary Table 18 shows the list of all introns with TRs.

**Supplementary Figure 6. Evolution of 12 regions filled with the 27-mer tandem repeat unit and its variants.** The upper right panel shows the self-dot plot of the tandem repeat at the above locus that are filled with the 27-mer unit and its variants. The table below shows the strings of 27-mer units present in the focal region, their frequencies, mutations, indels, and their names such as type1, type2, and type3 (see their details in Supplementary Table 4). The phylogenetic tree at the left-bottom shows the proximity of variants in terms of sequence similarity. The right bottom shows the locations of unit variants to illustrate how units are distributed inside the tandem repeat. Type 1 units are almost evenly distributed in all regions, while type 2 units are found evenly only in the five regions in the blue box of Figure 3. In addition to prevalent type 1 and type 2 units, another unit and its variants, denoted by type 3 and type 3’ respectively, are enriched in the group of regions in the green box of Figure 3. In addition, each region has its own unique unit variants. The characteristics of each unit represent the evolutionary footprint of an individual region.

**Supplementary Figure 7. Comparison between four CGC1 genomic regions with large tandem repeats, the lengths of which are confirmed by Nanopore ultralong reads, and their corresponding regions in the GALA assembly. a**. A dot plot of the CGC1 assembly region (chrI:4,715,580-4,817,023) and its corresponding region in the GALA assembly. The CGC1 assembly in this region has two TRs of ∼29 kb and ∼72 kb in size; however, the GALA assembly has shorter TRs of ∼18 kb and ∼22 kb in size. **b-c**. Similarly, here are dot plots of the two CGC1 regions, chrII:14,800,218-14,842,635 (**b**) and chrX:441,085-594,508 (**c**), and their corresponding regions in the GALA assembly. In both cases, TRs in CGC1 are longer than those in the GALA assembly. Specifically, in Figure **b**, TRs of CGC1 are of length ∼9 kb and ∼33 kb, while TRs in GALA assembly are ∼5 kb and ∼7 kb in size. In the bottom-right TRs of Figure 1**c**, TRs of CGC1 and GALA assembly are of length ∼153 kb and ∼19 kb, respectively. **d-f**. A dot plot of the CGC1 region (chrV:1,221,871-1,243,359) and its corresponding region in the GALA assembly (**d**). The sizes of the TRs in the two regions are nearly equal, but the white areas show local differences between the two TRs. To show the validity of the CGC1 region, two dot plots are displayed in Figures **e** and **f**, demonstrating that the CGC1 region are consistent with its corresponding regions in the earlier VC2010 assembly of Yoshimura et al. (YOSHIMURA *et al*. 2019) (**e**) and in a Nanopore ultralong read (**f**).

**Supplementary Figure 8. Fragmented assembled contigs in the hifiasm assembly in the 45S rDNA array.** The six assembled contigs generated by hifiasm 0.14 are shown at the bottom, and their approximate positions in the rDNA array of the CGC1 genome assembly are indicated by the dotted lines that link between SNVs specific to representative repeat units of 45S rDNA. The six hifiasm contigs are much shorter than the actual rDNA array.

**Supplementary Figure 9. Dot plot between the pSX1 region of the hifiasm assembly (horizontal axis) and the CGC1 genome assembly (vertical axis).**

**Supplementary Figure 10. Error correction of regions that span the pSX1 and 5S rDNA arrays. a-b**. The top rows show the self dot plots of two Nanopore ultralong reads that span the pSX1 array in chromosome X (441,085-594,508) (**a**) and 5S rDNA array in chromosome V (17,811,675-18,028,889) (**b**). **c-d**. Rows from the top to the bottom show spanning Nanopore ultralong reads (red), read coverage of Nanopore ultralong reads that were aligned with the red spanning reads, the consensus of the aligned Nanopore reads (green), read coverage of PacBio HiFi reads mapped to the Nanopore consensus, read coverage of Nanopore reads to the Nanopore consensus in the right column, the consensus of HiFi reads mapped to the Nanopore consensus (blue), and read coverage of HiFi and Nanopore reads mapped to the HiFi consensus. Nucleotide distribution at each base is displayed.

**Supplementary Figure 11. Evolution in the 5S rDNA array.** Similar to Figure 4, the upper right panel shows the self-dot plot of a region with the 5S rDNA array of length ∼212 kb in chromosome V (nt 17,811,675-18,028,889), the bottom-left table shows characteristics of four frequent variants of the reference 5S rDNA (976 nt, red) including the statistical significance codes in the last column for p-values (***<=0.1%, **<=1%, and *<=5%) such that each variant occurs adjacent to each other according to the Wald–Wolfowitz runs test. On the right side of the table, the colored bars indicate the locations of each variant.

**Supplementary Figure 12. Chromosome V has a 115-kb region (269,089-384,258) homologous to the pSX1 array in chromosome X (441,085-594,508). a**. The dotplot shows that the 115-kb region in the x-axis is spanned by a 152-kb Nanopore ultralong read in the y-axis, which confirms the presence of the 115 kb region. **b**. The dotplot indicates that the 115 kb region in the x-axis is homologous to the pSX1 array in chromosome X (441,085-594,508) in the y-axis. Their surrounding regions however do not match, showing that the two regions are distinct. **c**. Read coverage of Nanopore reads and PacBio HiFi reads mapped to the 115 kb region. The coverage by Nanopore ultralong reads is almost even, showing the long scale correctness of the assembly in this 115-kb region except for a number of erroneous bases that are colored by multiple base calls. These errors are corrected by PacBio HiFi reads.

**Supplementary Figure 13. The pSX1 array in chrV:260,245-414,258.** The upper panel shows characteristics of ten frequent variants of the reference pSX1 (172 nt) in the top: the frequency of each variant, the ratio of frequency to total, number of mismatches with the reference pSX1, percentage match with the reference, positions of base substitutions (e.g., 28A means the base at position 28 is substituted with A), and positions with deletion bases (e.g., 164del shows the base at position 164 is deleted). Blue colored substitutions are found in all of the ten pSX1 variants. Note that the match ratios of the ten variants are lower than 92%. Most substitutions are shared in common among the variants, making it hard to find a variant unique to each variant. The phylogenetic tree on the left side of the table shows the proximity of variants in terms of sequence similarity. In the bottom table, the colored bars indicate the locations of the occurrences of each variant.

**Supplementary Figure 14. Dot plots between pairs of 10 kb genomic sequences taken from the left end (top two rows) and right end (bottom two rows) of each chromosome.** The horizontal axis shows 10 kb sequences from the CGC1 assembly, and the vertical axis shows alternating 10 kb sequences from the VC2010 (Yoshimura et al., 2019) and CGC1 assemblies. The black colored rectangle at the left end or right end of each dot plot displays telomeric repeats. The bottom left self-dot plot displays the 10 kb sequence of the right end of chromosome I, and the many lines parallel to the diagonal show the tail of the rDNA array. The VC2010 assembly lacks telomere repeats at the right ends of chromosomes I and III, and hence dot plots between the missing regions and the corresponding regions in the CGC1 assembly are not displayed in the matrix.

**Supplementary Figure 15. Transcription from tandem repeats from three loci in various tissue types.** Three instances are shown where full-length RNA-seq data was mapped to TRs in the CGC1 assembly and also showed splicing in the genomic vicinity of a TR (demonstrating that the RNA-seq reads represented actual RNA rather than contaminant genomic DNA). For all three instances, the top of the figure shows genome annotations, as follows: N2 denotes structures for protein-coding genes lifted over from N2 to CGC1; AUG denotes de novo predictions of protein-coding genes with AUGUSTUS; ST2 denotes StringTie2 transcripts (which can arise either from mRNA or from ncRNA); and TR denotes tandem repeats of CGC1. Below these genome annotations, detailed alignments of long RNA-seq reads to TRs and their flanking genomic DNA sequences are shown. The first three tracks show alignments of tissue-specific RNA-seq reads from Li et al. (LI *et al*. 2020) (GSE130044, NCBI Gene Expression Omnibus); six further tracks show alignments of tissue-specific RNA-seq reads from Roach et al. (ROACH *et al*. 2020) (PRJEB31791, ENA). Supplementary Figures 15a, 15b, and 15c respectively display RNA-seq alignments with TRs in chrIII:12,198,925-12,210,304, chrX:16,263,528-16,273,490, and chrX:17,980,227-17,991,566. Supplementary Figures 15a-15b have their gene annotations to the same scale as the long RNA-seq alignments, while 15c has its gene annotations to smaller scale, as can be seen by comparing the upper TR annotation to the lower TR boundary lines.

**Supplementary Figure 16. Two ultralong reads (named 1 and 2) from different molecules that span the pSX1 array.** The left figure shows a dot lot of read 1 and an assembled region of the pSX1 array and its surrounding regions. The right is a dot plot of read 2 and the assembled region.

**Supplementary Figure 17. Confirmation of pSX1 variant occurrences in two Nanopore ultralong reads that span the pSX1 array. a**. The three pSX1 variants have single nucleotide variant (SNV) 171A (replacement of C at the 171th position with A) and 51TCT (insertion of TCT at the 51th position) that are shown in the leftmost column. For example, the variant in the top row has both 51TCT and 171A. The top row represents occurrences of the variant by green vertical bars. Occurrences of pSX1 variants are displayed when their complete sequences perfectly match the assembled pSX1 array. It is essential to find perfect matches because a few SNVs and indels are characteristic of each variant and must be aligned perfectly. The concern remains that assembling the reads may remove the necessary SNVs and indels unique to each variant. To ensure reliability of occurrences of a variant in the assembly, one approach would be to search “unassembled” reads for perfect matches (or, closest partial matches) of the variant; however, this task is difficult to accomplish because Nanopore’s ultra-long reads have a sequence error rate of ∼5% (∼8 nt of the 172 nt pSX1 are incorrect). **b**. We instead attempt to locate SNVs and indels unique to the variant, and list perfect matches of a short motif string around a focal SNV (or indel) because a shorter string less likely hit sequencing errors. For example, <u>TCTTCT</u>TCT<u>GAA</u> is such a motif around the insertion 51TCT, where the underlined bases are present in the representative pSX1 sequence in Figure **d**. The top, middle, and bottom rows show occurrence distributions of <u>TCTTCT</u>TCT<u>GAA</u> in the assembly, read 1 and read 2, respectively. The set of all occurrences of two pSX1 variants with 51TCT in Figure “a” are almost consistent with the set of <u>TCTTCT</u>TCT<u>GAA</u>, though the latter set is larger than the former because other less frequent variants with 51TCT are not considered in this analysis. Overall, the occurrences of the short motif in two different reads 1 and 2 show a clear pattern of reproducibility in the occurrences of the two variants with 51TCT in the pSX1 array. **c**. Similarly, we examined short motif <u>GAAAAAAA</u>A<u>A</u> surrounding substitution 171A, and the set of its occurrences are also consistent with the set of the two pSX1 variants with 171A in Figure **a**. **d**. The representative pSX1 sequence of length 172 nt. The two underlined substrings are used for designing short motifs <u>TCTTCT</u>TCT<u>GAA</u> with insertion 51TCT and <u>GAAAAAAA</u>A<u>A</u> with substitution 171A.

**Supplementary Figure 18. Confirmation of pSX1 variant occurrences in two Nanopore ultralong reads that span the pSX1 array. a**. The two pSX1 variants have SNVs 23G and 114A shown in the leftmost column. Vertical bars display occurrences of the two pSX1 variants. **b-c**. Vertical bars show distributions of two motifs <u>TGA</u>G<u>TAT</u> (**b**) and <u>AAT</u>T<u>TAA</u> (**c**) around respective SNVs 23G and 114T. They are almost consistent with distributions of the two pSX1 variants with 23G and 114T in Figure **a**. Bars absent in Figure **a** are found in minor pSX1 variants. **d**. The representative pSX1 sequence of length 172 nt. The two underlined substrings are used for designing short motifs <u>TGA</u>G<u>TAT</u> (in Figure **b**) and <u>AAT</u>T<u>TAA</u> (in Figure **c**) .

**Supplementary Figure 19. Confirmation of pSX1 variant occurrences in two Nanopore ultralong reads that span the pSX1 array. a**. The pSX1 variant had SNVs 58C and 131T in the leftmost column. **b-c**. Vertical bars show distributions of two motifs <u>TCT</u>C<u>AAA</u> (**b**) and <u>ATA</u>T<u>TTT</u> (**c**) around respective SNVs 58C and 131T. **d**. The representative pSX1 sequence of length 172 nt. The two underlined substrings are used for designing short motifs <u>TCT</u>C<u>AAA</u> (in Figure **b**) and <u>ATA</u>T<u>TTT</u> (in Figure **c**) .

**Supplementary Figure 20. Confirmation of pSX1 variant occurrences in two Nanopore ultralong reads that span the pSX1 array. a**. The four pSX1 variants have SNVs 72G, 84T, and 170T in the leftmost column. **b-d**. Vertical bars show distributions of two motifs <u>AAA</u>G<u>TTG</u> (**b**), <u>TTT</u>T<u>TAG</u> (**c**), and <u>AAA</u>T<u>CAT</u> (**d**) around respective SNVs 72G, 84T, and 170T. **d**. The representative pSX1 sequence of length 172 nt. The three underlined substrings are used for designing short motifs <u>AAA</u>G<u>TTG</u> (Figure **b**), <u>TTT</u>T<u>TAG</u> (Figure **c**), and <u>AAA</u>T<u>CAT</u> (Figure **d**) .

**Supplementary Figure 21. Confirmation of pSX1 variant occurrences in Nanopore ultralong reads that span the pSX1 array. a**. The focal pSX1 variant has SNVs 90G, 91A, and 170T in the leftmost column. **b**. Vertical bars show distributions of motif <u>GCT</u>GA<u>GAC</u> around consecutive SNVs 90G and 91A. Although the motif distributions of the assembly and read 1 are consistent, additional motif occurrences are seen in read 2. This discrepancy motivated us to confirm occurrences of <u>GCT</u>GA<u>GAC</u> in HiFi reads that aligned to the region in the assembly. **c**. Distributions of the short motif <u>GCT</u>GA<u>GAC</u> in 88 HiFi reads are largely consistent with the motif distribution in read 1 of Figure b, but do not support the additional motif occurrences in read 2 that seem to be caused sequencing errors in read. **d-f**. To further confirm the distribution of the focal pSX1 variant, we also examine the distributions of motif <u>GCTAGGAC</u> that is absent in the variant but is present in the representative pSX1 and the other variants except for the focal variant. As expected, the distributions of <u>GCTAGGAC</u> in the assembly, read 1 (Figure **d**), and 88 HiFi reads (Figure **e**) complement the distribution of <u>GCT</u>GA<u>GAC</u> in Figure **b**, and nearly occupy the white area where the pSX1 variant does not exist in Figure **a**. Two motifs <u>GCTAGGAC</u> and <u>GCT</u>GA<u>GAC</u> appear less frequently in the red-colored region that actually has many occurrences of another motif <u>GCTA</u>A<u>GAC</u> which replaces G with A at position 91 (Figure **f**). **g**. The representative pSX1 sequence of length 172 nt. The underlined substring is used for designing short motif <u>GCT</u>GA<u>GAC</u> (in Figure **b**).

## Supplementary Table Legends

**Supplementary Table 1.** Lengths and length distribution of long reads.

**Supplementary Table 2.** Statistics of contigs assembled with hiCanu from HiFi reads (excluding Nanopore reads)

**Supplementary Table 3.** Tandem repeats in the CGC1 genome.

**Supplementary Table 4.** Representative units and their variants.

**Supplementary Table 5.** Sequencing the pSX1, 5S rDNA, and 45S rDNA arrays.

**Supplementary Table 6.** Variants in 107 copies of 45S rDNA and their confirmation by RNA-seq data.

**Supplementary Table 7.** Estimation of the 45S rDNA array.

**Supplementary Table 8.** Single nucleotide variants found in CGC1, Bristol N2, CB4856 (Hawaiian strain), ALT1, and ATL2 strains within the 45S rDNA representative repeat unit of size 7197 nt.

**Supplementary Table 9.** Positions and lengths of telomeric repeats in each chromosome.

**Supplementary Table 10.** Allele frequency at the heteroplasmic position of mitochondrial genome.

**Supplementary Table 11.** Annotations of *C. elegans* protein-coding genes originally in the N2 genome (WormBase release WS292). Its data columns are as follows.

**Gene:** A given protein-coding gene, originally annotated in the N2 genome assembly, and lifted over (when possible) into the CGC1 genome assembly. All further data columns are pertinent to that particular gene.

**Max_prot_size:** This lists the size of the largest predicted protein product.

**Prot_size:** This lists the full range of sizes for all protein products from a gene’s predicted isoforms.

**Lifted_to_CGC1:** This shows whether an N2 protein-coding gene could be lifted to the CGC1 genome (“Lifted_to_CGC1”) or not (“Not_lifted”).

**Mapped_CGC1_gene:** This lists *de novo* AUGUSTUS protein-coding gene predictions for which the coding DNA (CDS DNA) exons of a lifted-over N2 gene have at least 1 nucleotide of overlap with the AUGUSTUS gene’s or genes’ CDS DNA exons.

**CGC1_trans_status:** This lists the status of predicted protein (translation) products of the lifted-over N2 genes in the CGC1 genome. “CGC1_ident_trans” indicates that the lifted-over N2 gene encodes at least one protein in the CGC1 genome that is identical to its N2 sequence; “Non_ident_trans” indicates that the lifted-over N2 gene does encode a protein, but does not encode any protein identical to its N2 sequence; “Not_translated” indicates that the lifted-over N2 gene cannot be translated from the CGC1 genome with gffread (PERTEA AND PERTEA 2020).

**Tandem_copies:** This lists protein-coding genes that were mapped by LiftOff (SHUMATE AND SALZBERG 2021) from a single locus in the N2 assembly to two or more tandemly repeated copies in the CGC1 genome, with LiftOff being run to require perfect (100%) identity between tandem repeats. Most N2 protein-coding genes that could be mapped at all were mapped to a unique locus in the CGC1 assembly, and thus are blank in this annotation column. Genes that were mapped to two or more CGC1 tandem loci were annotated with “Tandem_copies[x2]”, “Tandem_copies[x3]”, “Tandem_copies[x4]”, or “Tandem_copies[x7]”.

**Novel_CGC1_exon:** This lists *de novo* AUGUSTUS protein-coding gene predictions for which the coding DNA (CDS DNA) exons of a lifted-over N2 assembly gene have at least 1 nucleotide of overlap with the AUGUSTUS gene’s or genes’ CDS DNA exons, and for which the AUGUSTUS gene also has at least one CDS DNA exon that falls into novel regions of the CGC1 assembly (i.e., genome regions not uniquely mappable from the N2 genome) and that also does not overlap lifted-over protein-coding genes. When such an overlap occurs, the most parsimonious explanation is that a gene annotated in the earlier assembly can be lifted to CGC1, but that reannotating it in the CGC1 genome will reveal at least one CDS DNA exon that could not be predicted in the earlier N2 assembly (because its genomic DNA was absent from the N2 genome assembly).

**StringTie2_overlaps:** This lists transcripts (either protein- or ncRNA-coding) predicted in the CGC1 genome by StringTie2 (KOVAKA *et al*. 2019) from ribodepleted *C. elegans* RNA-seq data that have at least 1 nucleotide of overlap with CDS DNA exons of a given lifted-over protein-coding gene.

**Mass_spec:** This indicates whether protein sequences predicted to be encoded by this gene (pre-liftover) were detectable in spectra from published mass spectrometric *C. elegans* proteomics data (denoted with “Mass_spec”).

**Pfam:** This lists predicted protein domains from Pfam 37.0 (MISTRY *et al*. 2021), detected with family-specific significance thresholds, that are encoded by the gene’s protein product(s).

**InterPro:** This lists predicted protein domains from InterProScan 5.57-90.0 (PAYSAN-LAFOSSE *et al*. 2023) that are encoded by the gene’s protein product(s).

**OFind_Summary** and **OFind_Full:** This lists results for our OrthoFinder analysis (EMMS AND KELLY 2019) of orthologies between protein-coding genes from the *C. elegans* CGC1 genome (both *de novo* AUGUSTUS and lifted-over N2 genes), protein-coding genes from *Caenorhabditis nigoni* (YIN *et al*. 2018) and *Caenorhabditis remanei* (TETERINA *et al*. 2020), and protein sequences encoded by *C. elegans* transposons in WormBase WS292 (STERNBERG *et al*. 2024). Two different views of these results are given: the summary lists taxa and gene counts, while the full results give individual gene names.

**Supplementary Table 12.** Annotations of *C. elegans* protein-coding genes predicted *de novo* in the CGC1 genome by AUGUSTUS. Its data columns are as follows.

**Gene:** A given protein-coding gene predicted *de novo* in the CGC1 genome assembly. All further data columns are pertinent to that particular gene.

**Gen_DNA_overlap:** This lists whether a gene’s CDS DNA exons exclusively overlap CGC1 genomic DNA that is shared with the N2 assembly (“N2_gen_DNA”), exclusively overlap CGC1 sequences that are unique to the CGC1 assembly (“CGC1_gen_DNA”), or overlap both assemblies (“CGC1_gen_DNA; N2_gen_DNA”). AUGUSTUS predictions with “CGC1_gen_DNA” exonic annotations and that lack N2 (WS292) gene equivalents represent possible new protein-coding genes at least partially arising from CGC1-assembly-specific DNA.

**N2_gene_equiv:** This lists possible N2 protein-coding gene equivalents: i.e., lifted-over N2 (WS292) protein-coding genes for which the coding DNA (CDS DNA) exons of an AUGUSTUS gene have at least 1 nucleotide of overlap with the N2 (WS292) gene’s or genes’ CDS DNA exons. AUGUSTUS predictions with N2 equivalents are likely to be repredictions of those protein-coding genes.

**StringTie2_overlaps:** This lists *de novo* AUGUSTUS genes whose CDS DNA exons have at least 1 nucleotide of overlap with a transcript (either protein- or ncRNA-coding) predicted in the CGC1 assembly by StringTie2 (KOVAKA *et al*. 2019) from ribodepleted *C. elegans* RNA-seq data. Overlaps with StringTie2 transcripts provide evidence for the RNA-seq expression (and thus, the reality) of a AUGUSTUS prediction.

**Mass_spec:** This indicates whether protein sequences predicted to be encoded by this gene were detectable in spectra from published mass spectrometric *C. elegans* proteomics data (denoted with “Mass_spec”). Matches to proteomics spectra provide evidence for the translation (and thus, the reality) of a AUGUSTUS prediction.

**N2_pseudogene:** This lists possible N2 pseudogene equivalents: i.e., lifted-over N2 pseudogenes for which the coding DNA (CDS DNA) exons of an AUGUSTUS gene have at least 1 nucleotide of overlap with the N2 pseudogene’s or pseudogenes’ exons. Overlaps of this type are particularly relevant for AUGUSTUS predictions falling exclusively into N2-shared genomic DNA, because they suggest that the predictions in fact represent pseudogenes rather than protein-coding genes.

**Rep.DNA_overlap:** This lists overlaps (of at least 1 nucleotide) between an AUGUSTUS gene’s CDS DNA exons and repetitive DNA regions annotated in N2 (from WormBase WS292) and lifted over to CGC1. Overlaps of this type may disqualify AUGUSTUS predictions as being coding sequences of repetitive DNA rather than endogeneous protein-coding genes.

**TR_overlap:** This lists overlaps (of at least 1 nucleotide) between an AUGUSTUS gene’s CDS DNA exons and tandemly repeated (TR) regions annotated in CGC1 with mTR (MORISHITA *et al*. 2021). Overlaps of this type may disqualify AUGUSTUS predictions as overlapping TRs rather than being endogeneous protein-coding genes.

**Phobius:** This denotes predictions of signal and transmembrane sequences made with Phobius 1.01 (KÄLL *et al*. 2004) . ‘SigP’ indicates a predicted signal sequence, and ‘TM’ indicates one or more transmembrane-spanning helices, with N helices indicated with ‘(Nx)’. Varying predictions from different isoforms are listed.

**Max_prot_size:** The size of the largest predicted protein product.

**Prot_size:** This shows the full range of sizes for all protein products from a gene’s predicted isoforms.

**NCoils:** This shows coiled-coil domains, predicted by Ncoils 2002.08.22 (LUPAS 1996). Both the proportion of such sequence (ranging from 0.01 to 1.00) and the exact ratio of coiled residues to total residues are given. Proteins with no predicted coiled residues are blank.

**Psegs:** This shows what fraction of a protein is low-complexity sequence, as detected by PSEG 1999.06.10 (WOOTTON 1994). As with Ncoils, relative and absolute fractions of low-complexity residues are shown

**Pseg_val:** This gives the numerical value only from “Psegs”. This column enables easy searches for putative protein-coding genes whose Pseg fractions are suspiciously high, and may reflect mispredictions from repetitive genomic DNA.

**Pfam:** This lists predicted protein domains from Pfam 37.0 (MISTRY *et al*. 2021), detected with family-specific significance thresholds, that are encoded by the gene’s protein product(s).

**InterPro:** This lists predicted protein domains from InterProScan 5.57-90.0 (PAYSAN-LAFOSSE *et al*. 2023) that are encoded by the gene’s protein product(s).

**OFind_Summary** and **OFind_Full:** Results for our OrthoFinder analysis (EMMS AND KELLY 2019) of orthologies between protein-coding genes from the *C. elegans* CGC1 genome (both *de novo* AUGUSTUS and lifted-over N2 genes), protein-coding genes from *Caenorhabditis nigoni* (YIN *et al*. 2018) and *Caenorhabditis remanei* (TETERINA *et al*. 2020), and protein sequences encoded by *C. elegans* transposons in WormBase WS292 (STERNBERG *et al*. 2024). Two different views of these results are given: the summary lists taxa and gene counts, while the full results give individual gene names.

**Supplementary Table 13.** Annotations of *C. elegans* transcript loci (from both protein-coding and ncRNA genes) predicted *de novo* in the CGC1 assembly by StringTie2. Its data columns are as follows.

**StringTie2 genomic locus:** This lists predicted StringTie2 transcripts that map to a unique locus in the CGC1 assembly. All further data columns are pertinent to that particular locus.

**N2_genDNA:** This lists blocks of genomic DNA that overlap the StringTie2 genomic locus by at least 1 nucleotide and that were uniquely mapped from the N2 genome (i.e., blocks of N2-shared sequence in the CGC1 assembly). Such overlaps indicate a StringTie2 transcript locus that is either fully or partially encoded by genomic DNA already present in the N2 assembly.

**CGC1_genDNA:** This lists blocks of genomic DNA that overlap the StringTie2 genomic locus by at least 1 nucleotide and that were not uniquely mappable from the N2 genome (i.e., blocks of sequence specific to the CGC1 assembly). Such overlaps indicate a StringTie2 transcript locus that is either fully or partially encoded by sequences only present in the CGC1 assembly.

**N2_protcod:** This lists protein-coding genes lifted over from N2 to CGC1 whose CDS DNA exons overlap the StringTie2 genomic locus by at least 1 nucleotide. Such overlaps indicate a StringTie2 transcript locus that may arise from the N2 protein-coding gene.

**CGC1.AUG_protcod:** This lists protein-coding genes predicted *de novo* in CGC1 with AUGUSTUS whose CDS DNA exons overlap the StringTie2 genomic locus by at least 1 nucleotide. Such overlaps indicate a StringTie2 transcript locus that may arise from the CGC1 AUGUSTUS protein-coding gene.

**N2_ncRNA:** This lists ncRNA genes lifted over from N2 to CGC1 whose CDS DNA exons overlap the StringTie2 genomic locus by at least 1 nucleotide. Such overlaps indicate a StringTie2 transcript locus that may arise from the N2 ncRNA gene.

**N2_pseudogene:** This lists possible N2 pseudogene equivalents: i.e., lifted-over N2 pseudogenes for which the StringTie2 genomic locus has at least 1 nucleotide of overlap with the N2 pseudogene’s or pseudogenes’ exons. Overlaps of this type are particularly relevant for StringTie2 predictions falling exclusively into N2 -shared genomic DNA, because they suggest that the predictions in fact represent pseudogenes rather than protein-coding genes.

**Rfam:** This lists predicted ncRNA motifs from Rfam 14.10 (KALVARI *et al*. 2021), detected with family-specific significance thresholds, that are encoded by one or more of the StringTie2 transcripts mapped to this locus.

**Supplementary Table 14.** A set of 163 StringTie2 transcript loci that may be novel ncRNA genes at least partially encoded by CGC1-unique regions of the CGC1 assembly. They are a subset of the transcript loci in Supplementary Table 13, and have identical data columns.

**Supplementary Table 15.** A set of 183 genes predicted by AUGUSTUS that may be novel protein-coding genes at least partially encoded by CGC1-unique regions of the CGC1 assembly. They are a subset of the protein-coding genes in Supplementary Table 12, and have identical data columns.

**Supplementary Table 16.** A set of 314 genes predicted by AUGUSTUS that may be novel protein-coding genes despite being fully encoded by N2 -shared regions of the CGC1 assembly. They are a subset of the protein-coding genes in Supplementary Table 12, and have identical data columns.

**Supplementary Table 17.** A set of 3,300 StringTie2 transcript loci that may be novel ncRNA genes despite being fully encoded by N2-shared regions of the CGC1 assembly. They are a subset of the transcript loci in Supplementary Table 13, and have identical data columns.

**Supplementary Table 18.** List of introns with tandem repeats.

**Supplementary Table 19.** Software used in this study. “Software” provides the name of each computer program or suite of computer programs; “Purpose” describes why this software was used here; “Main web site (URL) or code location” gives the primary Web site for this software; “Bioconda source (if used)” gives the bioconda web site for programs that were installed as bioconda environments; “Online documentation” gives the web site for detailed online manuals, if a given program has one. Publications and arguments for each program are cited and described in Methods.

**Supplementary Table 20.** Published genomic data used in this study. Genome sequences, coding sequences, and proteomes of relevant nematodes were downloaded from WormBase (DAVIS et al. 2022) or ParaSite (HOWE et al. 2017; LEE et al. 2017) and used for transcriptomic or proteomic analyses. Four separate spreadsheets are given for data files of genomes, proteomes, CDS DNA sets, and gene annotations in GFF format. For each data file, “Species” gives the biological species described by the file, “Comments” describes the particular use(s) to which the data file was put in this study, and “URL” gives the Web source of the file.

## Supplementary Data Files

**Supplementary Data File 1.** GFF3 file of tandem repeat (TR) coordinates in the CGC1 assembly. TR coordinates here and in Supplementary Data Files 2-3 were computed with mTR (MORISHITA *et al*. 2021).

**Supplementary Data File 2.** GFF3 file of tandem repeat (TR) coordinates in the VC2010 assembly (Yoshimura et al., 2019), for comparison with CGC1.

**Supplementary Data File 3.** GFF3 file of tandem repeat (TR) coordinates computed in the N2 assembly, for comparison with CGC1.

**Supplementary Data File 4.** GTF file of N2 protein-coding genes lifted-over from the WS292 release of the N2 genome into the CGC1 assembly.

**Supplementary Data File 5.** GTF file of N2 ncRNA genes lifted-over from the WS292 release of the N2 genome into the CGC1 assembly.

**Supplementary Data File 6.** GFF3 file of *de novo* AUGUSTUS protein-coding gene predictions in the CGC1 assembly.

**Supplementary Data File 7.** GTF file of one set of StringTie2 transcripts predicted in the CGC1 assembly.

**Supplementary Data File 8.** GTF file of another set of StringTie2 transcripts predicted in the CGC1 assembly.

**Supplementary Data File 9.** BED file of StringTie2 exons predicted in the CGC1 assembly, extracted from Supplementary Data Files 4 and 5.

**Supplementary Data File 10.** GTF file of N2 pseudogenes lifted-over from the WS292 release of the N2 genome into the CGC1 assembly.

**Supplementary Data File 11.** BED file of N2 genomic sequence blocks mapped uniquely to the CGC1 assembly.

**Supplementary Data File 12.** BED file of CGC1-specific sequence blocks (without uniquely mapped N2 alignments) in the CGC1 assembly.

