## Supplementary Figure 1 for "CGC1, a new reference genome for *Caenorhabditis elegans*"

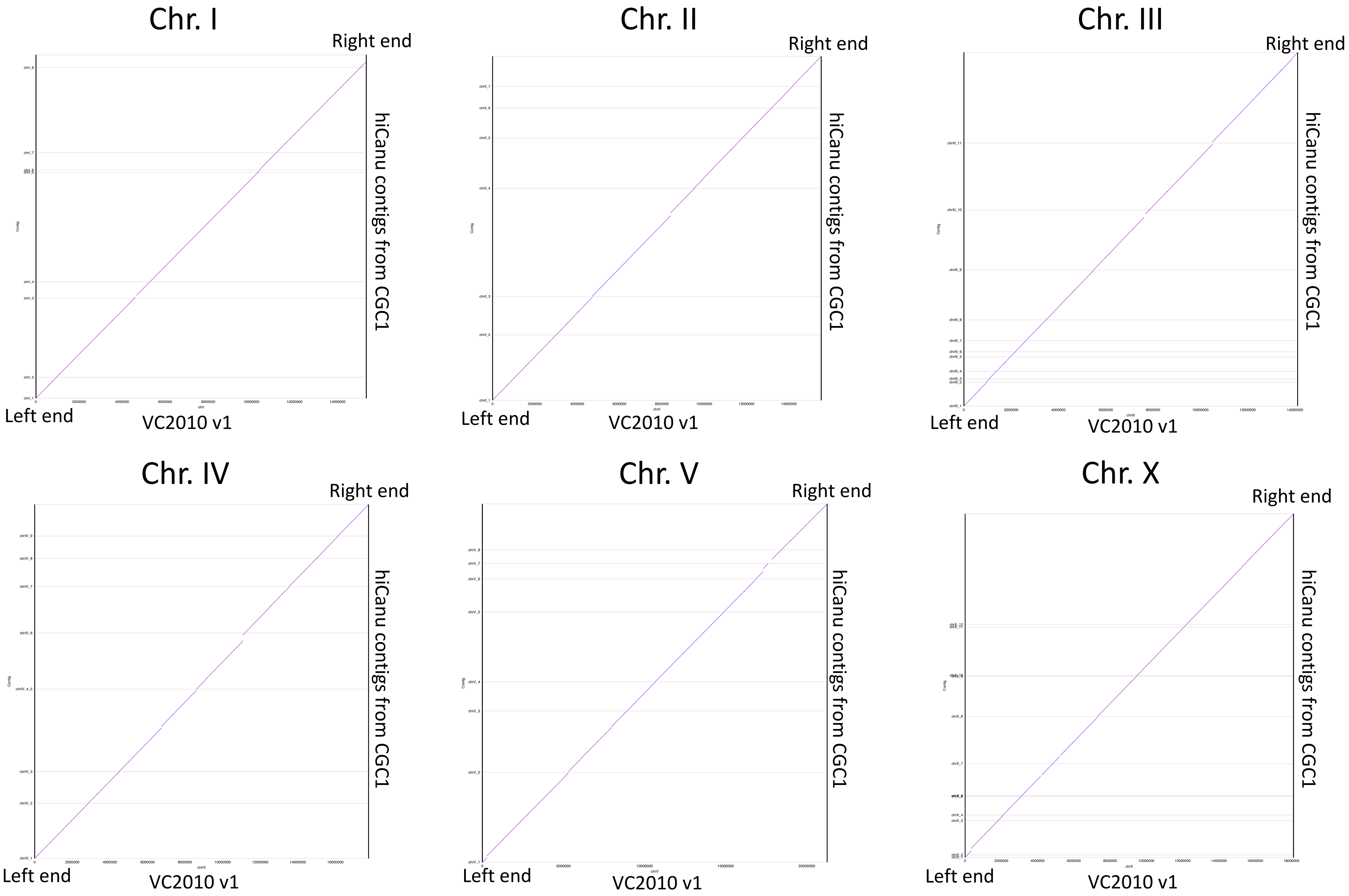


**Supplementary Figure 1. Alignment of hiCanu non-redundant contigs from CGC1 in the vertical axis to the VC2010 v1 genome in the horizontal axis.** We aligned hiCanu contigs using nucmer in MUMmer4.0.0rc1 and generated the dot plots using mummerplot in MUMmer4.0.0rc1.
