## Supplementary Figure 2 for "CGC1, a new reference genome for *Caenorhabditis elegans*"

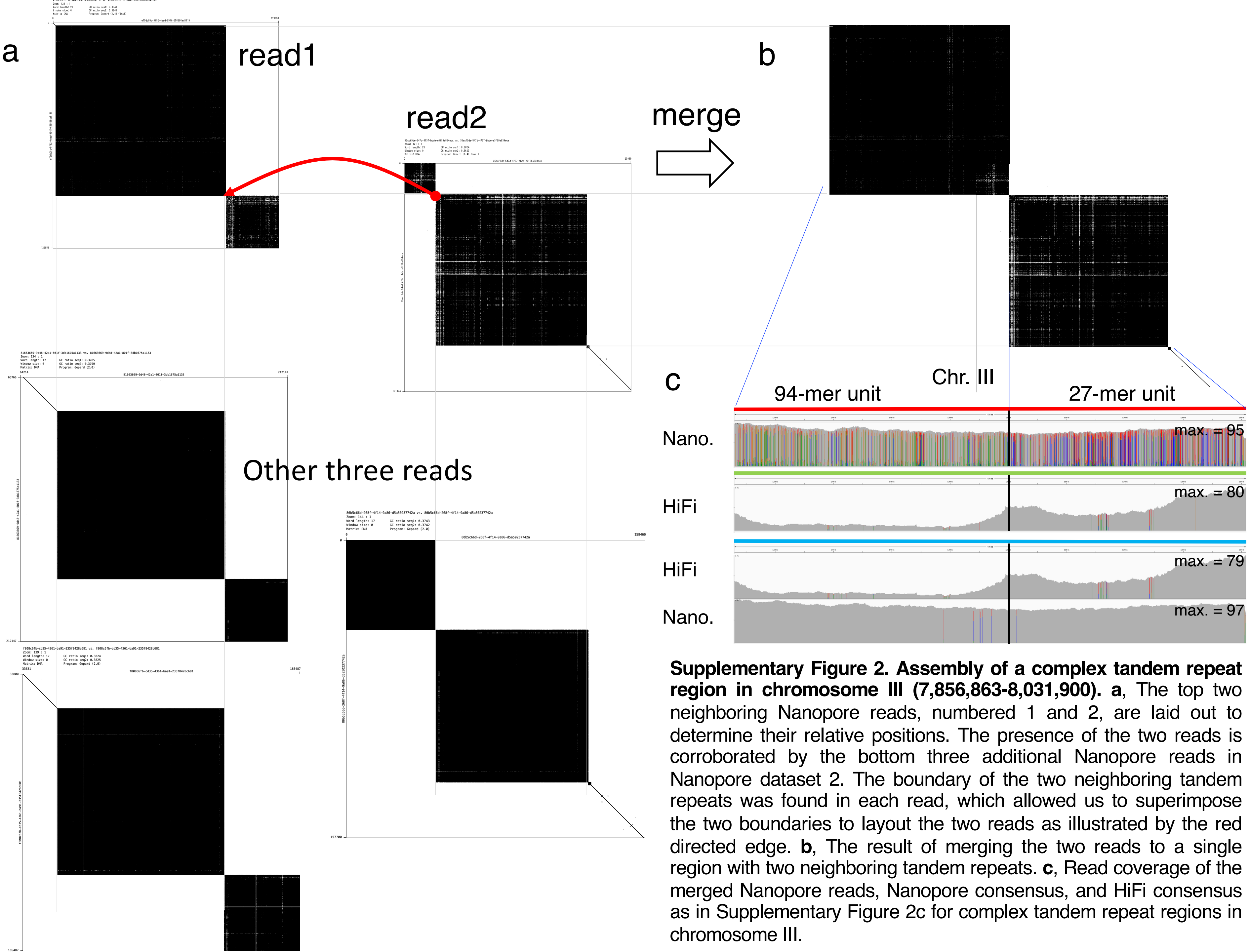


**Supplementary Figure 2. Assembly of a complex tandem repeat region in chromosome III (7,856,863-8,031,900). a**, The top two neighboring Nanopore reads, numbered 1 and 2, are laid out to determine their relative positions. The presence of the two reads is corroborated by the bottom three additional Nanopore reads in Nanopore dataset 2. The boundary of the two neighboring tandem repeats was found in each read, which allowed us to superimpose the two boundaries to layout the two reads as illustrated by the red directed edge. **b**, The result of merging the two reads to a single region with two neighboring tandem repeats. **c**, Read coverage of the merged Nanopore reads, Nanopore consensus, and HiFi consensus as in Supplementary Figure 2c for complex tandem repeat regions in chromosome III.
