## Supplementary Figure 3 for "CGC1, a new reference genome for *Caenorhabditis elegans*"

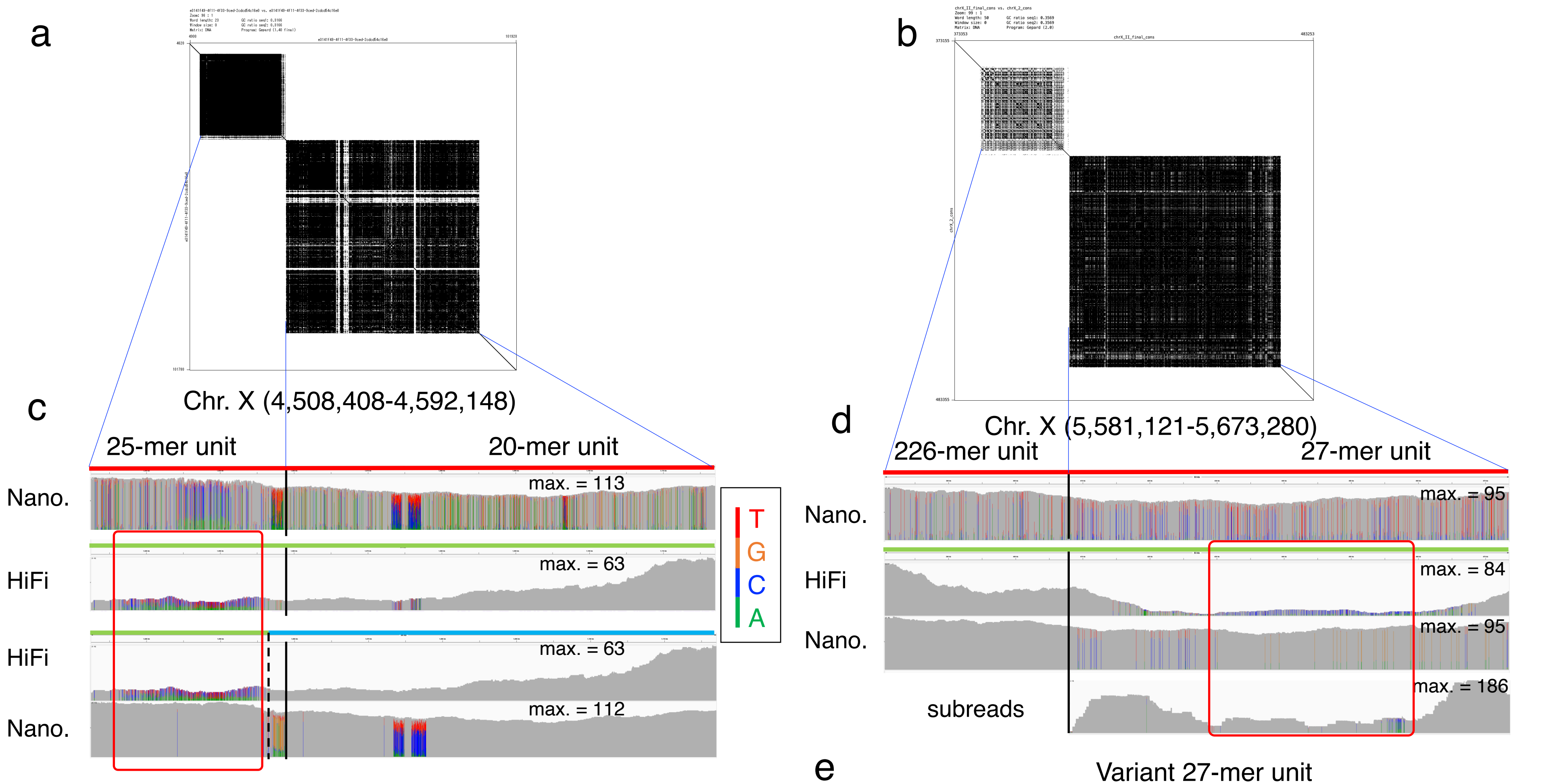


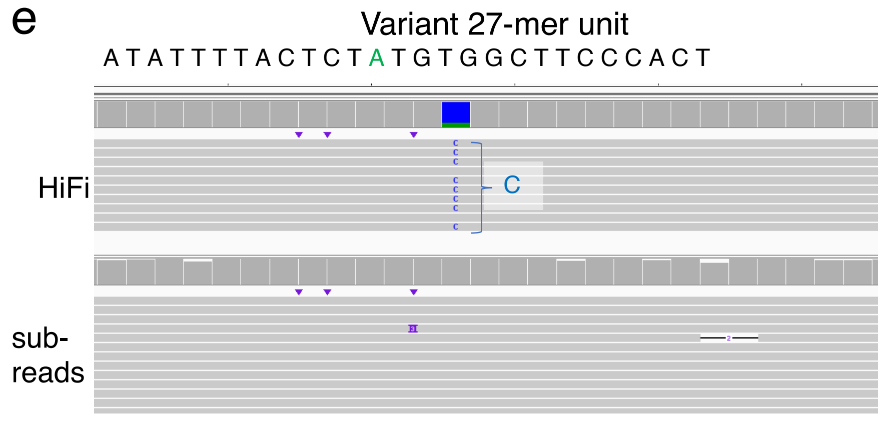


**Supplementary Figure 3. Assembly of two complex tandem repeat regions in chromosome X.** **a-b**, Dot plots of two Nanopore reads that span two tandem repeats in chromosome X, 4,508,408-4,592,148 (**a**) and 5,581,121-5,673,280 (**b**). **c**, The red box highlights that HiFi reads in the second row have a number of bases that disagree with the first-half of Nanopore consensus (green); however, Nanopore reads in the 4th row nearly concord with the consensus, suggesting HiFi reads have sequence bias in the first half. Thus, we used the Nanopore consensus (green) in the first-half and the HiFi consensus (blue) in the last-half as the final consensus. **d**, The red box emphasizes that HiFi reads in the 2nd row have many bases that are inconsistent with the last half of Nanopore consensus (green) where Nanopore reads in the 3rd row and subreads in the 4th row are nearly consistent, suggesting sequence bias in HiFi reads (see Figure e). **e**, We thus mapped PacBio subreads to the Nanopore consensus, and confirmed the lack of sequence bias in the PacBio subreads and a high accuracy of the Nanopore consensus because the consensus in the subread multiple alignment was occupied by a single base at each column. Specifically, alignments of HiFi reads and subreads to the variant 27-mer unit such that C in the representative is substituted by A. Subreads free from sequence bias support A, but HiFi reads support C, which can be a false-positive call. Thus, we used the Nanopore consensus as the reference sequence in place of the HiFi consensus.
