## Supplementary Figure 4 for "CGC1, a new reference genome for *Caenorhabditis elegans*"

### chrI:673,252-690,349

**Supplementary Figure 4 . Confirmation of 173 tandem repeat sequences in 152 >5-kb repeat regions (excluding 45S rDNA) in the CGC1 assembly**. The upper shows the self dot plot of the tandem repeat in the region. The lower two tracks show the read coverages of HiFi reads and Nanopore ultralong reads that are aligned with the tandem repeat. Some of 152 regions can have more than one tandem repeat, and therefore, a total of 173 tandem repeats are shown in the figure. Nanopore's ultralong reads have nearly uniform read coverage but discrepancies are found in some genomic regions due to recurrent sequencing errors. In contrast, the read coverage of HiFi reads can be uneven if the occurrences of tandem repeats are highly similar to each other and HiFi reads can map to different locations, thereby generating mismatches. The genomic region of chrI:952,541-1,043,792 on page 2 shows such an example. Many other example regions are found on pages 10, 13, 14, 16, 30, 38, 50, 52, 54, 55, 56, 57, 60, 62, 66, 70, 72, 77, 80, 81, 85, 86, 89, 92, 104, 113, 116, 118, 119, 121, 126, 127, 130, 132, 134, 135, 136, 138, 140, 144, 147, and 148. The consensus sequence of individual tandem repeat is selected if there is little discrepancy in one of the coverage information. Otherwise, a consensus was generated by manual inspection.

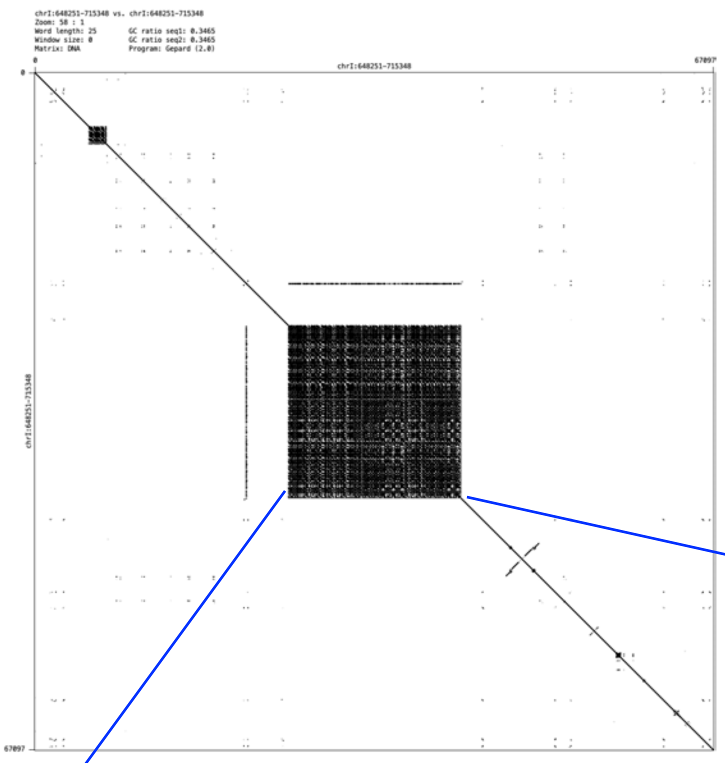


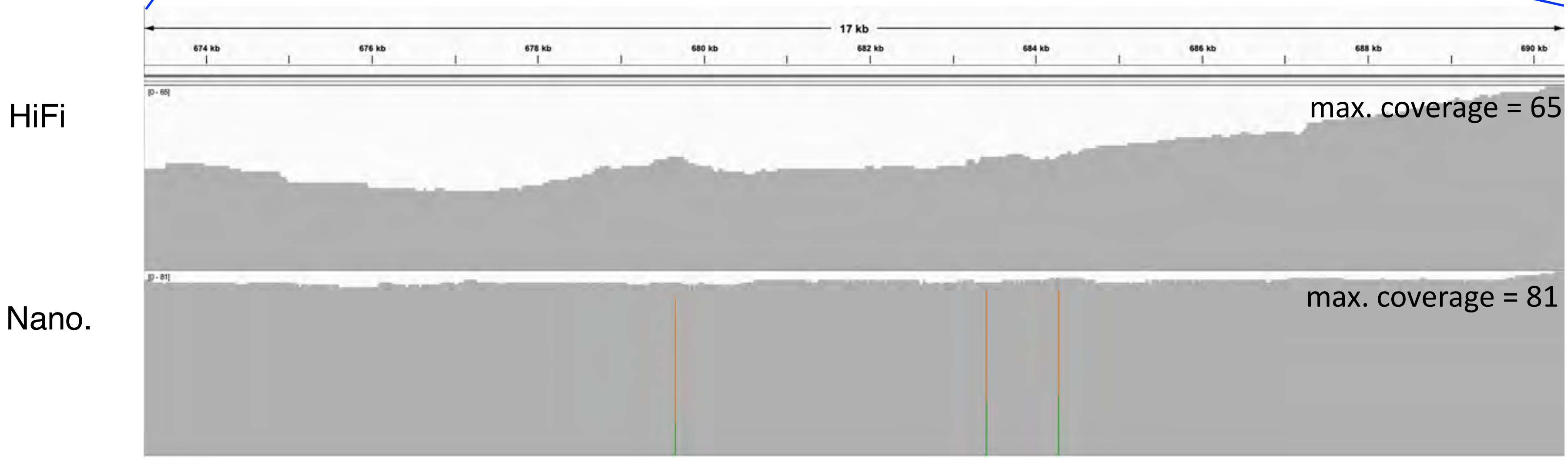

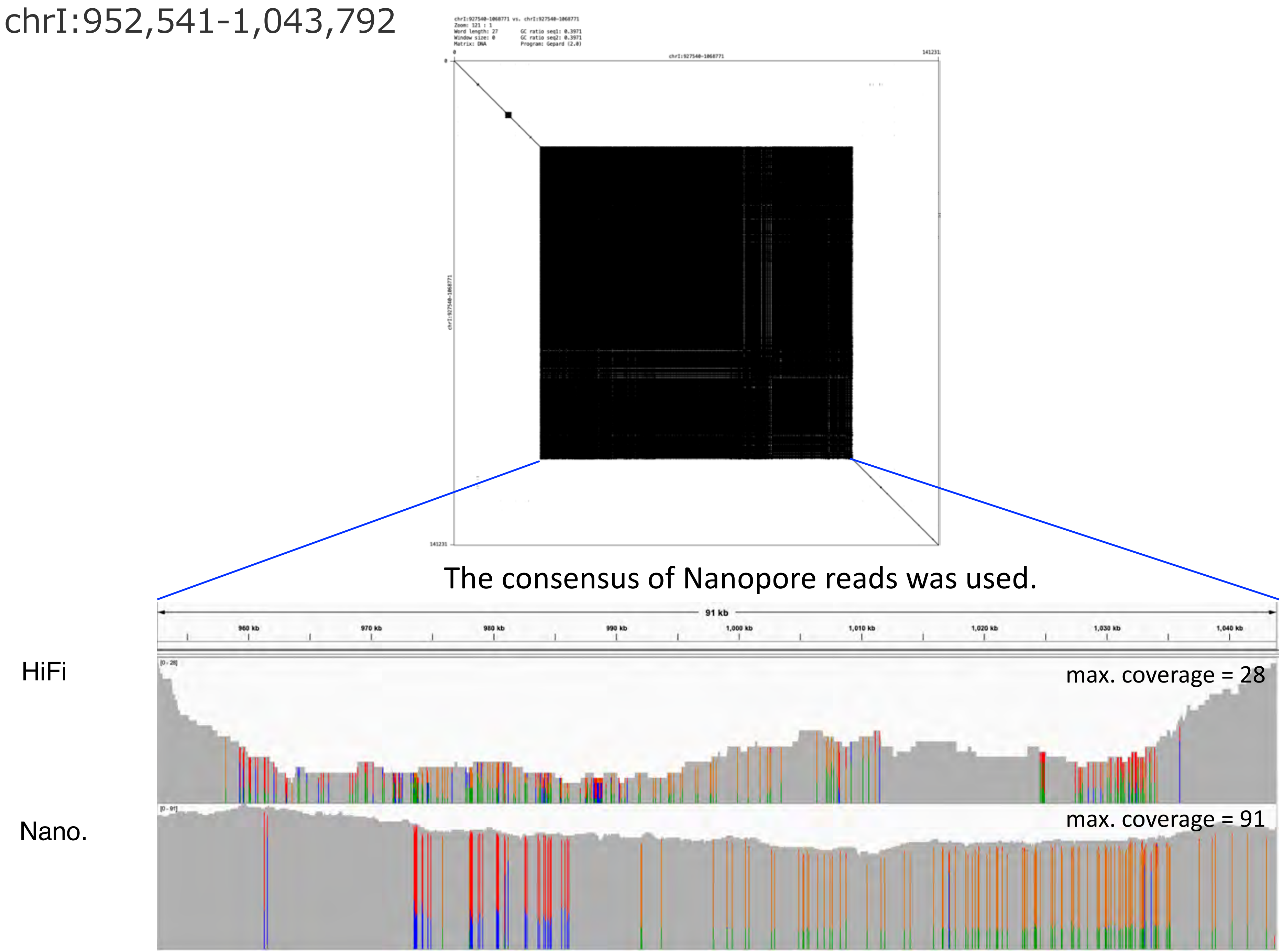
chrI:952,541-1,043,792
The consensus of Nanopore reads was used.
91 kb
960 kb
970 kb
980 kb
990 kb
1,000 kb
1,010 kb
1,020 kb
1,030 kb
1,040 kb
HiFi
max. coverage = 28
Nano.
max. coverage = 91

### chrI:1,227,303-1,233,212

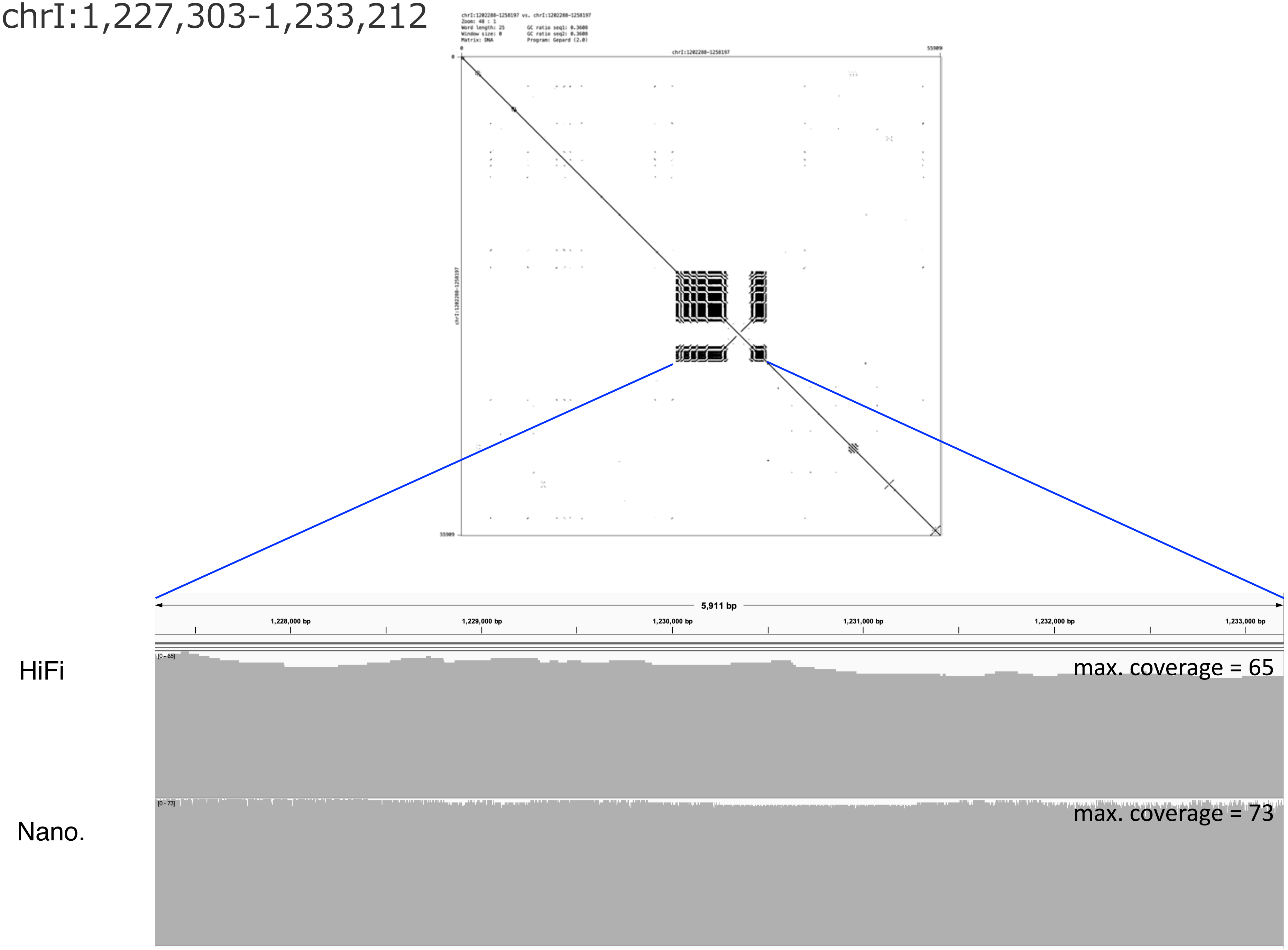

### chrI:2,052,110-2,058,866

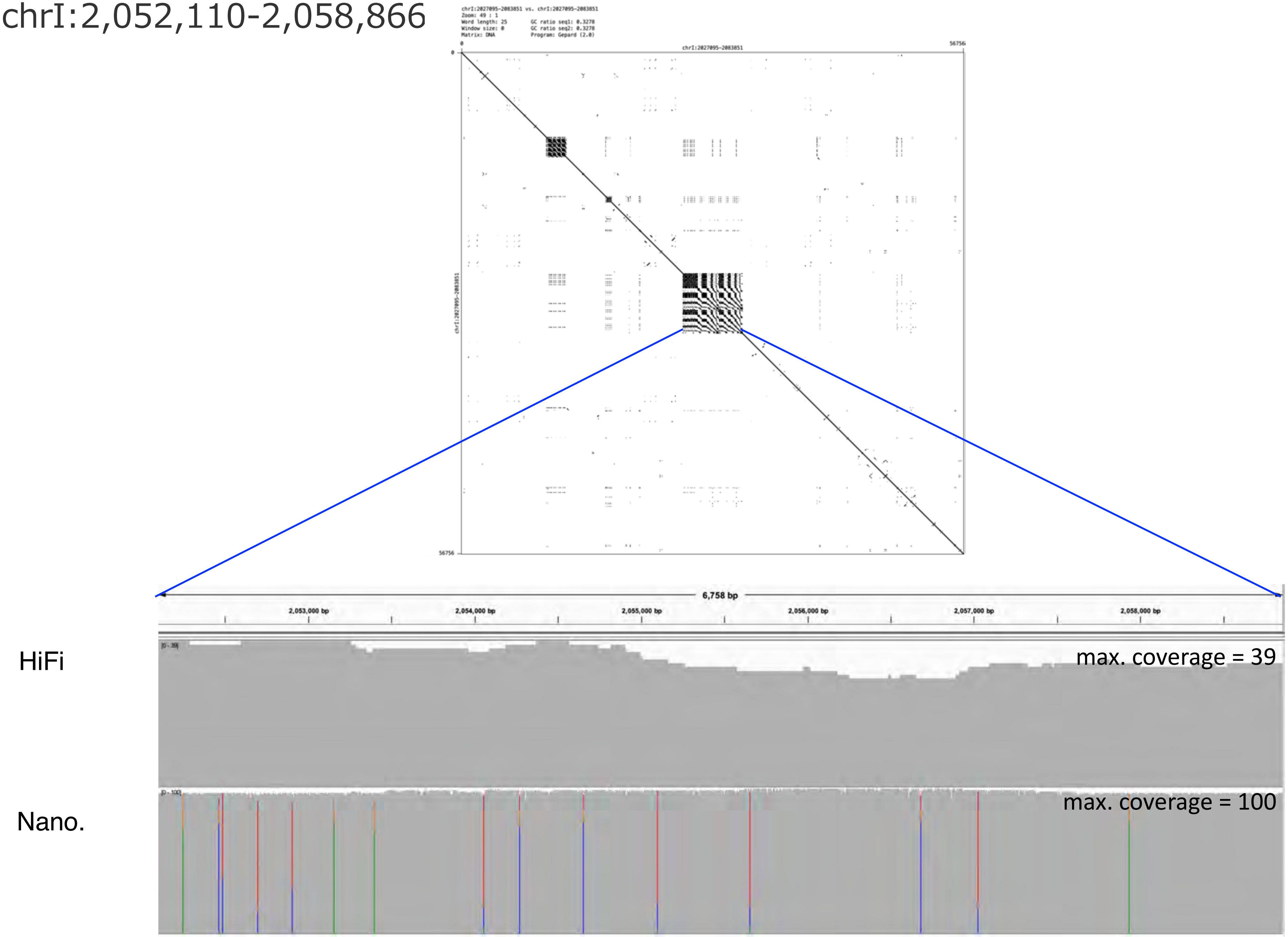

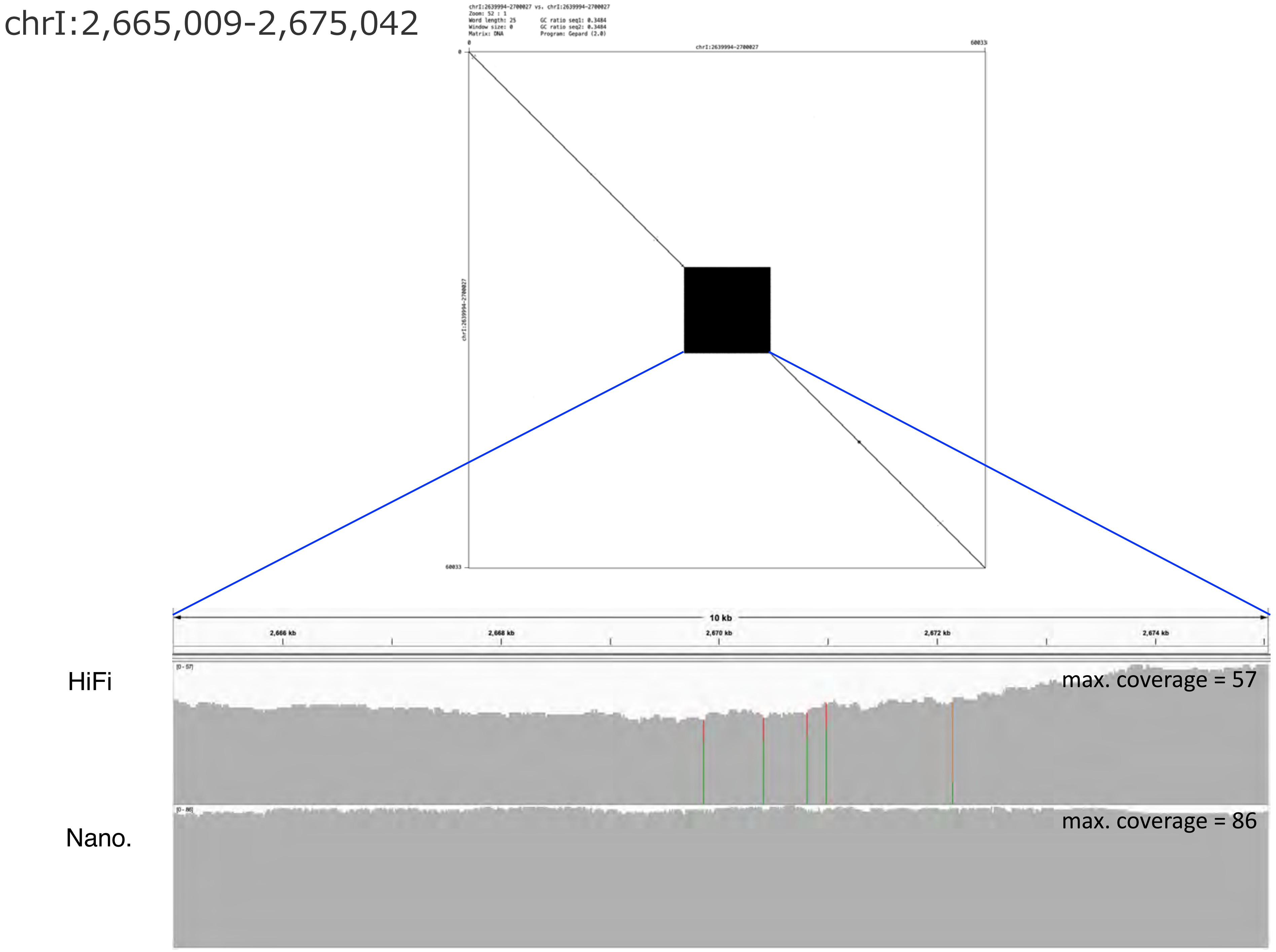
chrI:2,665,009-2,675,042
chrI:2639994-2700027 vs. chrI:2639994-2700027
Zoom: 52 : 1
Word length: 25
GC ratio seq1: 0.3484
Window size: 0
GC ratio seq2: 0.3484
Matrix: DNA
Program: Gepard (2.0)
0
chrI:2639994-2700027
60033
10 kb
2,666 kb
2,668 kb
2,670 kb
2,672 kb
2,674 kb
HiFi
[0 - 57]
max. coverage = 57
Nano.
[0 - 86]
max. coverage = 86

chrI:3,801,708-3,807,986

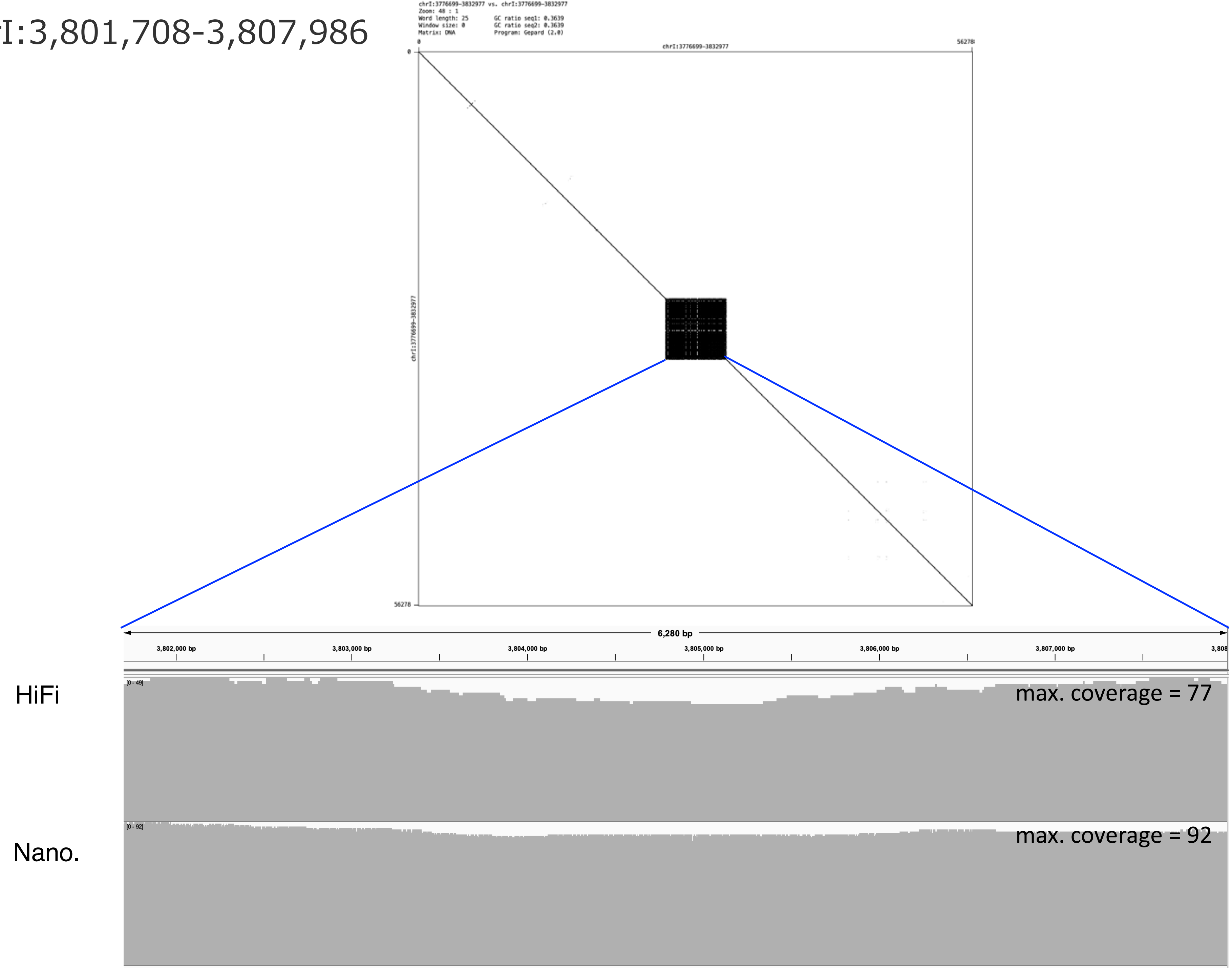

chrI:4,132,423-4,145,136

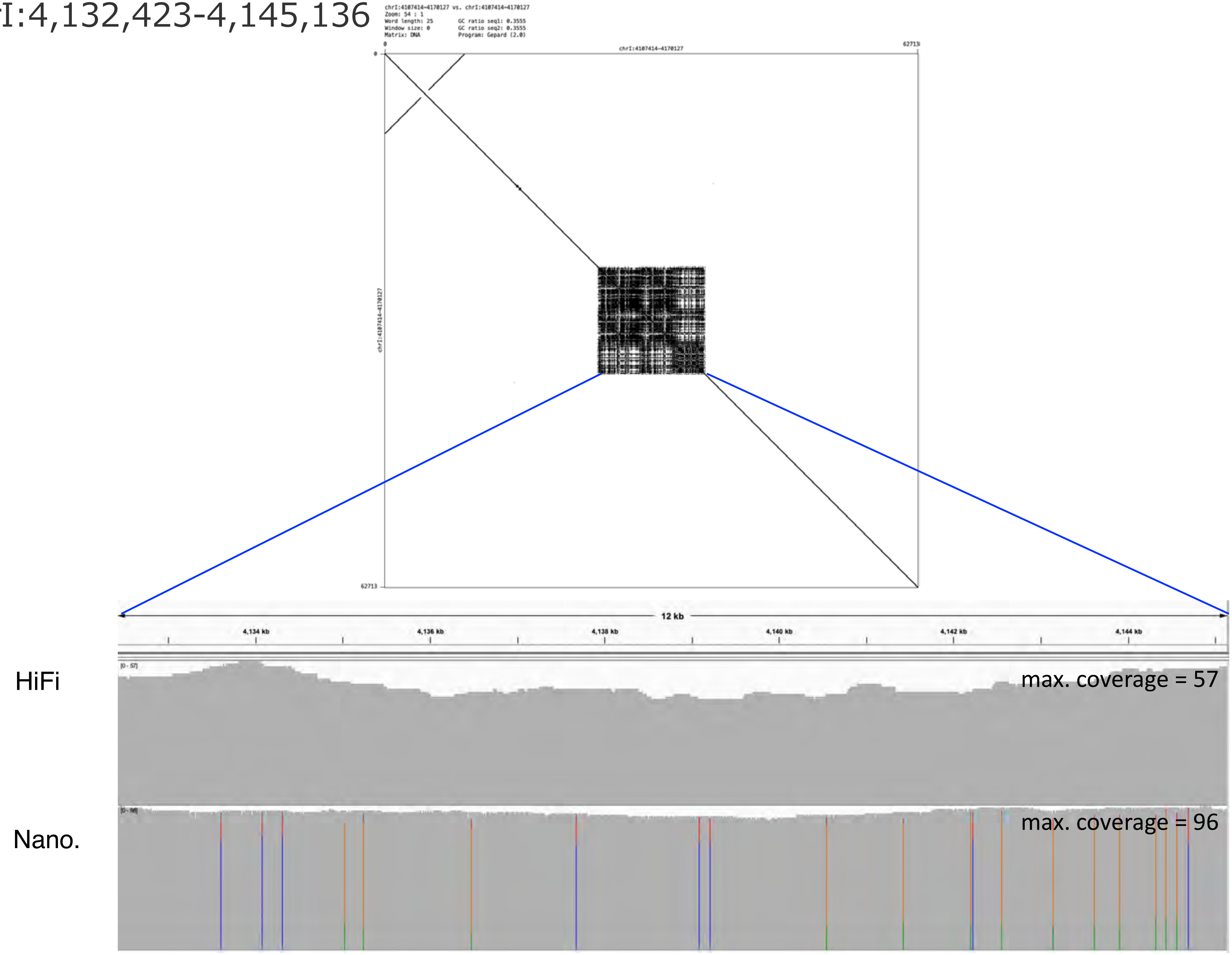

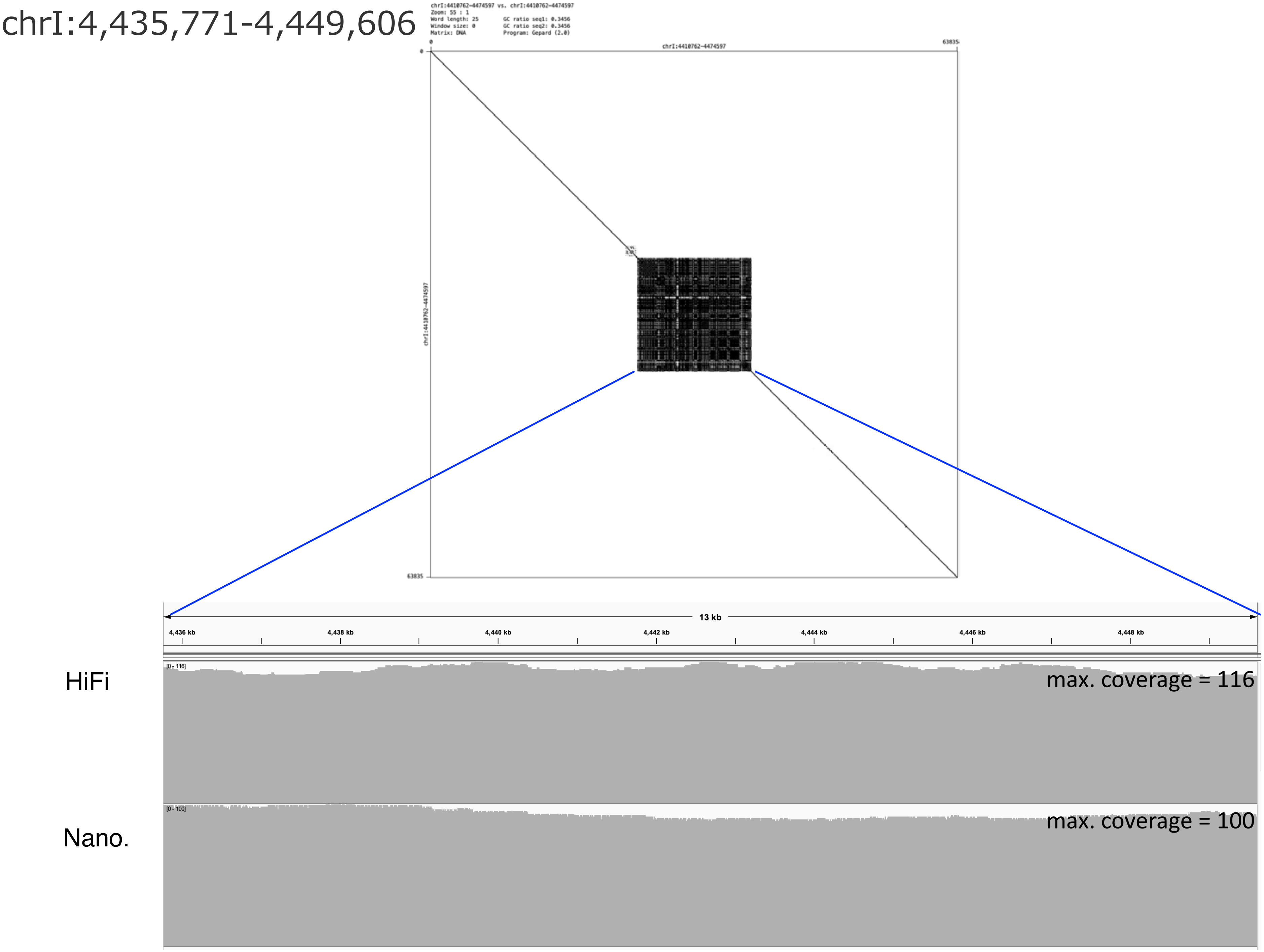

chrI:4,435,771-4,449,606
chrI:4410762-4474597 vs. chrI:4410762-4474597
Zoom: 55 : 1
Word length: 25
Window size: 0
Matrix: DNA
GC ratio seq1: 0.3456
GC ratio seq2: 0.3456
Program: Gepard (2.0)
chrI:4410762-4474597
0
63835
13 kb
4,436 kb
4,438 kb
4,440 kb
4,442 kb
4,444 kb
4,446 kb
4,448 kb
HiFi
[0 - 116]
max. coverage = 116
Nano.
[0 - 100]
max. coverage = 100

chrI:4,715,580-4,817,023

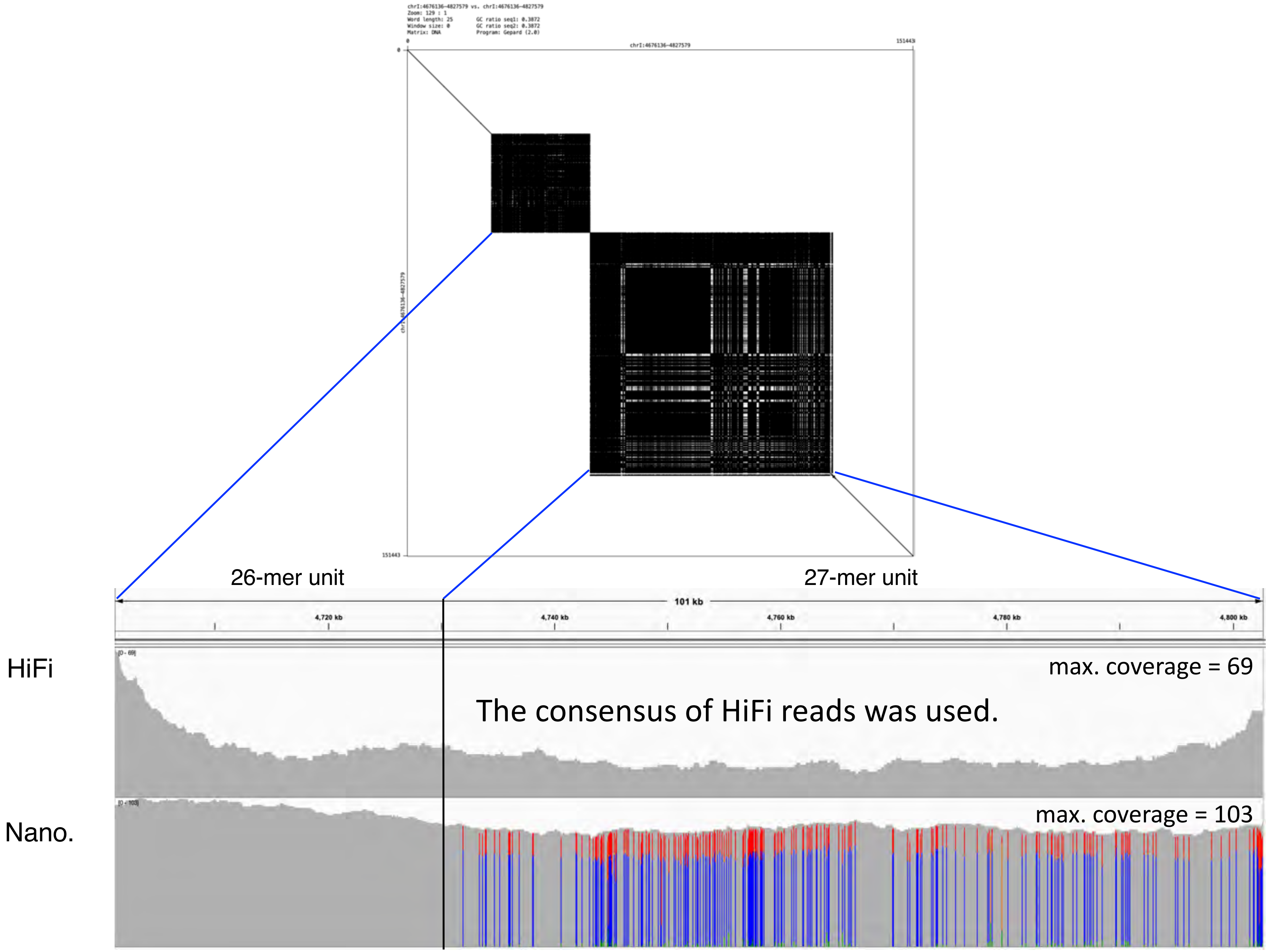

chrI:5,422,459-5,449,196

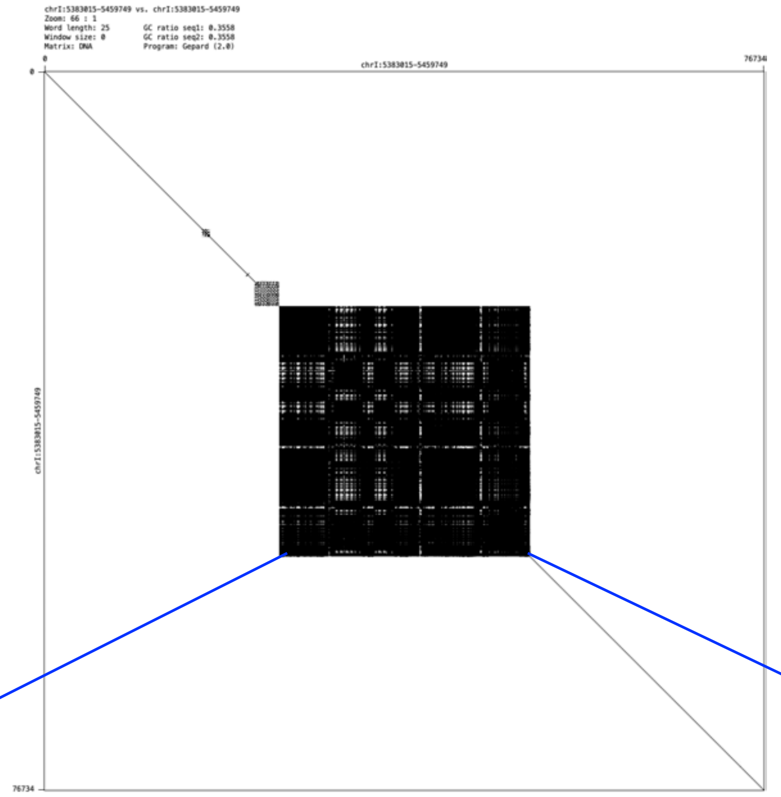


The consensus of Nanopore reads was used.

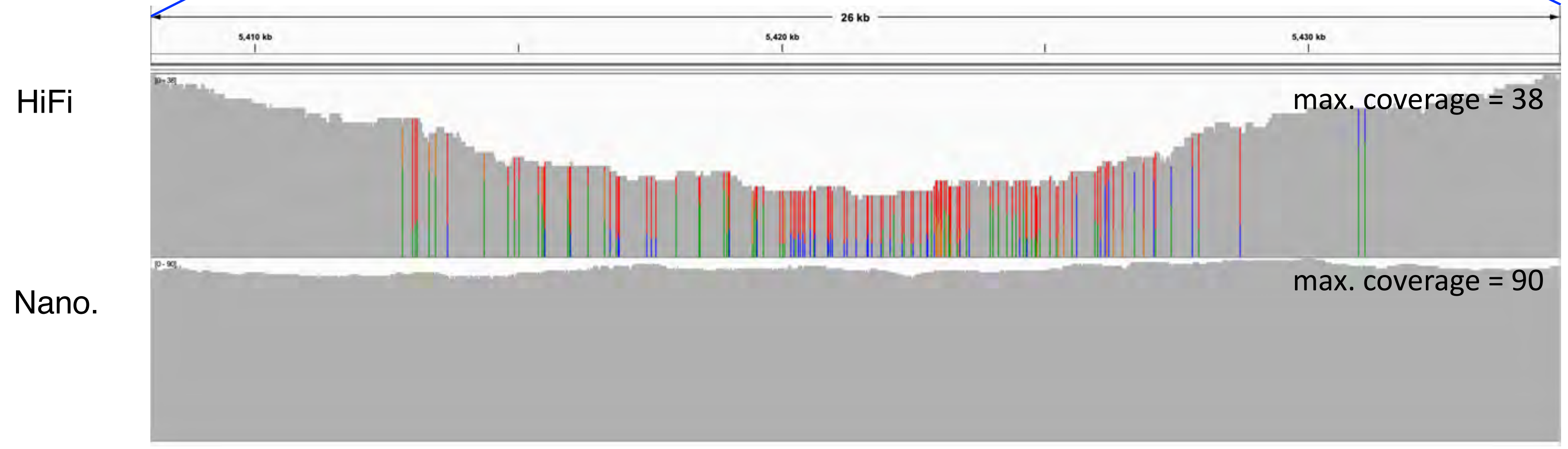

chrI:7,067,899-7,077,902

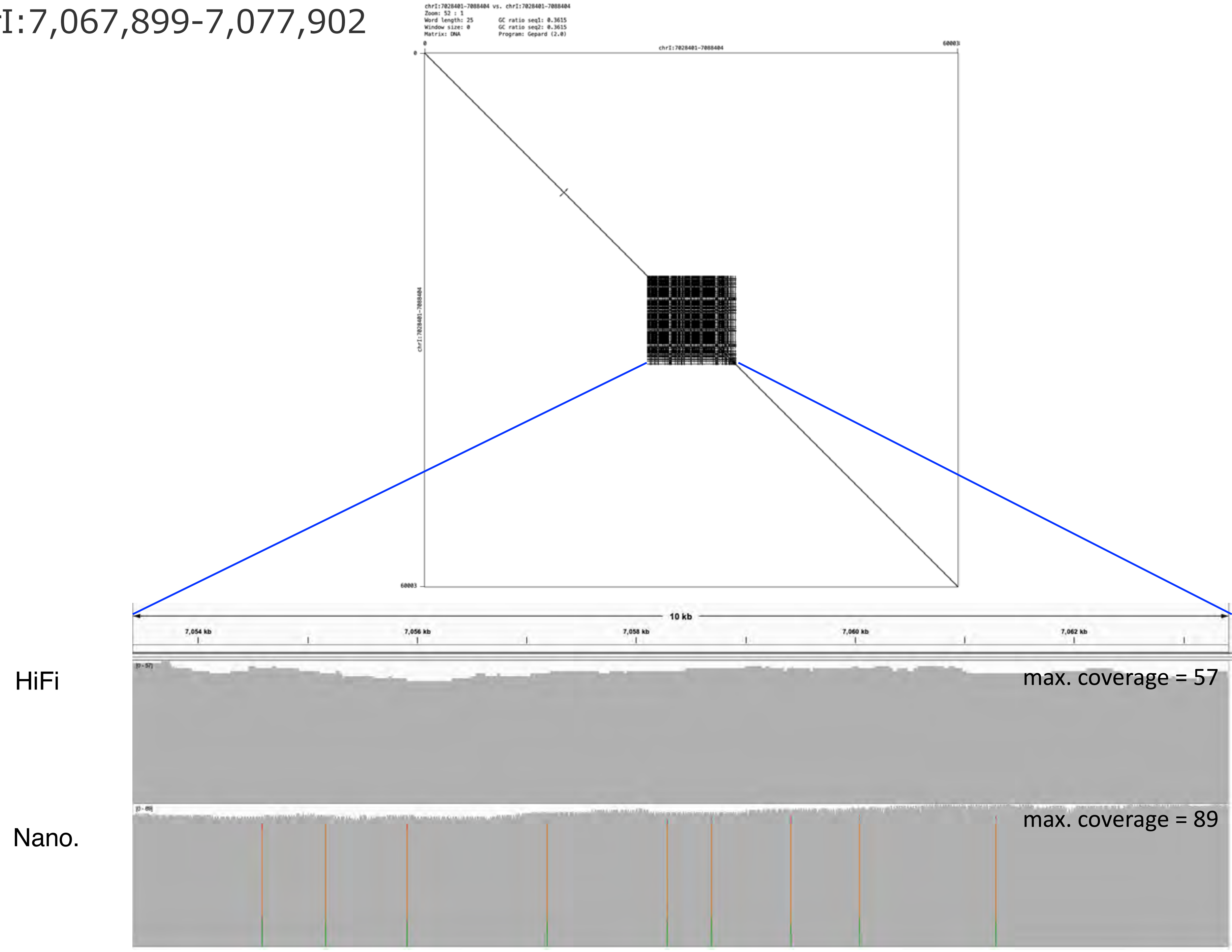

chrI:10,138,572-10,152,432

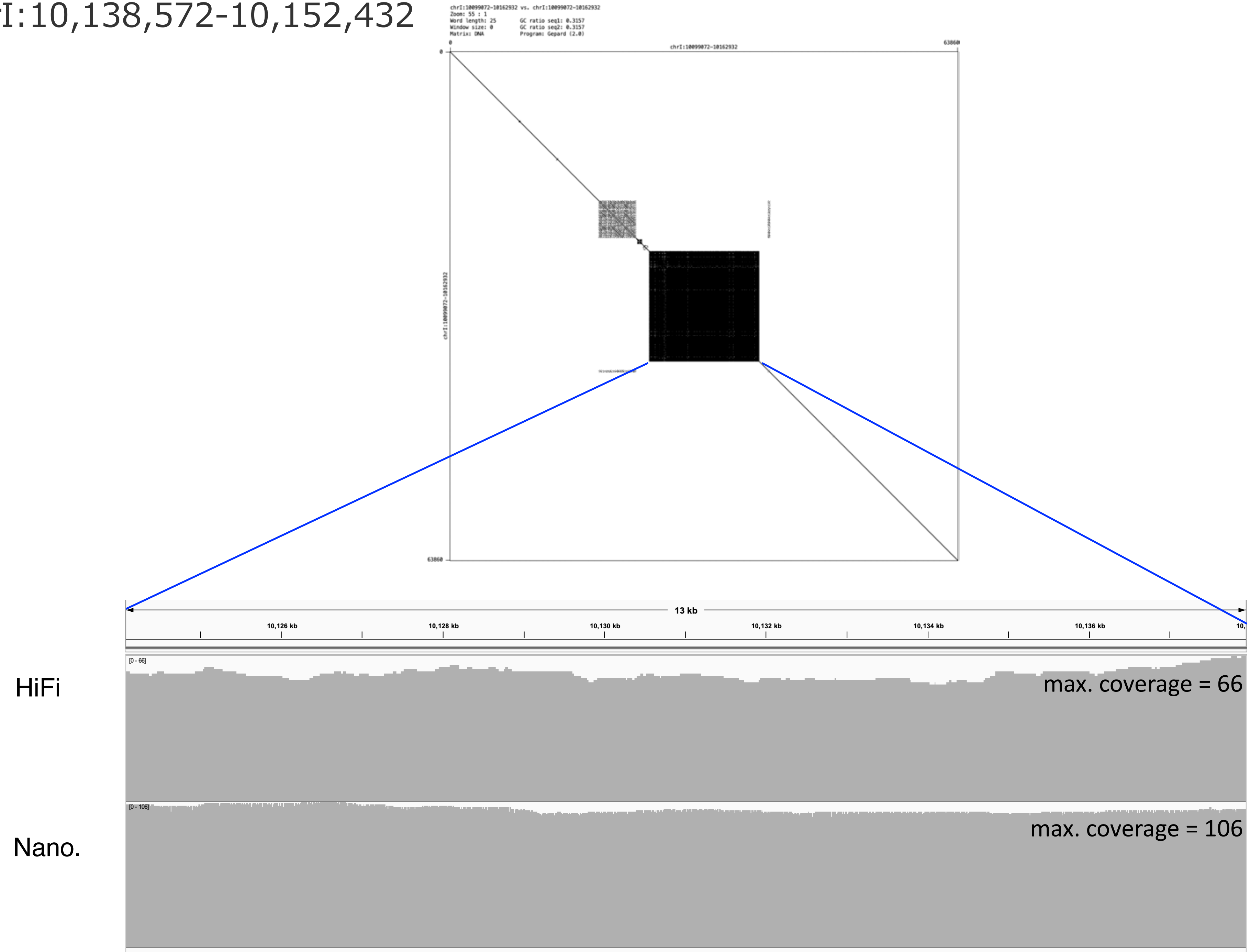

### chrI:10,460,225-10,477,707

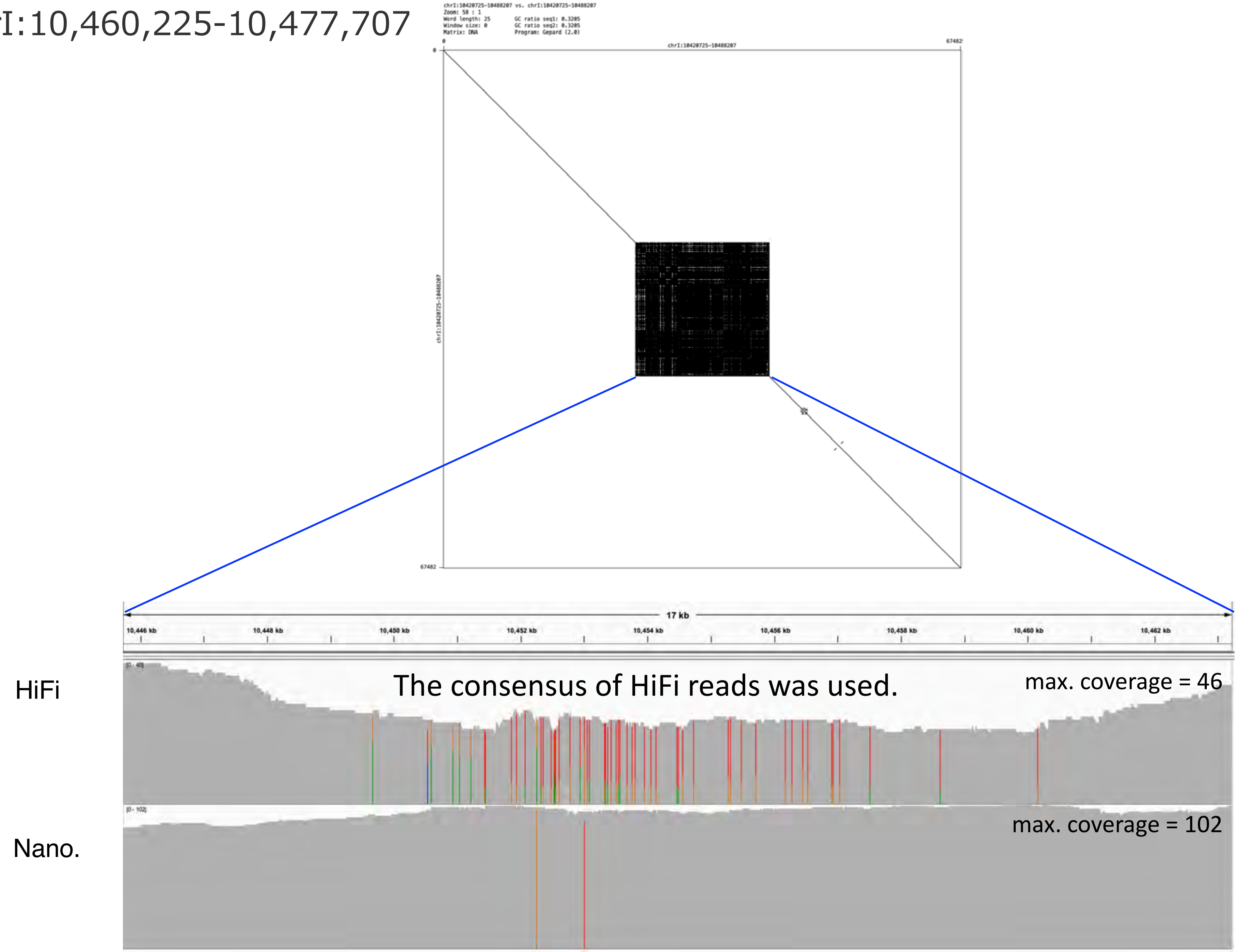

### chrI:10,549,319-10,685,899

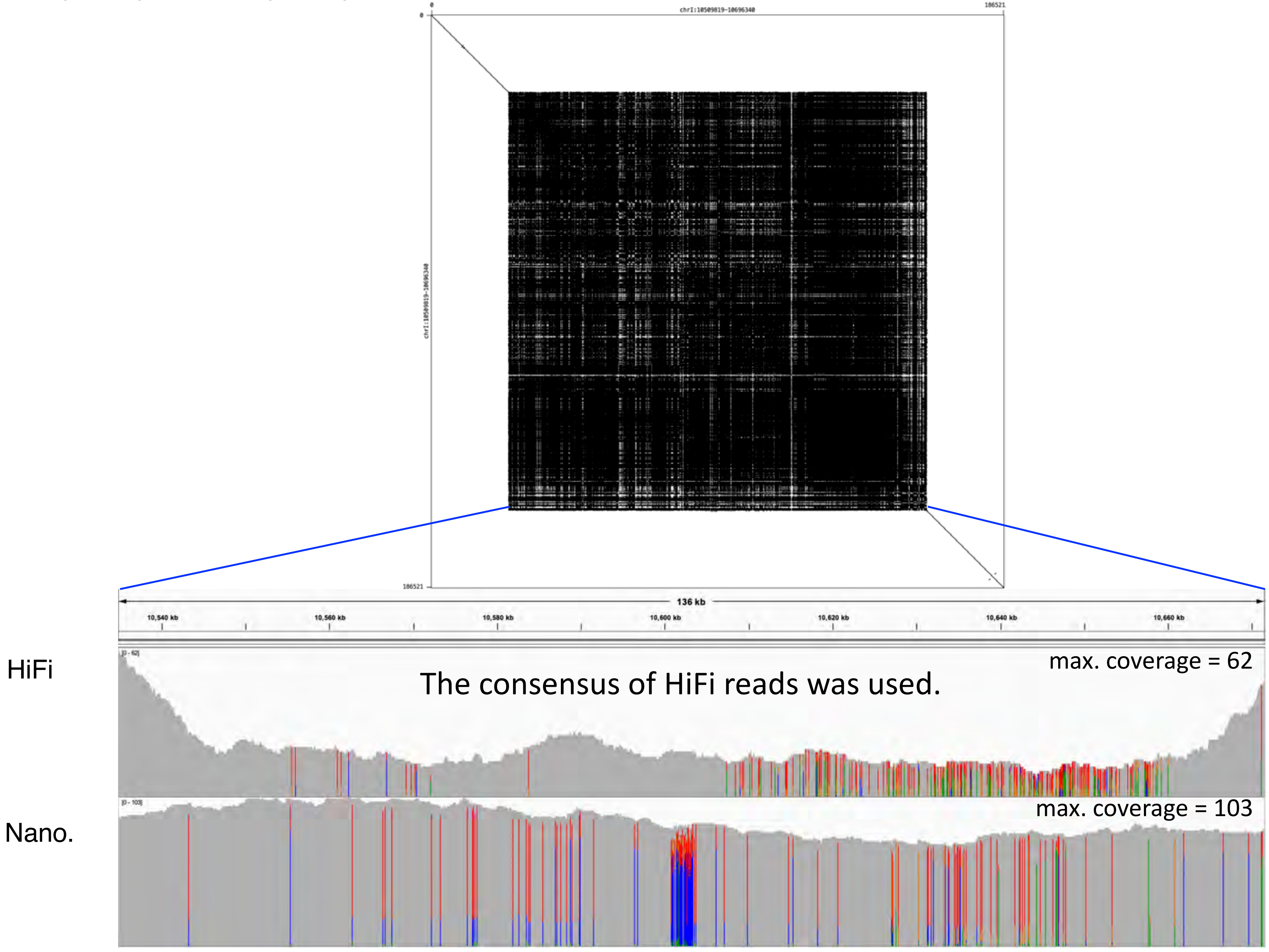

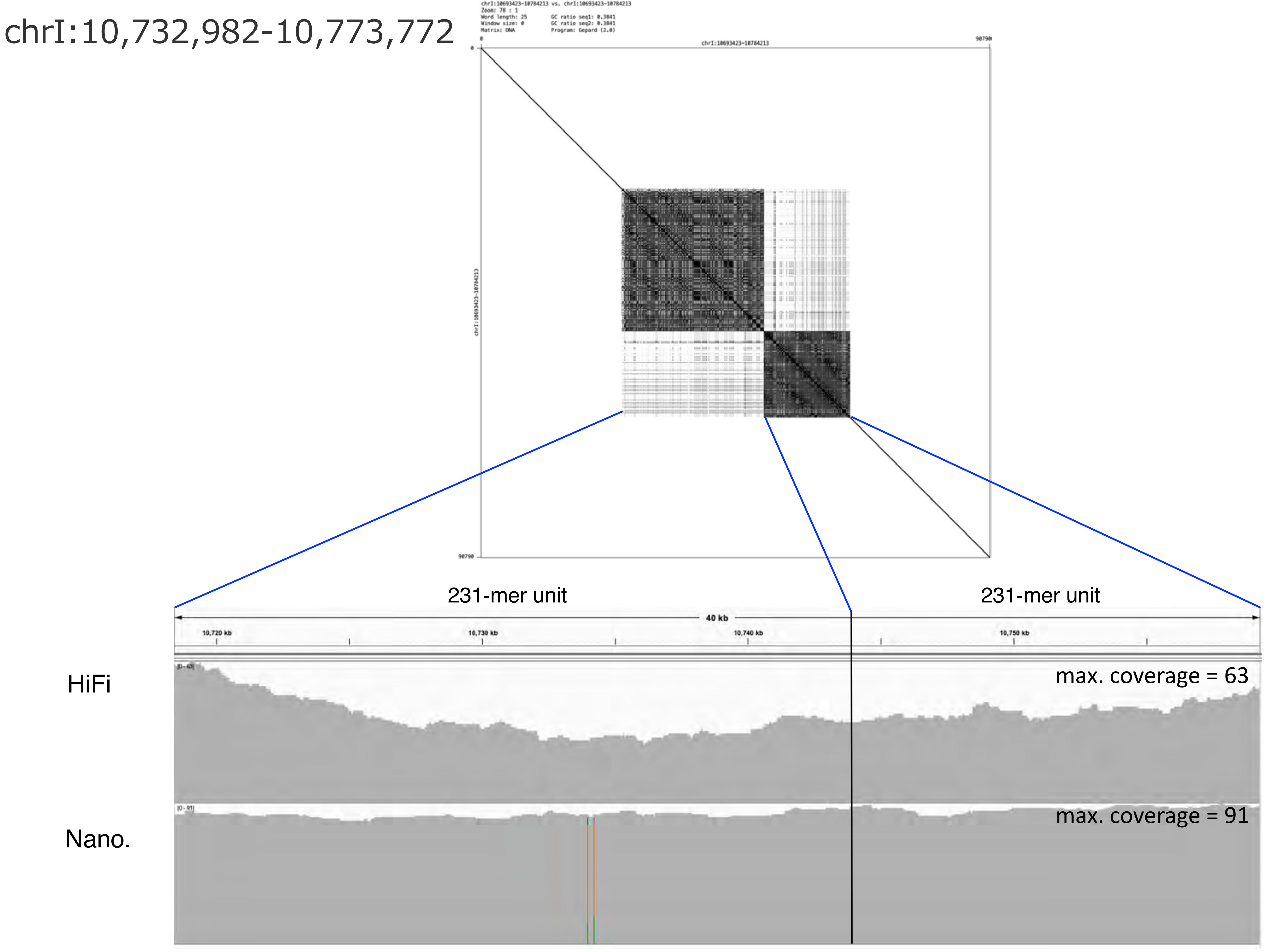
chrI:10,732,982-10,773,772
chrI:10693423-10784213 vs. chrI:10693423-10784213
Zoom: 78 : 1
Word length: 25
Window size: 0
Matrix: DNA
GC ratio seq1: 0.3841
GC ratio seq2: 0.3841
Program: Gepard (2.0)
chrI:10693423-10784213
0
90790
231-mer unit
231-mer unit
40 kb
10,720 kb
10,730 kb
10,740 kb
10,750 kb
HiFi
max. coverage = 63
Nano.
max. coverage = 91

chrI:11,449,789-11,578,375

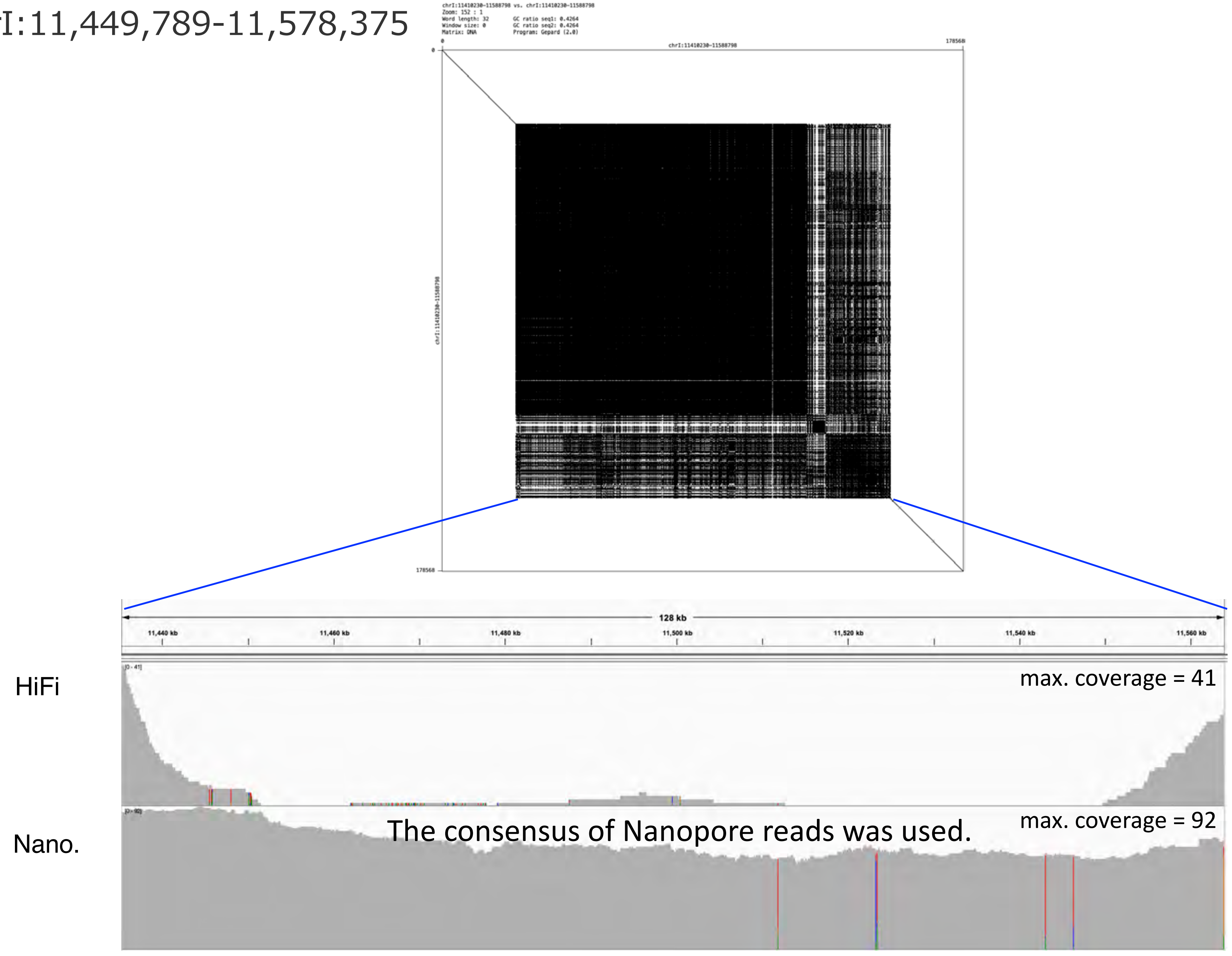

### chrI:12,011,119-12,016,749

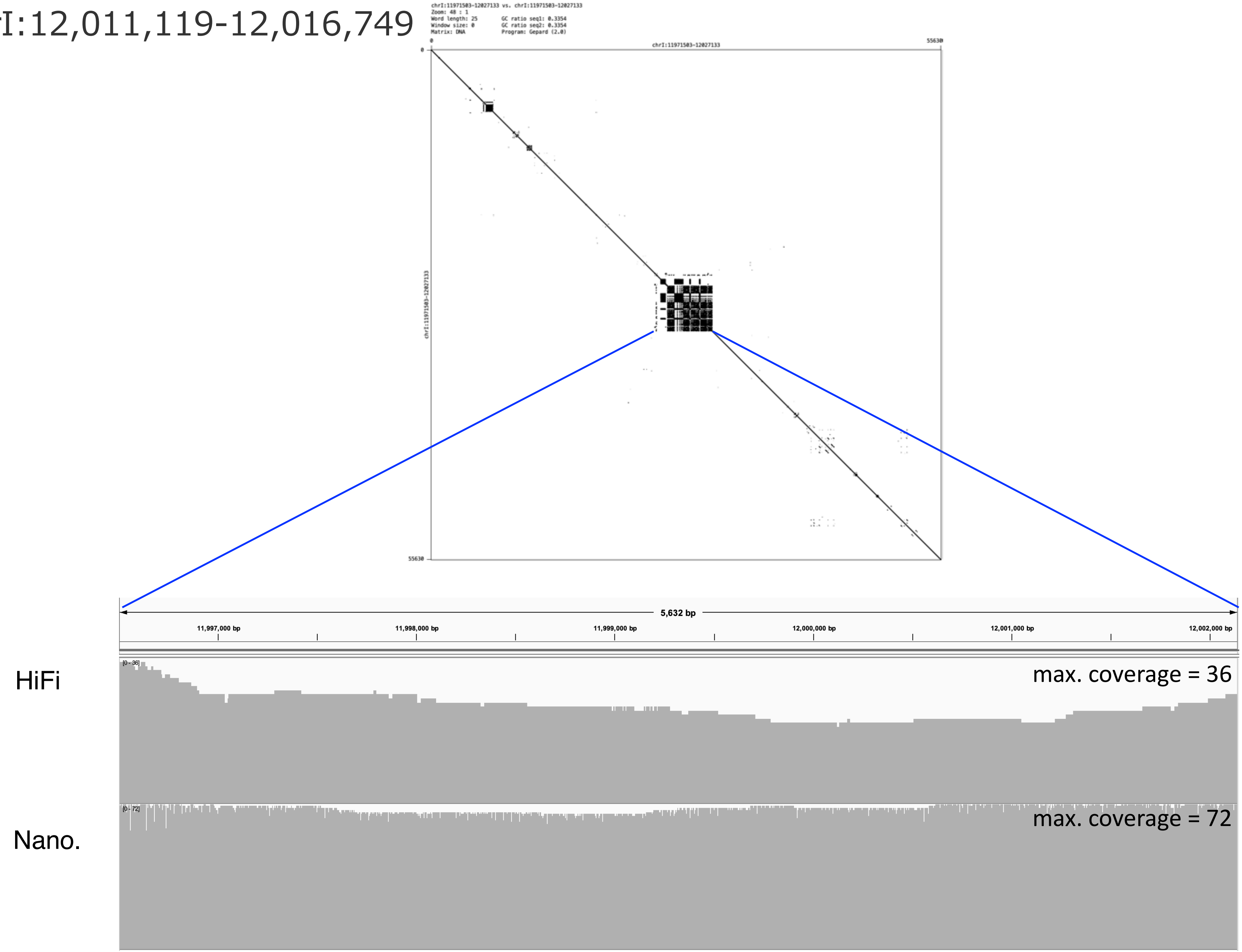

### chrI:13,790,040-13,796,824

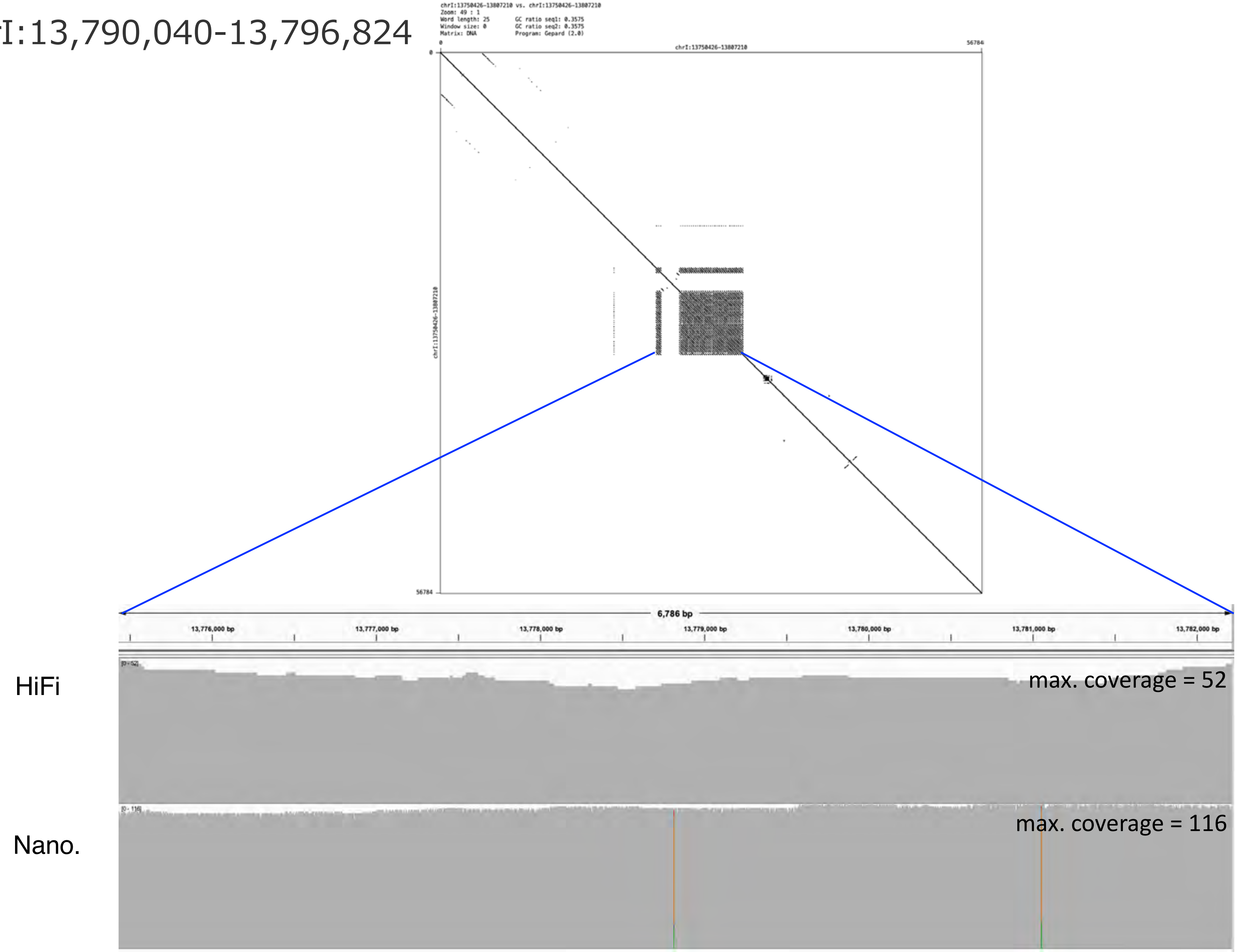

### chrI:13,875,677-13,880,843

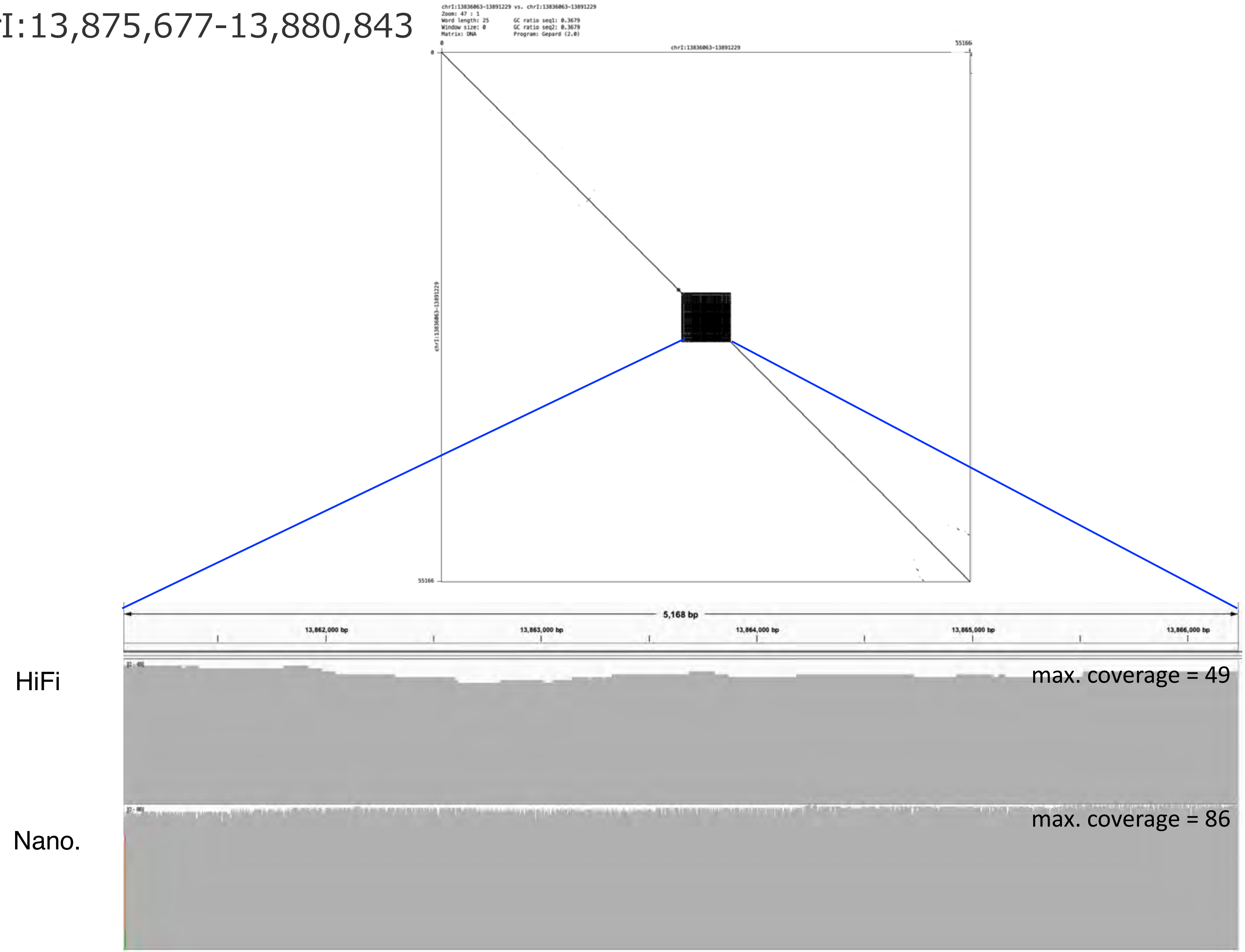

chrI:14,259,349-14,271,227

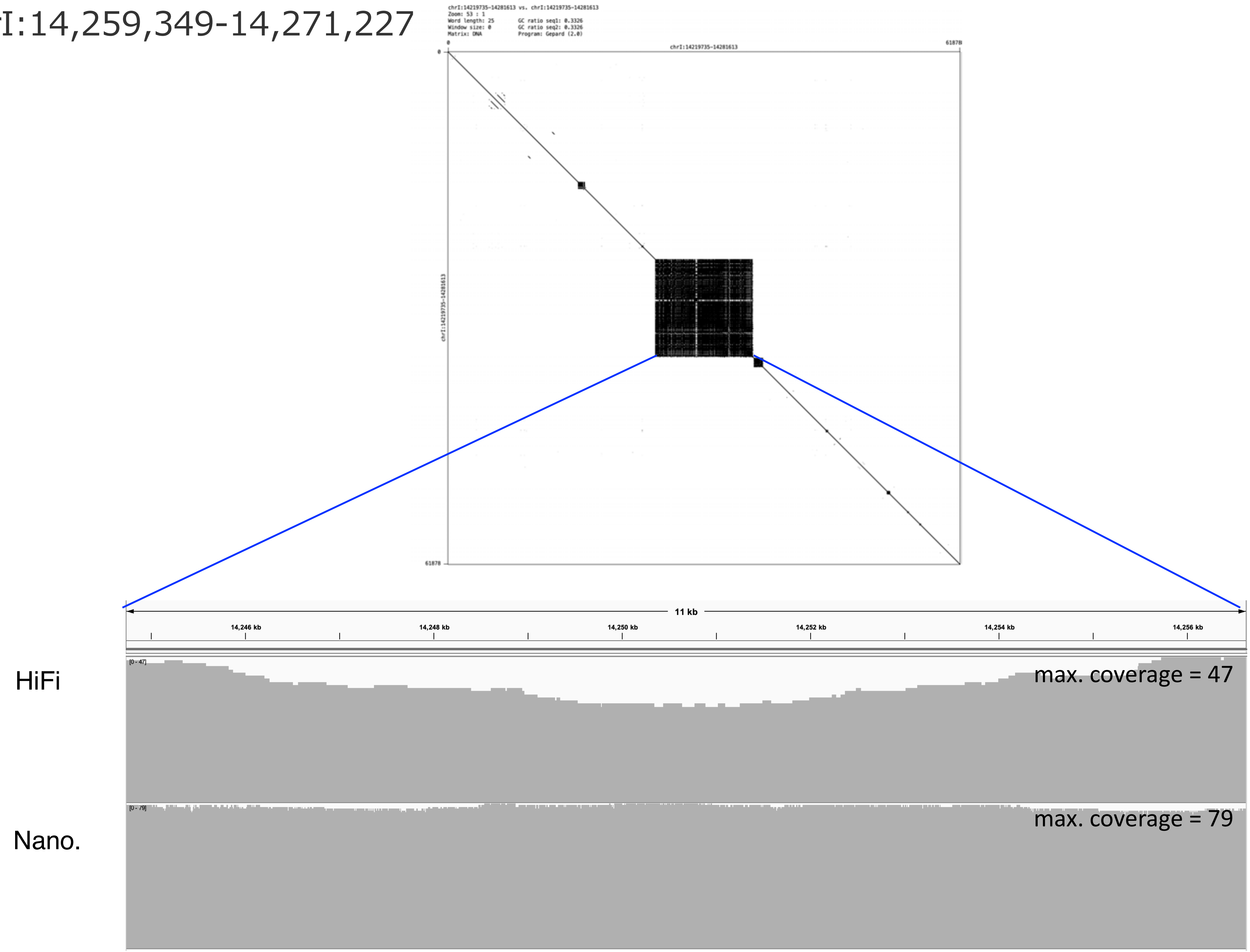

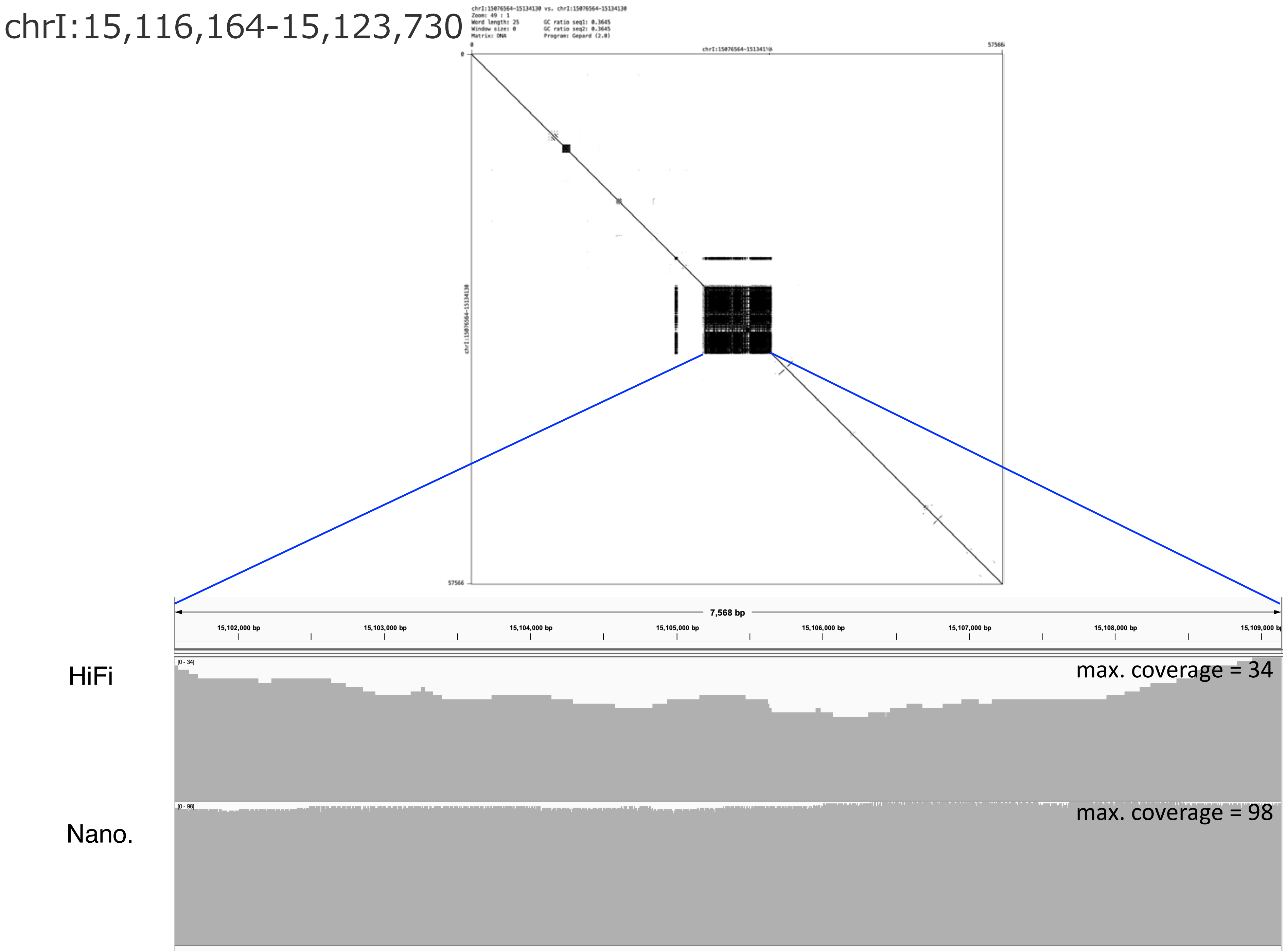
chrI:15,116,164-15,123,730
chrI:15076564-15134130 vs. chrI:15076564-15134130
Zoom: 49 : 1
Word length: 25
Window size: 0
Matrix: DNA
GC ratio seq1: 0.3645
GC ratio seq2: 0.3645
Program: Gepard (2.0)
0
57566
chrI:15076564-15134130
7,568 bp
15,102,000 bp
15,103,000 bp
15,104,000 bp
15,105,000 bp
15,106,000 bp
15,107,000 bp
15,108,000 bp
HiFi
[0 - 34]
max. coverage = 34
Nano.
[0 - 98]
max. coverage = 98

chrII:697,778-705,675

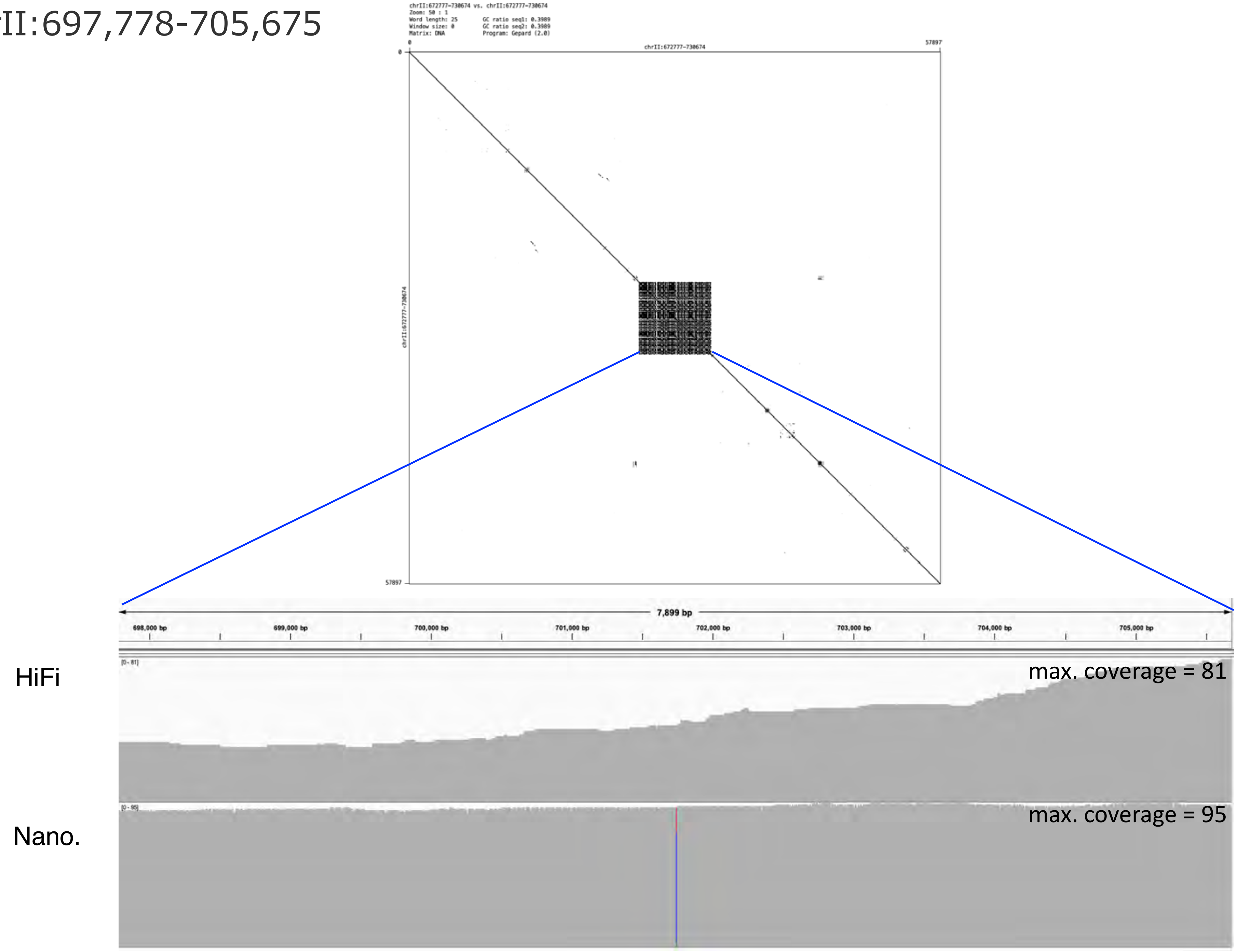

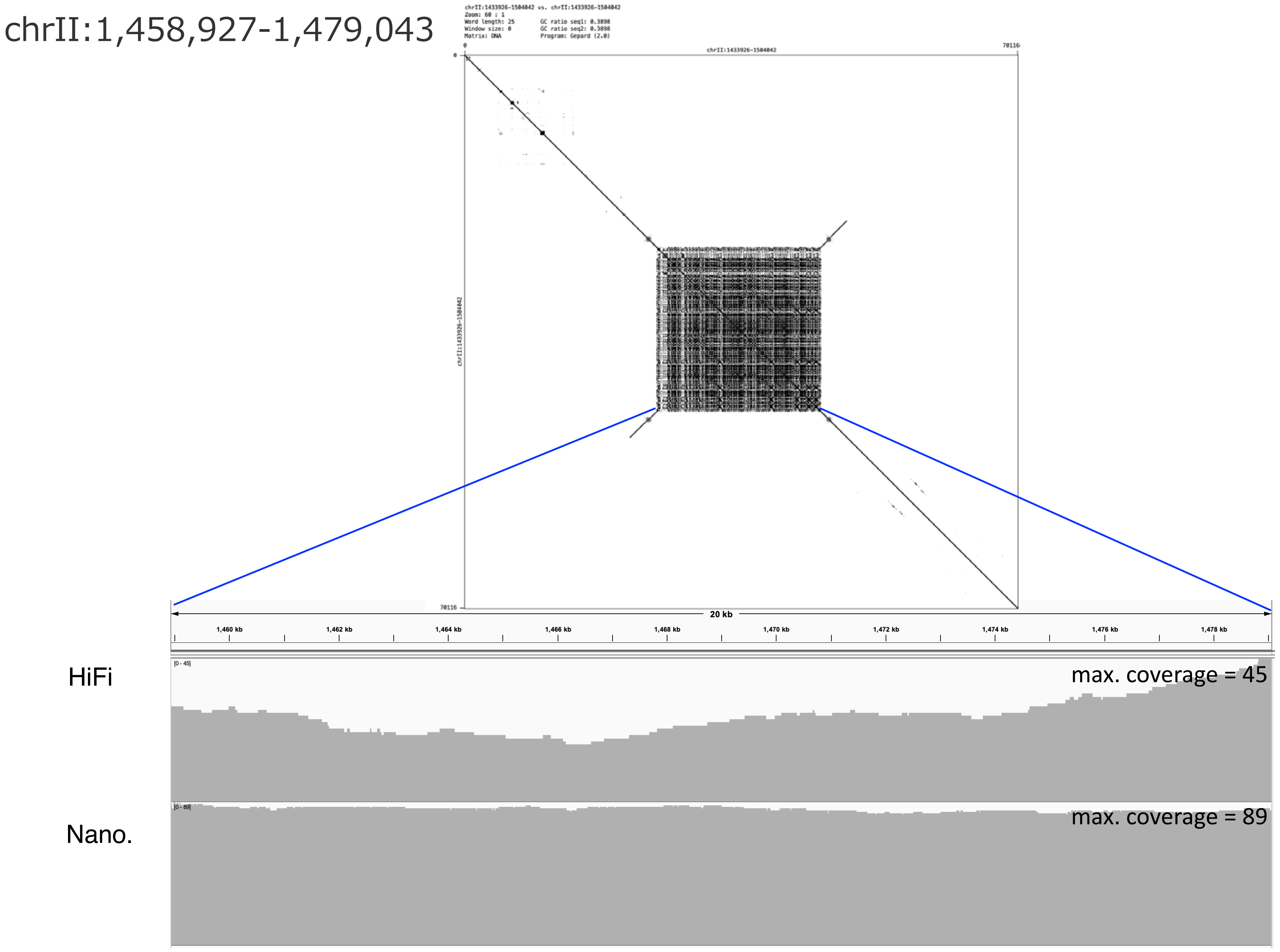
chrII:1,458,927-1,479,043
chrII:1433926-1504042 vs. chrII:1433926-1504042
Zoom: 60 : 1
Word length: 25
Window size: 0
Matrix: DNA
GC ratio seq1: 0.3898
GC ratio seq2: 0.3898
Program: Gepard (2.0)
chrII:1433926-1504042
0
70116
chrII:1433926-1504042
70116
20 kb
1,460 kb
1,462 kb
1,464 kb
1,466 kb
1,468 kb
1,470 kb
1,472 kb
1,474 kb
1,476 kb
1,478 kb
HiFi
[0 - 45]
max. coverage = 45
Nano.
[0 - 89]
max. coverage = 89

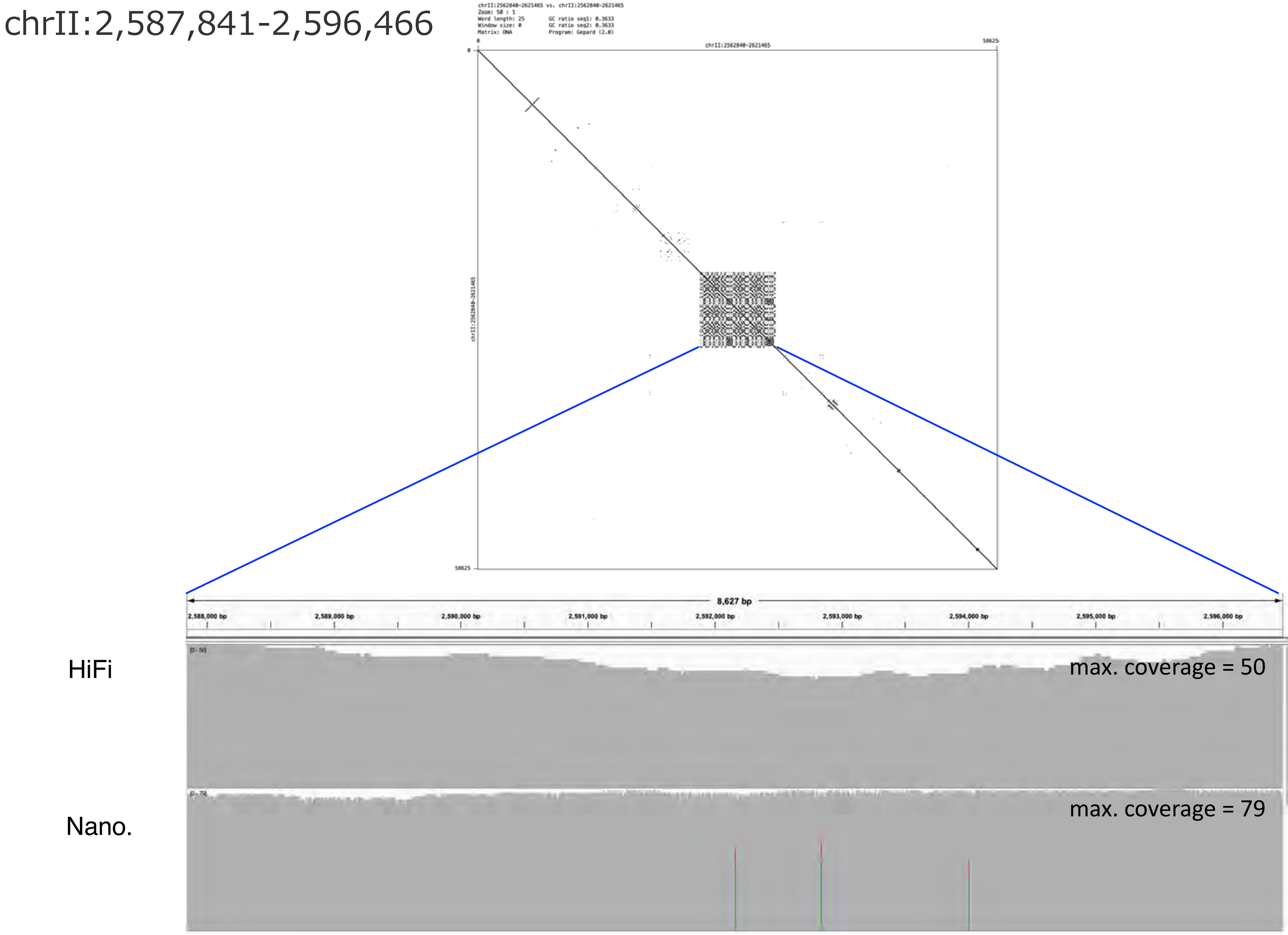

chrII:2,587,841-2,596,466
chrII:2562840-2621465 vs. chrII:2562840-2621465
Zoom: 50 : 1
Word length: 25
GC ratio seq1: 0.3633
Window size: 0
GC ratio seq2: 0.3633
Matrix: DNA
Program: Gepard (2.0)
0
chrII:2562840-2621465
58625
8,627 bp
2,588,000 bp
2,589,000 bp
2,590,000 bp
2,591,000 bp
2,592,000 bp
2,593,000 bp
2,594,000 bp
2,595,000 bp
2,596,000 bp
HiFi
max. coverage = 50
Nano.
max. coverage = 79

### chrII:2,923,257-2,929,270

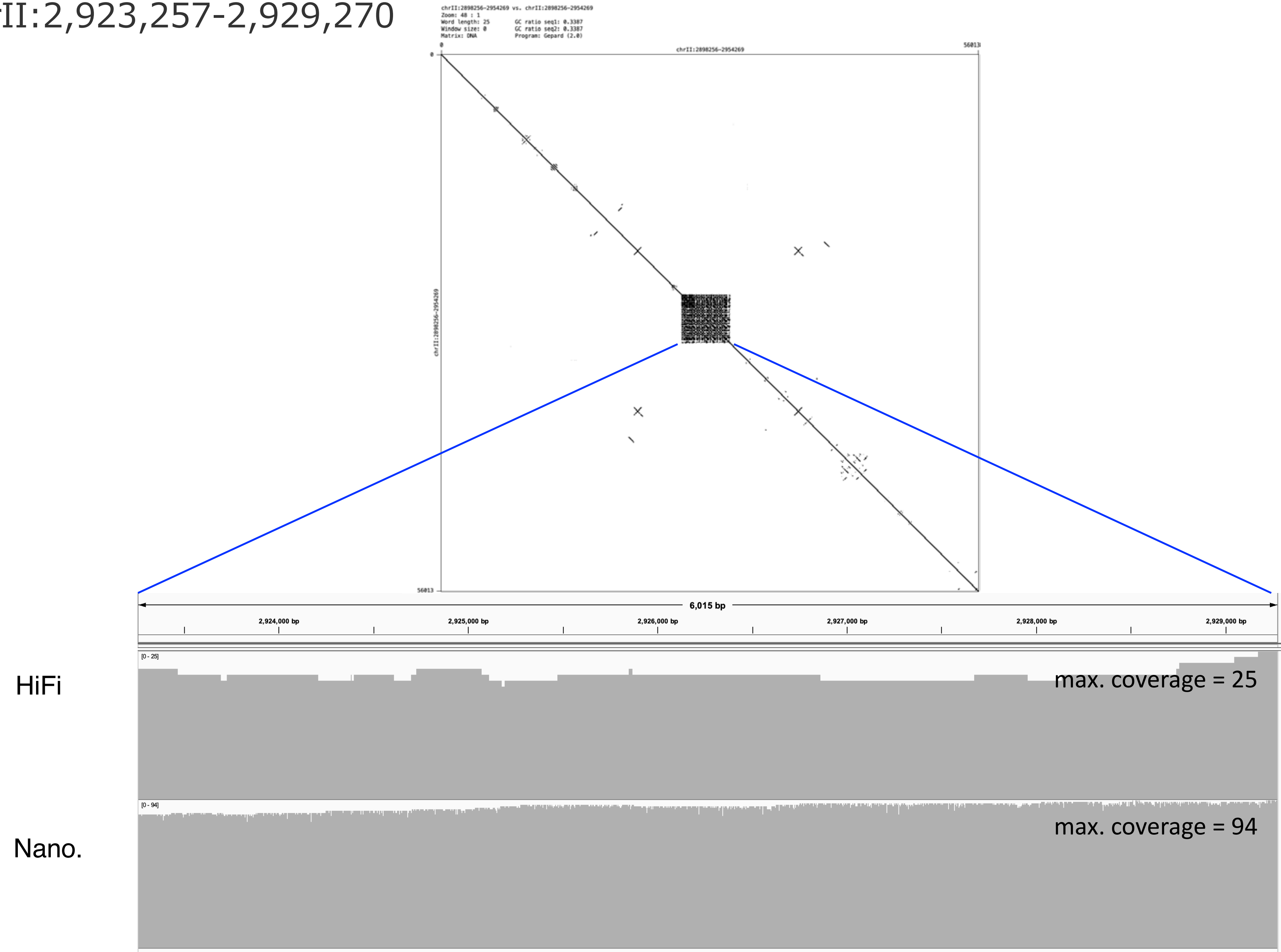

### chrII:3,189,552-3,209,877

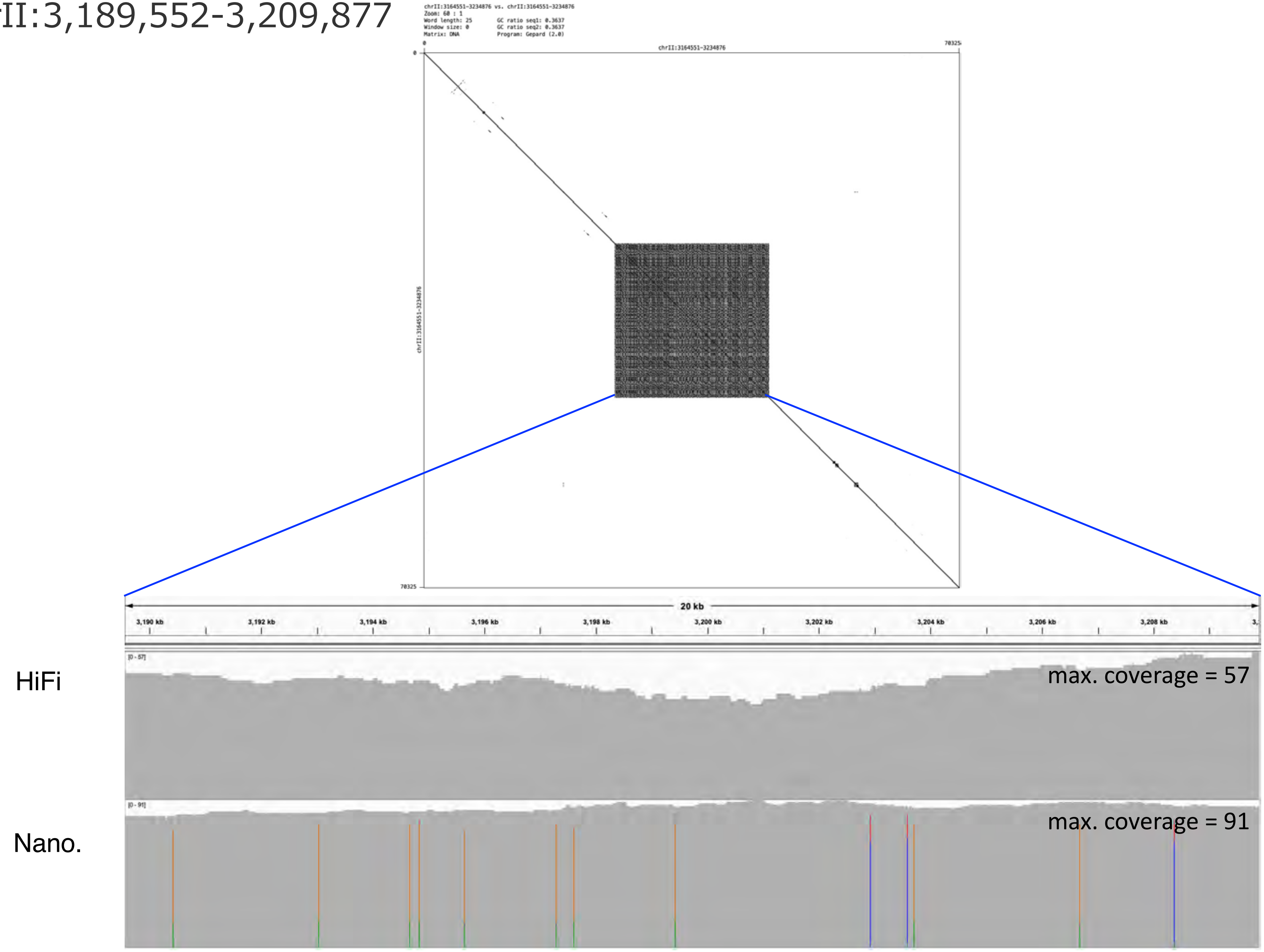

### chrII:3,506,184-3,514,999

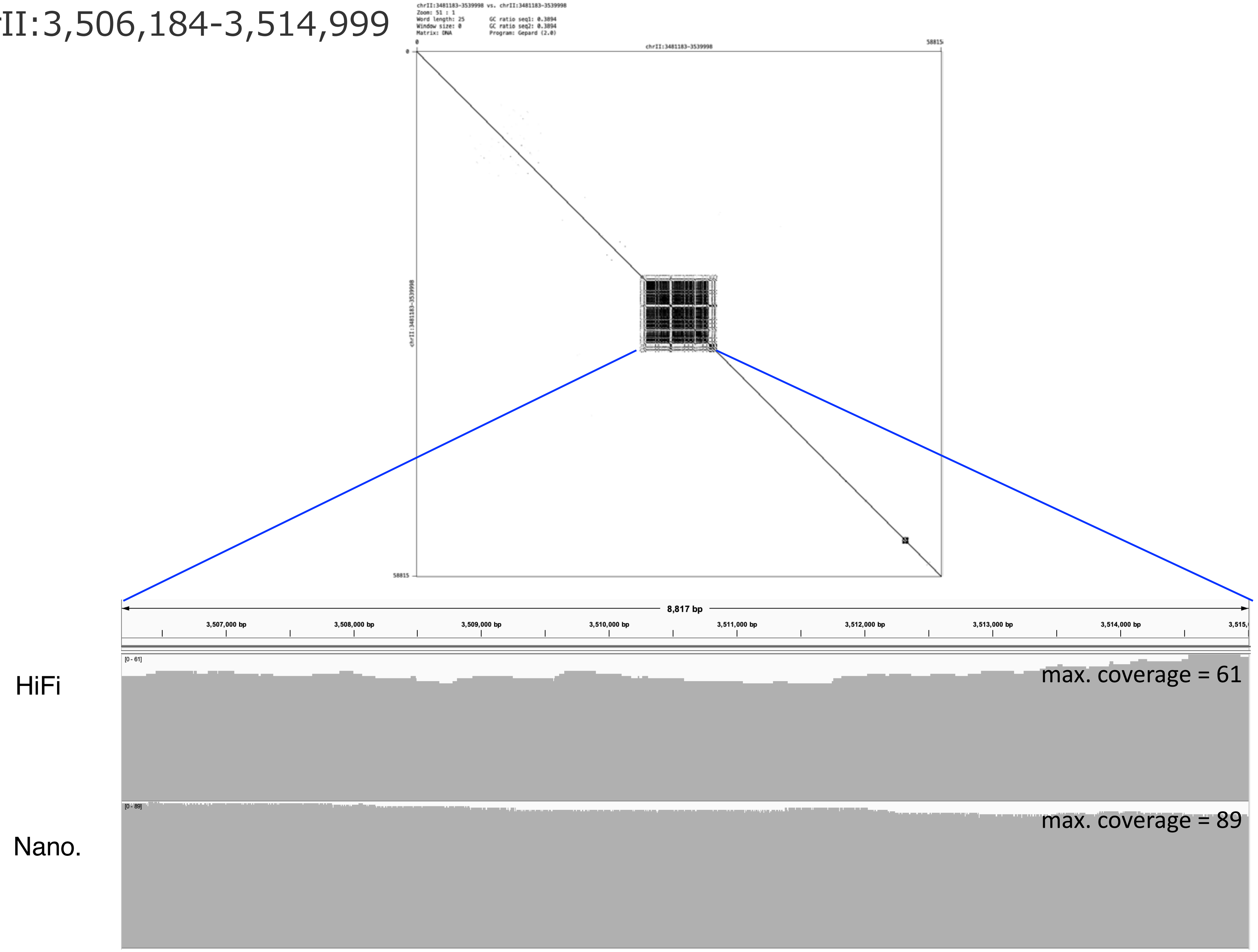

### chrII:3,842,207-3,852,973

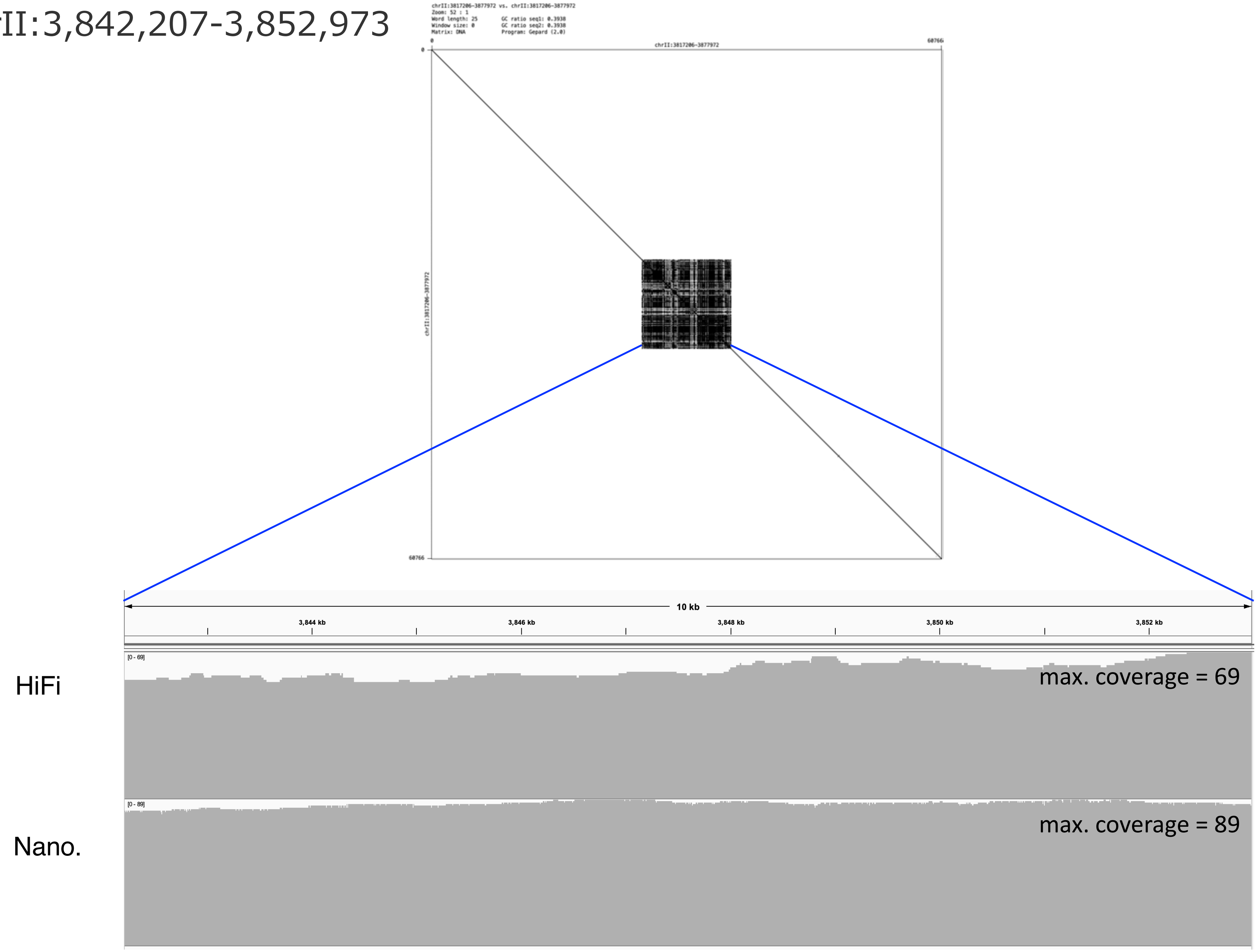

chrII:4,002,513-4,008,061

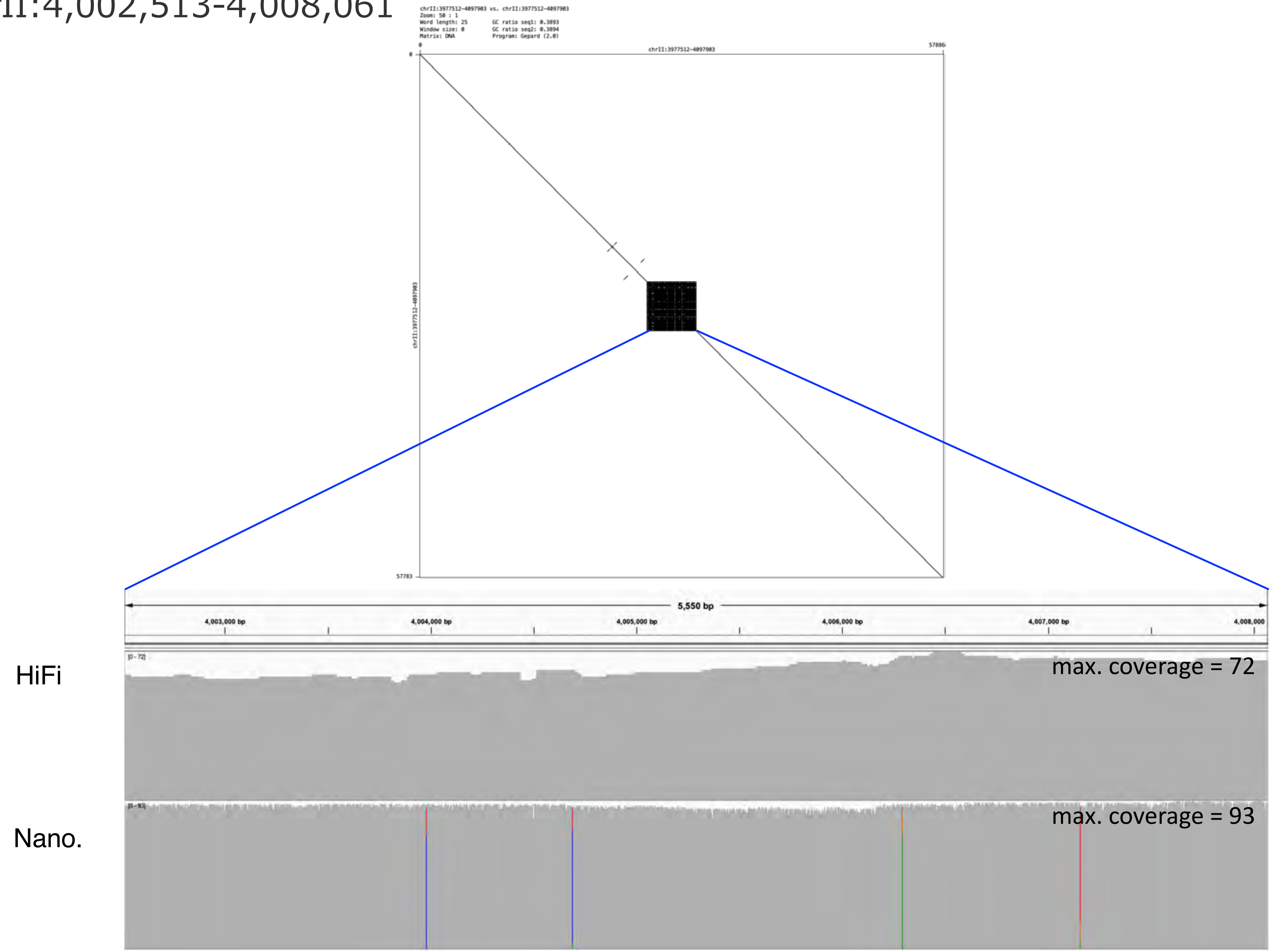

chrII:4,056,156-4,072,904

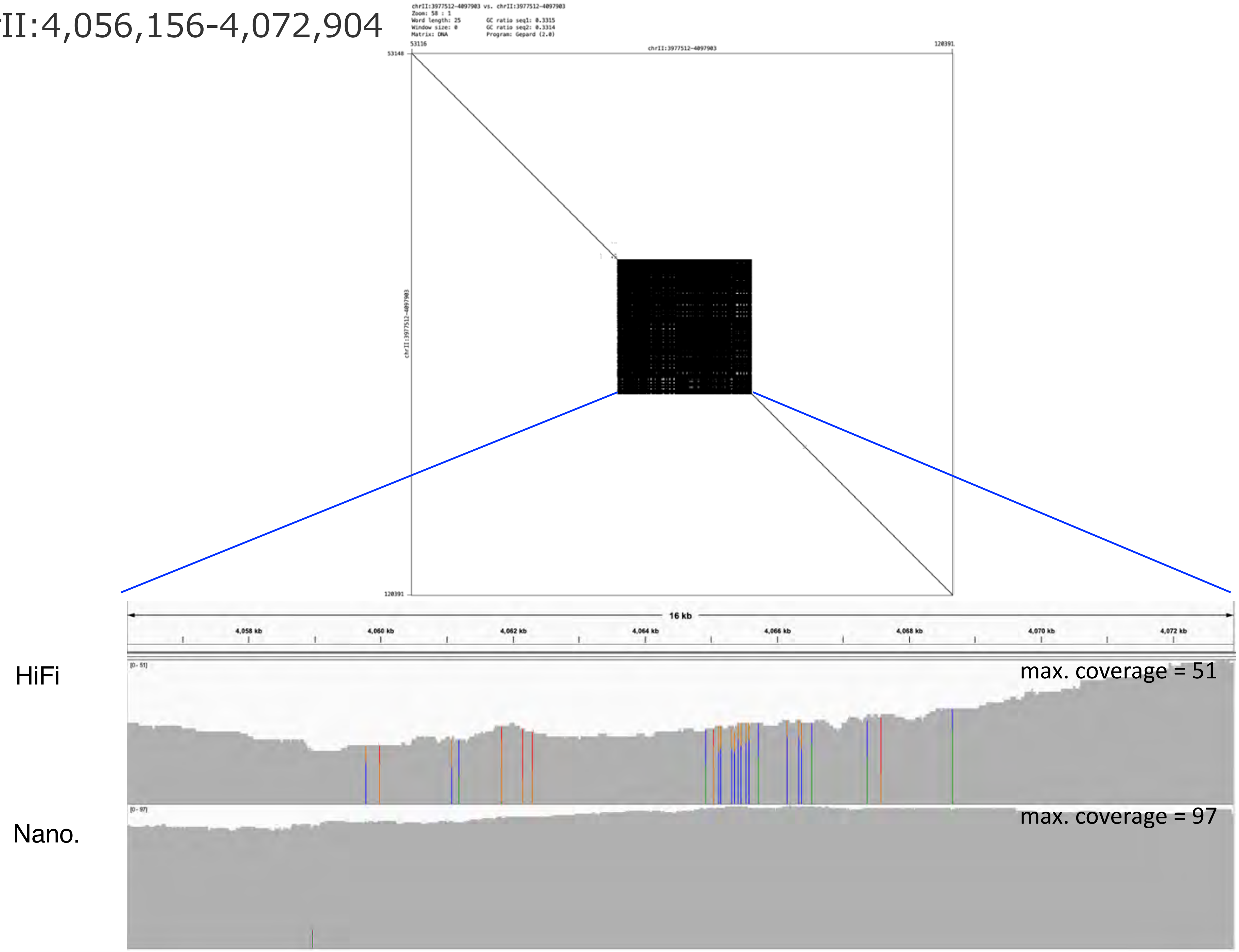

### chrII:4,386,656-4,392,377

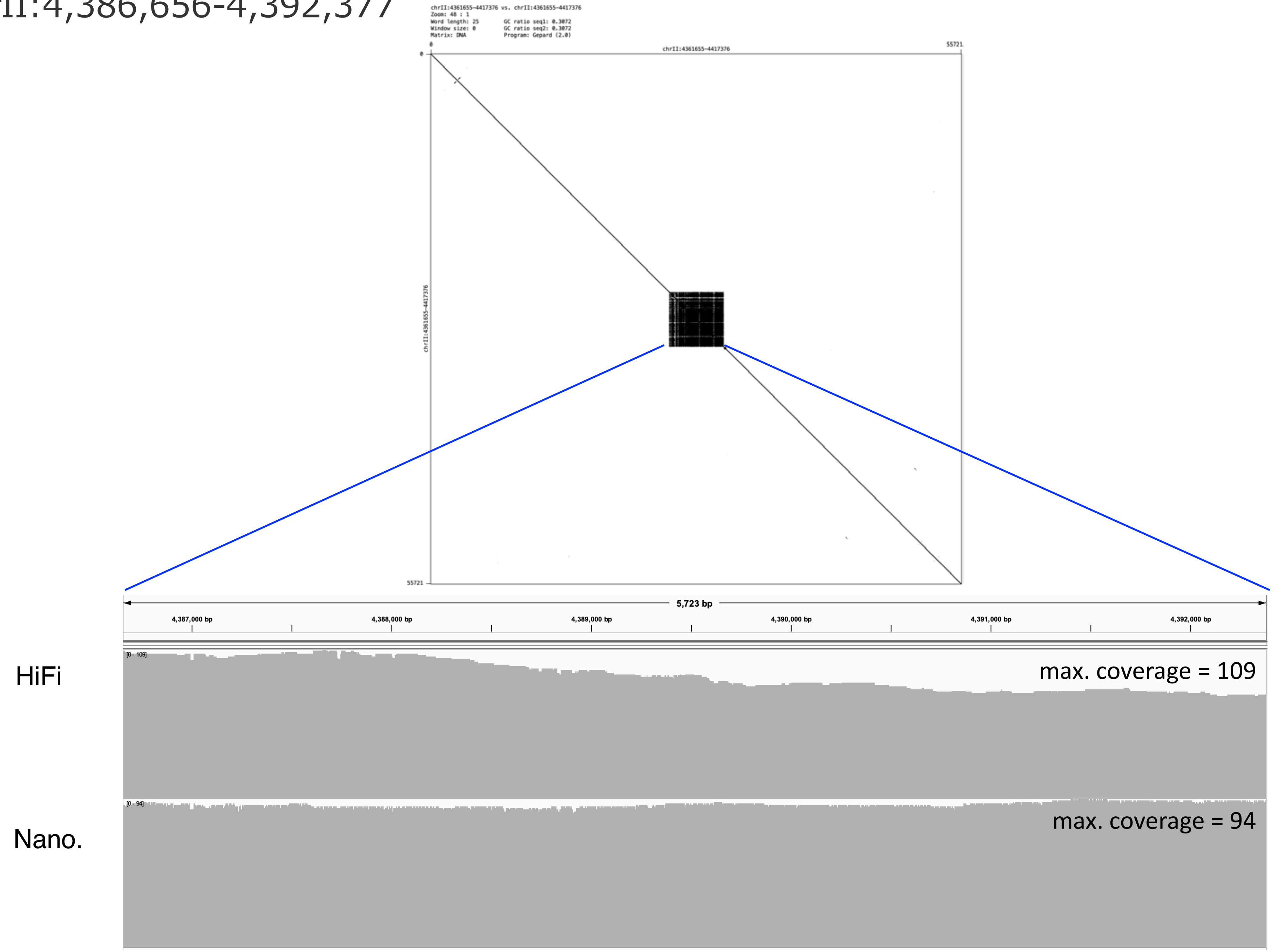

### chrII:4,724,786-4,779,090

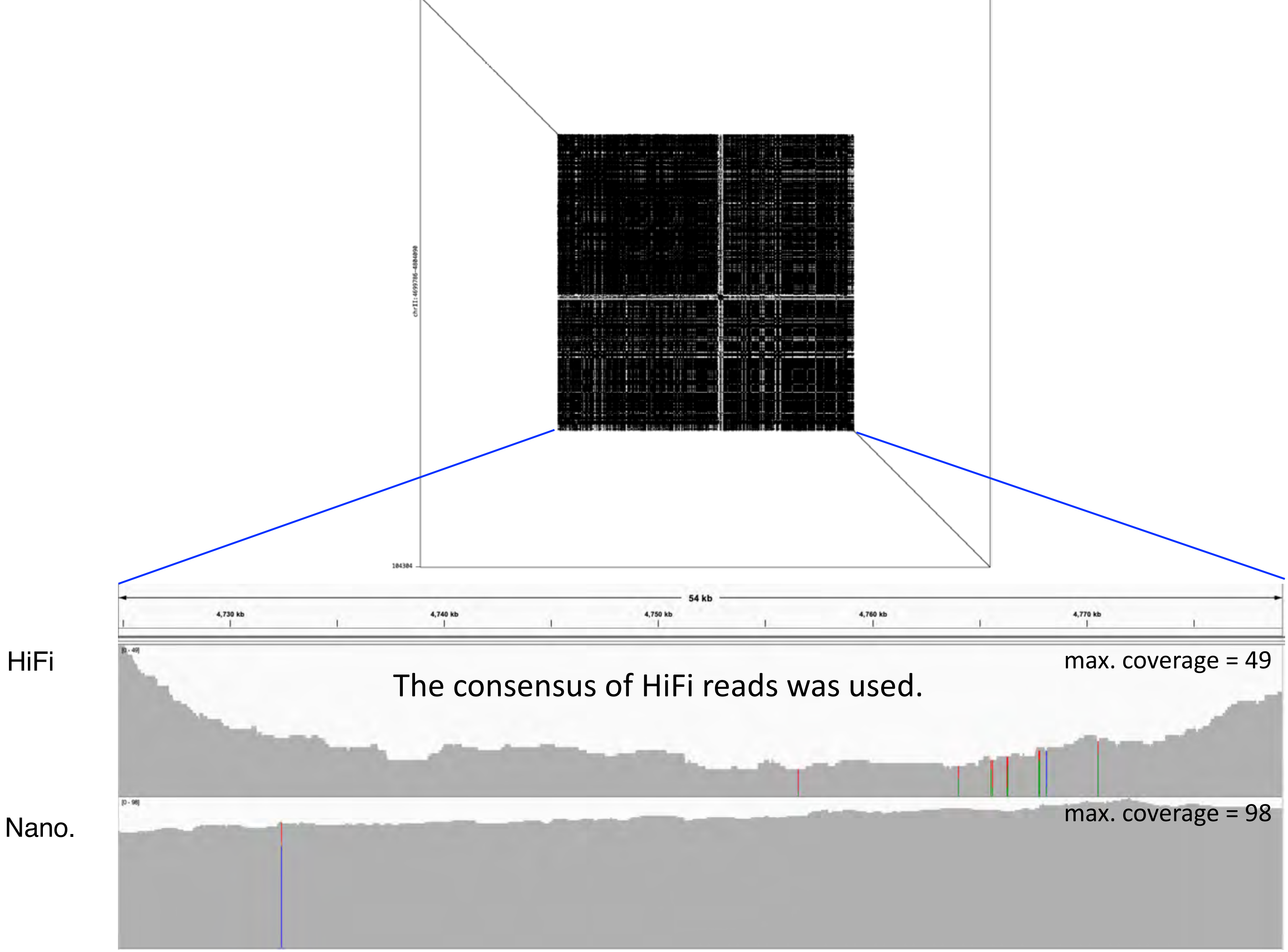

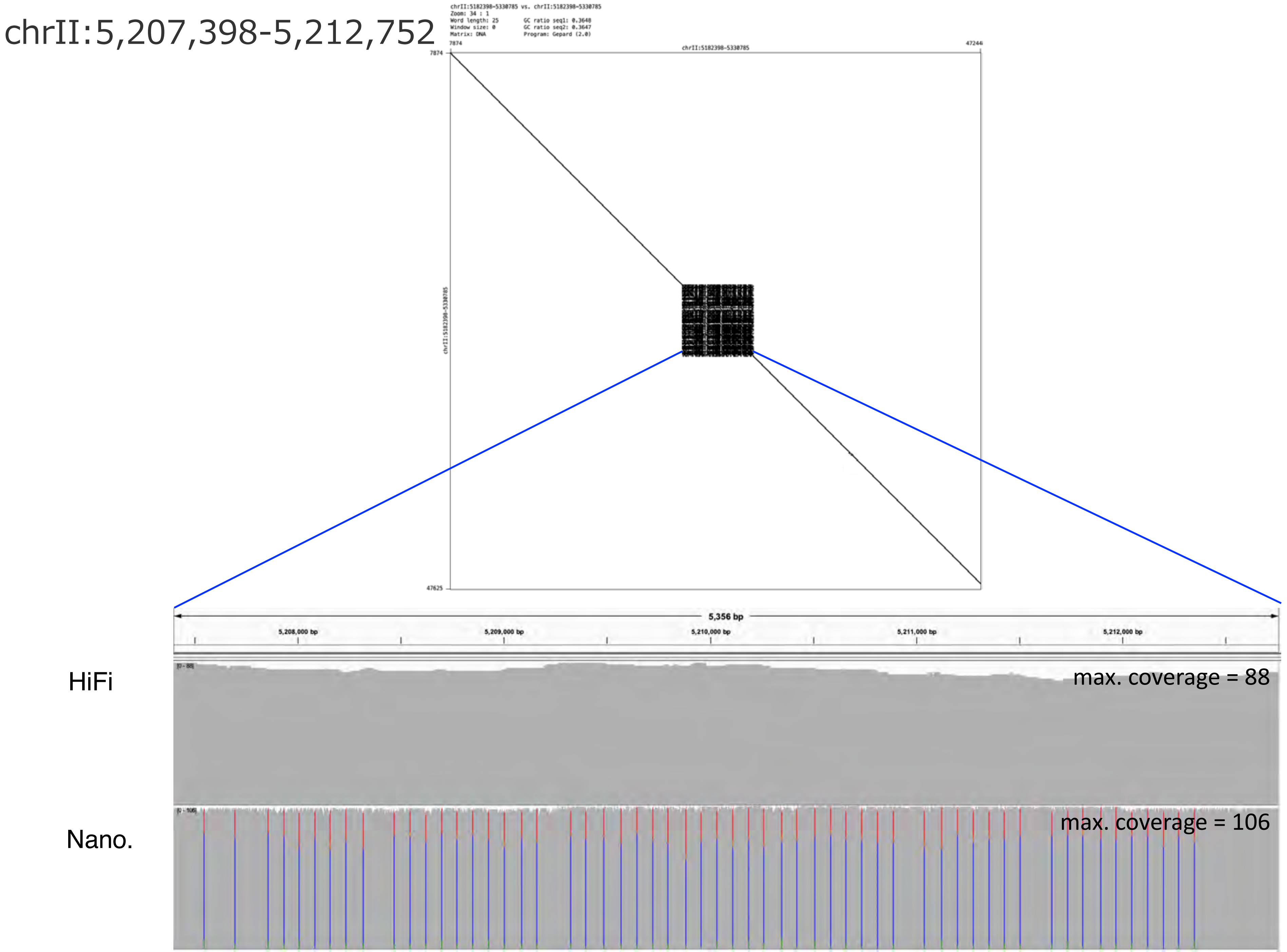
chrII:5,207,398-5,212,752
chrII:5182398-5330785 vs. chrII:5182398-5330785
Zoom: 34 : 1
Word length: 25
Window size: 0
Matrix: DNA
GC ratio seq1: 0.3648
GC ratio seq2: 0.3647
Program: Gepard (2.0)
7874
chrII:5182398-5330785
47244
7874
chrII:5182398-5330785
47625
5,356 bp
5,208,000 bp
5,209,000 bp
5,210,000 bp
5,211,000 bp
5,212,000 bp
HiFi
max. coverage = 88
Nano.
max. coverage = 106

### chrII:5,232,530-5,238,503

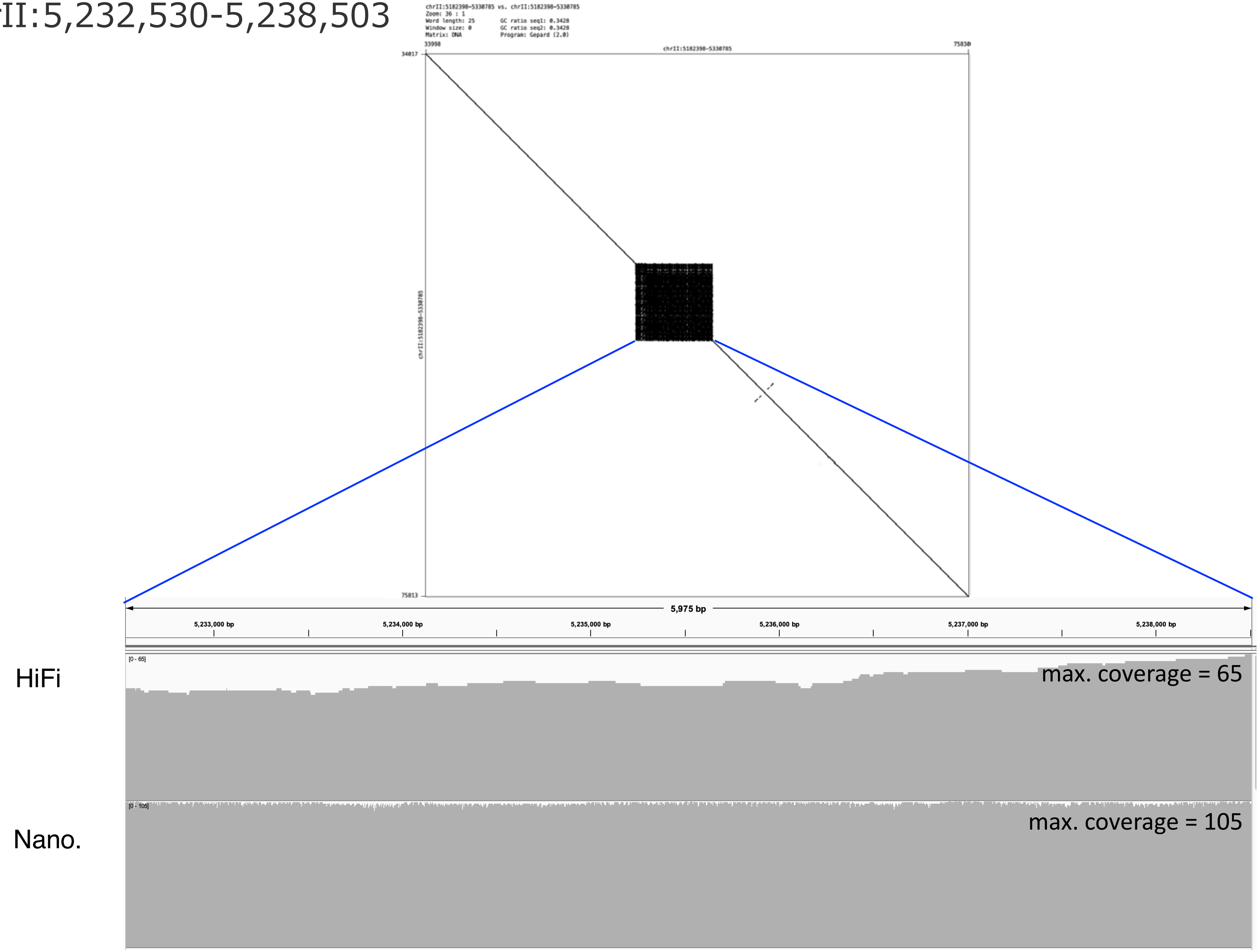

### chrII:5,285,584-5,305,785

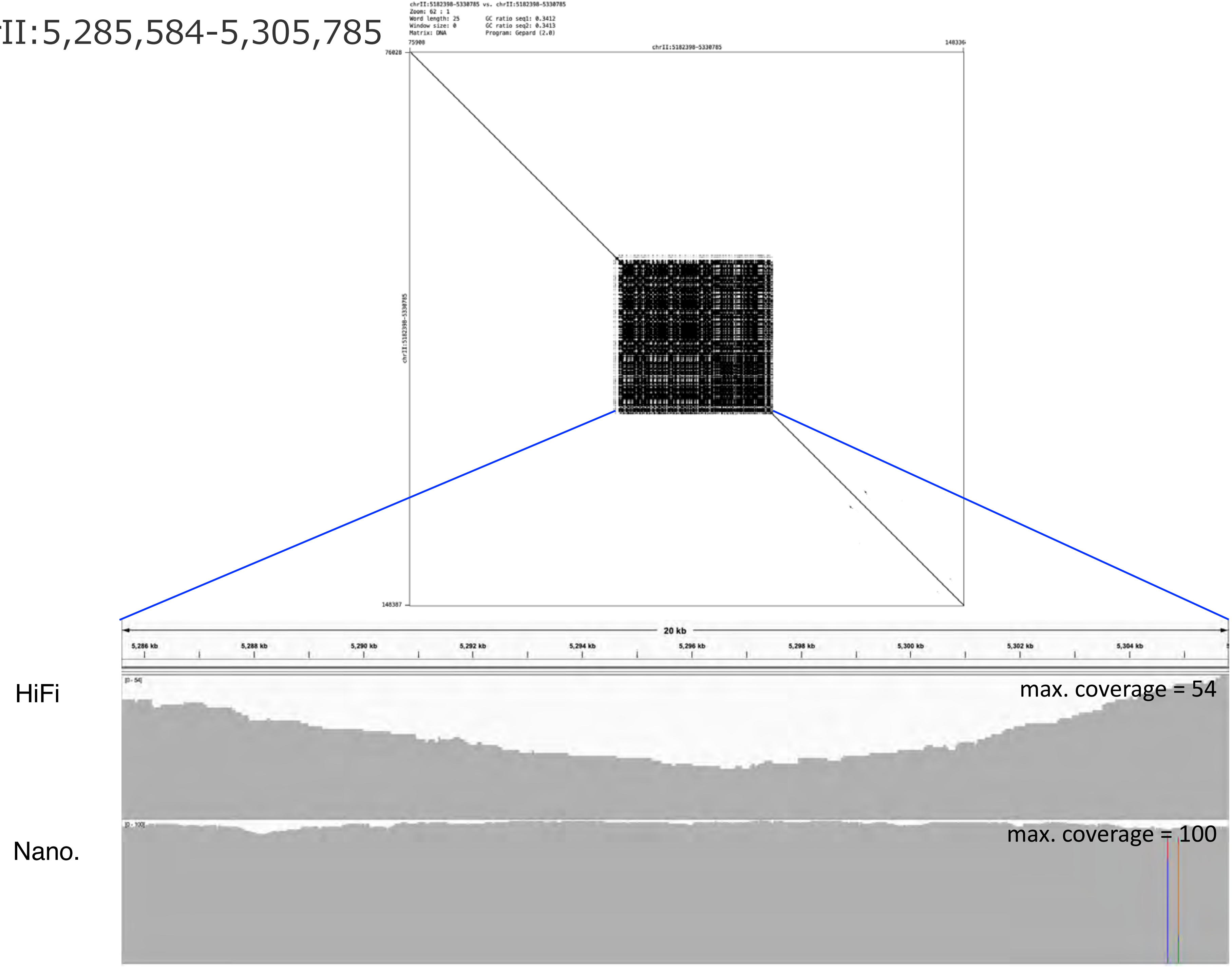

### chrII:6,667,858-6,686,160

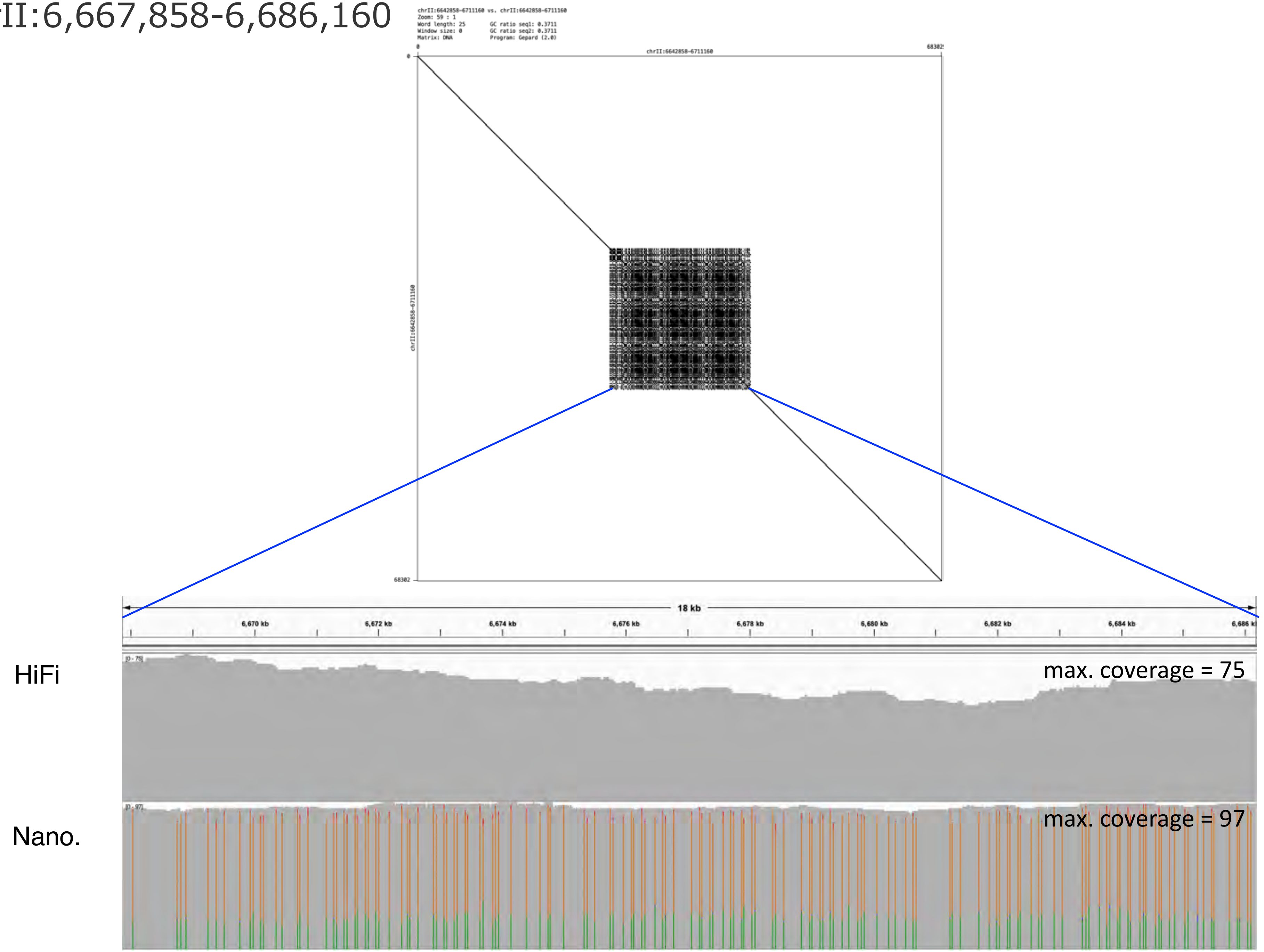

### chrII:8,471,747-8,615,728

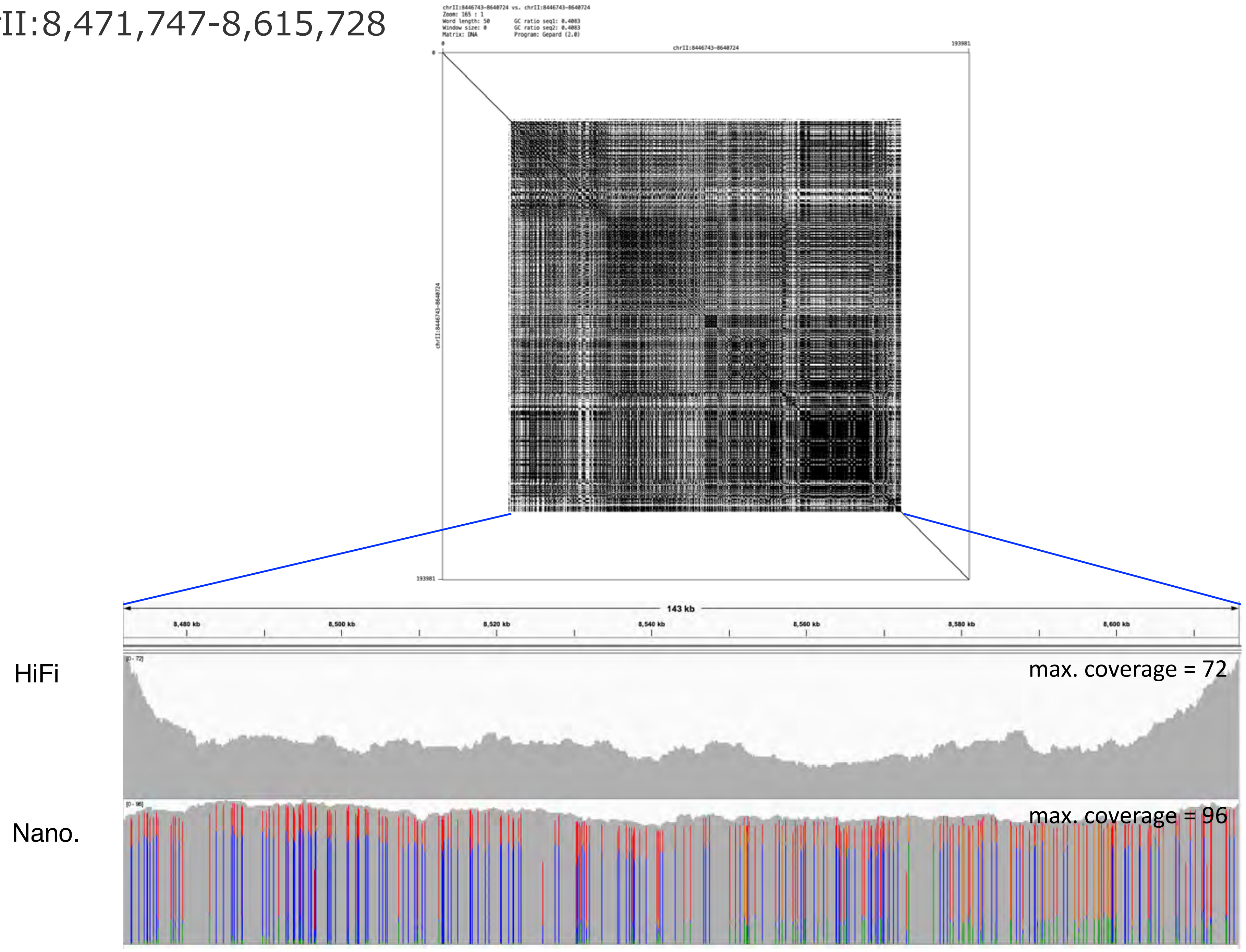

chrII:9,728,104-9,751,427

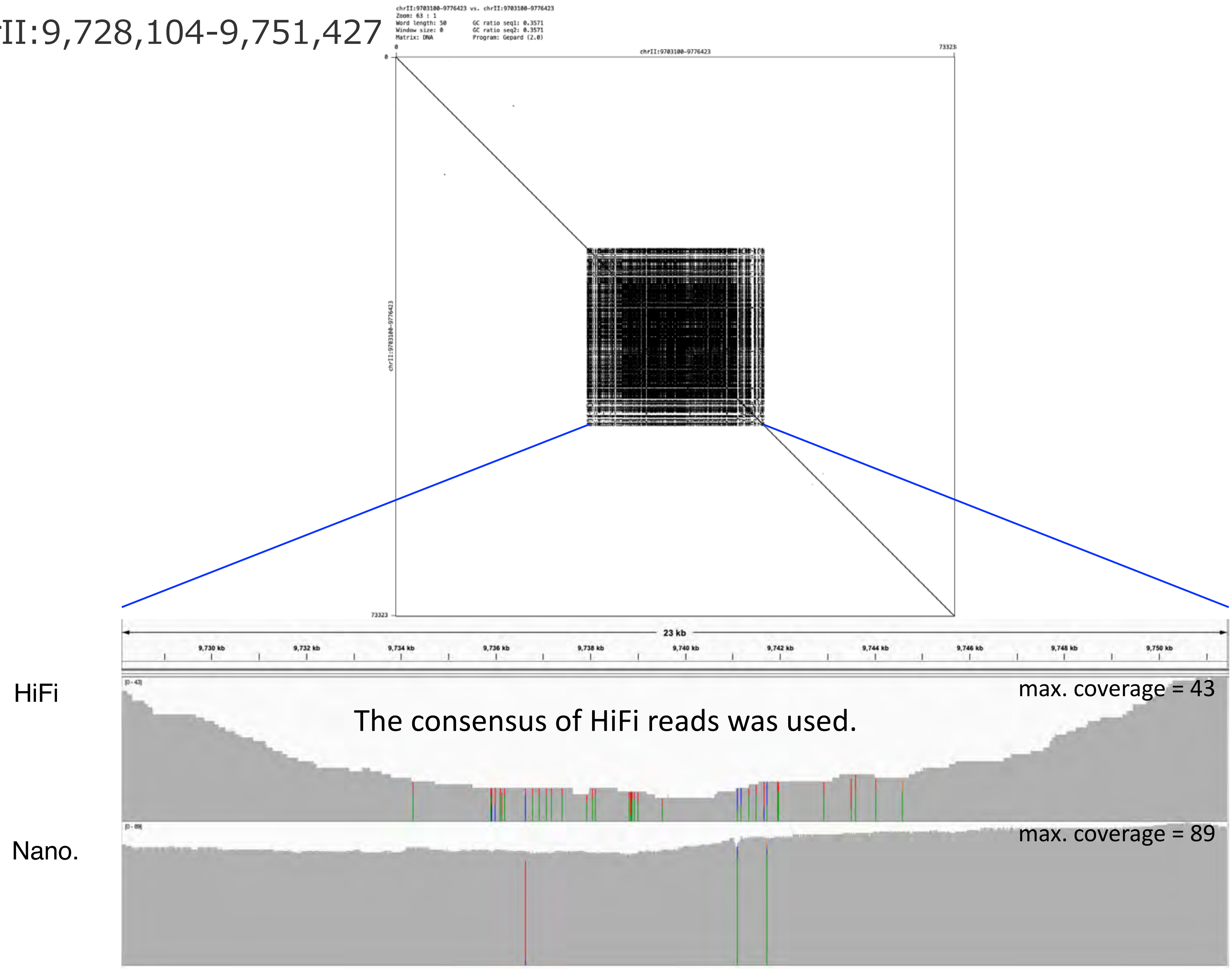

chrII:10,153,536-10,184,748

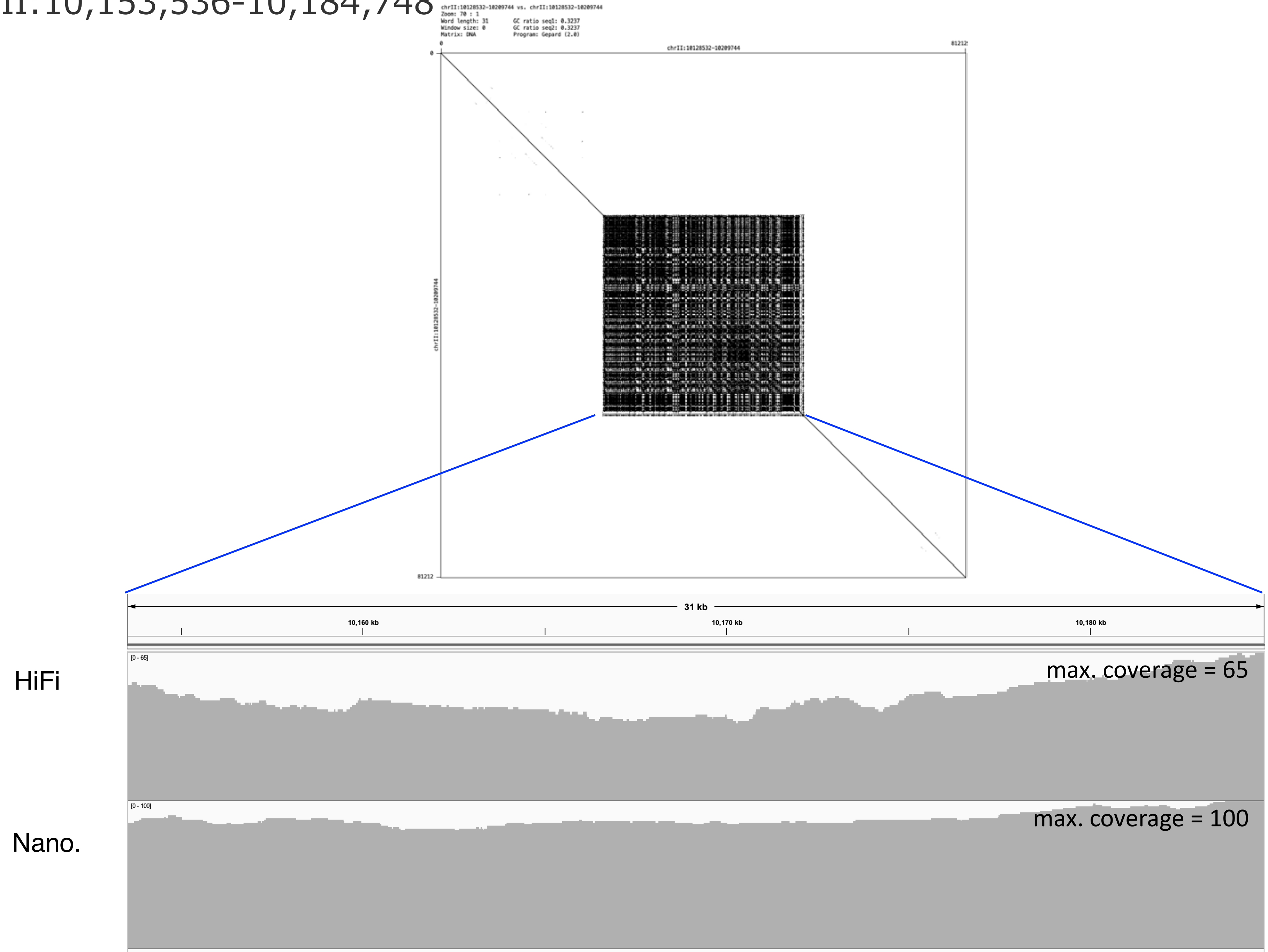

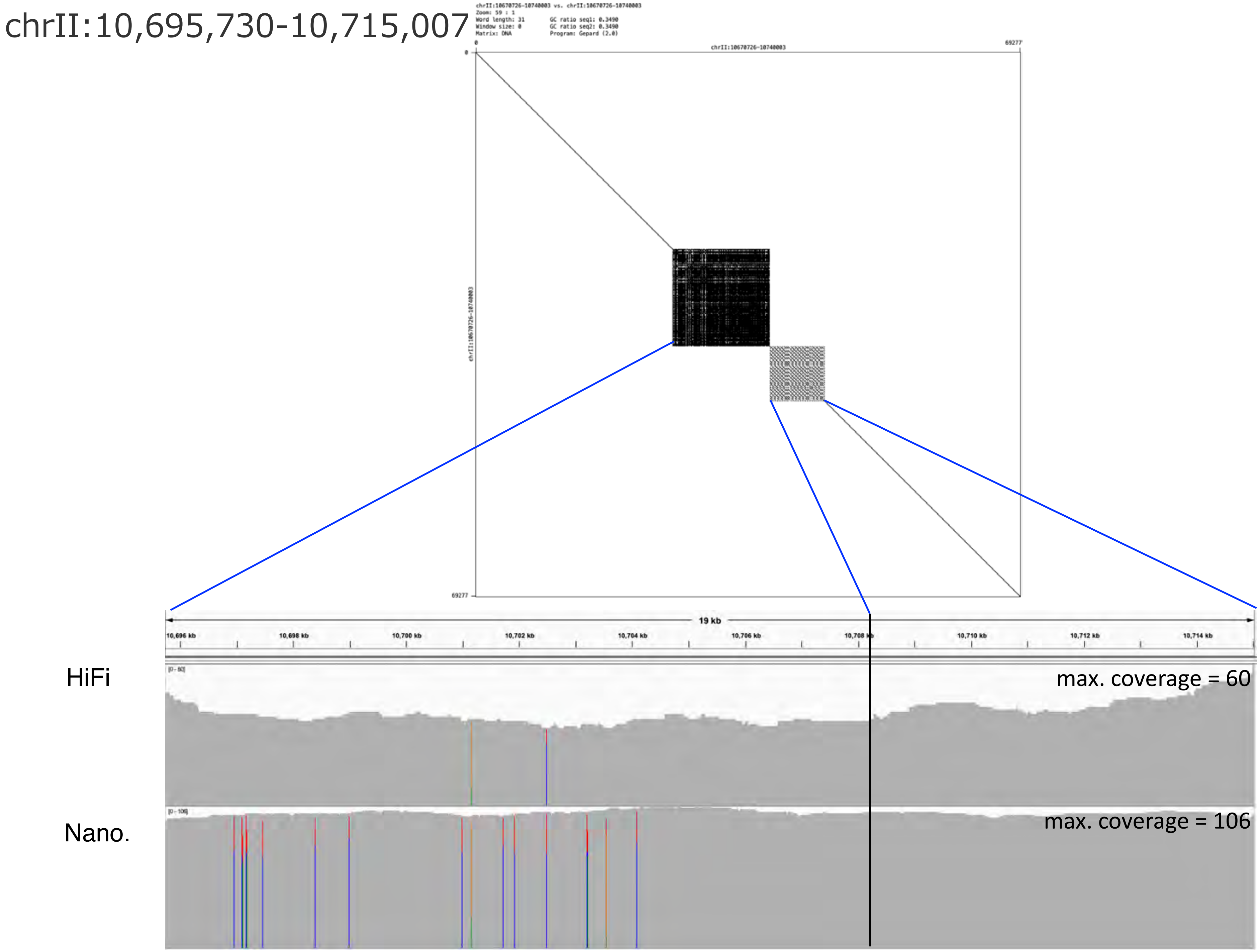
chrII:10,695,730-10,715,007
chrII:10670726-10740003 vs. chrII:10670726-10740003
Zoom: 59 : 1
Word length: 31
Window size: 0
Matrix: DNA
GC ratio seq1: 0.3490
GC ratio seq2: 0.3490
Program: Gepard (2.0)
0
chrII:10670726-10740003
69277
chrII:10670726-10740003
69277
19 kb
10,696 kb
10,698 kb
10,700 kb
10,702 kb
10,704 kb
10,706 kb
10,708 kb
10,710 kb
10,712 kb
10,714 kb
HiFi
max. coverage = 60
Nano.
max. coverage = 106

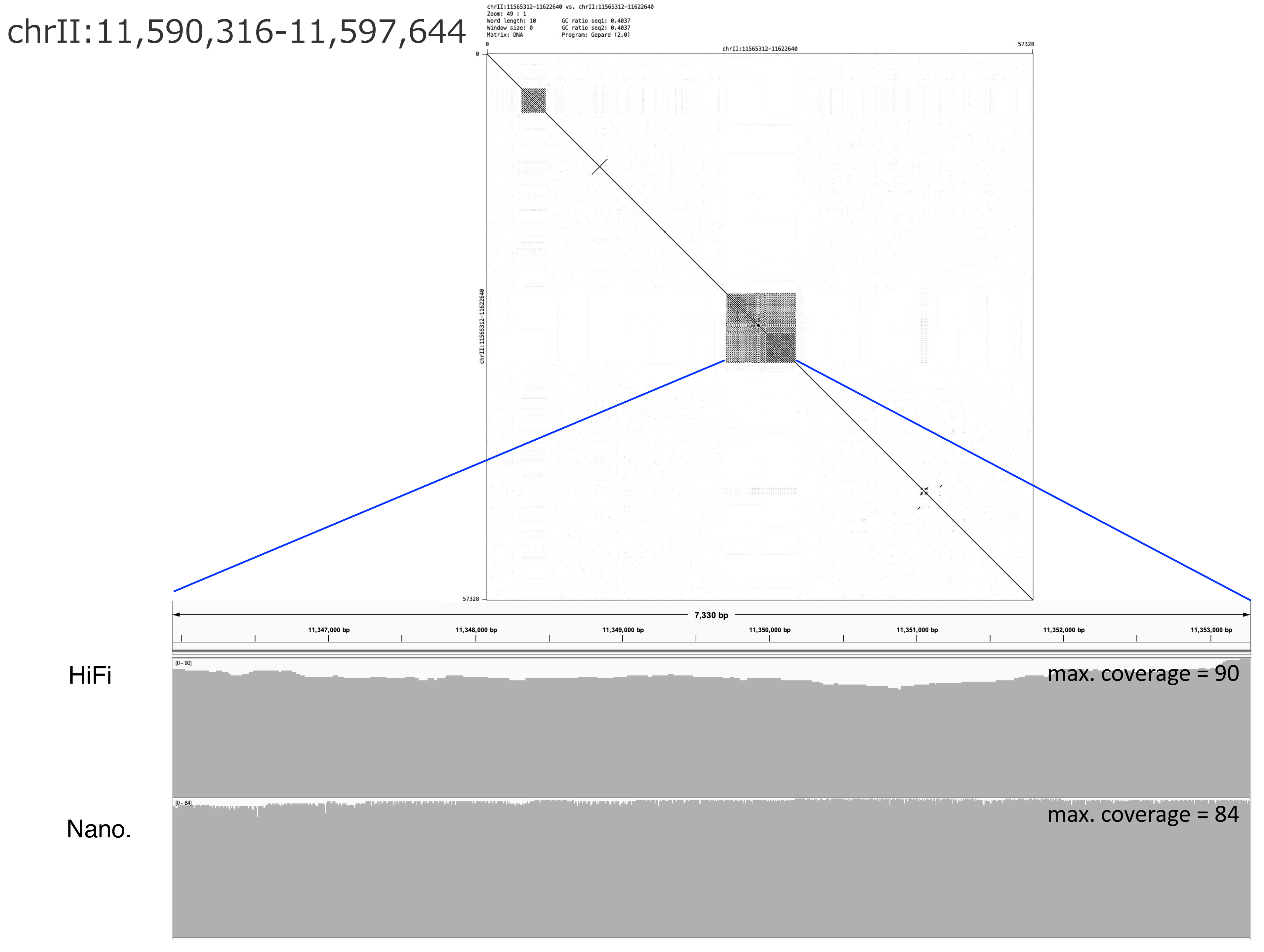
chrII:11,590,316-11,597,644
chrII:11565312-11622640 vs. chrII:11565312-11622640
Zoom: 49 : 1
Word length: 10
Window size: 0
Matrix: DNA
GC ratio seq1: 0.4037
GC ratio seq2: 0.4037
Program: Gepard (2.0)
chrII:11565312-11622640
0
57328
7,330 bp
11,347,000 bp
11,348,000 bp
11,349,000 bp
11,350,000 bp
11,351,000 bp
11,352,000 bp
11,353,000 bp
HiFi
[0 - 90]
max. coverage = 90
Nano.
[0 - 84]
max. coverage = 84

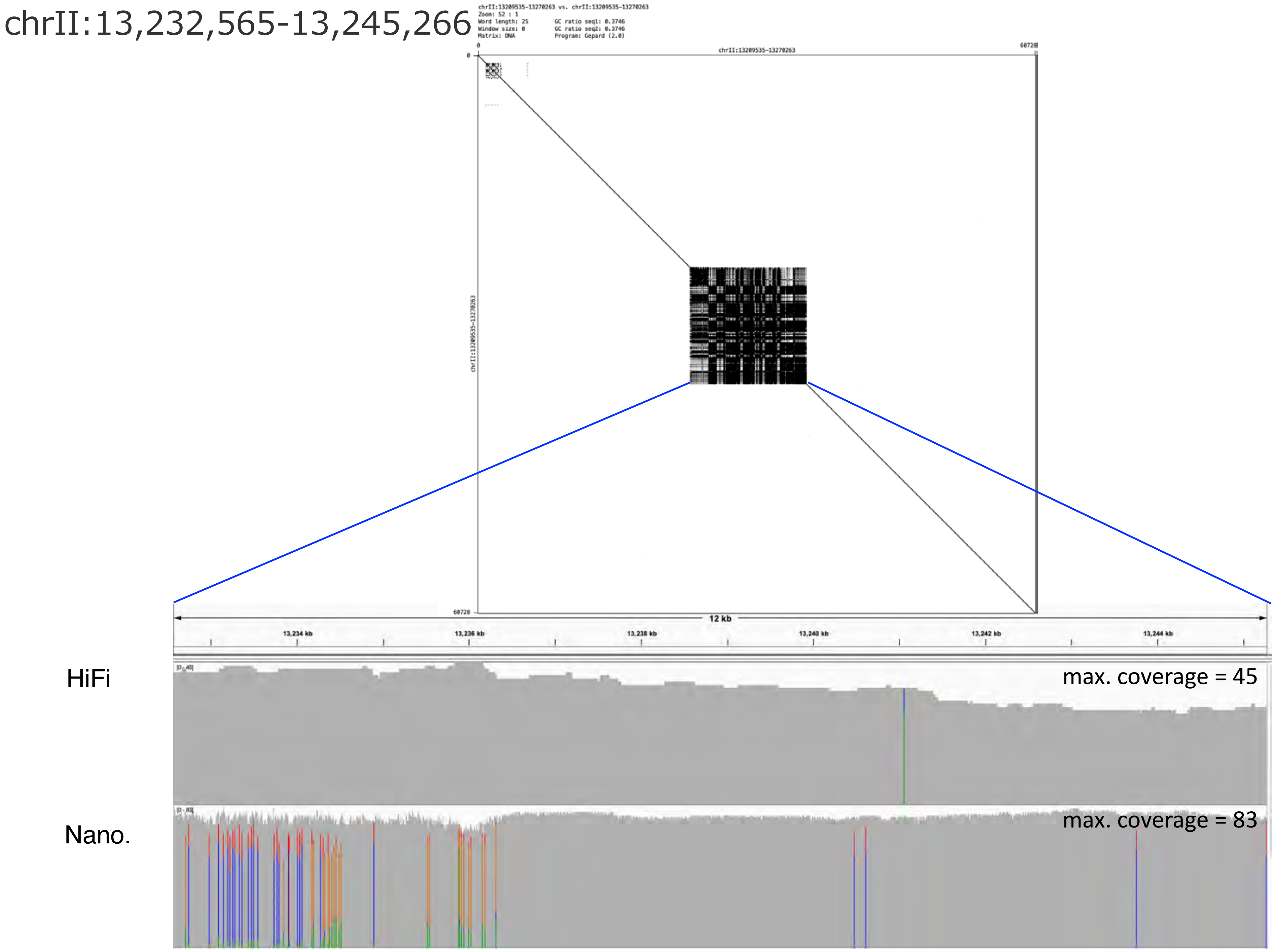
chrII:13,232,565-13,245,266
chrII:13209535-13270263 vs. chrII:13209535-13270263
Zoom: 52 : 1
Word length: 25
Window size: 0
Matrix: DNA
GC ratio seq1: 0.3746
GC ratio seq2: 0.3746
Program: Gepard (2.0)
chrII:13209535-13270263
60728
12 kb
13,234 kb
13,236 kb
13,238 kb
13,240 kb
13,242 kb
13,244 kb
HiFi
max. coverage = 45
Nano.
max. coverage = 83

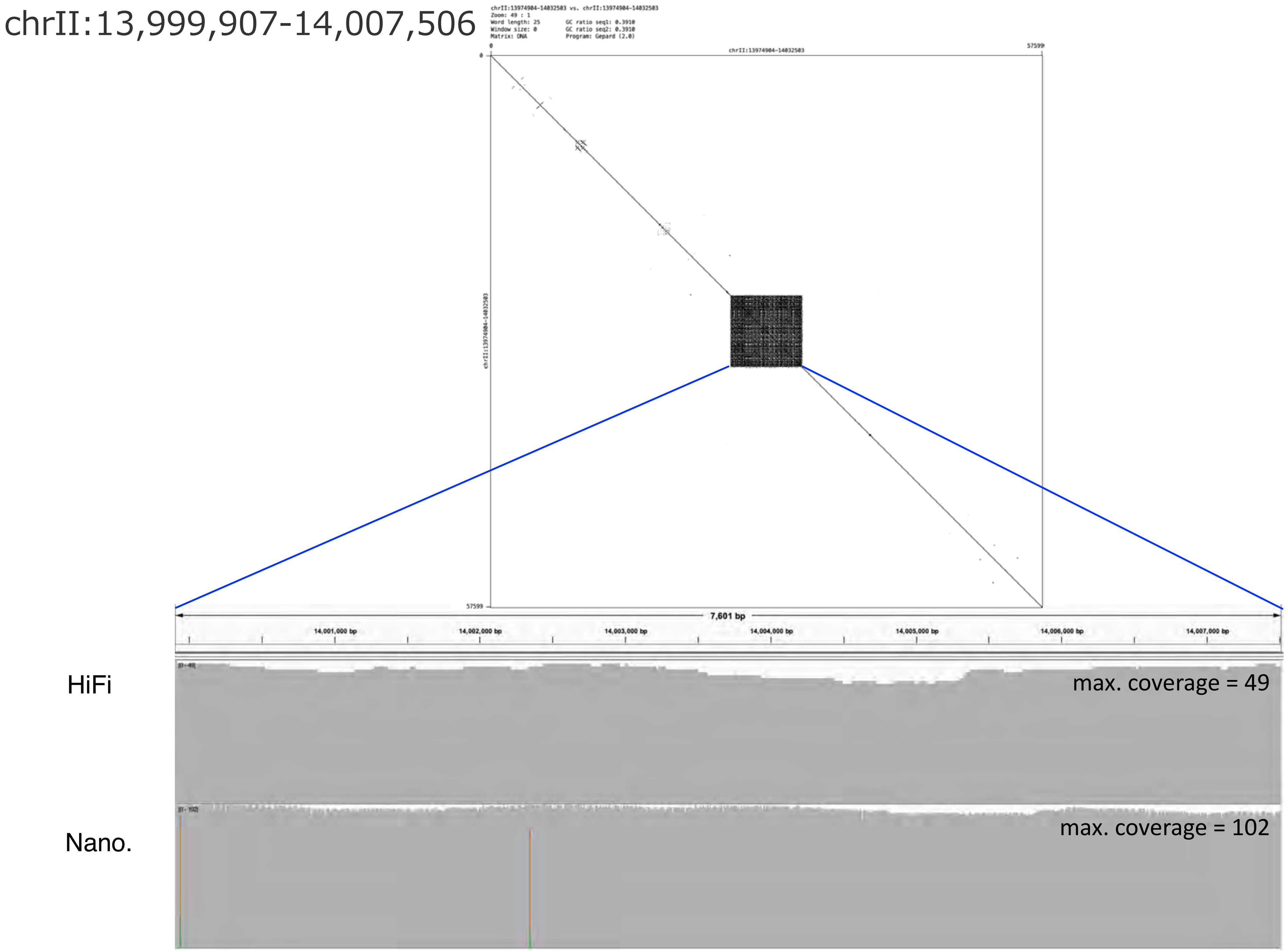

chrII:13,999,907-14,007,506
chrII:13974904-14032503 vs. chrII:13974904-14032503
Zoom: 49 : 1
Word length: 25
Window size: 0
Matrix: DNA
GC ratio seq1: 0.3910
GC ratio seq2: 0.3910
Program: Gepard (2.0)
0
chrII:13974904-14032503
57599
7,601 bp
14,001,000 bp
14,002,000 bp
14,003,000 bp
14,004,000 bp
14,005,000 bp
14,006,000 bp
14,007,000 bp
HiFi
max. coverage = 49
Nano.
max. coverage = 102

chrII:14,403,563-14,449,398

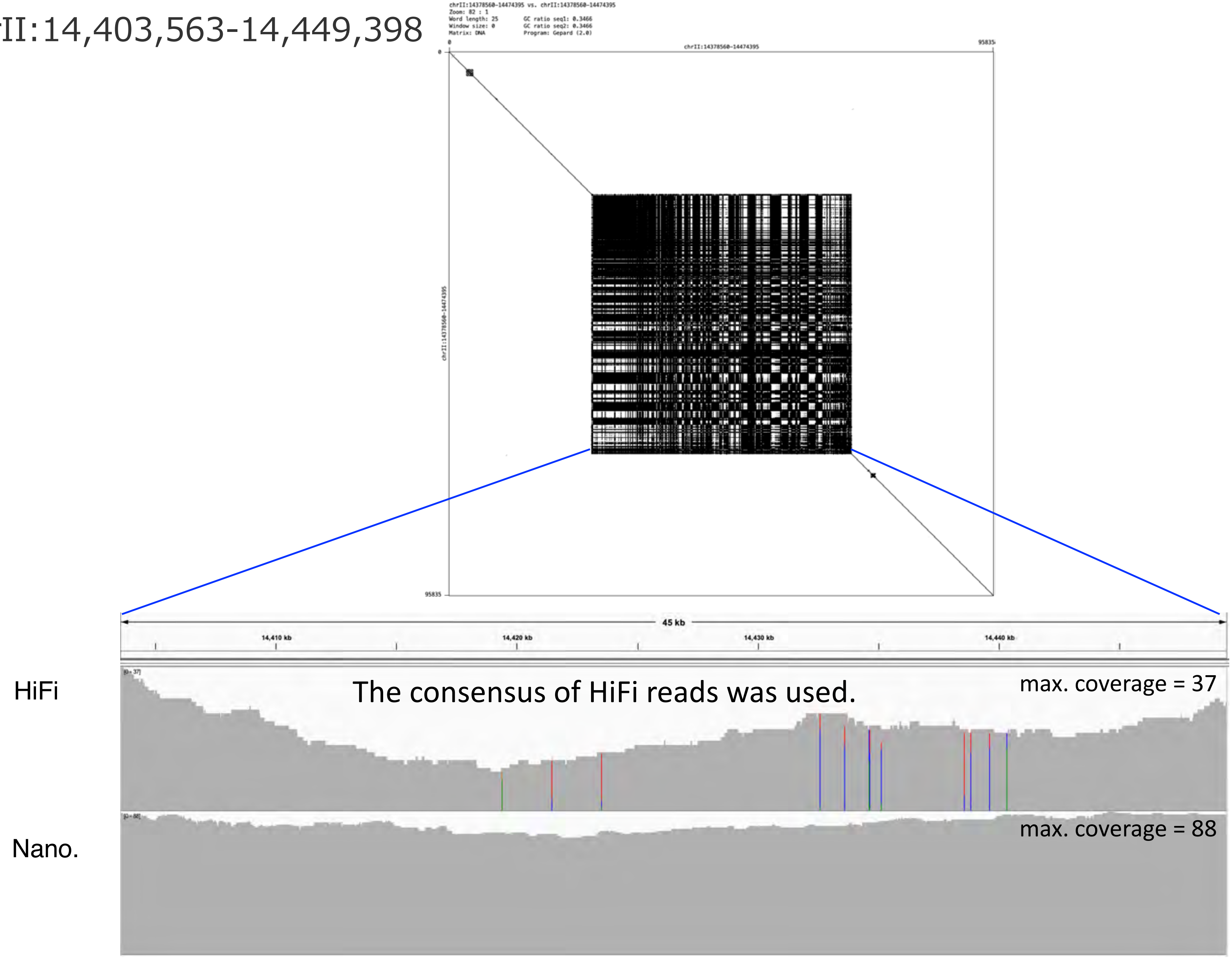

chrII:14,787,048-14,842,635

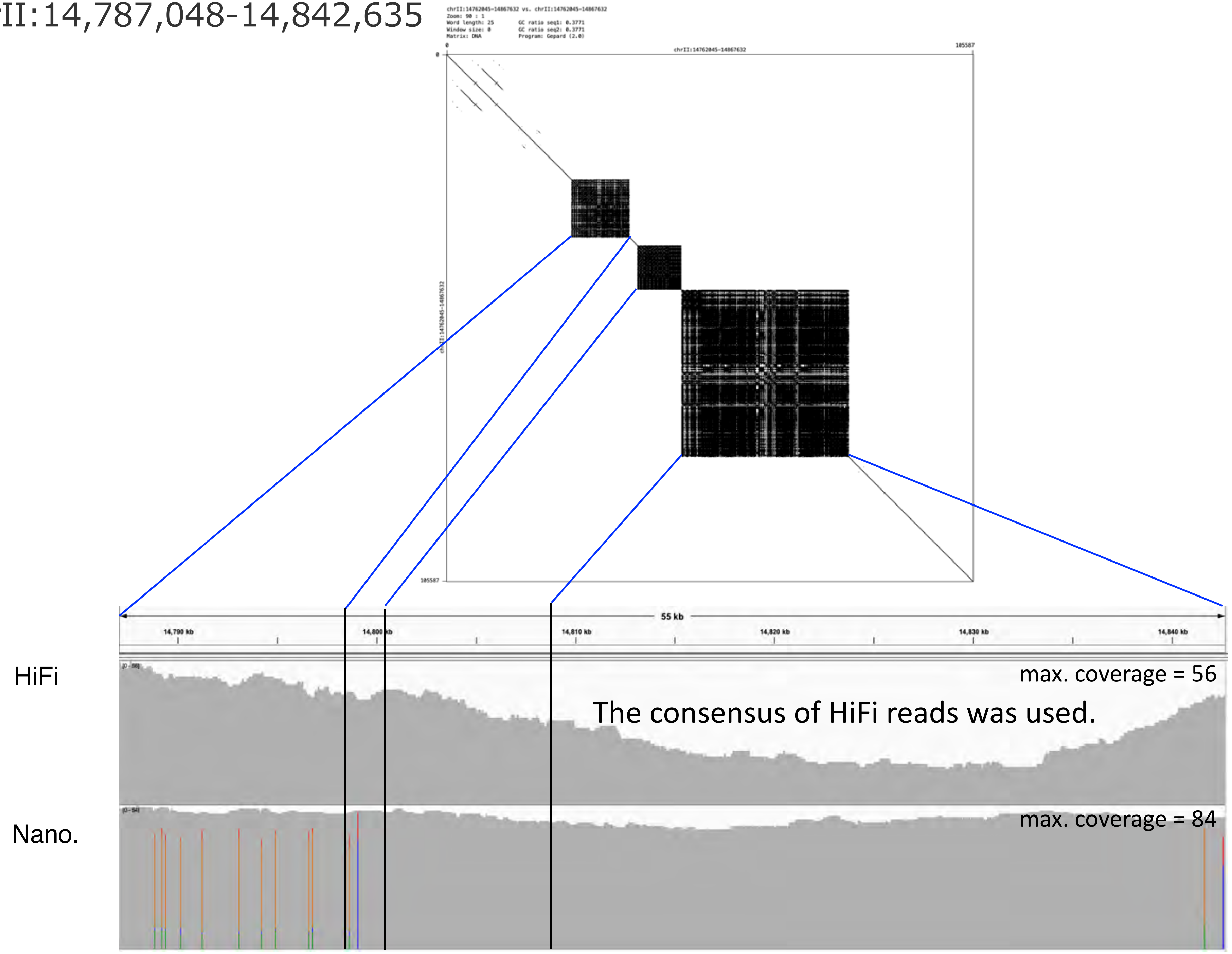

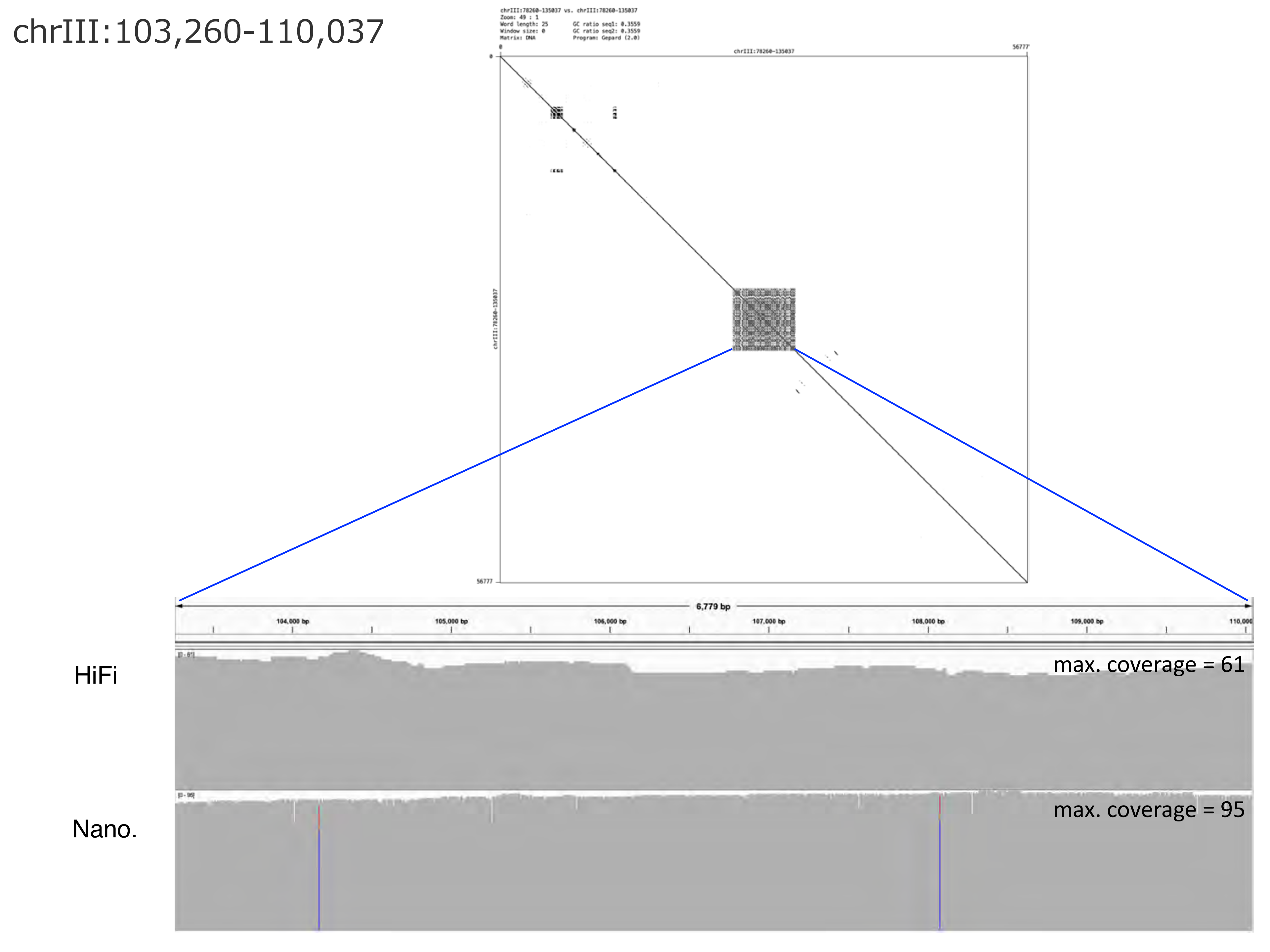

chrIII:103,260-110,037
chrIII:78260-135037 vs. chrIII:78260-135037
Zoom: 49 : 1
Word length: 25
Window size: 0
Matrix: DNA
GC ratio seq1: 0.3559
GC ratio seq2: 0.3559
Program: Gepard (2.0)
0
56777
chrIII:78260-135037
6,779 bp
104,000 bp
105,000 bp
106,000 bp
107,000 bp
108,000 bp
109,000 bp
HiFi
max. coverage = 61
Nano.
max. coverage = 95

### chrIII:279,439-284,776

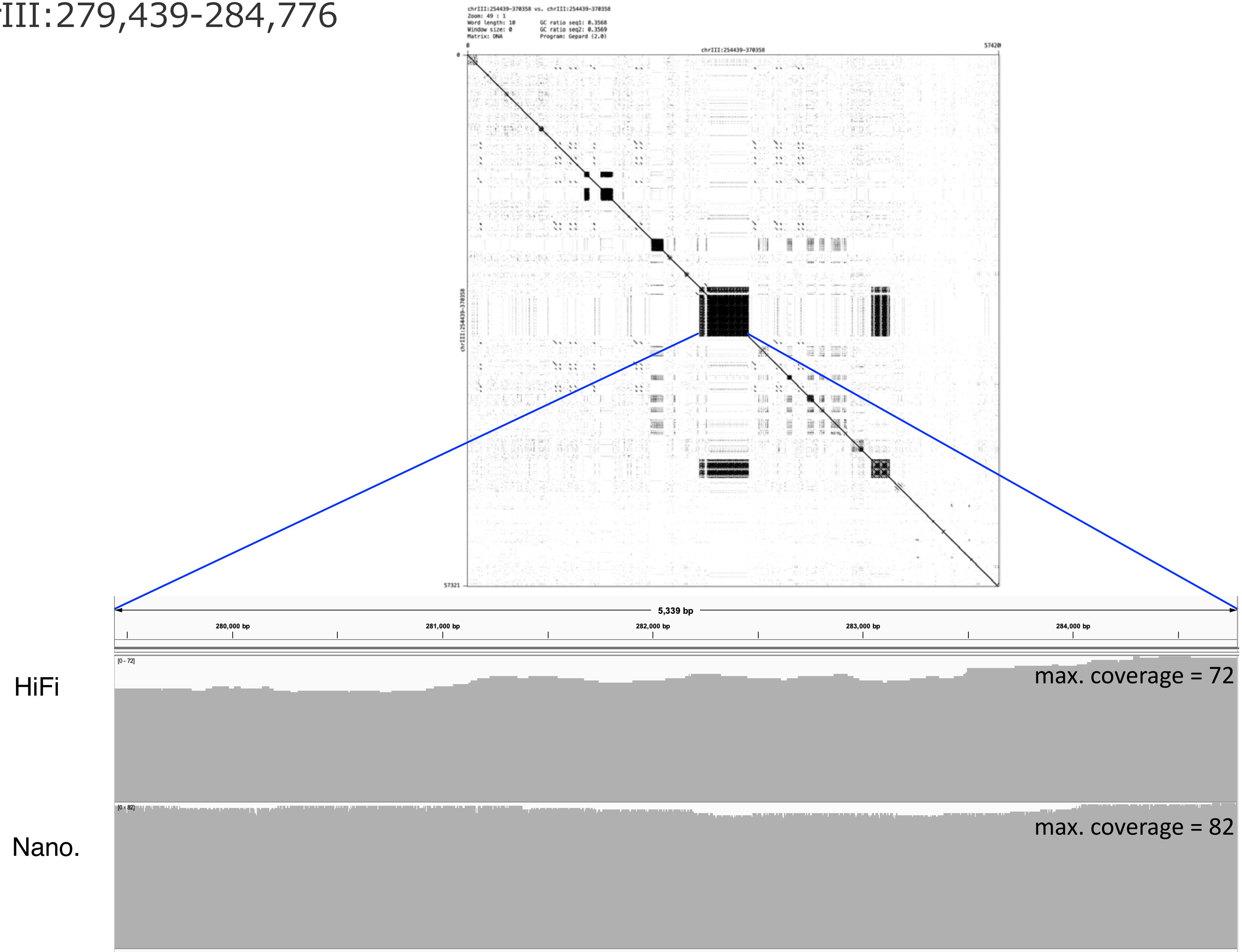

chrIII:329,510-345,358

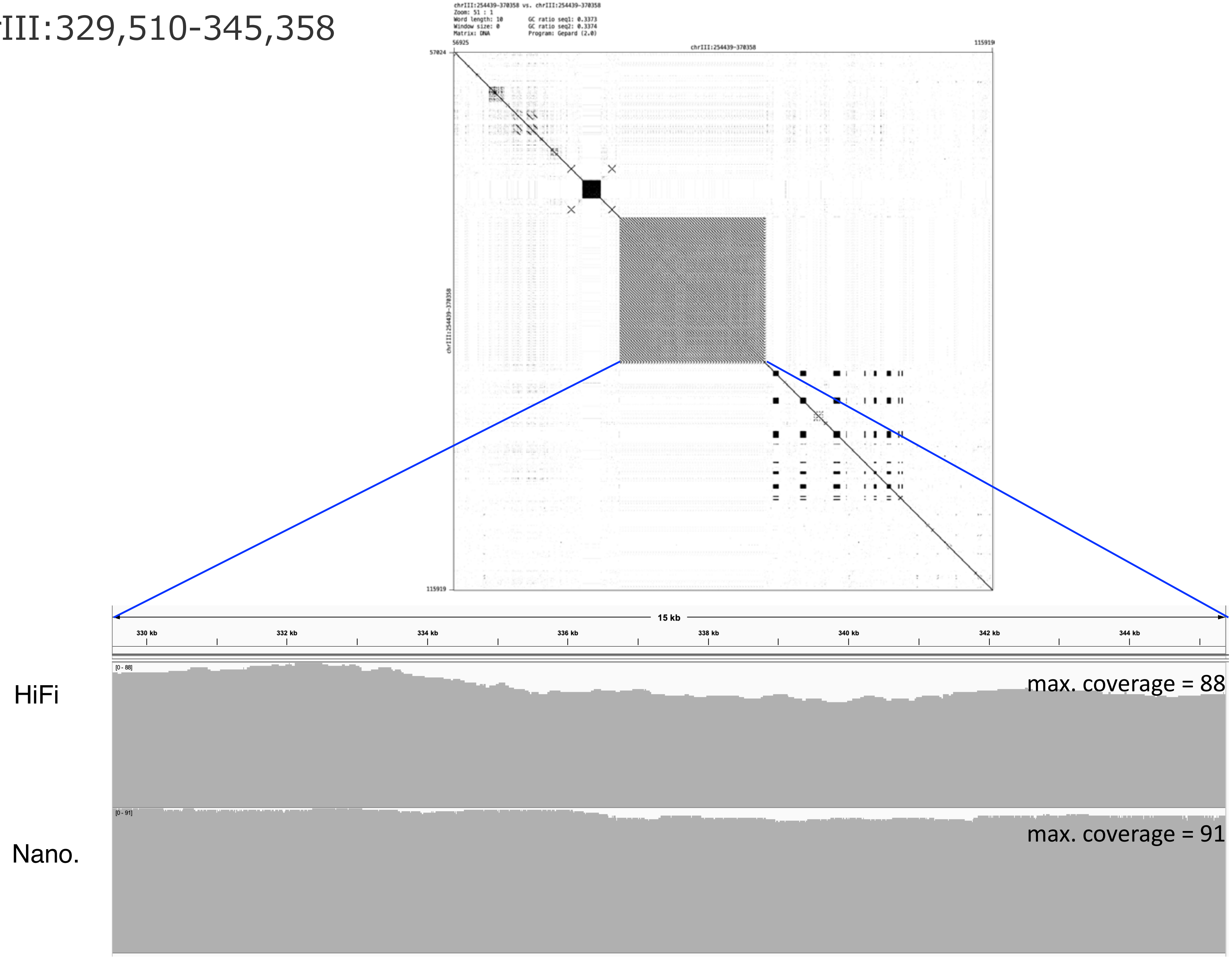

chrIII:466,232-479,321

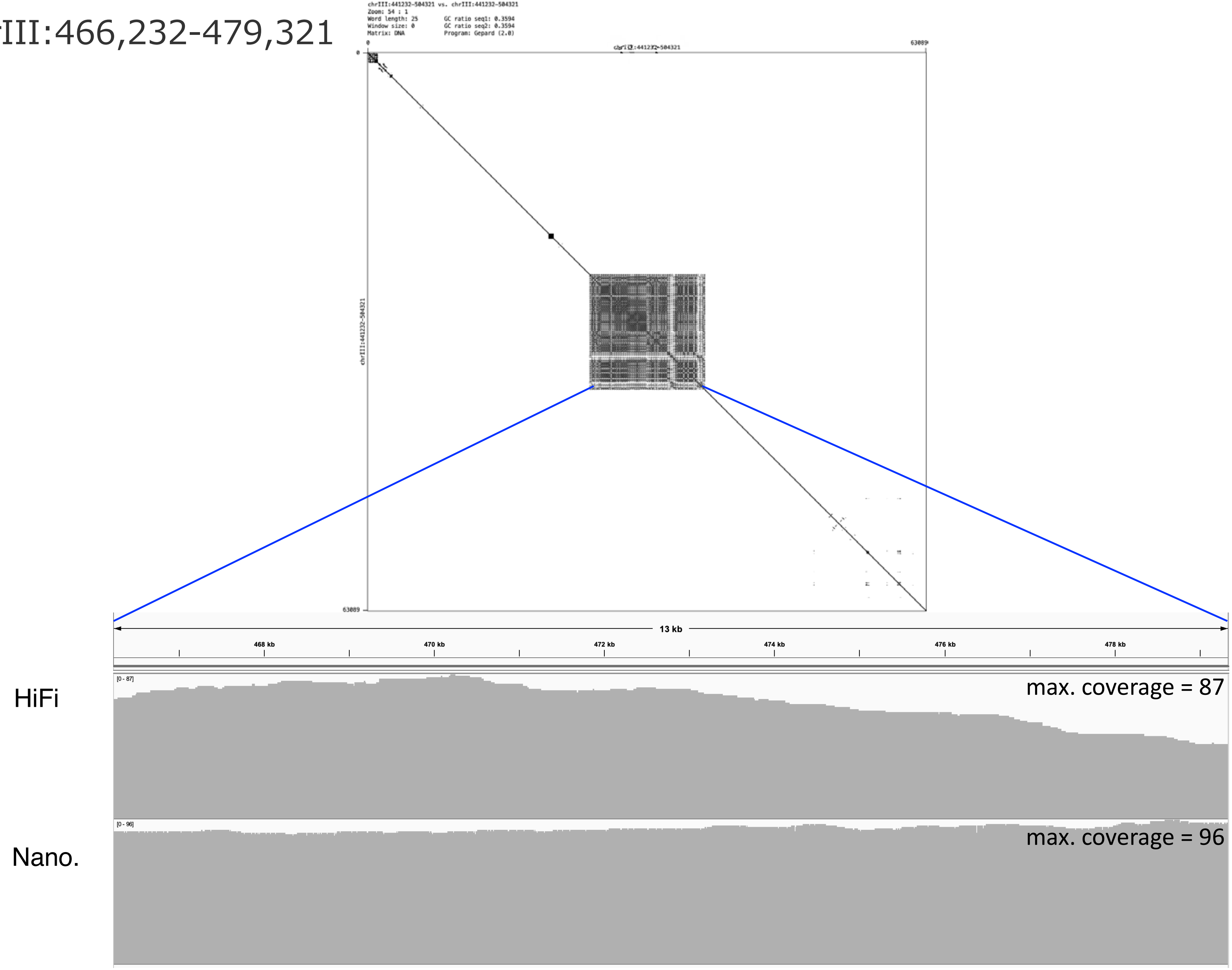

### chrIII:960,210-1,010,857

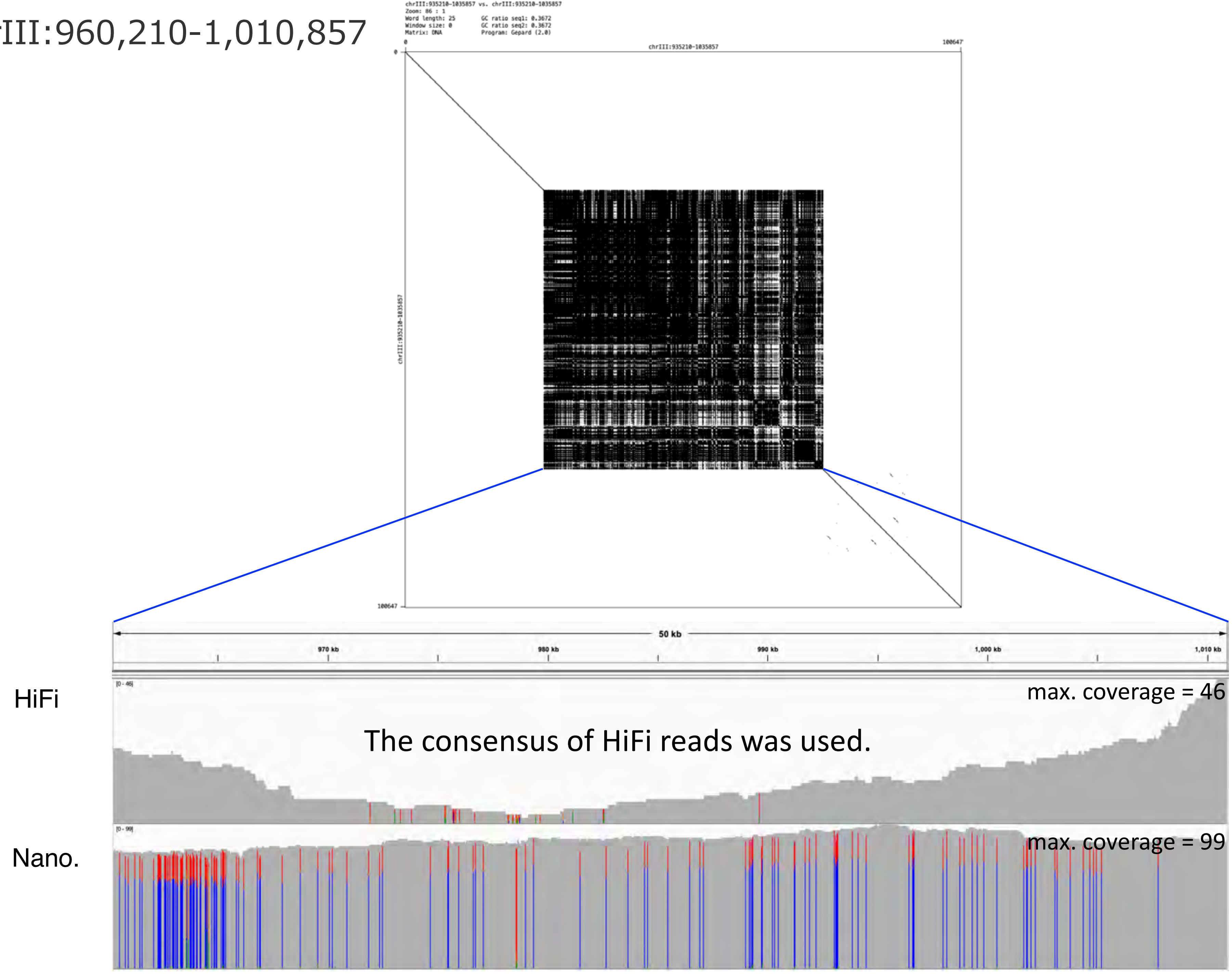

### chrIII:1,096,478-1,169,029

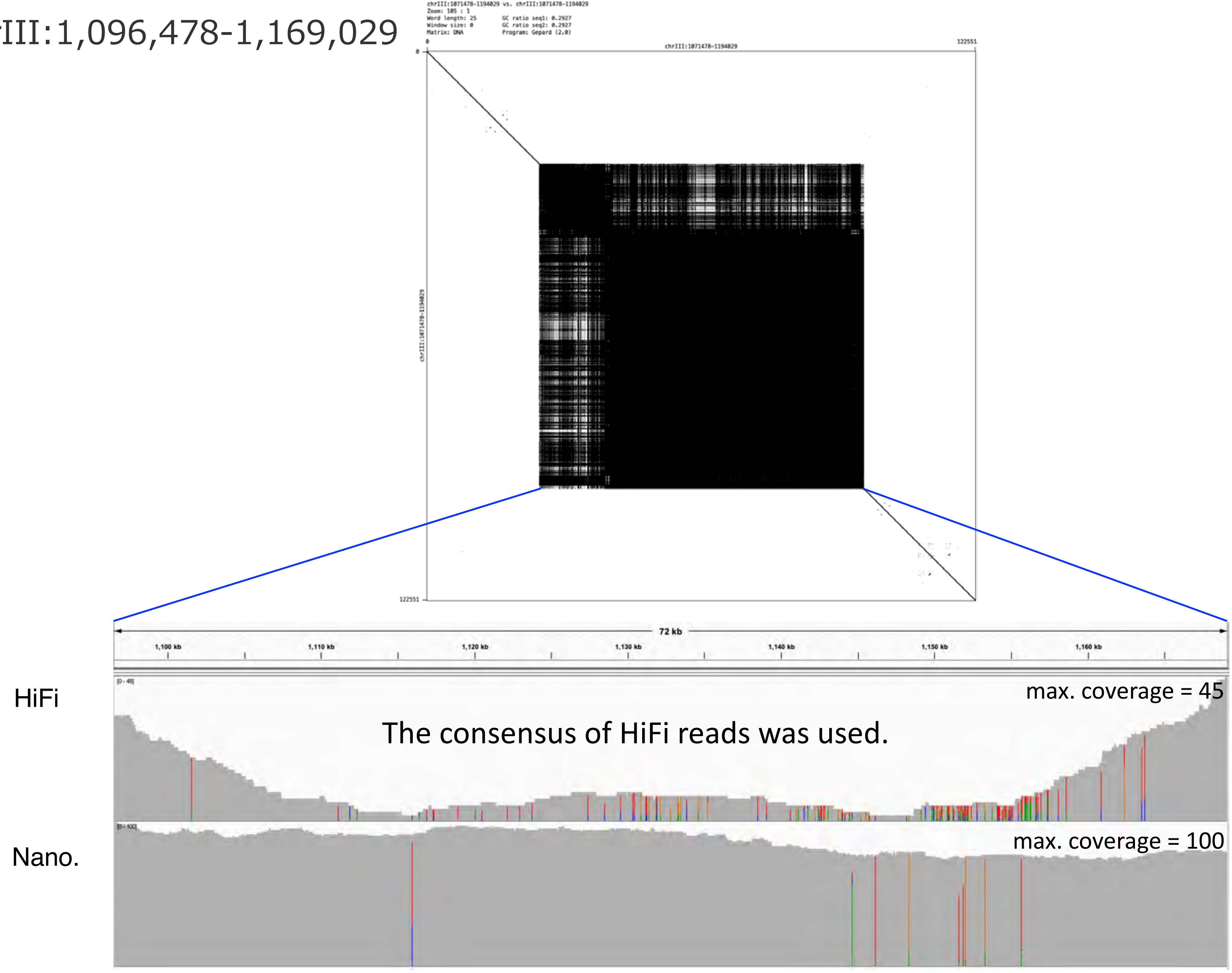

### chrIII:1,457,429-1,555,246

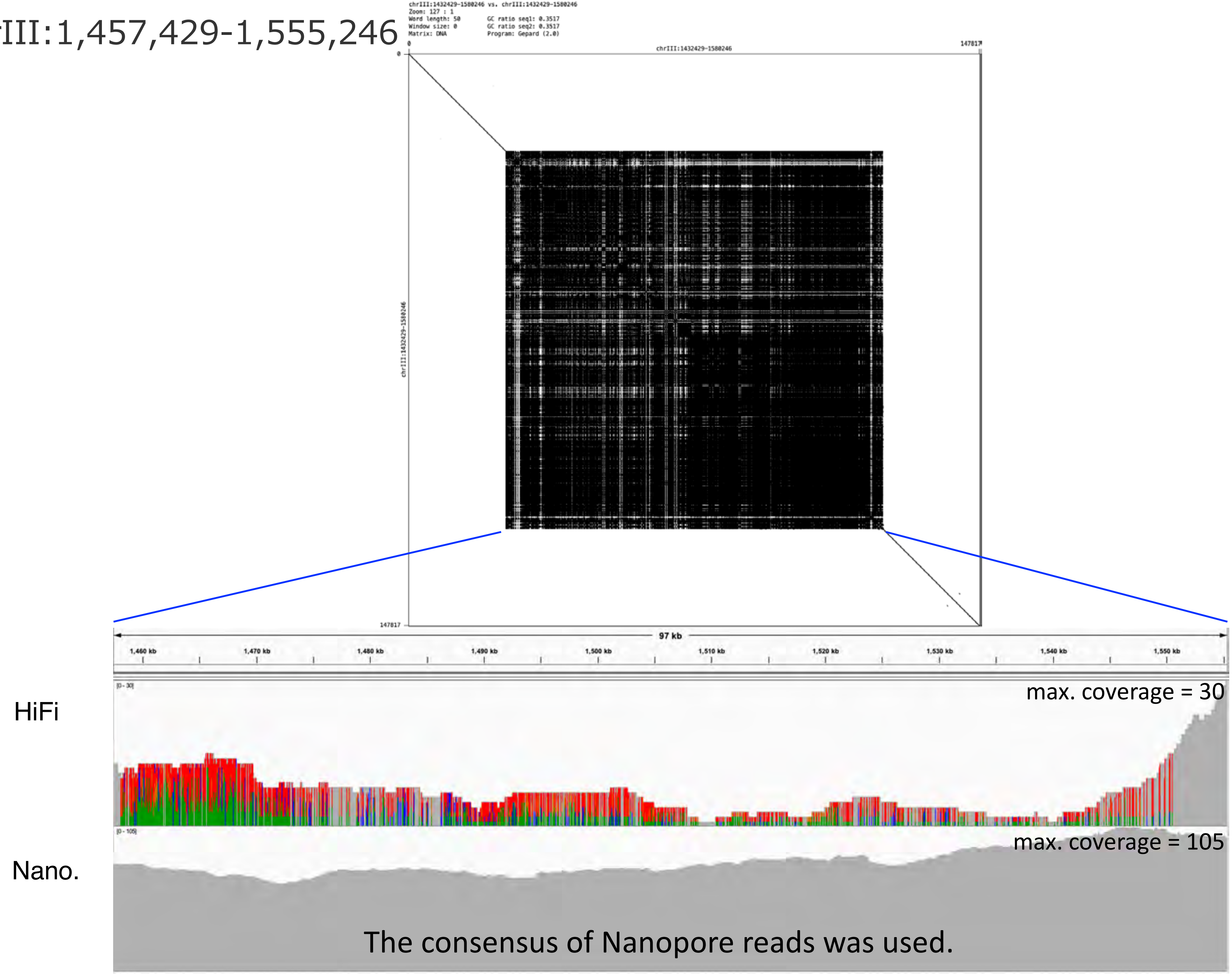

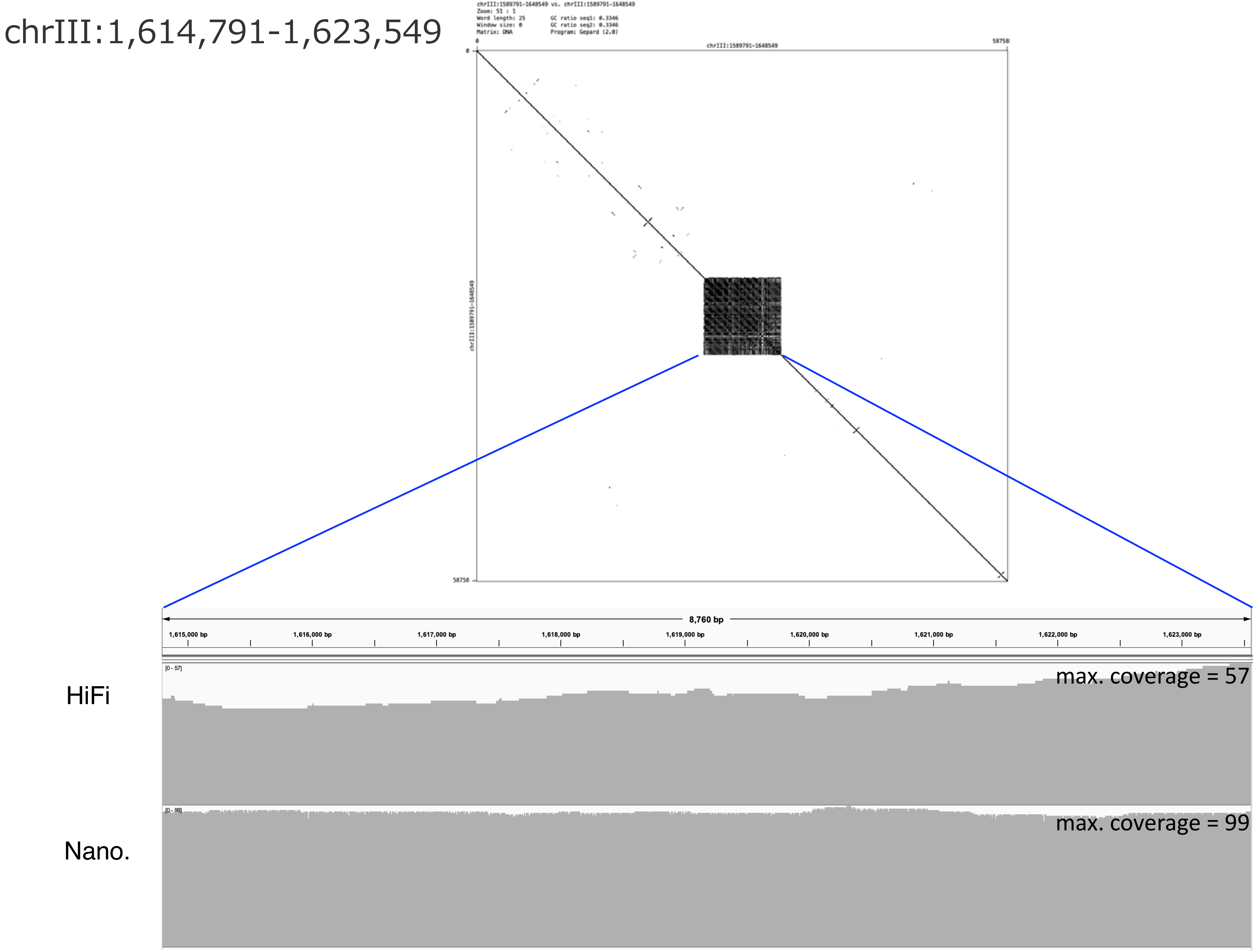
chrIII:1,614,791-1,623,549
chrIII:1589791-1648549 vs. chrIII:1589791-1648549
Zoom: 51 : 1
Word length: 25
Window size: 0
Matrix: DNA
GC ratio seq1: 0.3346
GC ratio seq2: 0.3346
Program: Gepard (2.0)
chrIII:1589791-1648549
0
58758
8,760 bp
1,615,000 bp
1,616,000 bp
1,617,000 bp
1,618,000 bp
1,619,000 bp
1,620,000 bp
1,621,000 bp
1,622,000 bp
1,623,000 bp
HiFi
[0 - 57]
max. coverage = 57
Nano.
[0 - 99]
max. coverage = 99

chrIII:2,332,942-2,371,214

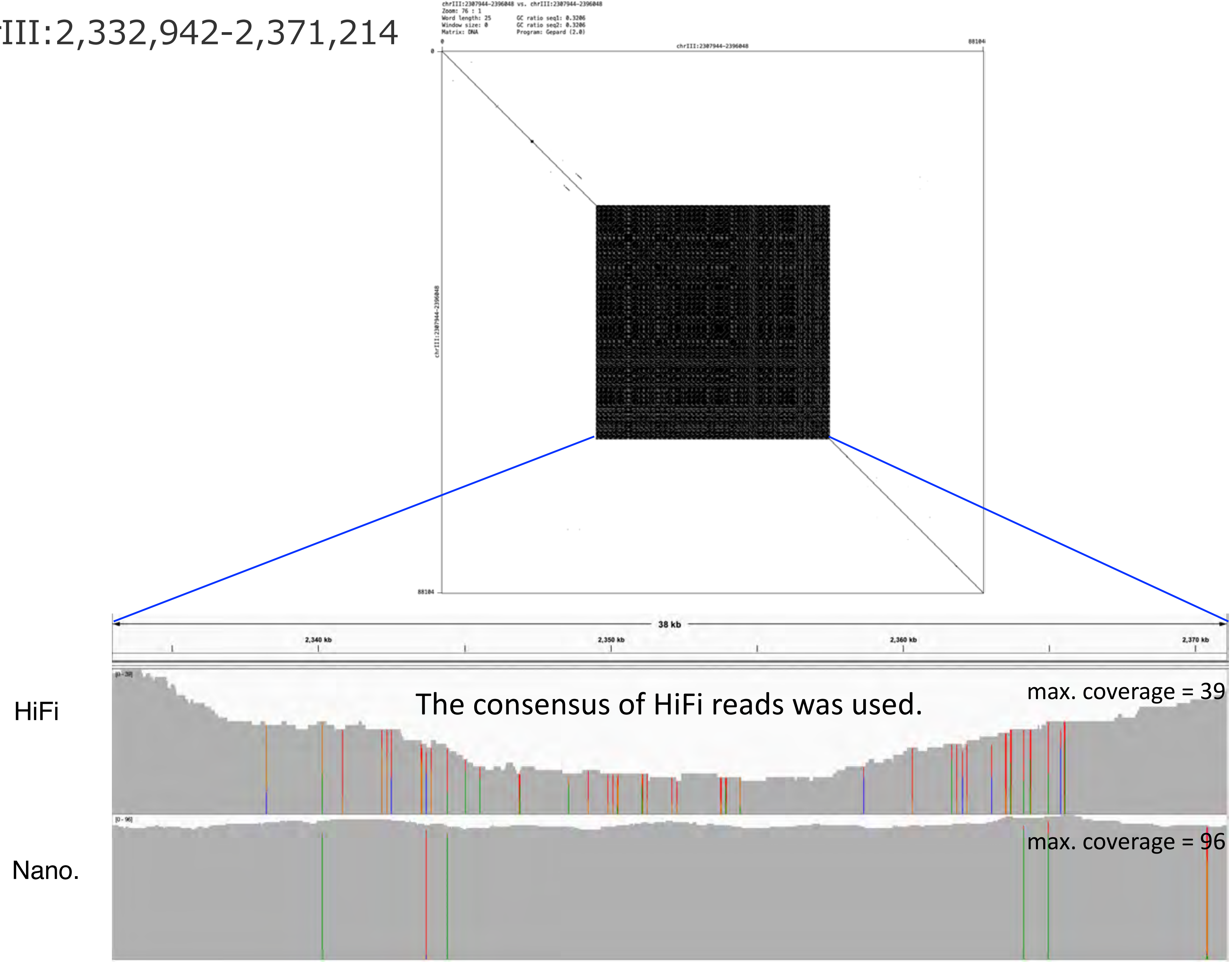

### chrIII:2,791,638-2,846,089

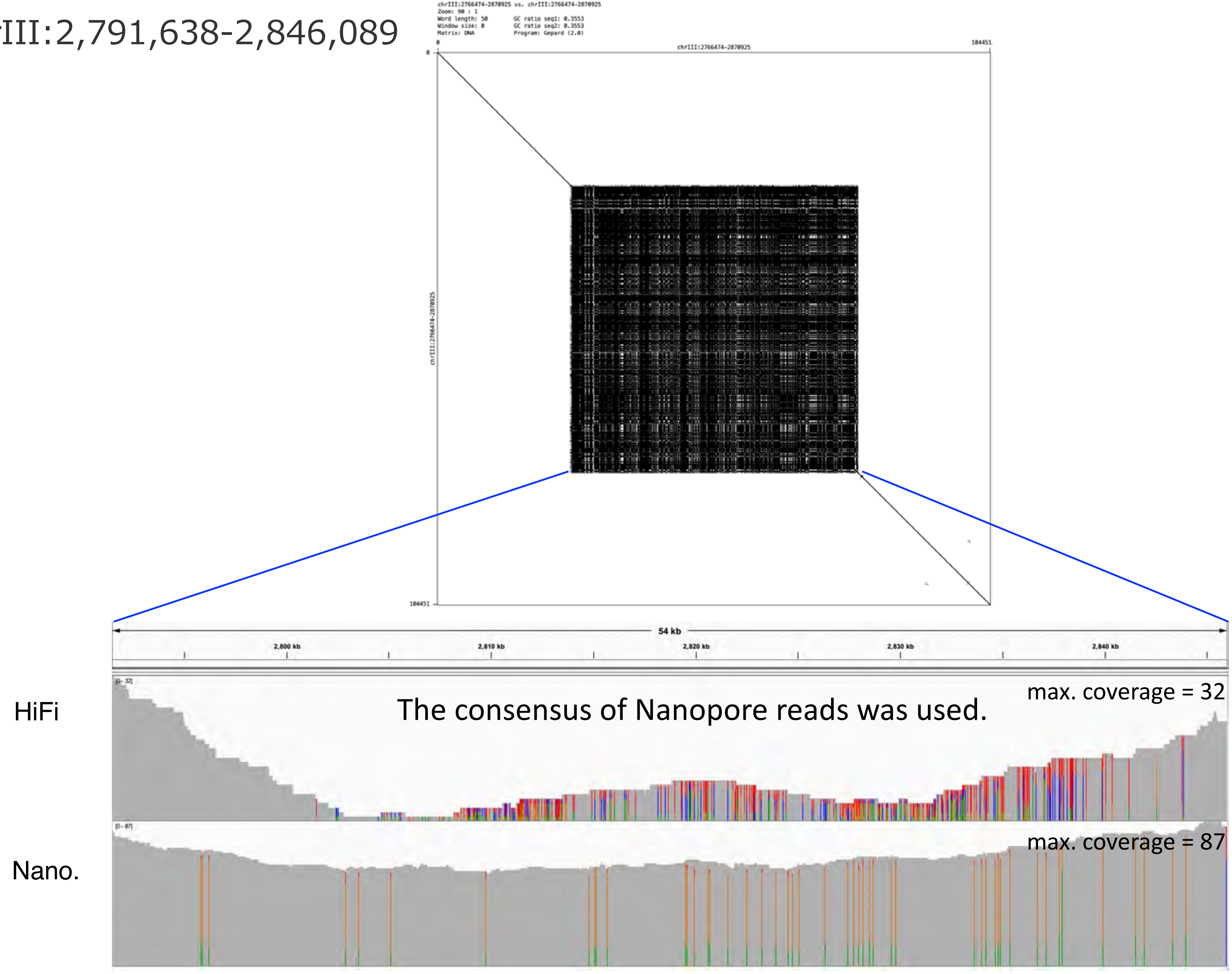

chrIII:3,681,598-3,704,204

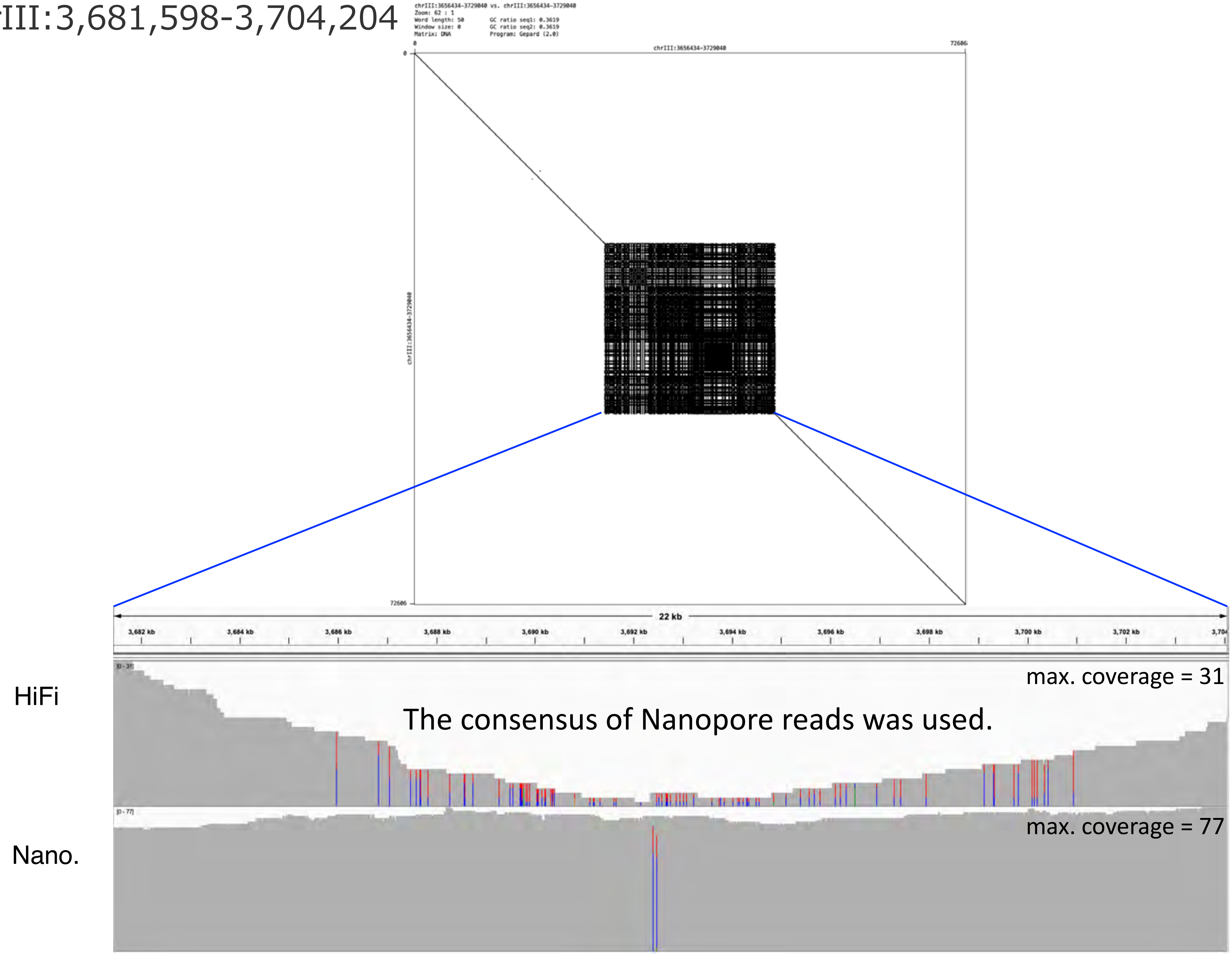

### chrIII:5,728,283-5,804,341

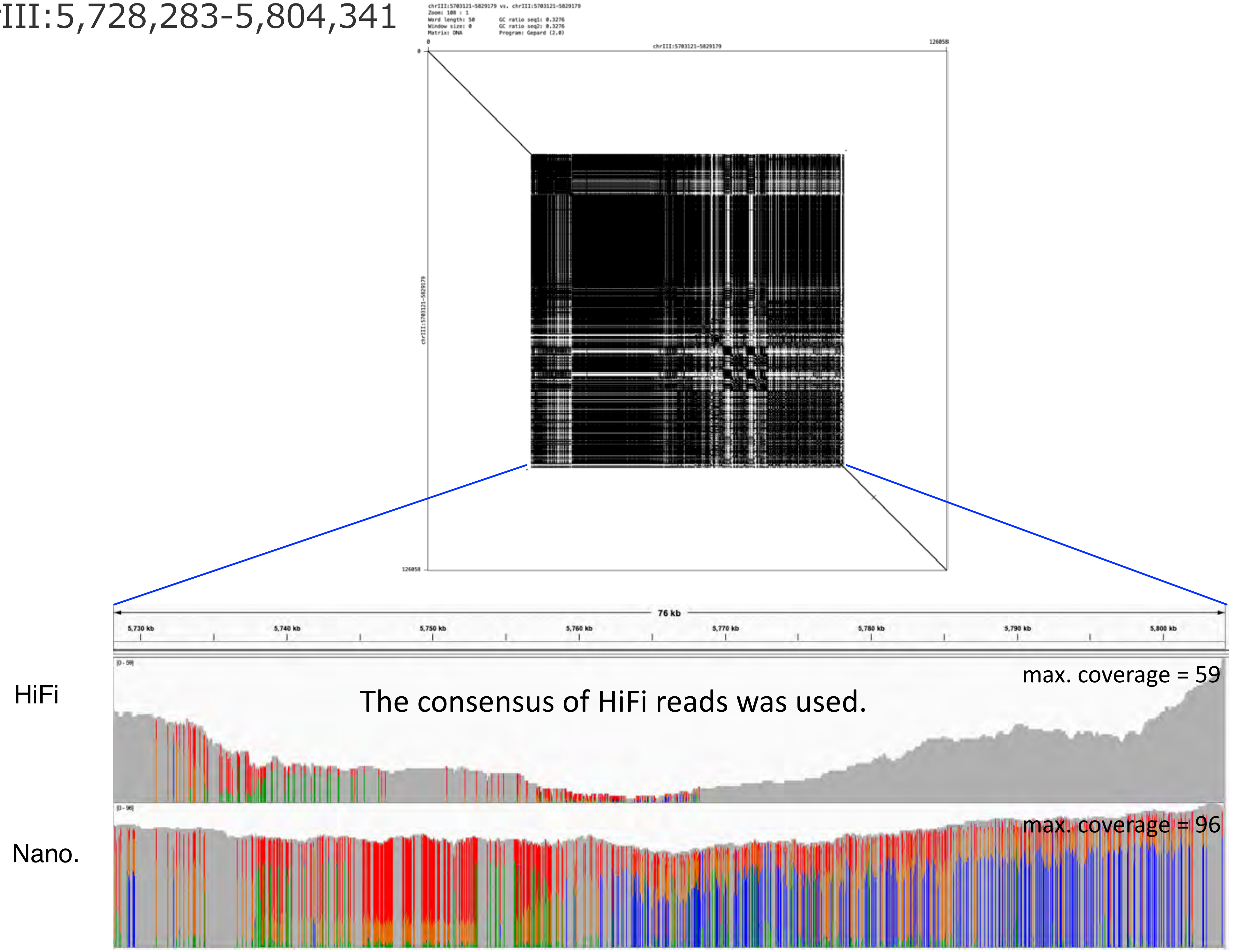

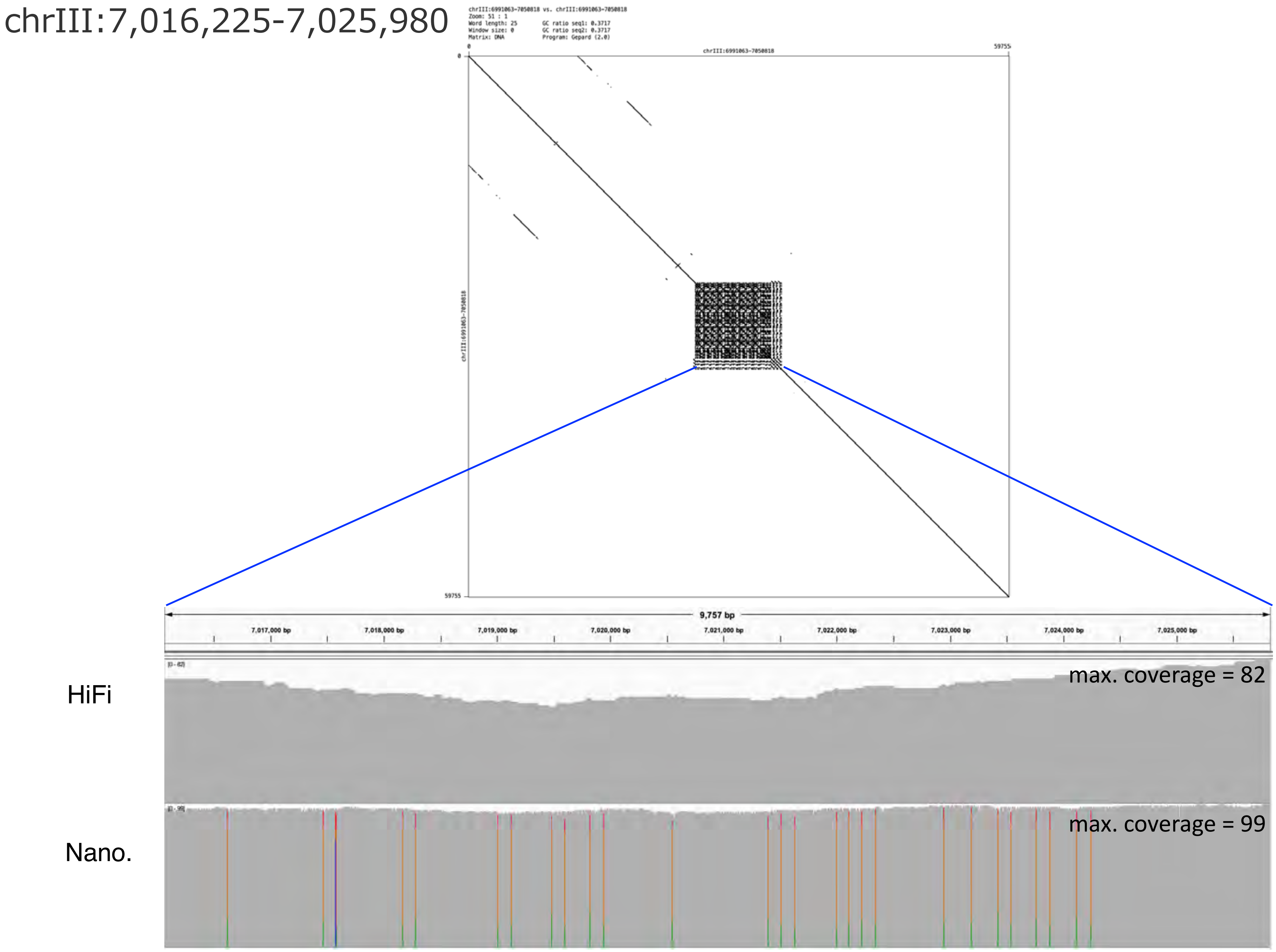

chrIII:7,016,225-7,025,980
chrIII:6991063-7050818 vs. chrIII:6991063-7050818
Zoom: 51 : 1
Word length: 25
Window size: 0
Matrix: DNA
GC ratio seq1: 0.3717
GC ratio seq2: 0.3717
Program: Gepard (2.0)
chrIII:6991063-7050818
0
59755
9,757 bp
7,017,000 bp
7,018,000 bp
7,019,000 bp
7,020,000 bp
7,021,000 bp
7,022,000 bp
7,023,000 bp
7,024,000 bp
7,025,000 bp
HiFi
max. coverage = 82
Nano.
max. coverage = 99

### chrIII:7,856,863-8,031,900

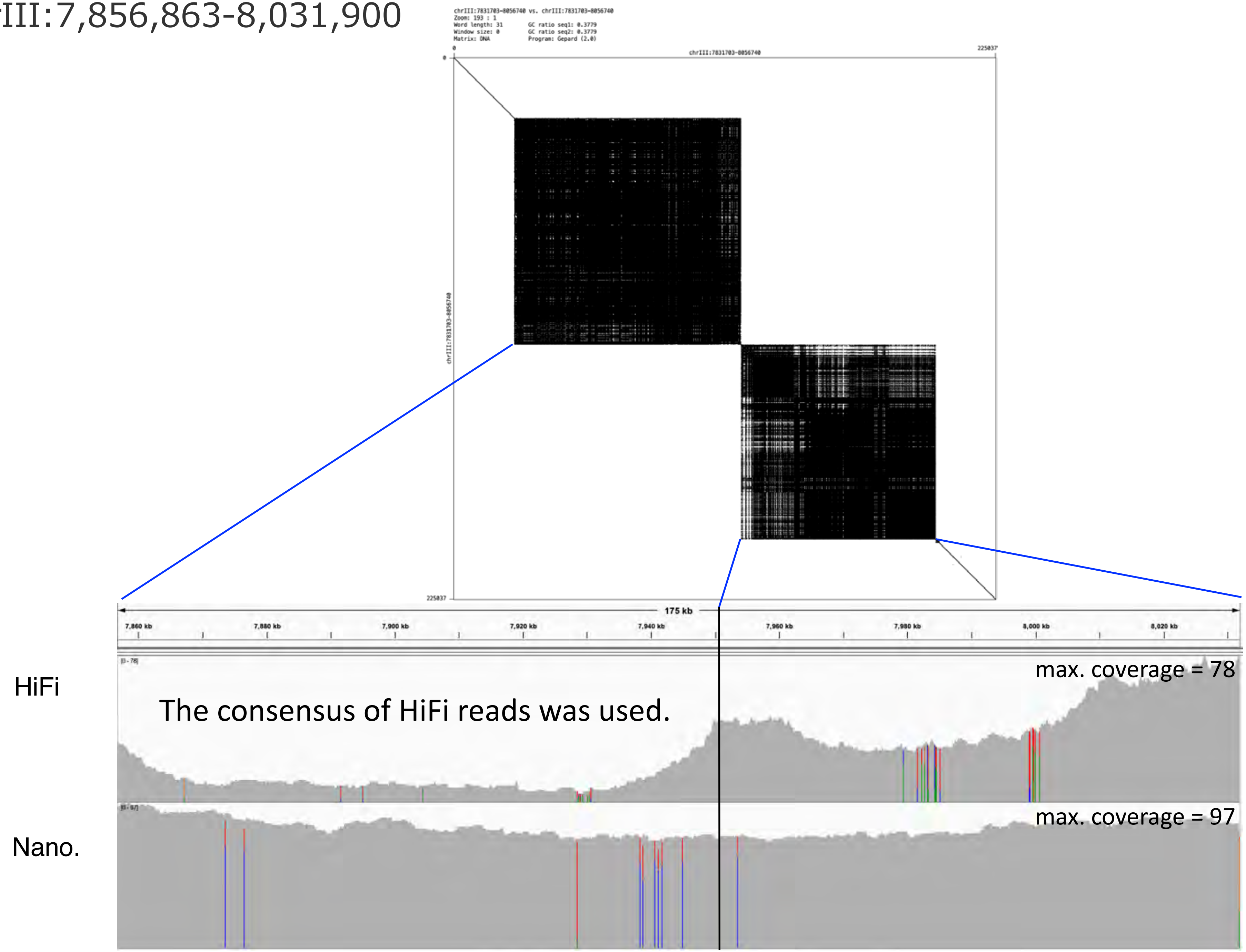

### chrIII:8,174,772-8,214,520

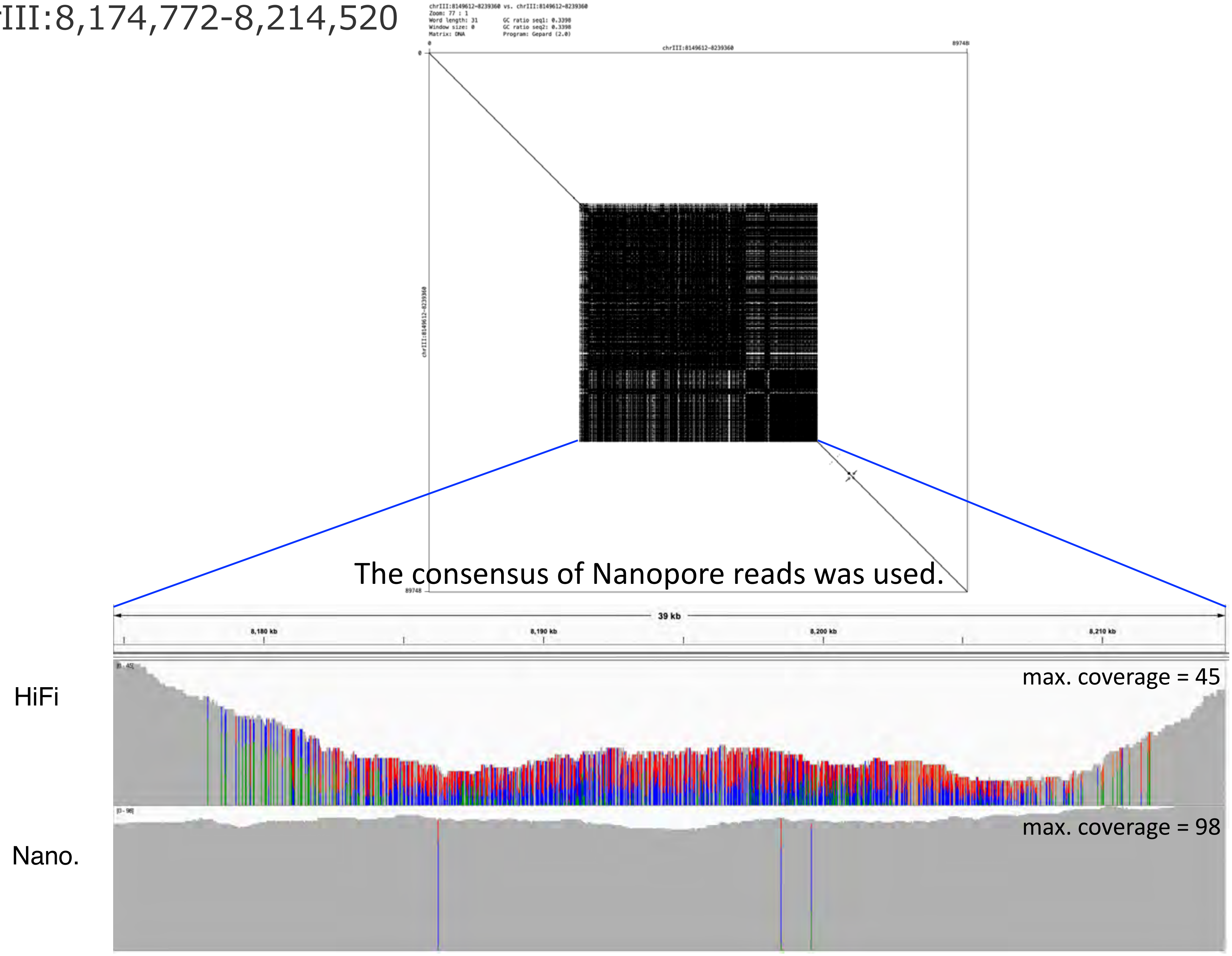

### chrIII:9,497,286-9,512,310

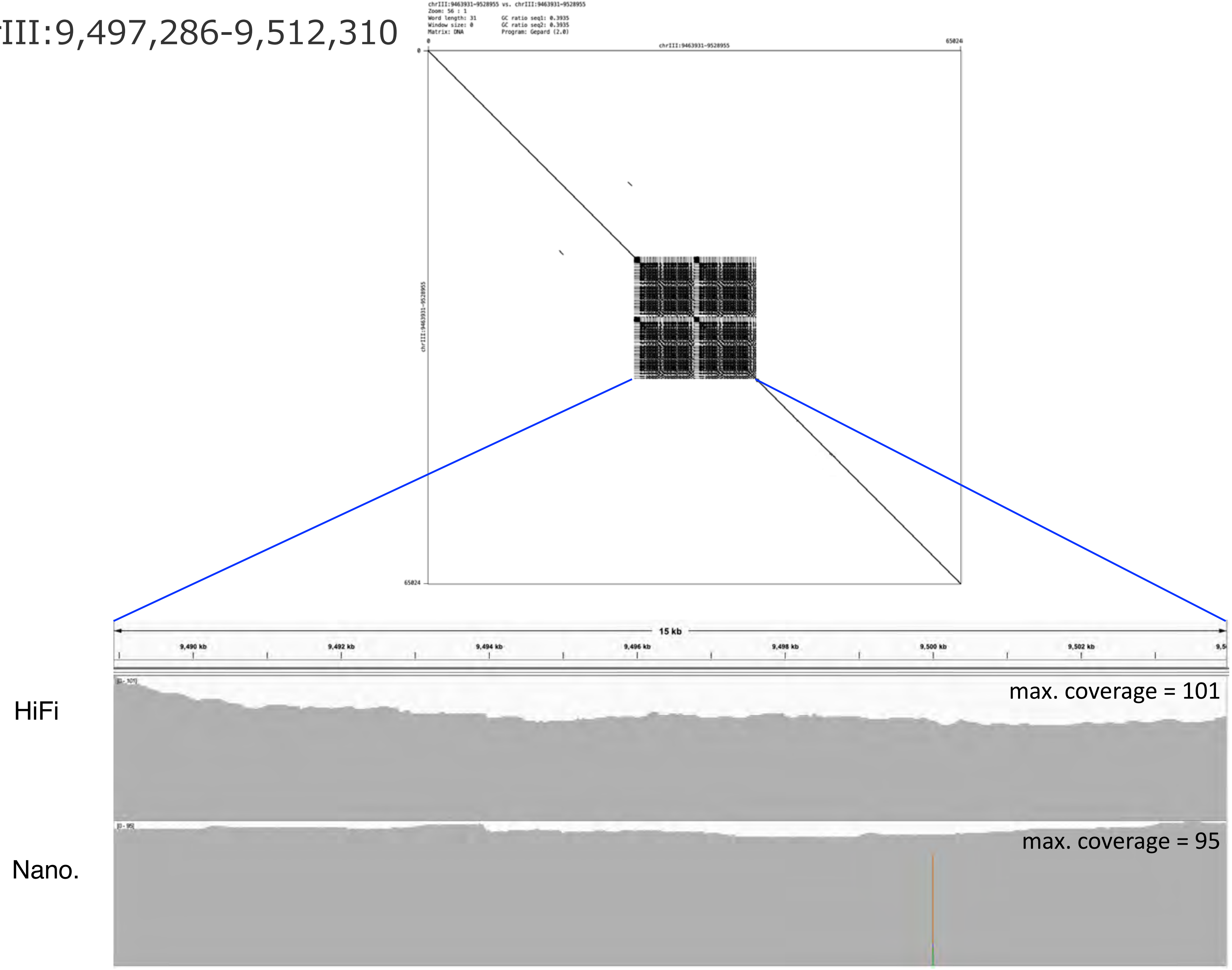

chrIII:10,873,648-10,972,710

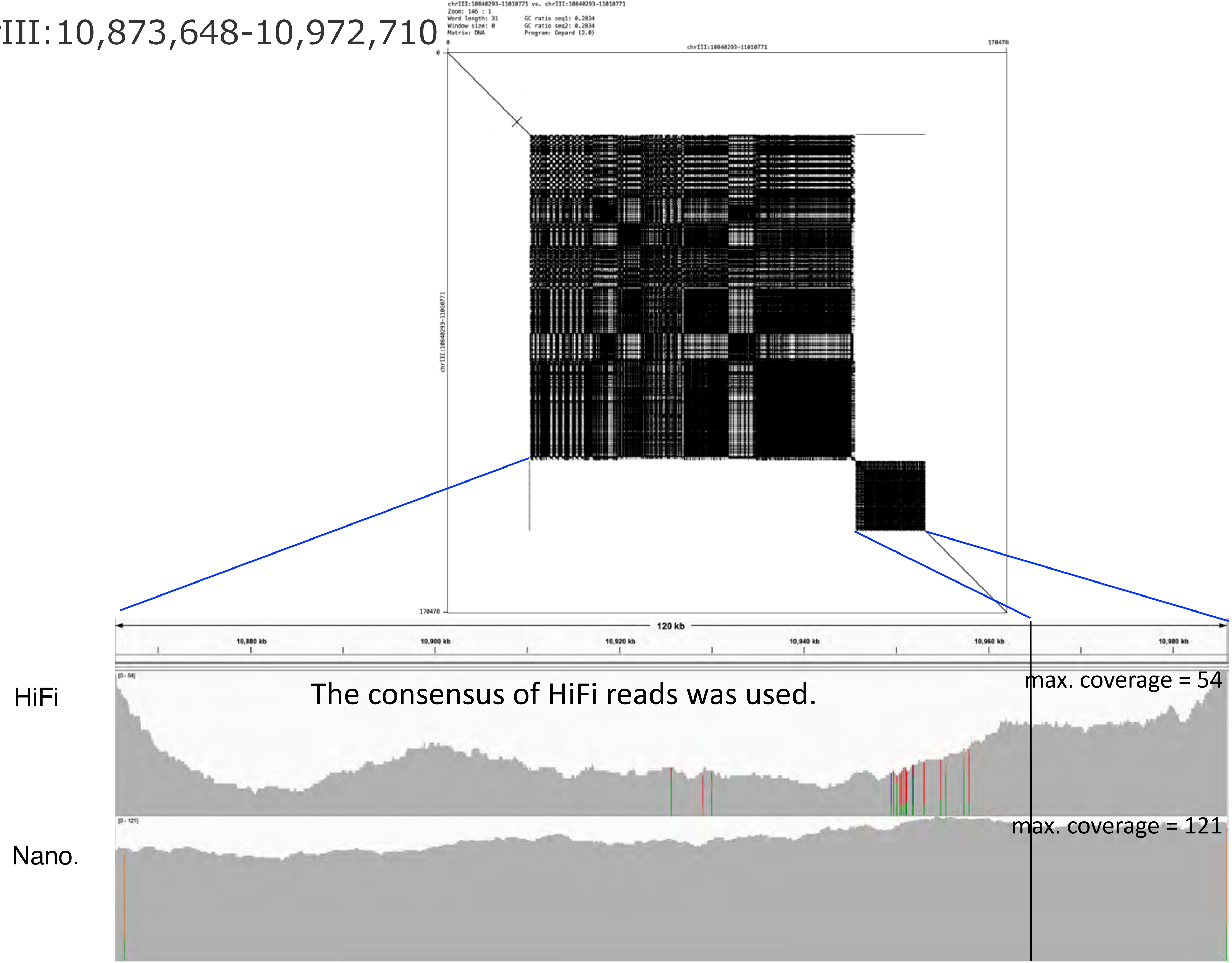

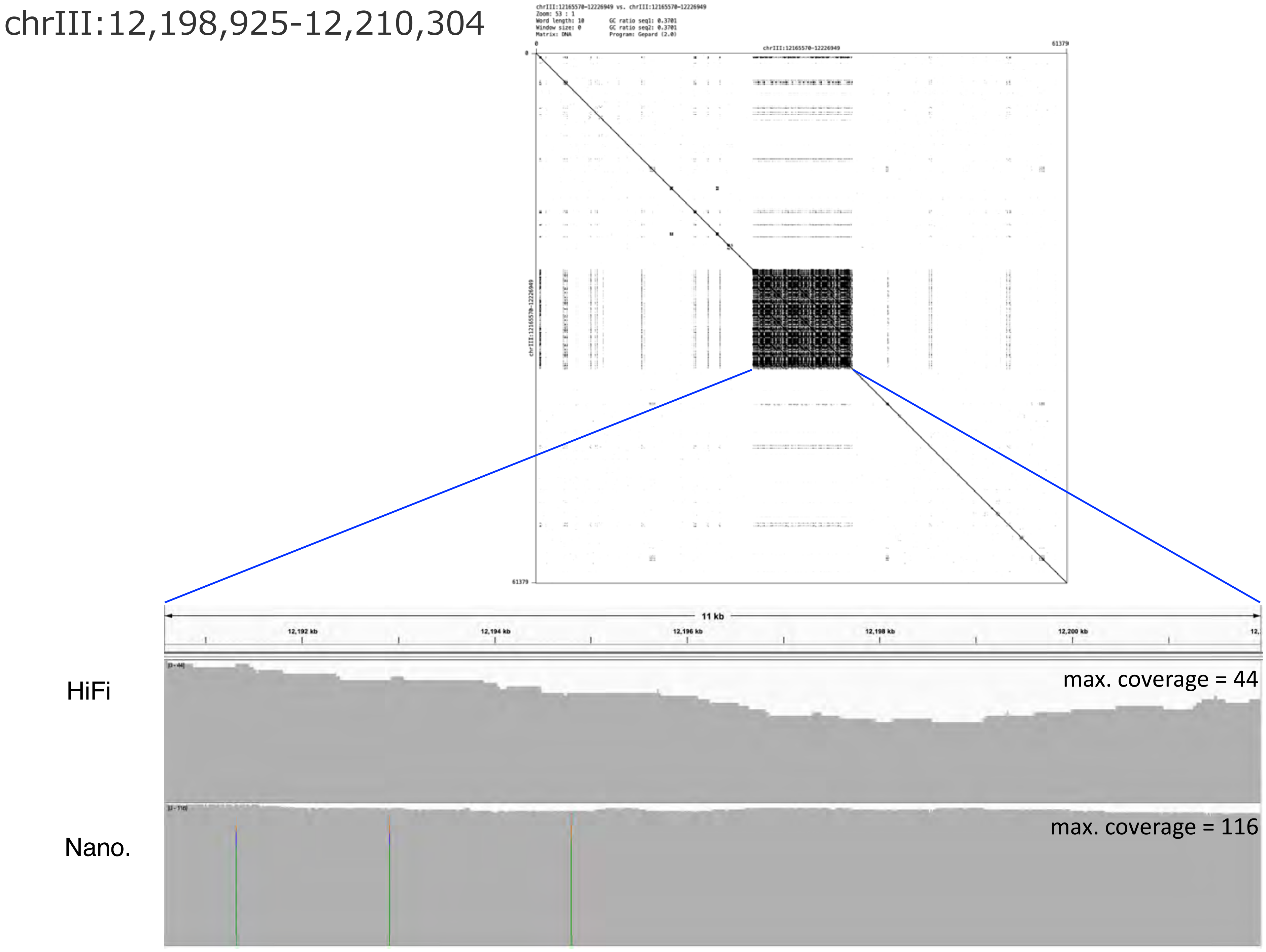
chrIII:12,198,925-12,210,304
chrIII:12165570-12226949 vs. chrIII:12165570-12226949
Zoom: 53 : 1
Word length: 10
Window size: 0
Matrix: DNA
GC ratio seq1: 0.3701
GC ratio seq2: 0.3701
Program: Gepard (2.0)
0
chrIII:12165570-12226949
61379
0
chrIII:12165570-12226949
61379
11 kb
12,192 kb
12,194 kb
12,196 kb
12,198 kb
12,200 kb
HiFi
max. coverage = 44
Nano.
max. coverage = 116

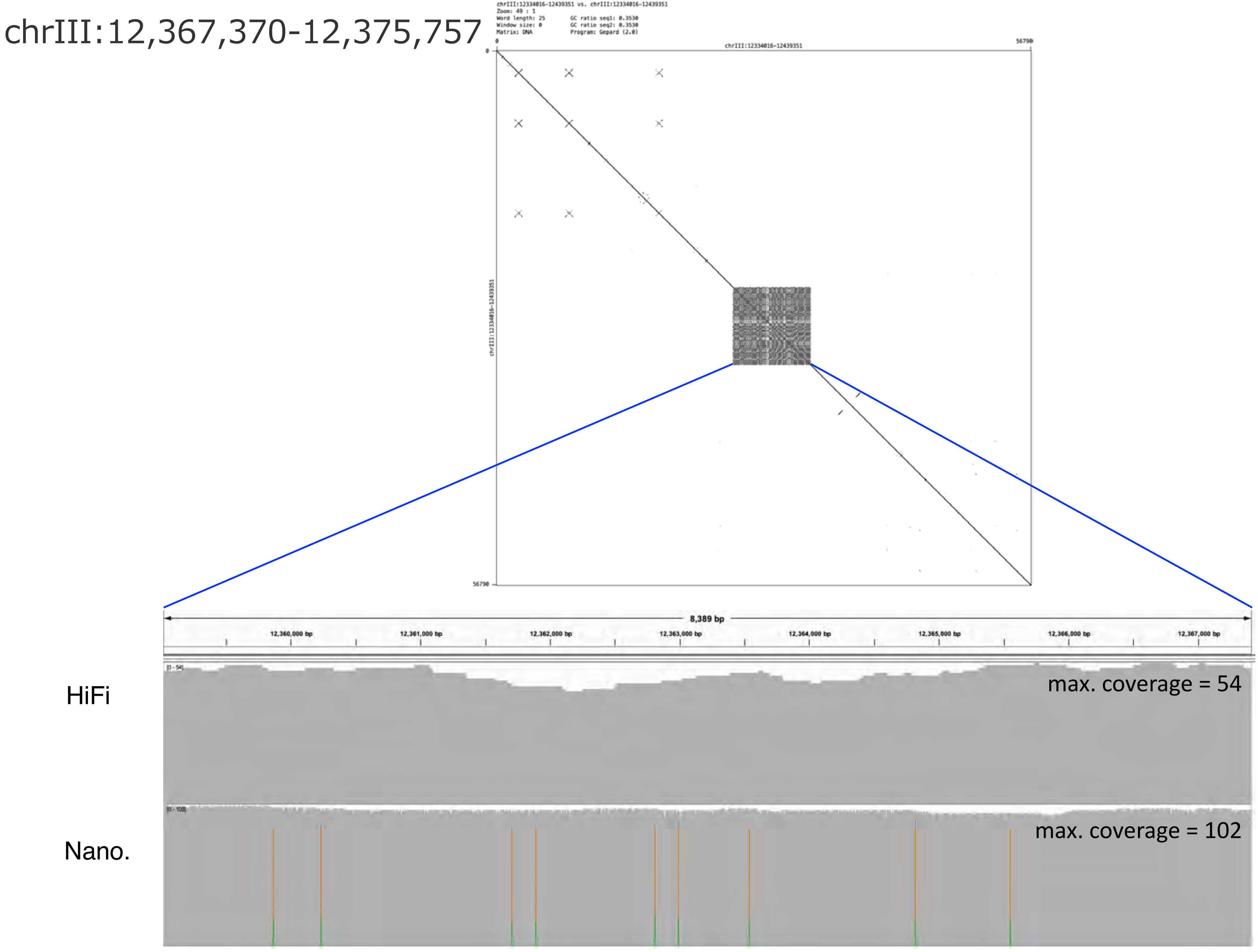
chrIII:12,367,370-12,375,757
chrIII:12334016-12439351 vs. chrIII:12334016-12439351
Zoom: 49 : 1
Word length: 25
Window size: 0
Matrix: DNA
GC ratio seq1: 0.3530
GC ratio seq2: 0.3530
Program: Gepard (2.0)
chrIII:12334016-12439351
0
56790
8,389 bp
12,360,000 bp
12,361,000 bp
12,362,000 bp
12,363,000 bp
12,364,000 bp
12,365,000 bp
12,366,000 bp
12,367,000 bp
HiFi
max. coverage = 54
Nano.
max. coverage = 102

### chrIII:12,415,497-12,422,705

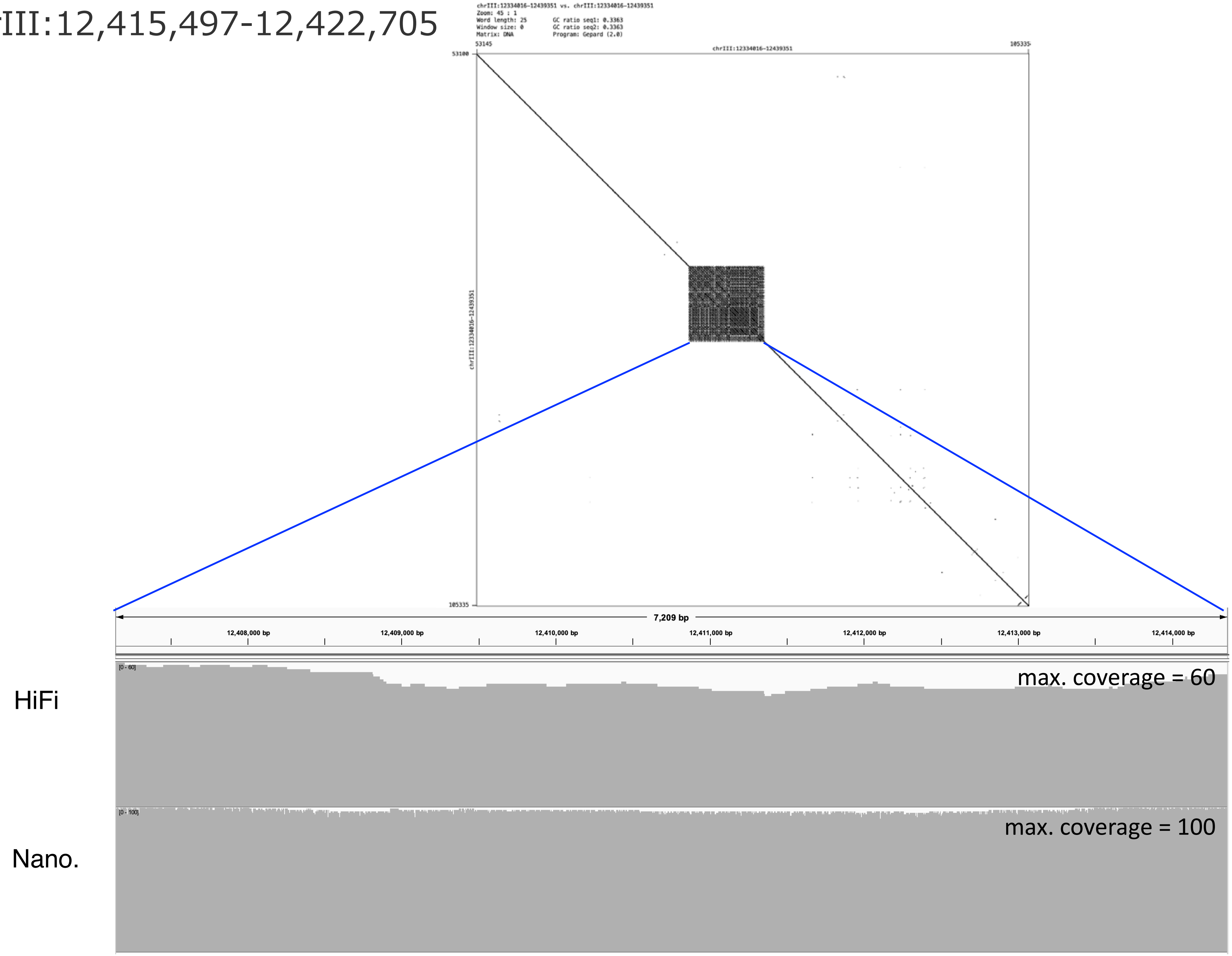

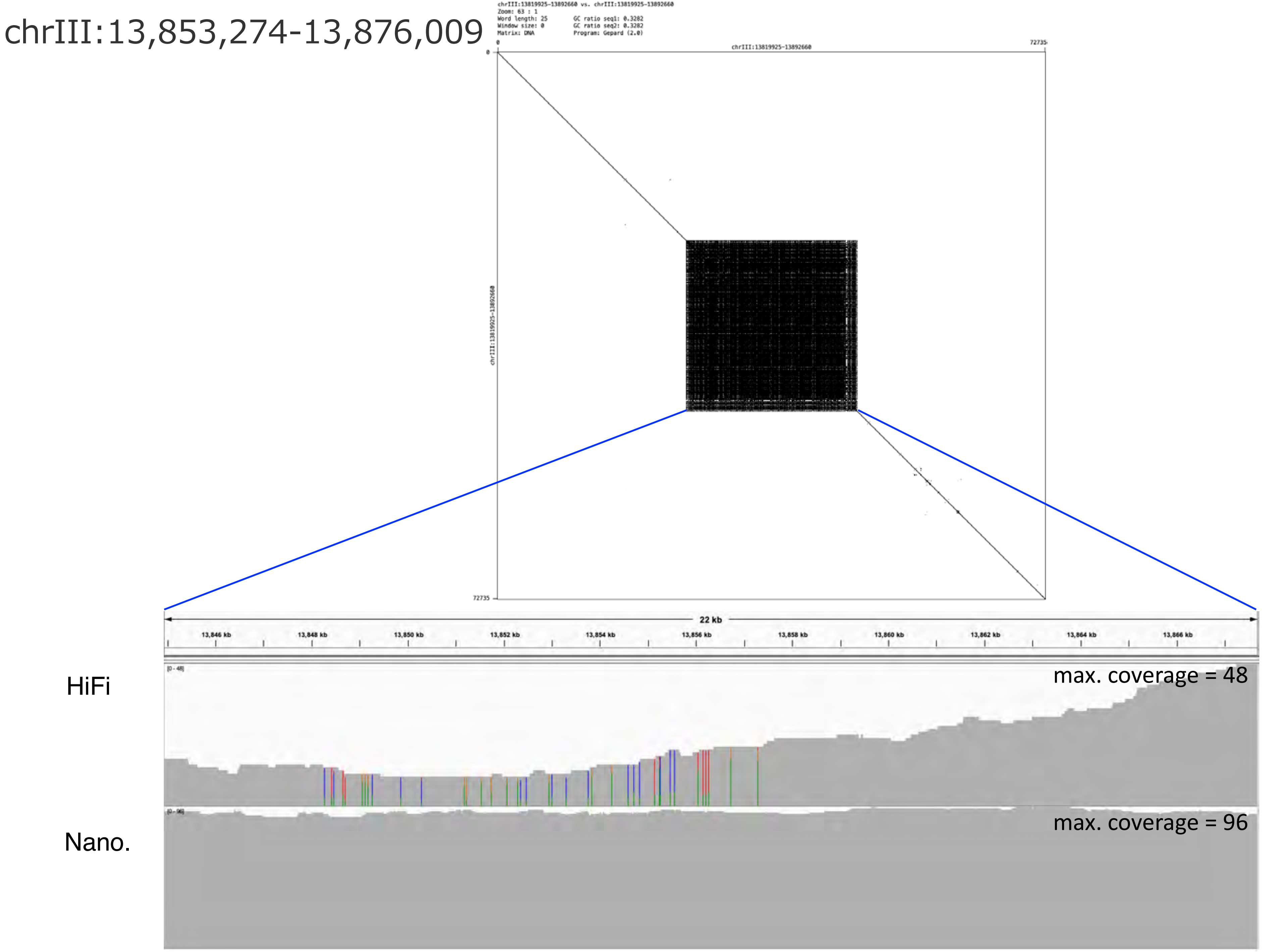
chrIII:13,853,274-13,876,009
chrIII:13819925-13892660 vs. chrIII:13819925-13892660
Zoom: 63 : 1
Word length: 25
Window size: 0
Matrix: DNA
GC ratio seq1: 0.3282
GC ratio seq2: 0.3282
Program: Gepard (2.0)
chrIII:13819925-13892660
0
72735
chrIII:13819925-13892660
22 kb
13,846 kb
13,848 kb
13,850 kb
13,852 kb
13,854 kb
13,856 kb
13,858 kb
13,860 kb
13,862 kb
13,864 kb
13,866 kb
HiFi
max. coverage = 48
Nano.
max. coverage = 96

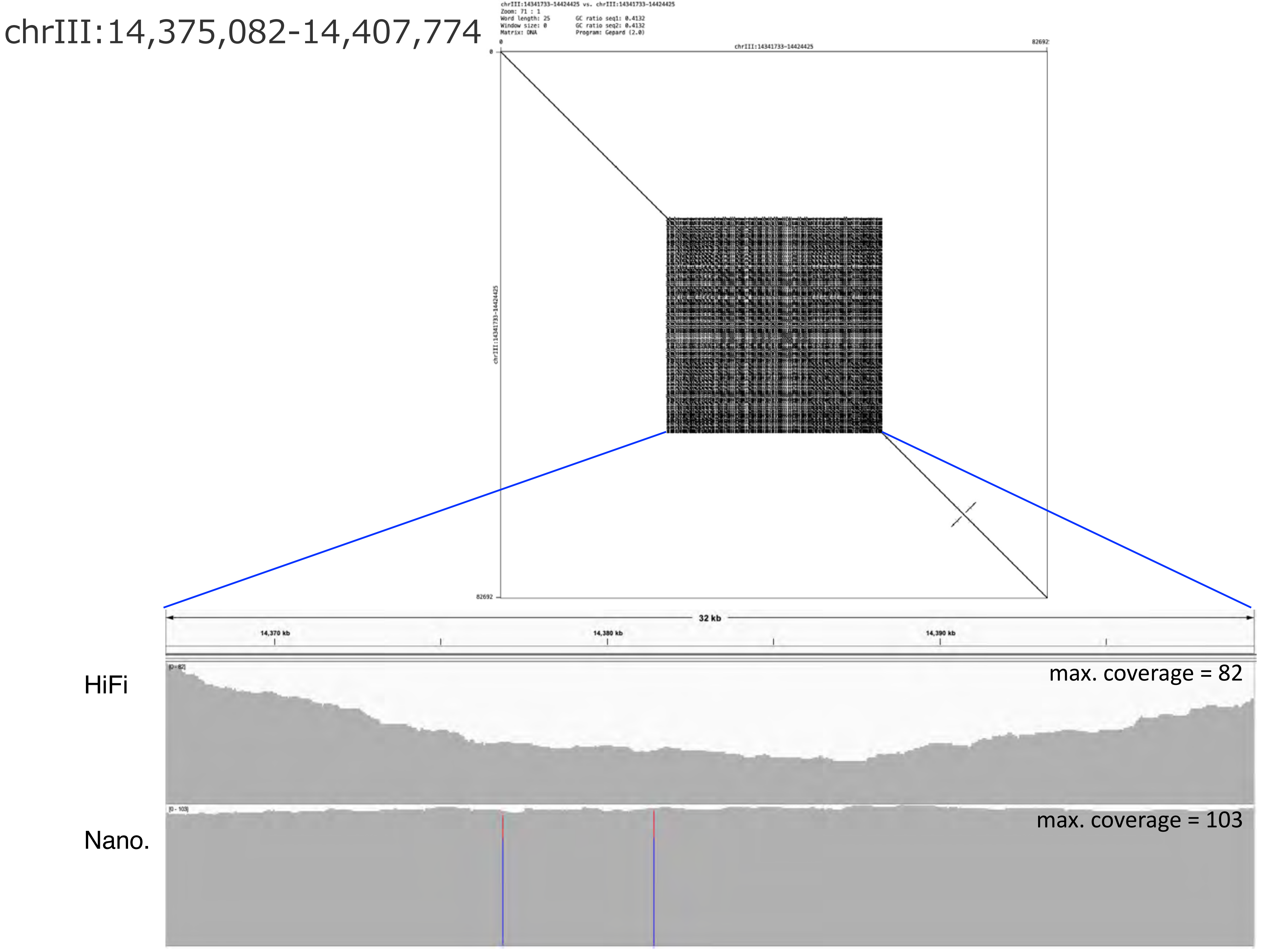
chrIII:14,375,082-14,407,774
chrIII:14341733-14424425 vs. chrIII:14341733-14424425
Word length: 25
Matrix: DNA
Program: Gepard (2.0)
chrIII:14341733-14424425
0
82692
32 kb
14,370 kb
14,380 kb
14,390 kb
HiFi
max. coverage = 82
Nano.
max. coverage = 103

chrIV:696,328-706,244

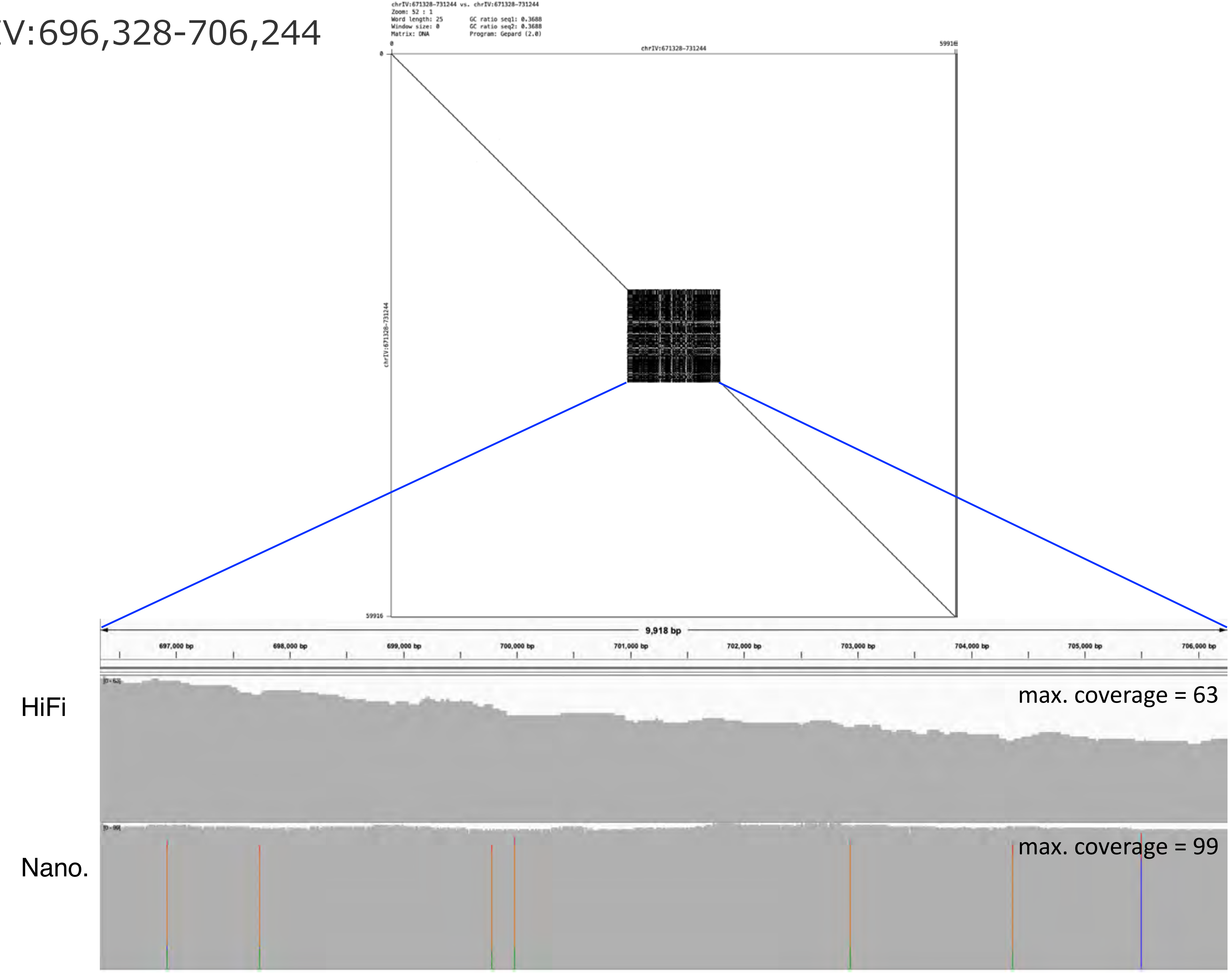

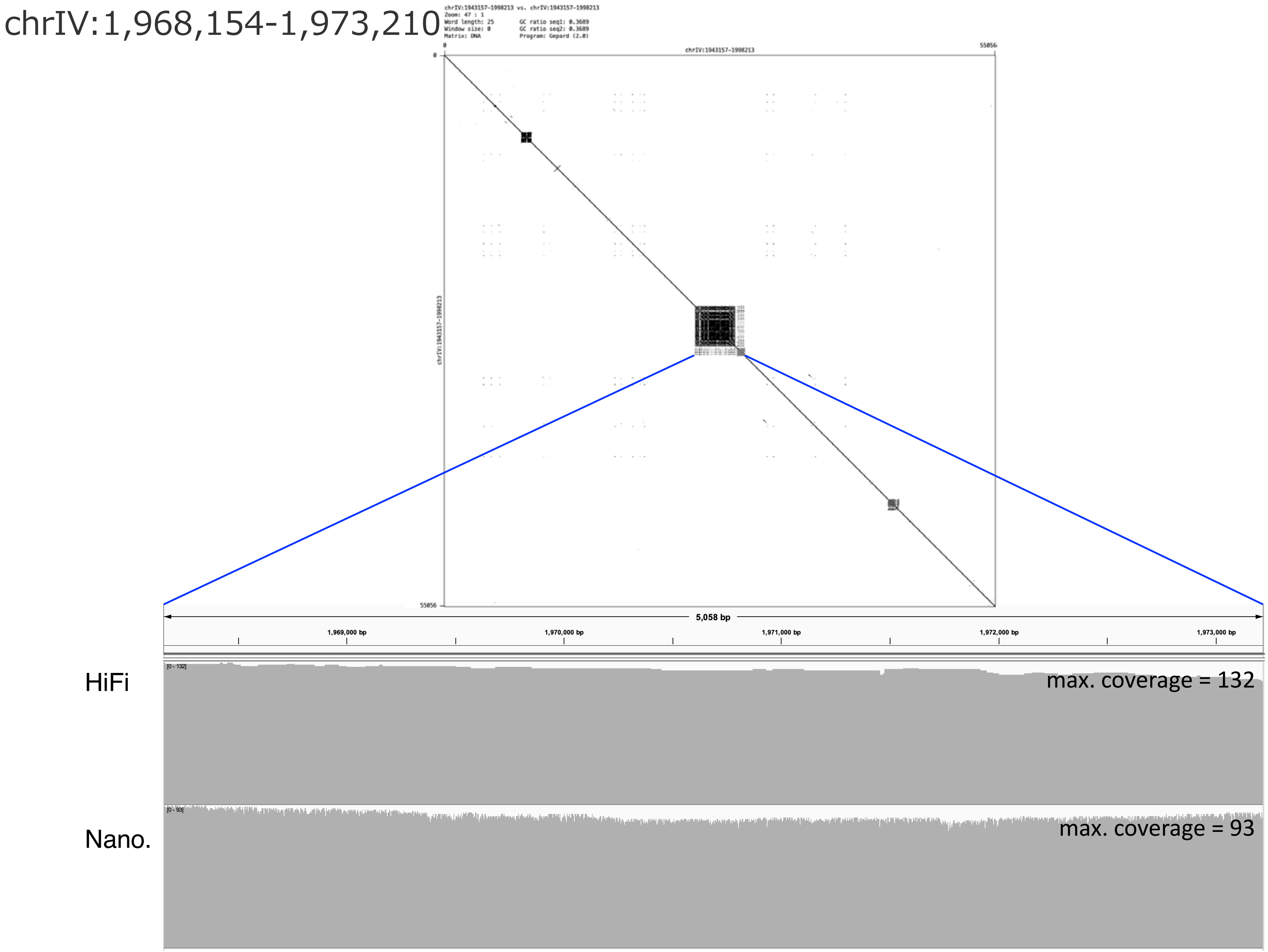
chrIV:1,968,154-1,973,210
chrIV:1943157-1998213 vs. chrIV:1943157-1998213
Zoom: 47 : 1
Word length: 25
Window size: 0
Matrix: DNA
GC ratio seq1: 0.3689
GC ratio seq2: 0.3689
Program: Gepard (2.0)
chrIV:1943157-1998213
0
55056
0
55056
5,058 bp
1,969,000 bp
1,970,000 bp
1,971,000 bp
1,972,000 bp
1,973,000 bp
HiFi
[0 - 132]
max. coverage = 132
Nano.
[0 - 93]
max. coverage = 93

### chrIV:2,857,216-2,919,705

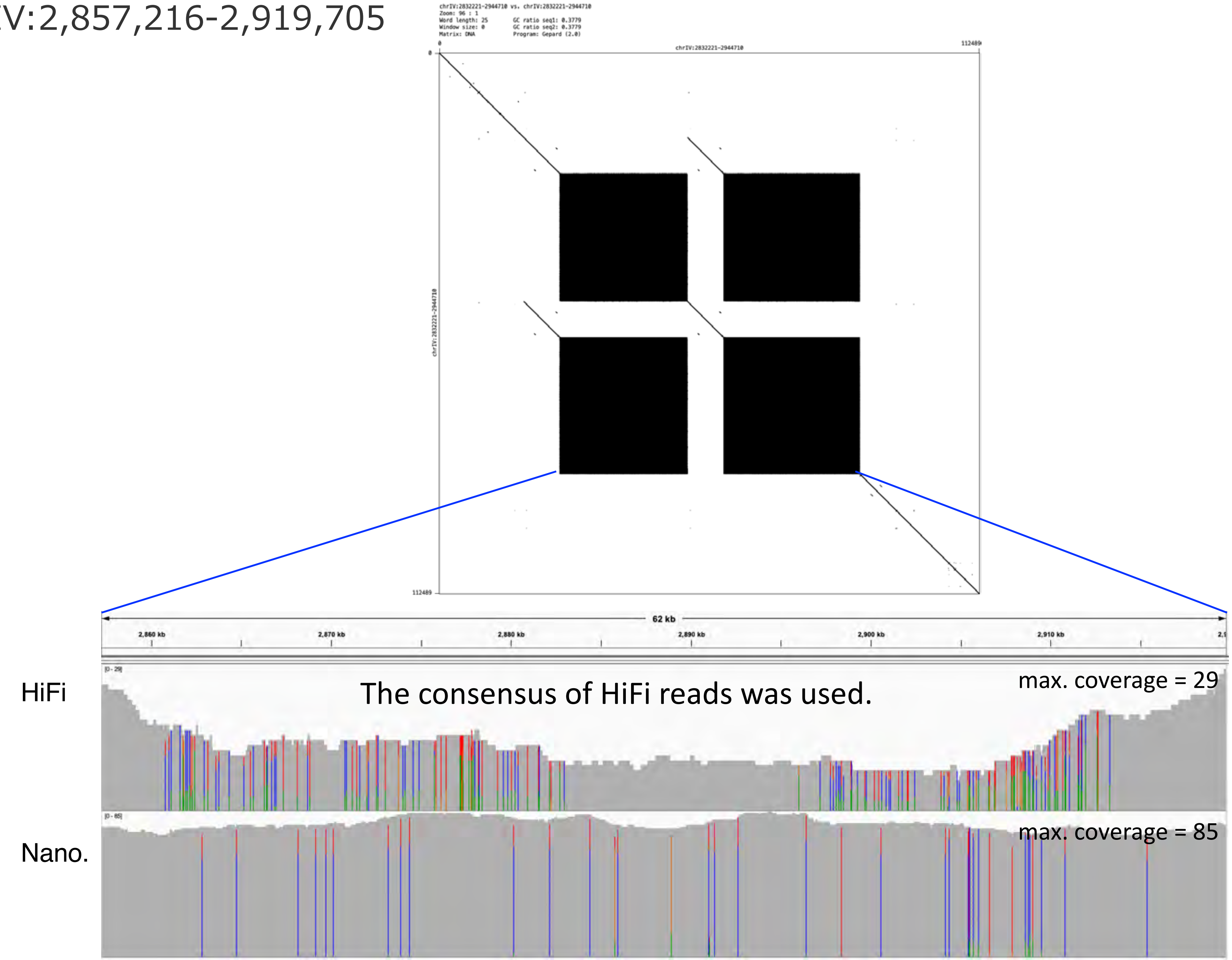

chrIV:3,295,670-3,323,947

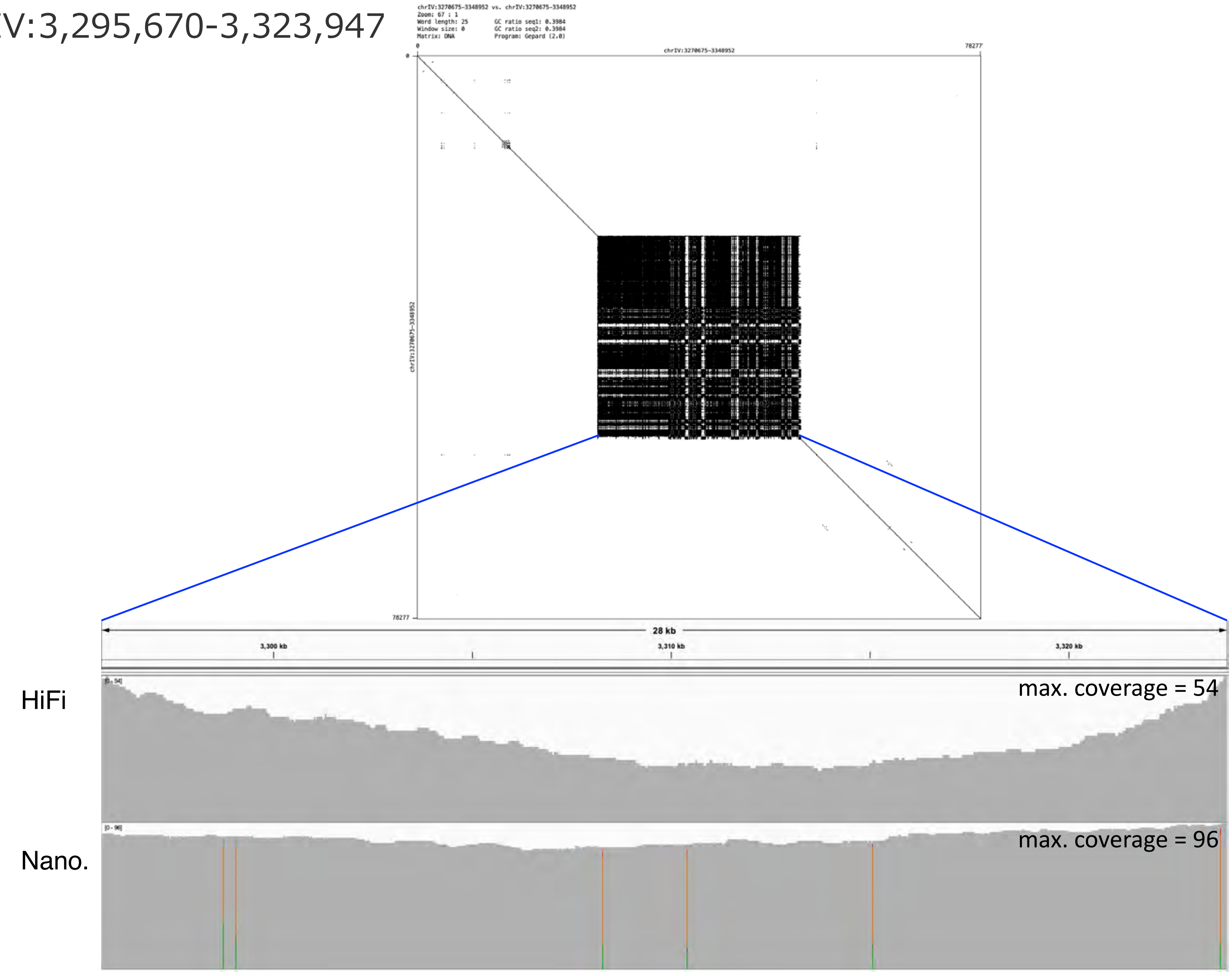

### chrIV:4,531,638-4,564,245

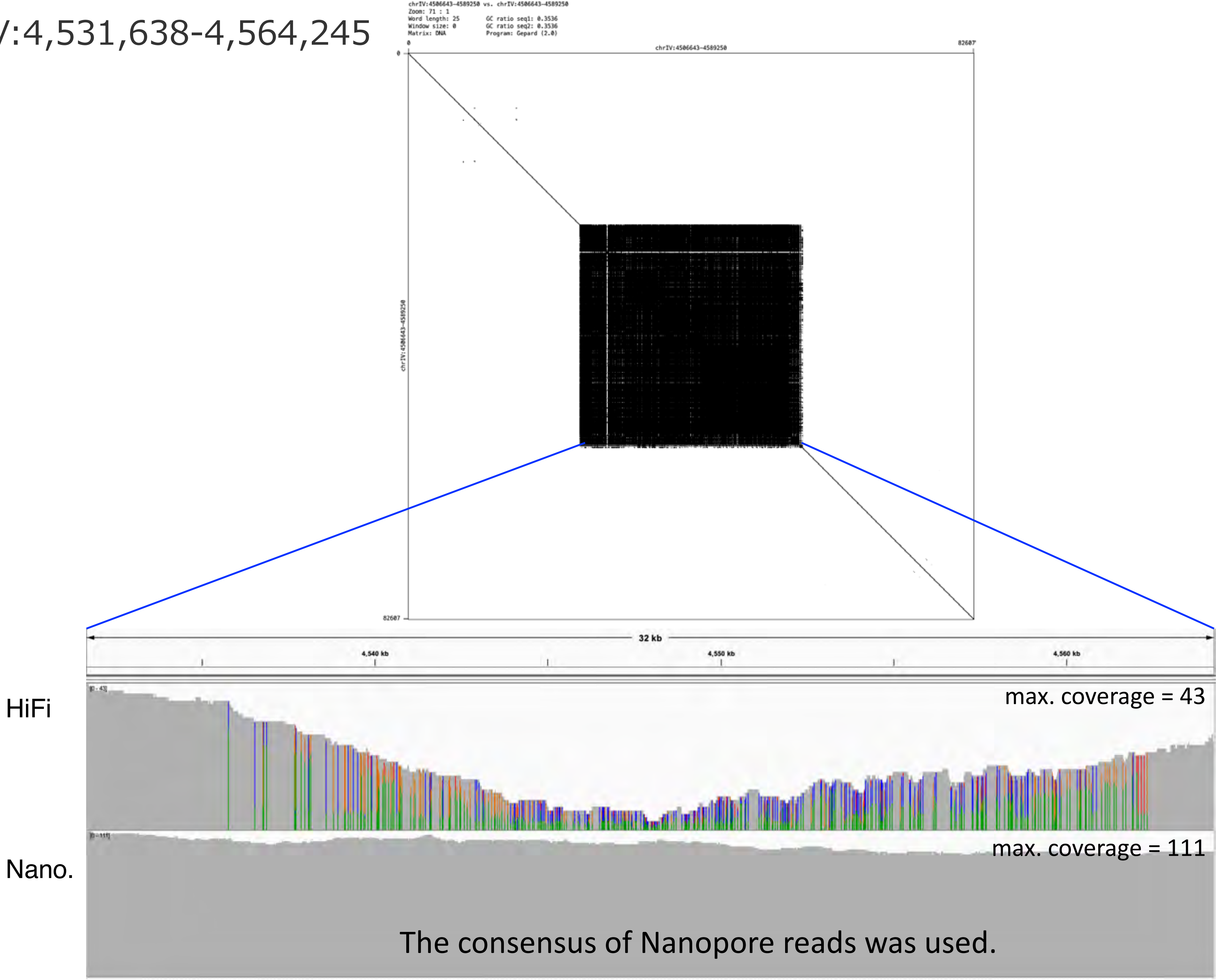


The consensus of Nanopore reads was used.

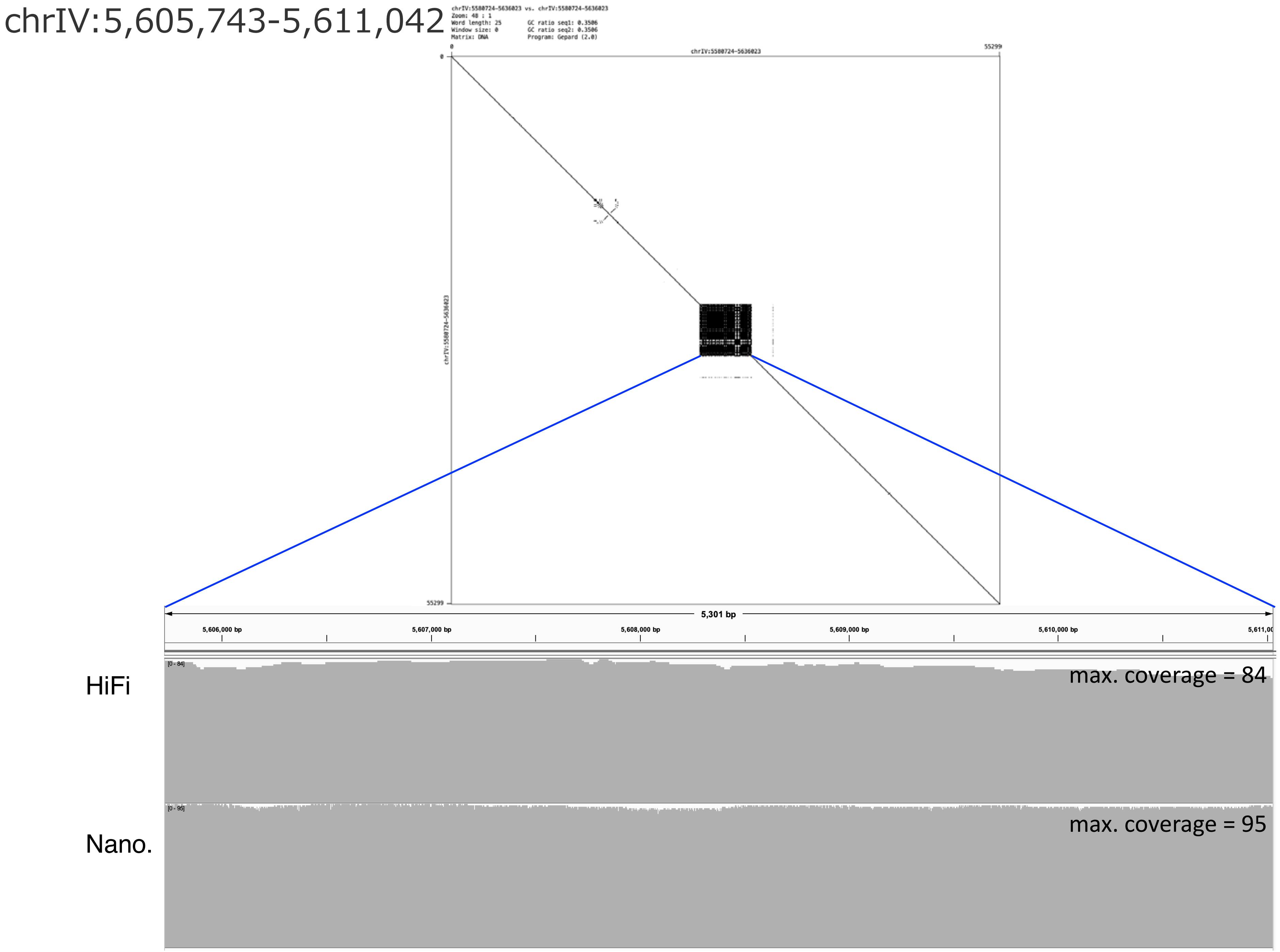
chrIV:5,605,743-5,611,042
chrIV:5580724-5636023 vs. chrIV:5580724-5636023
Zoom: 48 : 1
Word length: 25
Window size: 0
Matrix: DNA
GC ratio seq1: 0.3506
GC ratio seq2: 0.3506
Program: Gepard (2.0)
chrIV:5580724-5636023
55299
5,301 bp
5,606,000 bp
5,607,000 bp
5,608,000 bp
5,609,000 bp
5,610,000 bp
HiFi
max. coverage = 84
Nano.
max. coverage = 95

### chrIV:6,501,916-6,516,317

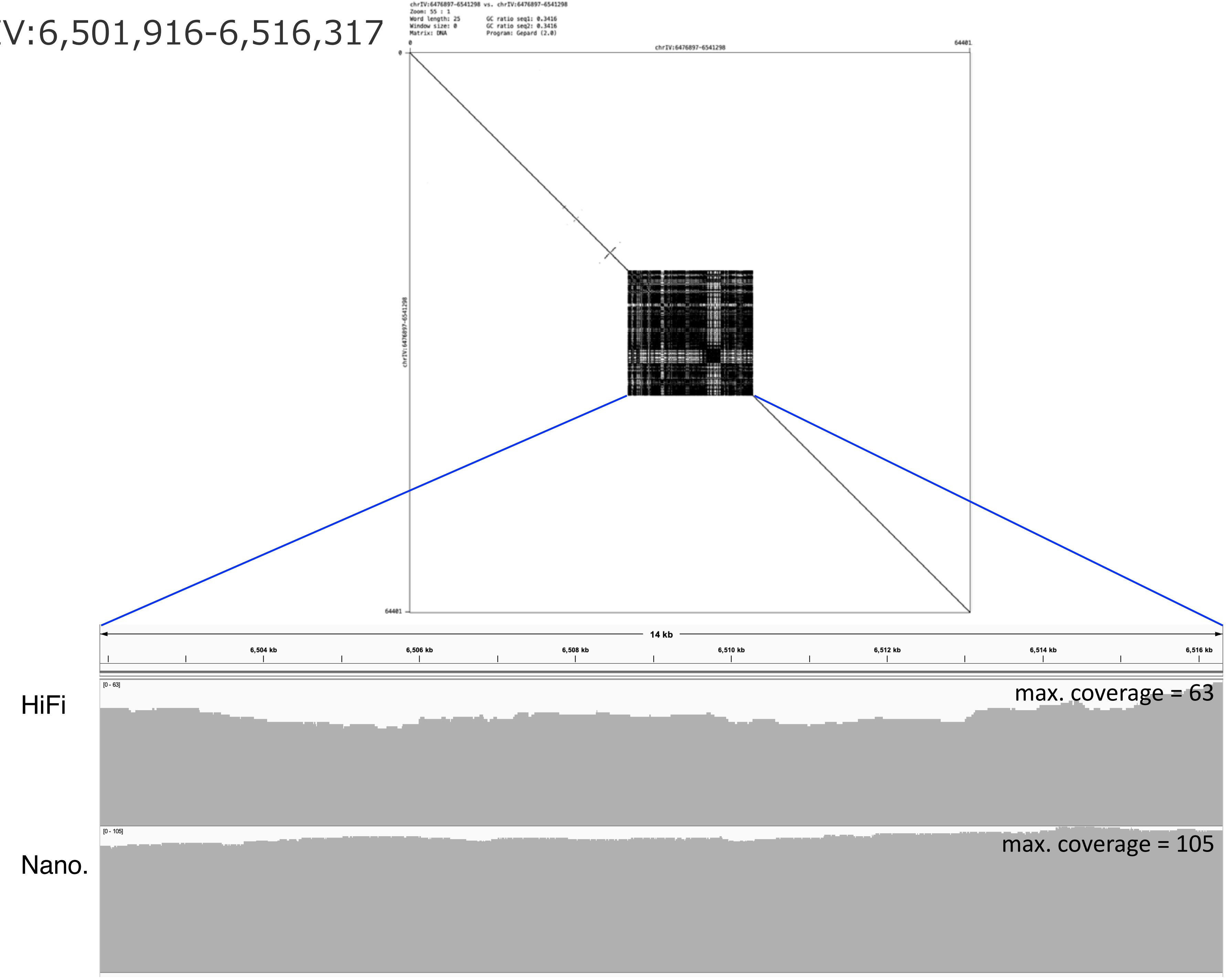

chrIV:6,824,094-6,928,586

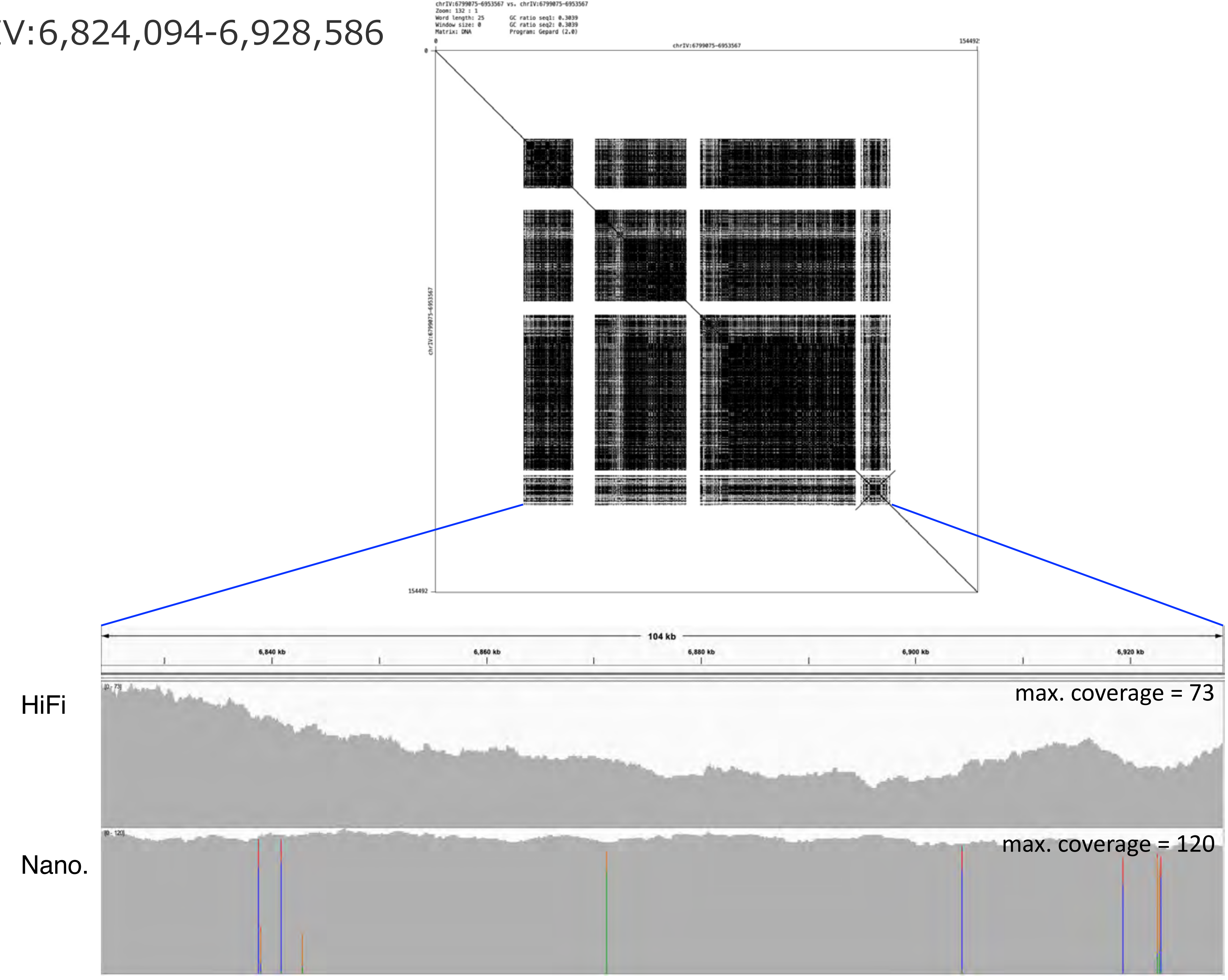

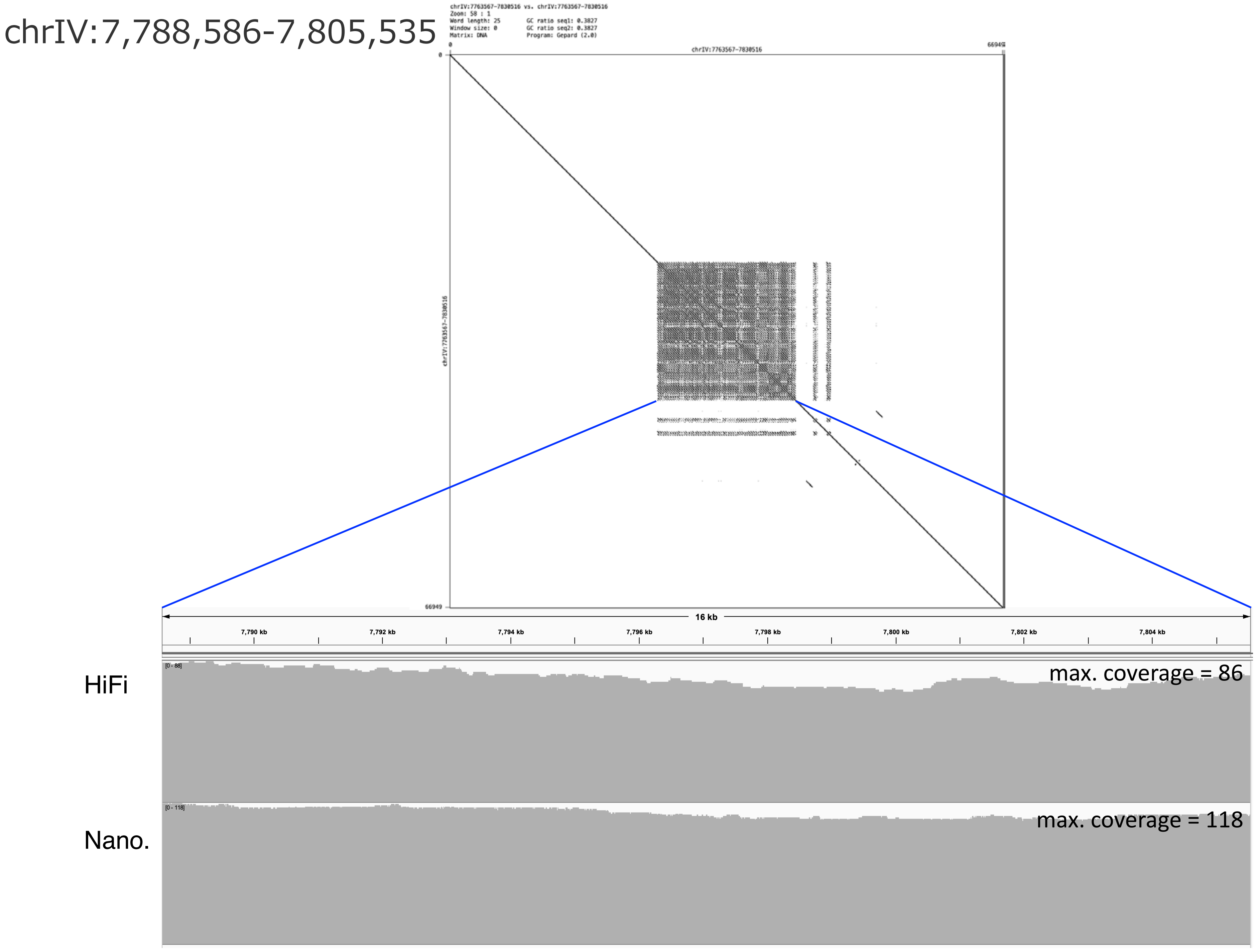
chrIV:7,788,586-7,805,535
chrIV:7763567-7830516 vs. chrIV:7763567-7830516
Zoom: 58 : 1
Word length: 25
Window size: 0
Matrix: DNA
GC ratio seq1: 0.3827
GC ratio seq2: 0.3827
Program: Gepard (2.0)
chrIV:7763567-7830516
0
66949
16 kb
7,790 kb
7,792 kb
7,794 kb
7,796 kb
7,798 kb
7,800 kb
7,802 kb
7,804 kb
HiFi
max. coverage = 86
Nano.
max. coverage = 118

chrIV:8,780,652-8,862,273

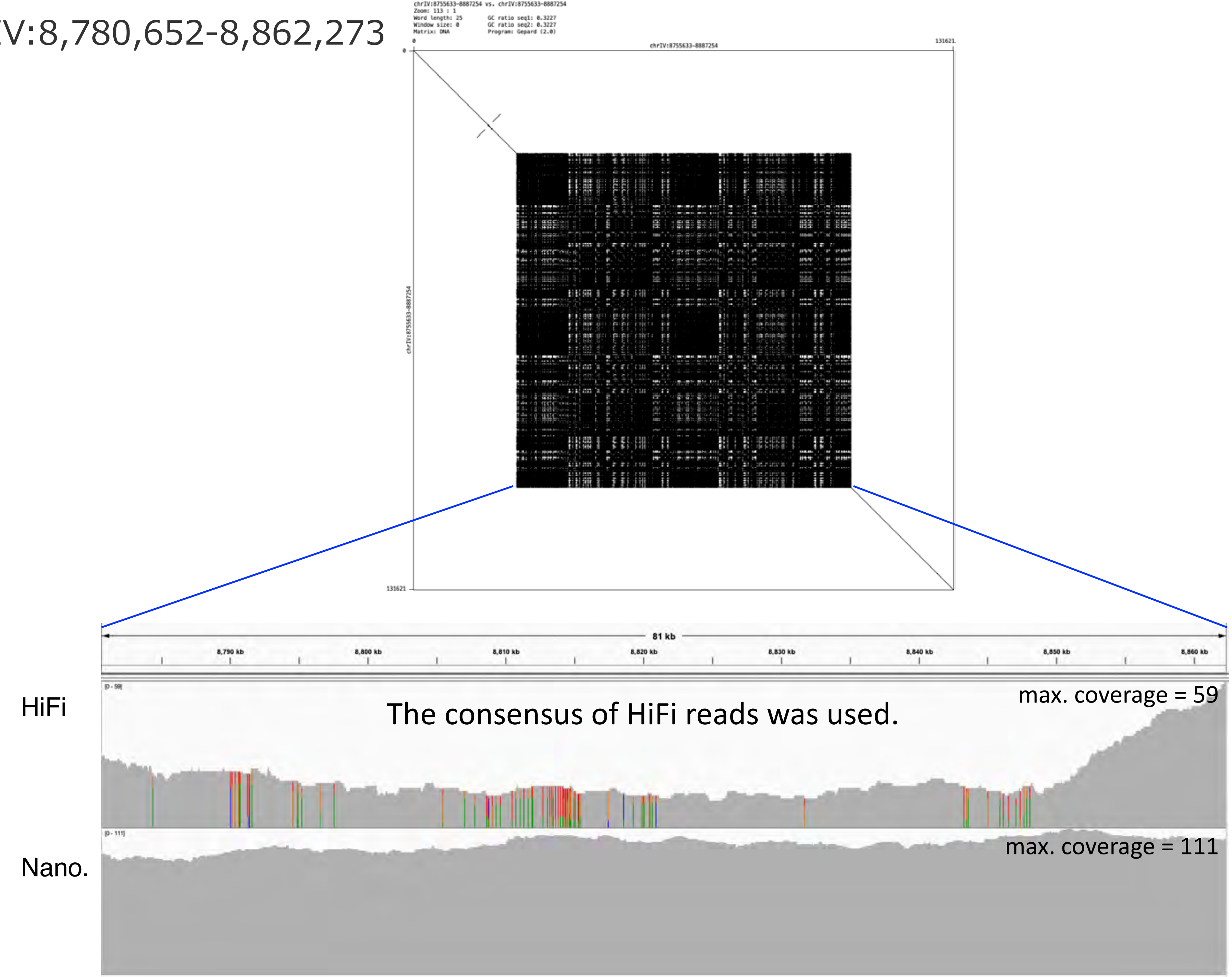


The consensus of HiFi reads was used.

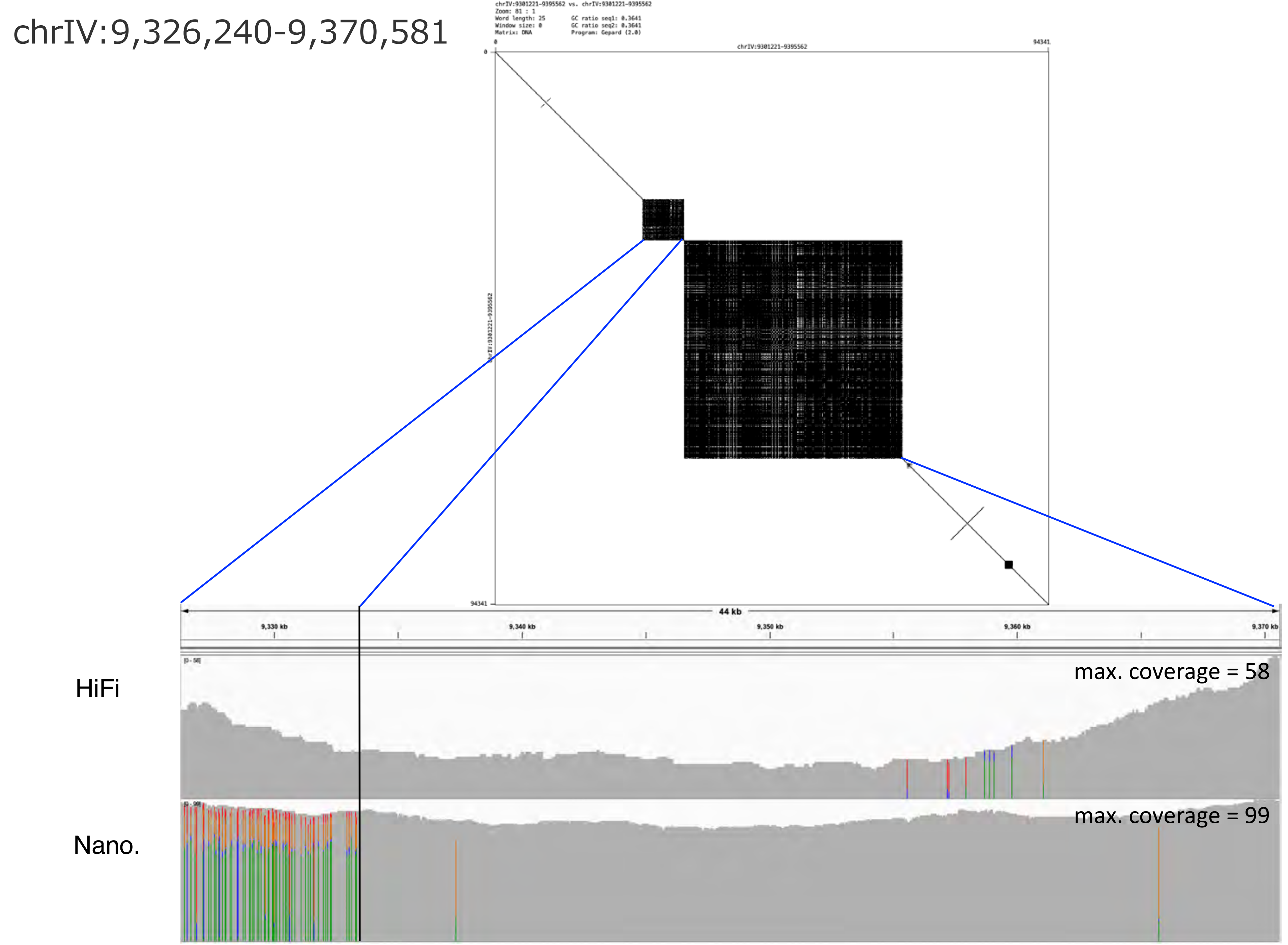
chrIV:9,326,240-9,370,581
chrIV:9301221-9395562 vs. chrIV:9301221-9395562
Zoom: 81 : 1
Word length: 25
Window size: 0
Matrix: DNA
GC ratio seq1: 0.3641
GC ratio seq2: 0.3641
Program: Gepard (2.0)
chrIV:9301221-9395562
0
94341
44 kb
9,330 kb
9,340 kb
9,350 kb
9,360 kb
9,370 kb
HiFi
max. coverage = 58
Nano.
max. coverage = 99

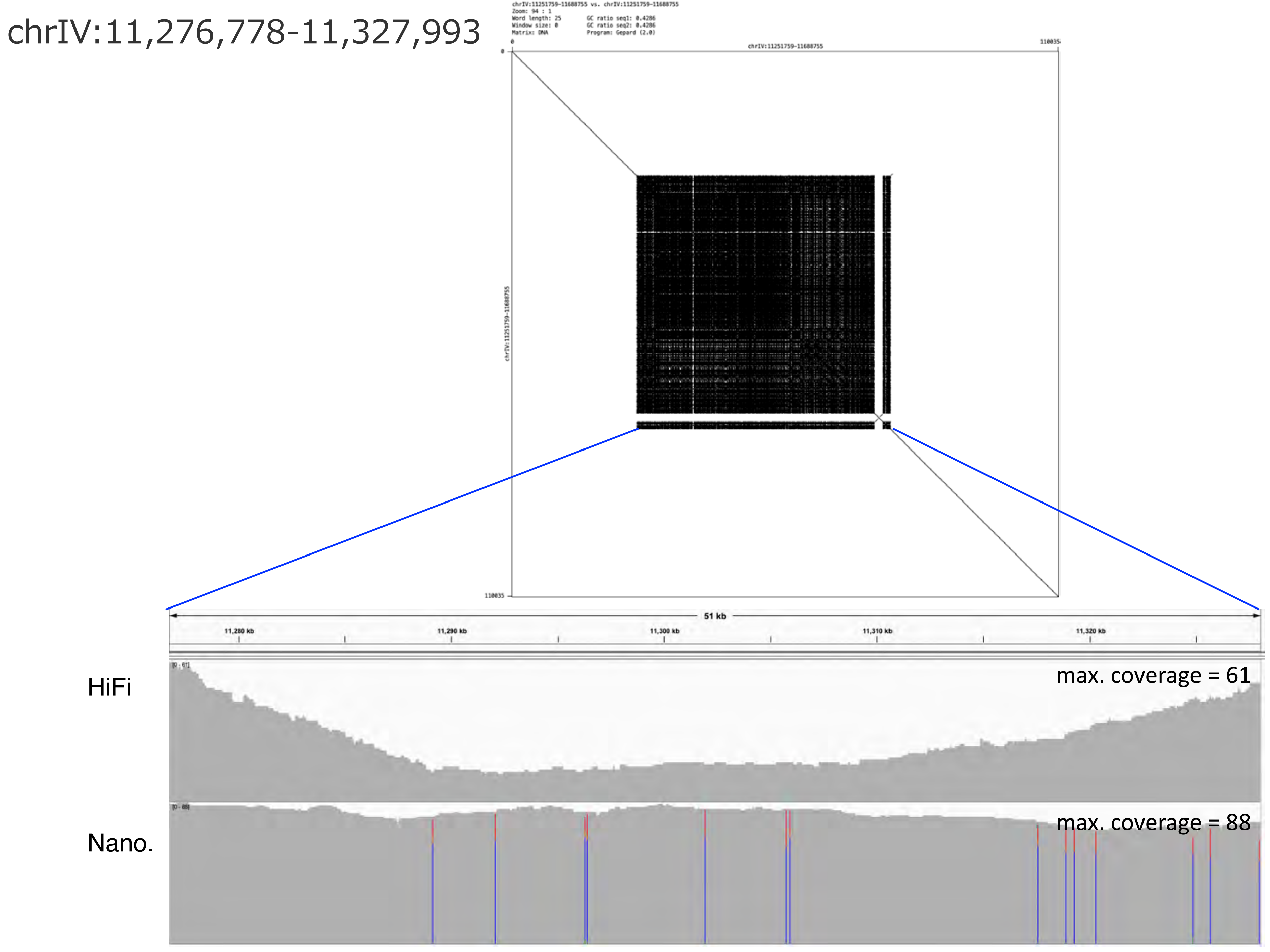
chrIV:11,276,778-11,327,993
chrIV:11251759-11688755 vs. chrIV:11251759-11688755
Zoom: 94 : 1
Word length: 25
Window size: 0
Matrix: DNA
GC ratio seq1: 0.4286
GC ratio seq2: 0.4286
Program: Gepard (2.0)
chrIV:11251759-11688755
0
110035
51 kb
11,280 kb
11,290 kb
11,300 kb
11,310 kb
11,320 kb
HiFi
max. coverage = 61
Nano.
max. coverage = 88

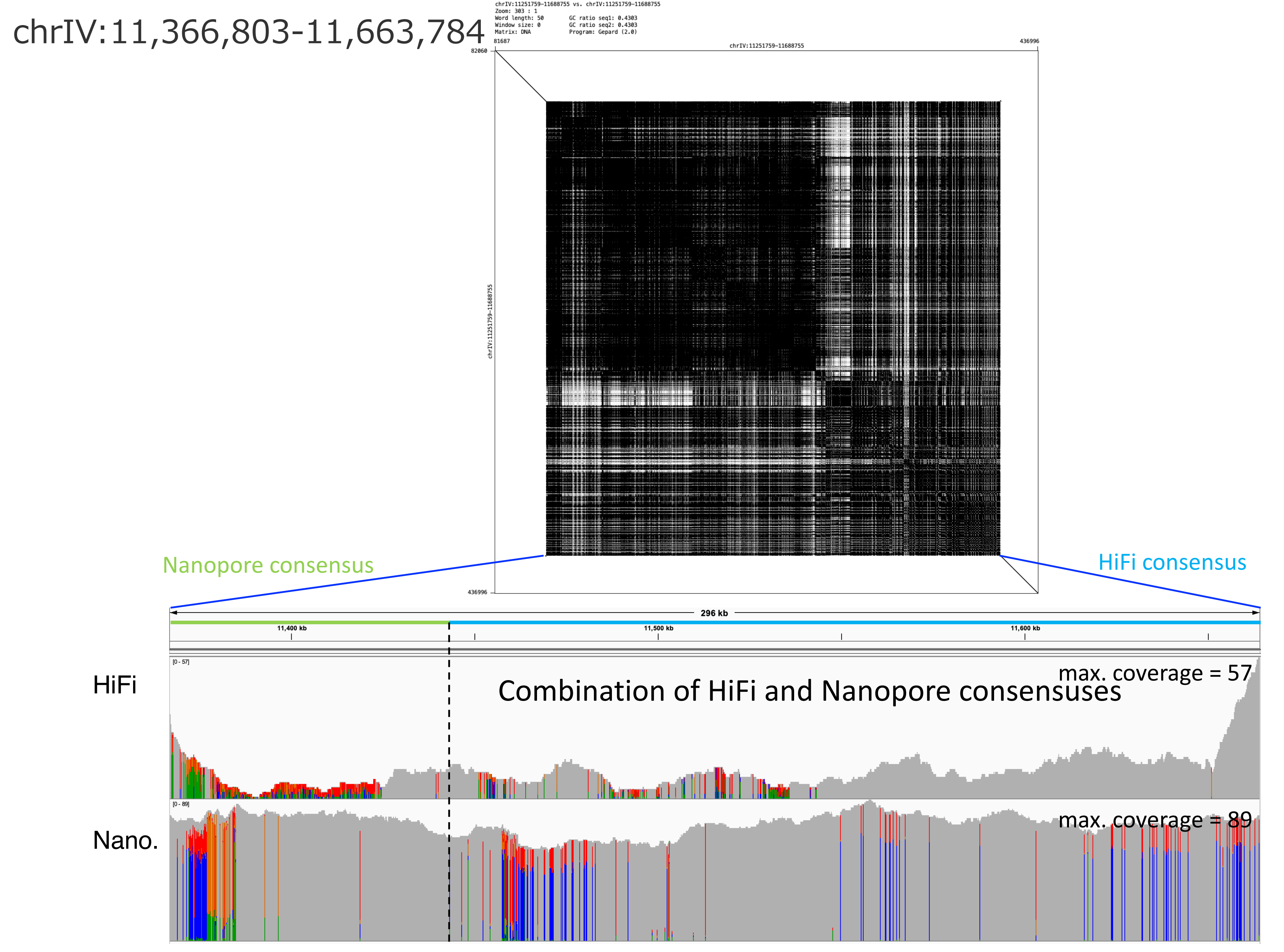
chrIV:11,366,803-11,663,784
chrIV:11251759-11688755 vs. chrIV:11251759-11688755
Zoom: 303 : 1
Word length: 50
Window size: 0
Matrix: DNA
GC ratio seq1: 0.4303
GC ratio seq2: 0.4303
Program: Gepard (2.0)
81687
chrIV:11251759-11688755
436996
82060
chrIV:11251759-11688755
436996
Nanopore consensus
HiFi consensus
296 kb
11,400 kb
11,500 kb
11,600 kb
HiFi
[0 - 57]
max. coverage = 57
Combination of HiFi and Nanopore consensuses
Nano.
[0 - 89]
max. coverage = 89

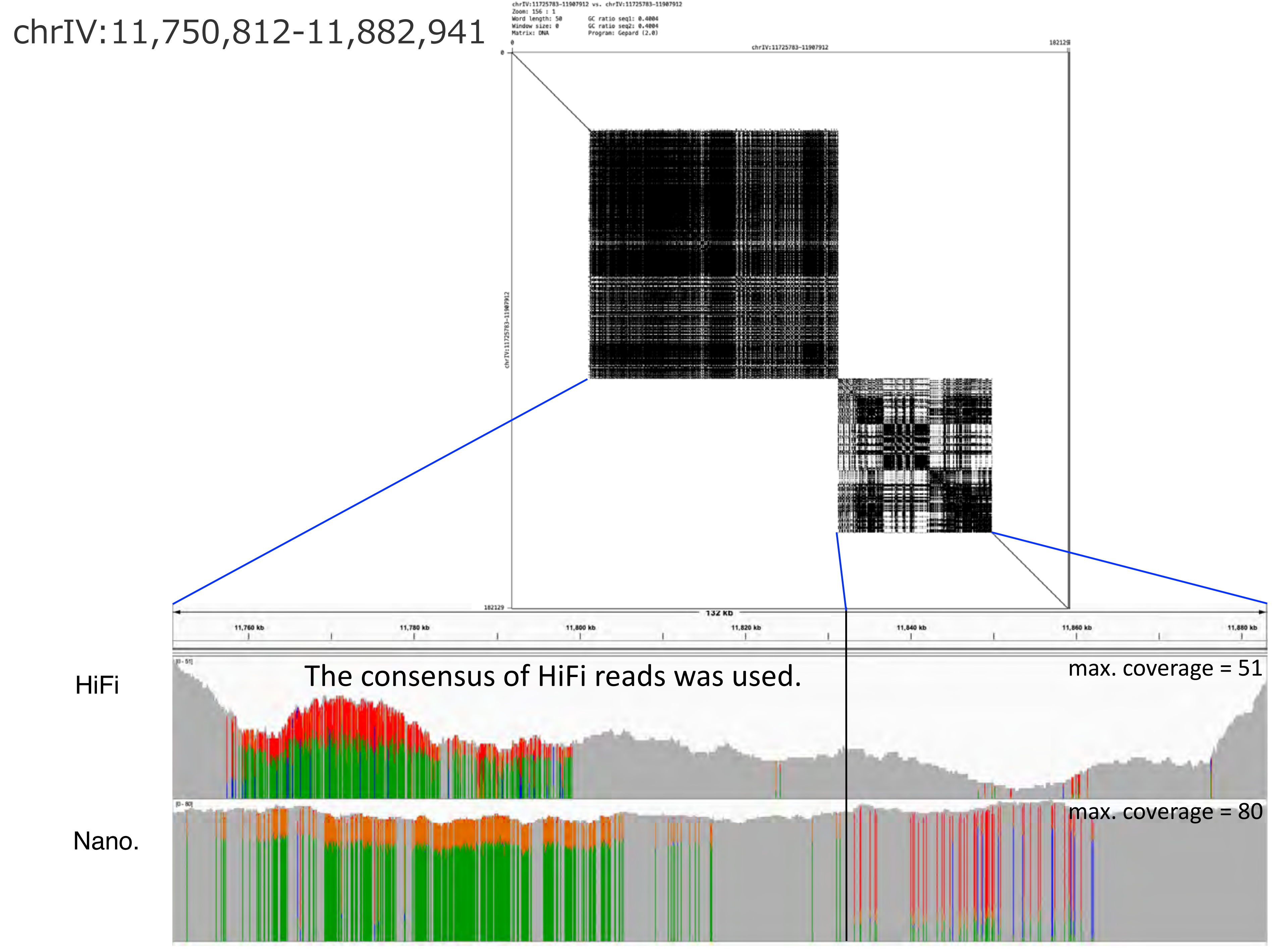

chrIV:11,750,812-11,882,941
chrIV:11725783-11907912 vs. chrIV:11725783-11907912
Zoom: 156 : 1
Word length: 50
Window size: 0
Matrix: DNA
GC ratio seq1: 0.4004
GC ratio seq2: 0.4004
Program: Gepard (2.0)
chrIV:11725783-11907912
0
182129
132 kb
11,760 kb
11,780 kb
11,800 kb
11,820 kb
11,840 kb
11,860 kb
11,880 kb
HiFi
The consensus of HiFi reads was used.
max. coverage = 51
Nano.
max. coverage = 80

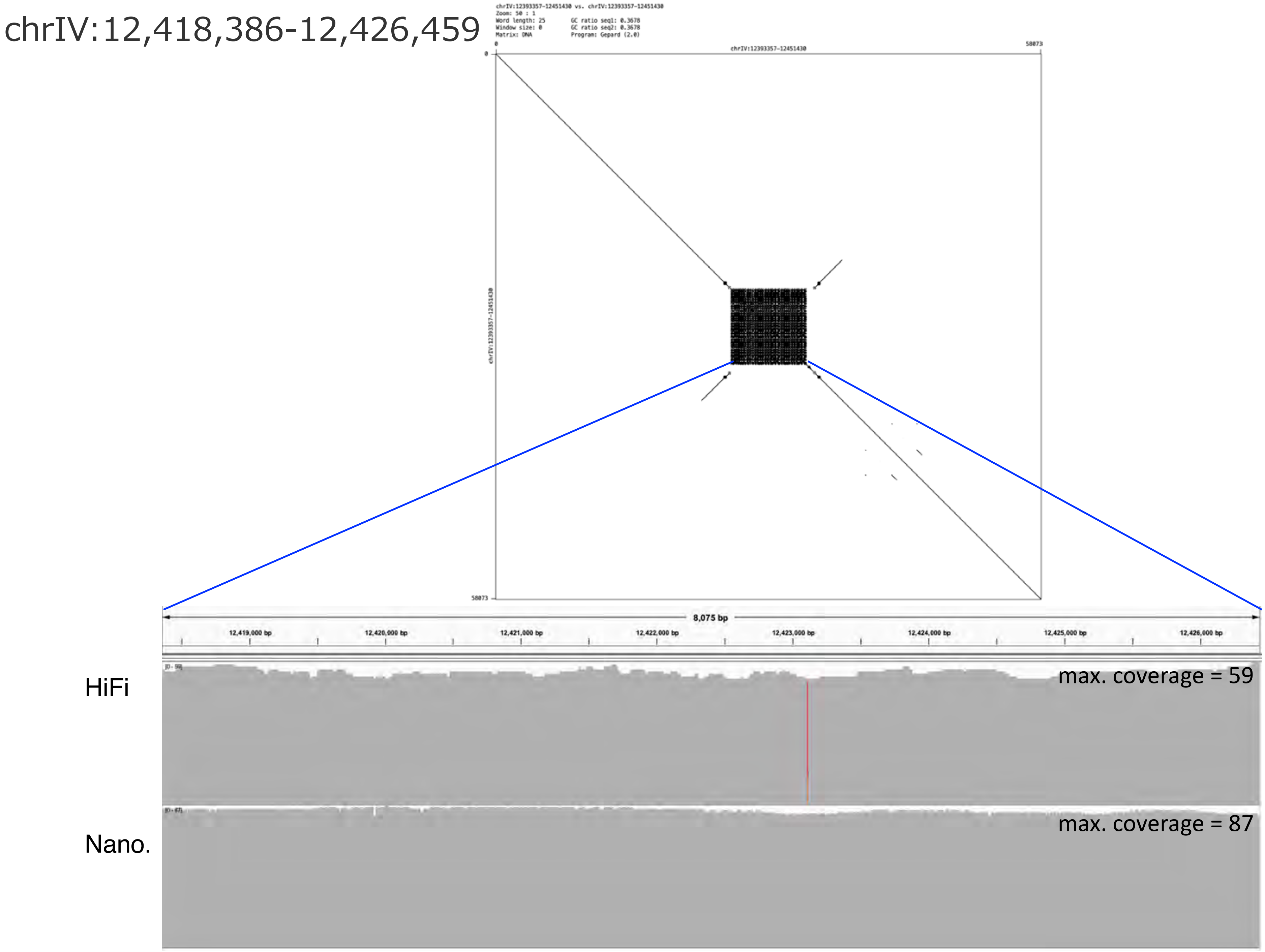

chrIV:12,418,386-12,426,459
chrIV:12393357-12451430 vs. chrIV:12393357-12451430
Zoom: 50 : 1
Word length: 25
Window size: 0
Matrix: DNA
GC ratio seq1: 0.3678
GC ratio seq2: 0.3678
Program: Gepard (2.0)
chrIV:12393357-12451430
0
58073
8,075 bp
12,419,000 bp
12,420,000 bp
12,421,000 bp
12,422,000 bp
12,423,000 bp
12,424,000 bp
12,425,000 bp
12,426,000 bp
HiFi
max. coverage = 59
Nano.
max. coverage = 87

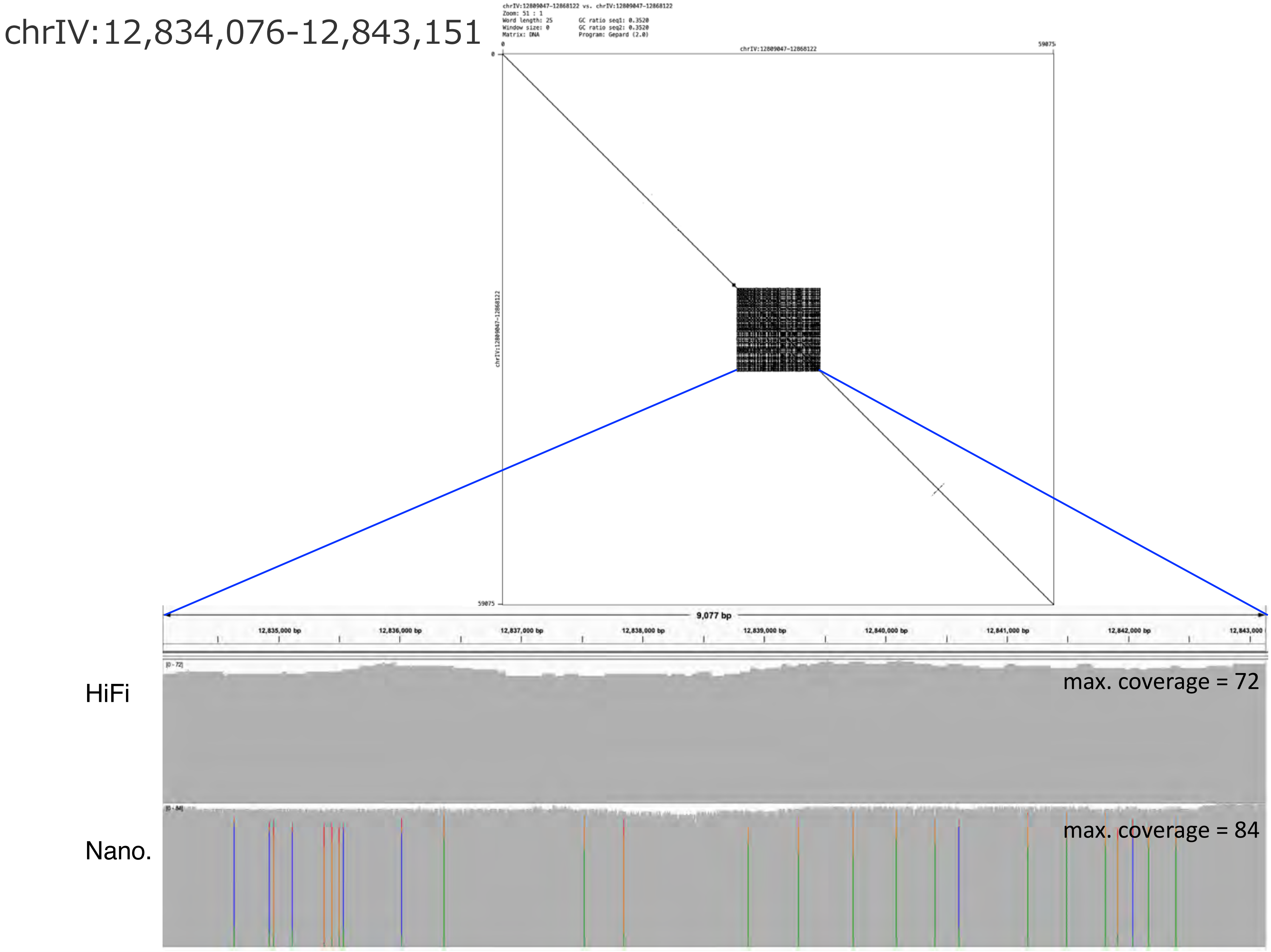

chrIV:12,834,076-12,843,151
chrIV:12809047-12868122 vs. chrIV:12809047-12868122
Zoom: 51 : 1
Word length: 25
Window size: 0
Matrix: DNA
GC ratio seq1: 0.3520
GC ratio seq2: 0.3520
Program: Gepard (2.0)
0
chrIV:12809047-12868122
59075
chrIV:12809047-12868122
9,077 bp
12,835,000 bp
12,836,000 bp
12,837,000 bp
12,838,000 bp
12,839,000 bp
12,840,000 bp
12,841,000 bp
12,842,000 bp
12,843,000
HiFi
max. coverage = 72
Nano.
max. coverage = 84

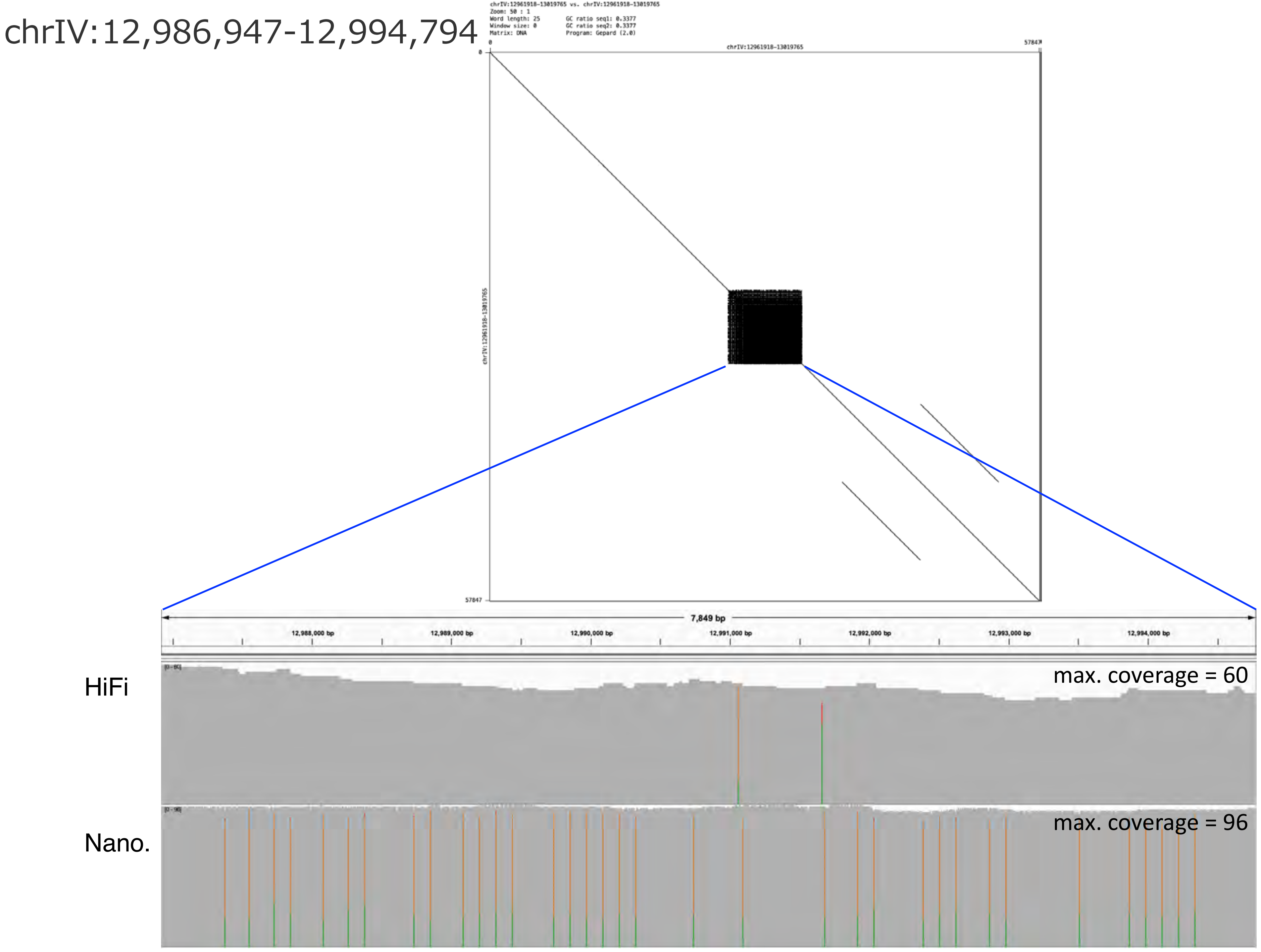
chrIV:12,986,947-12,994,794
chrIV:12961918-13019765 vs. chrIV:12961918-13019765
Zoom: 50 : 1
Word length: 25
Window size: 0
Matrix: DNA
GC ratio seq1: 0.3377
GC ratio seq2: 0.3377
Program: Gepard (2.0)
chrIV:12961918-13019765
57847
7,849 bp
12,988,000 bp
12,989,000 bp
12,990,000 bp
12,991,000 bp
12,992,000 bp
12,993,000 bp
12,994,000 bp
HiFi
max. coverage = 60
Nano.
max. coverage = 96

### chrIV:13,138,562-13,180,419

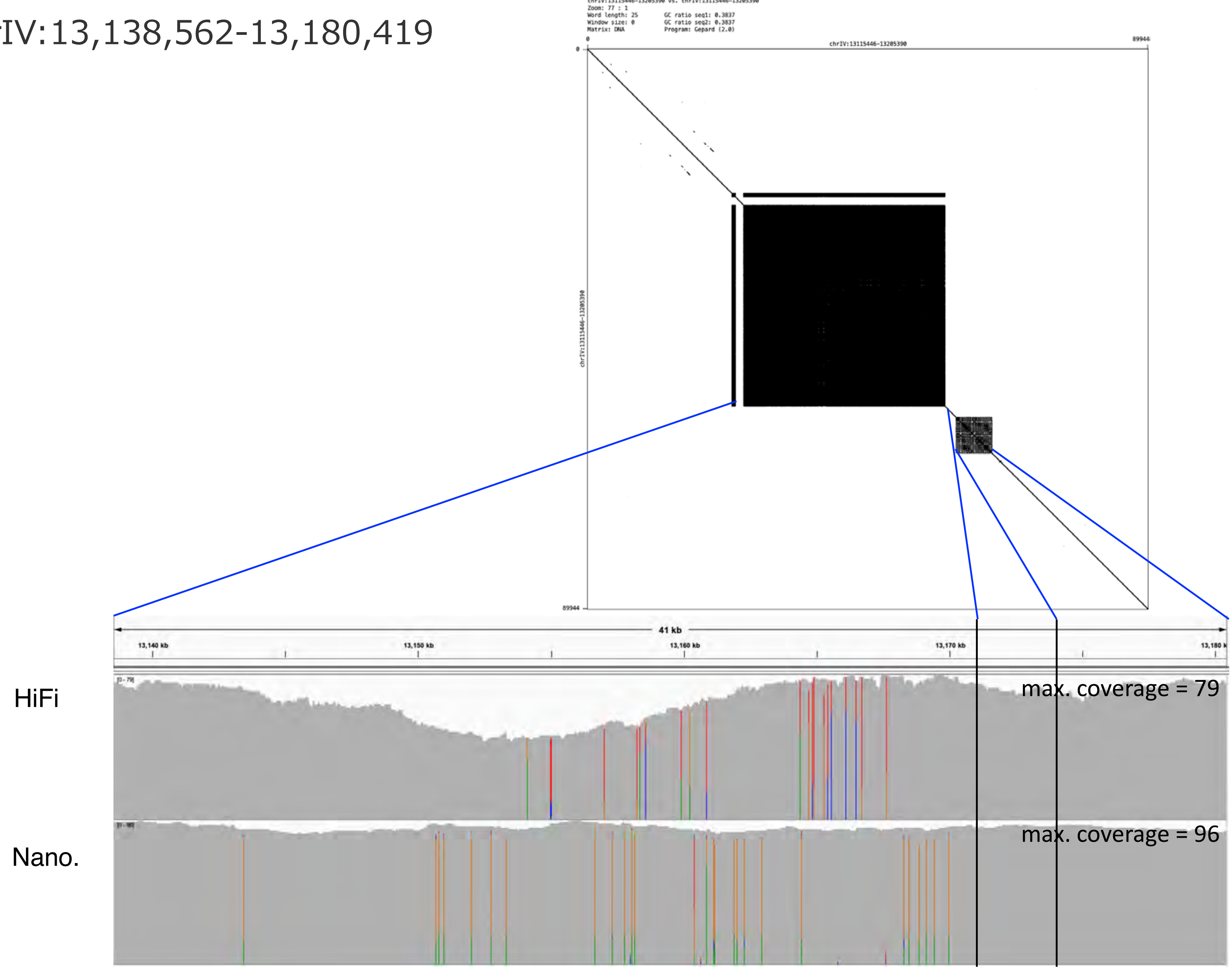

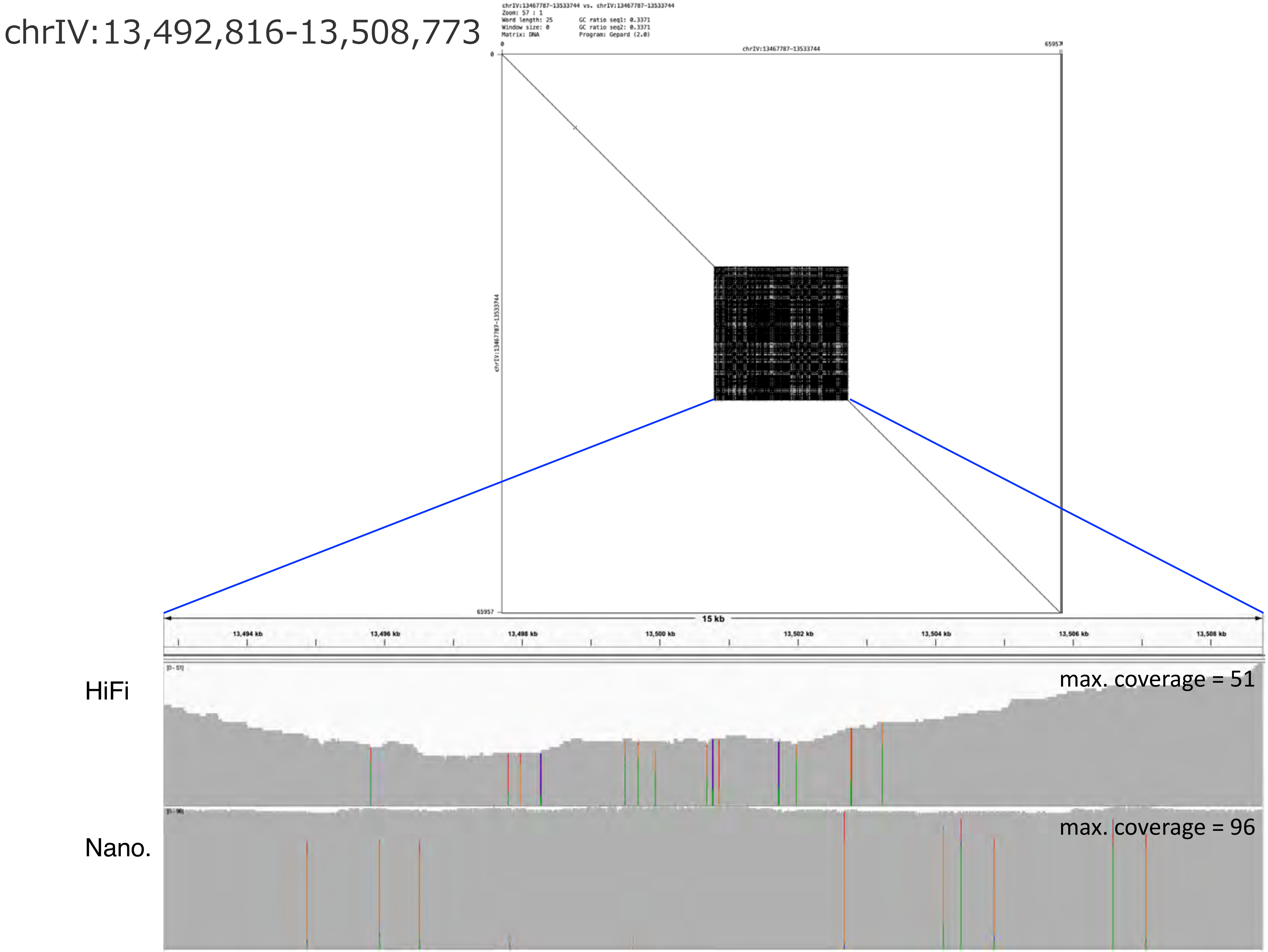
chrIV:13,492,816-13,508,773
chrIV:13467787-13533744 vs. chrIV:13467787-13533744
Zoom: 57 : 1
Word length: 25
Window size: 0
Matrix: DNA
GC ratio seq1: 0.3371
GC ratio seq2: 0.3371
Program: Gepard (2.0)
chrIV:13467787-13533744
0
65957
15 kb
13,494 kb
13,496 kb
13,498 kb
13,500 kb
13,502 kb
13,504 kb
13,506 kb
13,508 kb
HiFi
max. coverage = 51
Nano.
max. coverage = 96

chrIV:13,600,816-13,607,907

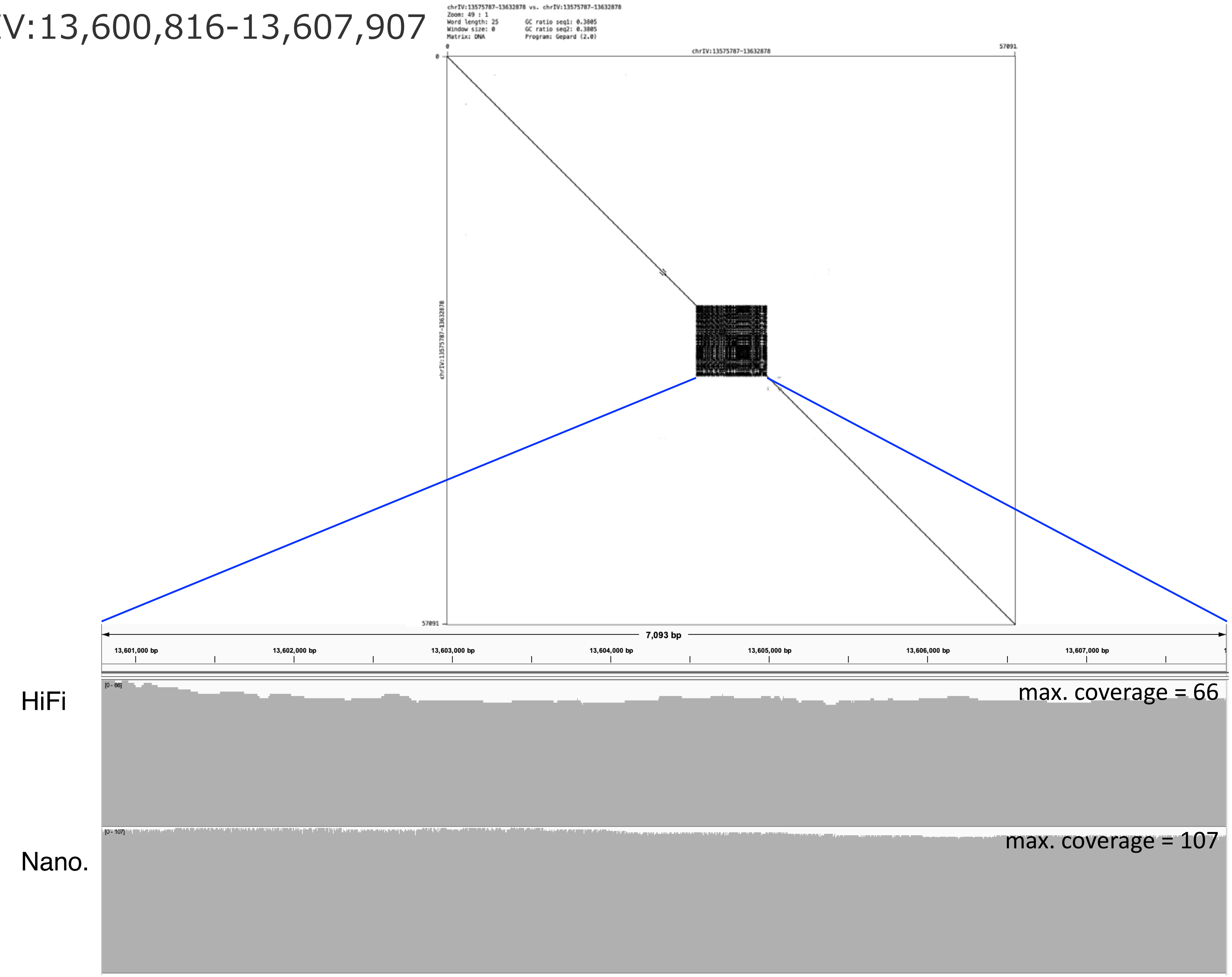

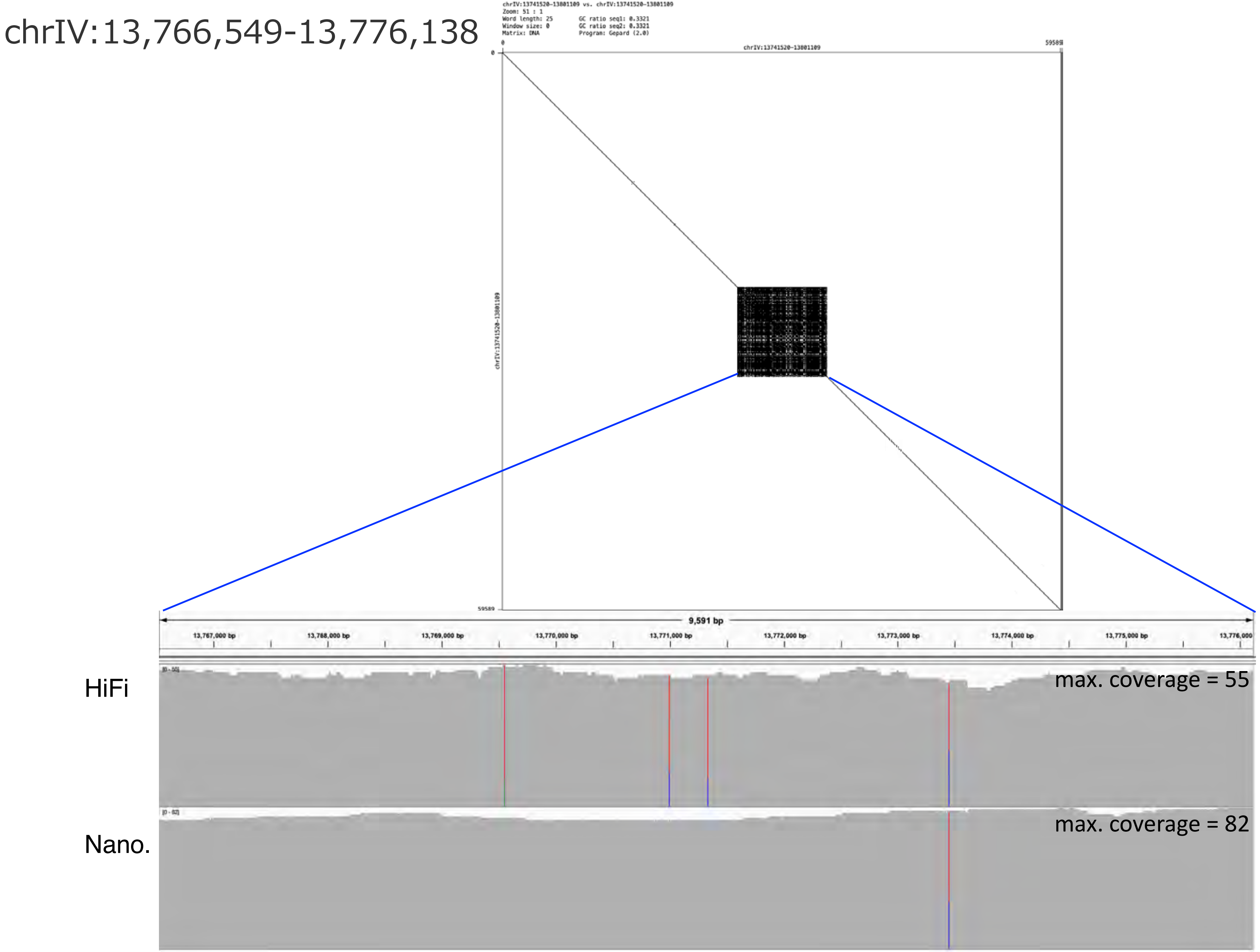
chrIV:13,766,549-13,776,138
chrIV:13741520-13801109 vs. chrIV:13741520-13801109
Zoom: 51 : 1
Word length: 25
Window size: 0
Matrix: DNA
GC ratio seq1: 0.3321
GC ratio seq2: 0.3321
Program: Gepard (2.0)
chrIV:13741520-13801109
9,591 bp
13,767,000 bp
13,768,000 bp
13,769,000 bp
13,770,000 bp
13,771,000 bp
13,772,000 bp
13,773,000 bp
13,774,000 bp
13,775,000 bp
13,776,000
HiFi
max. coverage = 55
Nano.
max. coverage = 82

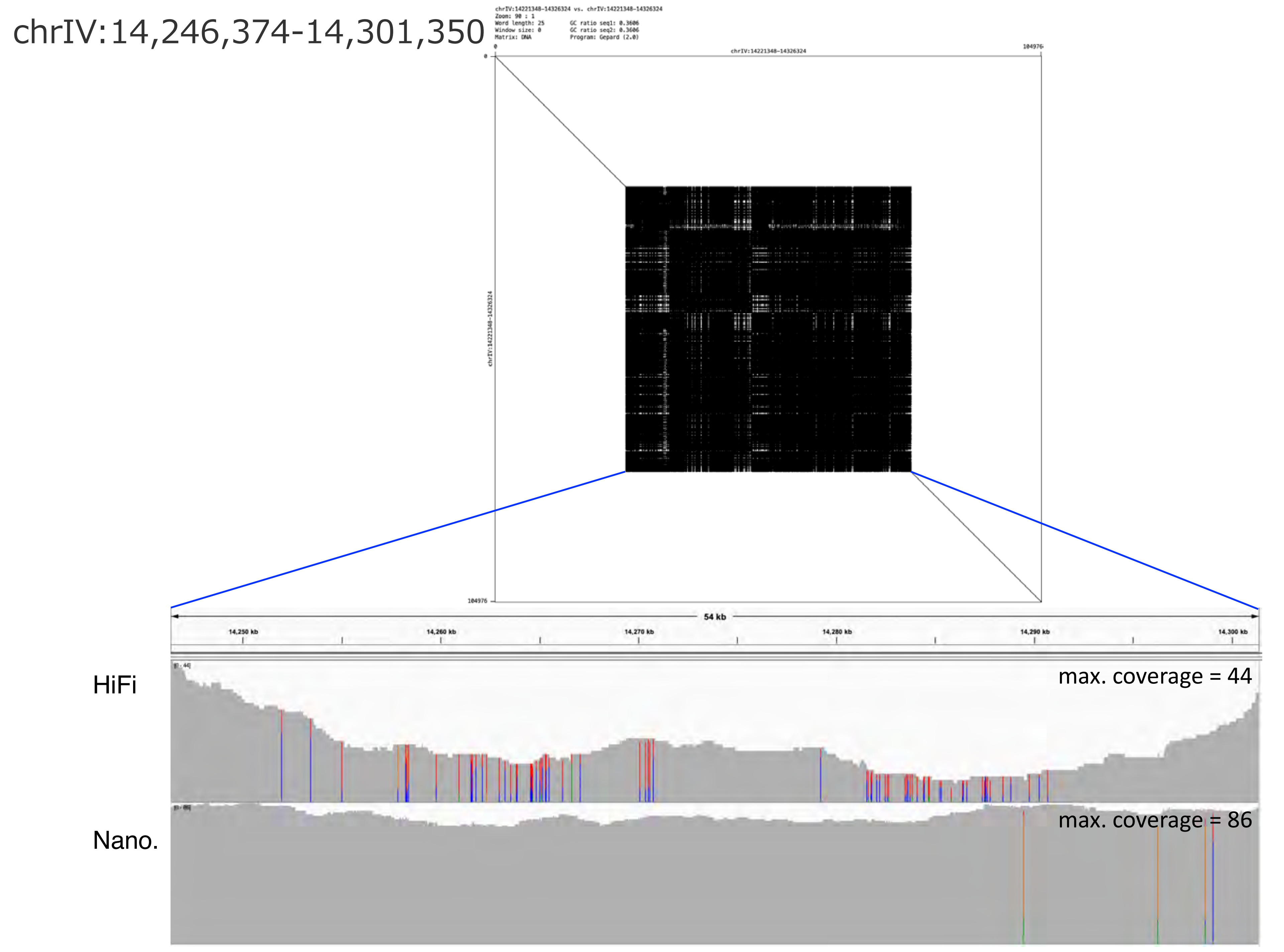
chrIV:14,246,374-14,301,350
chrIV:14221348-14326324 vs. chrIV:14221348-14326324
Zoom: 90 : 1
Word length: 25
Window size: 0
Matrix: DNA
GC ratio seq1: 0.3606
GC ratio seq2: 0.3606
Program: Gepard (2.0)
chrIV:14221348-14326324
0
104976
chrIV:14221348-14326324
54 kb
14,250 kb
14,260 kb
14,270 kb
14,280 kb
14,290 kb
14,300 kb
HiFi
max. coverage = 44
Nano.
max. coverage = 86

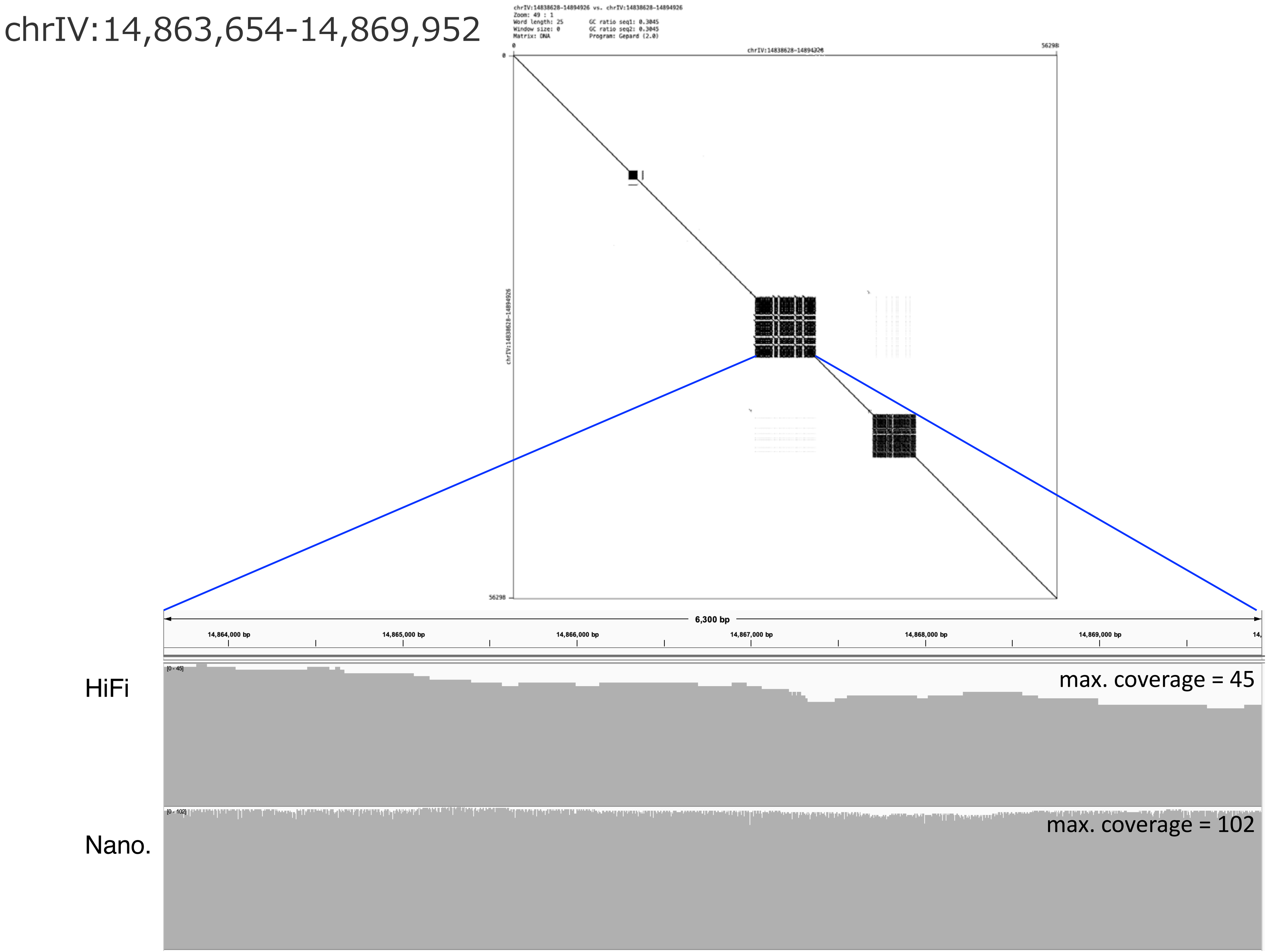
chrIV:14,863,654-14,869,952
chrIV:14838628-14894926 vs. chrIV:14838628-14894926
Zoom: 49 : 1
Word length: 25
Window size: 0
Matrix: DNA
GC ratio seq1: 0.3045
GC ratio seq2: 0.3045
Program: Gepard (2.0)
0
56298
chrIV:14838628-14894926
0
chrIV:14838628-14894926
56298
6,300 bp
14,864,000 bp
14,865,000 bp
14,866,000 bp
14,867,000 bp
14,868,000 bp
14,869,000 bp
HiFi
[0 - 45]
max. coverage = 45
Nano.
[0 - 102]
max. coverage = 102

chrIV:15,004,142-15,015,460

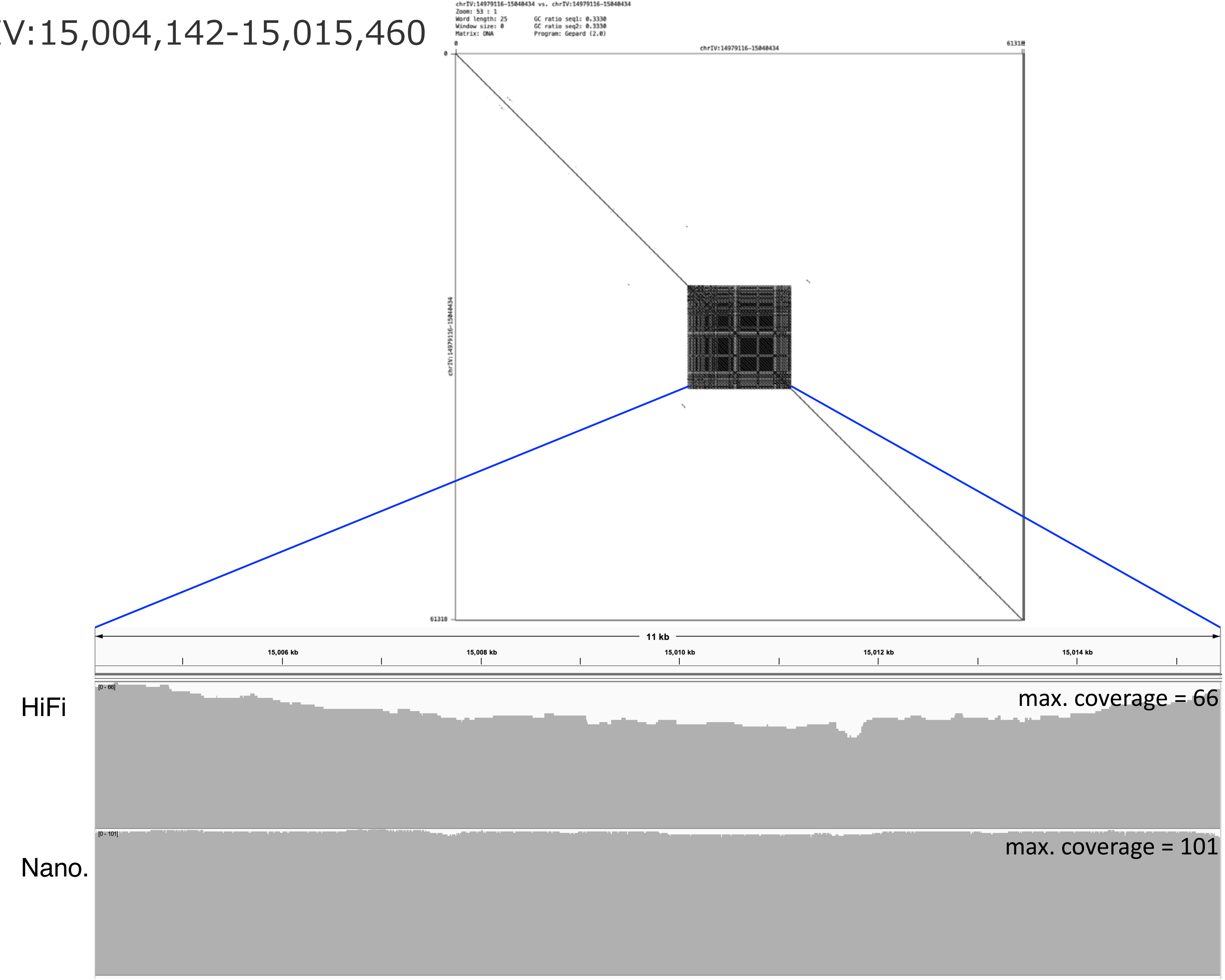

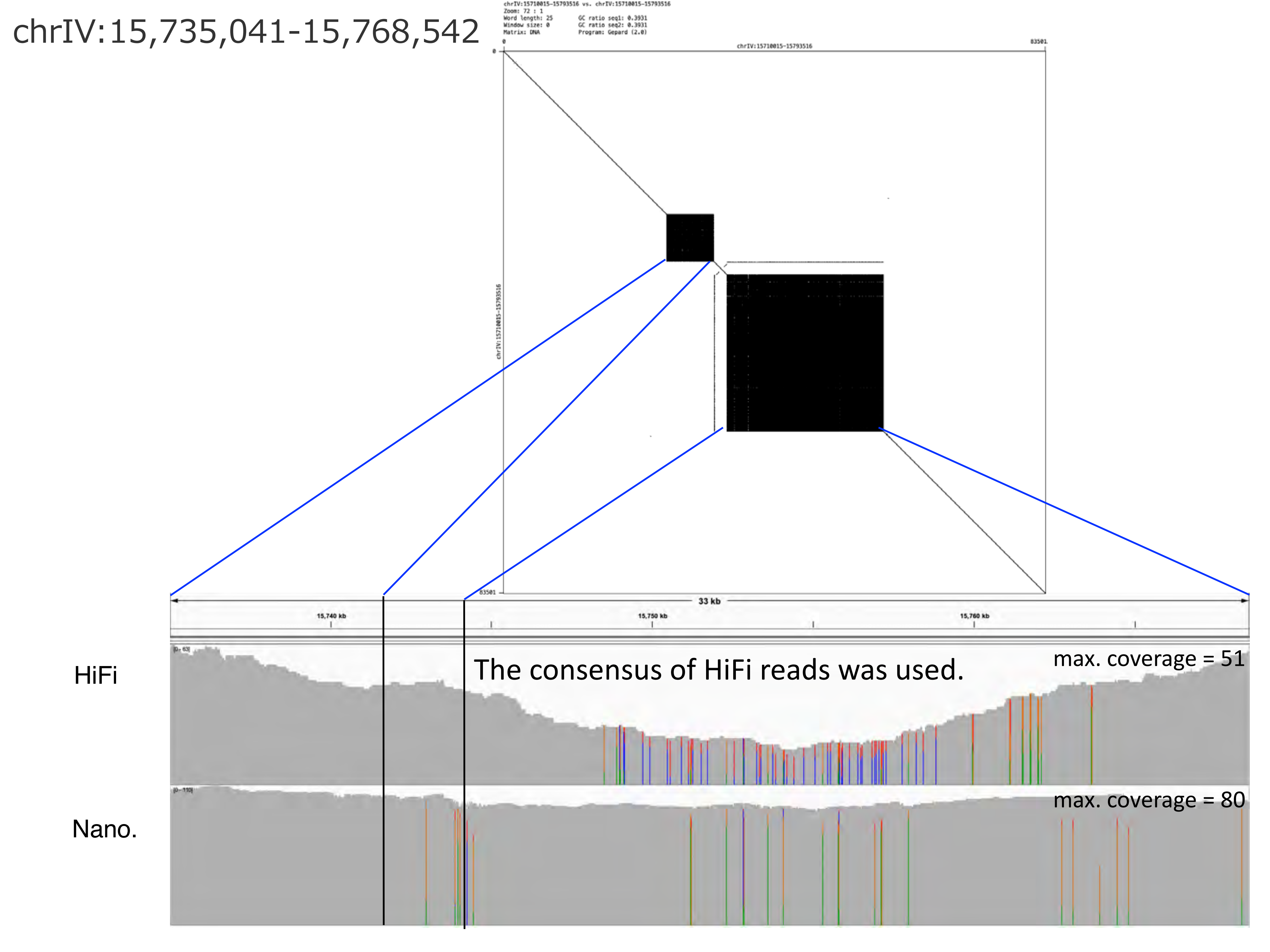
chrIV:15,735,041-15,768,542
chrIV:15710015-15793516 vs. chrIV:15710015-15793516
Zoom: 72 : 1
Word length: 25
Window size: 0
Matrix: DNA
GC ratio seq1: 0.3931
GC ratio seq2: 0.3931
Program: Gepard (2.0)
0
chrIV:15710015-15793516
83501
0
chrIV:15710015-15793516
83501
33 kb
15,740 kb
15,750 kb
15,760 kb
HiFi
The consensus of HiFi reads was used.
max. coverage = 51
Nano.
max. coverage = 80

chrIV:16,067,203-16,082,648

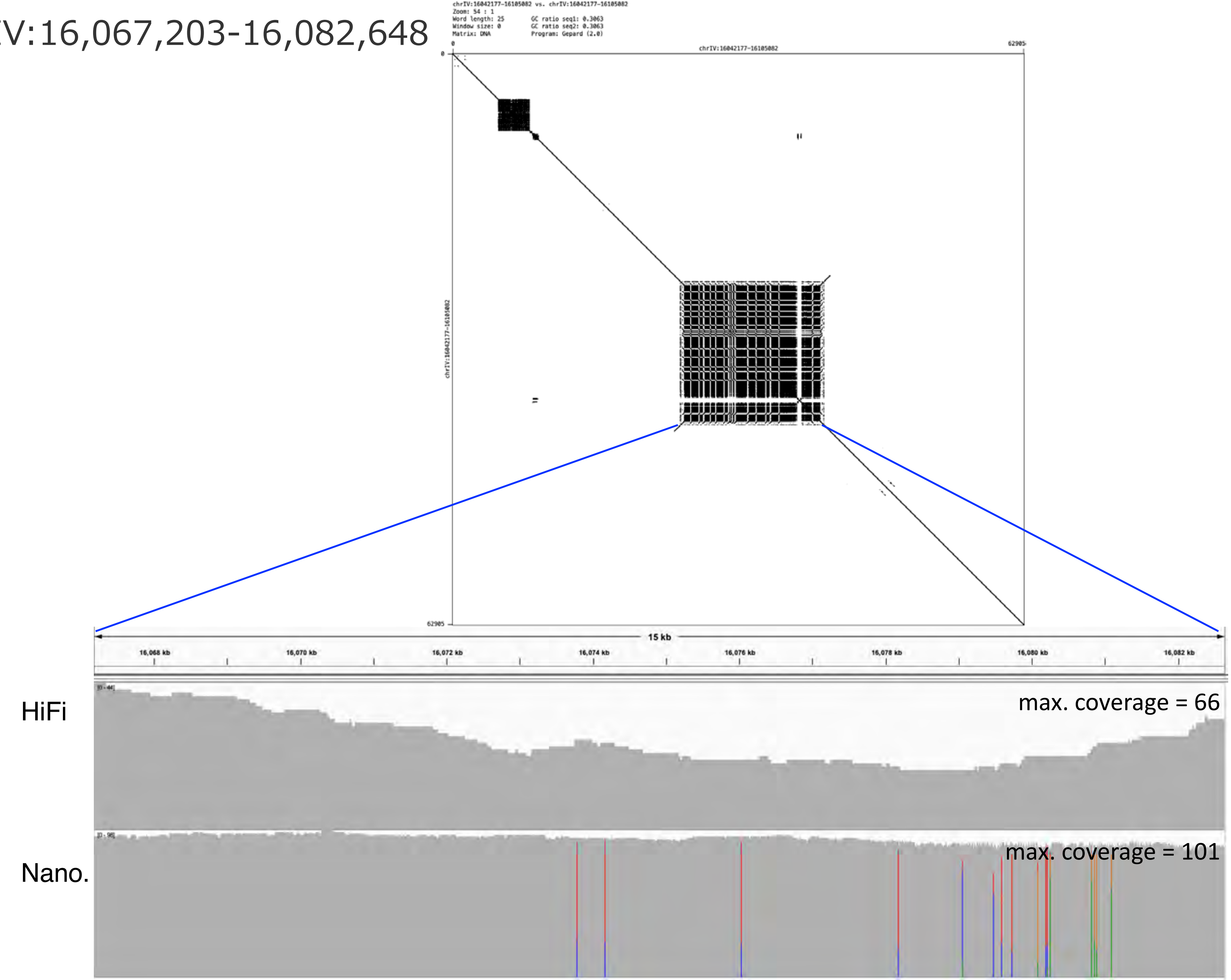

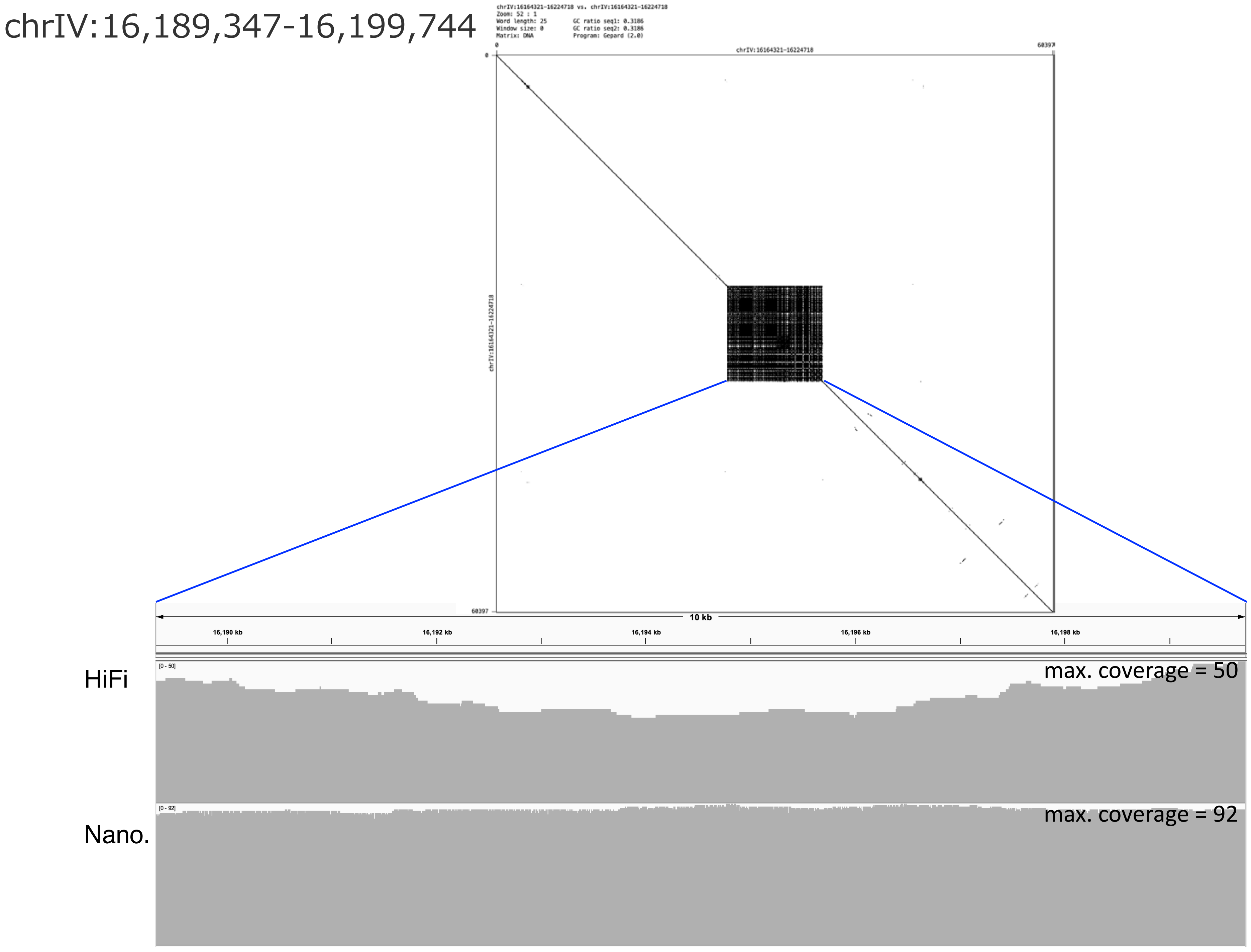
chrIV:16,189,347-16,199,744
chrIV:16164321-16224718 vs. chrIV:16164321-16224718
Zoom: 52 : 1
Word length: 25
Window size: 0
Matrix: DNA
GC ratio seq1: 0.3186
GC ratio seq2: 0.3186
Program: Gepard (2.0)
chrIV:16164321-16224718
60397
10 kb
16,190 kb
16,192 kb
16,194 kb
16,196 kb
16,198 kb
HiFi
[0 - 50]
max. coverage = 50
Nano.
[0 - 92]
max. coverage = 92

chrIV:16,904,413-16,942,307

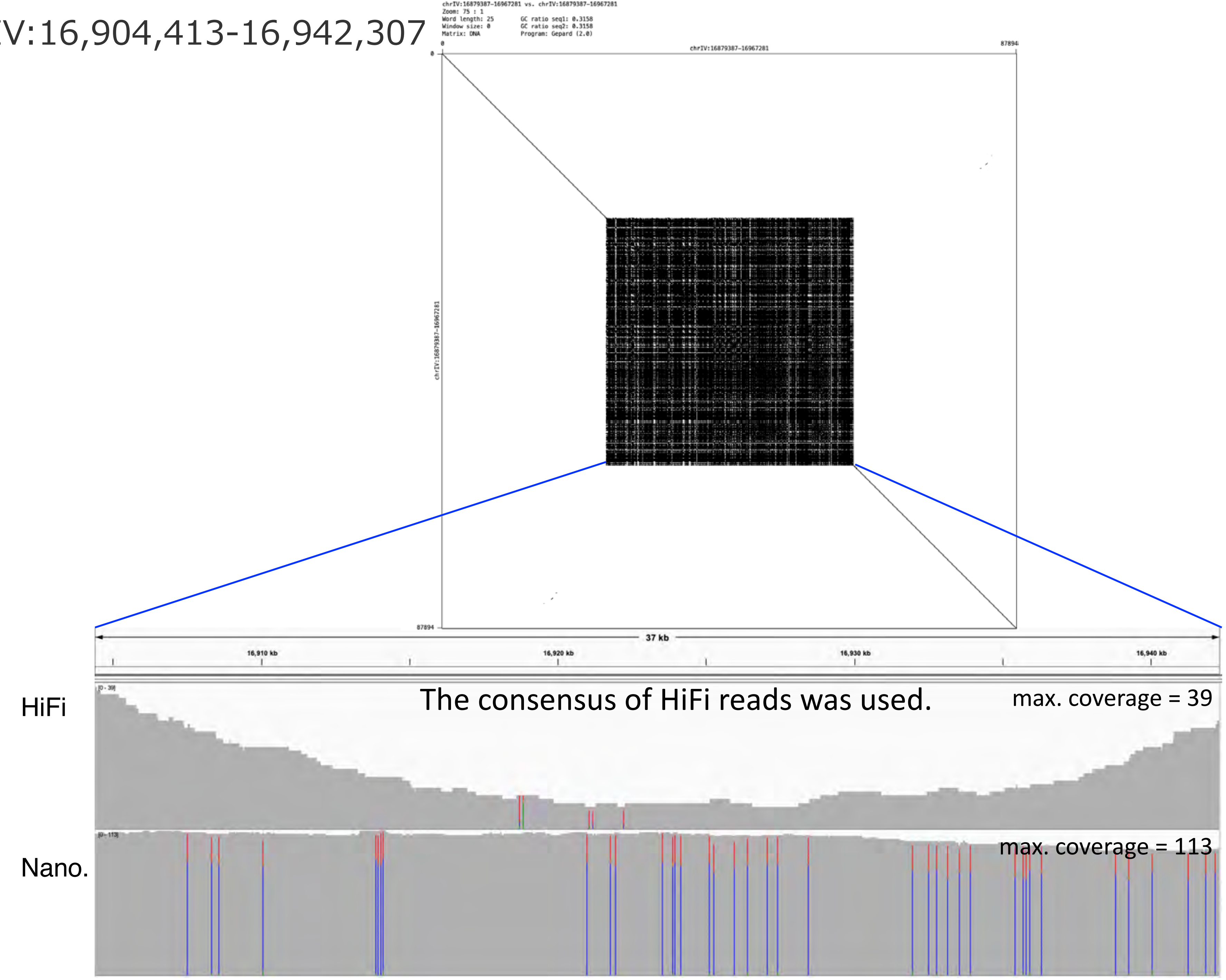

chrV:269,089-384,258

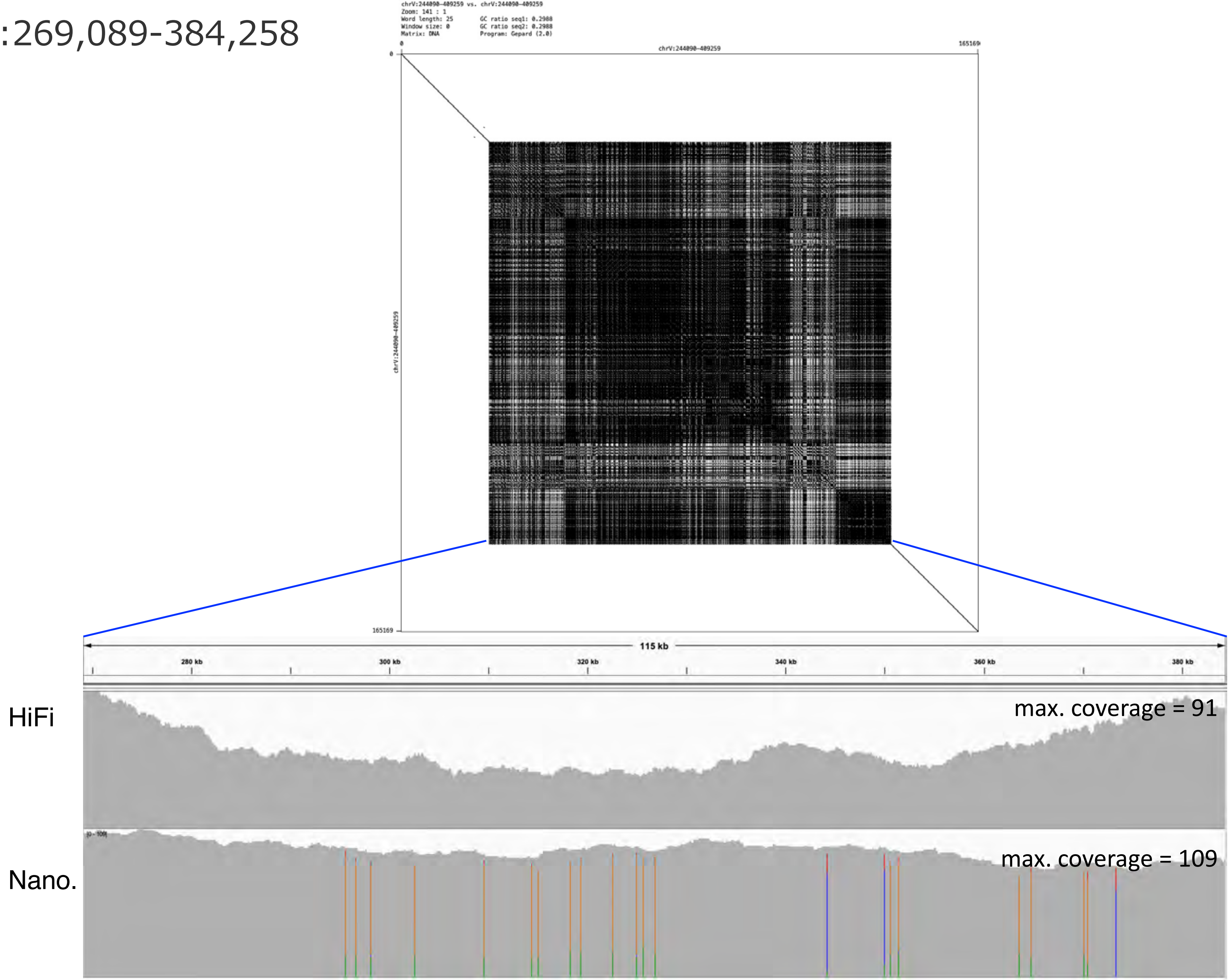

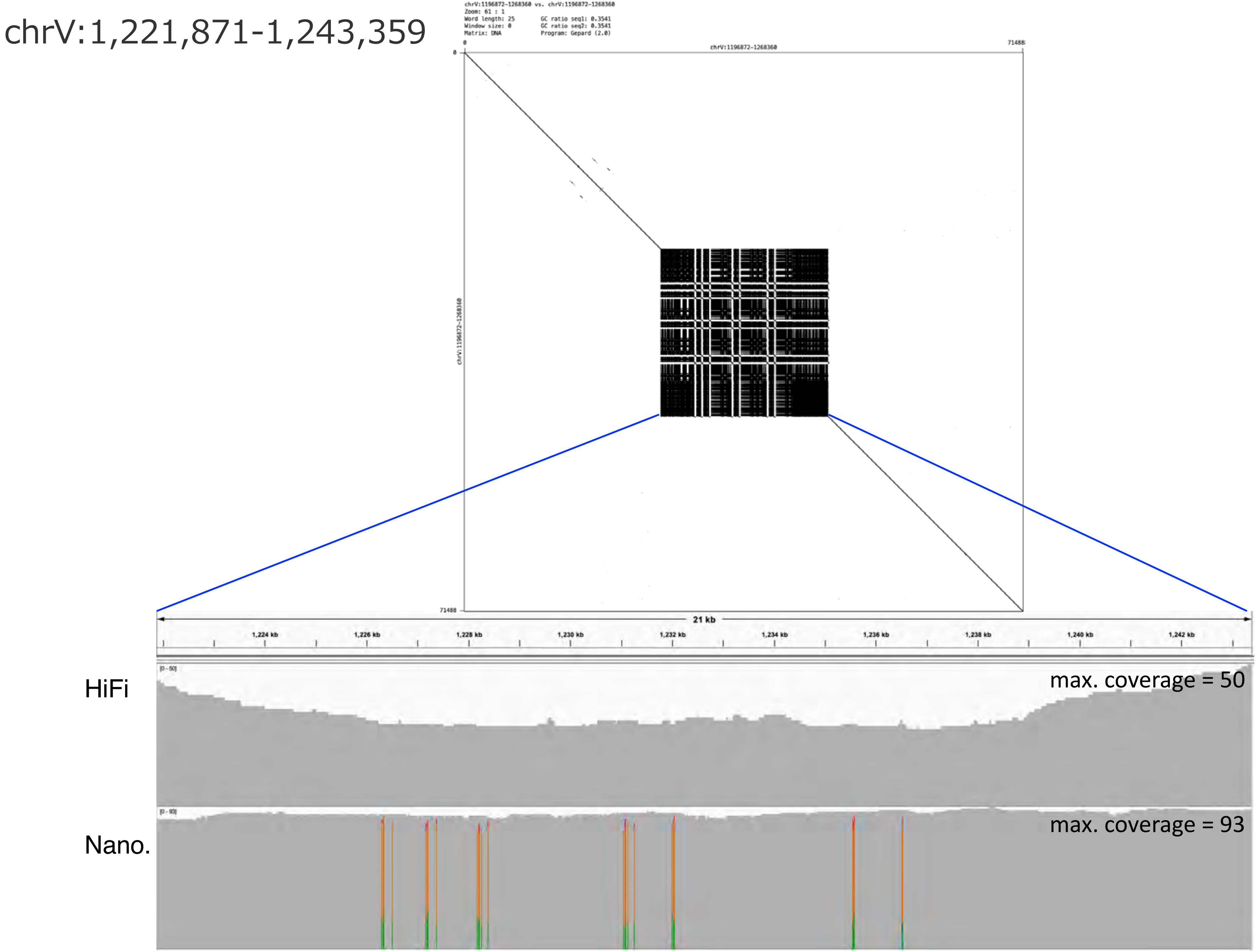
chrV:1,221,871-1,243,359
chrV:1196872-1268360 vs. chrV:1196872-1268360
Zoom: 61 : 1
Word length: 25
Window size: 0
Matrix: DNA
GC ratio seq1: 0.3541
GC ratio seq2: 0.3541
Program: Gepard (2.0)
chrV:1196872-1268360
0
71488
21 kb
1,224 kb
1,226 kb
1,228 kb
1,230 kb
1,232 kb
1,234 kb
1,236 kb
1,238 kb
1,240 kb
1,242 kb
HiFi
max. coverage = 50
Nano.
max. coverage = 93

chrV:1,783,925-1,808,427

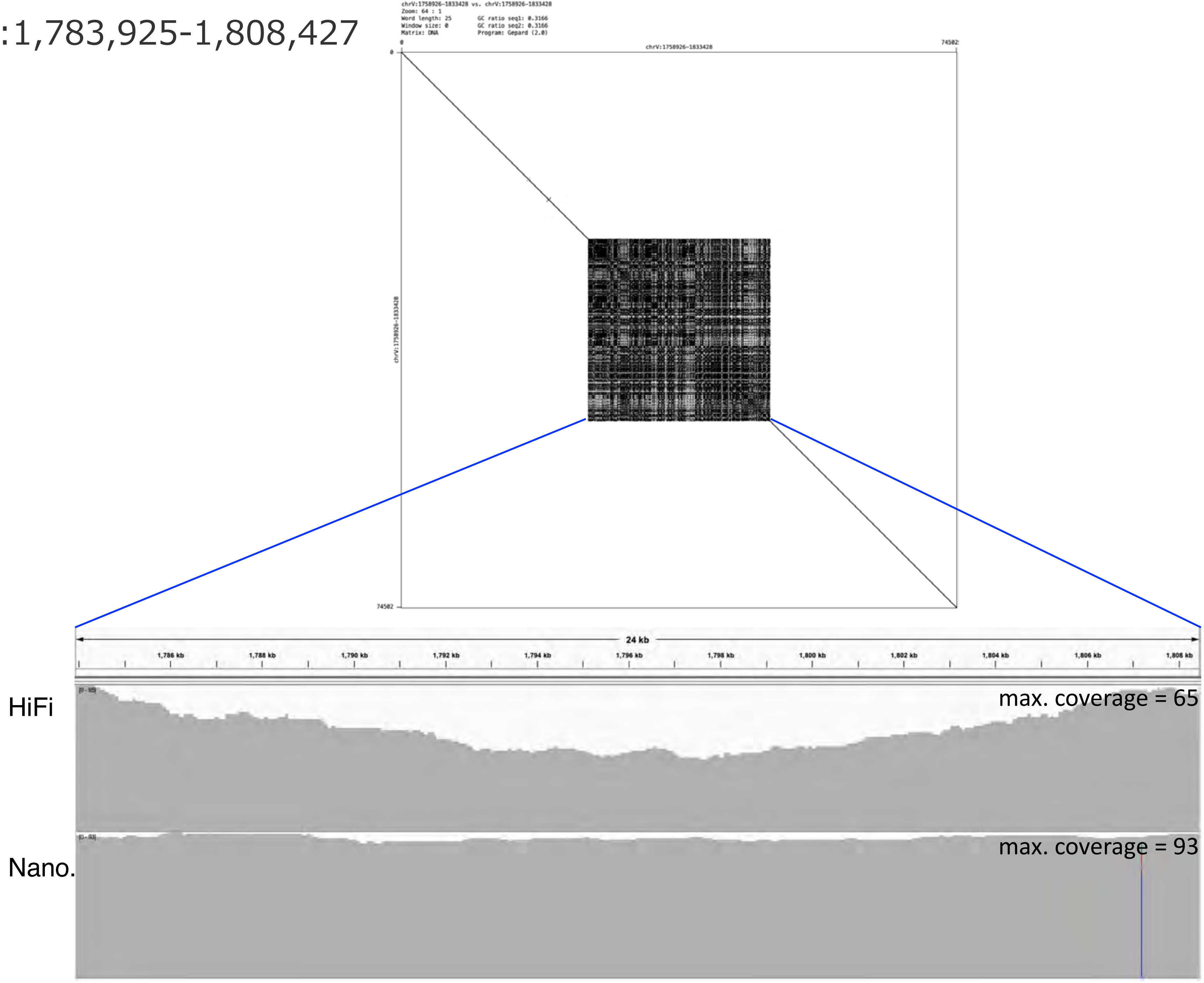

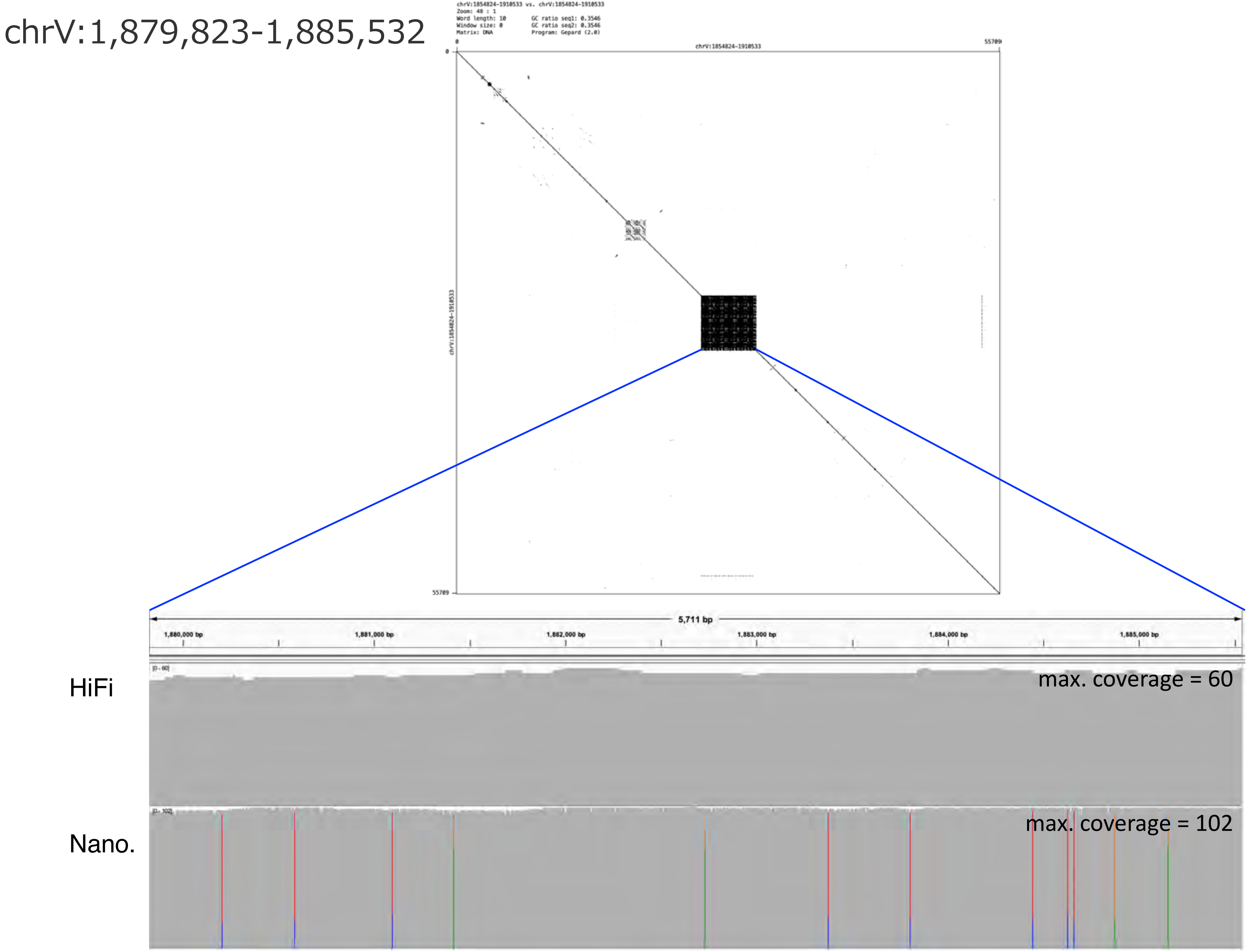
chrV:1,879,823-1,885,532
chrV:1854824-1910533 vs. chrV:1854824-1910533
Zoom: 48 : 1
Word length: 10
Window size: 0
Matrix: DNA
GC ratio seq1: 0.3546
GC ratio seq2: 0.3546
Program: Gepard (2.0)
chrV:1854824-1910533
0
55709
5,711 bp
1,880,000 bp
1,881,000 bp
1,882,000 bp
1,883,000 bp
1,884,000 bp
1,885,000 bp
HiFi
max. coverage = 60
Nano.
max. coverage = 102

### chrV:3,173,258-3,180,846

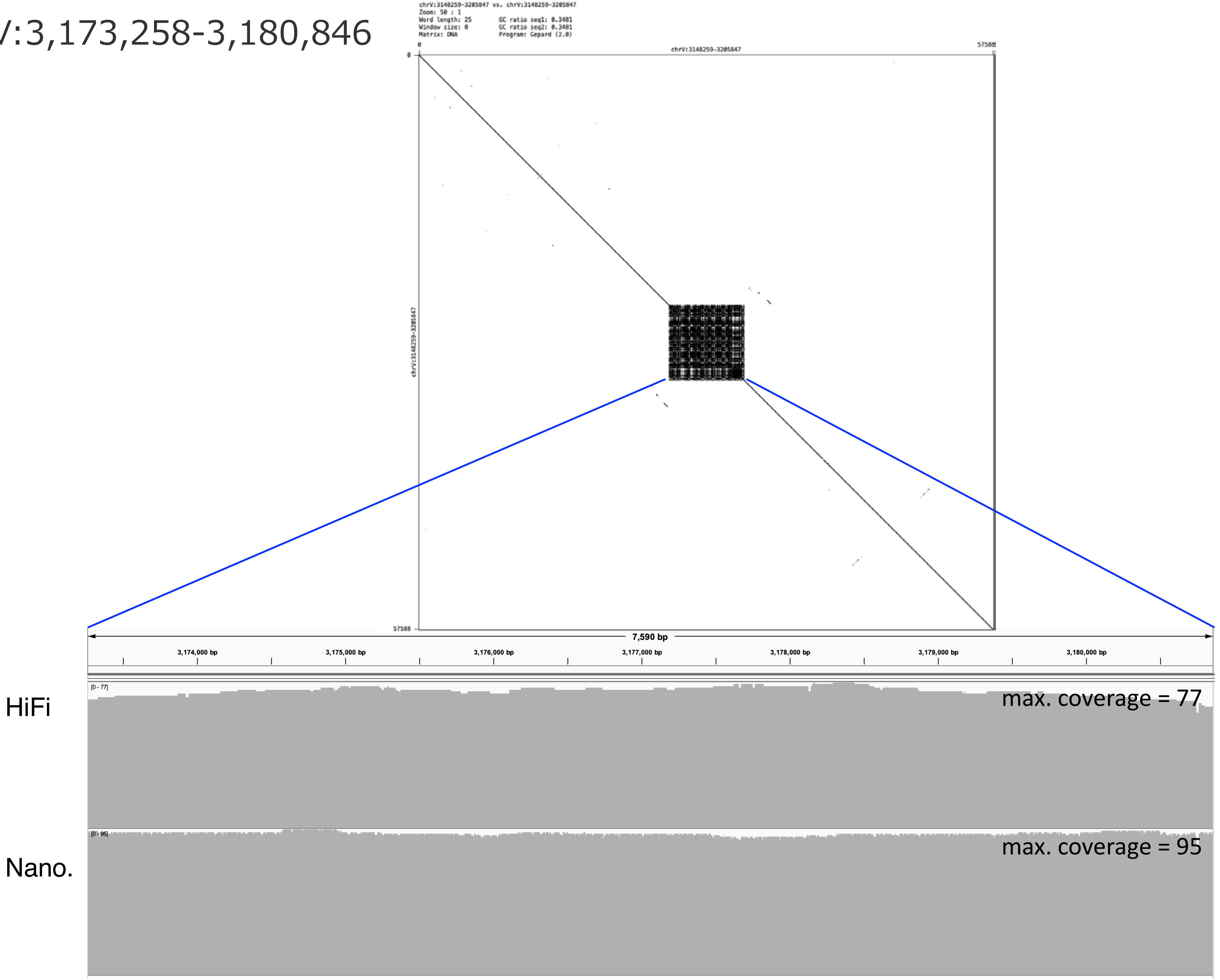

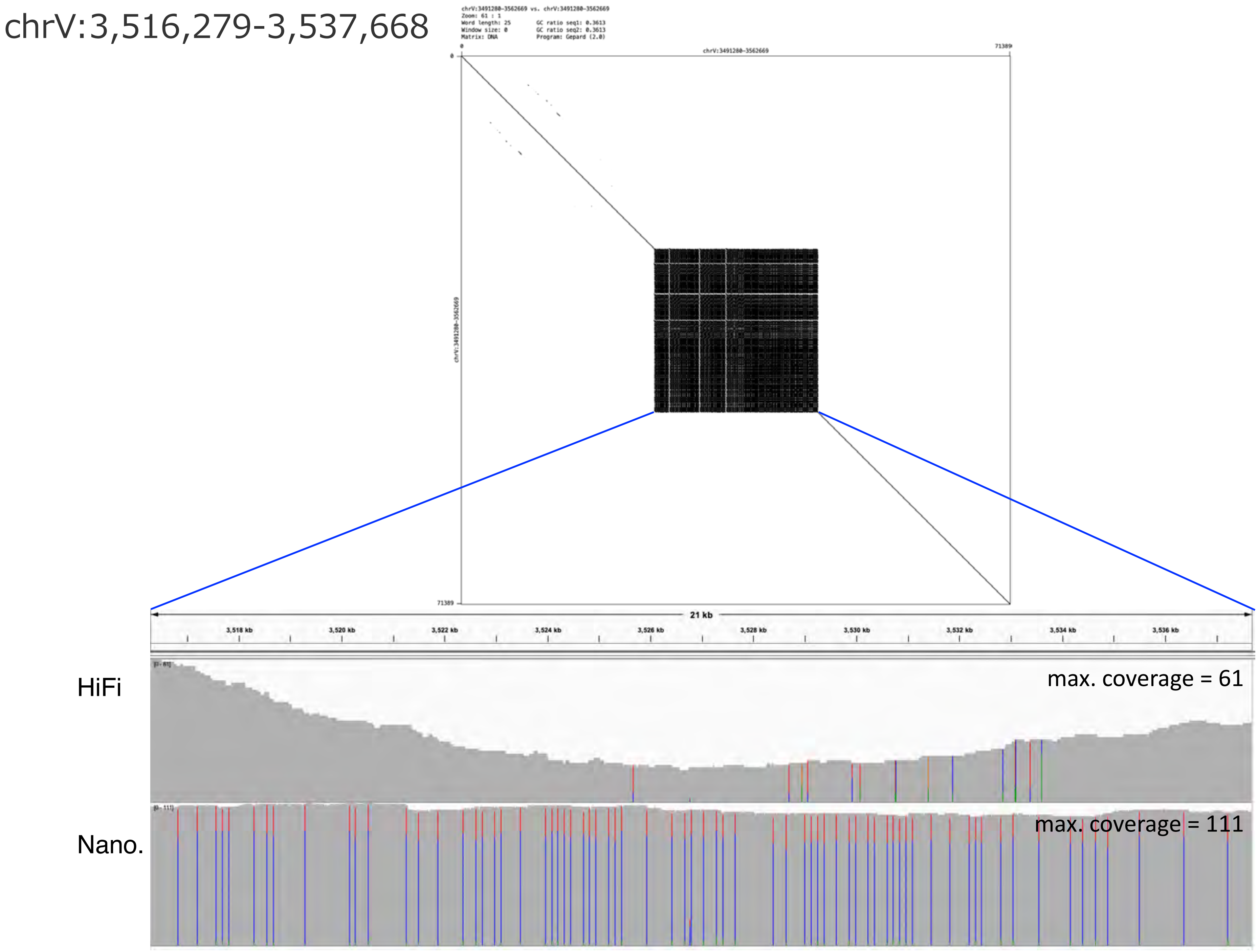

chrV:3,516,279-3,537,668
chrV:3491280-3562669 vs. chrV:3491280-3562669
Zoom: 61 : 1
Word length: 25
Window size: 0
Matrix: DNA
GC ratio seq1: 0.3613
GC ratio seq2: 0.3613
Program: Gepard (2.0)
chrV:3491280-3562669
21 kb
3,518 kb
3,520 kb
3,522 kb
3,524 kb
3,526 kb
3,528 kb
3,530 kb
3,532 kb
3,534 kb
3,536 kb
HiFi
max. coverage = 61
Nano.
max. coverage = 111

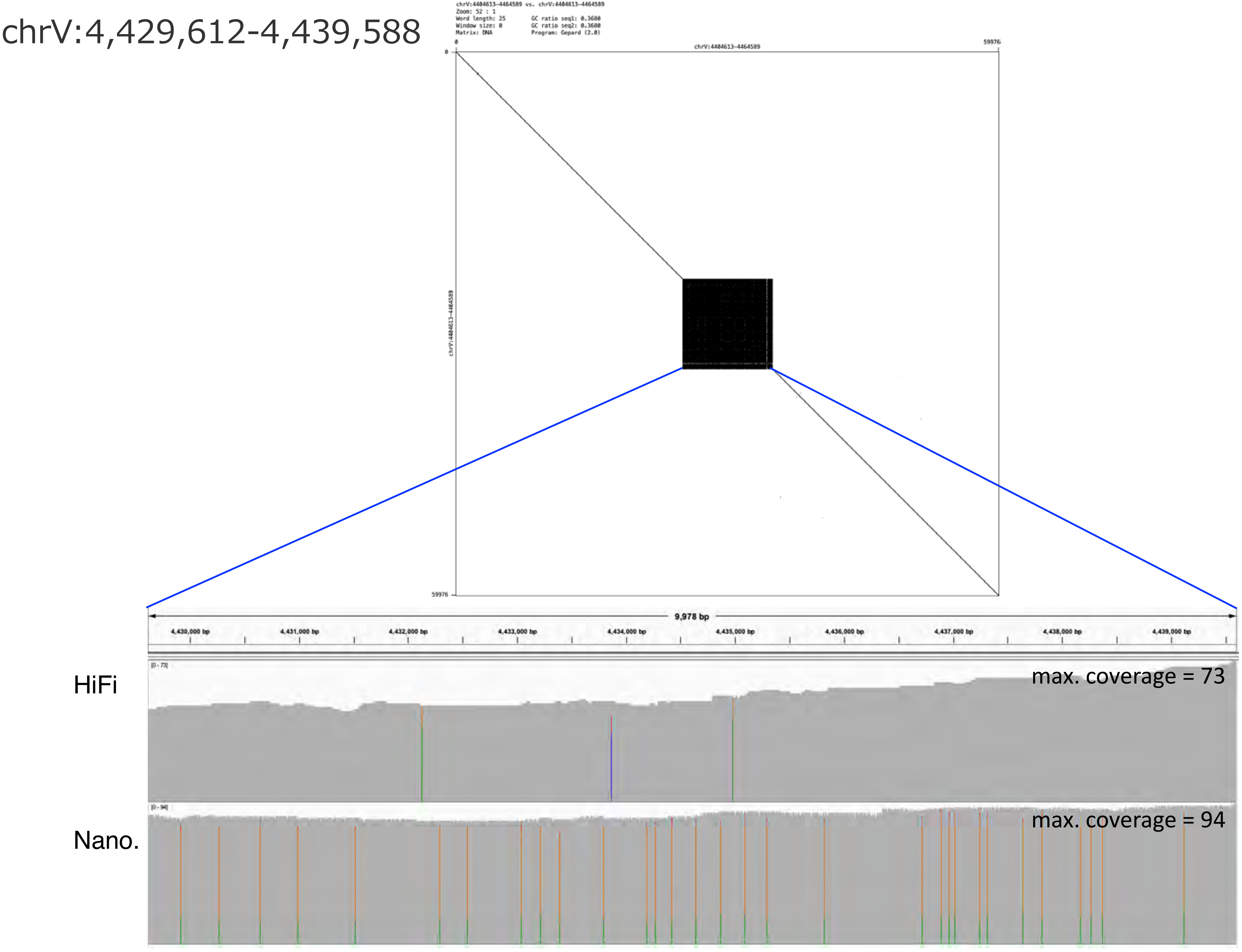
chrV:4,429,612-4,439,588
chrV:4404613-4464589 vs. chrV:4404613-4464589
Zoom: 52 : 1
Word length: 25
Window size: 0
Matrix: DNA
GC ratio seq1: 0.3680
GC ratio seq2: 0.3680
Program: Gepard (2.0)
chrV:4404613-4464589
0
59976
chrV:4404613-4464589
9,978 bp
4,430,000 bp
4,431,000 bp
4,432,000 bp
4,433,000 bp
4,434,000 bp
4,435,000 bp
4,436,000 bp
4,437,000 bp
4,438,000 bp
4,439,000 bp
HiFi
[0 - 73]
max. coverage = 73
Nano.
[0 - 94]
max. coverage = 94

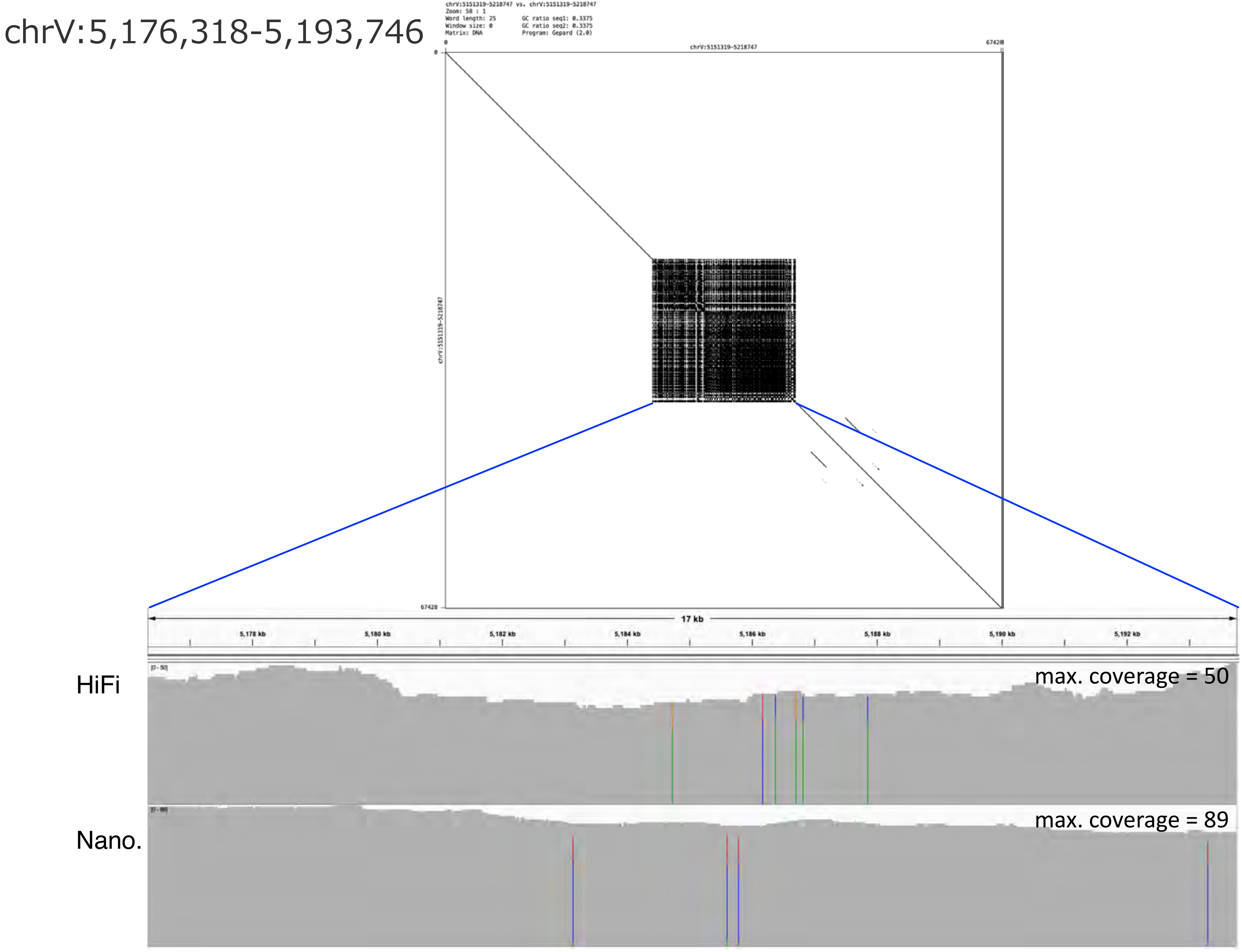
chrV:5,176,318-5,193,746
chrV:5151319-5218747 vs. chrV:5151319-5218747
Zoom: 58 : 1
Word length: 25
Window size: 0
Matrix: DNA
GC ratio seq1: 0.3375
GC ratio seq2: 0.3375
Program: Gepard (2.0)
chrV:5151319-5218747
0
67428
17 kb
5,178 kb
5,180 kb
5,182 kb
5,184 kb
5,186 kb
5,188 kb
5,190 kb
5,192 kb
HiFi
[0 - 50]
max. coverage = 50
Nano.
max. coverage = 89

chrV:5,400,257-5,506,731

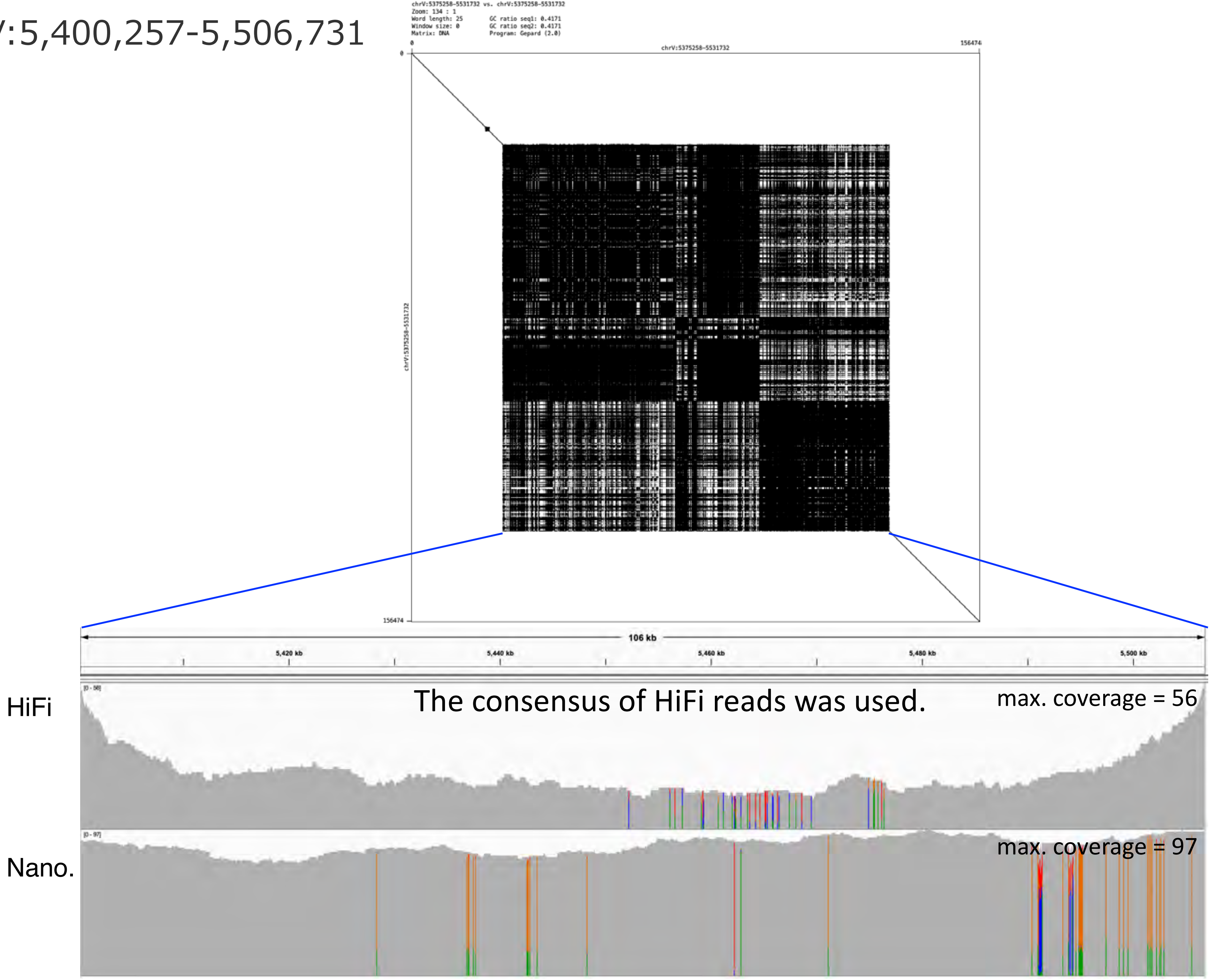

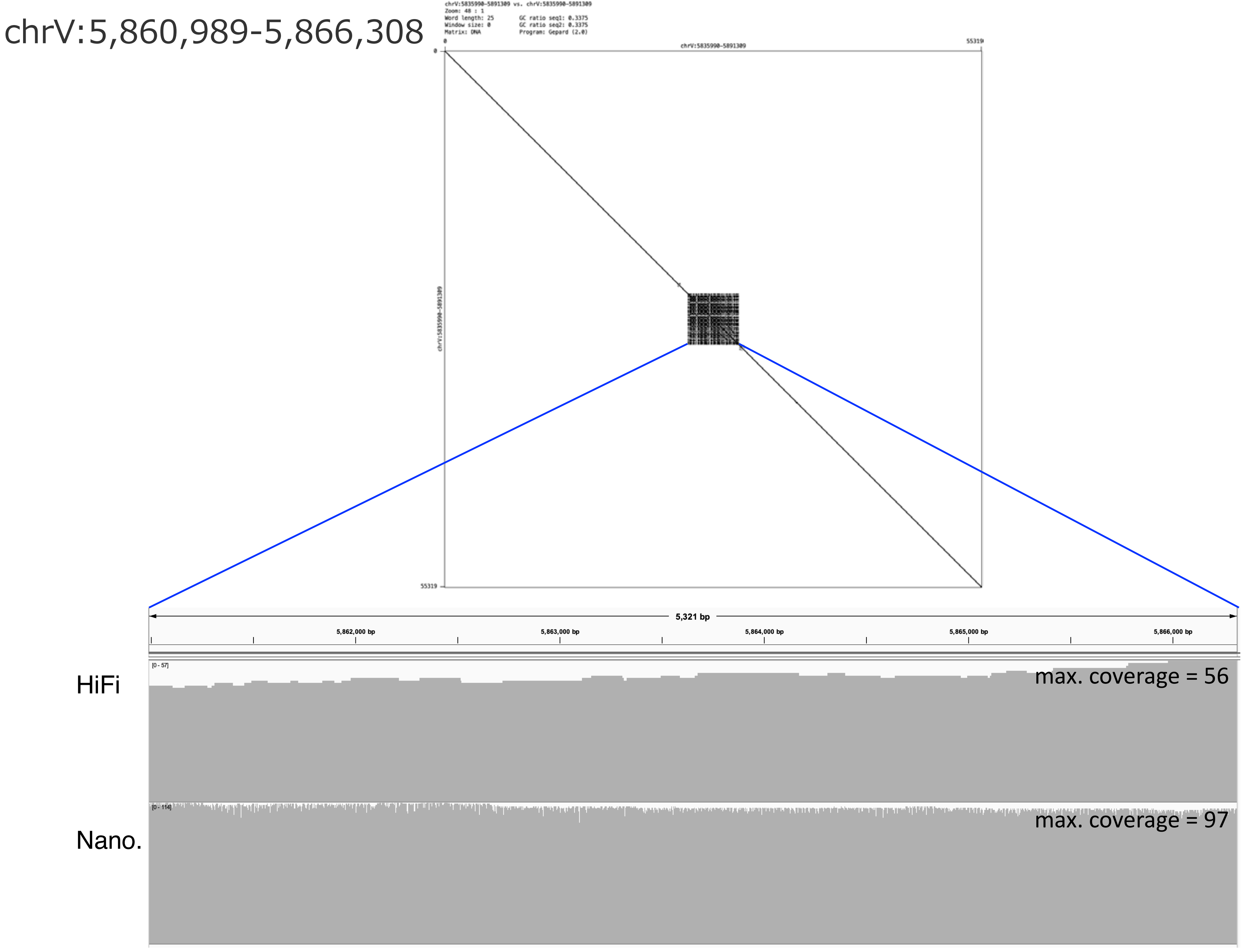
chrV:5,860,989-5,866,308
chrV:5835990-5891309 vs. chrV:5835990-5891309
Zoom: 48 : 1
Word length: 25
Window size: 0
Matrix: DNA
GC ratio seq1: 0.3375
GC ratio seq2: 0.3375
Program: Gepard (2.0)
0
chrV:5835990-5891309
55319
0
chrV:5835990-5891309
55319
5,321 bp
5,862,000 bp
5,863,000 bp
5,864,000 bp
5,865,000 bp
5,866,000 bp
HiFi
[0 - 57]
max. coverage = 56
Nano.
[0 - 114]
max. coverage = 97

chrV:6,295,268-6,326,116

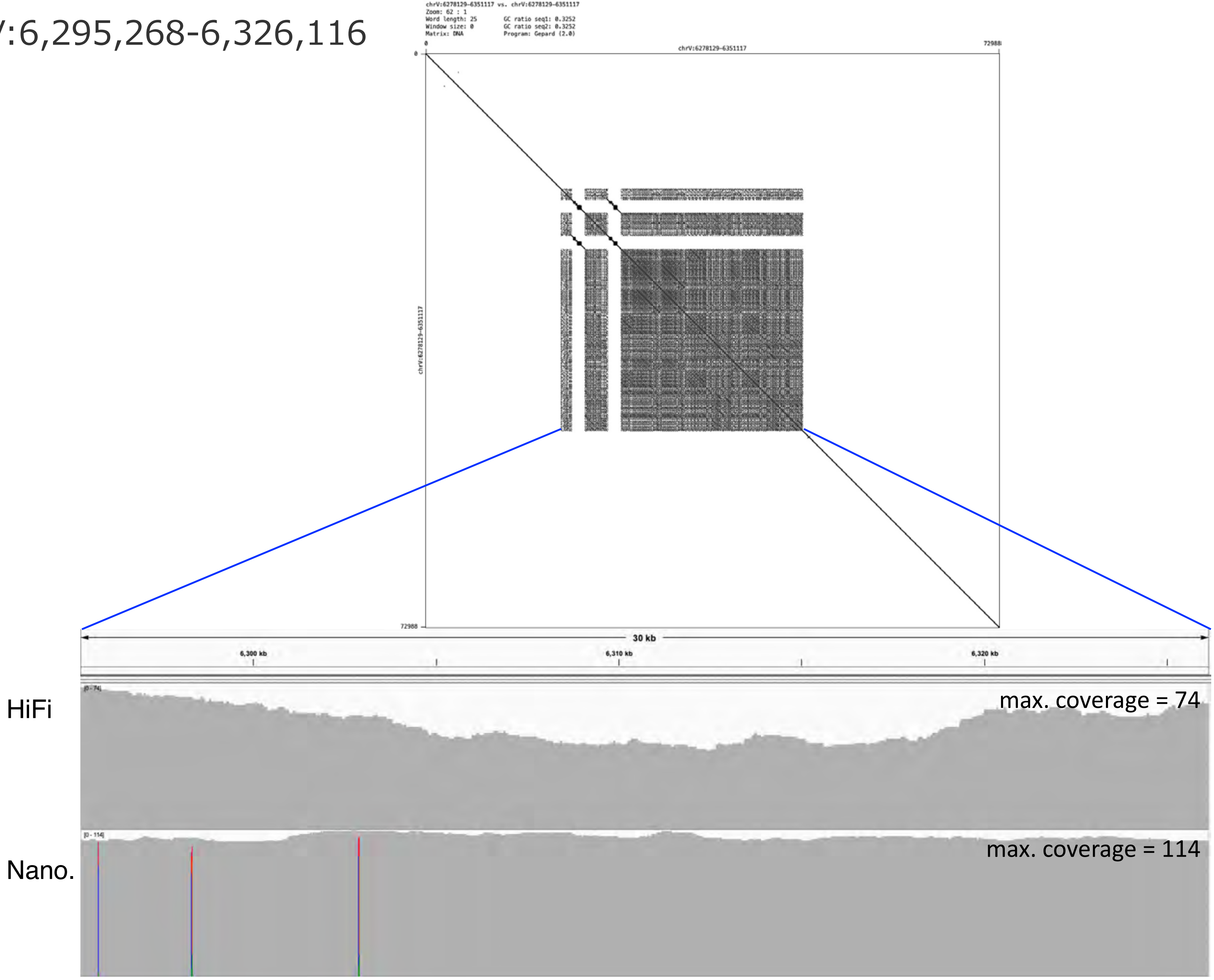

chrV:6,402,973-6,437,584

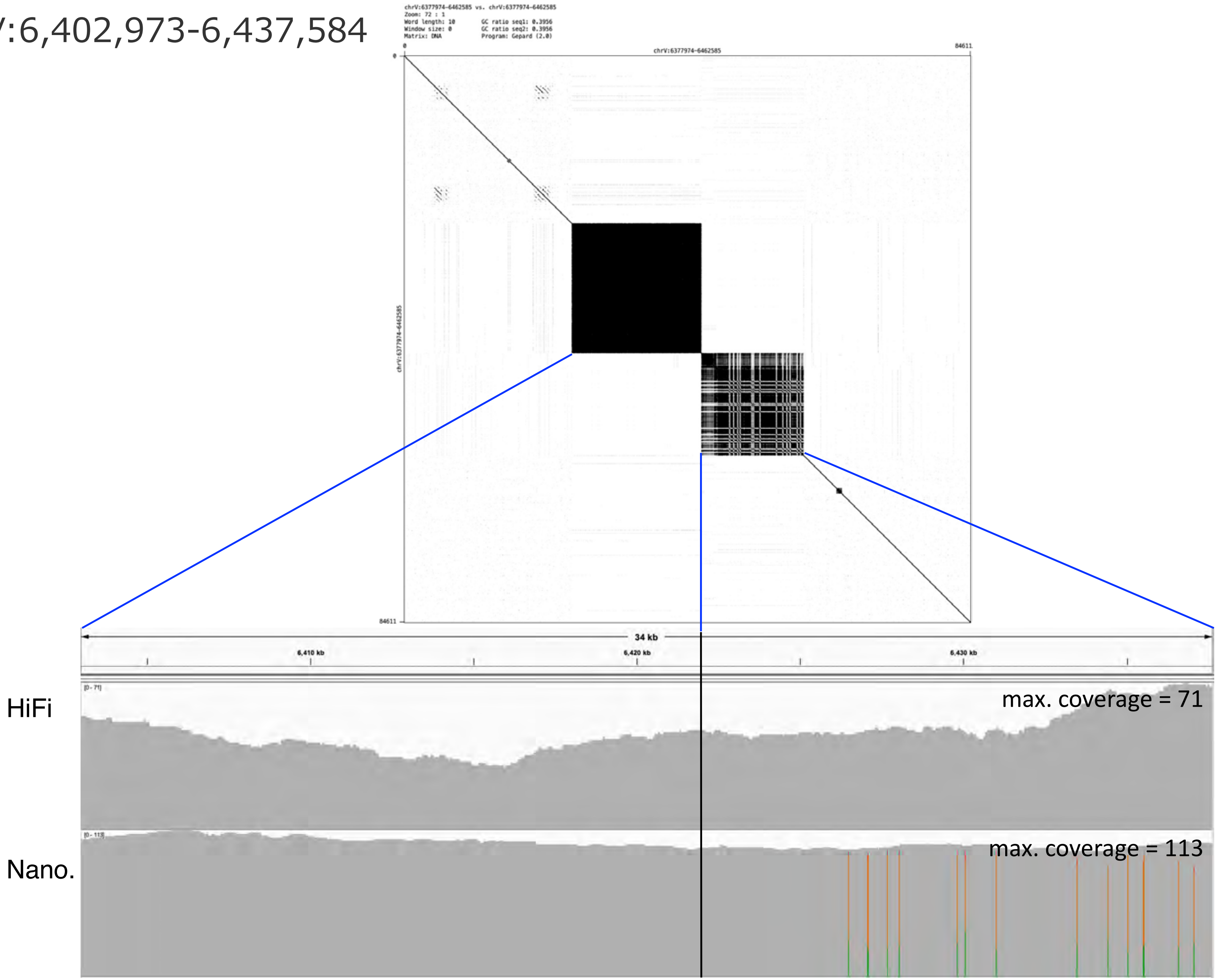

chrV:7,198,682-7,225,639

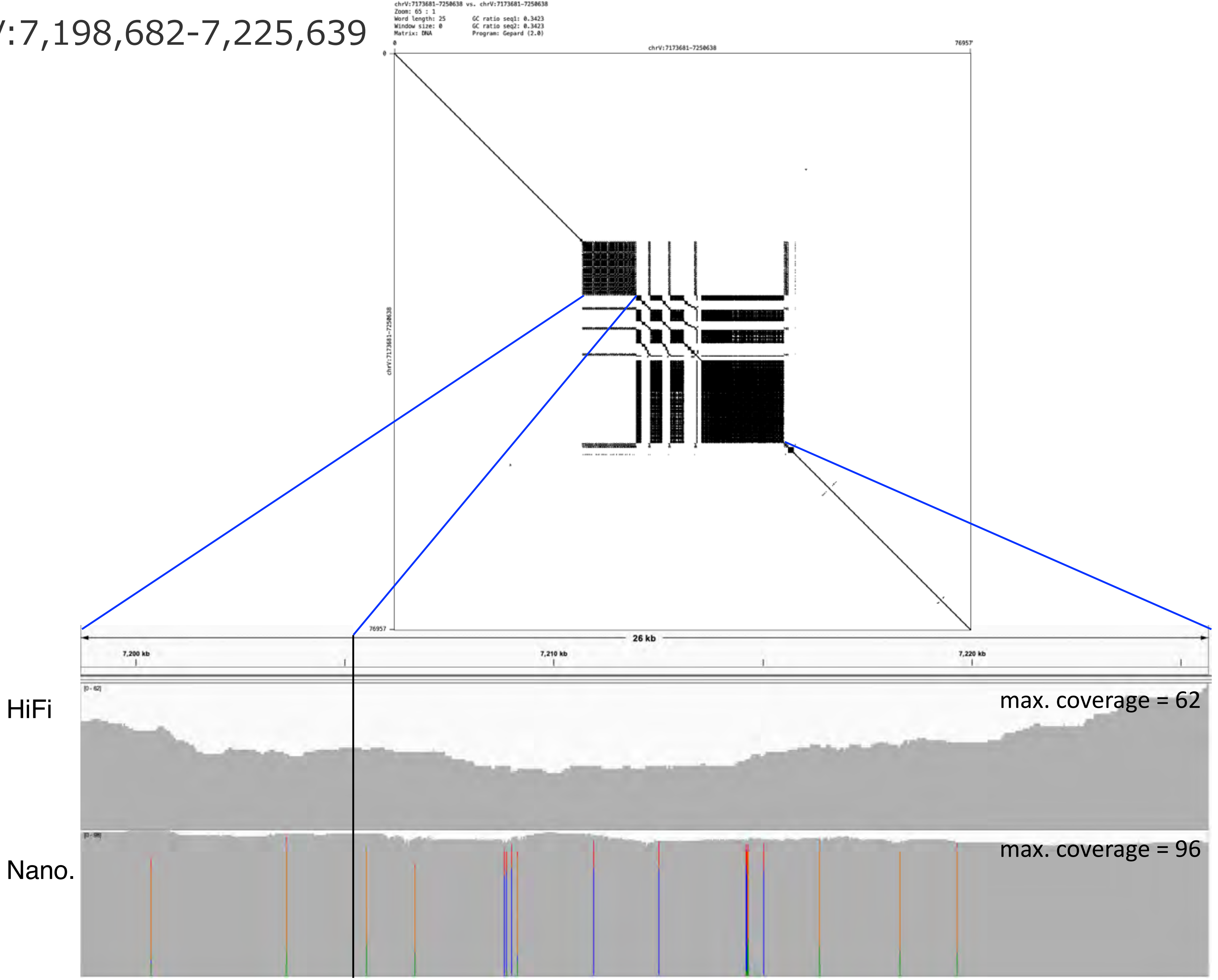

### chrV:8,199,864-8,267,523

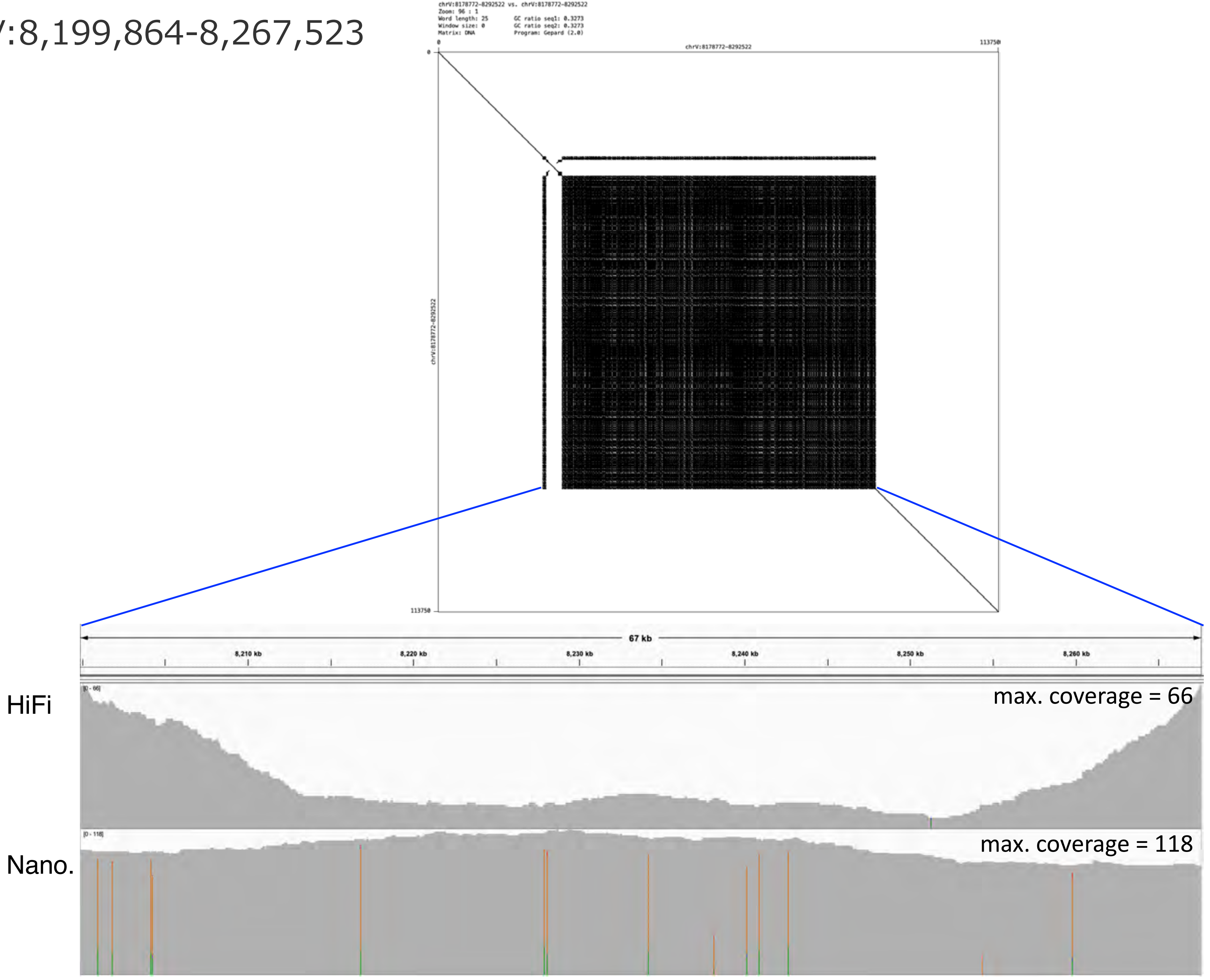

### chrV:8,331,409-8,346,763

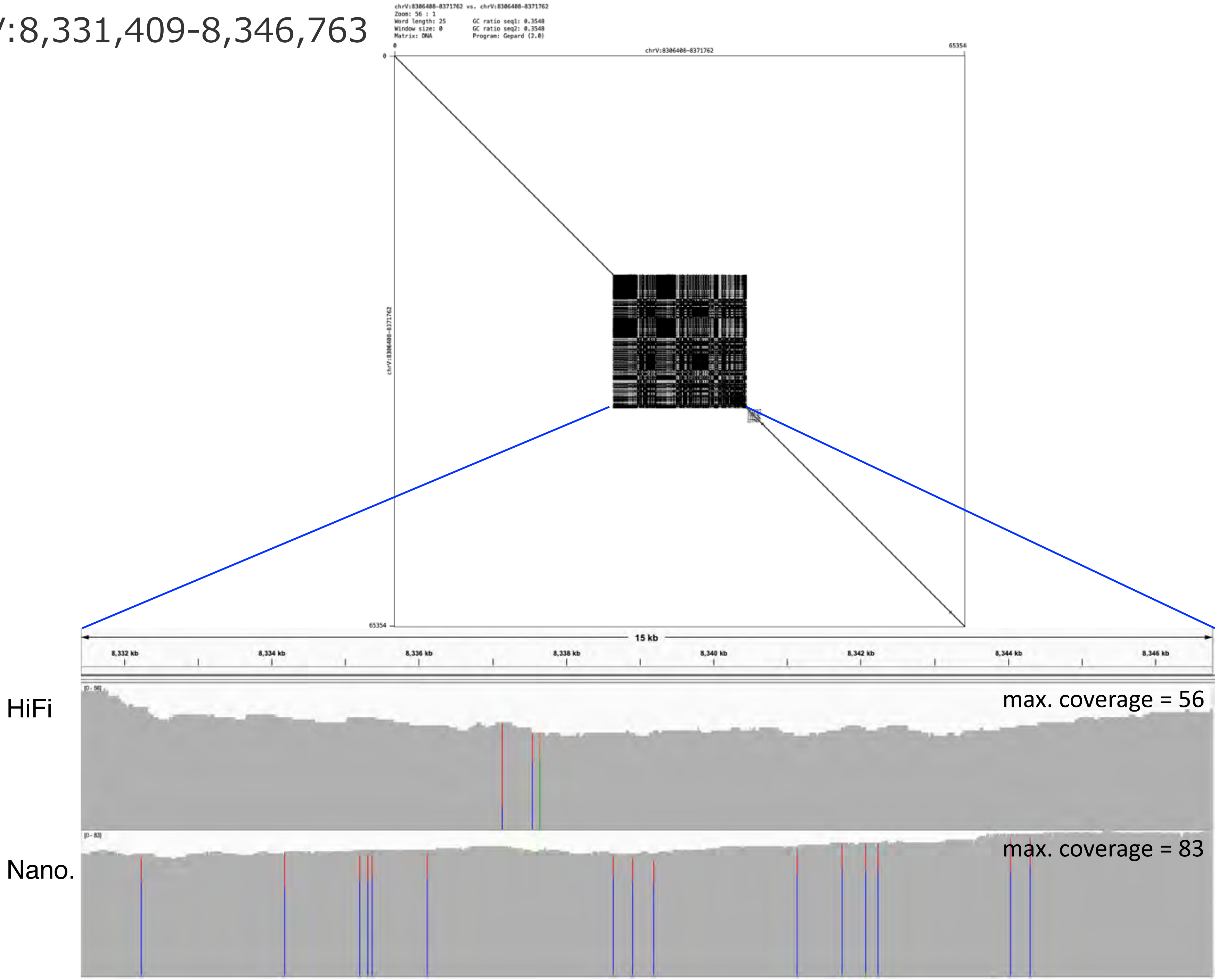

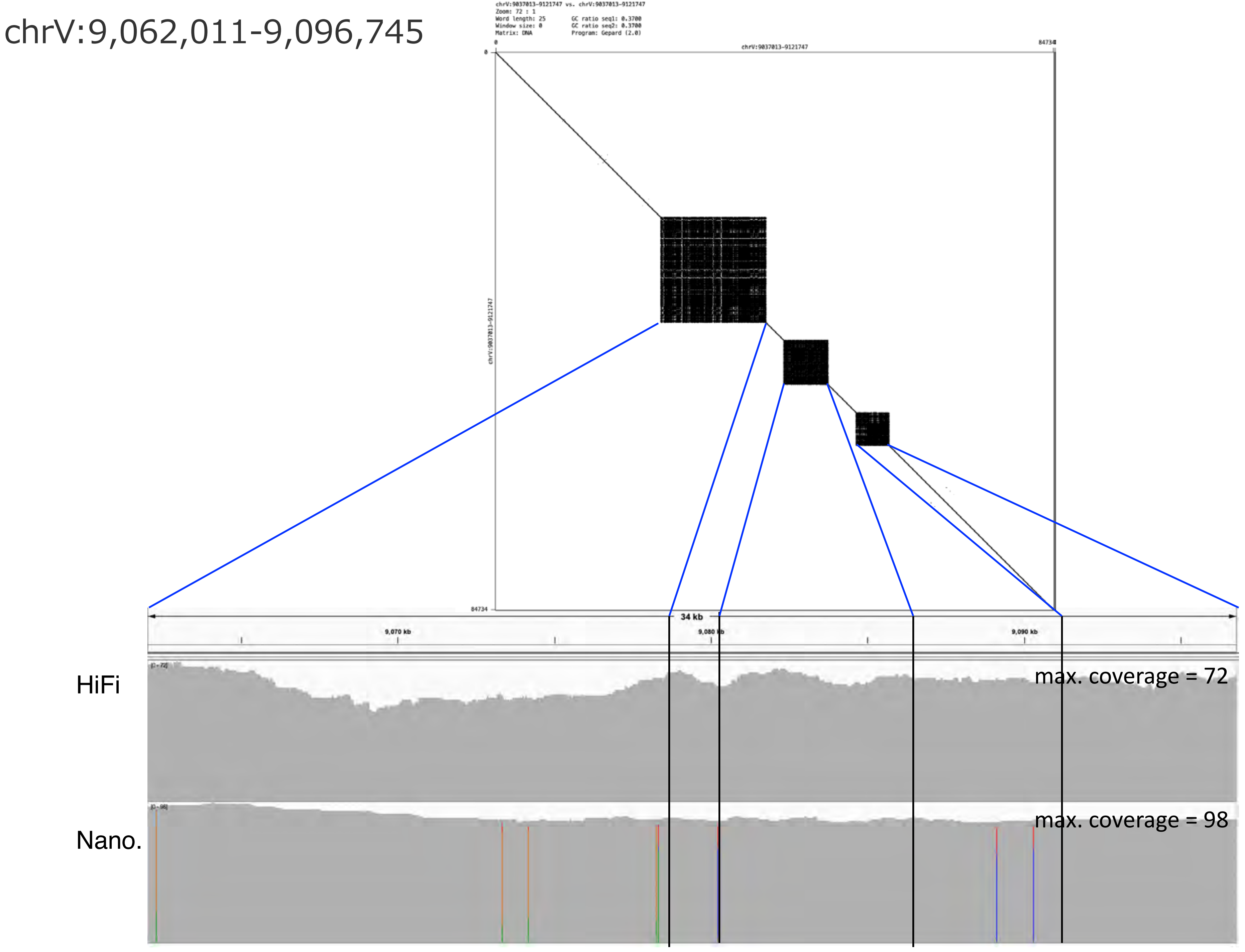
chrV:9,062,011-9,096,745
chrV:9037013-9121747 vs. chrV:9037013-9121747
Zoom: 72 : 1
Word length: 25
GC ratio seq1: 0.3700
Window size: 0
GC ratio seq2: 0.3700
Matrix: DNA
Program: Gepard (2.0)
0
chrV:9037013-9121747
84734
chrV:9037013-9121747
34 kb
9,070 kb
9,080 kb
9,090 kb
HiFi
max. coverage = 72
Nano.
max. coverage = 98

chrV:9,811,530-9,840,166

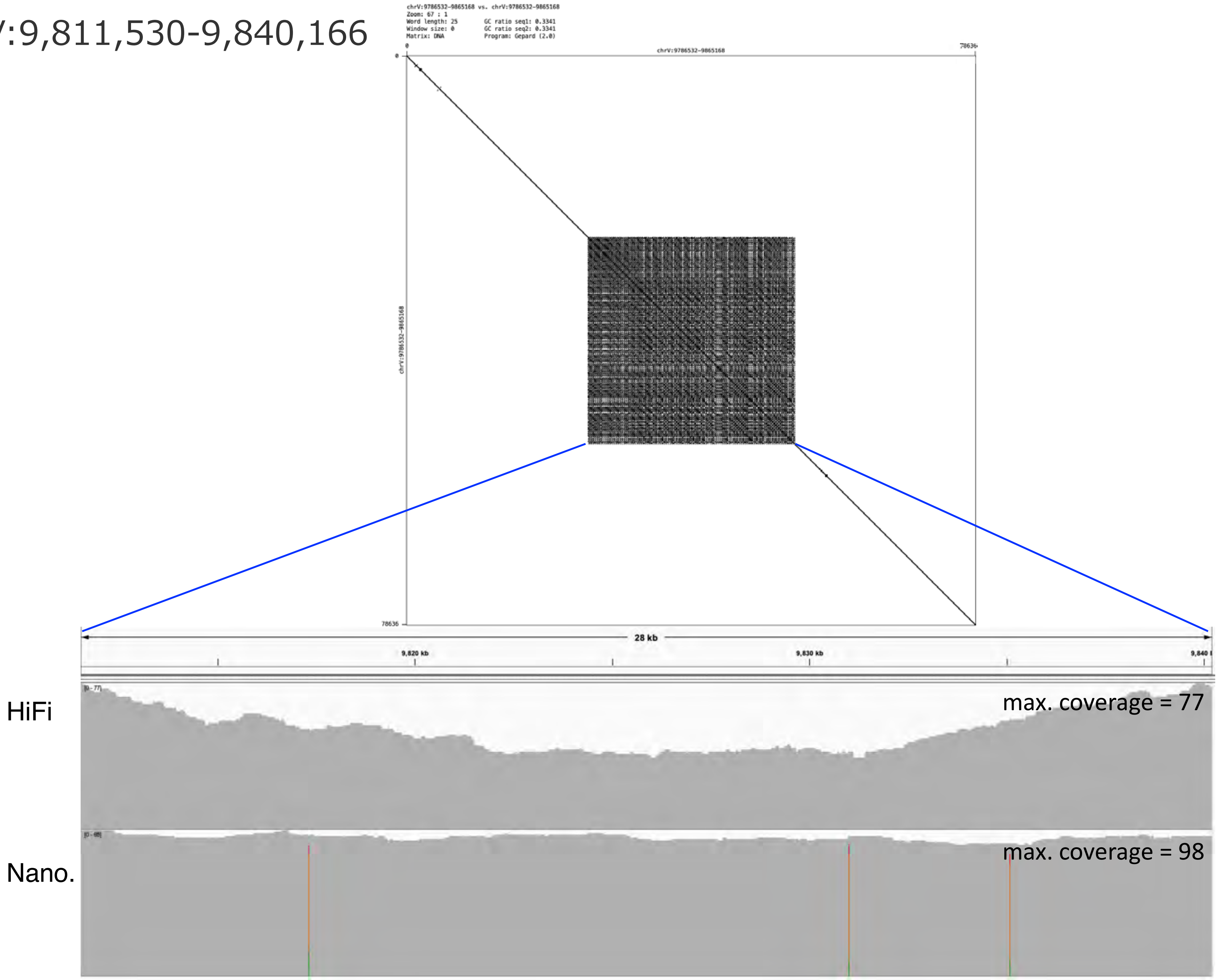

### chrV:11,019,781-11,083,946

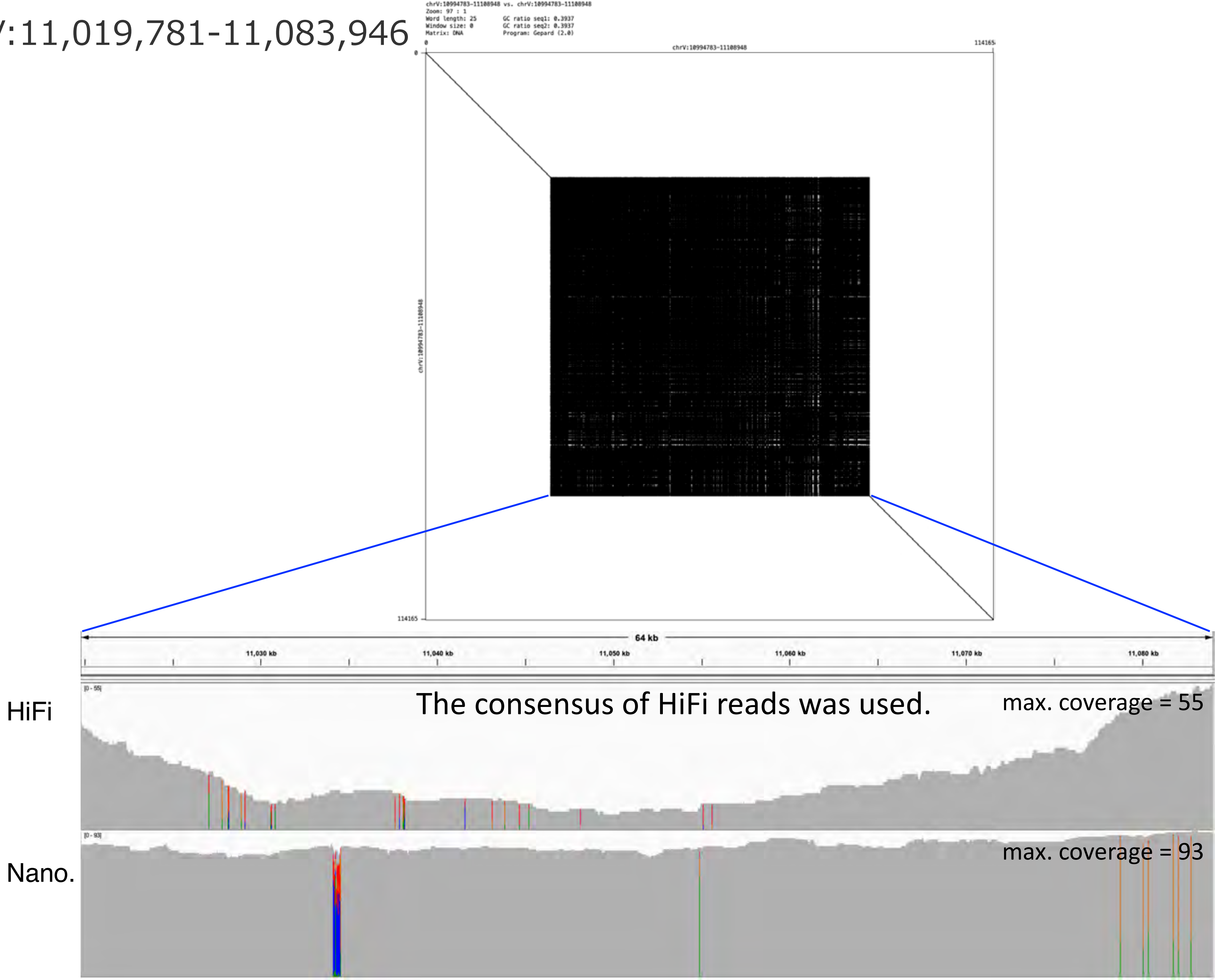

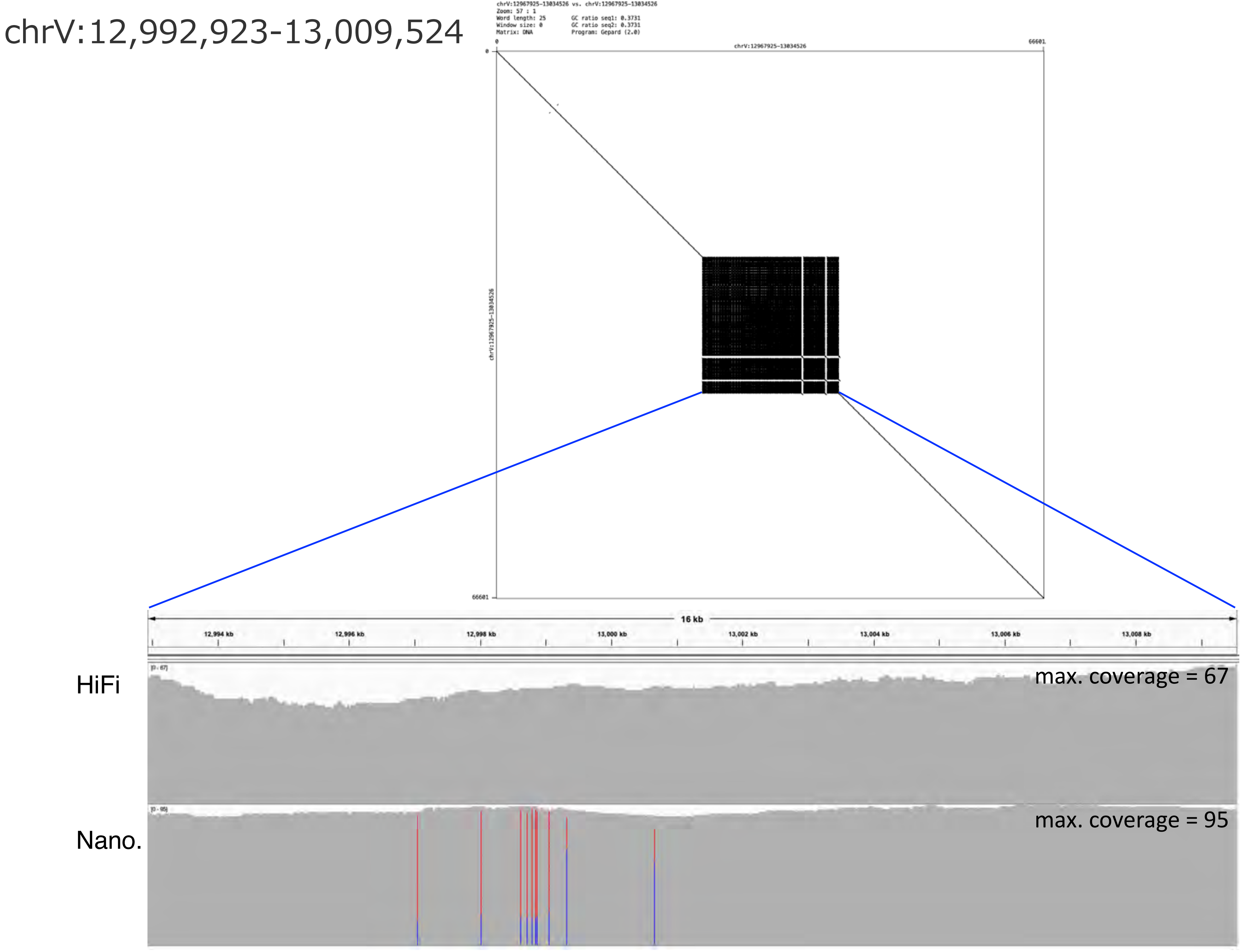

chrV:12,992,923-13,009,524
chrV:12967925-13034526 vs. chrV:12967925-13034526
Zoom: 57 : 1
Word length: 25
Window size: 0
Matrix: DNA
GC ratio seq1: 0.3731
GC ratio seq2: 0.3731
Program: Gepard (2.0)
0
chrV:12967925-13034526
66601
chrV:12967925-13034526
66601
16 kb
12,994 kb
12,996 kb
12,998 kb
13,000 kb
13,002 kb
13,004 kb
13,006 kb
13,008 kb
HiFi
[0 - 67]
max. coverage = 67
Nano.
[0 - 95]
max. coverage = 95

### chrV:15,100,929-15,110,476

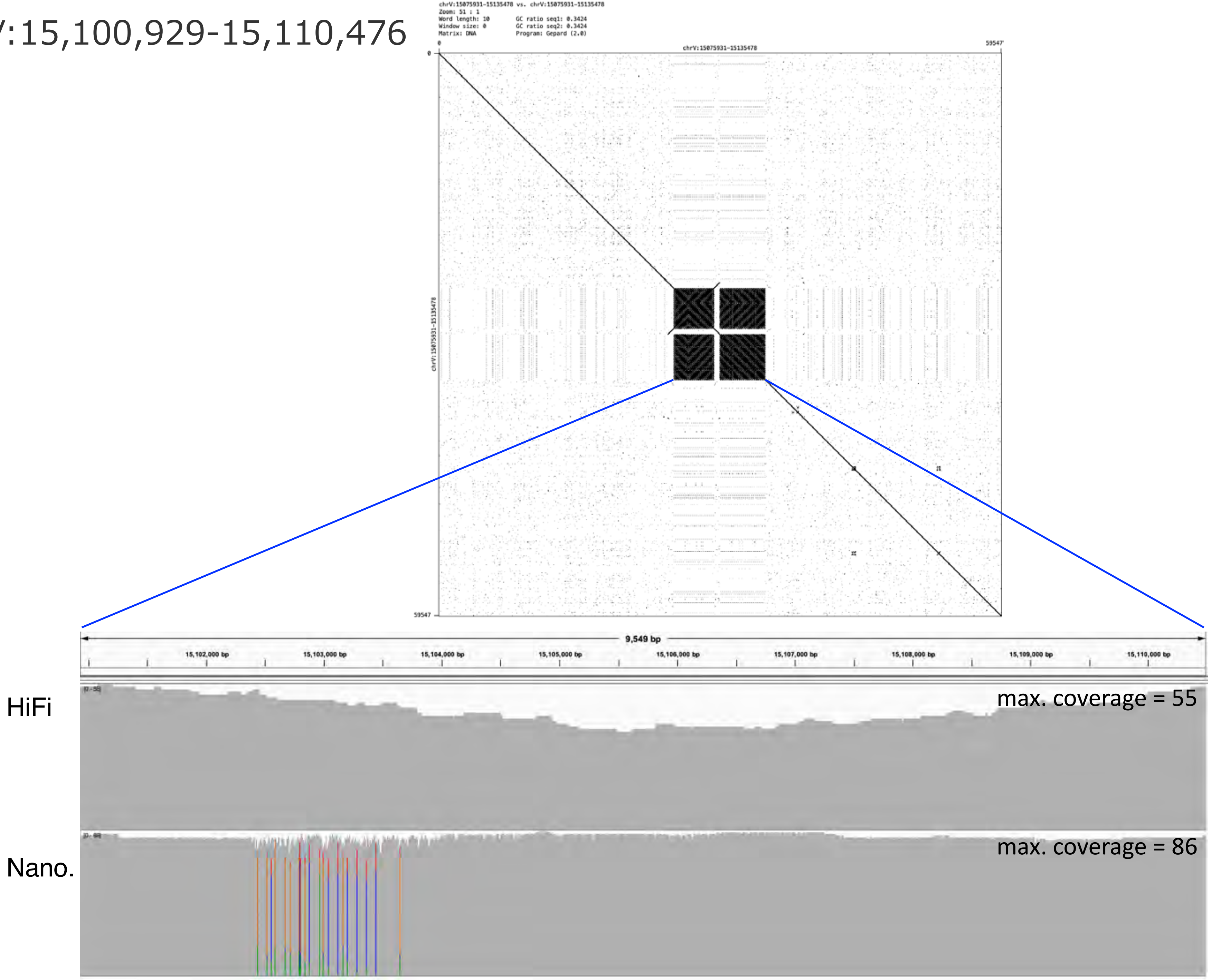

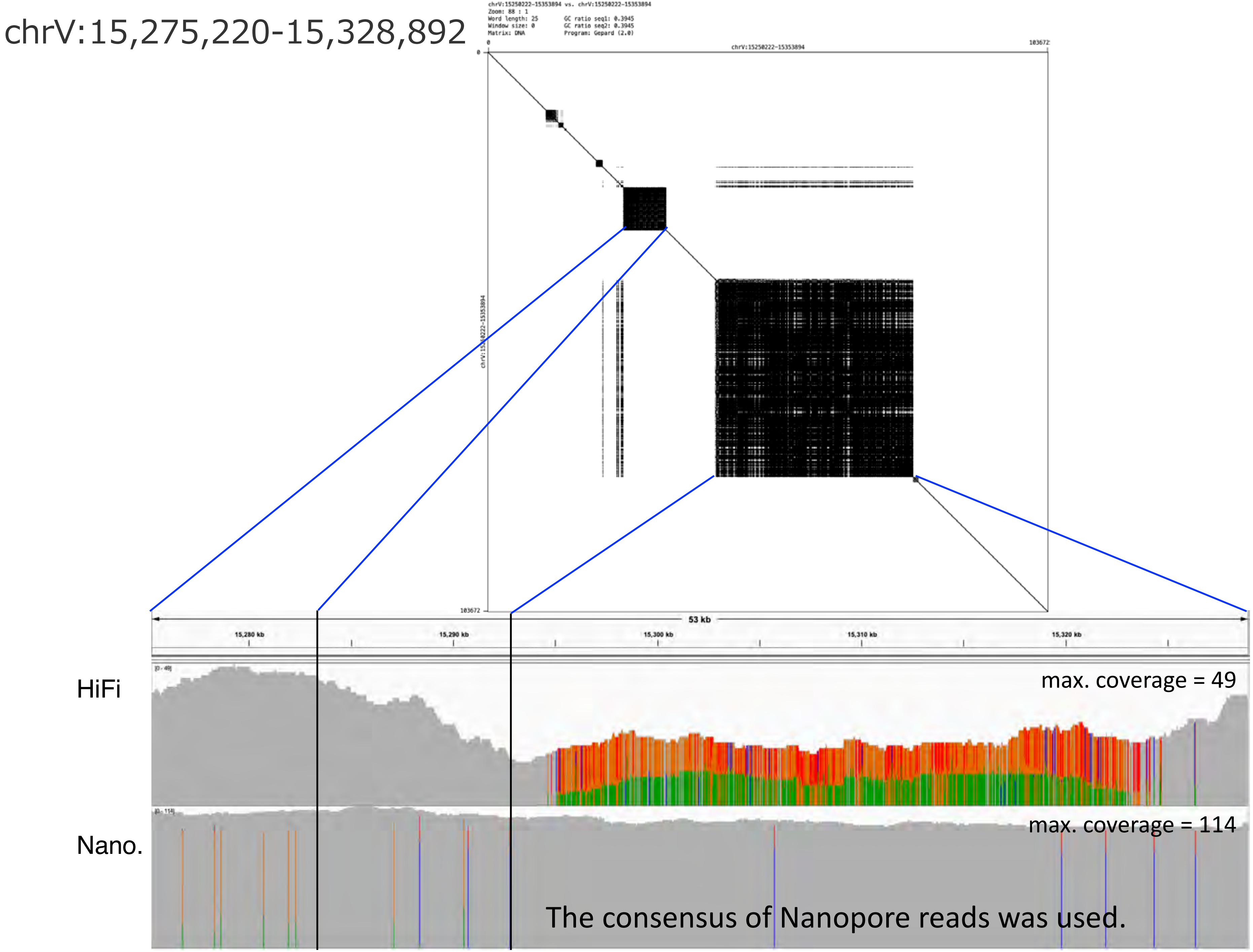

chrV:15,275,220-15,328,892
chrV:15250222-15353894 vs. chrV:15250222-15353894
Zoom: 88 : 1
Word length: 25
Window size: 0
Matrix: DNA
GC ratio seq1: 0.3945
GC ratio seq2: 0.3945
Program: Gepard (2.0)
0
chrV:15250222-15353894
103672
103672
53 kb
15,280 kb
15,290 kb
15,300 kb
15,310 kb
15,320 kb
HiFi
[0 - 49]
max. coverage = 49
Nano.
[0 - 114]
max. coverage = 114
The consensus of Nanopore reads was used.

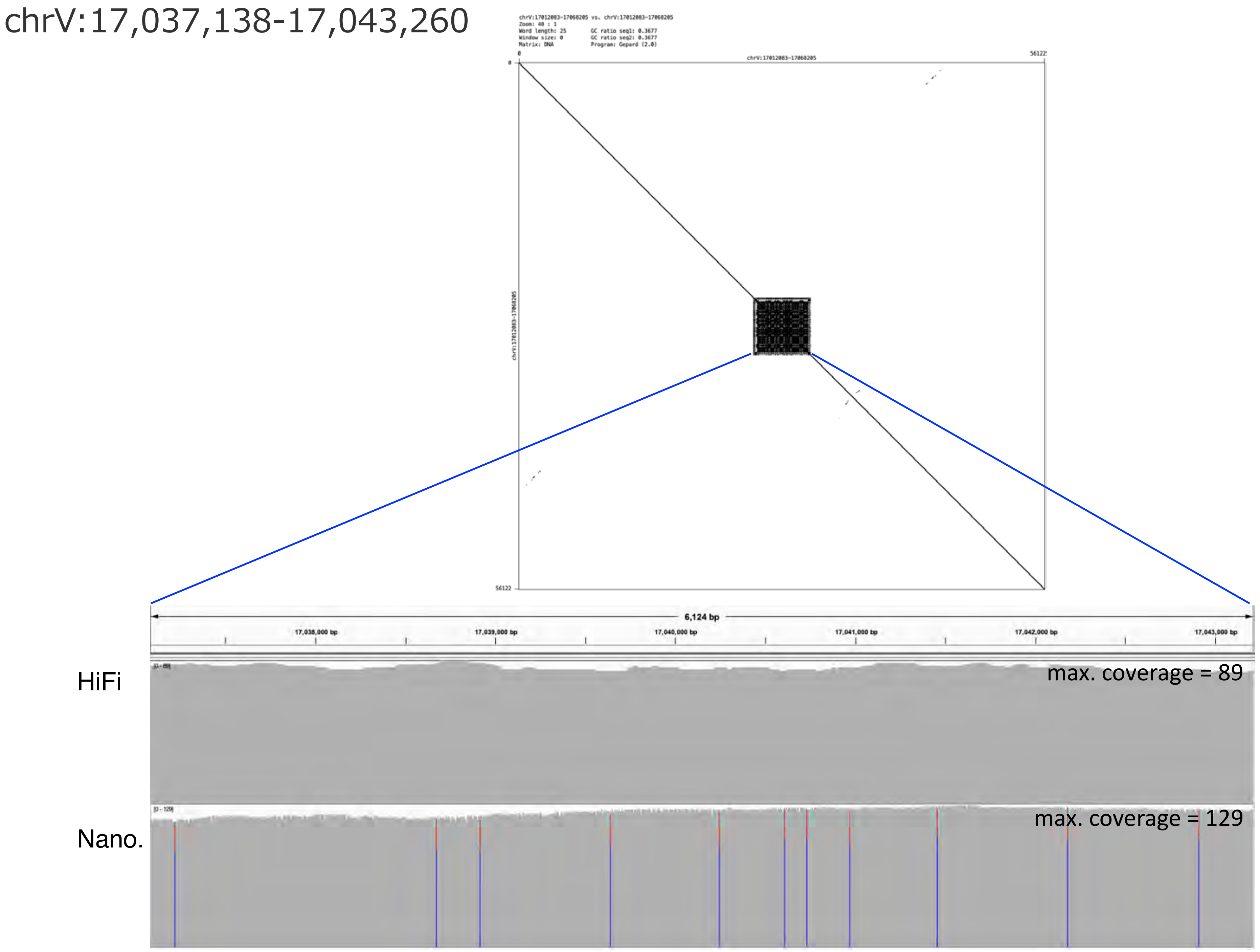
chrV:17,037,138-17,043,260
chrV:17012083-17068205 vs. chrV:17012083-17068205
Zoom: 48 : 1
Word length: 25
Window size: 0
Matrix: DNA
GC ratio seq1: 0.3677
GC ratio seq2: 0.3677
Program: Gepard (2.0)
chrV:17012083-17068205
0
56122
6,124 bp
17,038,000 bp
17,039,000 bp
17,040,000 bp
17,041,000 bp
17,042,000 bp
17,043,000 bp
HiFi
max. coverage = 89
Nano.
max. coverage = 129

chrV:17,299,125-17,405,283

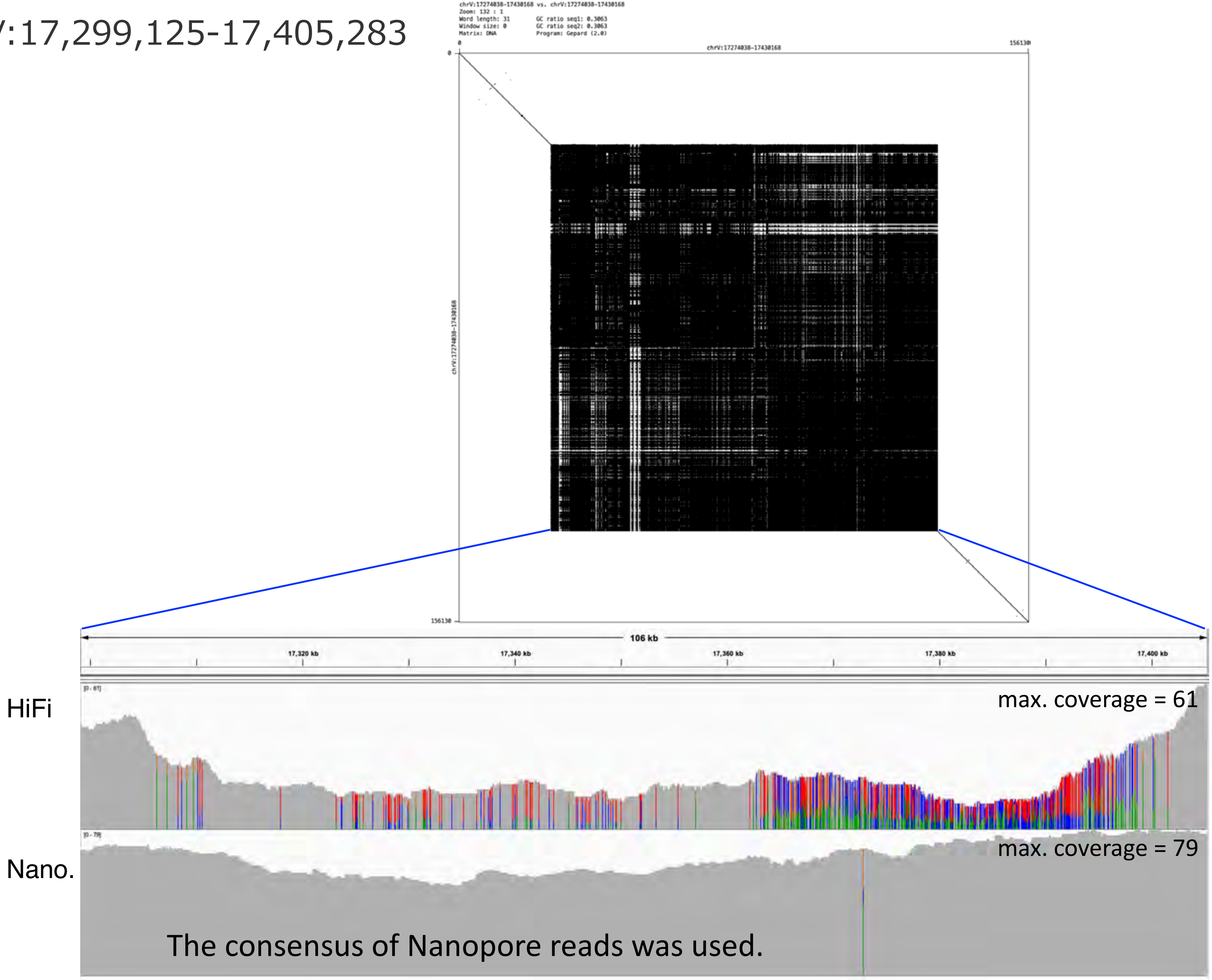

chrV:17,811,675-18,028,889

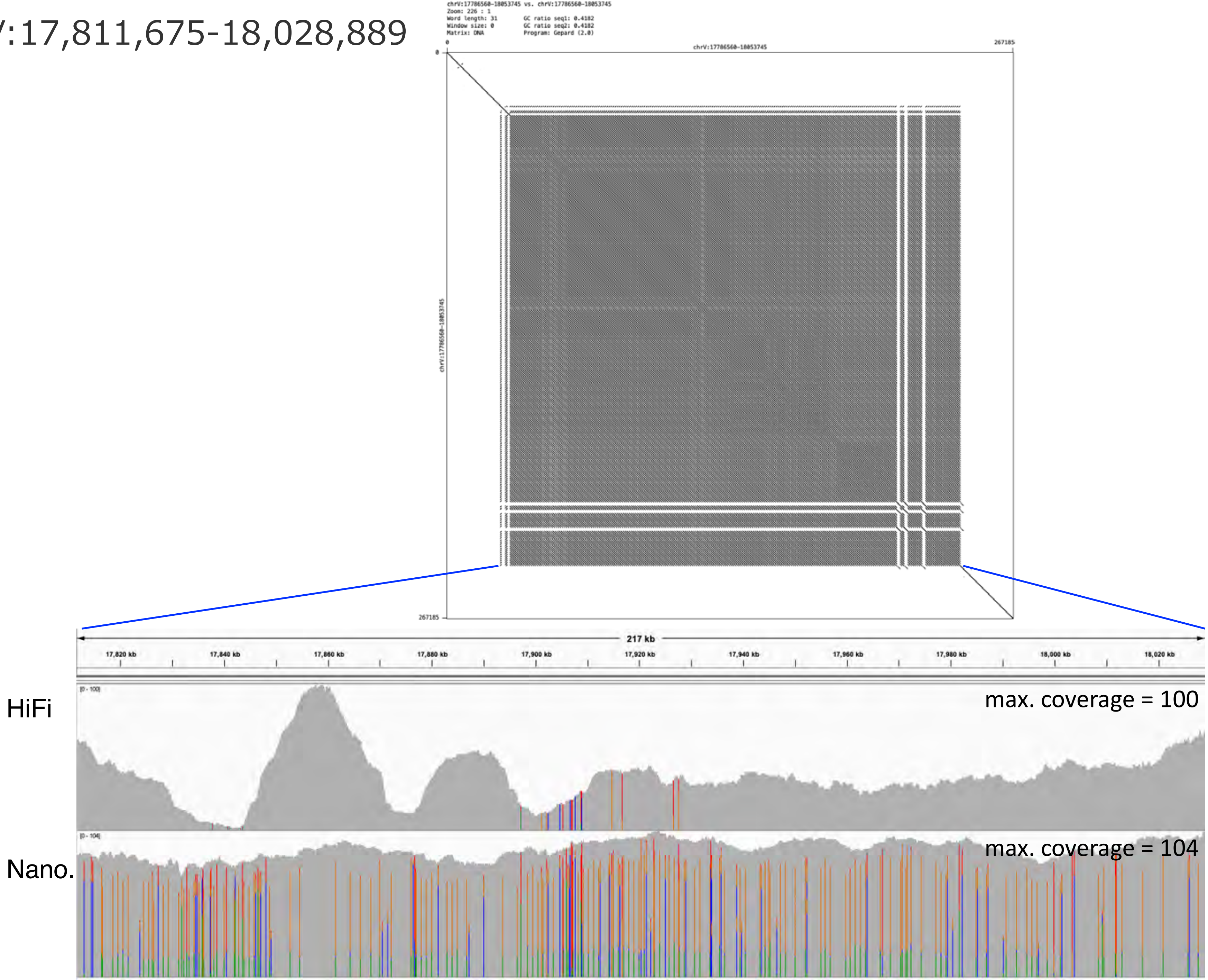

chrV:18,205,300-18,247,161

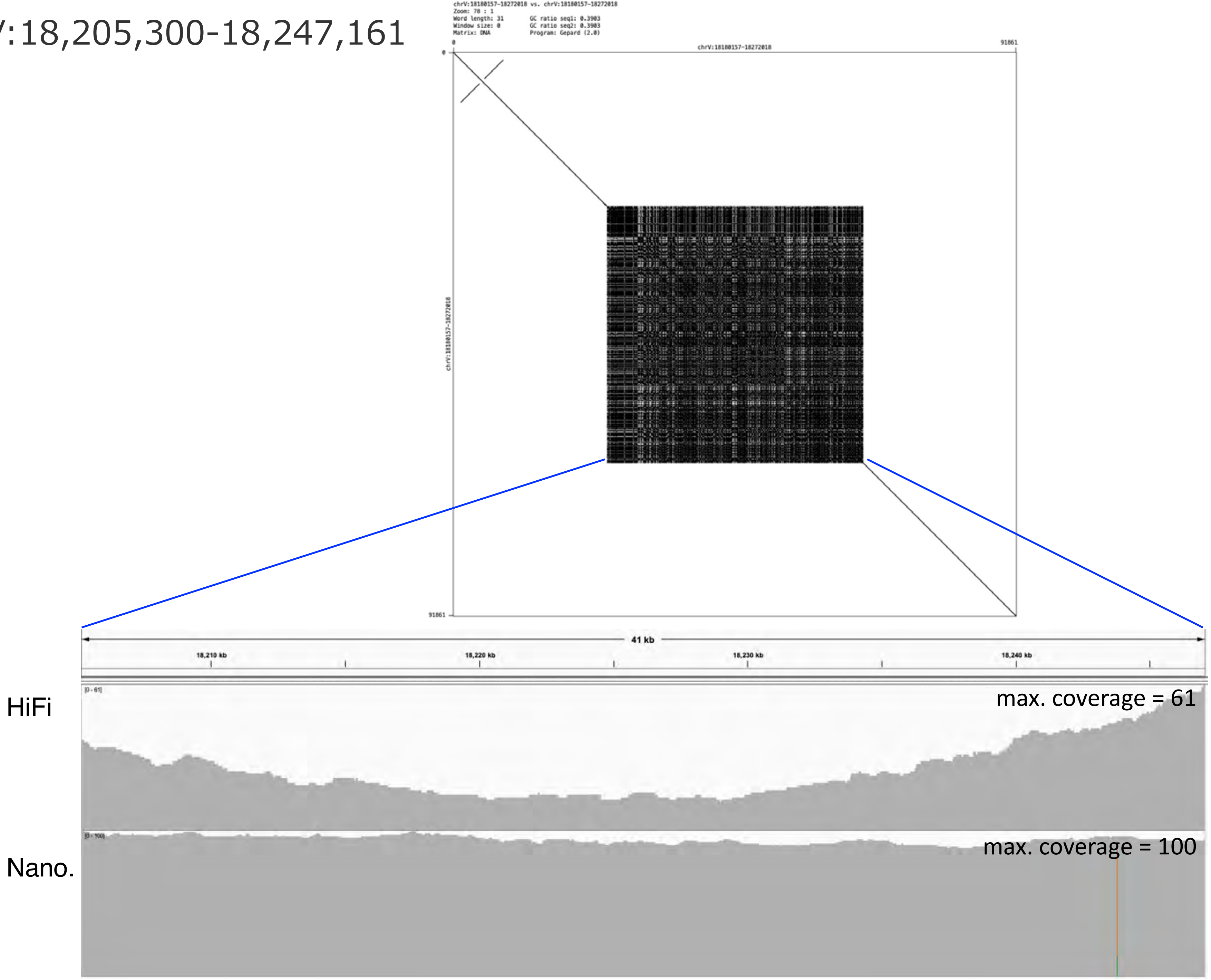

chrV:18,319,740-18,377,175

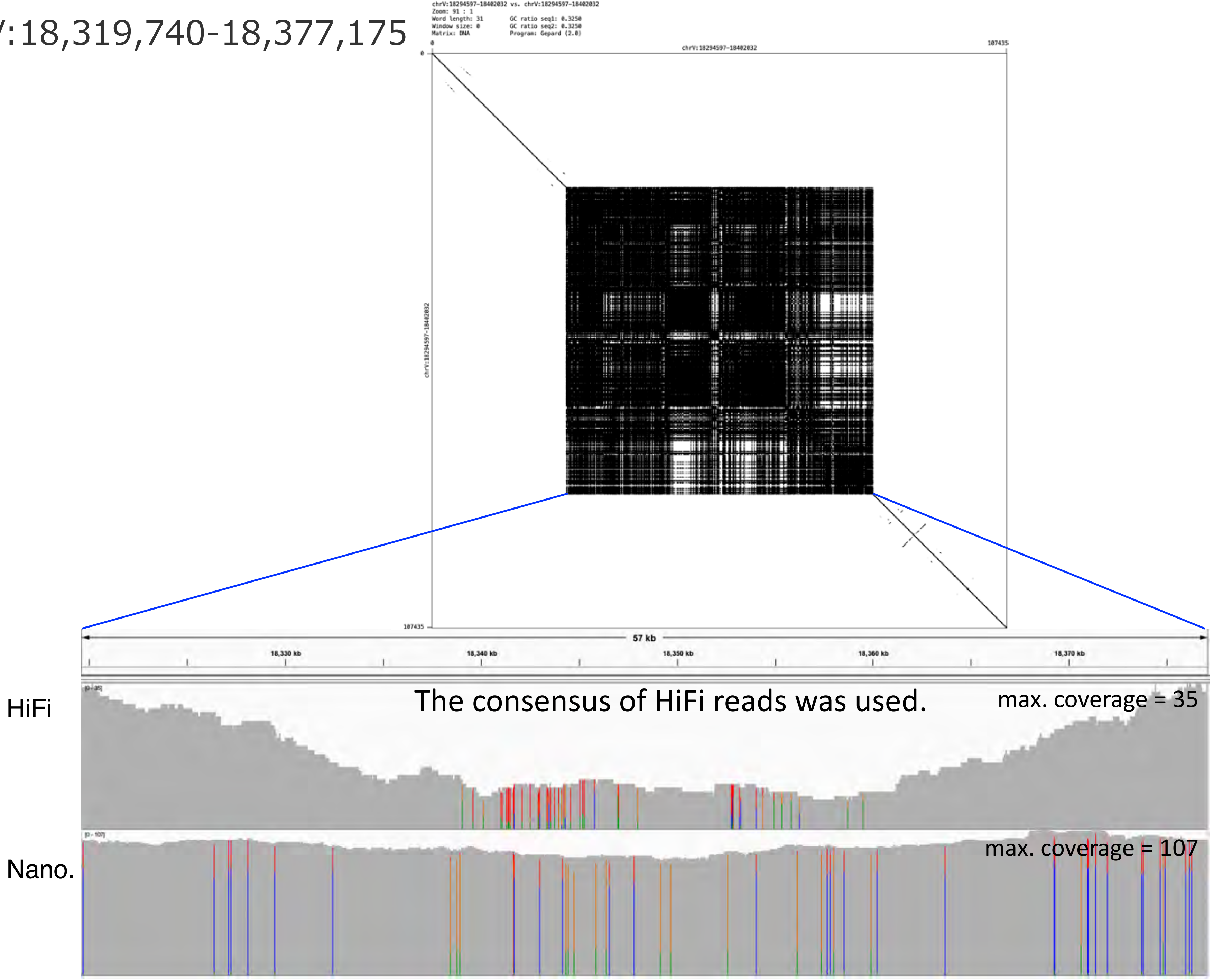

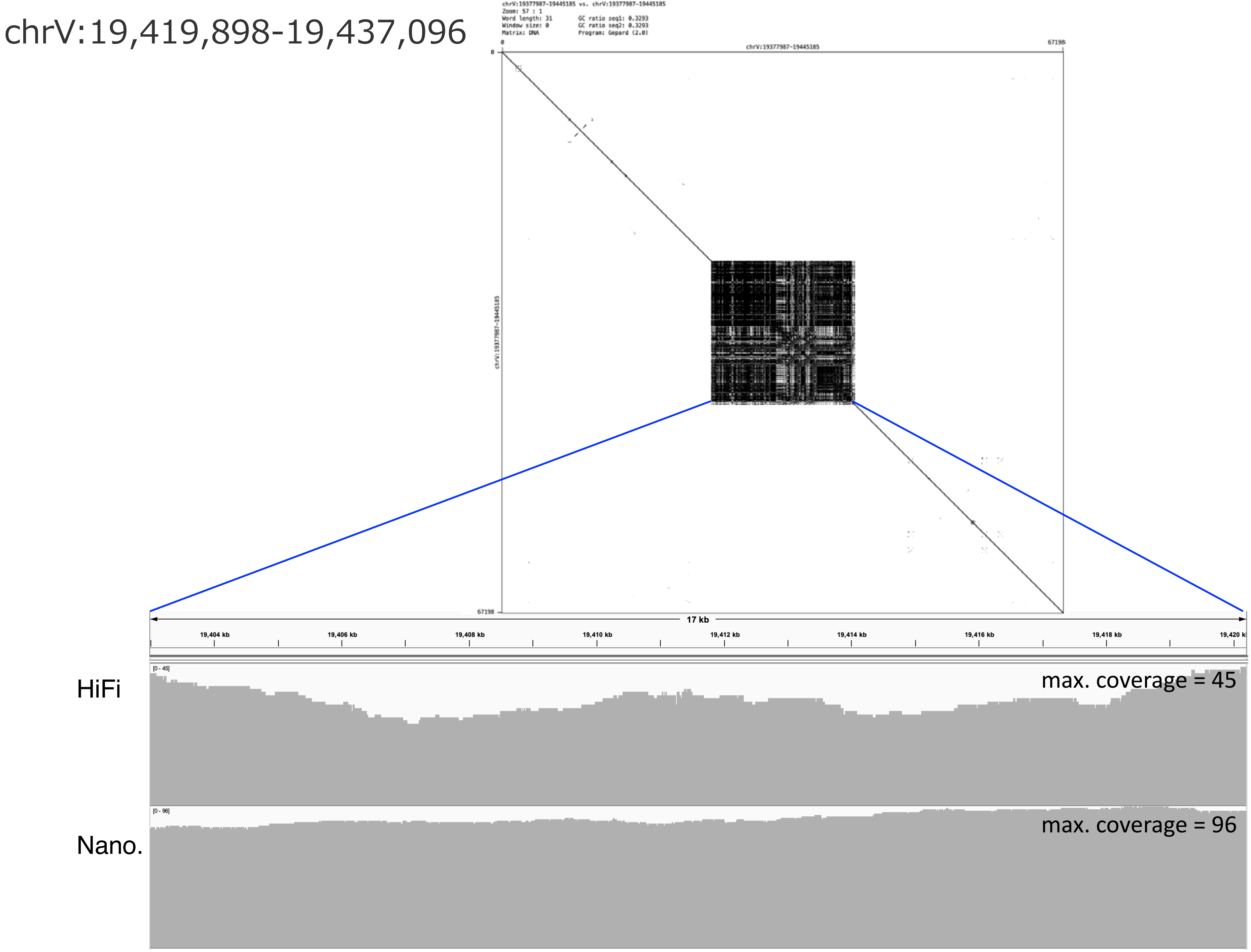
chrV:19,419,898-19,437,096
chrV:19377987-19445185 vs. chrV:19377987-19445185
Zoom: 57 : 1
Word length: 31
Window size: 0
Matrix: DNA
GC ratio seq1: 0.3293
GC ratio seq2: 0.3293
Program: Gepard (2.0)
0
67198
chrV:19377987-19445185
0
chrV:19377987-19445185
67198
17 kb
19,404 kb
19,406 kb
19,408 kb
19,410 kb
19,412 kb
19,414 kb
19,416 kb
19,418 kb
HiFi
[0 - 45]
max. coverage = 45
Nano.
[0 - 96]
max. coverage = 96

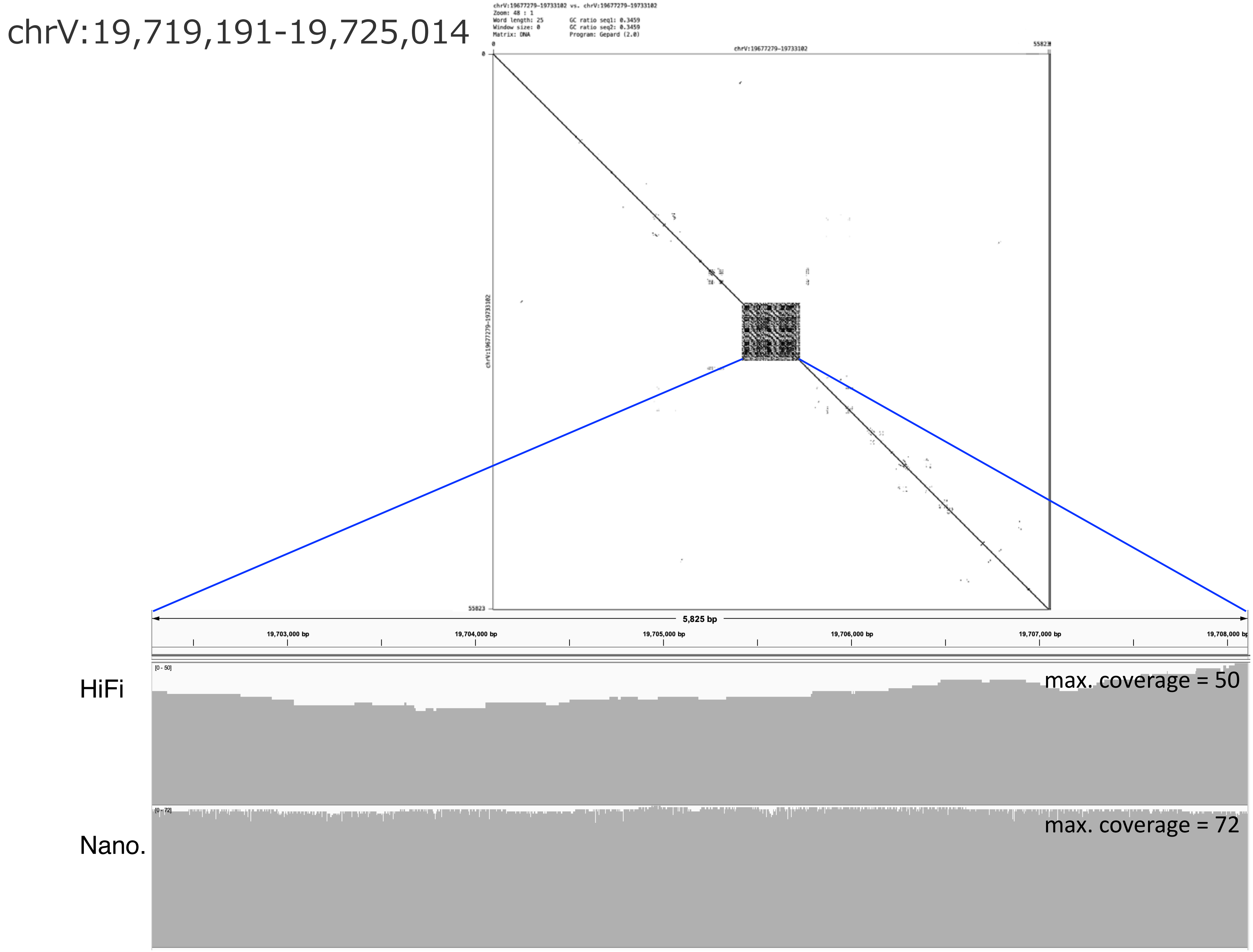

chrV:19,719,191-19,725,014
chrV:19677279-19733102 vs. chrV:19677279-19733102
Zoom: 48 : 1
Word length: 25
Window size: 0
Matrix: DNA
GC ratio seq1: 0.3459
GC ratio seq2: 0.3459
Program: Gepard (2.0)
0
55823
chrV:19677279-19733102
0
55823
chrV:19677279-19733102
5,825 bp
19,703,000 bp
19,704,000 bp
19,705,000 bp
19,706,000 bp
19,707,000 bp
19,708,000 bp
HiFi
[0 - 50]
max. coverage = 50
Nano.
[0 - 72]
max. coverage = 72

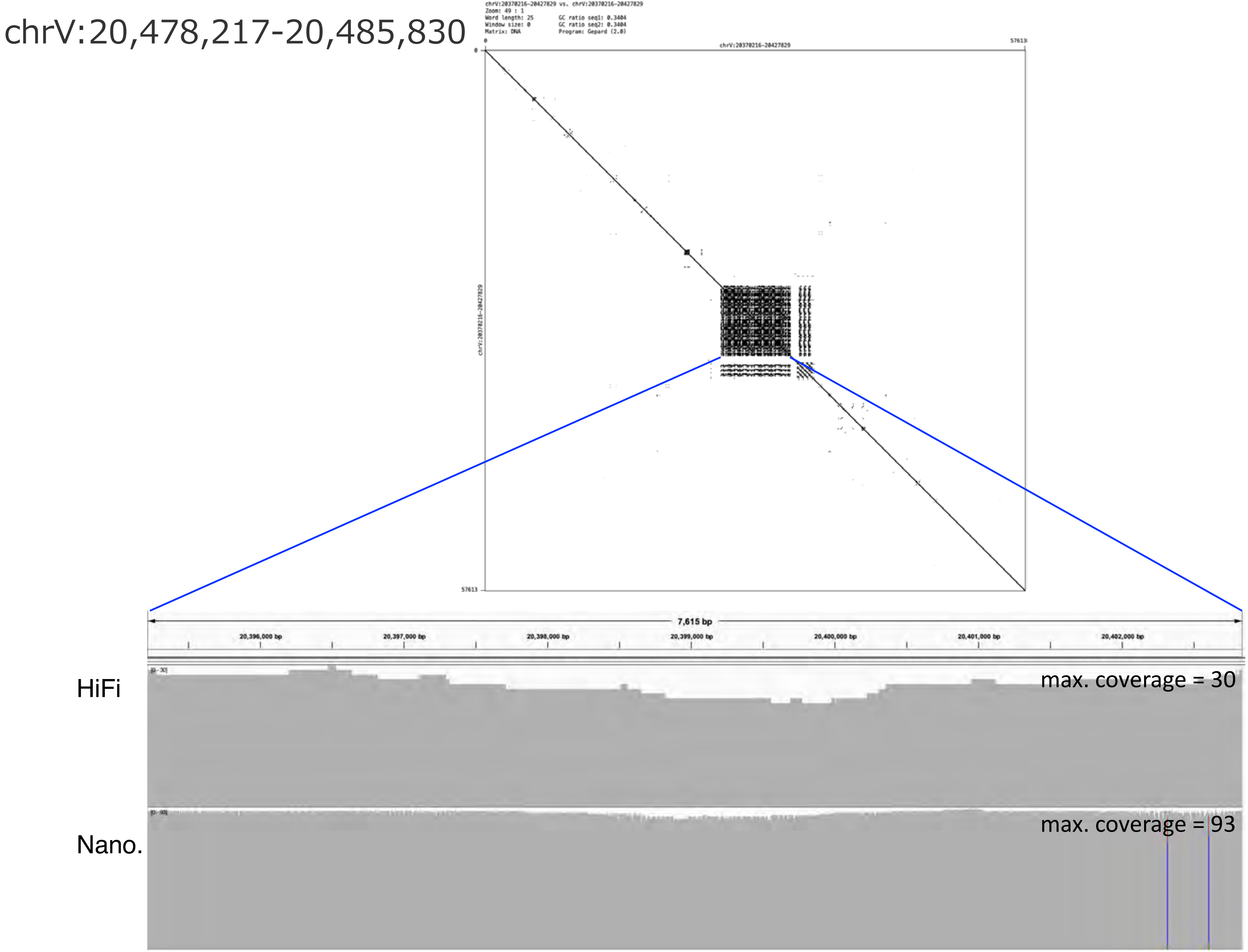
chrV:20,478,217-20,485,830
chrV:20370216-20427829 vs. chrV:20370216-20427829
Zoom: 49 : 1
Word length: 25
Window size: 0
Matrix: DNA
GC ratio seq1: 0.3404
GC ratio seq2: 0.3404
Program: Gepard (2.0)
chrV:20370216-20427829
0
57613
7,615 bp
20,396,000 bp
20,397,000 bp
20,398,000 bp
20,399,000 bp
20,400,000 bp
20,401,000 bp
20,402,000 bp
HiFi
max. coverage = 30
Nano.
max. coverage = 93

chrV:21,571,705-21,578,725

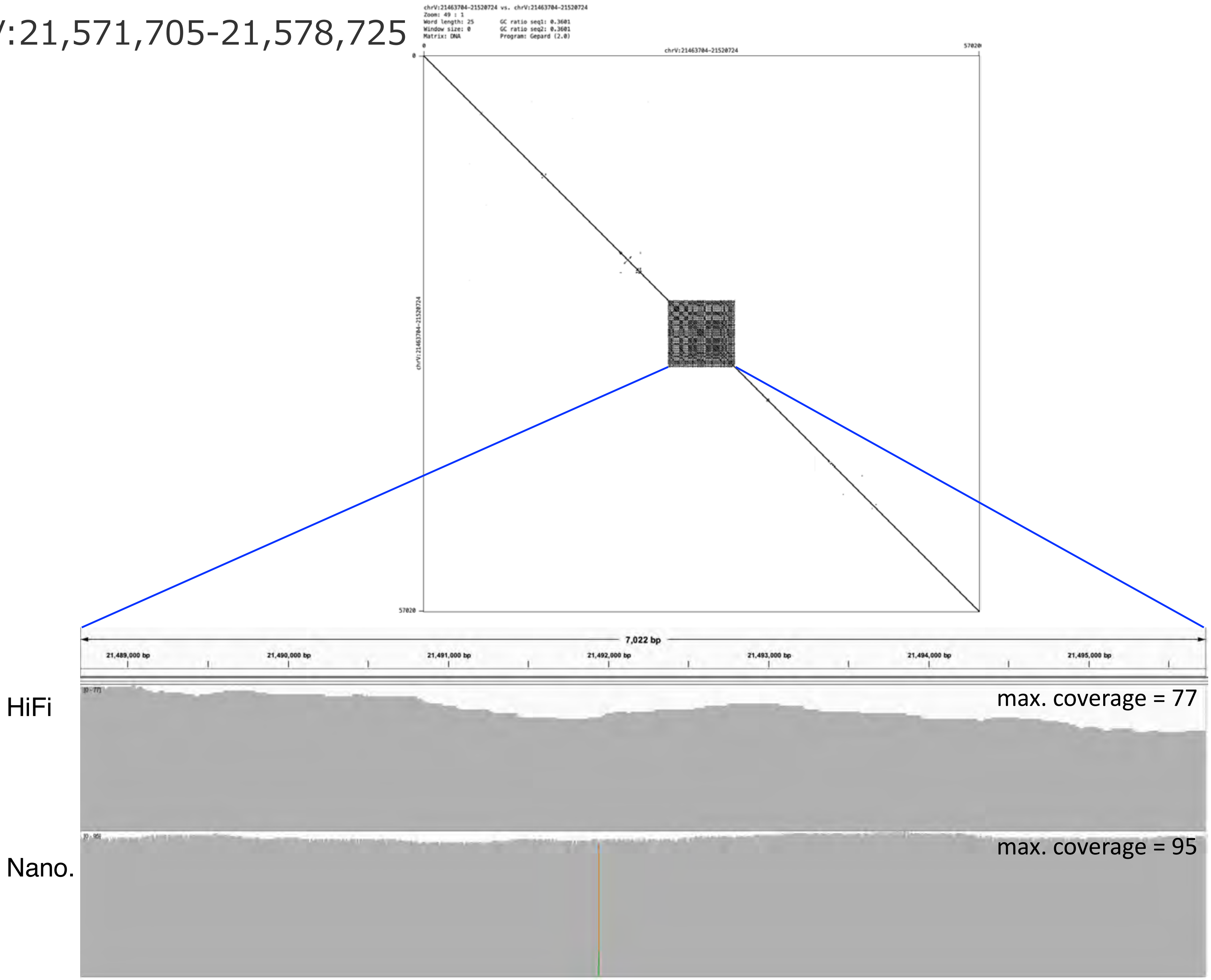

### chrX:122,781-250,556

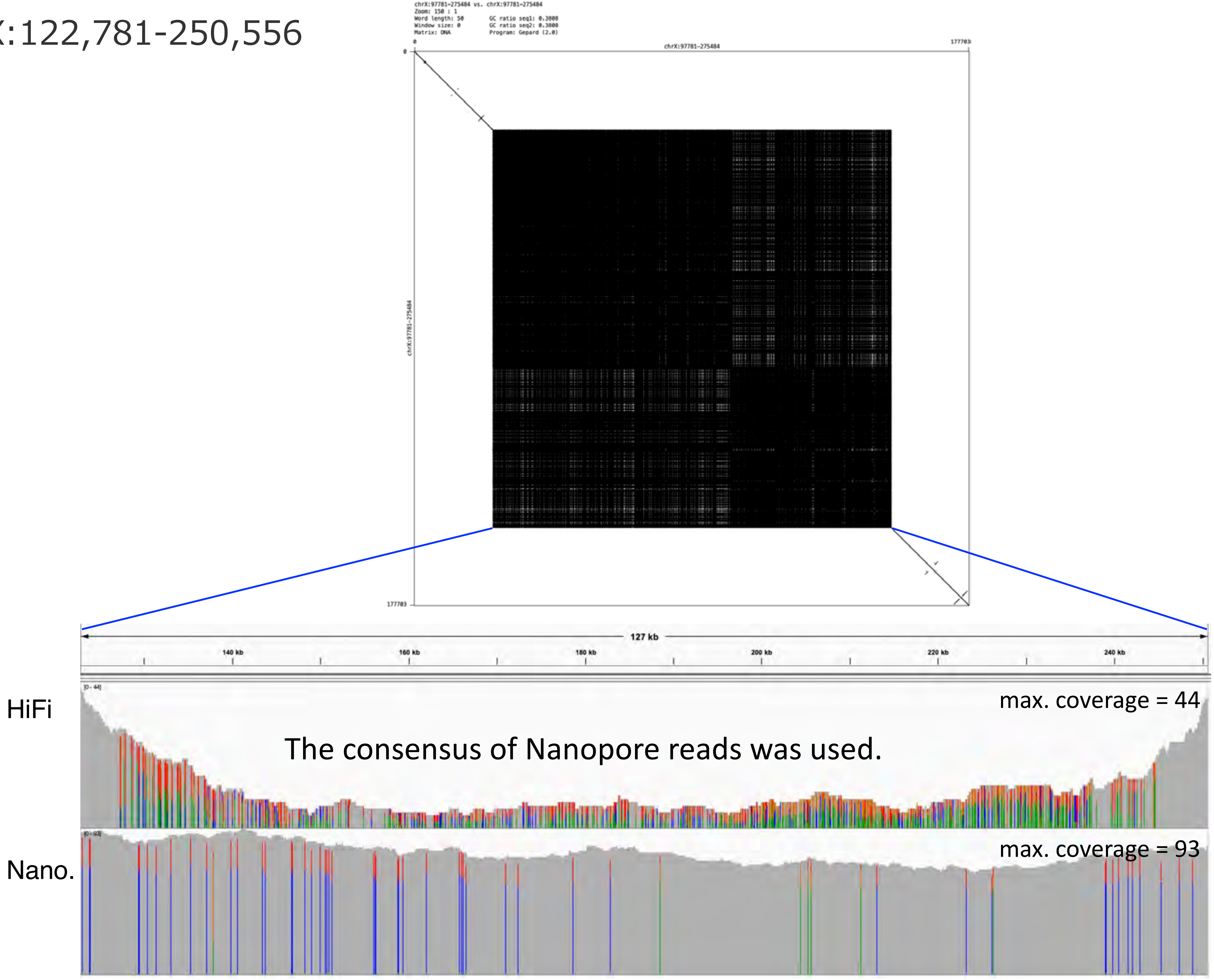

### chrX:427,000-594,551

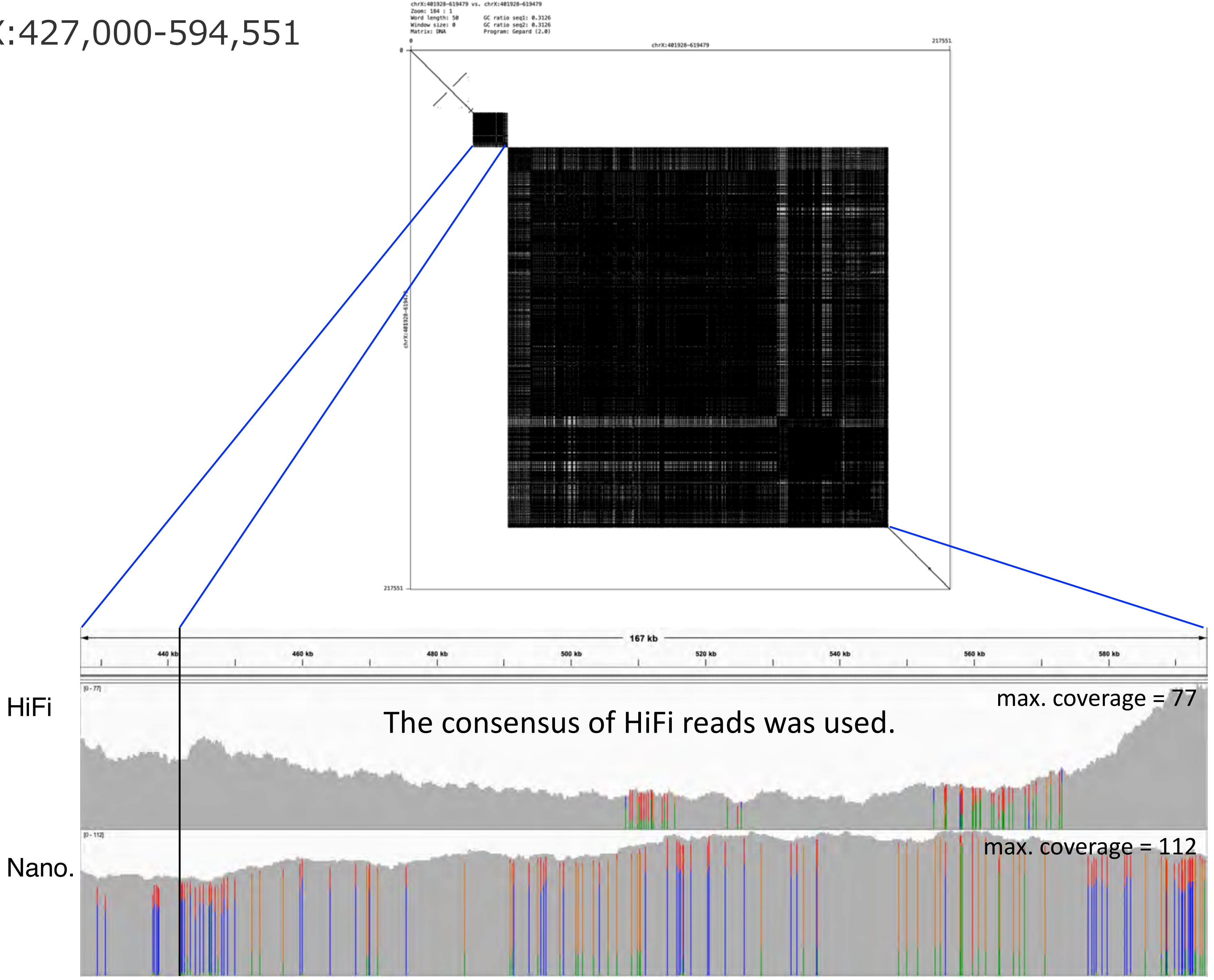

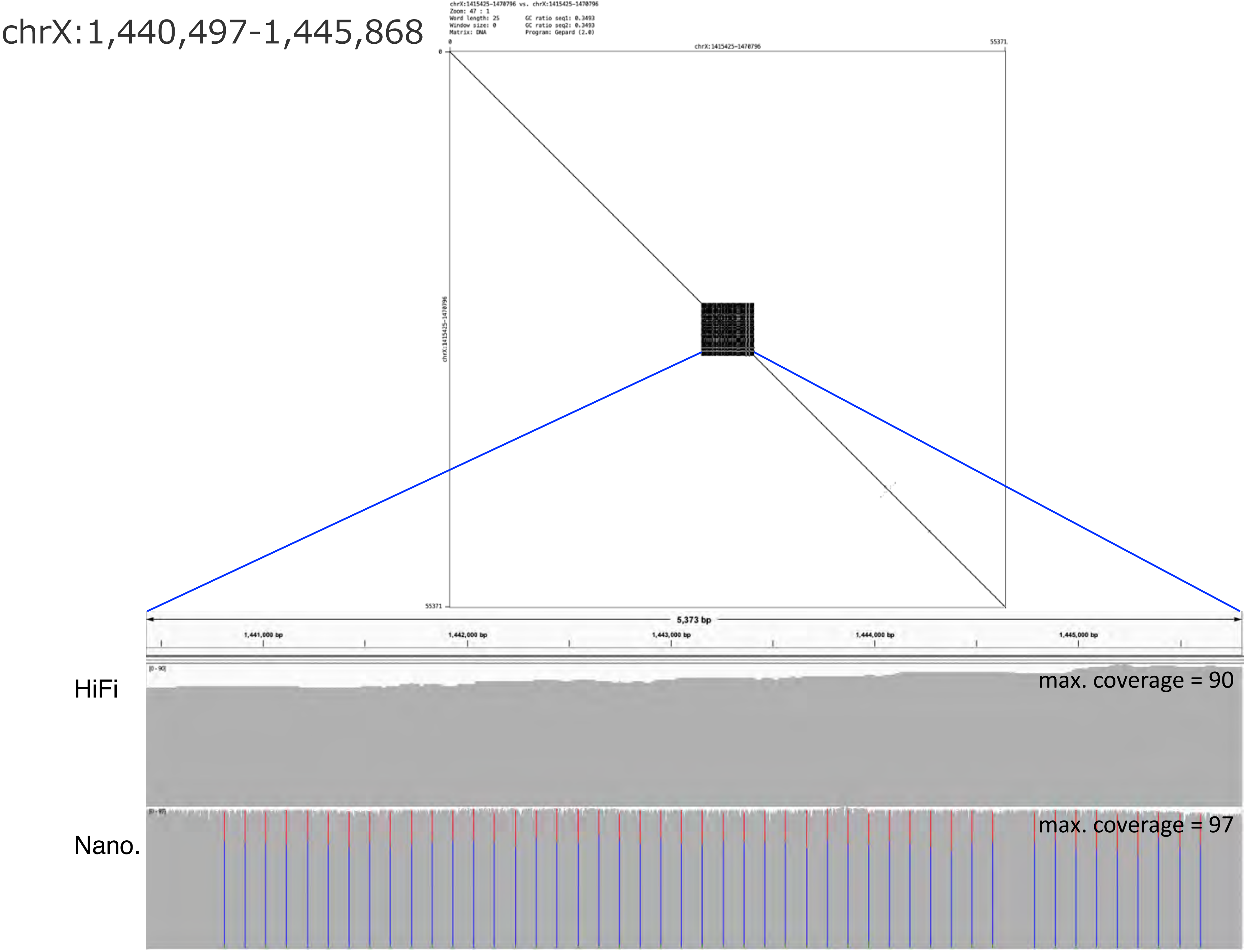
chrX:1,440,497-1,445,868
chrX:1415425-1470796 vs. chrX:1415425-1470796
Zoom: 47 : 1
Word length: 25
Window size: 0
Matrix: DNA
GC ratio seq1: 0.3493
GC ratio seq2: 0.3493
Program: Gepard (2.0)
0
chrX:1415425-1470796
55371
0
chrX:1415425-1470796
55371
5,373 bp
1,441,000 bp
1,442,000 bp
1,443,000 bp
1,444,000 bp
1,445,000 bp
HiFi
max. coverage = 90
Nano.
max. coverage = 97

chrX:1,942,721-1,953,297

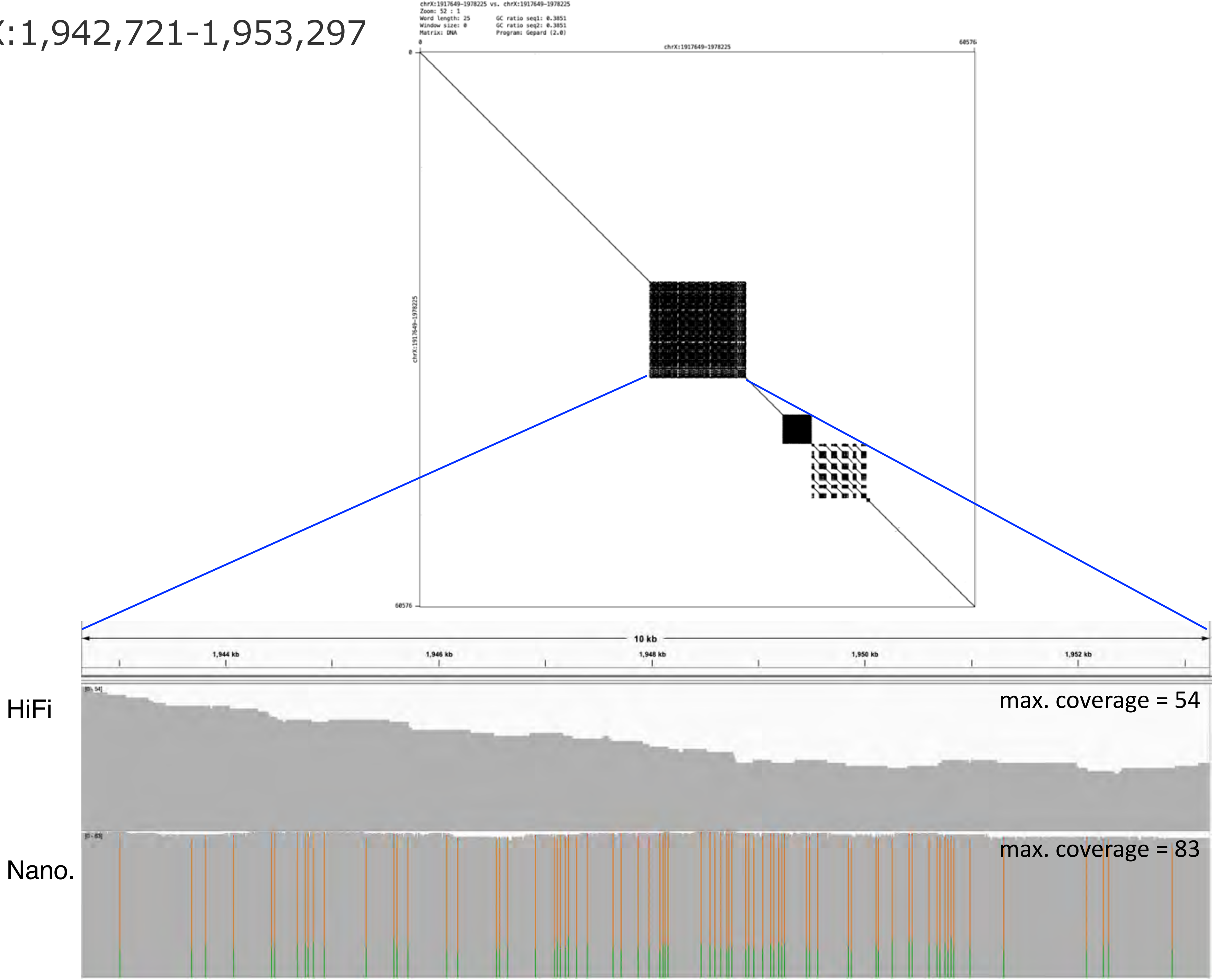

### chrX:2,074,379-2,088,168

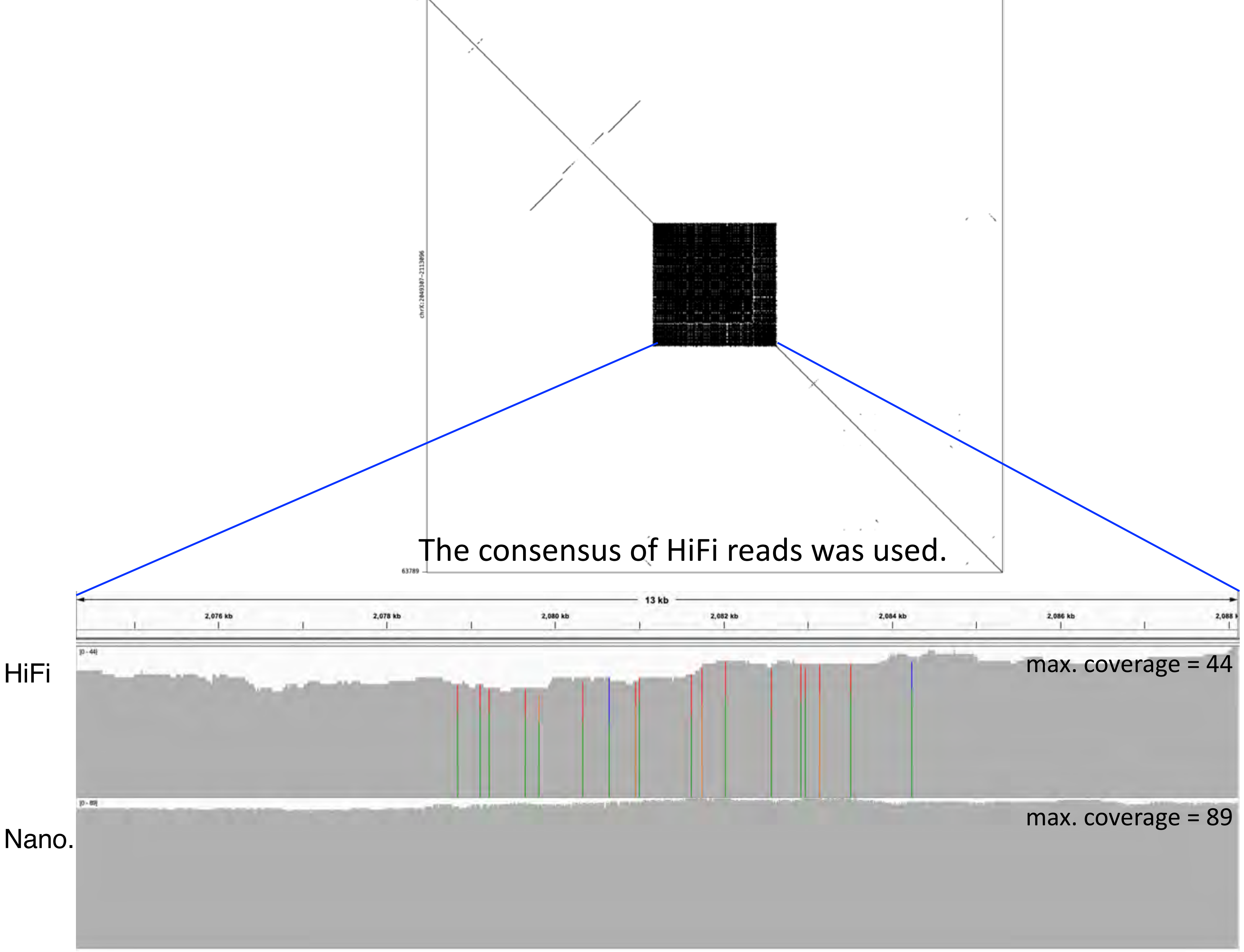

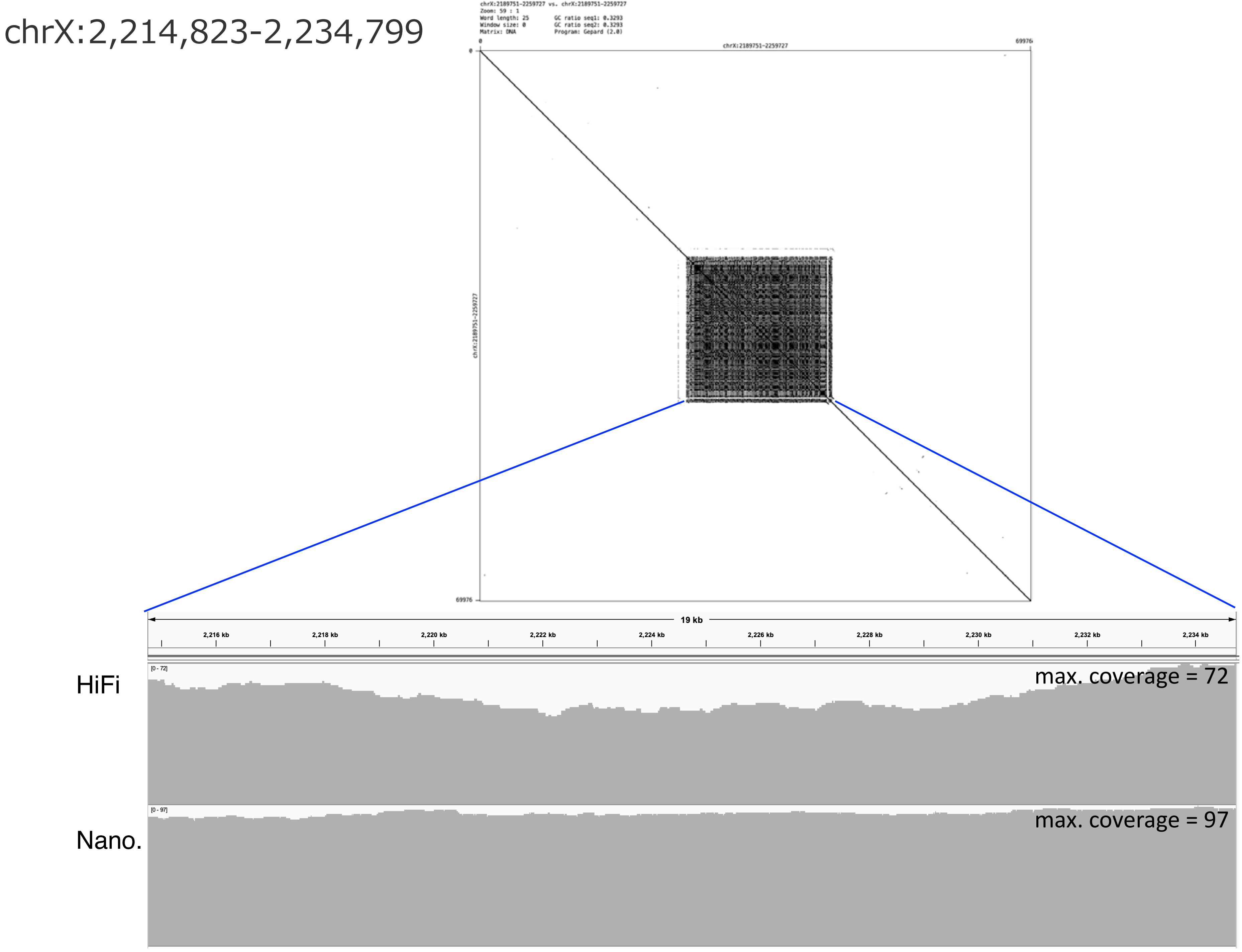
chrX:2,214,823-2,234,799
chrX:2189751-2259727 vs. chrX:2189751-2259727
Zoom: 59 : 1
Word length: 25
Window size: 0
Matrix: DNA
GC ratio seq1: 0.3293
GC ratio seq2: 0.3293
Program: Gepard (2.0)
0
chrX:2189751-2259727
69976
0
chrX:2189751-2259727
69976
19 kb
2,216 kb
2,218 kb
2,220 kb
2,222 kb
2,224 kb
2,226 kb
2,228 kb
2,230 kb
2,232 kb
2,234 kb
HiFi
[0 - 72]
max. coverage = 72
Nano.
[0 - 97]
max. coverage = 97

### chrX:2,371,565-2,420,299

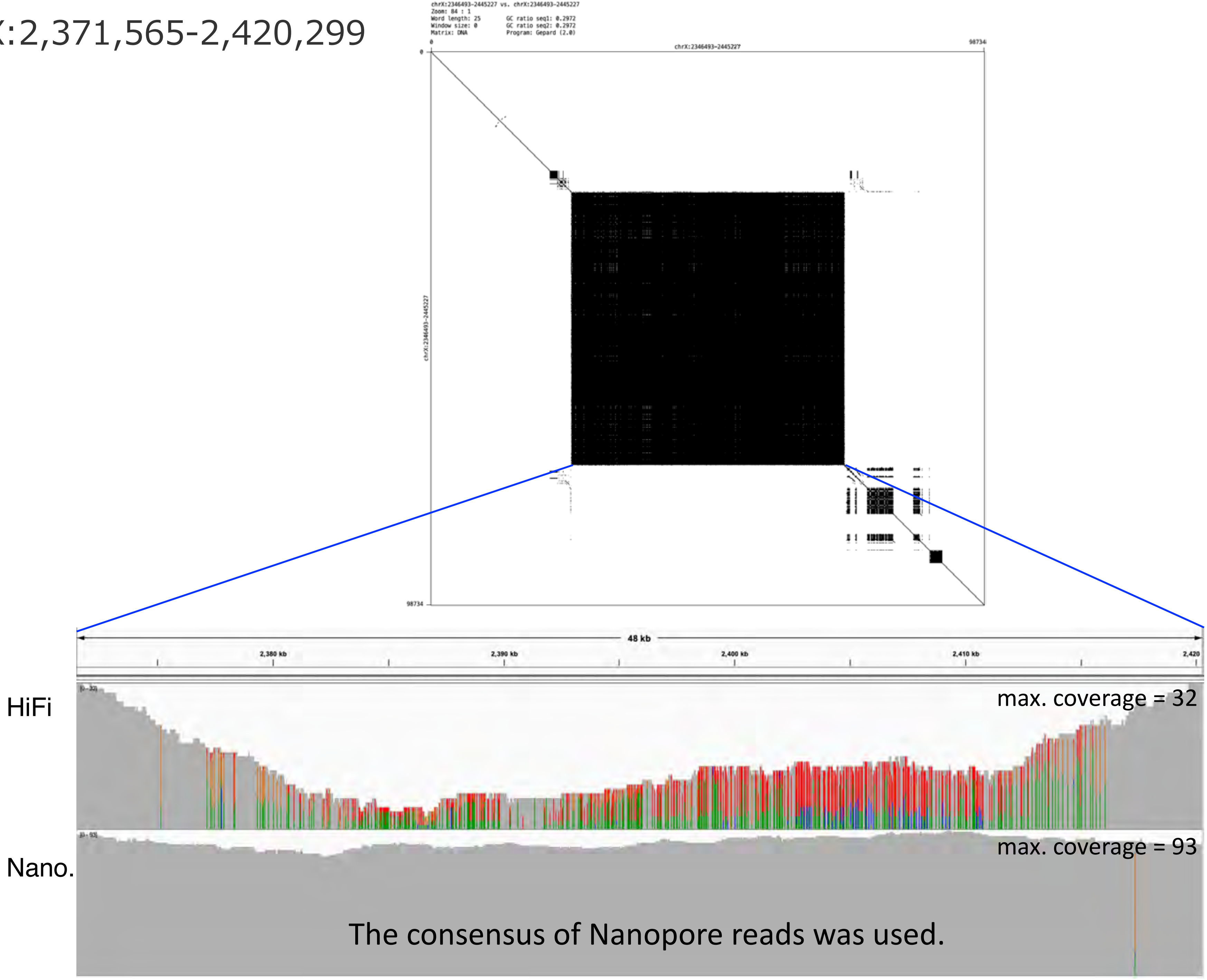

chrX:2,484,213-2,496,772

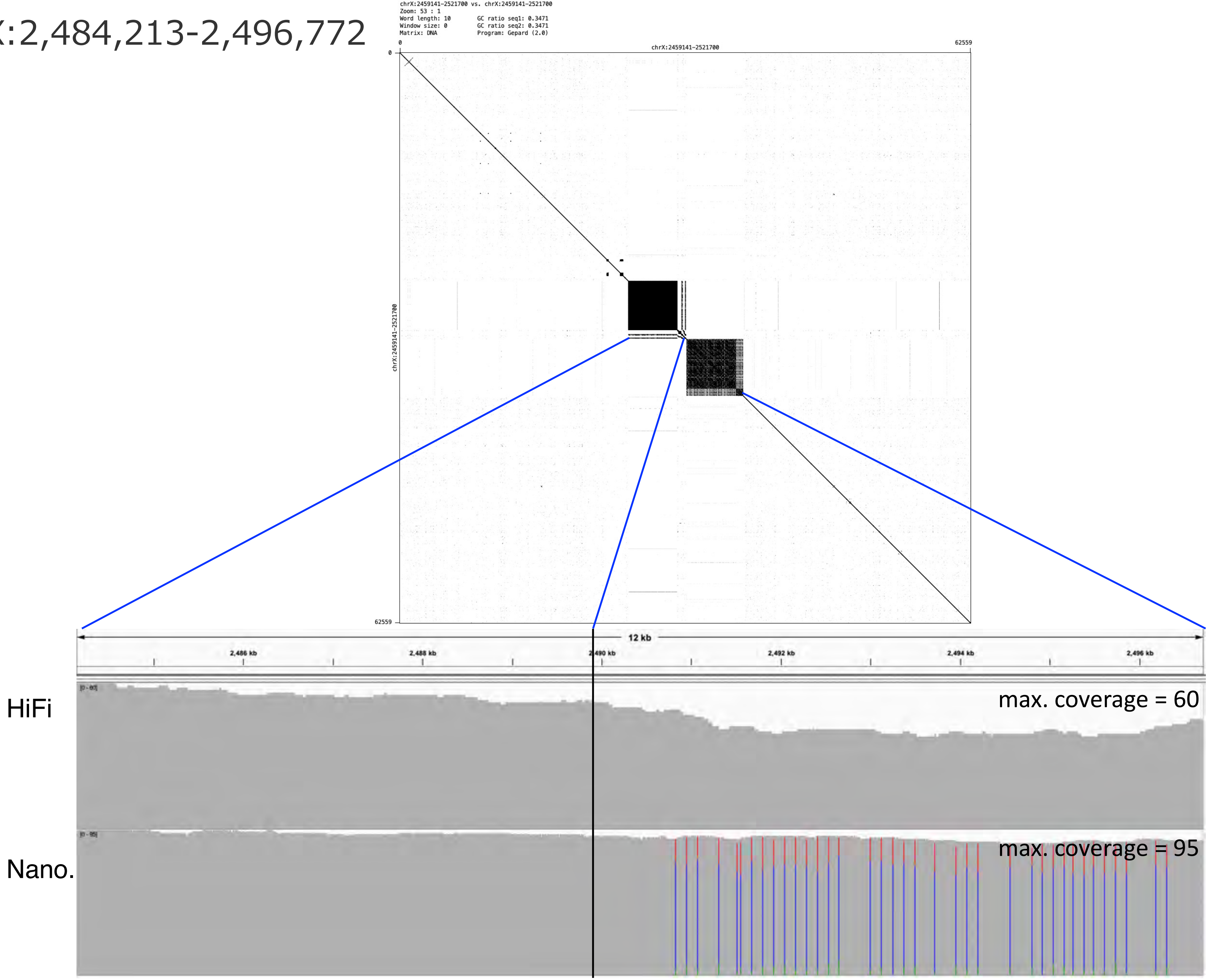

### chrX:3,442,470-3,481,980

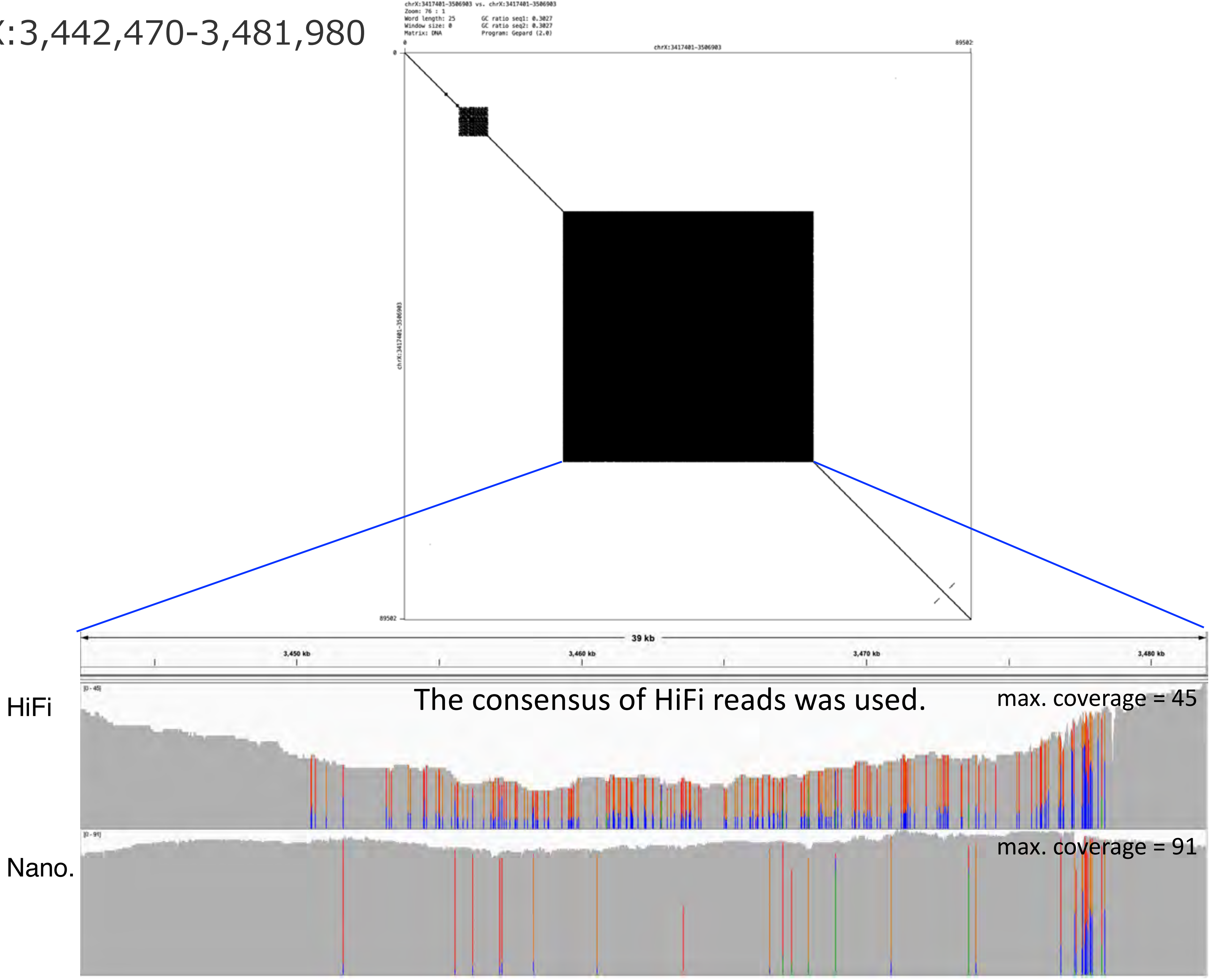

### chrX:4,508,408-4,592,148

Combination of consensuses of HiFi and Nanopore reads

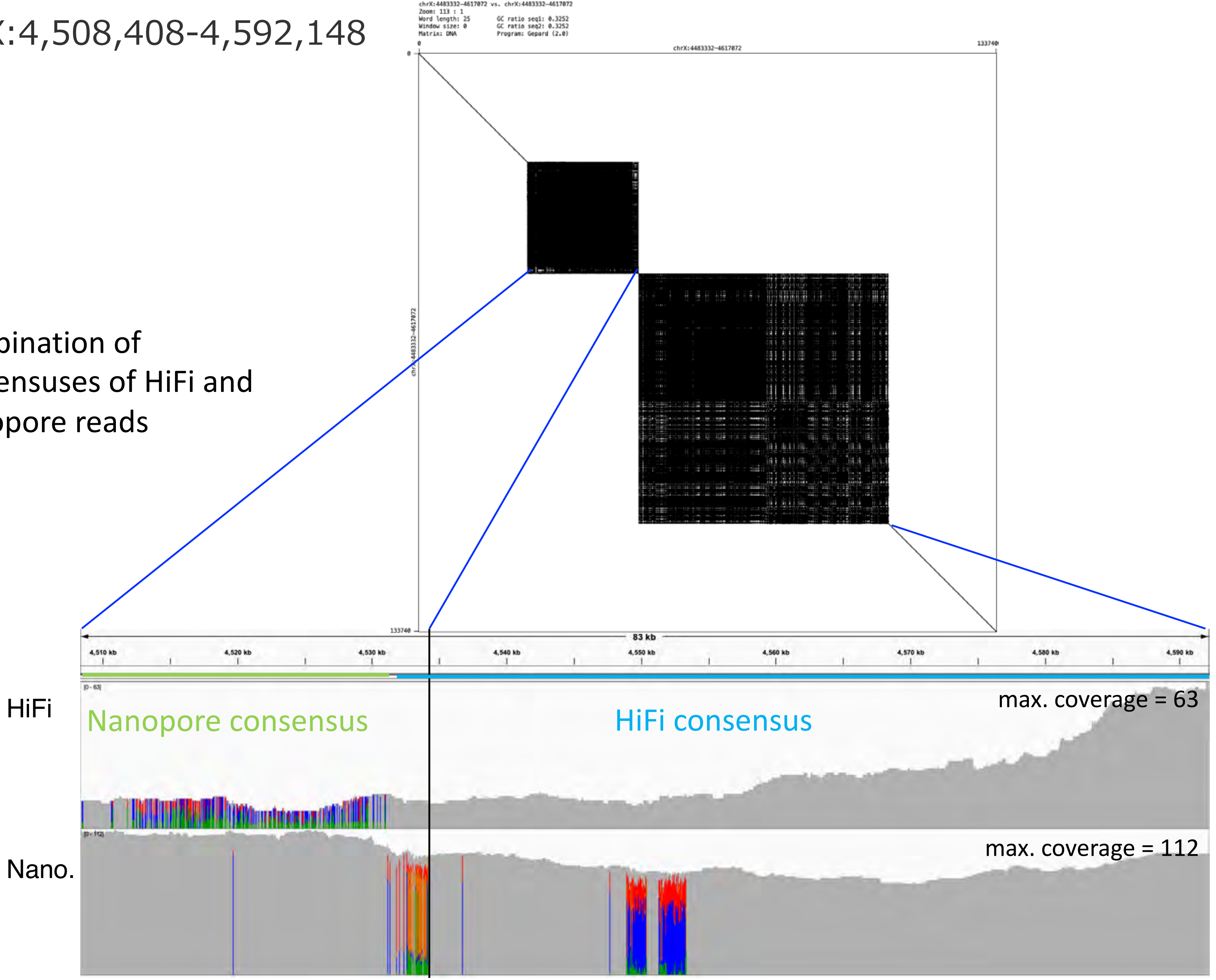

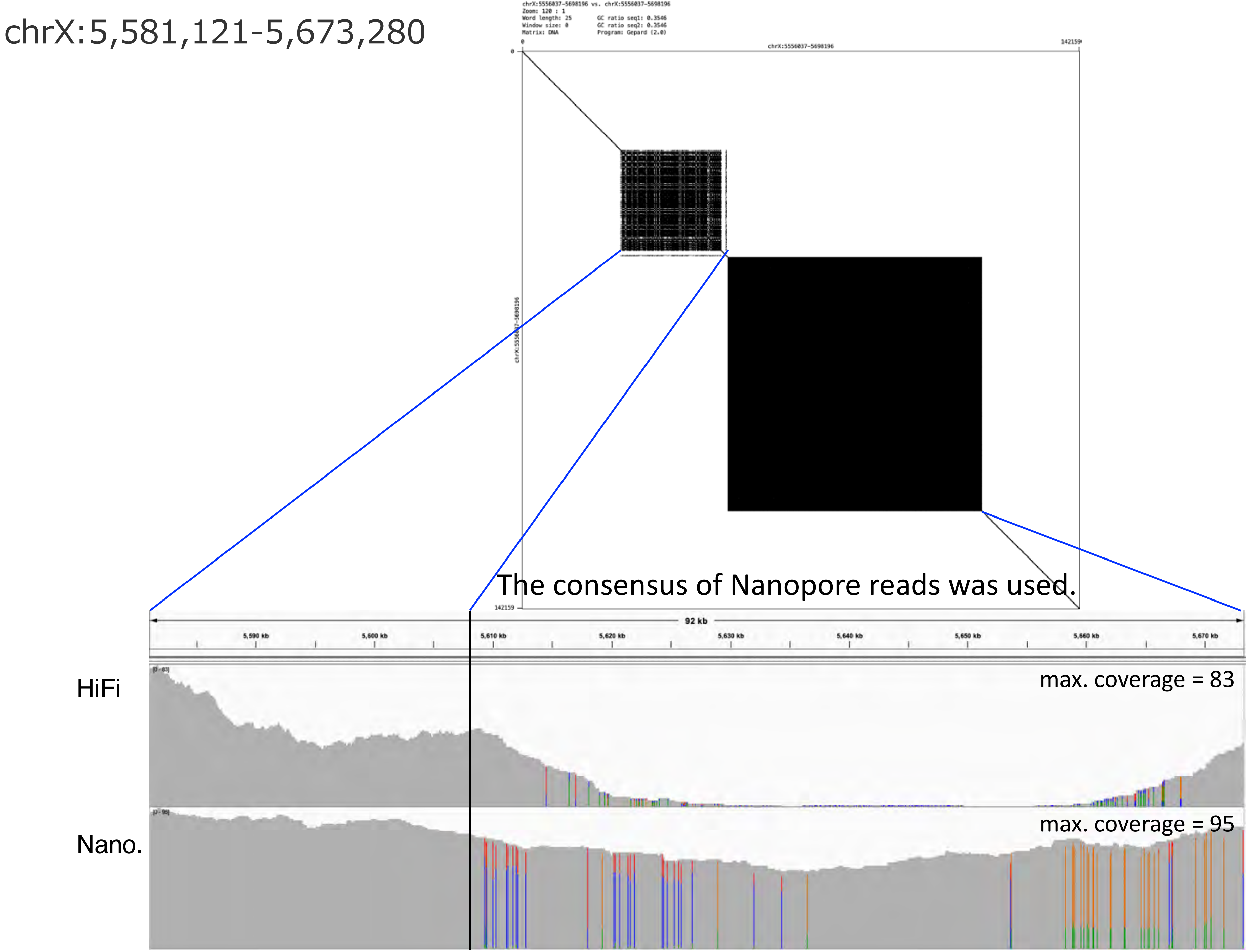
chrX:5,581,121-5,673,280
chrX:5556037-5698196 vs. chrX:5556037-5698196
Zoom: 120 : 1
Word length: 25
Window size: 0
Matrix: DNA
GC ratio seq1: 0.3546
GC ratio seq2: 0.3546
Program: Gepard (2.0)
chrX:5556037-5698196
0
142159
The consensus of Nanopore reads was used.
92 kb
5,590 kb
5,600 kb
5,610 kb
5,620 kb
5,630 kb
5,640 kb
5,650 kb
5,660 kb
5,670 kb
HiFi
max. coverage = 83
Nano.
max. coverage = 95

chrX:6,524,962-6,533,573

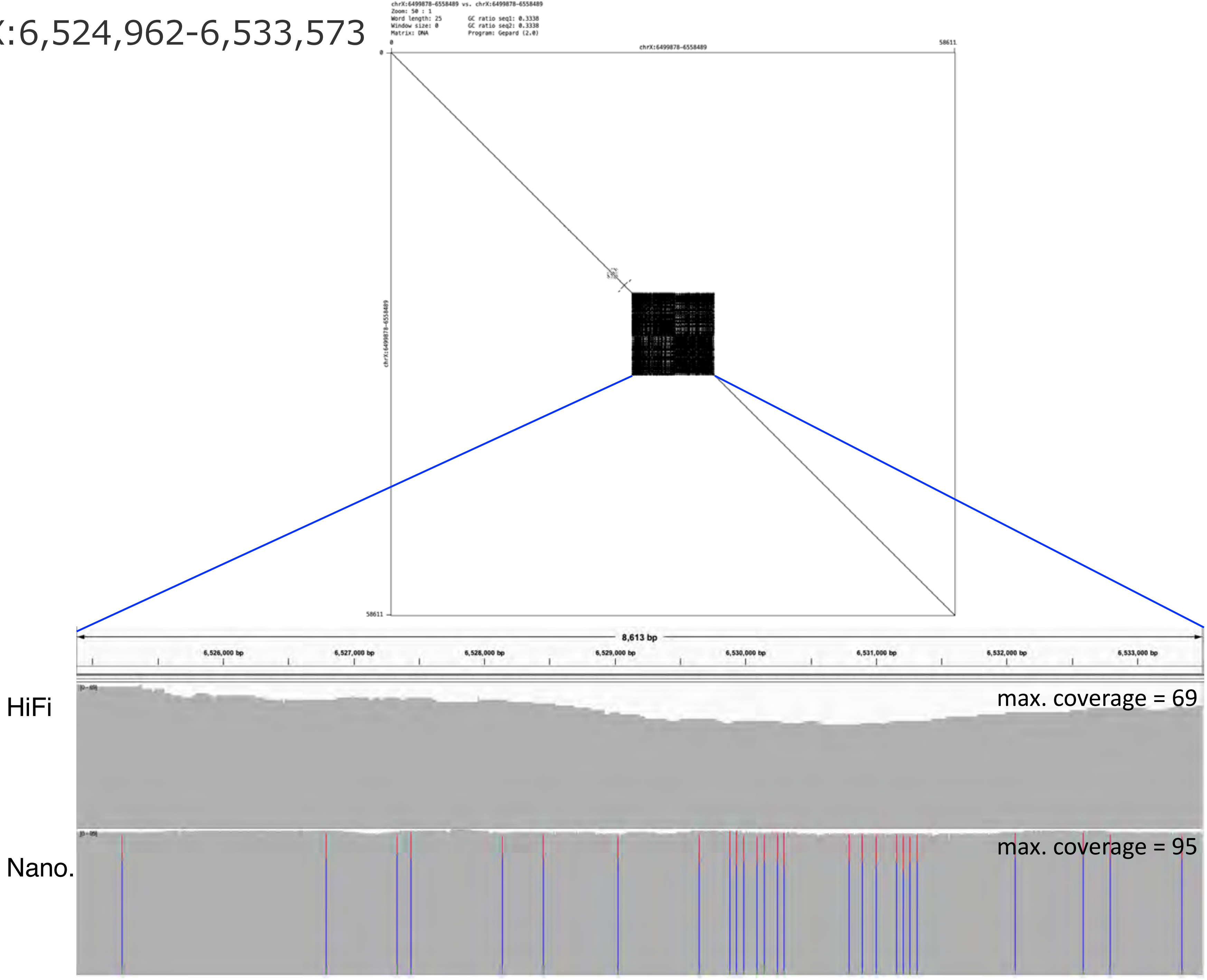

### chrX:7,695,338-7,750,454

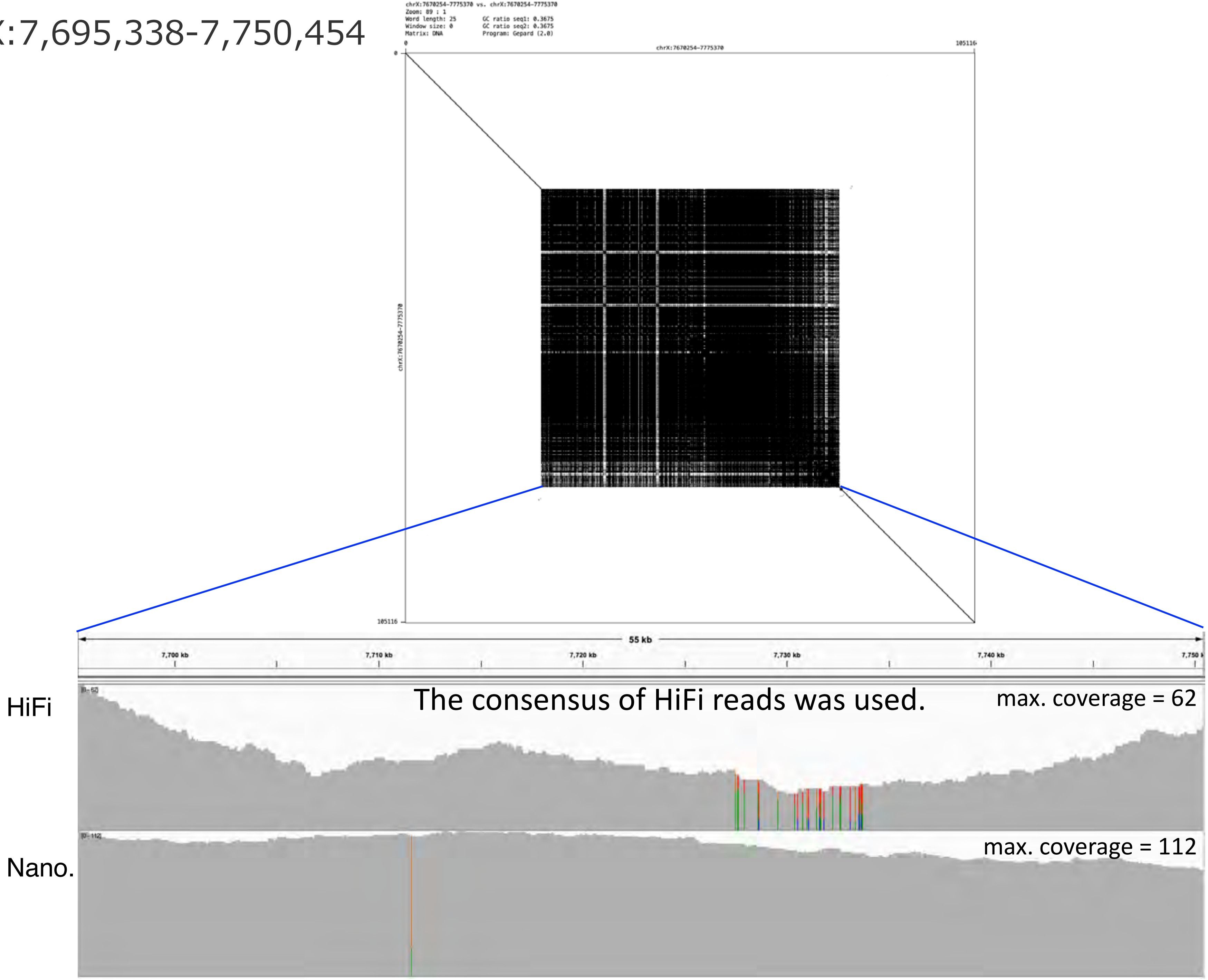

chrX:8,022,827-8,033,469

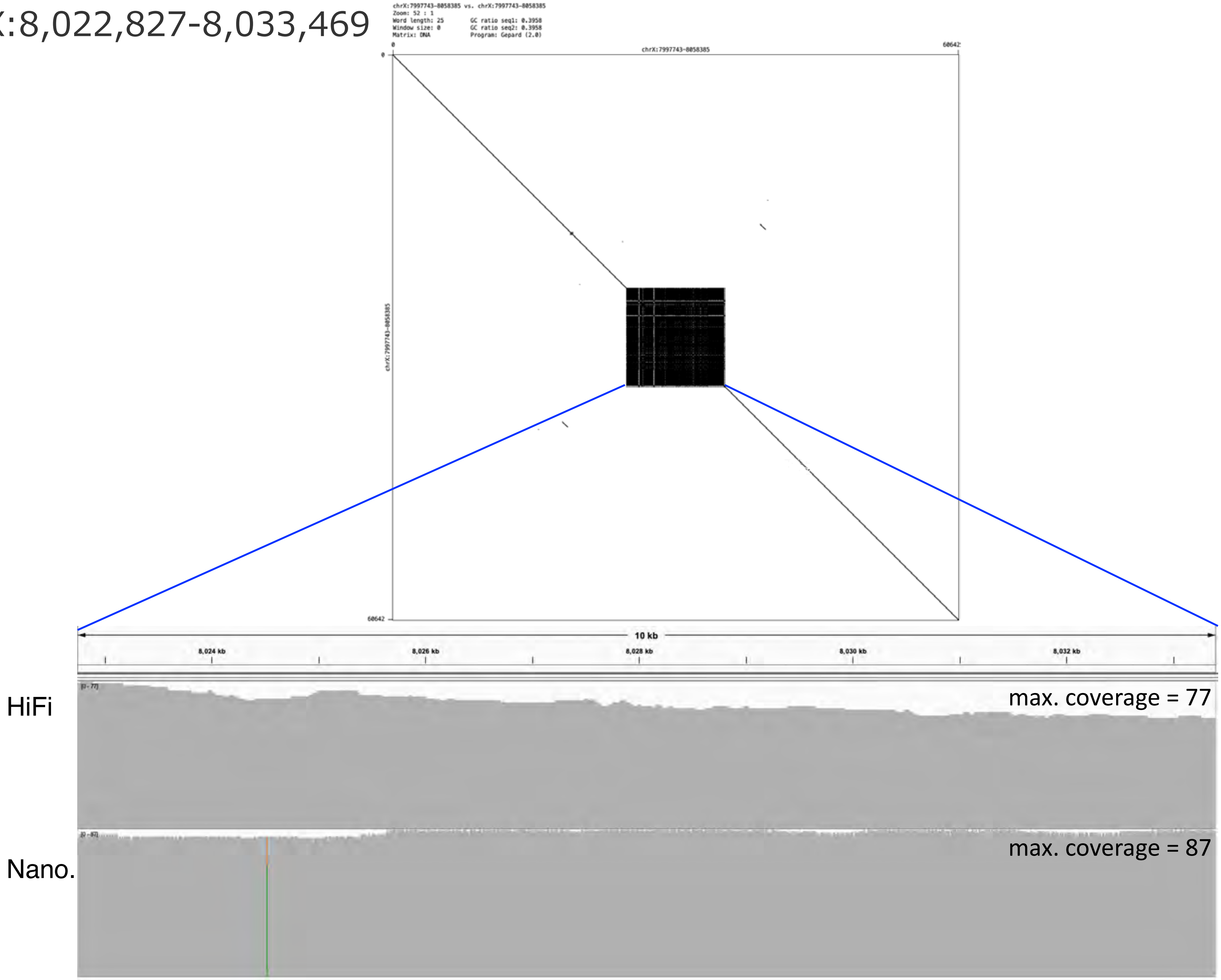

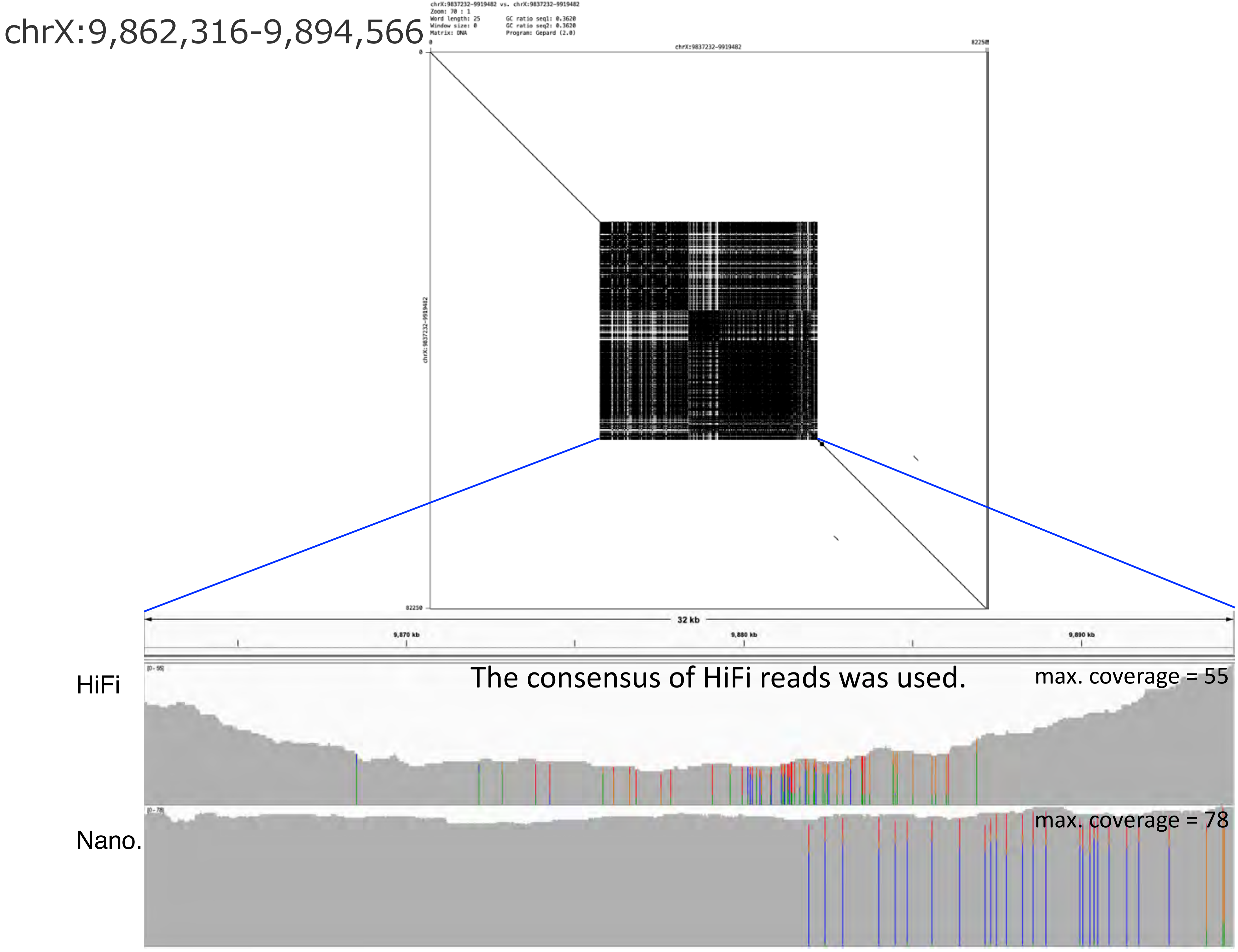
chrX:9,862,316-9,894,566
chrX:9837232-9919482 vs. chrX:9837232-9919482
Zoom: 70 : 1
Word length: 25
Window size: 0
Matrix: DNA
GC ratio seq1: 0.3620
GC ratio seq2: 0.3620
Program: Gepard (2.0)
chrX:9837232-9919482
0
82250
32 kb
9,870 kb
9,880 kb
9,890 kb
HiFi
[0 - 55]
The consensus of HiFi reads was used.
max. coverage = 55
Nano.
[0 - 78]
max. coverage = 78

chrX:10,144,324-10,159,458

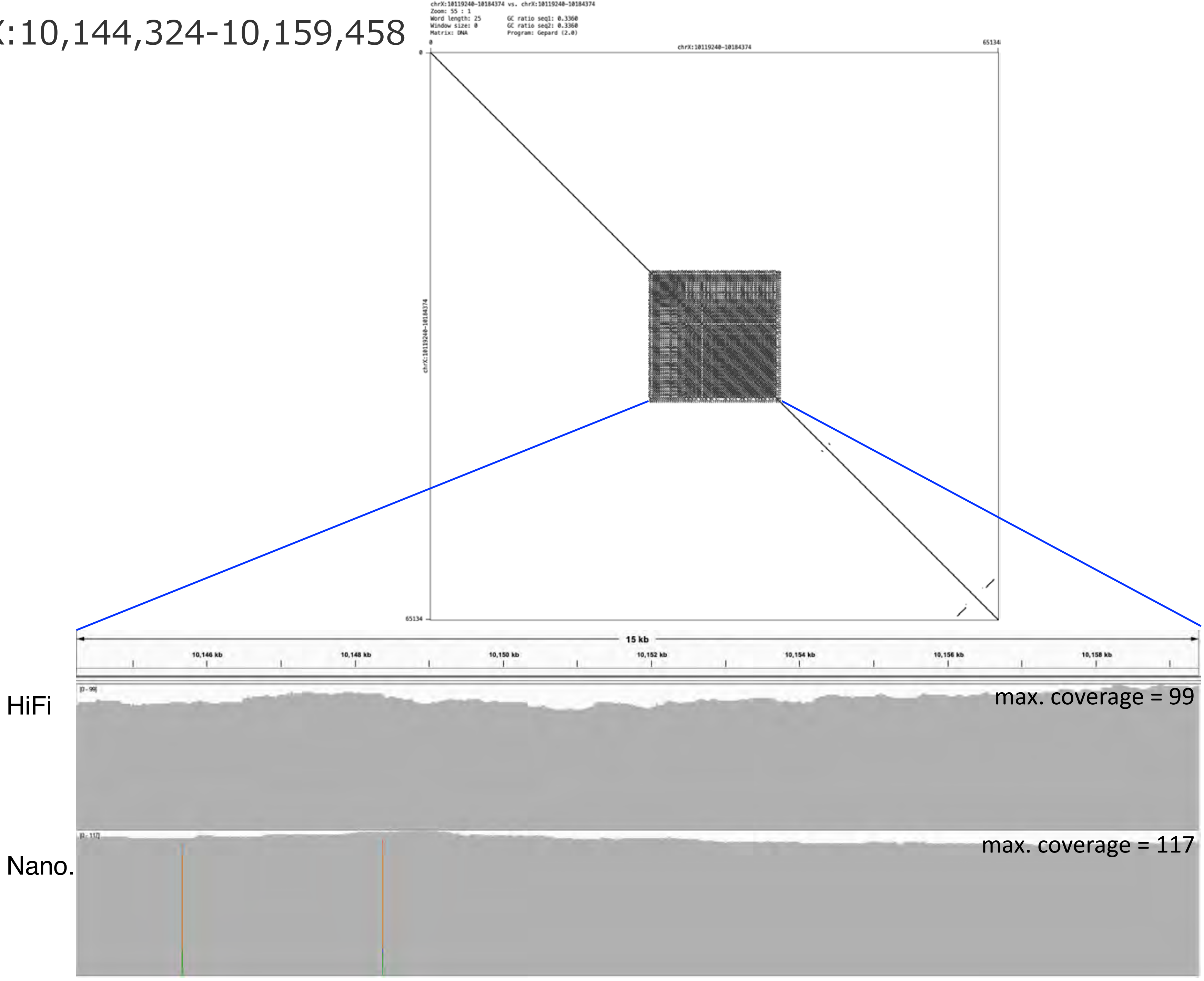

chrX:11,080,698-11,109,263

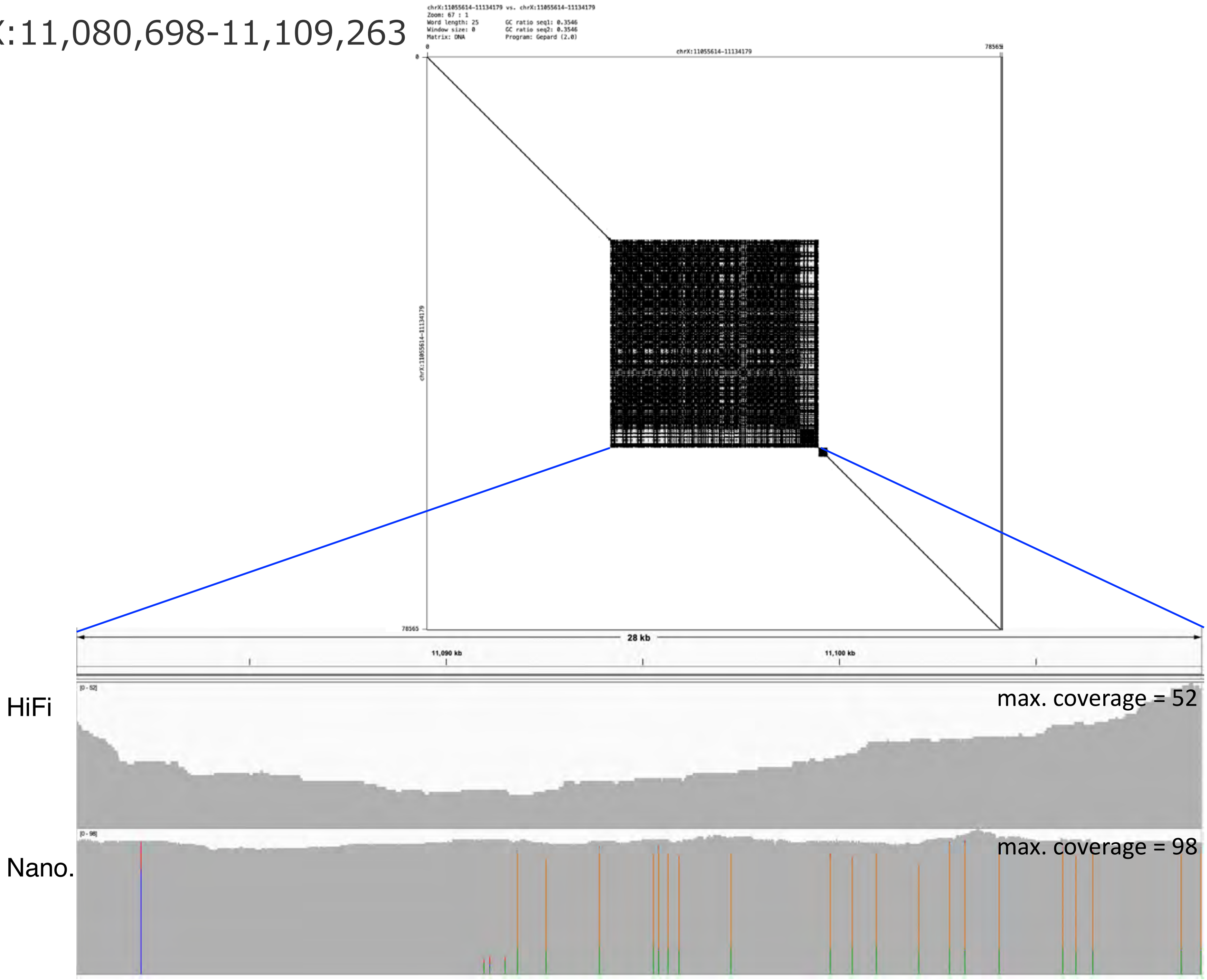

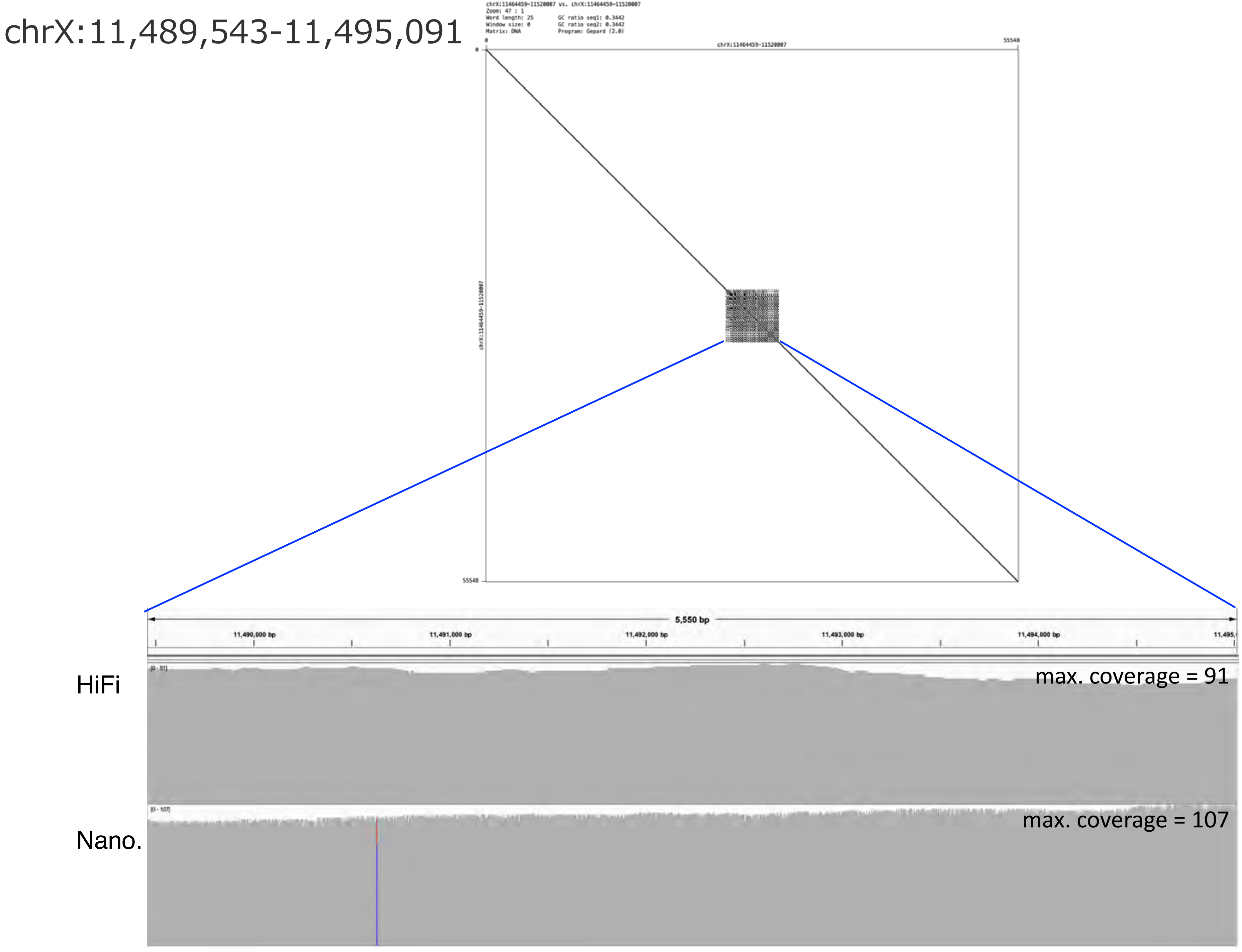
chrX:11,489,543-11,495,091
chrX:11464459-11520007 vs. chrX:11464459-11520007
Zoom: 47 : 1
Word length: 25
Window size: 0
Matrix: DNA
GC ratio seq1: 0.3442
GC ratio seq2: 0.3442
Program: Gepard (2.0)
chrX:11464459-11520007
0
55548
5,550 bp
11,490,000 bp
11,491,000 bp
11,492,000 bp
11,493,000 bp
11,494,000 bp
HiFi
max. coverage = 91
Nano.
max. coverage = 107

### chrX:12,540,490-12,691,681

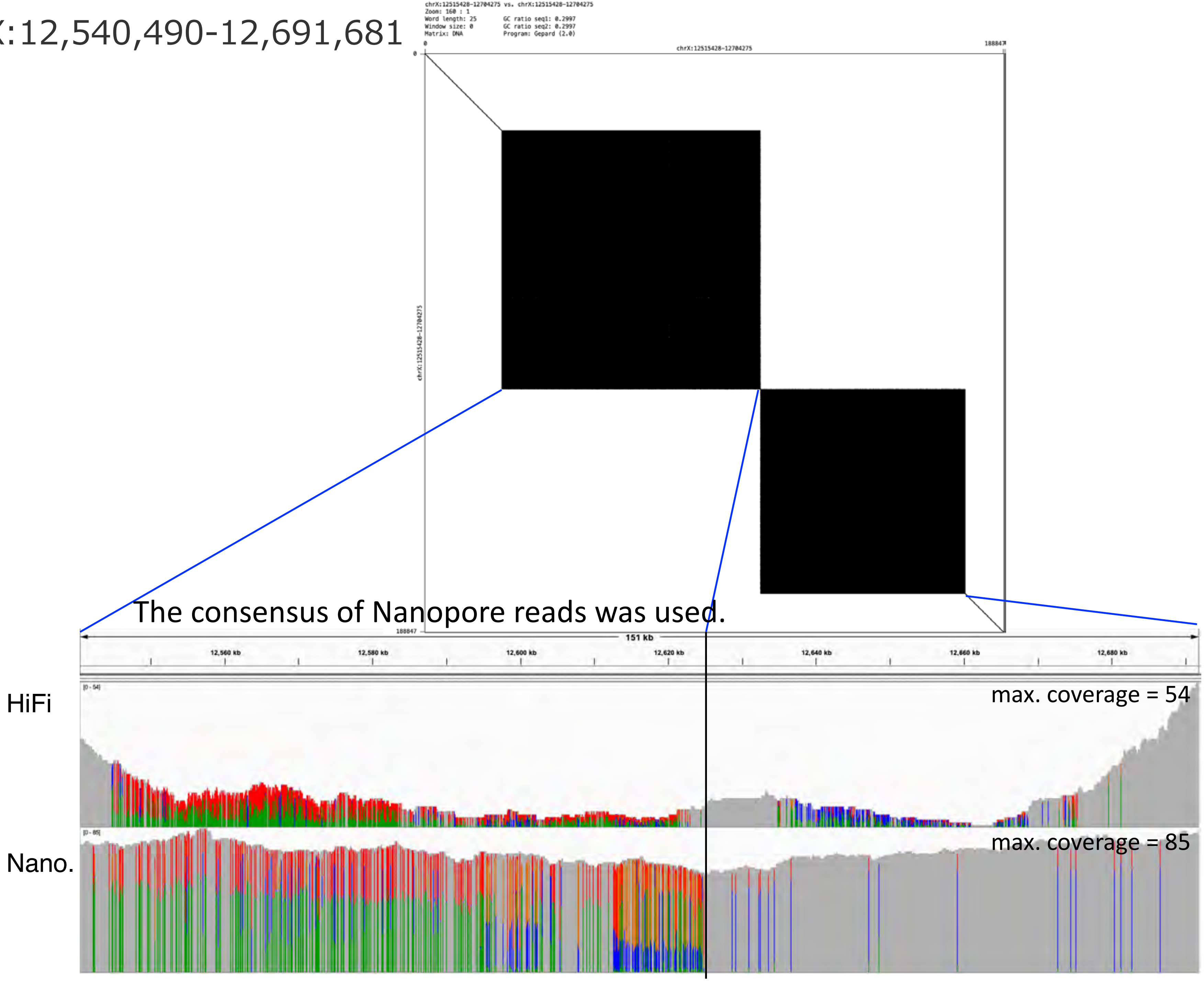

chrX:12,788,546-12,843,414

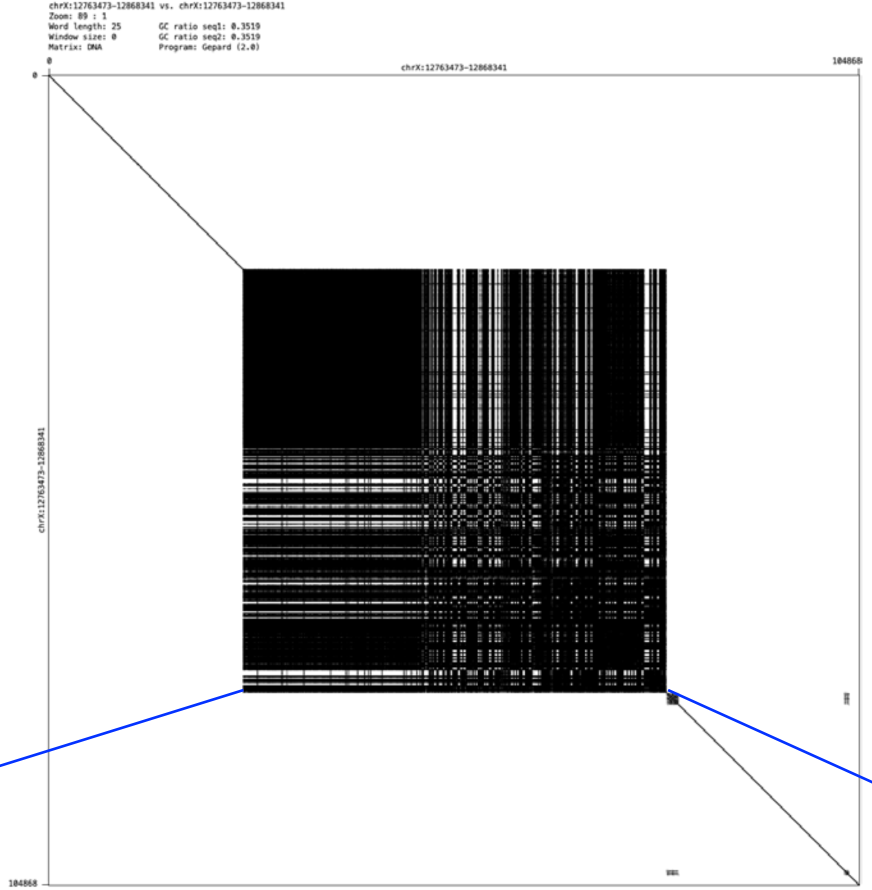


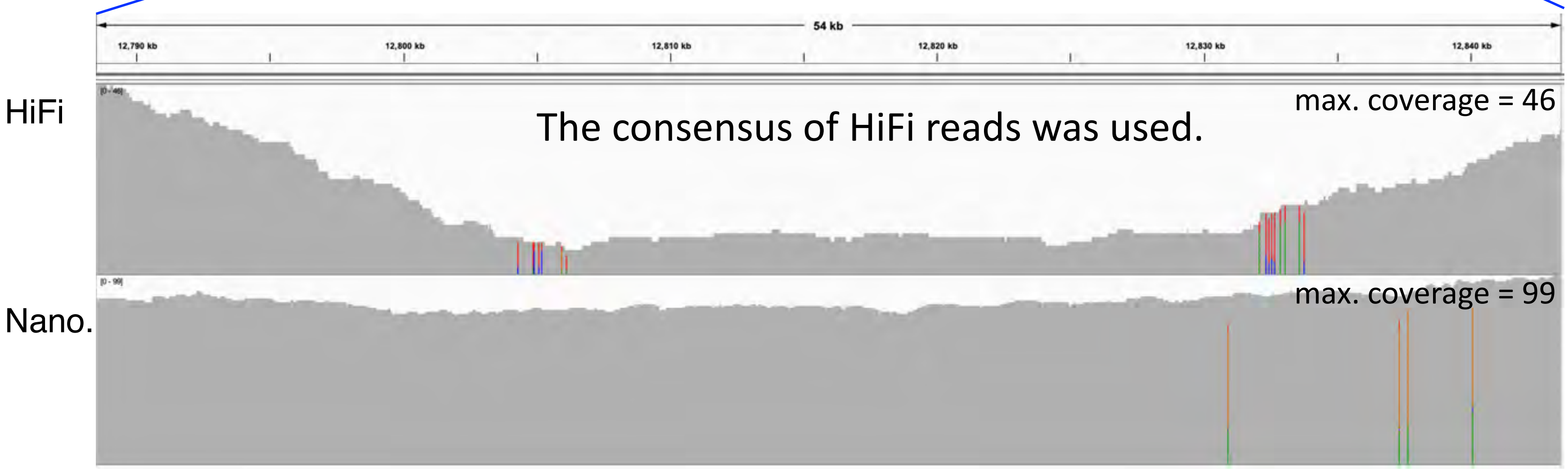

chrX:13,232,904-13,246,781

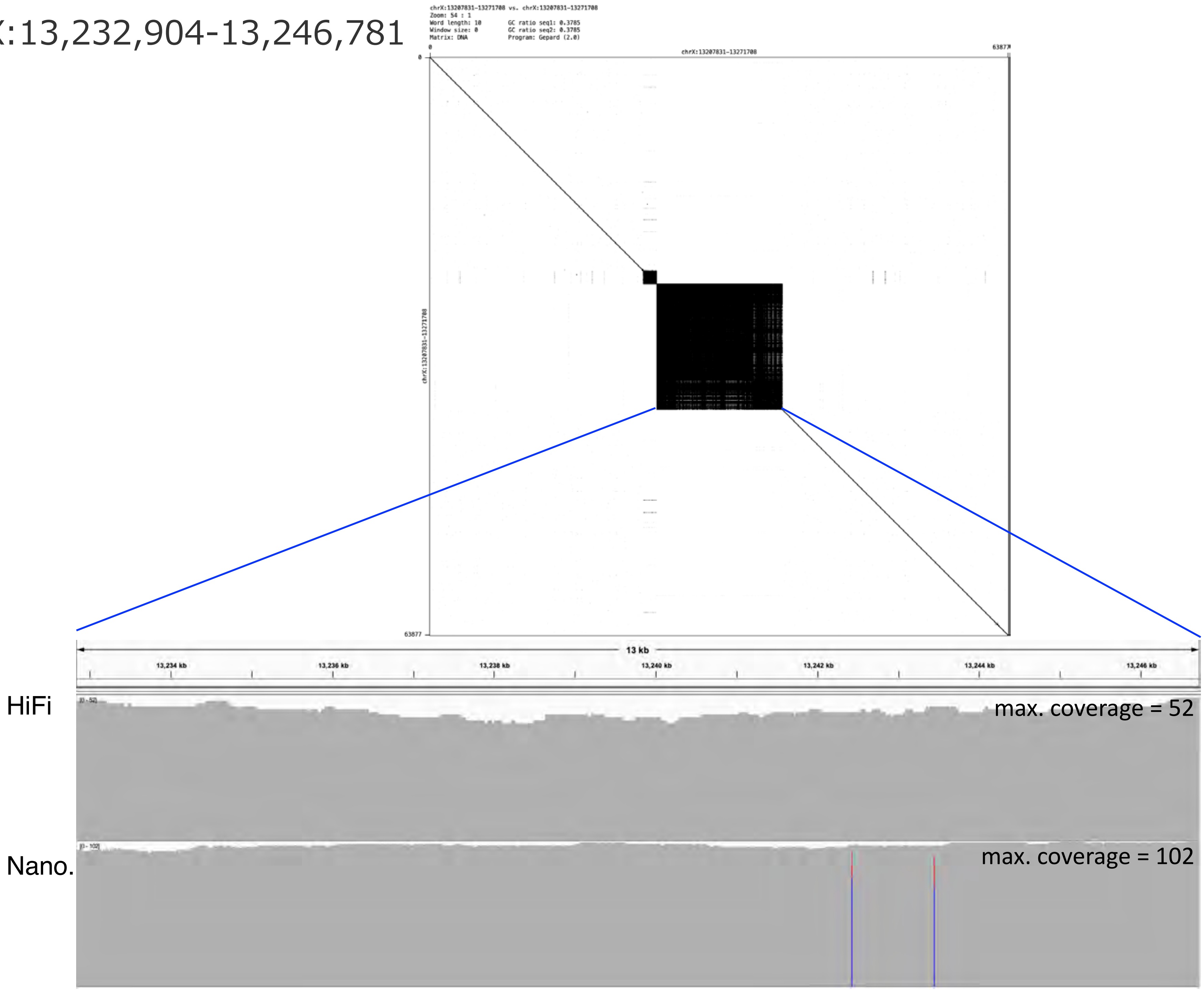

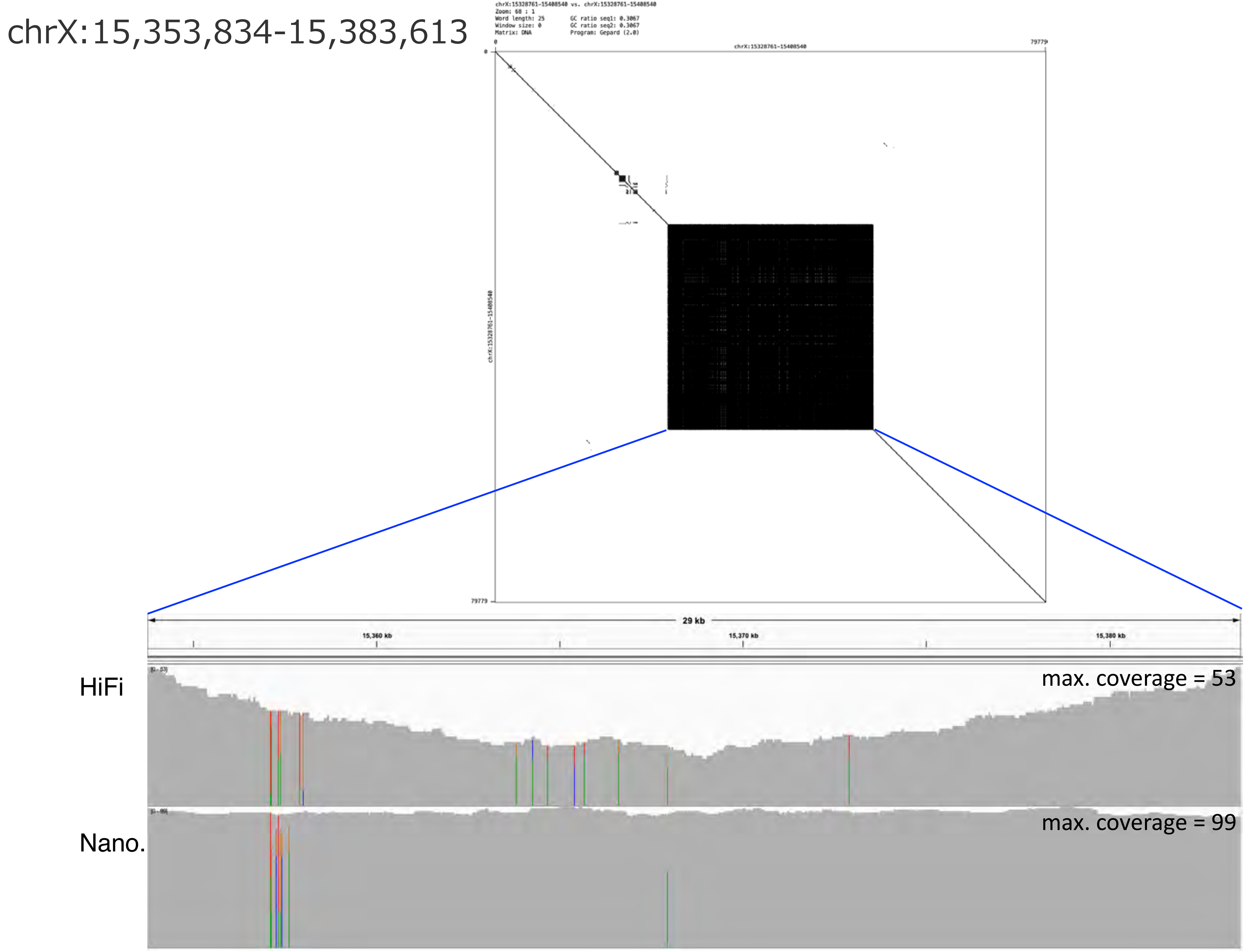
chrX:15,353,834-15,383,613
chrX:15328761-15408540 vs. chrX:15328761-15408540
Zoom: 68 : 1
Word length: 25
Window size: 0
Matrix: DNA
GC ratio seq1: 0.3067
GC ratio seq2: 0.3067
Program: Gepard (2.0)
chrX:15328761-15408540
0
79779
29 kb
15,360 kb
15,370 kb
15,380 kb
HiFi
max. coverage = 53
Nano.
max. coverage = 99

chrX:15,902,101-15,945,941

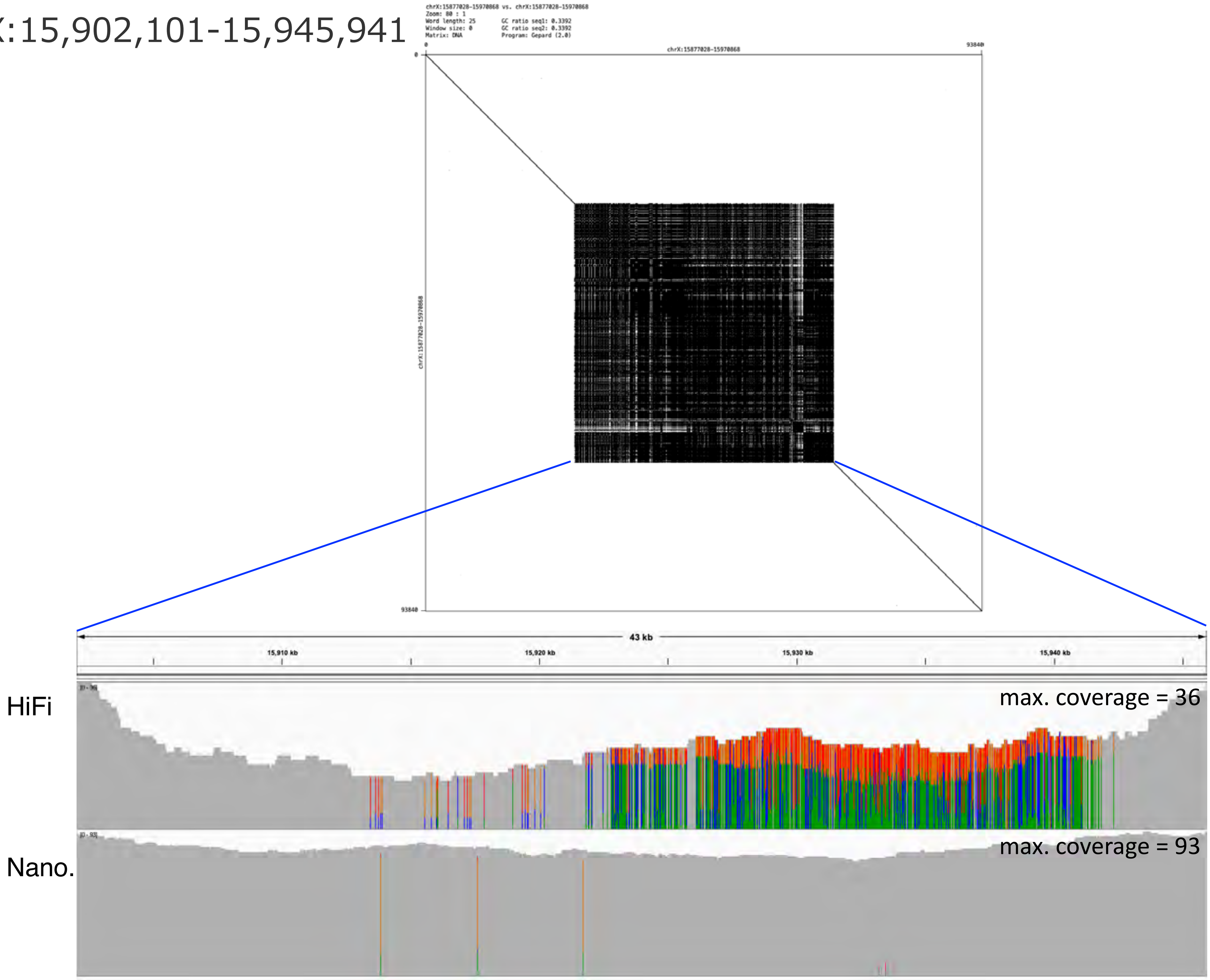

chrX:16,263,528-16,273,490

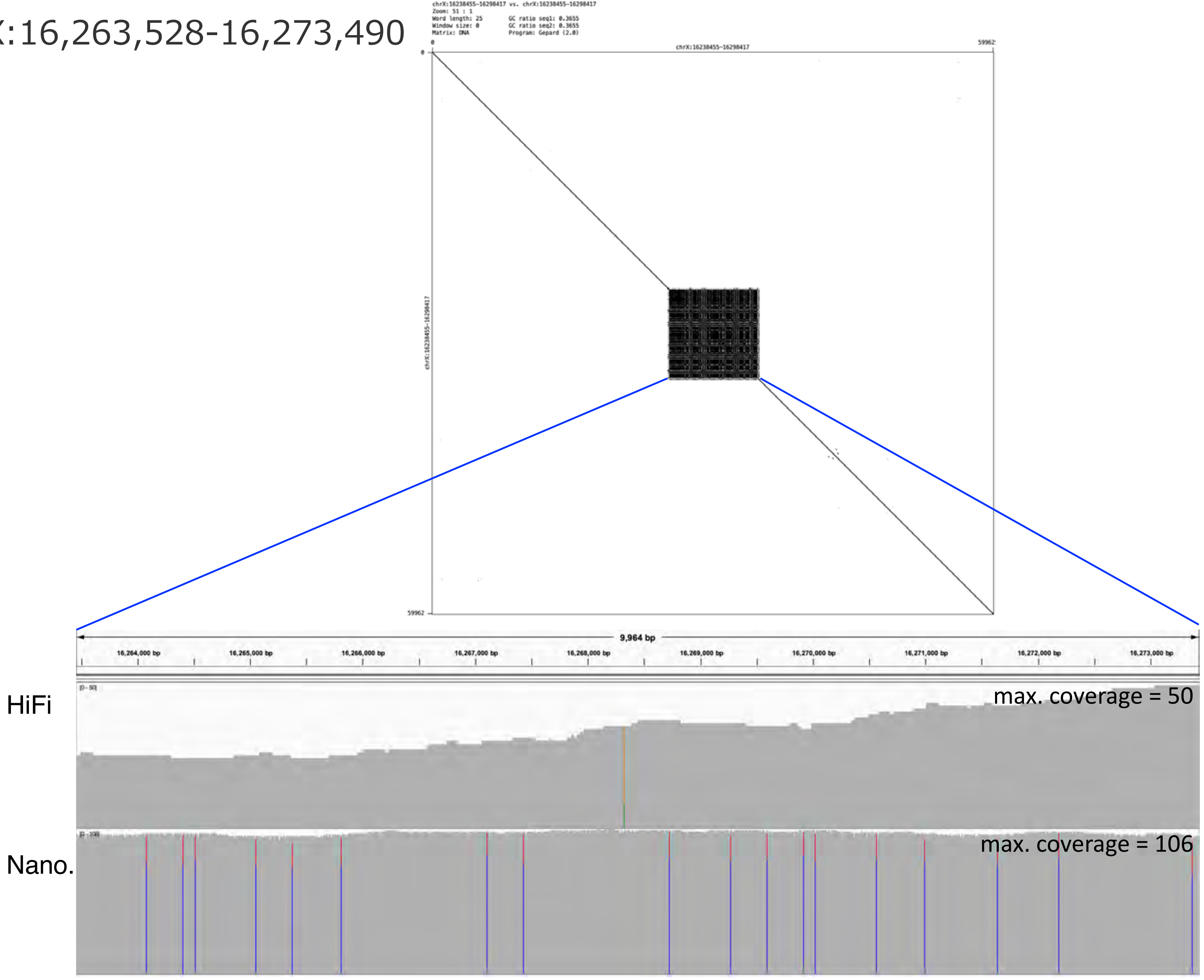

chrX:17,805,304-17,810,435

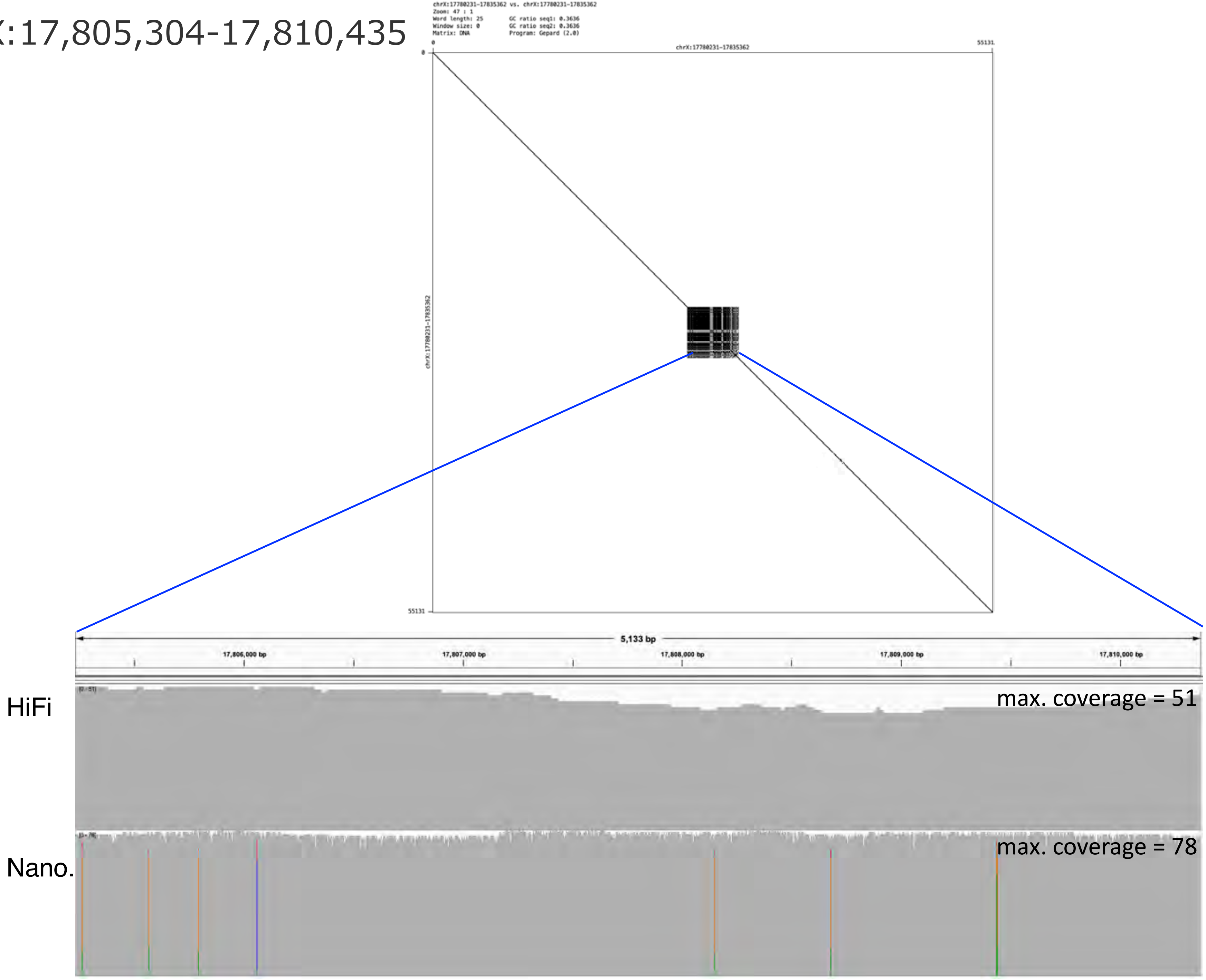

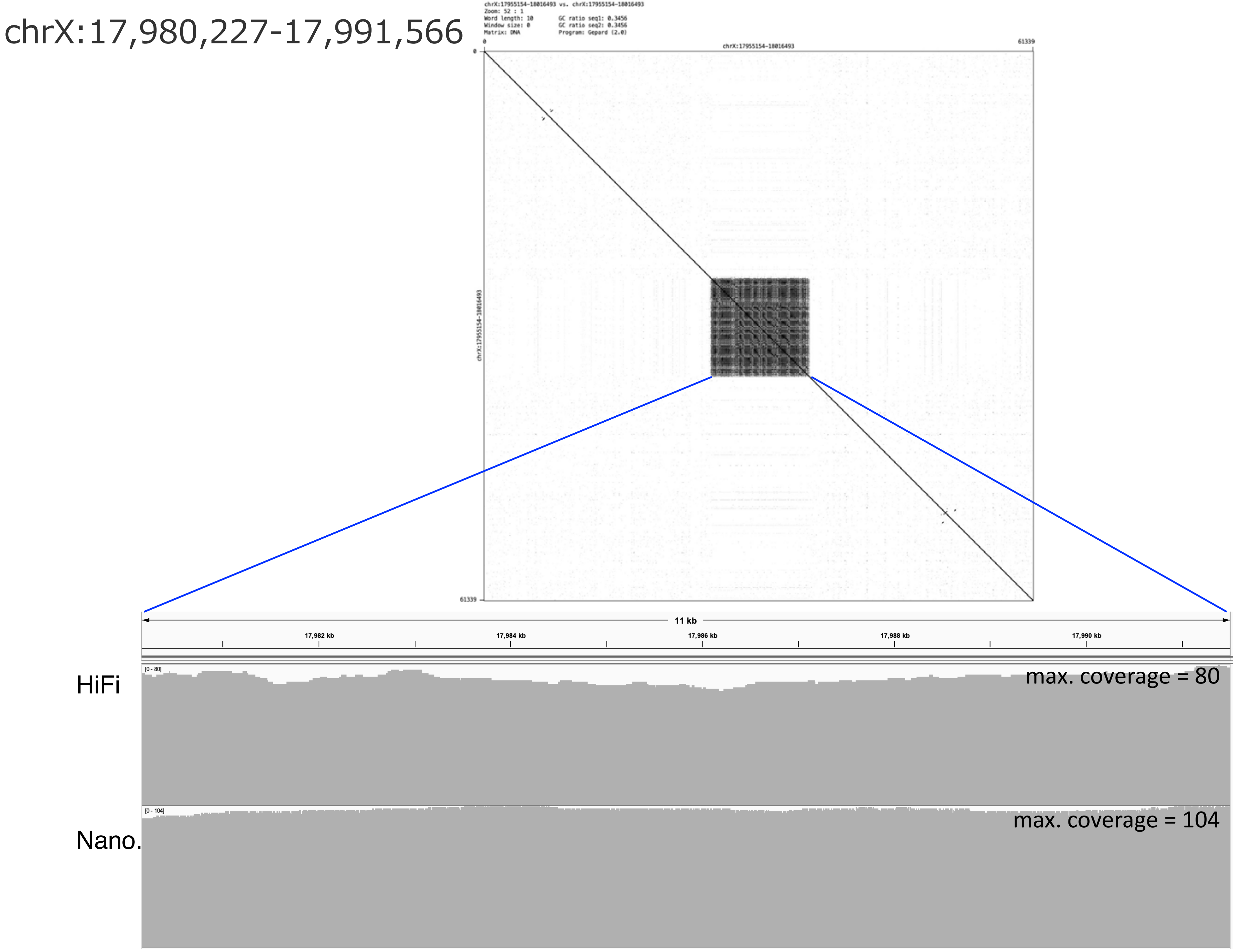
chrX:17,980,227-17,991,566
chrX:17955154-18016493 vs. chrX:17955154-18016493
Zoom: 52 : 1
Word length: 10
Window size: 0
Matrix: DNA
GC ratio seq1: 0.3456
GC ratio seq2: 0.3456
Program: Gepard (2.0)
chrX:17955154-18016493
0
61339
chrX:17955154-18016493
11 kb
17,982 kb
17,984 kb
17,986 kb
17,988 kb
17,990 kb
HiFi
[0 - 80]
max. coverage = 80
Nano.
[0 - 104]
max. coverage = 104

chrX:18,495,757-18,511,331

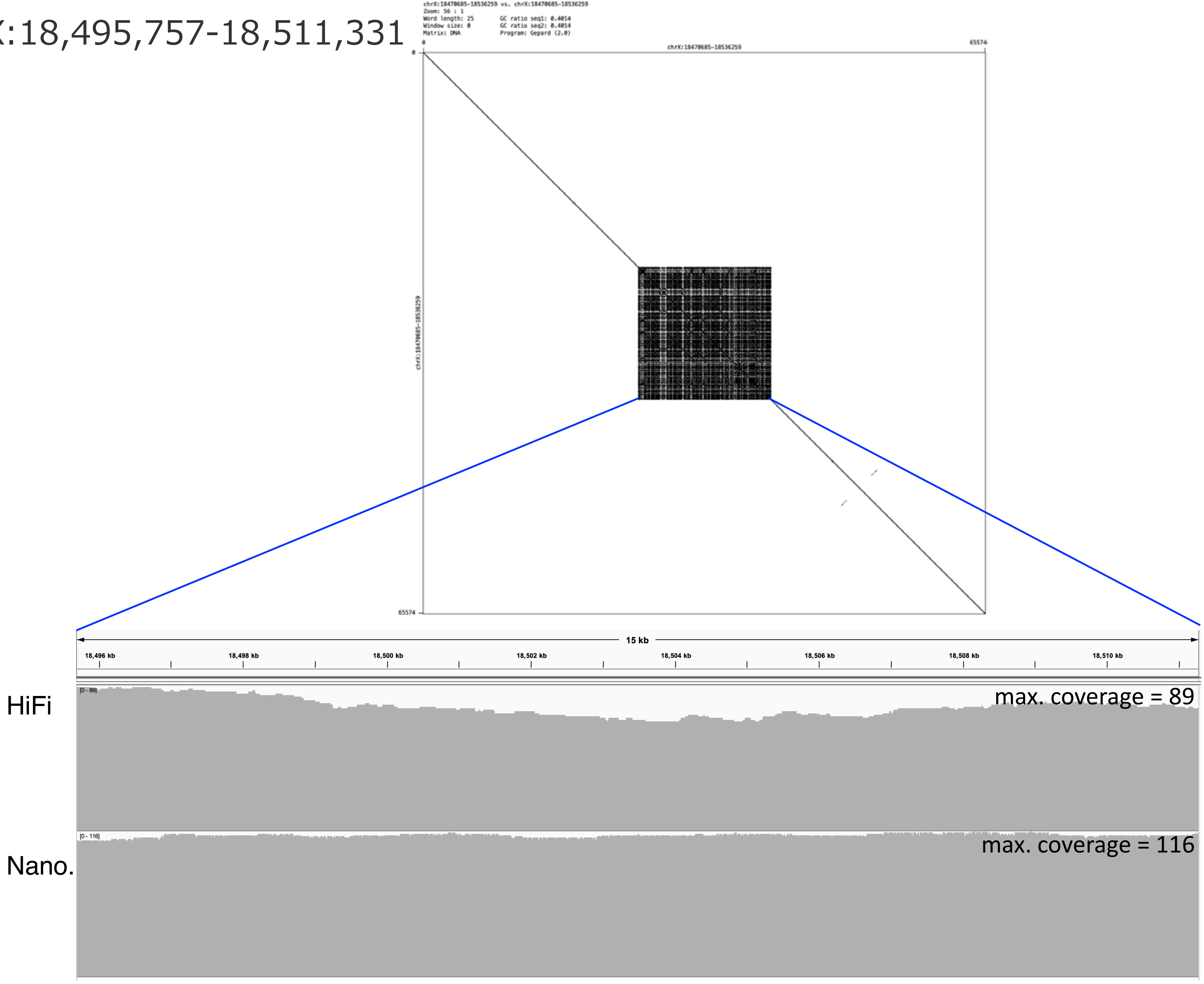
