## Supplementary Figure 5 for "CGC1, a new reference genome for *Caenorhabditis elegans*"

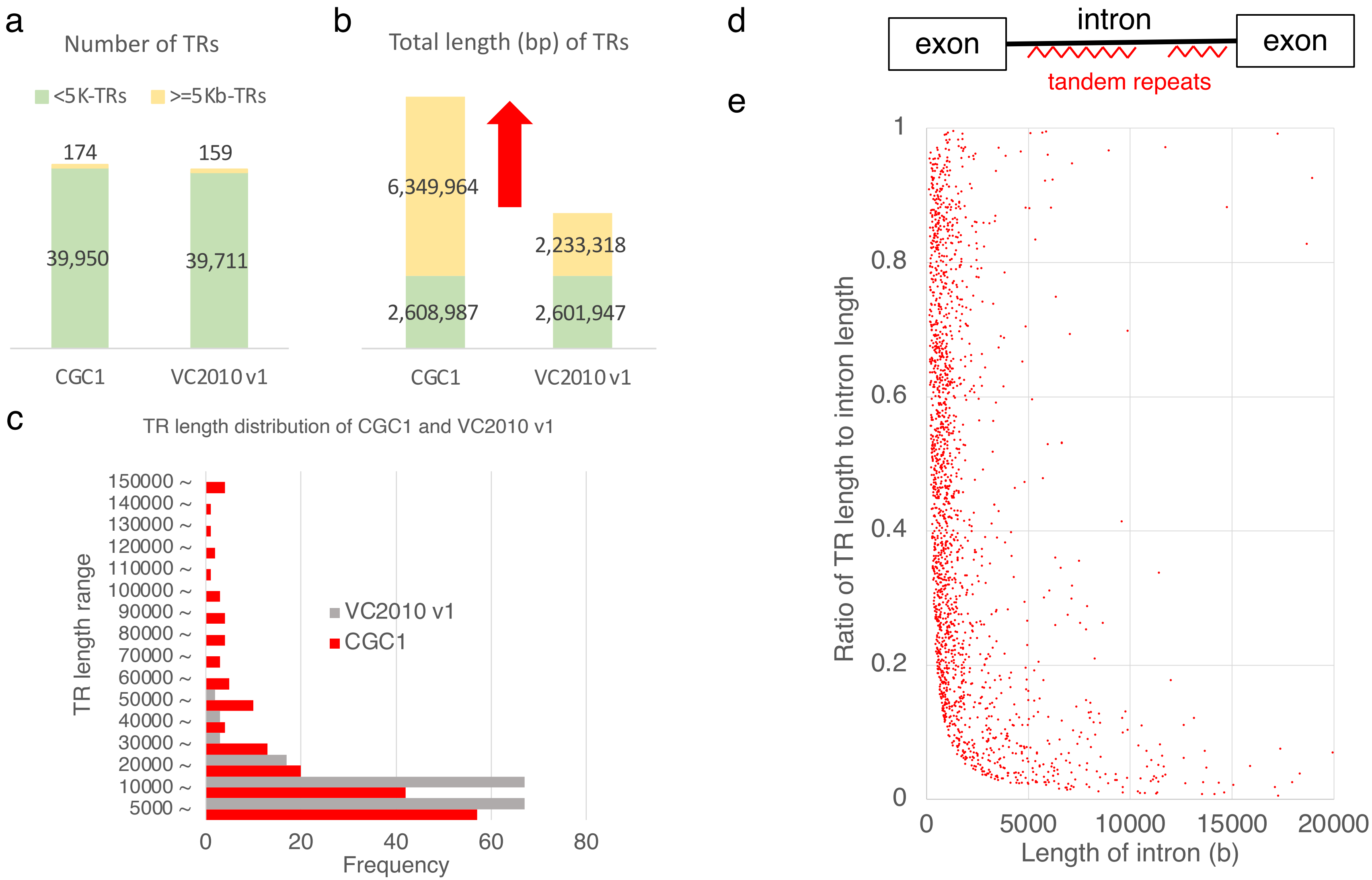


**Supplementary Figure 5. Statistics of tandem repeats (TRs). a-b**. TRs are clustered into two groups: one group has TRs shorter than 5 kb (green), and the other has the remaining longer TRs (yellow). Figure **a** shows the number of each group and Figure **b** displays the total length of TRs in each group in the respective assemblies of CGC1 and VC2010 (v1). Of note, the longer TR group (of length ≥5 kb) dominates the total length of TRs in the new CGC1 assembly. **c**. To highlight that CGC1 has longer TRs than VC2010, the histogram shows the length distribution of TRs of length 5 kb or more in the two assemblies. **d-e**. To examine whether TRs in introns dominate the sizes of introns, Figure **d** illustrates an intron with a tandem repeat. Each dot in Figure **e** shows the pair of the length of an intron (x-axis) and the ratio of the total length of TRs in the intron to the intron length (y-axis). Some introns are largely filled with TRs, but a number of introns are almost free from TRs. For example, among introns of length 1 kb or more, 113 have ratio 0.8 or more, while 432 have ratio 0.2 or less. Supplementary Table 18 shows the list of all introns with TRs.
