## Supplementary Figure 6 for "CGC1, a new reference genome for *Caenorhabditis elegans*"

**Supplementary Figure 6. Evolution of 12 regions filled with the 27-mer tandem repeat unit and its variants.** The upper right panel shows the self-dot plot of the tandem repeat at the above locus that are filled with the 27-mer unit and its variants. The table below shows the strings of 27-mer units present in the focal region, their frequencies, mutations, indels, and their names such as type1, type2, and type3 (see their details in Supplementary Table 4). The phylogenetic tree at the left-bottom shows the proximity of variants in terms of sequence similarity. The right bottom shows the locations of unit variants to illustrate how units are distributed inside the tandem repeat. Type 1 units are almost evenly distributed in all regions, while type 2 units are found evenly only in the five regions in the blue box of Figure 3. In addition to prevalent type 1 and type 2 units, another unit and its variants, denoted by type 3 and type 3' respectively, are enriched in the group of regions in the green box of Figure 3. In addition, each region has its own unique unit variants. The characteristics of each unit represent the evolutionary footprint of an individual region.

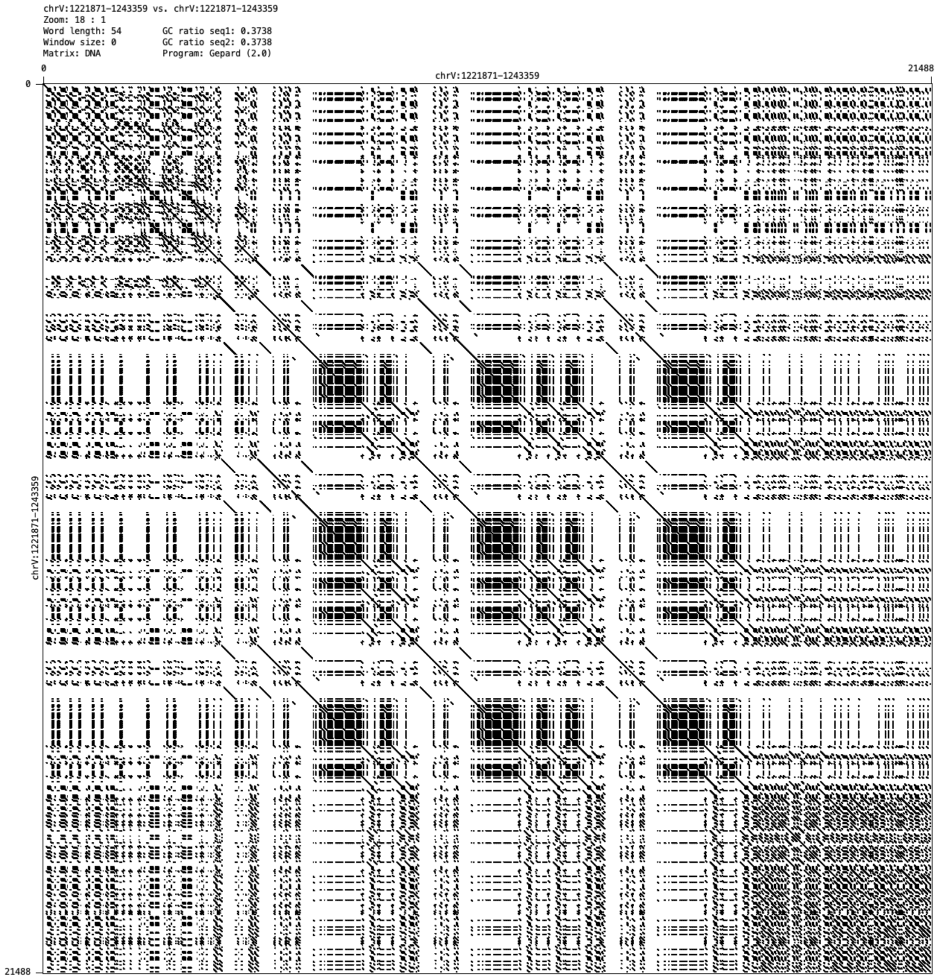


### chrV: 1,221,871-1,243,359

| chrV_1 | | | | |
|---|---|---|---|---|
| Representative unit and its variant | Freq. | mutation | indel | |
| ACTCTCTGTGGCTTCCCACTATATTTT | 238 | | | type1 |
| ACTCTCTGTGGCTTCACAGTATATTTT | 323 | 16A 19G | | |
| ACTCTCTGTGGCTTCACAGTATTTTTT | 57 | 16A 19G 23T | | |

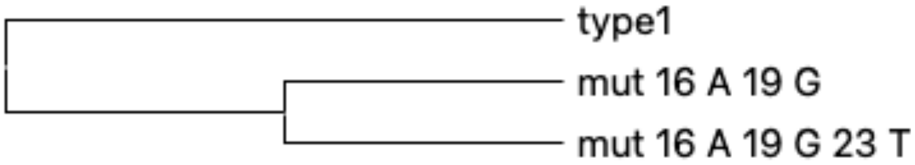


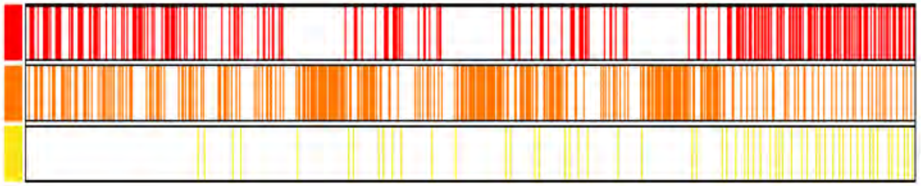

### chrIV: 2,857,216-2,919,705

| chrIV_1 | | | | |
|---|---|---|---|---|
| Representative unit and its variant | Freq. | mutation | indel | |
| ACTCTCTGTGGCTTCCCACTATATTTT | 448 | | | type1 |
| ACTCTCTGTGGCTTCCCACTATAGTTT | 90 | 24G | | |
| ACTCTCTGTGGCTTCACCAATTATTTT | 1203 | 16A 18-21 CAAT | | type3’ |
| ACTCTCTGAGGCTTCACCAATTATTTT | 48 | 9A 16A 18-21 CAAT | | |
| ACTCTCTGTGGCTTCACC-ATTTATTTT | 48 | 16A 18C 20A 23T | 19 del | |
| ACTCTCTGTGGCTTCACCAAATATTTT | 34 | 16A 18-20 CAA | | |

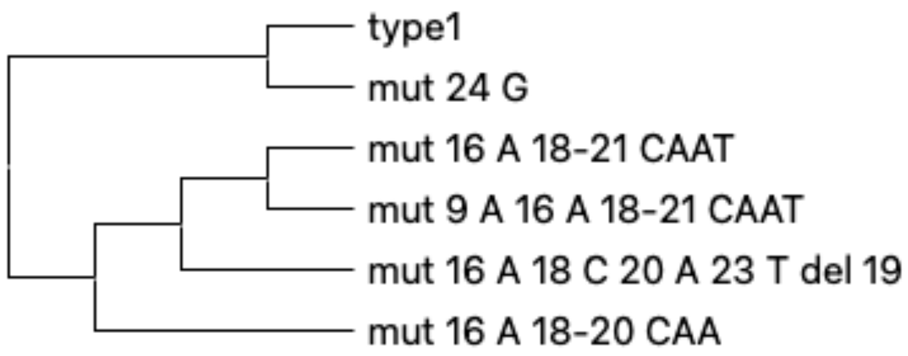


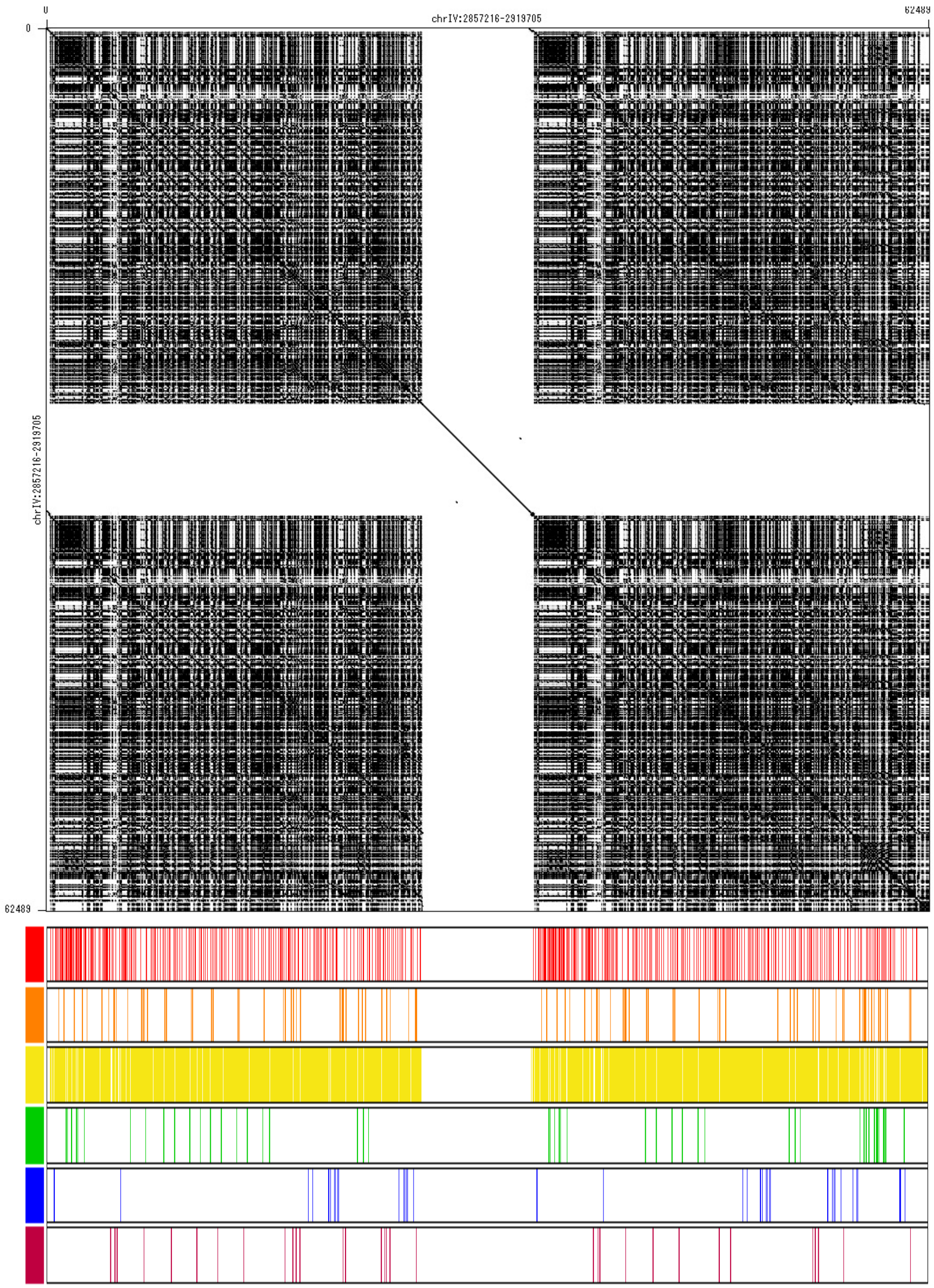

### chrI:952,541-1,043,792

| chrl_1 | | | | |
|---|---|---|---|---|
| Representative unit and its variant | Freq. | mutation | indel | |
| ACTCTCTGTGGCTTCCCACTATATTTT | 1780 | | | type1 |
| -CTCTCTGTGGCTTCCCACTATATTTT | 178 | | 1del | |
| ACTCTCTGTGGCTTCACCAACTATTTT | 372 | 16A 18-21CAAC | | type3’ |
| ACTCTCTGTGGCTTCACCAATTATTTT | 338 | 16A 18-21CAAT | | type3’ |
| ACTCTCTATGGCTTCACCTATTATTTT | 178 | 8A 16A 18-21CTAT | | type3’ |
| ACTCTCTATGGCTTCACCAATTATCTTT | 128 | 8A 16A 18-21CAAT | 25C | type3’ |

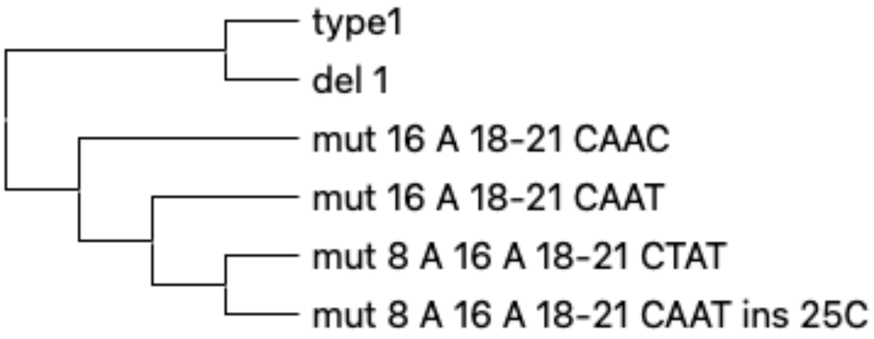


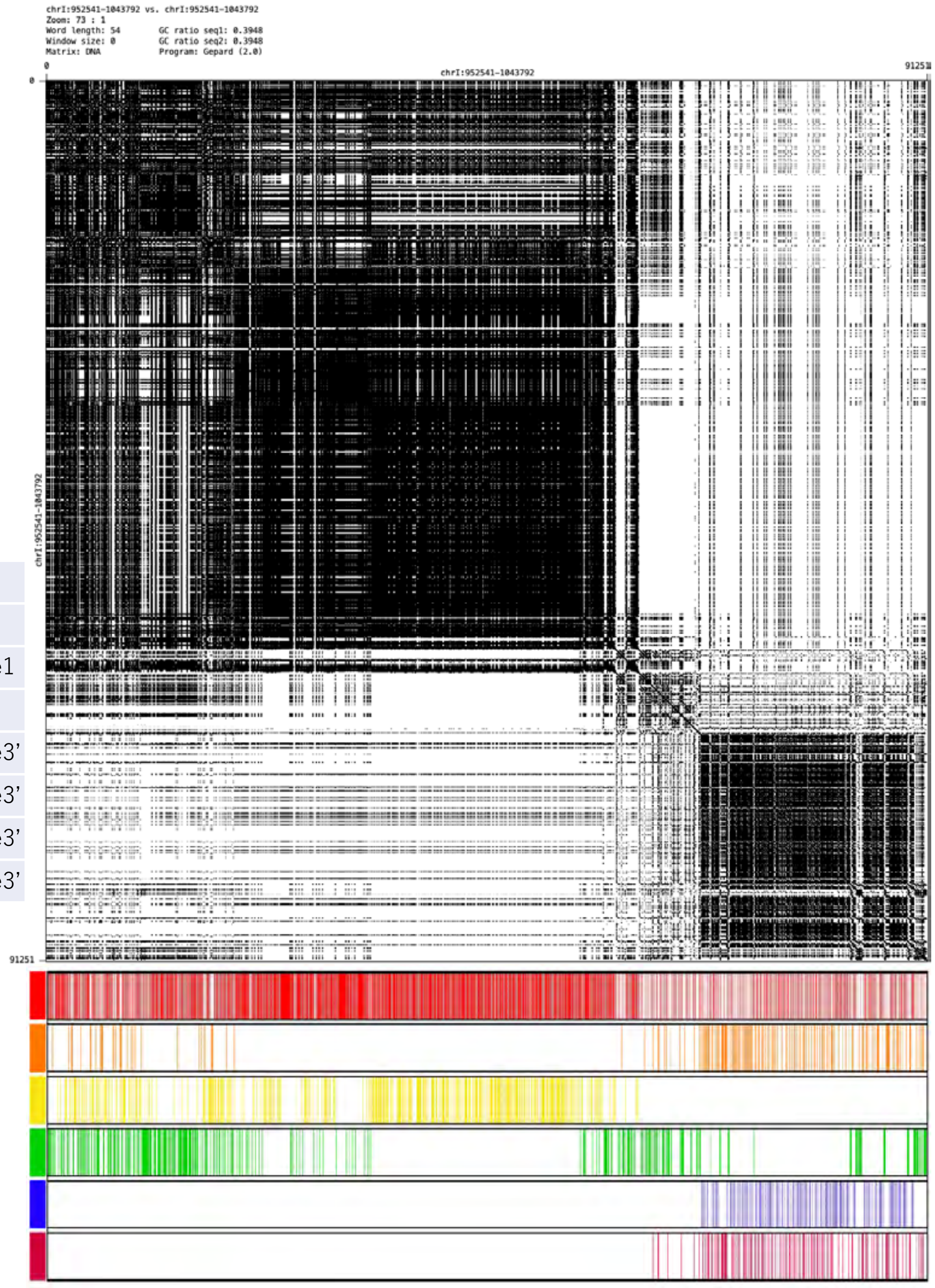

### chrX: 122,781-250,556

| chrX_1 | | | | |
|---|---|---|---|---|
| Representative unit and its variant | Freq. | mutation | indel | |
| ACTCTCTGTGGCTTCCCACTATATTTT | 2714 | | | type1 |
| ACTCTCTGTGGCTTCCCACTATGTTTT | 108 | 23G | | |
| ACTTTCTGTGGCTTCCCACTATATTTT | 192 | 4T | | |
| ACTCTCTGTGGCTTCATCGATTATTTT | 1230 | 16-21ATCGAT | | type3’ |
| ACTTTCTGTGGCTTCATCGATTATTTT | 117 | 4T 16-21ATCGAT | | type3’ |
| ACTCTCTGTGGCTTCATCGTTTATTTT | 52 | 16-19ATCG 21T | | type3’ |

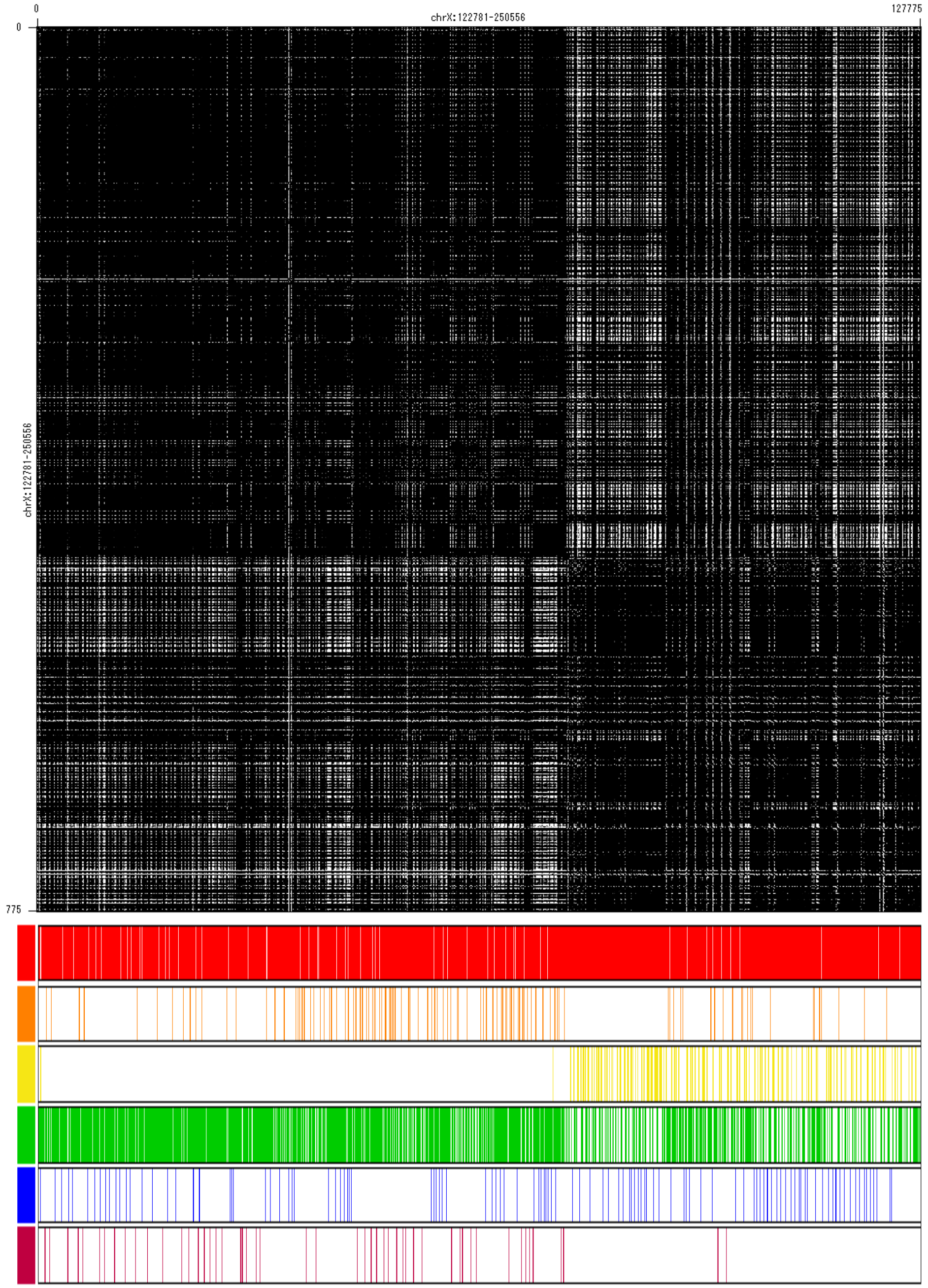


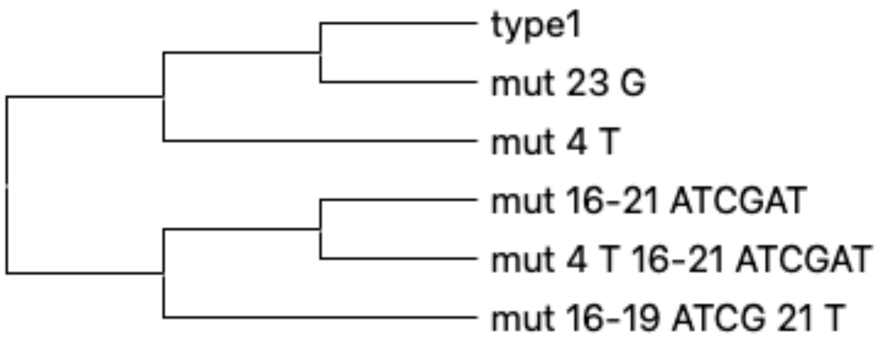

### chrI:4,745,128-4,817,023

| chrI_2 | | | | |
|---|---|---|---|---|
| Representative unit and its variant | Freq. | mutation | indel | |
| ACTCTCTGTGGCTTCCCACTATATTTT | 1072 | | | type1 |
| ACTCTCTGTAGCTTCCCACTATATTTT | 259 | 10A | | |
| ACTCTCTGTGGCTTCACCGATTATTTT | 315 | 16A 18-21CGAT | | type3 |
| ACTCTCTGTGGCTTCACCGATTA-TTT | 146 | 16A 18-21CGAT | 24del | type3’ |
| ACTCTCTGTGGCTTCACCGATGATTTT | 279 | 16A 18-22CGATG | | type3’ |
| ACTCTCTGTGGATTCACCGATTATTTT | 419 | 12A 16A 18-21CGAT | | type3’ |

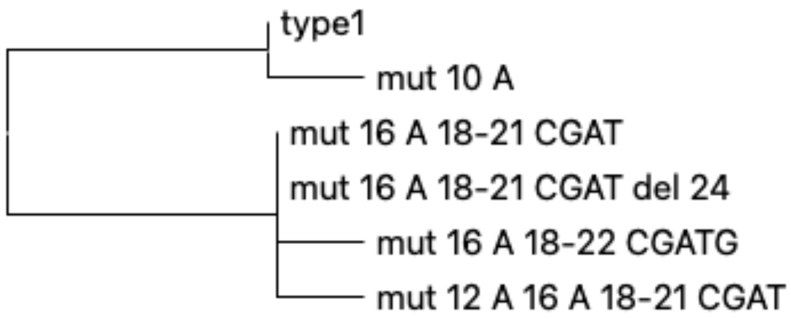


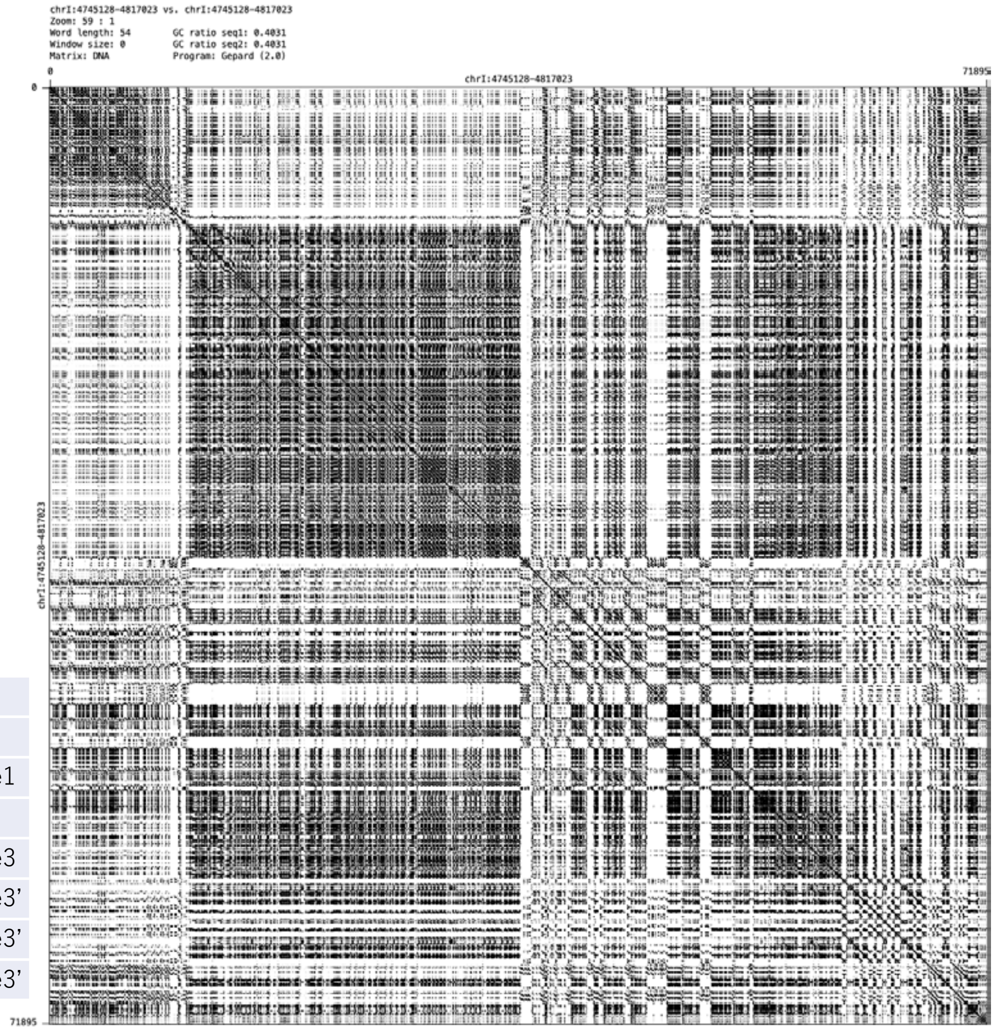


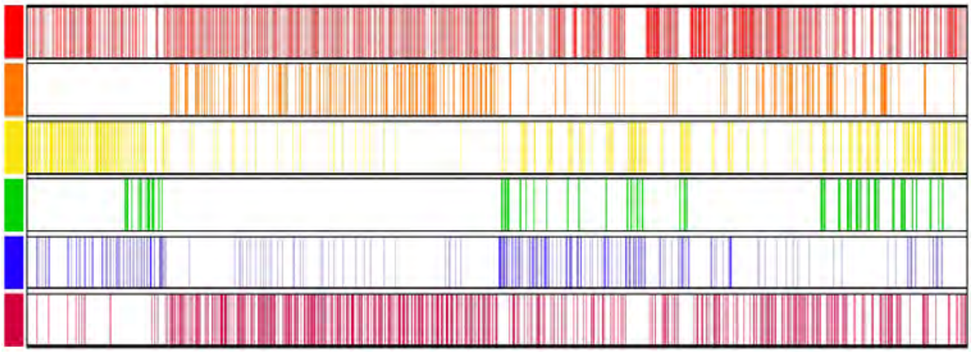

### chrX:5,608,553-5,673,280

| chrX_3 | | | | |
|---|---|---|---|---|
| Representative unit and its variant | Freq. | mutation | indel | |
| ACTCTCTGTGGCTTCCCACTATATTTT | 1188 | | | type1 |
| ACTCTCT--GGCTTCCCACTATATTTT | 607 | | del 8-9 | |
| ACTCTCTCTGGCTTCCCACTATATTTT | 16 | 8C | | |
| ACTCTATGTGGCTTCCCACTATATTTT | 489 | 6A | | |
| ACTCTATGTGGCTTTCCACTATATTTT | 32 | 6A 15T | | |
| ACTCTCTGTGGCTTCACCGATTATTTT | 51 | 16A 18-21 CGAT | | type3’ |

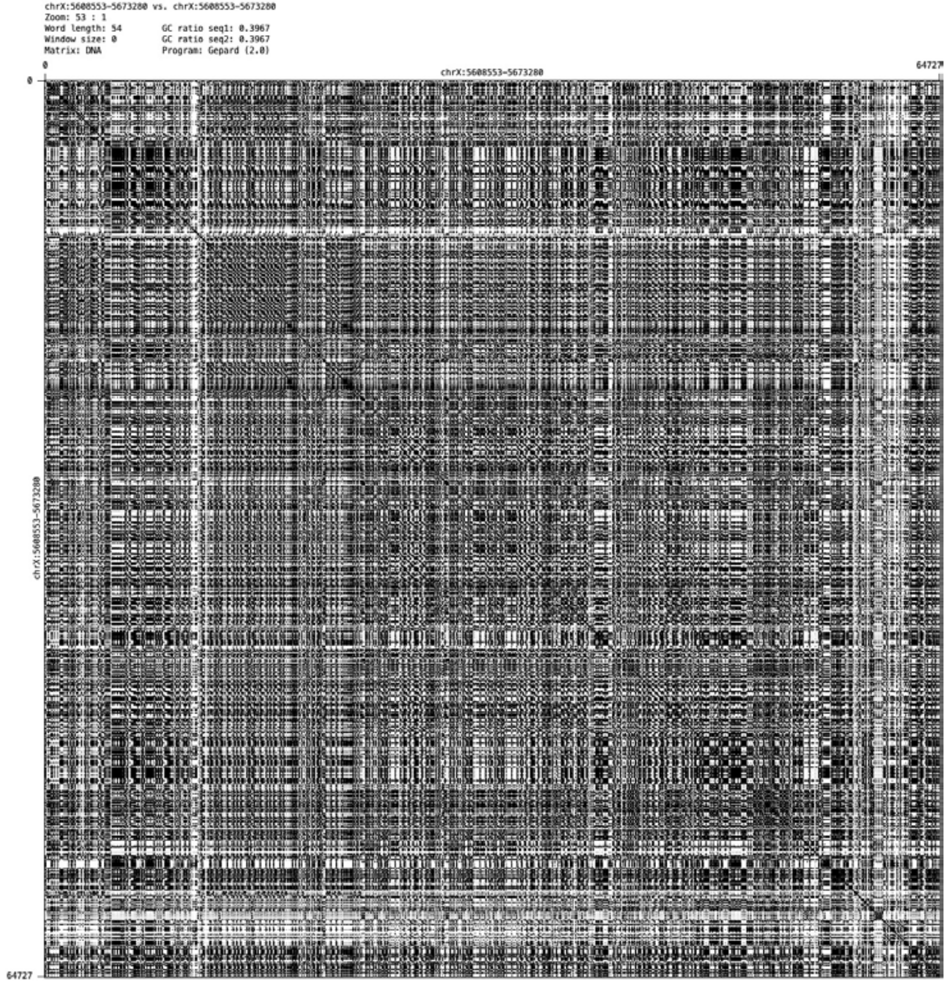


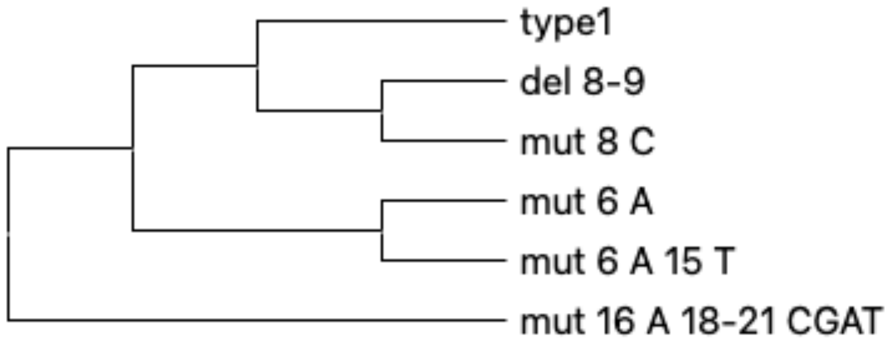


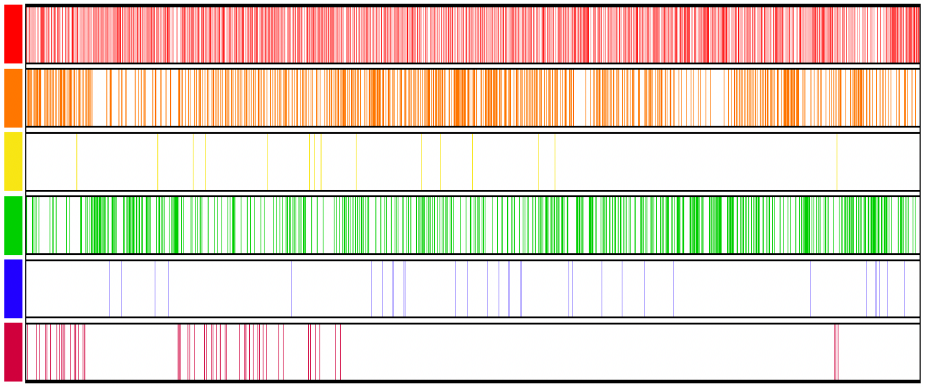

### chrIII:960,210-1,010,857

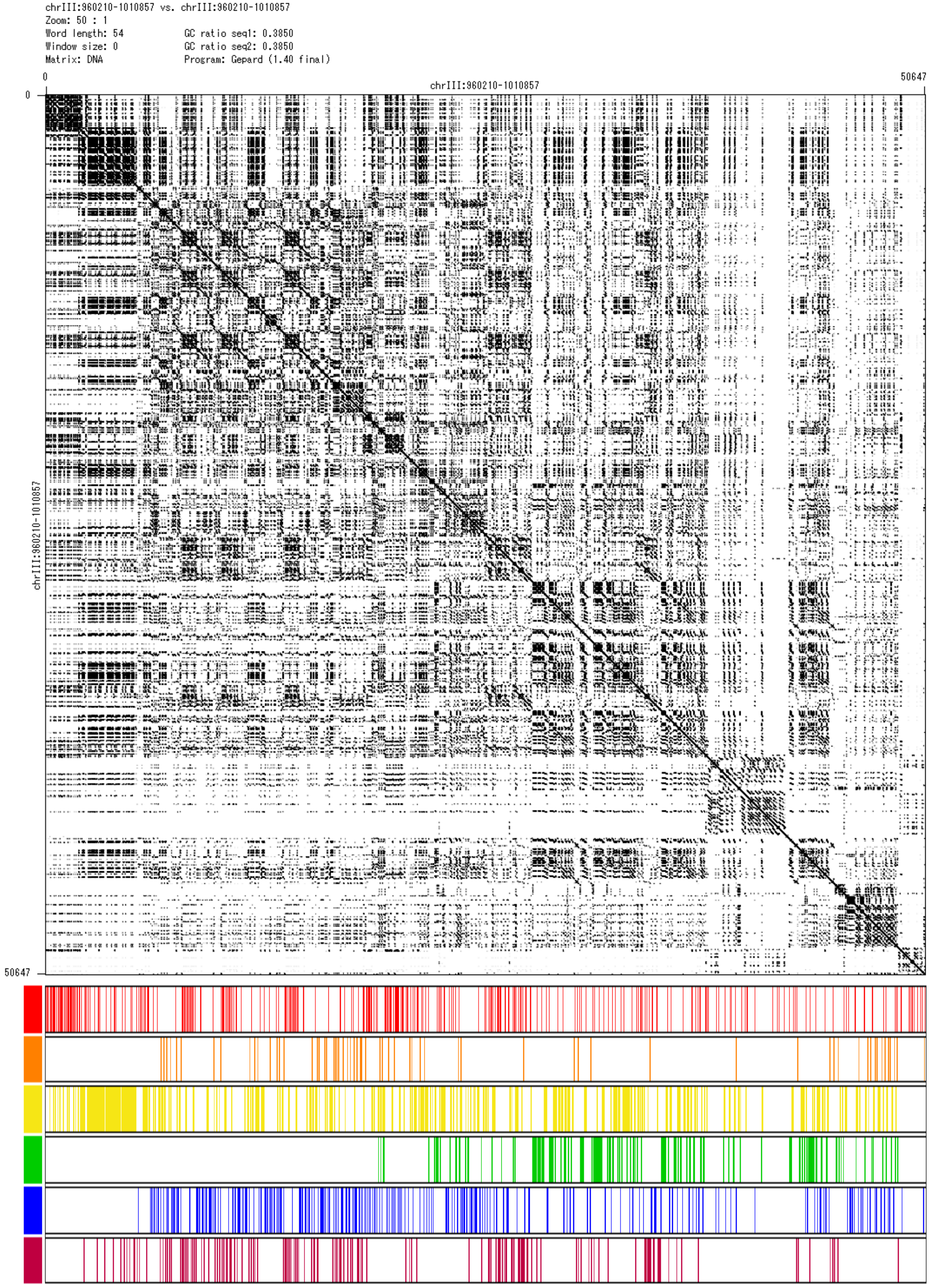


| chrIII_1 | | | | |
|---|---|---|---|---|
| Representative unit and its variant | Freq. | mutation | indel | |
| ACTCTCTGTGGCTTCCCACTATATTTT | 340 | | | type1 |
| ACTCTCTGTGGCTTCTCACCATATTTT | 64 | 16T 20C | | |
| ACTCTCTGTGGCTTCCCAATATATTTT | 421 | 19A | | |
| AATCTCTGTGGCTTCCCAATATATTTT | 178 | 2A 19A | | |
| ACTCTCTCTGGCTTCCCACTATATTTT | 337 | 8C | | |
| ACTCTCTCTGGCTTCCCAATATATTTT | 147 | 8C 19A | | |

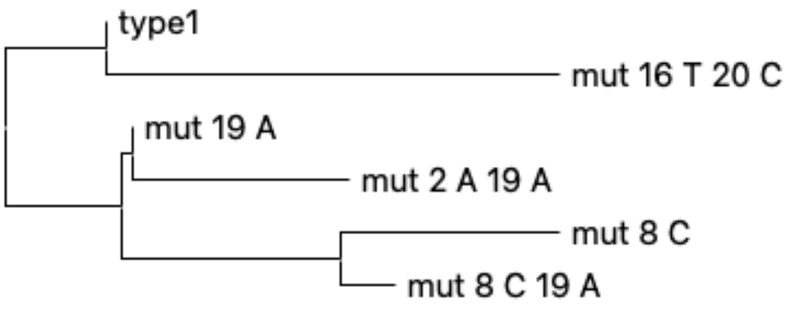

### chrII:14,809,101-14,842,635

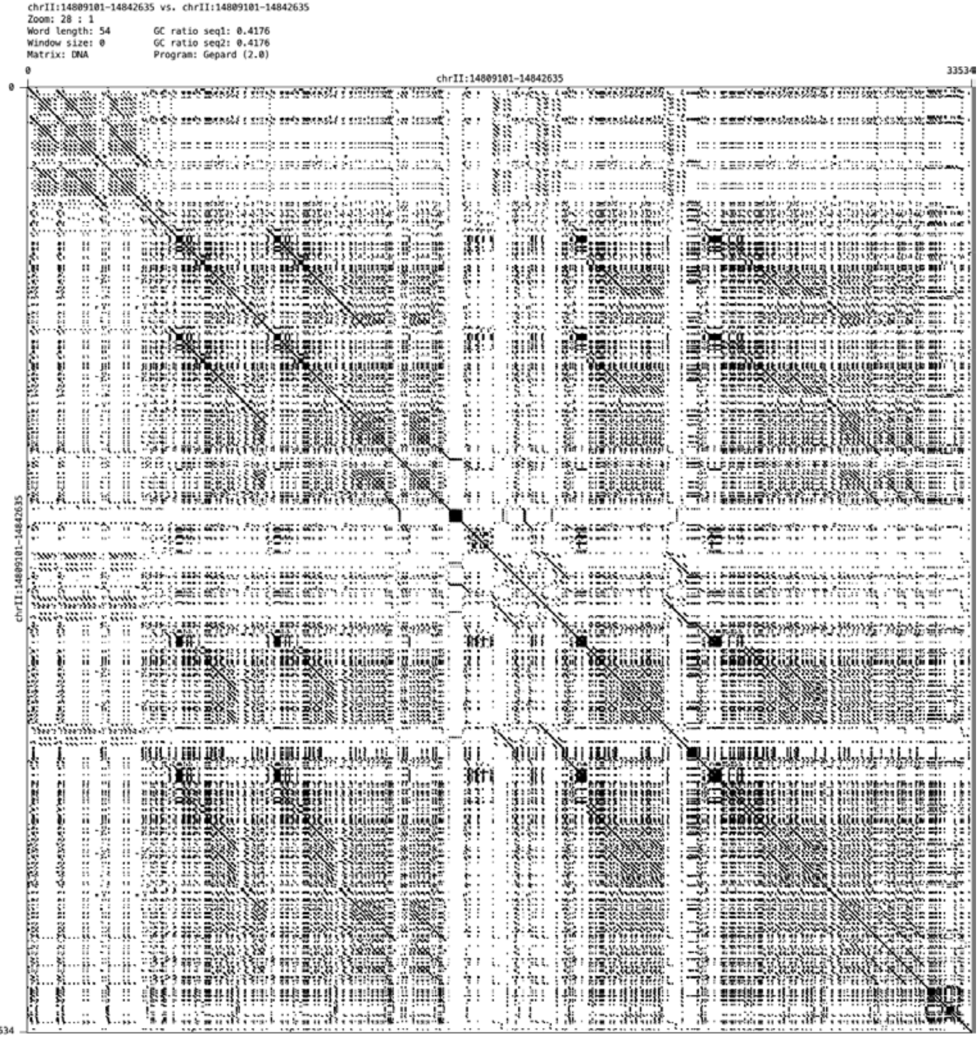


| chrII_1 | | | | |
|---|---|---|---|---|
| Representative unit and its variant | Freq. | mutation | indel | |
| ACTCTCTGTGGCTTCCCACTATATTTT | 227 | | | type1 |
| ACTCTCTGTGACTTCCCACTATATTTT | 61 | 11A | | |
| ACTCTCTTTGGCTTCCCACTATATTTT | 61 | 8T | | |
| ACTCTCTGTGGCTTCCCACCATATTTT | 396 | 20C | | type2 |
| ACTCT--GTGGCTTCCCACTATATTCT | 125 | 26C | 6-7del | |
| ACTCTCTGTGGCTTCTCACCATATTTT | 38 | 16T 20C | | |

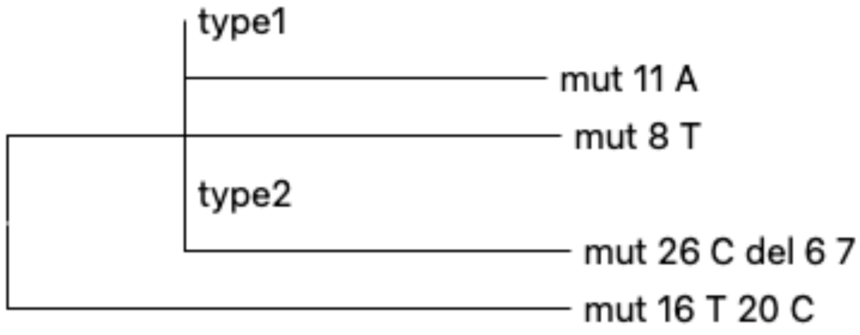


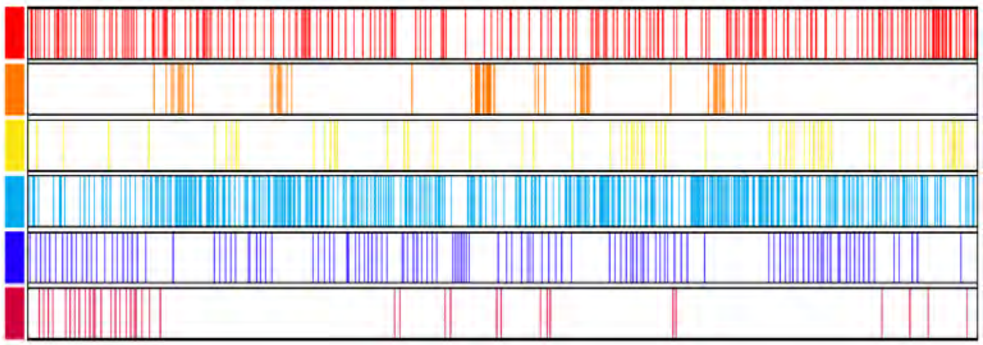

### chrIV: 9,333,371-9,370,581

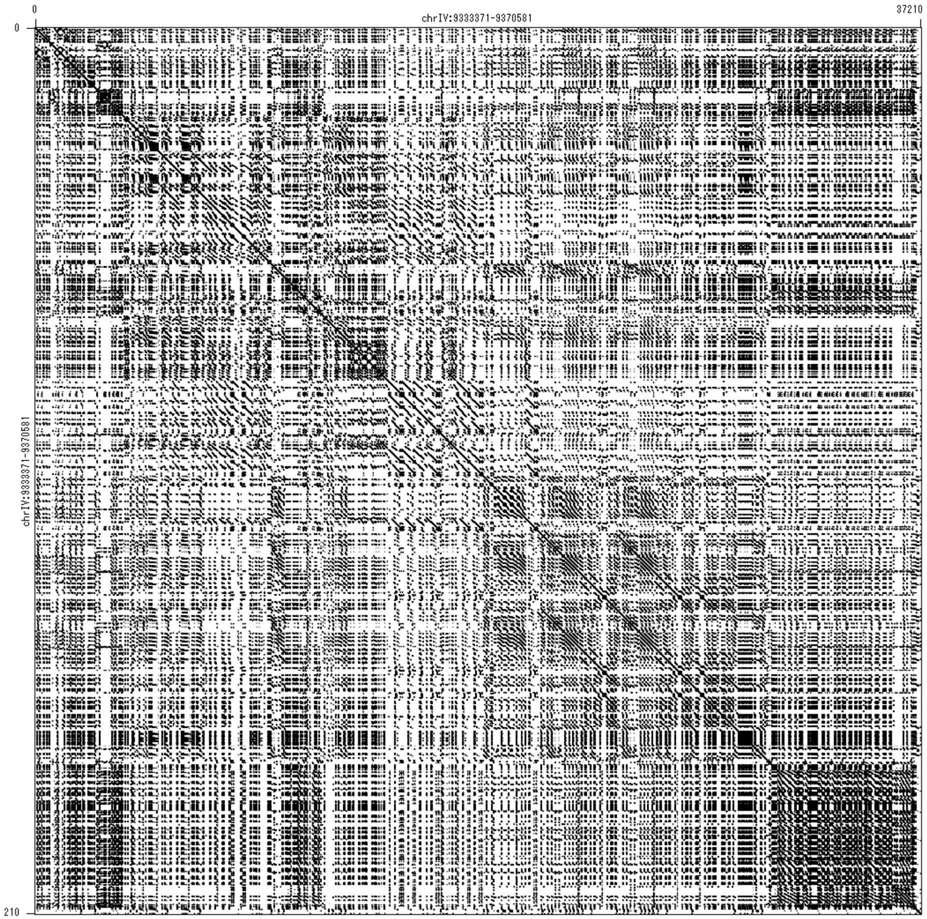


| chrIV_2 | | | | |
|---|---|---|---|---|
| Representative unit and its variant | Freq. | mutation | indel | |
| ACTCTCTGTGGCTTCCCACTATATTTT | 101 | | | type1 |
| ACTCTCTGTGGCTTCCCACCATATTTT | 230 | 20C | | type2 |
| ACTCTCTGTGACTTCCCACCATATTTT | 93 | 11A 20C | | |
| ACTCTCTGTGACTTCCCACTATTATTT | 106 | 11A 23-24TA | | |
| ACTCTCTGTGGCTTCCCACTATTATTT | 587 | 23-24TA | | |
| ACTCTCTTTGGCTTCCCACTACTATTT | 77 | 8T 22-24CTA | | |

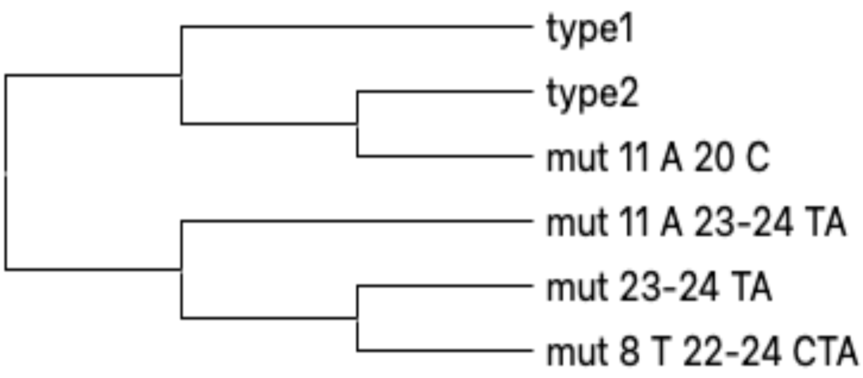


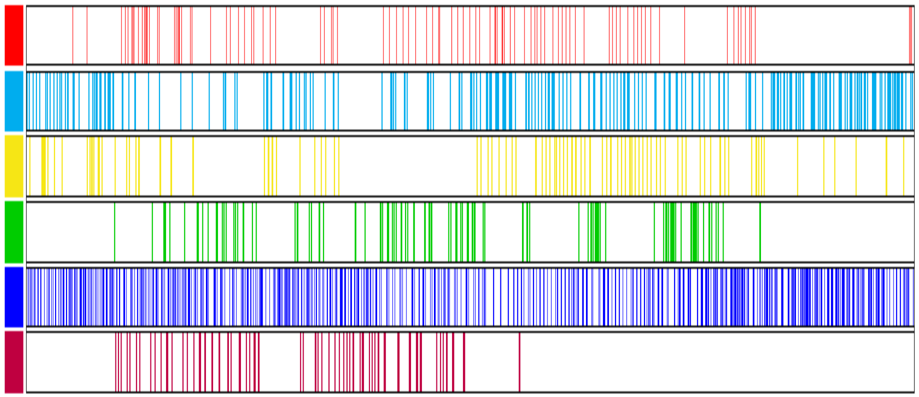

### chrIII: 7,856,863-7,950,893

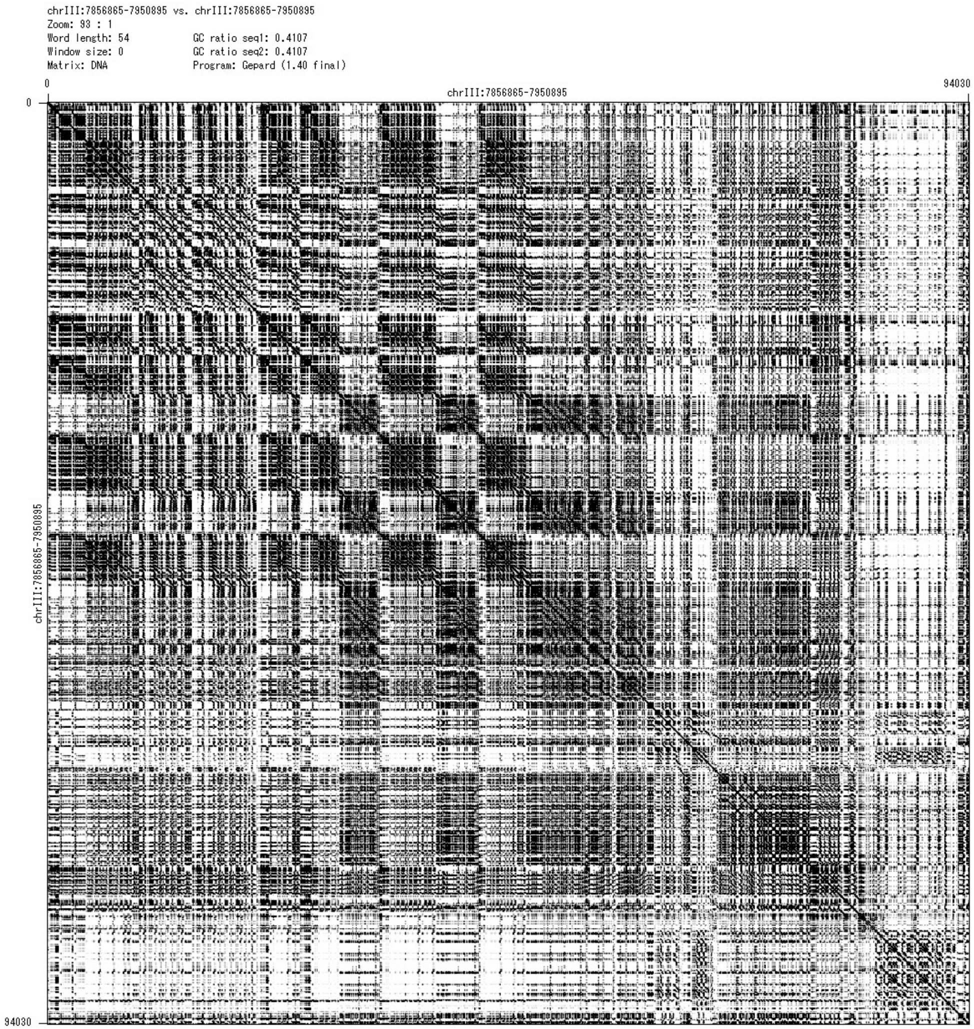


| chrIII_2 | | | | |
|---|---|---|---|---|
| Representative unit and its variant | Freq. | mutation | indel | |
| ACTCTCTGTGGCTTCCCACTATATTTT | 971 | | | type1 |
| ACTCTCTGTGGCTTCCCACCATATTTT | 1201 | 20C | | type2 |
| ACTCTCTGTGACTTCCCACCATATTTT | 139 | 11A 20C | | |
| ACTCTCTGTGGCTTCCCACTATTTTTT | 441 | 23T | | |
| ACTCTCTGTGGCTTCTCACTATTTTTT | 103 | 16T 23T | | |
| ACTATCTGTGGCTTCCCACTATTTTTT | 57 | 4T 23T | | |

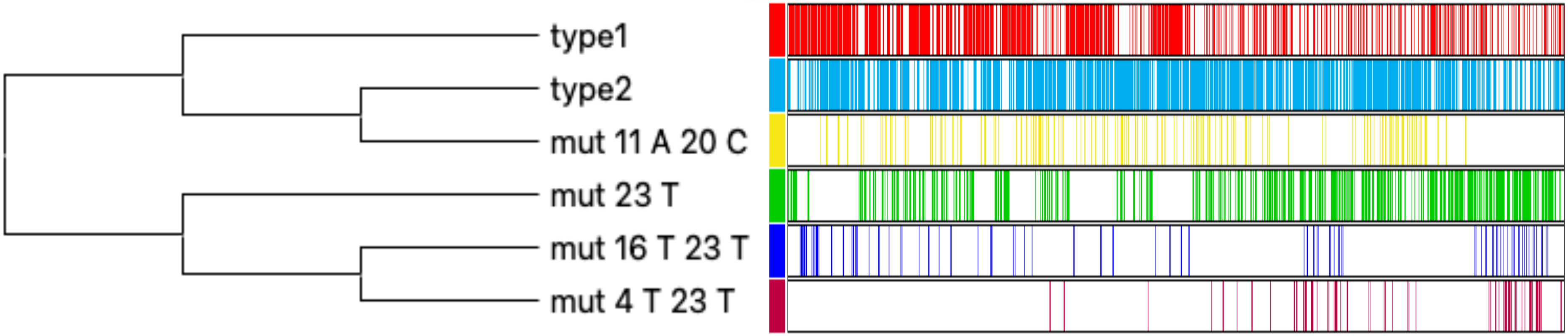

### chrV: 15,292,245-15,328,892

| chrV_2 | | | | |
|---|---|---|---|---|
| Representative unit and its variant | Freq. | mutation | indel | |
| ACTCTCTGTGGCTTCCCACTATATTTT | 169 | | | type1 |
| ACTCTCTGTGGCTTCCAACCATATTTT | 254 | 17A 20C 22T | | |
| ACTCTCTGTGGCTTCCAACTATATTTT | 99 | 16-17 CA | | |
| ACTCTCTGTGGCTTCCCACCATATTTT | 525 | 20C | | type2 |
| ACTCTCTGTGGCTTCCCACCATATTCT | 98 | 20C 26C | | |
| ACTCCCTGTGGCTTCCCACCATATTTT | 52 | 5C 20C | | |

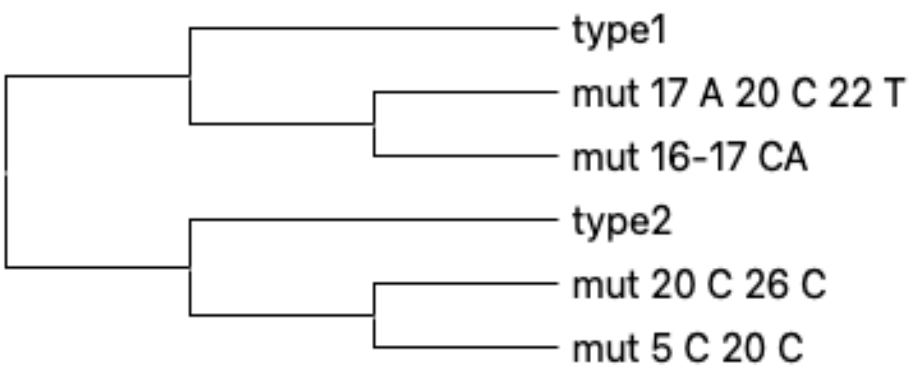


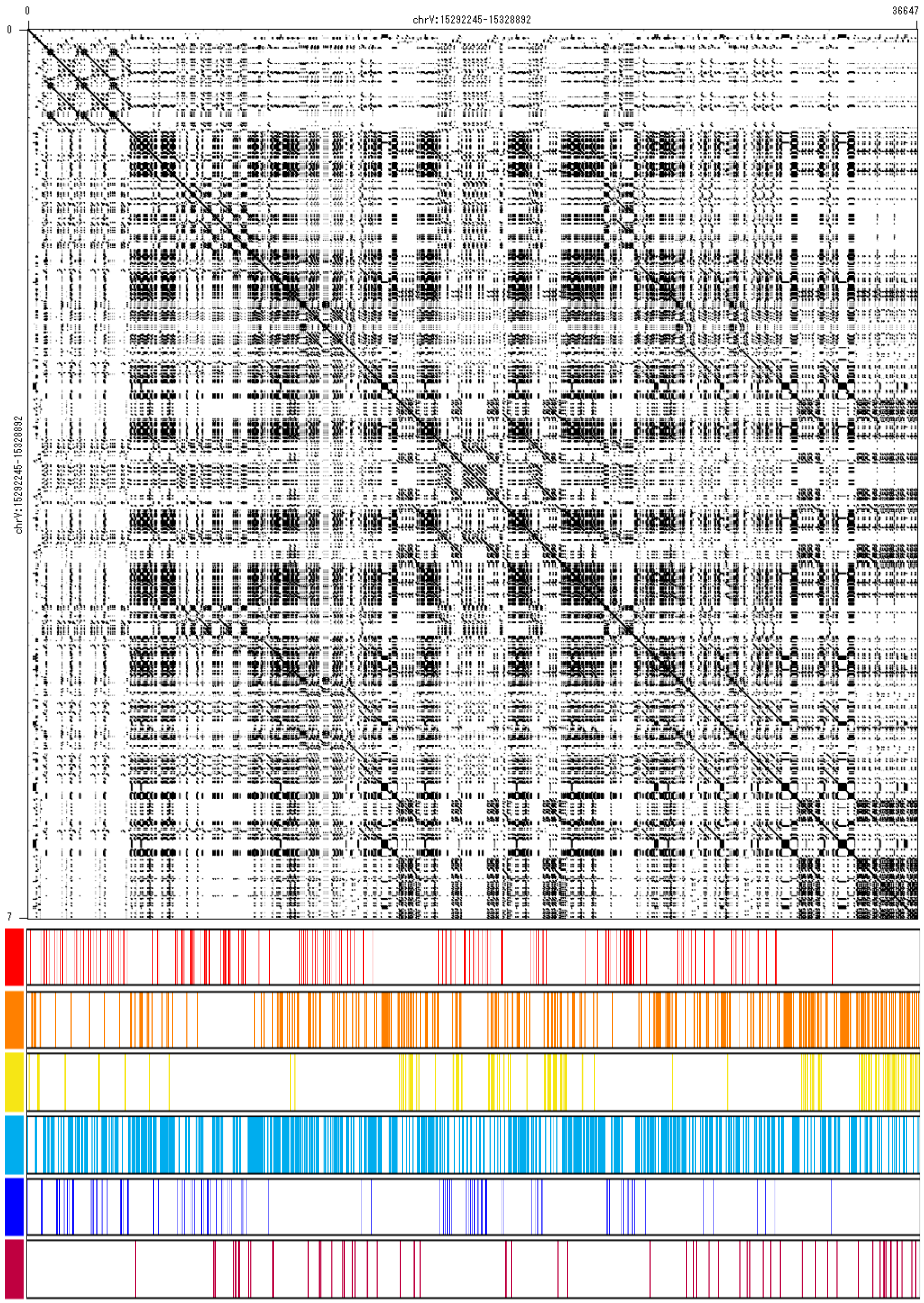

### chrX: 2,074,379-2,088,168

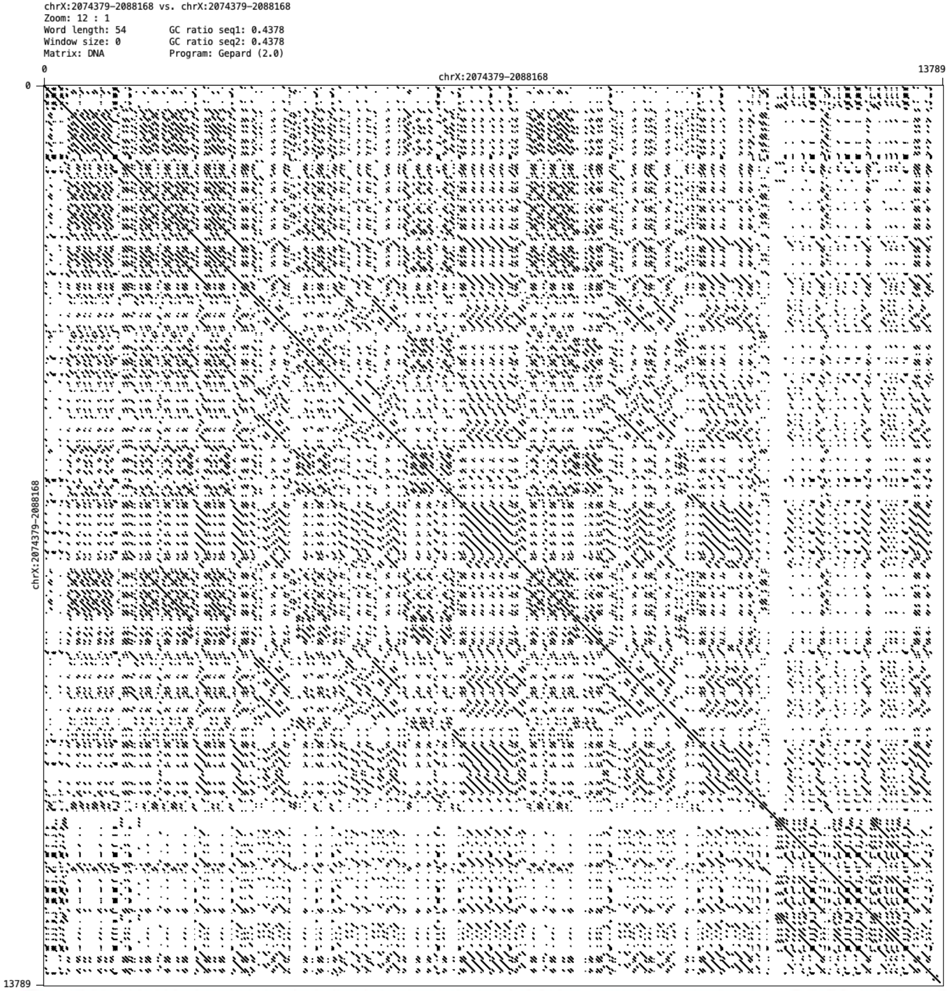


| chrX_2 | | | | |
|---|---|---|---|---|
| Representative unit and its variant | Freq. | mutation | indel | |
| ACTCTCTGTGGCTTCCCACTATATTTT | 161 | | | type1 |
| ACTCTCTGTGGCATCCCACTATATTTT | 53 | 13A | | |
| ACTCTCTGTGGCTTCCCACCATATTTT | 161 | 20C | | type2 |
| ACTCTCTGTGGCTTCCCACCATAGTTT | 69 | 20C 24G | | |
| ACTCTCTGTGGCTTCCCACCATATGTT | 55 | 20C 25G | | |

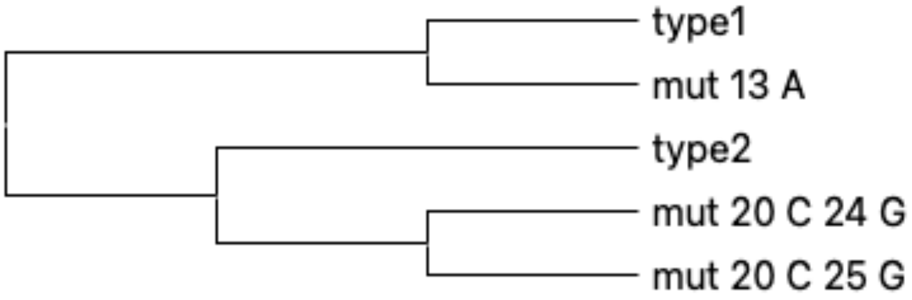


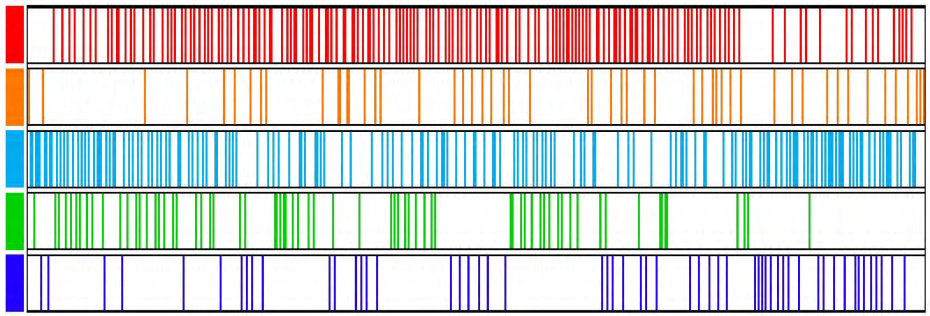
