## Supplementary Figure 7 for "CGC1, a new reference genome for *Caenorhabditis elegans*"

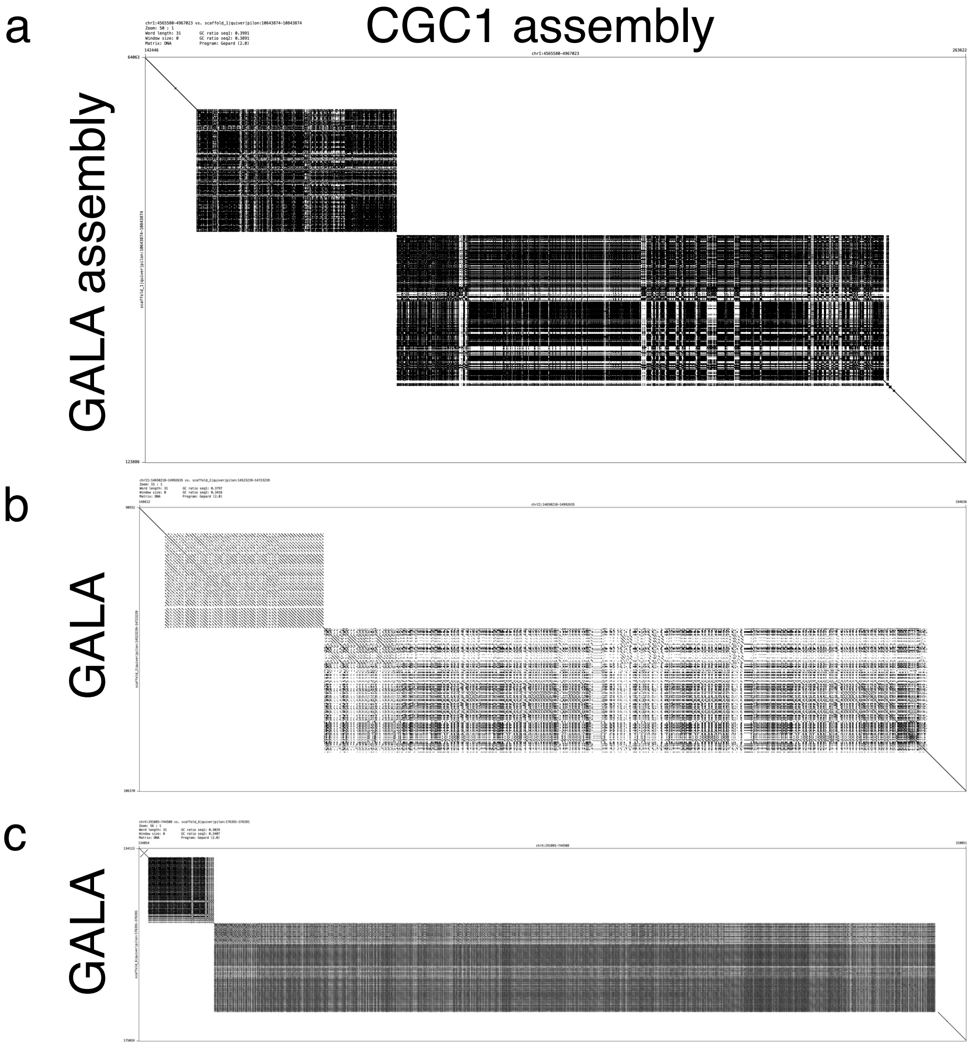

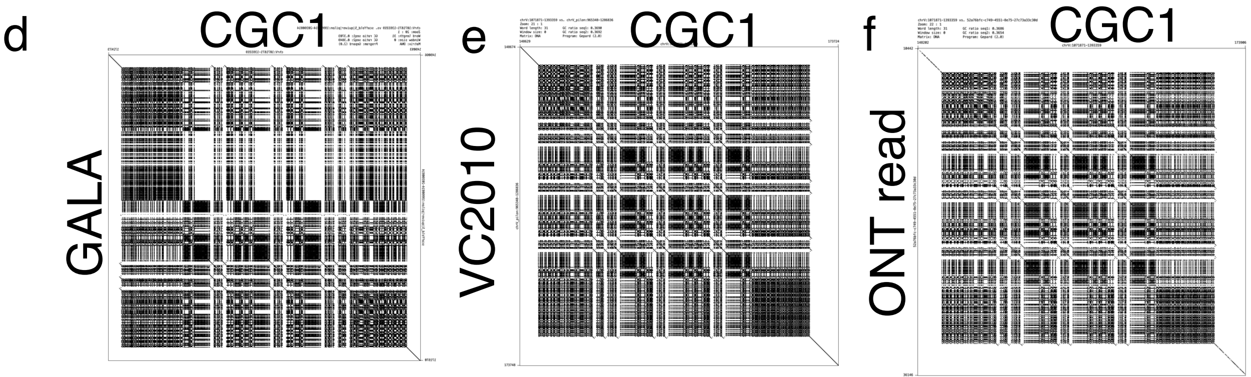


**Supplementary Figure 7. Comparison between four CGC1 genomic regions with large tandem repeats, the lengths of which are confirmed by Nanopore ultralong reads, and their corresponding regions in the GALA assembly. a**. A dot plot of the CGC1 region (chrI:4,715,580-4,817,023) and its corresponding region in the GALA assembly. The CGC1 region has two TRs of ~29 kb and ~72 kb in size; however, the GALA region has shorter TRs of ~18 kb and ~22 kb in size. **b-c**. Similarly, here are dot plots of the two CGC1 regions, chrII:14,800,218-14,842,635 (**b**) and chrX:441,085-594,508 (**c**), and their corresponding regions in the GALA assembly. In both cases, TRs in CGC1 are longer than those in GALA. Specifically, in Figure **b**, TRs of CGC1 are of length ~9 kb and ~33 kb, while TRs in GALA are ~5 kb and ~7 kb in size. In the bottom-right TRs of Figure 1**c**, TRs of CGC1 and GALA are of length ~153 kb and ~19 kb, respectively. **d-f**. A dot plot of the CGC1 region (chrV:1,221,871-1,243,359) and its corresponding region in the GALA assembly (**d**). The sizes of the TRs in the two regions are nearly equal, but the white areas show local differences between the two TRs. To show the validity of the CGC1 region, two dot plots are displayed in Figures **e** and **f**, demonstrating that the CGC1 region are consistent with its corresponding regions in VC2010 (**e**) and in a Nanopore ultralong read (**f**).
