## Supplementary Figure 8 for "CGC1, a new reference genome for *Caenorhabditis elegans*"

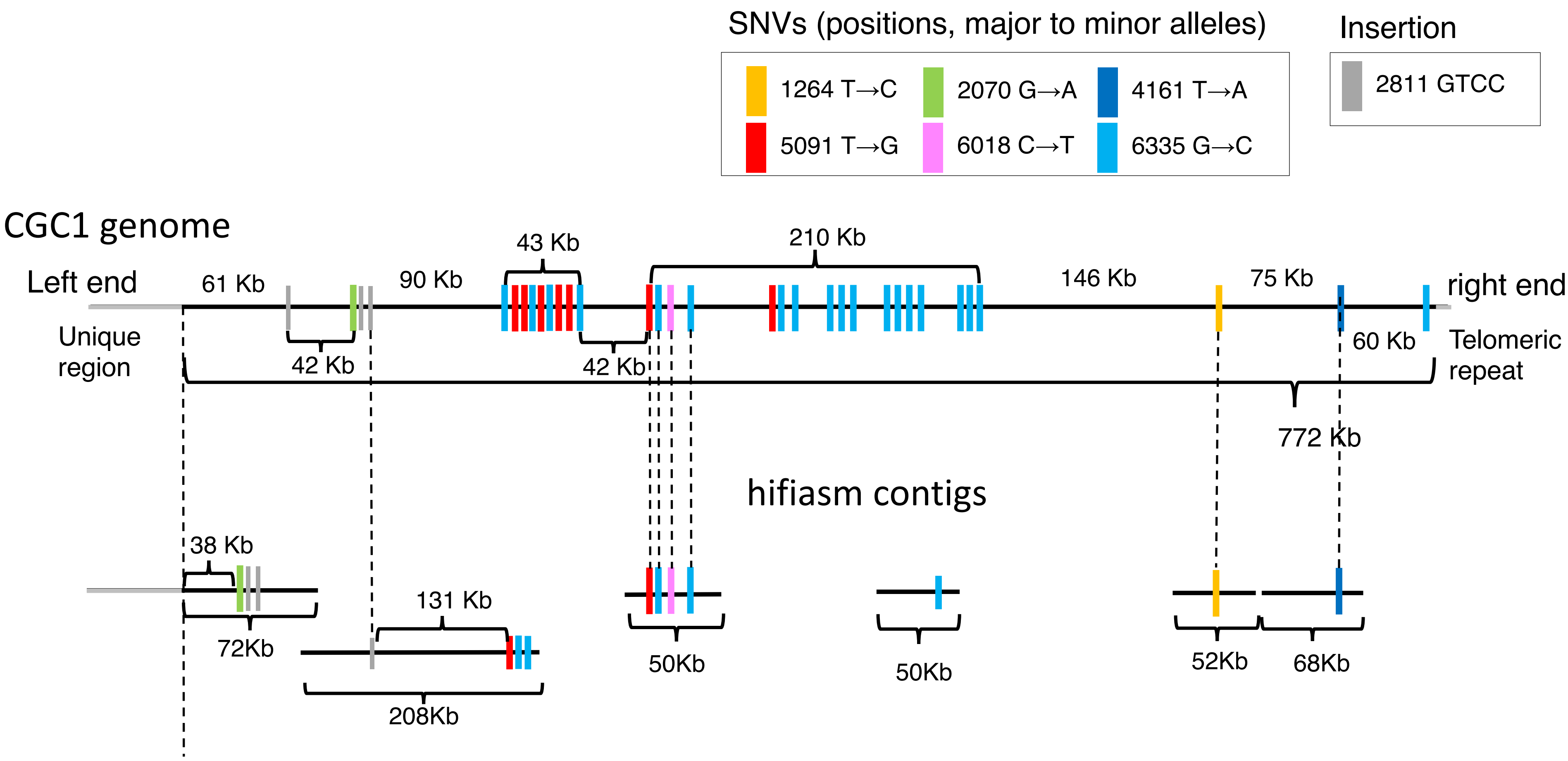


**Supplementary Figure 8. Fragmented assembled contigs in the hifiasm assembly in the 45S rDNA array.** The six assembled contigs generated by hifiasm 0.14 are shown at the bottom, and their approximate positions in the rDNA array of the CGC1 genome assembly are indicated by the dotted lines that link between SNVs specific to representative repeat units of 45S rDNA. These six contigs are much shorter than the rDNA array.
