## Supplementary Figure 9 for "CGC1, a new reference genome for *Caenorhabditis elegans*"

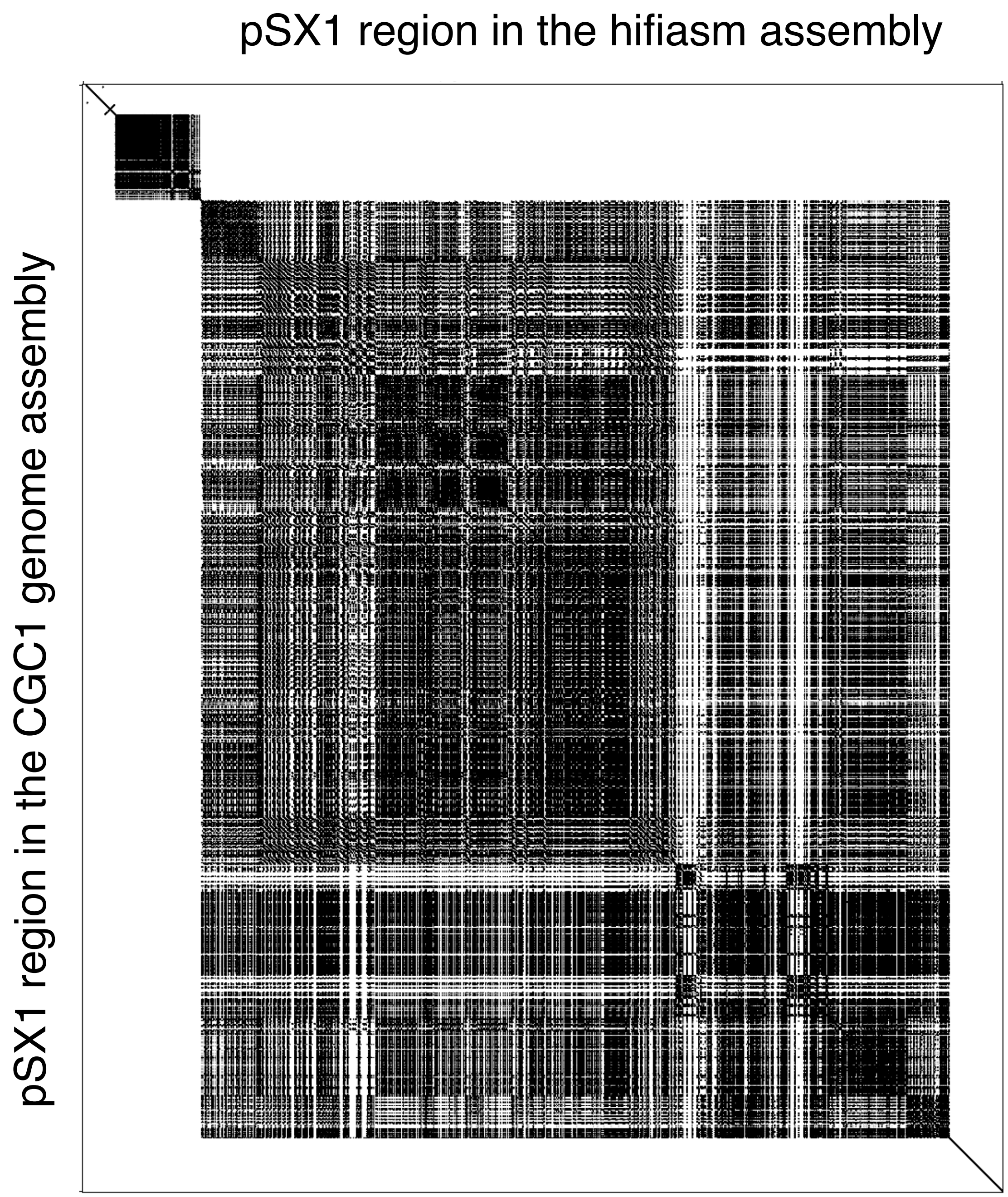


**Supplementary Figure 9. Dot plot between the pSX1 region of the hifiasm assembly (horizontal axis) and the CGC1 genome assembly (vertical axis).**
