## Supplementary Figure 10 for "CGC1, a new reference genome for *Caenorhabditis elegans*"

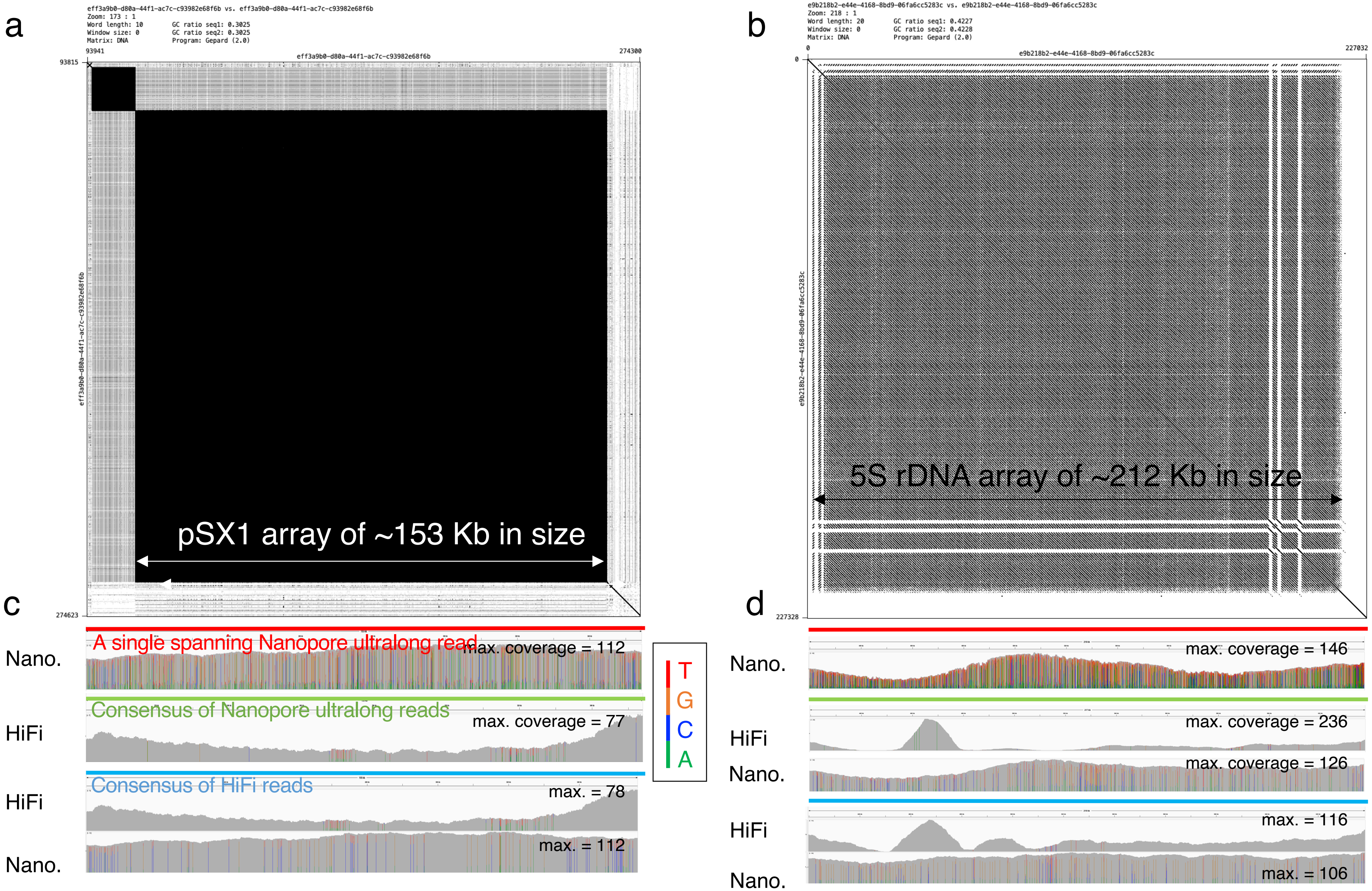


**Supplementary Figure 10. Error correction of regions that span the pSX1 and 5S rDNA arrays. a-b**. The top rows show the self dot plots of two Nanopore ultralong reads that span the pSX1 array in chromosome X (441,085-594,508) (**a**) and 5S rDNA array in chromosome V (17,811,675-18,028,889) (**b**). **c-d**. Rows from the top to the bottom show spanning Nanopore ultralong reads (red), read coverage of Nanopore ultralong reads that were aligned with the red spanning reads, the consensus of the aligned Nanopore reads (green), read coverage of PacBio HiFi reads mapped to the Nanopore consensus, read coverage of Nanopore reads to the Nanopore consensus in the right column, the consensus of HiFi reads mapped to the Nanopore consensus (blue), and read coverage of HiFi and Nanopore reads mapped to the HiFi consensus. Nucleotide distribution at each base is displayed.
