## Supplementary Figure 11 for "CGC1, a new reference genome for *Caenorhabditis elegans*"

**Supplementary Figure 11. Evolution in the 5S rDNA array.** Similar to Figure 4, the upper right panel shows the self-dot plot of a region with the 5S rDNA array of length ~212 kb in chromosome V (nt 17,811,675-18,028,889), the bottom-left table shows characteristics of four frequent variants of the reference 5S rDNA (976 nt, red) including the statistical significance codes in the last column for p-values (***<=0.1%, **<=1%, and *<=5%) such that each variant occurs adjacent to each other according to the Wald–Wolfowitz runs test. On the right side of the table, the colored bars indicate the locations of the each variant.

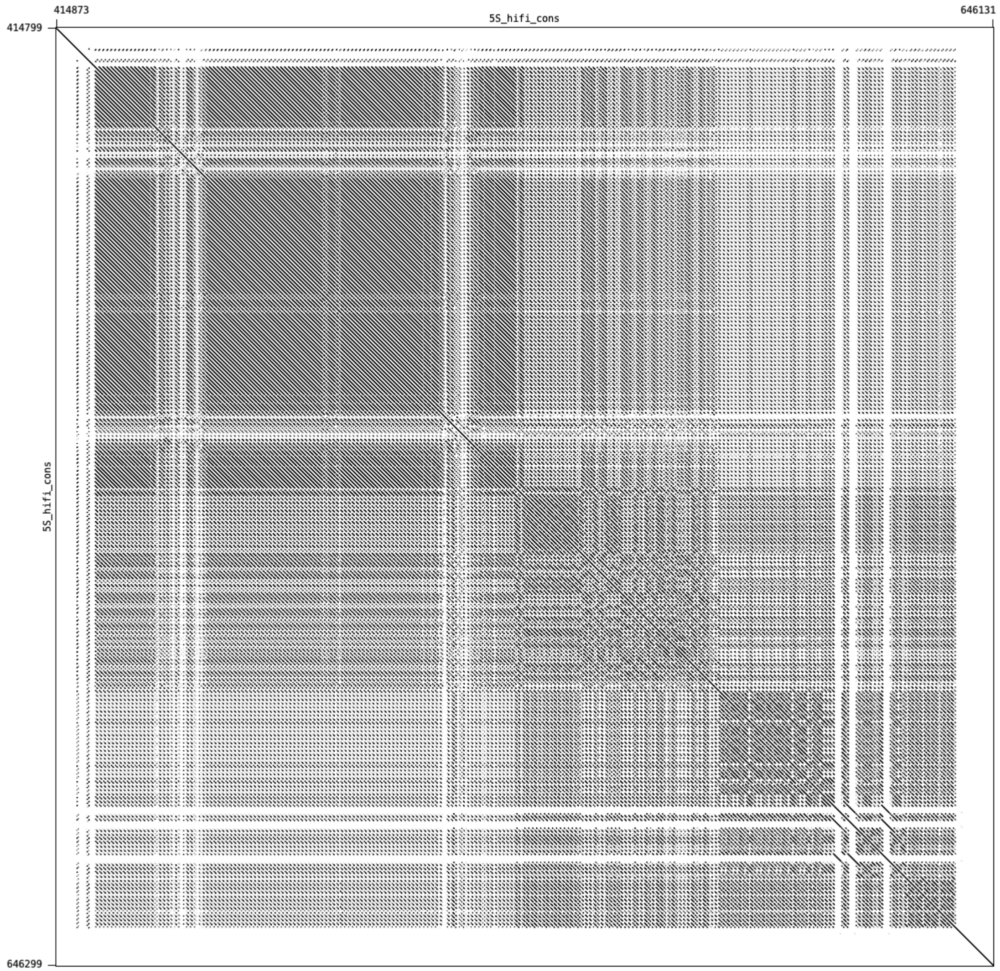


| Frequency | Ratio of frequency | # mis-matches | match ratio | substitutions | deletions | insertions | p-value |
|---|---|---|---|---|---|---|---|
| 73 | 36.5 | 0 | 100 | | | | *** |
| 12 | 6 | 9 | 99.08 | 3A,162G,431G,621G | 92-95 | 318A | *** |
| 8 | 4 | 9 | 99.08 | 162G | 92-95 | 302TTTT | |
| 8 | 4 | 9 | 99.08 | 162G,621G | 92-95 | 302TTT | *** |

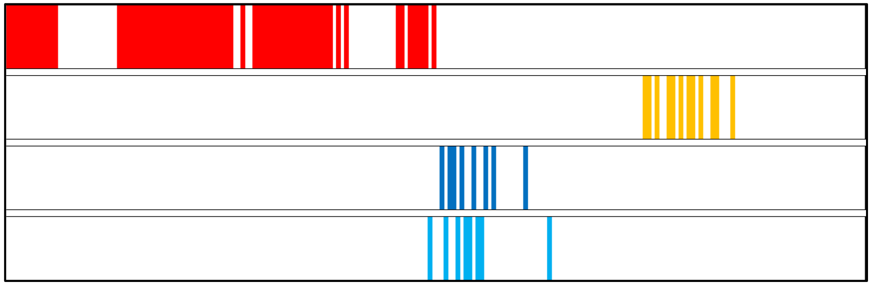
