## Supplementary Figure 12 for "CGC1, a new reference genome for *Caenorhabditis elegans*"

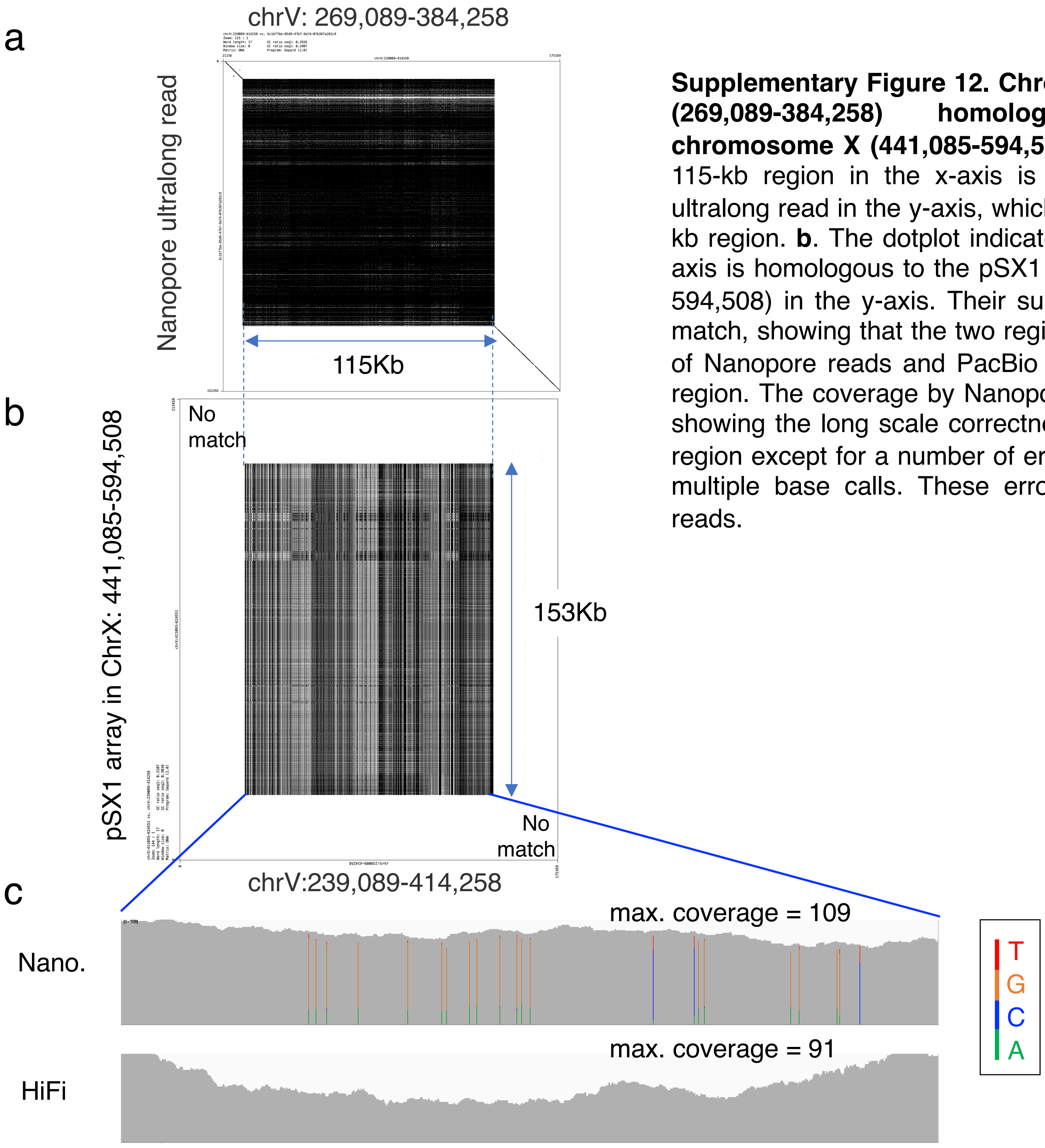


**Supplementary Figure 12. Chromosome V has a 115-kb region (269,089-384,258) homologous to the pSX1 array in chromosome X (441,085-594,508). a**. The dotplot shows that the 115-kb region in the x-axis is spanned by a 152-kb Nanopore ultralong read in the y-axis, which confirms the presence of the 115 kb region. **b**. The dotplot indicates that the 115 kb region in the x-axis is homologous to the pSX1 array in chromosome X (441,085-594,508) in the y-axis. Their surrounding regions however do not match, showing that the two regions are distinct. **c**. Read coverage of Nanopore reads and PacBio HiFi reads mapped to the 115 kb region. The coverage by Nanopore ultralong reads is almost even, showing the long scale correctness of the assembly in this 115-kb region except for a number of erroneous bases that are colored by multiple base calls. These errors are corrected by PacBio HiFi reads.
