## Supplementary Figure 13 for "CGC1, a new reference genome for *Caenorhabditis elegans*"

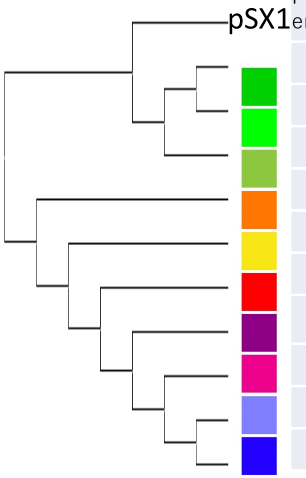


| Frequency | Ratio of Frequency | #mis-matches | Match ratio | Substitutions and deletions |
|---|---|---|---|---|
| 8 | 1.19 | 17 | 90.12 | 28A,45T,53T,94G,103A,104A,113C,117G,145A,146T,172T,7A,58T,80G,131T,159C |
| 10 | 1.49 | 15 | 91.28 | 28A,45T,53T,94G,103A,104A,113C,117G,145A,146T,172T,7A,131T,164del,165del |
| 20 | 2.98 | 15 | 91.28 | 28A,45T,53T,94G,103A,104A,113C,117G,145A,146T,172T,16A,131T,164del,165del |
| 7 | 1.04 | 21 | 87.79 | 28A,45T,53T,94G,103A,104A,113C,117G,145A,146T,172T,65T,68C,74A,89A,92A,100C,131T,132A,164del,168C |
| 16 | 2.38 | 18 | 89.53 | 28A,45T,53T,94G,103A,104A,113C,117G,145A,146T,172T,16A,58A,74A,92A,128G,164del,168C |
| 38 | 5.65 | 18 | 89.53 | 28A,45T,53T,94G,103A,104A,113C,117G,145A,146T,172T,7A,58A,74A,92A,128G,164del,168C |
| 7 | 1.04 | 24 | 86.05 | 28A,45T,53T,94G,103A,104A,113C,117G,145A,146T,172T,7A,33A,56T,72C,73C,74A,75C,92A,128G,157G,164del,168C,171T |
| 13 | 1.93 | 22 | 87.21 | 28A,45T,53T,94G,103A,104A,113C,117G,145A,146T,172T,7A,58A,65T,72C,73C,74A,75C,83C,92A,164del,168C |
| 14 | 2.08 | 23 | 86.63 | 28A,45T,53T,94G,103A,104A,113C,117G,145A,146T,172T,7A,58A,65T,72C,73C,74A,75C,83C,92A,131T,164del,165del |
| 21 | 3.13 | 23 | 86.63 | 28A,45T,53T,94G,103A,104A,113C,117G,145A,146T,172T,16A,58A,65T,72C,73C,74A,75C,83C,92A,131T,164del,165del |

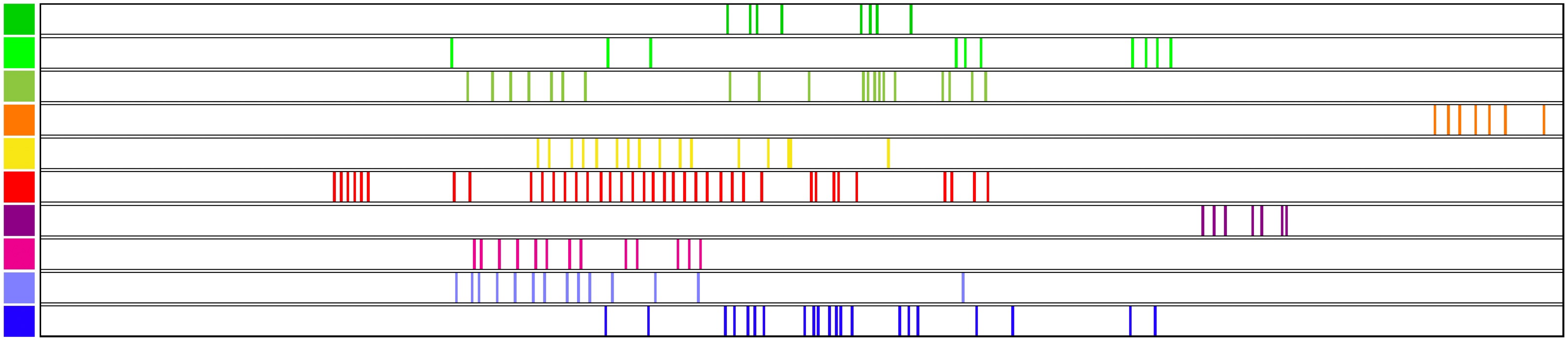

**Supplementary Figure 13. The pSX1 array in chrV:260,245-414,258.** The upper panel shows characteristics of ten frequent variants of the reference pSX1 (172 nt) in the top: the frequency of each variant, the ratio of frequency to total, number of mismatches with the reference pSX1, percentage match with the reference, positions of base substitutions (e.g., 28A means the base at position 28 is substituted with A), and positions with deletion bases (e.g., 164del shows the base at position 164 is deleted). Blue colored substitutions are found in all of the ten pSX1 variants. Note that the match ratios of the ten variants are lower than 92%. Most substitutions are shared in common among the variants, making it hard to find a variant unique to each variant. The phylogenetic tree on the left side of the table shows the proximity of variants in terms of sequence similarity. In the bottom table, the colored bars indicate the locations of the occurrences of each variant.
