## Supplementary Figure 14 for "CGC1, a new reference genome for *Caenorhabditis elegans*"

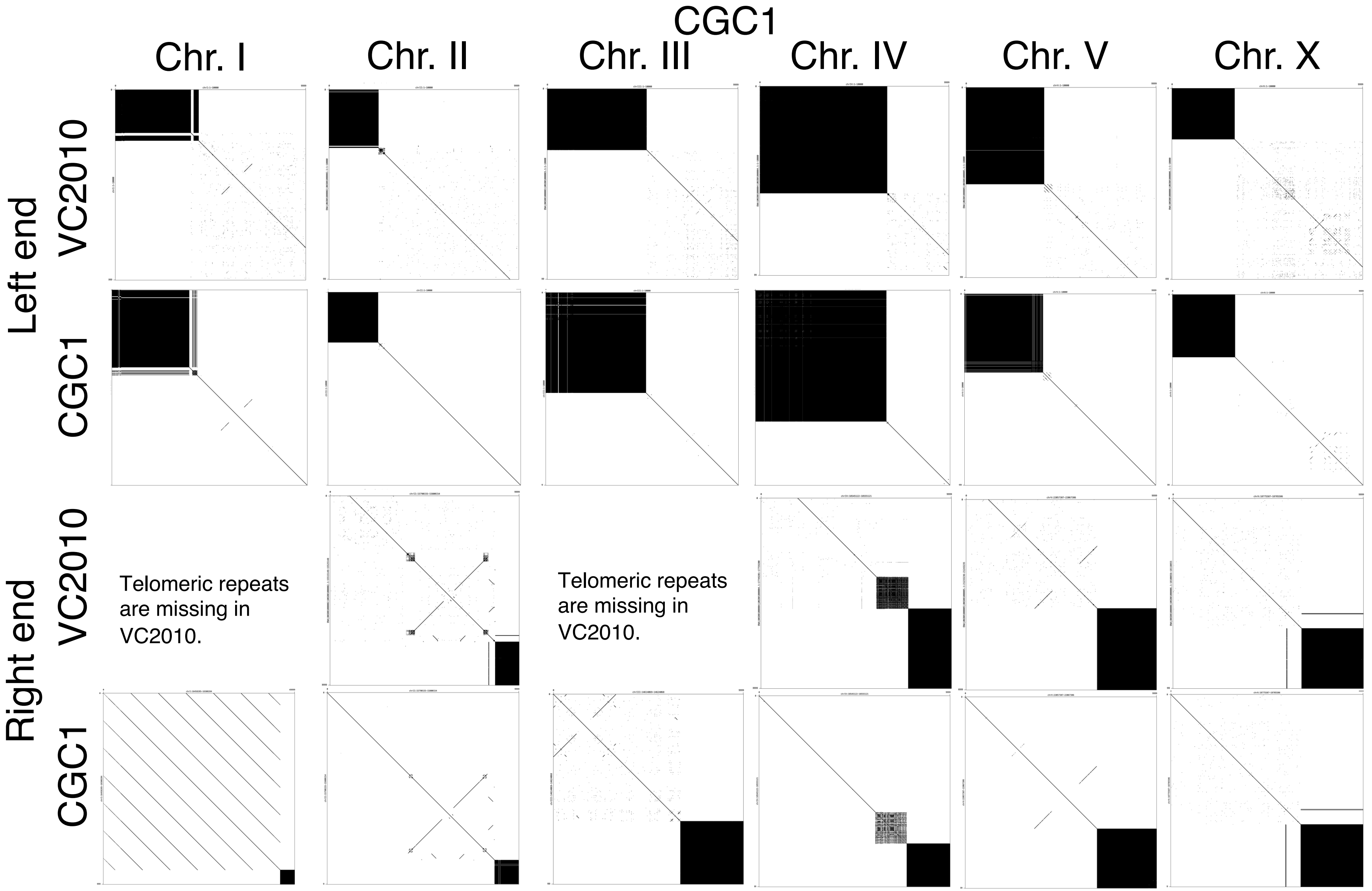


**Supplementary Figure 14. Dot plots between pairs of 10 kb genomic sequences taken from the left end (top two rows) and right end (bottom two rows) of each chromosome.** The horizontal axis shows 10 kb sequences from the CGC1 assembly, and the vertical axis shows alternating 10 kb sequences from the VC2010 (Yoshimura et al., 2019) and CGC1 assemblies. The black colored rectangle at the left end or right end of each dot plot displays telomeric repeats. The bottom left self-dot plot displays the 10 kb sequence of the right end of chromosome I, and the many lines parallel to the diagonal show the tail of the rDNA array. The VC2010 assembly lacks telomere repeats at the right ends of chromosomes I and III, and hence dot plots between the missing regions and the corresponding regions in the CGC1 assembly are not displayed in the matrix.
