## Supplementary Figure 15 for "CGC1, a new reference genome for *Caenorhabditis elegans*"

**Supplementary Figure 15. Transcription from tandem repeats from three loci in various tissue types.** Three instances are shown where full-length RNA-seq data was mapped to TRs in the CGC1 assembly and also showed splicing in the genomic vicinity of a TR (demonstrating that the RNA-seq reads represented actual RNA rather than contaminant genomic DNA). For all three instances, the top of the figure shows genome annotations, as follows: N2 denotes structures for protein-coding genes lifted over from N2 to CGC1; AUG denotes de novo predictions of protein-coding genes with AUGUSTUS; ST2 denotes StringTie2 transcripts (which can arise either from mRNA or from ncRNA); and TR denotes tandem repeats of CGC1. Below these genome annotations, detailed alignments of long RNA-seq reads to TRs and their flanking genomic DNA sequences are shown. The first three tracks show alignments of tissue-specific RNA-seq reads from Li et al. (Li *et al.* 2020) (GSE130044, NCBI Gene Expression Omnibus); six further tracks show alignments of tissue-specific RNA-seq reads from Roach et al. (Roach *et al.* 2020) (PRJEB31791, ENA). Supplementary Figures 15a, 15b, and 15c respectively display RNA-seq alignments with TRs in chrIII:12,198,925-12,210,304, chrX:16,263,528-16,273,490, and chrX:17,980,227-17,991,566. Supplementary Figures 15a-15b have their gene annotations to the same scale as the long RNA-seq alignments, while 15c has its gene annotations to smaller scale, as can be seen by comparing the upper TR annotation to the lower TR boundary lines.

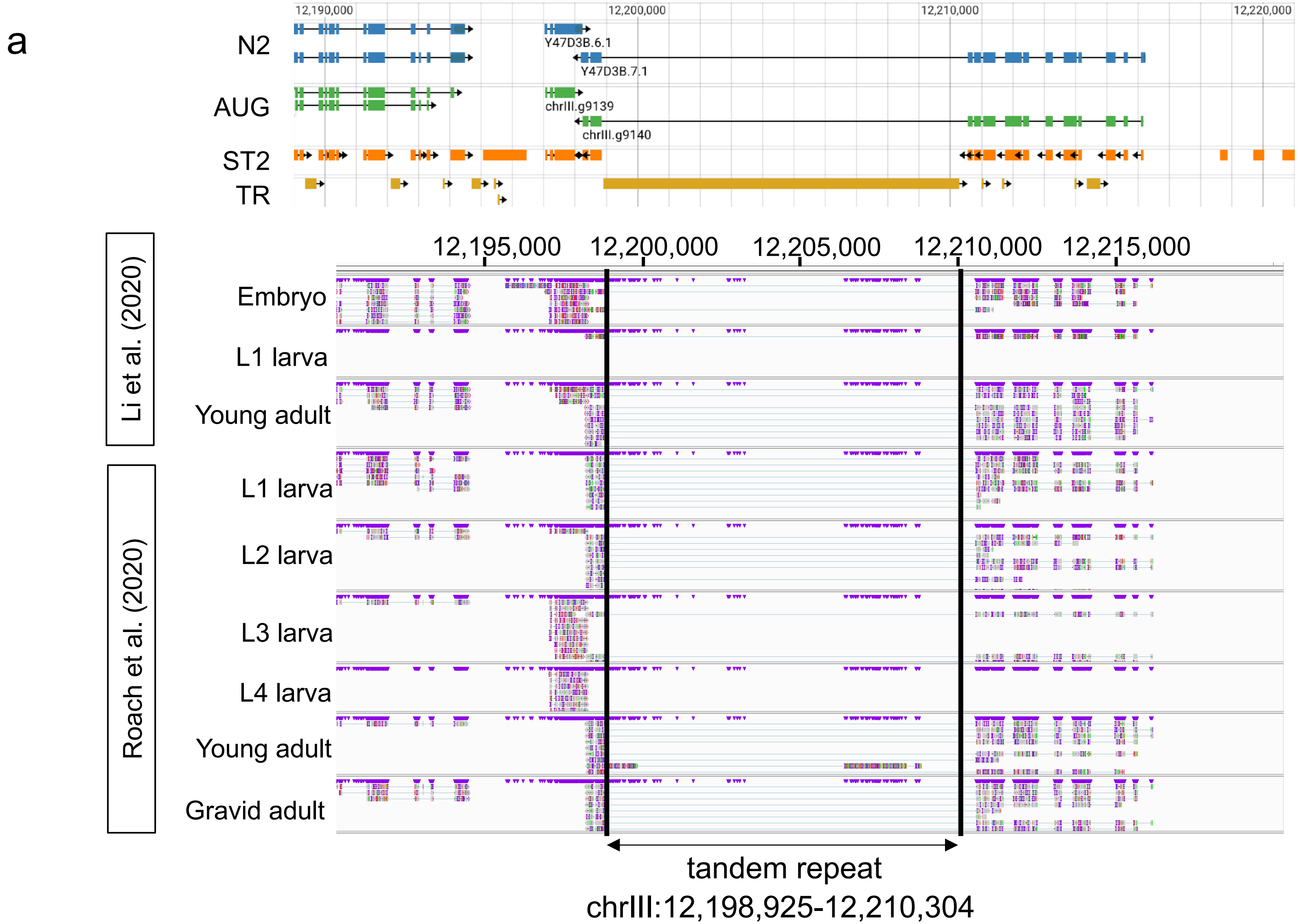

b

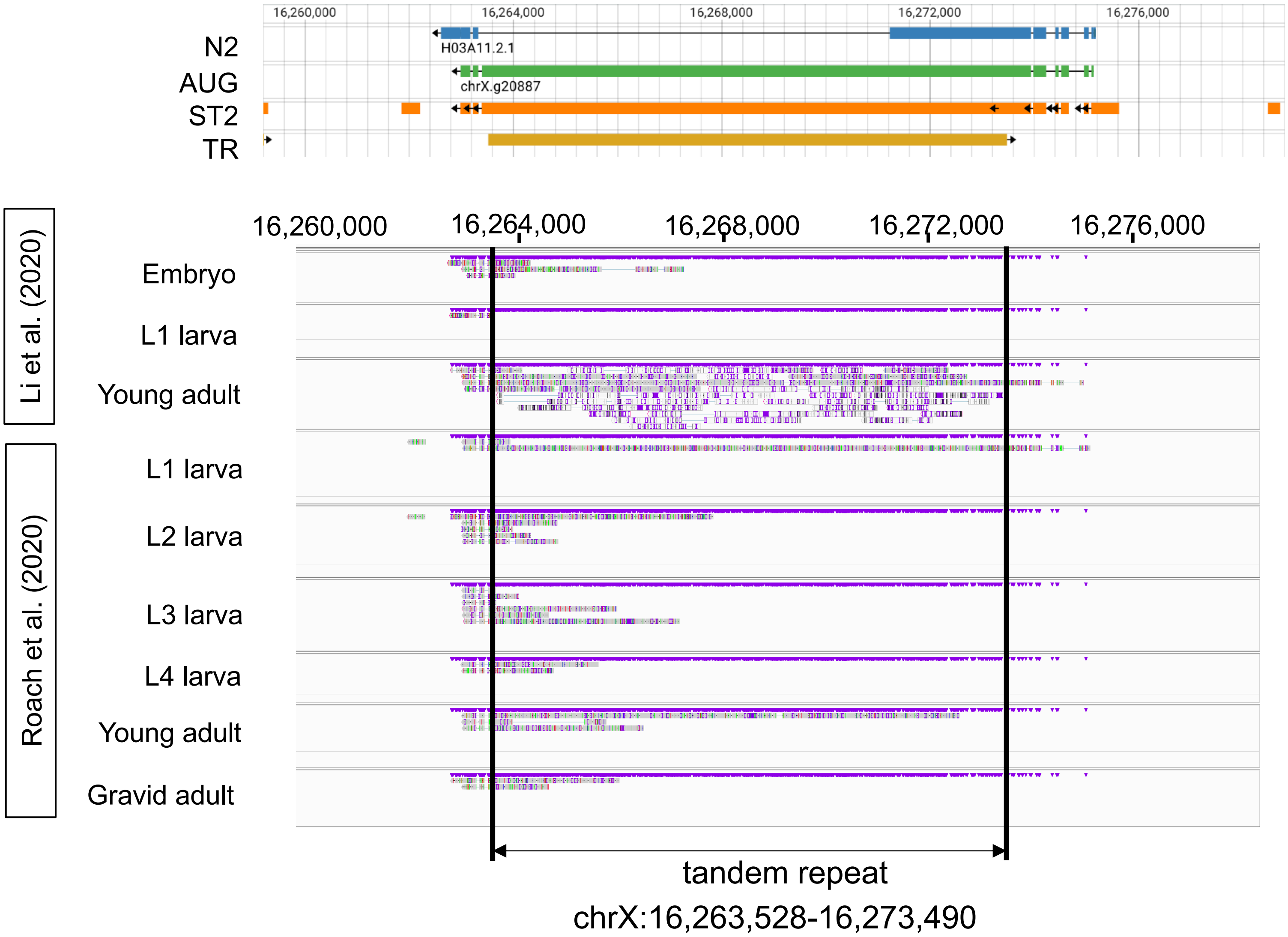
16,260,000
16,264,000
16,268,000
16,272,000
16,276,000
N2
H03A11.2.1
AUG
chrX.g20887
ST2
TR
Li et al. (2020)
Embryo
L1 larva
Young adult
Roach et al. (2020)
L1 larva
L2 larva
L3 larva
L4 larva
Young adult
Gravid adult
tandem repeat
chrX:16,263,528-16,273,490

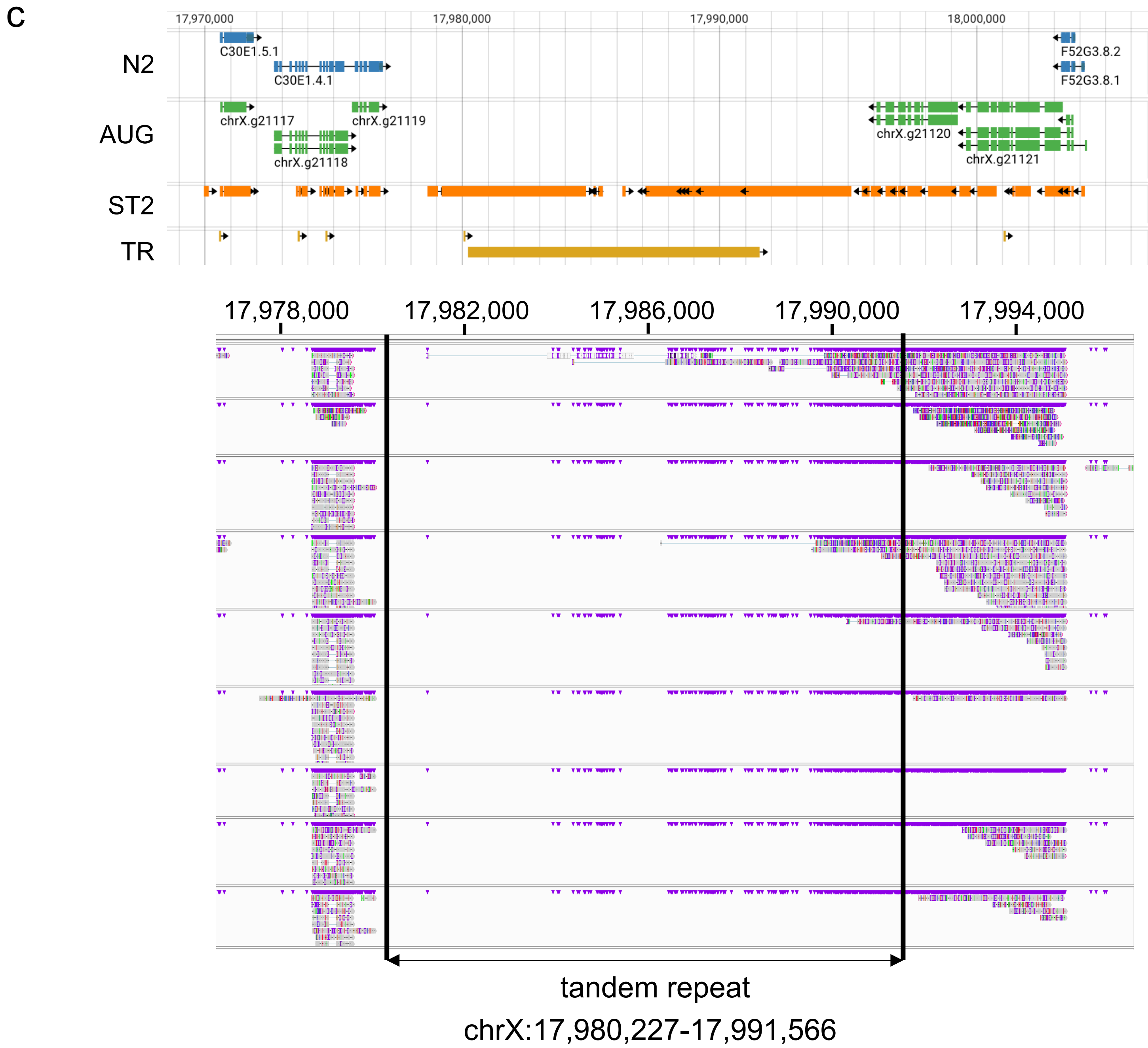
c
17,970,000
17,980,000
17,990,000
18,000,000
N2
C30E1.5.1
C30E1.4.1
F52G3.8.2
F52G3.8.1
AUG
chrX.g21117
chrX.g21119
chrX.g21118
chrX.g21120
chrX.g21121
ST2
TR
17,978,000
17,982,000
17,986,000
17,990,000
17,994,000
tandem repeat
chrX:17,980,227-17,991,566
