## Supplementary Figure 16 for "CGC1, a new reference genome for *Caenorhabditis elegans*"

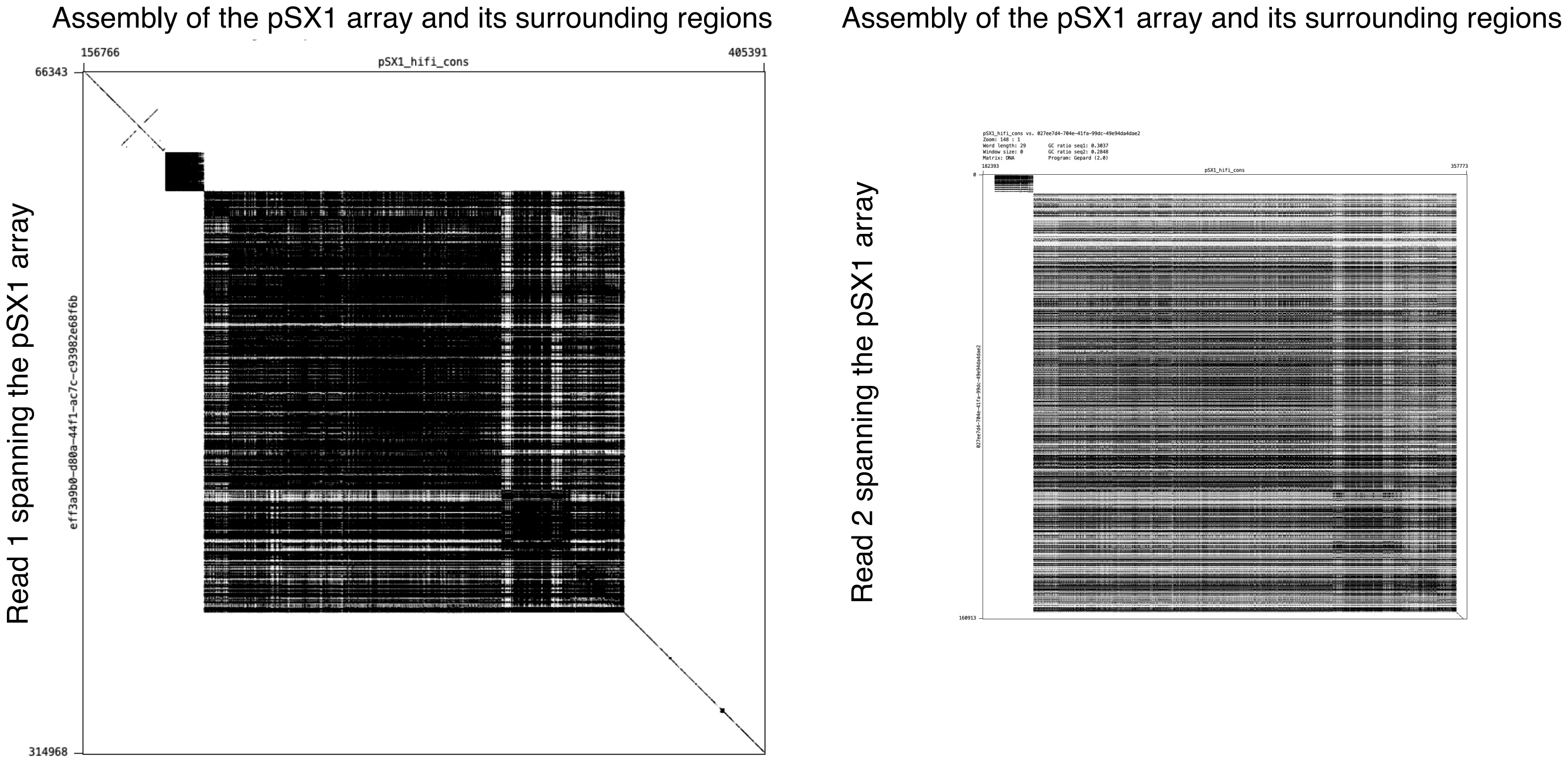


**Supplementary Figure 16. Two ultralong reads (named 1 and 2) from different molecules that span the pSX1 array.** The left figure shows a dot lot of read 1 and an assembled region of the pSX1 array and its surrounding regions. The right is a dot plot of read 2 and the assembled region.
