## Supplementary Figure 17 for "CGC1, a new reference genome for *Caenorhabditis elegans*"

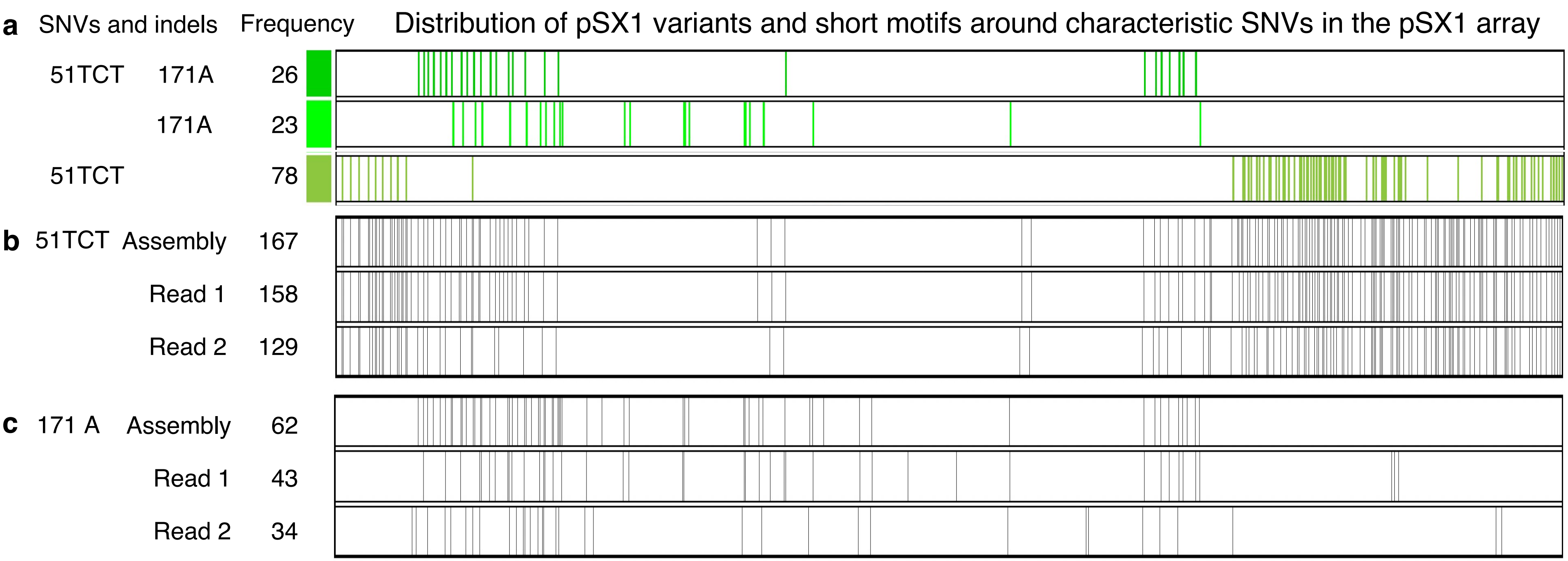


**d**

1 20
TTTTTCGGGTTTTTTGAAAT
GAATATCGTAGCTACAGAAA
CGGTAGTACACTCTTCTGAA
AATACAAAAAATTTGCAATT
TTTATAGCTAGGACACTTTT
TGTCTGCCCAAATATAAGCA
ACCAAAAATAATTTCCAAGT
TTTTGATGATTTGTTGCATA
TTGAAAAAAACA

51TCT TCTTCTTCTGAA
51
