## Supplementary Figure 18 for "CGC1, a new reference genome for *Caenorhabditis elegans*"

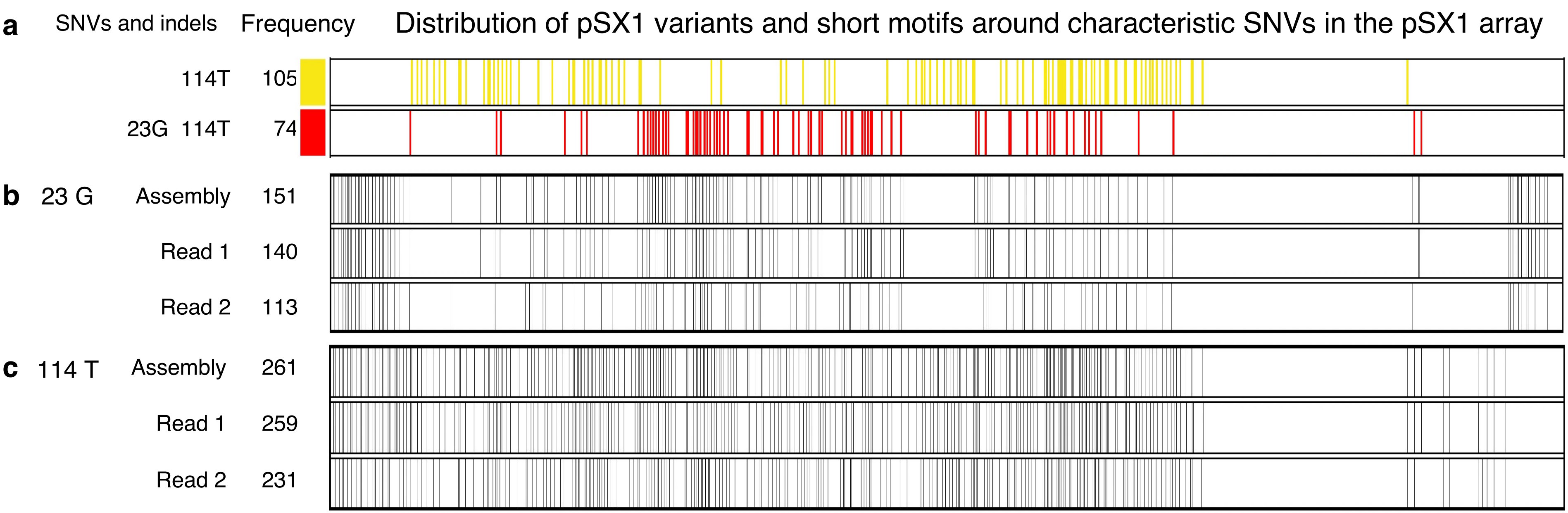


**d**

```
1                  20
TTTTTCGGGTTTTTTGAAAT
GAATATCGTAGCTACAGAAA
CGGTAGTACACTCTTCTGAA
AATACAAAAAATTTGCAATT
TTTATAGCTAGGACACTTTT
TGTCTGCCCAAATATAAGCA
ACCAAAAATAATTTCCAAGT
TTTTGATGATTTGTTGCATA
TTGAAAAAAACA

23G  TGAGTAT
        23

114T AATTTAA
        114
```

**Supplementary Figure 18: Confirmation of pSX1 variant occurrences in two Nanopore ultralong reads that span the pSX1 array. a.** The two pSX1 variants have SNVs 23G and 114A shown in the leftmost column. Vertical bars display occurrences of the two pSX1 variants. **b-c.** Vertical bars show distributions of two motifs TGAGTAT (b) and AATTTAA (c) around respective SNVs 23G and 114T. They are almost cosistent with distributions of the two pSX1 variants with 23G and 114T in Figure a. Bars absent in Figure a are found in minor pSX1 variants. **d.** The representative pSX1 sequence of length 172 nt. The two underlined substrings are used for designing short motifs TGAGTAT (in Figure b) and AATTTAA (in Figure c) .
