## Supplementary Figure 19 for "CGC1, a new reference genome for *Caenorhabditis elegans*"

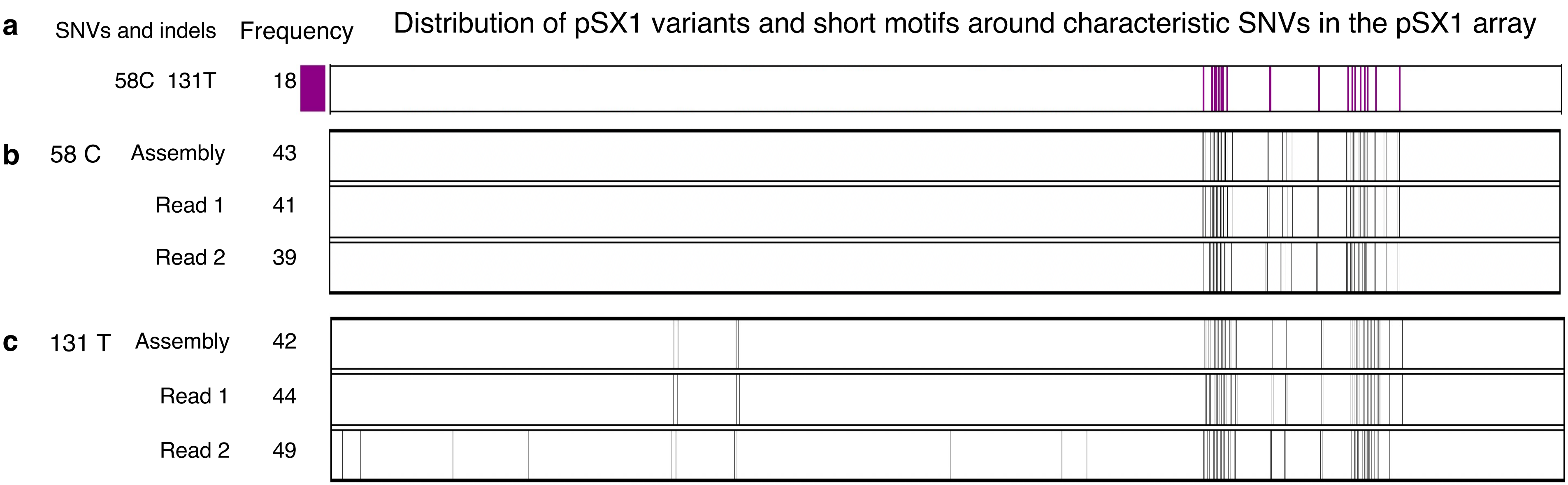


**d**

```
1                  20
TTTTTCGGGTTTTTTGAAAT
GAATATCGTAGCTACAGAAA
CGGTAGTACACTCTTCTGAA
AATACAAAAAATTTGCAATT
TTTATAGCTAGGACACTTTT
TGTCTGCCCAAATATAAGCA
ACCAAAAATAATTTCCAAGT
TTTTGATGATTTGTTGCATA
TTGAAAAAAACA

58C   TCTCAAA
         58

131T ATATTTT
         131
```

**Supplementary Figure 19: Confirmation of pSX1 variant occurrences in two Nanopore ultralong reads that span the pSX1 array. a.** The pSX1 variant had SNVs 58C and 131T in the leftmost column. **b-c.** Vertical bars show distributions of two motifs TCTCAAA (b) and ATATTTT (c) around respective SNVs 58C and 131T. **d.** The representative pSX1 sequence of length 172 nt. The two underlined substrings are used for designing short motifs TCTCAAA (in Figure b) and ATATTTT (in Figure c) .
