## Supplementary Figure 20 for "CGC1, a new reference genome for *Caenorhabditis elegans*"

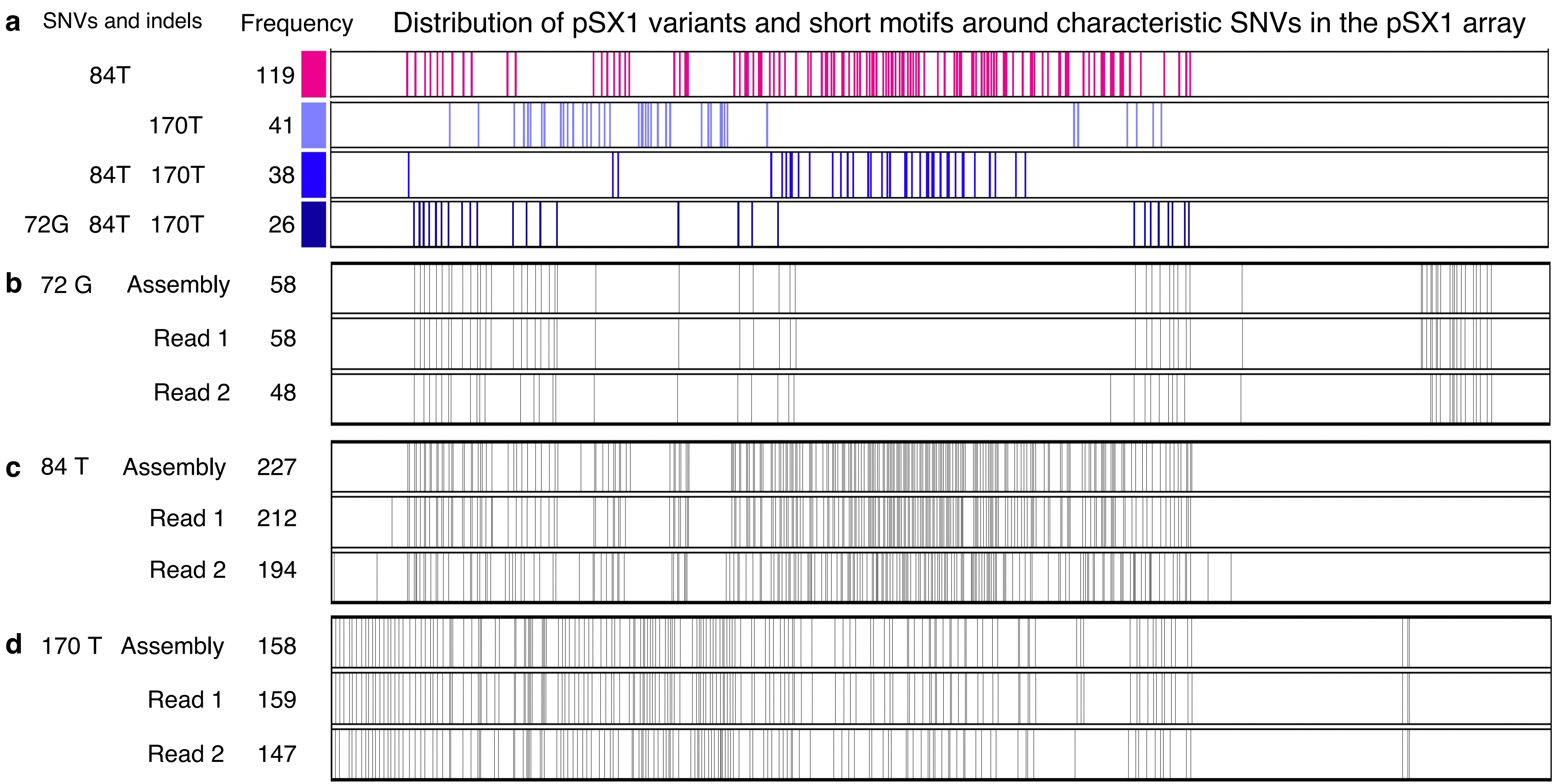


**e**

```
1                  20
TTTTTCGGGTTTTTTGAAAT     72G   AAAGTTG
GAATATCGTAGCTACAGAAA               72
CGGTAGTACACTCTTCTGAA
AATACAAAAAATTTGCAATT     84T   TTTTTAG
TTTATAGCTAGGACACTTTT               84
TGTCTGCCCAAATATAAGCA
ACCAAAAAATAATTTCCAAGT    170T  AAATCAT
TTTTGATGATTTGTTGCATA               170
TTGAAAAAAACAT
```

**Supplementary Figure 20: Confirmation of pSX1 variant occurrences in two Nanopore ultralong reads that span the pSX1 array. a.** The four pSX1 variants have SNVs 72G, 84T, and 170T in the leftmost column. **b-d.** Vertical bars show distributions of two motifs AAAGTTG (b), TTTTTAG (c), and AAATCAT (d) around respective SNVs 72G, 84T, and 170T . **d.** The representative pSX1 sequence of length 172 nt. The three underlined substrings are used for designing short motifs AAAGTTG (Figure b), TTTTTAG (Figure c), and AAATCAT (Figure d) .
