## Supplementary Figure 21 for "CGC1, a new reference genome for *Caenorhabditis elegans*"

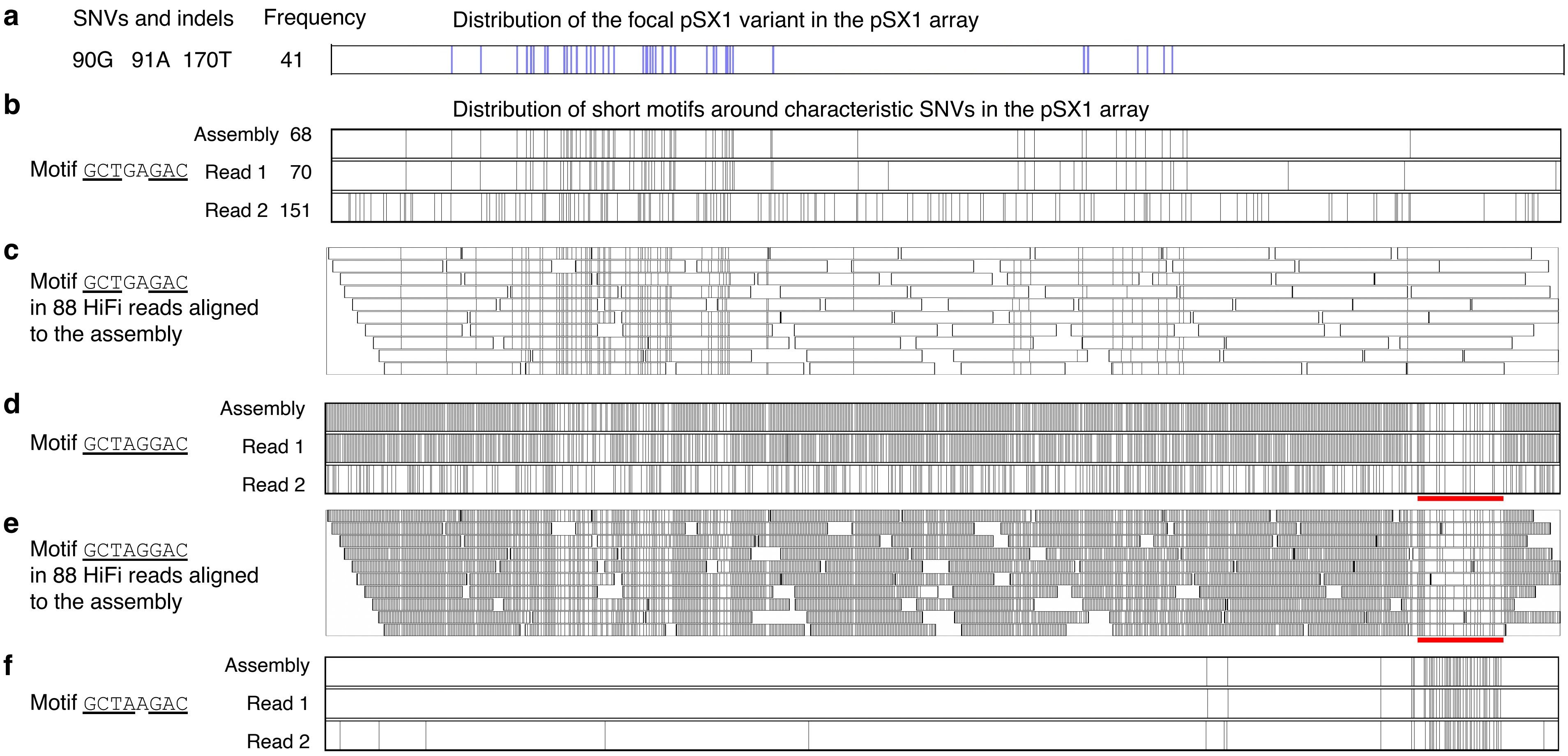


**g**

```
1                  20
TTTTTCGGGTTTTTTGAAAT
GAATATCGTAGCTACAGAAA
CGGTAGTACACTCTTCTGAA
AATACAAAAAATTTGCAATT
TTTATAGCTAGGACACTTTT
TGTCTGCCCAAATATAAGCA
ACCAAAAATAATTTCCAAGT
TTTTGATGATTTGTTGCATA
TTGAAAAAAACA
90G 91A   GCTGAGAC
             90,91
```

**Supplementary Figure 21: Confirmation of pSX1 variant occurrences in Nanopore ultralong reads that span the pSX1 array. a.** The focal pSX1 variant has SNVs 90G, 91A, and 170T in the leftmost column. **b.** Vertical bars show distributions of motif GCTGAGAC around consecutive SNVs 90G and 91A. Although the motif distributions of the assembly and read 1 are consistent, additional motif occurrences are seen in read 2. This discrepancy motivated us to confirm occurrences of GCTGAGAC in HiFi reads that aligned to the region in the assembly. **c**. Distributions of the short motif GCTGAGAC in 88 HiFi reads are largely consistent with the motif distribution in read 1 of Figure **b**, but do not support the additional motif occurrences in read 2 that seem to be caused sequencing errors in read. **d-f.** To further confirm the distribution of the focal pSX1 variant, we also examine the distributions of motif GCTAGGAC that is absent in the variant but is present in the representative pSX1 and the other variants except for the focal variant. As expected, the distributions of GCTAGGAC in the assembly, read 1 (Figure **d**), and 88 HiFi reads (Figure **e**) complement the distribution of GCTGAGAC in Figure **b**, and nearly occupy the white area where the pSX1 variant does not exist in Figure **a**. Two motifs GCTAGGAC and GCTGAGAC appear less frequently in the red-colored region that actually has many occurrences of another motif GCTAAGAC which replaces G with A at position 91 (Figure **f**). **g.** The representative pSX1 sequence of length 172 nt. The underlined substring is used for designing short motif GCTGAGAC (in Figure **b**).
